# Conformational maps of human 20S proteasomes reveal PA28- and immuno-dependent inter-ring crosstalks

**DOI:** 10.1101/2020.05.05.078170

**Authors:** Jean Lesne, Marie Locard-Paulet, Julien Parra, Dušan Zivković, Marie-Pierre Bousquet, Odile Burlet-Schiltz, Julien Marcoux

## Abstract

Here, we used for the first time Hydrogen-Deuterium eXchange coupled to Mass Spectrometry (HDX-MS) to investigate conformational differences between the human standard 20S (std20S) and immuno 20S (i20s) proteasomes alone or in complex with PA28αβ or PA28γ activators. Their solvent accessibility was analyzed through a dedicated bioinformatic pipeline including stringent statistical analysis and 3D visualization. These data confirmed the existence of allosteric differences between the std20S and i20S at the surface of the α-ring triggered from inside the catalytic β-ring. Additionally, binding of the PA28 regulators to the 20S proteasomes modified solvent accessibility due to conformational changes of the β-rings. This work is not only a proof-of-concept that HDX-MS can be used to get structural insights on large multi-protein complexes in solution, it also demonstrates that the binding of the std20S or i20S subtype to any of its PA28 activator triggers allosteric changes that are specific to this 20S/PA28 pair.

## Introduction

The ubiquitin proteasome system (UPS) is central to proteostasis. Its most downstream element is the 26S proteasome that clears the cells of abnormal, denatured or damaged proteins and regulates degradation of short-lived proteins, in most cases conjugated to ubiquitin. The 26S proteasome is a highly conserved compartmentalized multi-catalytic protease composed of more than 30 subunits constituting the 20S catalytic core and the 19S regulatory complex. Its activity is directly involved in many cytokines and hub proteins intracellular concentration, and regulates immunogenic peptide production. It is the focus of intense regulation and dynamically localizes within cells in response to physiological and external perturbations^1,2^ and its genetic polymorphism is directly responsible for numerous pathologies including cancer, heart disease and type 2 diabetes to name a few^3^.

The main mechanism regulating the proteasome activity is the substitution of the catalytic subunits β1, β2 and β5 constituting the standard 20S (std20S) with other subunits. For example, the immunoproteasome (i20S) contains the immuno-subunits β1i, β2i and β5i that have different cleavage specificities and generate immunogenic peptides^4^. It is strongly expressed in immune cells and can be induced in most tissues by interferon-γ.

The proteolytic activity of the 20S is also regulated by interacting proteins, the most common being the ATP-dependent 19S activator involved in the UPS degradation pathway. However, less abundant ubiquitin-independent regulators such as PA28αβ, PA28γ or PA200 also activate the 20S and modify its substrate specificity. In addition, a vast collection of Proteasome Interacting Proteins (PIPs) regulate or assist proteasomal functions^5–11^.

Besides its subunit composition and association with regulators and/or PIPs, the proteasome is subjected to a wide variety of post-translational modifications (PTMs) affecting its subunits activity^12–14^. The combination of these three levels of regulation (catalytic subunits, regulators, PTMs) implies a wide proteasome subtype heterogeneity, which probably results in specialized functions that can adapt protein degradation pathways to changing conditions in the cell. One of today’s main challenges is to understand how different proteasome subtypes influence its proteolytic activity and the cell function.

The proteasome has been intensely studied from a structural and functional point of view since its discovery in 1988^15^. Electron microscopy (EM) allowed to observe the 20S in complex with different regulators^16–21^ together with certain catalytic intermediate-states of the 26S^22^. Covalent cross-linking coupled to mass spectrometry (MS) helped refining 26S structures^23,24^ and generating structural models involving different partners, including the 19S^18^, Ecm29^11^ and other PIPs^11^. X-ray crystallography also provided high-resolution structures of proteasomes alone or in the presence of covalent inhibitors^25^, regulators^26,27^, and complexes of proteasome with their associated regulators^11,25,26^.

The complex size is obviously a limit for its analysis by Nuclear Magnetic Resonance (NMR). However, the structure of the eukaryotic proteasome from *Thermoplasma acidophilum*, containing only one type of subunit α and β, has been resolved. It showed evidence of a remodelling of the catalytic sites located in the center of the proteasome upon binding of a regulatory particle on its surface, which can be described as an outer-to-inner allosteric change^28^. A reverse inner-to-outer change was also observed at the binding interface when modifying the catalytic site, but this was not confirmed by the comparison of the std20S X-ray structure^29^ with the recent i20S X-ray^30^ and cryo-EM^31^ structures (RMSD=0.392 Å and 0.480 Å, respectively). However, the outer-to-inner mechanism was recently shown in the human 20S cryo-EM structure upon PA200 binding^20^: conformational changes were found not only at the binding interface (opening of the pore), but also down to the catalytic sites.

Despite these achievements, high-resolution methods are still limited by either the size, the heterogeneity, or the dynamics of the purified complexes. This is evidenced by the small number of structures of the 20S bound to regulators and their numerous unresolved regions.

Here, we optimized the emerging method Hydrogen-Deuterium eXchange coupled to MS (HDX-MS) to investigate conformational changes occurring upon binding of the std/i20S to P28 regulators. HDX-MS allows to differentiate flexible and/or accessible from rigid and/or protected regions of a complex and identifies interaction surfaces and allosteric changes in solution. We present the first HDX-MS analysis of the entire human std and i20S core particles as well as their PA28αβ and PA28γ regulators, mapping solvent accessibility/dynamics for each complex. The conformational maps of the std and i20S present significant differences, and their differential analysis with and without regulators provide a molecular rationale for their distinct functions. Our data provide evidence for the human 20S inner-to-outer allosteric change upon incorporation of the immuno-subunits and the reverse outer-to-inner transduction signal upon PA28 binding, illustrating the interplay between the different proteasome regulation pathways. Altogether, this work highlights the potential of HDX-MS as a new tool to generate low resolution but informative structural information on large hetero-oligomeric complexes. It opens the door to many other applications, including identifying PIPs binding surfaces and the stabilizing/destabilizing effects of other regulators of the 20S activity, including small molecules.

## Results

### Dynamic interfaces uniting the 4 rings of the 20S proteasome

HDX consists in the exchange of backbone amide hydrogens (H) with deuteriums (D) in proteins in solution. The exchange rate depends on the solvent accessibility and/or the flexibility of a given region, thereby providing low resolution but crucial information on protein conformation (**Fig. 1A**). We performed two distinct analyses: (i) identification of the most accessible and/or dynamic regions of the proteins in solution (**Fig. 1B-C**), and (ii) differential analysis of each protein alone or in complex (**Fig. 1D-E**). For differential analyses, we visualized the sum of differences of relative deuterium uptakes (RDU) across time points between conditions for peptides passing the statistical thresholds (see Methods). This allowed the identification of regions presenting significant differences of deuteration, which indicates a modification of their solvent accessibility upon complex formation that can be due to the presence of a binding interface or to allosteric changes. This approach was used to identify the conformational differences between the std20S and i20s, and to compare their structural changes upon binding to PA28αβ or PA28γ.

**Figure 1.**
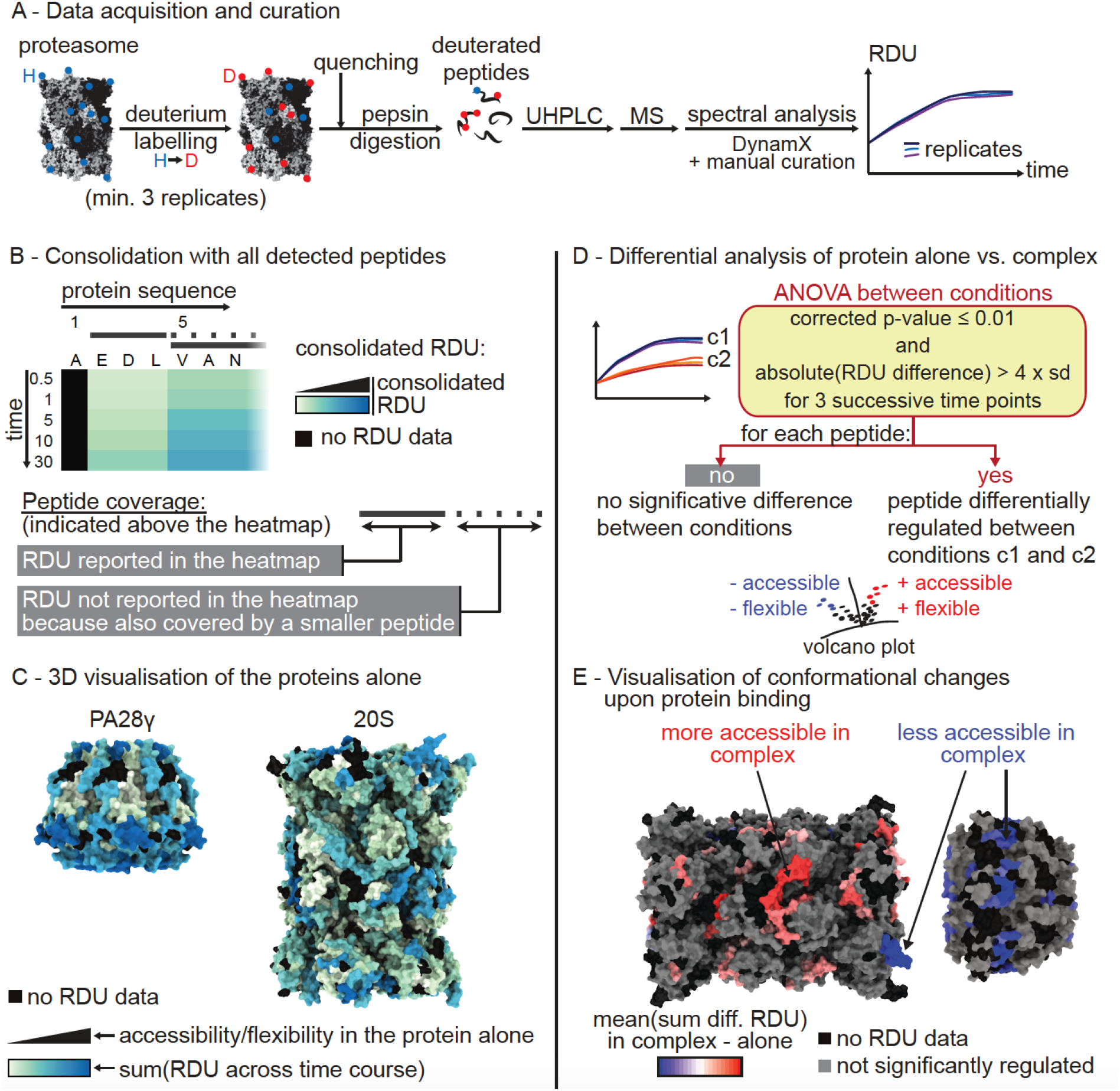
HDX workflow applied to the analysis of the 20S proteasome and its regulators. (**A**) Workflow applied for HDX sample preparation, data acquisition, inspection and curation. (**B**) Consolidation: projection of peptide RDUs to the protein sequence was performed by mapping the RDUs of the smallest peptides (solid lines) to the amino-acid portions they cover on the protein sequence. The RDUs of the longer peptides were not reported on the heatmap (dashed lines). (**C**) Visualization of the solvent accessibility of the protein analyzed alone through mapping of the sum of consolidated RDUs across all time points to their 3D structure. (**D**) Statistical analysis applied to the comparison of proteins alone or in complex (20S + PA28). (**E**) 3D visualization of the differences of RDU measured between the proteins alone or in complex. The consolidation is performed as described in (B) with the sum(mean difference of RDUs per time point). “H”: hydrogen; “D”: deuterium; “diff.”: difference; “RDU”: relative deuterium uptake.

We obtained 89% subunit sequence coverage on average for the 17 subunits of the std/i20S (**Fig. S1-S2, Table S1** and **Table S2**). HDX-MS results were mapped on the 3D structure of the human std20S (PDB: 5LE5, **Fig. 2A-B**). As expected, we found a faster and higher deuteration of the solvent-facing α-ring interface of the std20S compared to any other ring interfaces (**Fig. 2C**). This observation benchmarks our approach by confirming that high RDUs directly indicate regions of the proteasome that are more dynamic and potentially accessible for interaction with regulators or PIPs. The N-terminal (N-ter) part of the α-subunits constituting the α-ring were highly deuterated. These N-ter are located around the proteasome pore entry, and their high RDU strongly supports the hypothesis of a flexible pore entrance in the std20S that fluctuates between open and closed states^32–34^. Furthermore, the α-ring presented very dynamic patches consisting of non-contiguous peptides from the same subunit in α3, α4 and α5. These may interact with many different PIPs. Our analysis also identified dynamic patches at the interface between different subunits like α1/α2, α3/α4 and α5/α6 that correspond to the 1-2, 3-4 and 5-6 α-pockets^35^ known to accommodate the C-terminal (C-ter) tails of Rpt3, Rpt2 and Rpt5, respectively^18,35^. We also identified flexible patches on the interface of the α-ring facing the β-ring, where amino acid stretches of α4, α5 and α7 were slightly more flexible than the remaining of the interface (**Fig.2D** and **Fig.S3**).

**Figure 2.**
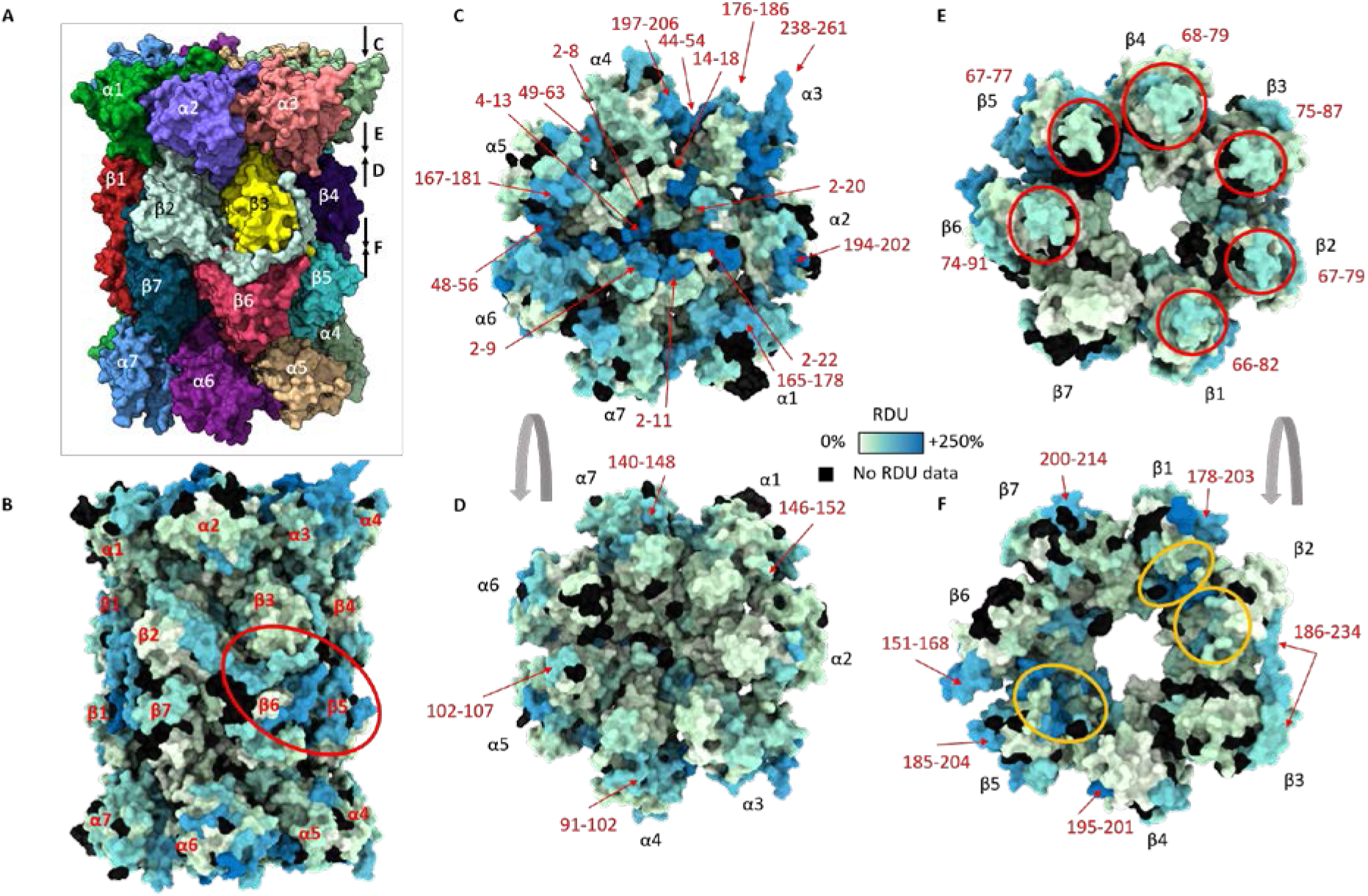
HDX-MS of the std20S reveals a dynamic pore entrance and inter-ring interface. (**A-B**) Structure of the human std20S proteasome (PDB: 5LE5)^29^ showing the four stacked αββα heptameric rings colored by subunit (**A**) or relative deuterium uptake (RDU) (**B**). (**C-F**) Their solvent accessibility is represented on each face of the α-ring (solvent interface (**C**) and β-ring interface (**D**)) and β-ring (α-ring interface (**E**) and β-ring interface (**F**)) as the sum of consolidated relative deuterium uptakes between 0.5 and 30 min (described in **Fig. 1B,C**). Regions discussed in the text are indicated with red arrows and the moderately flexible/accessible bulges at the interface between the α- and β-ring are circled in red in (**E**). The corresponding residues are indicated in red around the structures. Active sites are circled in orange (**F**). The ring faces visualized in D to F are indicated by arrows on the right of the structure in (**A**).

The β-ring was globally less deuterated than the α-ring (**Fig. 2E-F**), which was expected since it does not present a large solvent interface. However, after 30 minutes, some α-facing bulges were moderately deuterated in all β subunits but β7 (red circles in **Fig. 2E**, and **Fig. S3**). Interestingly, some of these dynamic loops face the highly deuterated regions of the α-ring described above and define flexible regions of the α-ring/β-ring interface. The most dynamic regions of the β-ring were located on its outer surface and were constituted of the C-ter of β1, β2, β4, β5 and β7 as well as loops between β-strands and α-helices, particularly between helices 4 and 5 of β1, β4, β5, β6, β7 and the C-ter of β2 (**Fig. 2F** in red and **Fig. S3**). These regions are particularly interesting since they can potentially interact with the numerous PIPs described in the literature.

### Conformational differences between the std and i20S active sites and substrate pockets

The std20S catalytic chamber localized at the β-ring/β-ring interface was locally very flexible (**Fig. 2F and Fig. 3**). The residues forming the S1 and S2 pockets^25^ of β1 were highly deuterated, together with the T23 of the S3 pocket (**Fig. 3B**). The catalytic residues of β2 were poorly deuterated compared to the remaining of this subunit (**Fig. 3C**), with the exception of 45-53 constituting its S1 and S2 pockets. Conversely, most of the residues forming β5 catalytic site and substrate pockets were highly deuterated (**Fig. 3D**).

**Figure 3.**
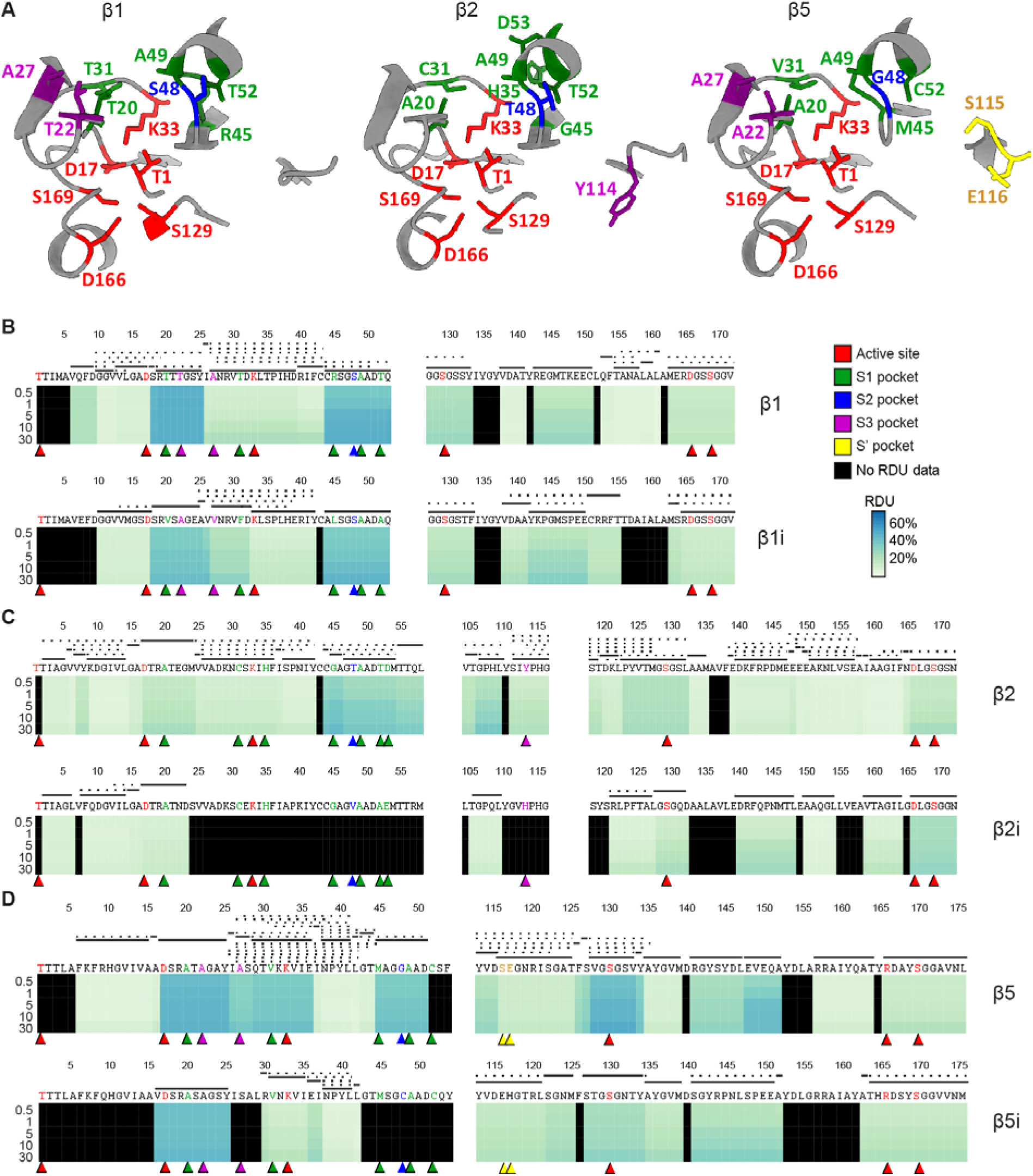
Comparison of the catalytic subunit solvent accessibilities. (**A**) Representation of the three catalytic subunits β1 (left), β2 (middle) and β5 (right) of the std20S (PDB: 5LE5). The residues forming the active site and the substrate-specific pockets are indicated and color-coded (active site, S1, S2, S3 and S’ are represented in red, green, blue, purple and yellow, respectively). (**B-D**) Deuteration heatmaps showing the relative deuterium uptake (RDU) for each timepoint of the kinetics, along the protein sequences of β1/β1i (**B**), β2/β2i (**C**) and β5/β5i (**D**). The peptide sequence coverage is presented above the heatmaps as explained in **Fig. 1B**. For the sake of clarity, only the sequence stretches containing residues of the active sites and substrate pockets are represented here.

The direct comparison of RDUs between the standard and immuno-catalytic subunits (β1/β1i, β2/β2i and β5/β5i) was not possible due to sequence differences and a relatively poor sequence coverage of the immuno-specific subunits (**Fig. S3** and **Table S1**). Thus, we cannot comment on the accessibility of the S1, S2 and S3 pockets of β2i (**Fig. 3B**). Nevertheless, our data showed that T20 (pocket S1) and T22 (pocket S3) of β1 were more deuterated than the corresponding region of β1i. The active site and S1 pocket of β5 also appeared more flexible in the standard than in the immuno-subunit (**Fig. 3D**). Although limited, these data inform on the differences of flexibility between std and i20S catalytic sites.

### An inner-to-outer conformational change upon substitution of standard to immuno-subunits

Although mice i20S and std20S catalytic subunits possess very similar β2 and β2i substrate binding channels, their S1 and S3 substrate pockets are smaller in β1i *vs*. β1; and β5i possesses smaller S2 and S3 pockets whereas its S1 pocket is larger than in β5^25^. However, very little structural difference is found between the α and non-catalytic β subunits of the i20S and std20S (Root-Mean Square Deviation of the Cα < 0.72 A°). The same is true when comparing the recent structure of the human i20S^30^ with the std20S^29^ (RMSD of the Cα < 0.67 Å). The inner-to-outer allosteric change observed on a prokaryotic 20S by NMR^28^ was not seen in these crystallographic structures, maybe due to crystal packing. In order to confirm or infirm this inner-to-outer allosteric change, we compared the RDUs of all the non-catalytic subunits of the i20S (**Fig. S2, S3** and **Table S2**) with those of the std20S.

The differences in solvent accessibility of the catalytic and substrate pockets between the std20S and i20S translated to significant conformational changes on the non-catalytic 20S subunits. The results of our statistical analysis are presented in **Fig. 4, Fig. S2**, **Fig. S4**, **Fig. S5**, **Tables S3-S4**. Overall, the non-catalytic β-ring subunits were more dynamic in the i20S than in the std20S, (> 50 peptides significantly more deuterated, **Fig. 4A-B** in red, **Fig. S4-S5**). The β-ring/β-ring interface and the channel were particularly more dynamic/accessible in the i20S *vs*. std20S (**Fig. 4A** and **Fig. S4**), together with two α-facing bulges of β6 and β7 (**Fig. 4B**, red circles). Four β-facing bulges of α2/3/5/6 were also significantly more dynamic in the i20S *vs*. std20S (**Fig.4C**, red circles).

**Figure 4.**
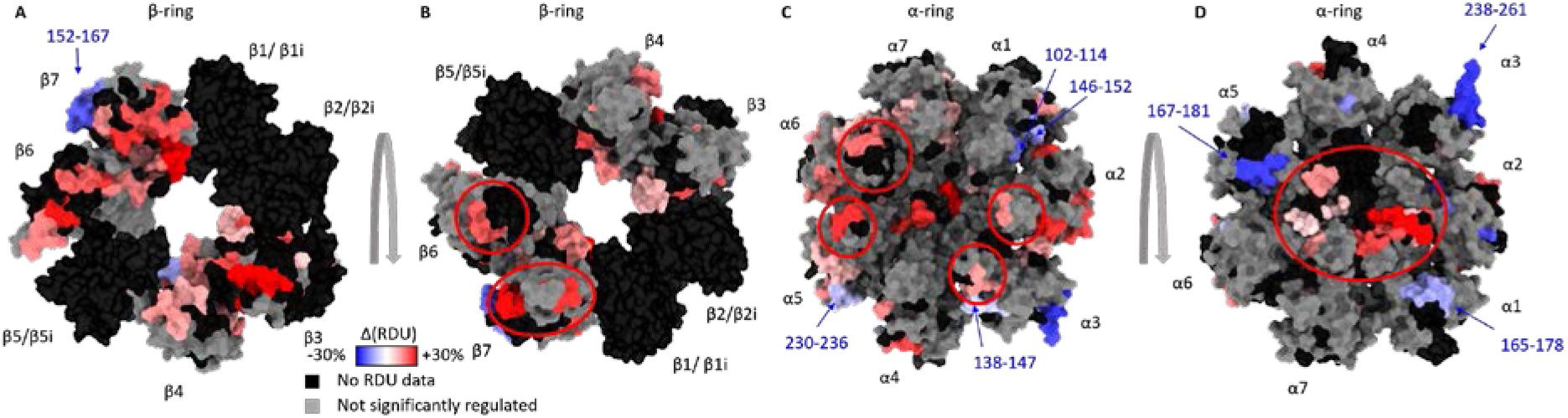
Differential analysis of std20S and i20S. (**A-D**) Sum of differential relative uptakes (Δ(RDU)) between the i20S and the std20S: β-ring/β-ring interface (**A**); β-ring facing the α-ring (**B**); α-ring facing the β-ring (**C**); solvent-facing α-ring (**D**). Color coded regions were significantly more accessible/dynamic in the std20S (blue=-30%) or i20S (red=+30%) and regions of interest are circled or annotated with the same colors. Regions that are not covered in one or both conditions, or do not pass the statistical threshold are represented in black and grey, respectively (see **Fig. 1D,E** for more details). “RDU”: relative deuterium uptake.

The pore entrance on the α-ring solvent-accessible surface (α subunits N-ter) was significantly more dynamic/accessible upon replacement of β1/2/5 by the immuno-subunits (**Fig. 4D** red circles, **Fig. S4-S5**). This suggests a long-range mechanism enabling the i20S pore to dwell longer in the open-state, and could explain its generally higher activity^36^. Conversely, three solvent-facing regions of the α-ring were significantly more dynamic/accessible in the std20S than in the i20S (**Fig. 4D** in blue, **Fig. S2**, **Fig. S5**). Since these constitute potential binding sites for the proteasome regulators, this could explain the preferential association of specific regulators to different 20S subtypes. Interestingly, these three subunits also contain regions that are more accessible in the std20S on the β-ring facing interface (**Fig. 4C**), suggesting that this dynamic behavior can be related to the presence of the standard catalytic subunits within the β-ring. More precisely, the α1 102-114 and 146-152, which were more deuterated in the std20S *vs*. i20S, are in direct contact with a portion of β2/β2i (69-72) also more accessible in the std20S (**Fig. S4**). Altogether, these results confirm the inner-to-outer allosteric effect following incorporation of the immuno-β subunits that lead to α-ring remodeling: more dynamic pore entrance but protected external anchor regions (α3 C-ter).

### An activation loop less dynamic in PA28β

There is no structure yet for PA28γ but those of the human homo-heptameric PA28α (non-physiological) and murine hetero-heptameric PA28α4β3 were resolved in 1997^26^ and 2017^27^, respectively. These monomers are composed of a four-helix bundle, assembled as a barrel-shaped heptamer (**Fig. 5**) that controls proteasome catalytic activity through a conserved activation loop^37,38^. The structures of PA28α, PA28β and PA28αβ miss a region of 15-31 residues bridging the α-helices 1 and 2, although they were present in the recombinant constructs (**Fig. 5A** and **Fig. S7** in red). These apical loops, most likely flexible, sit on top of the regulator and might thereby control the substrate access to the proteasome. The human PA28γ chain has a similar loop (35 residues), and no information on its structure has ever been reported. Since the sequence identity of human PA28γ is closer to PA28α (40%) than PA28β (35%) or *PfPA28* (34%) (**Table S5**), we generated a homology model of human PA28γ (**Fig. 5D**) using the structure of human PA28α^27^ as a template with the SWISS-MODEL server^39^.

**Figure 5.**
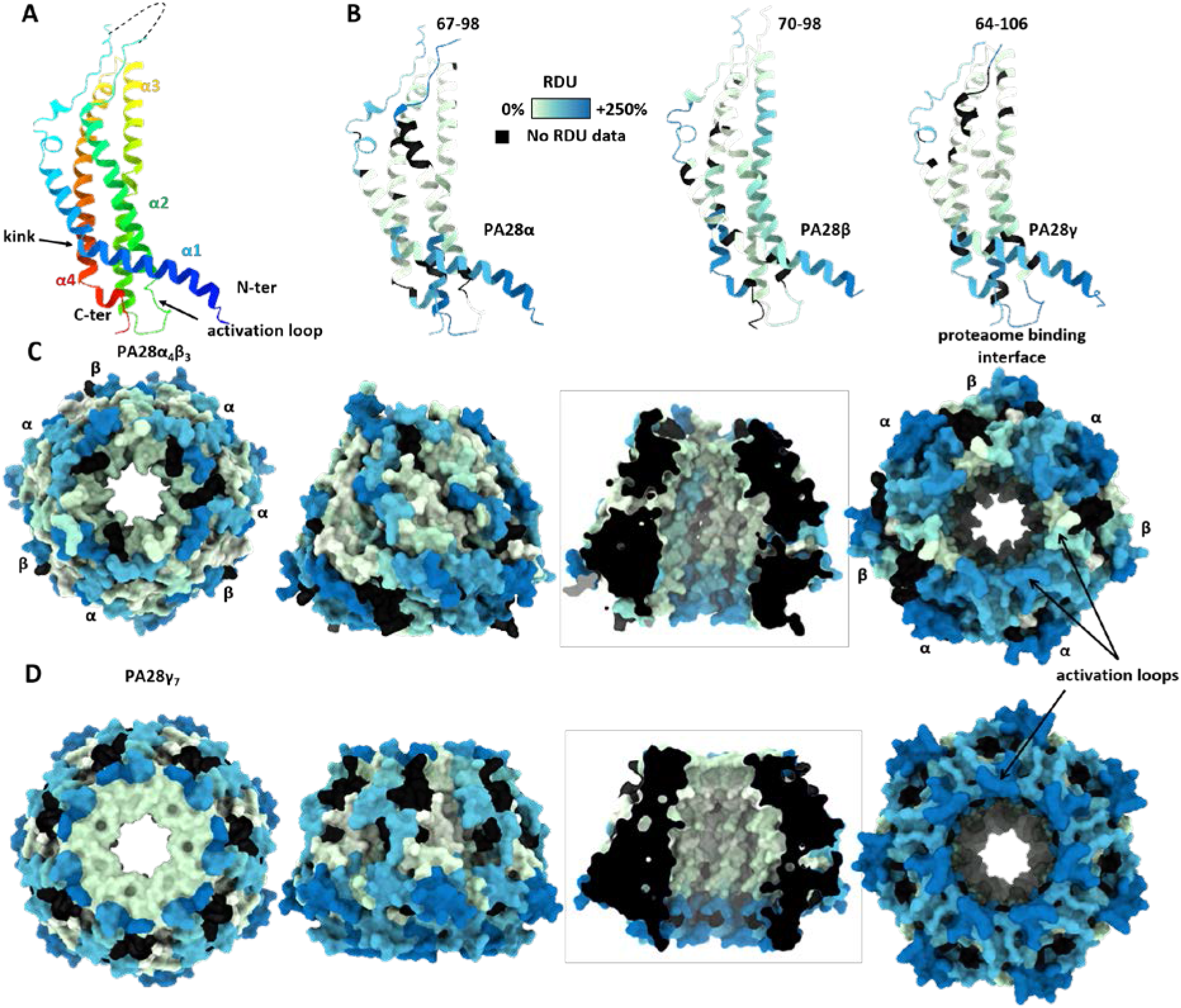
PA28 regulators present flexible loops on both ends of the channel. (**A-B**) Structure of monomeric PA28α showing the 4-helix-bundle colored based on amino-acid numbering (**A**) or sum of hydrogen-deuterium relative uptake accumulated over 30 min (**B). (C-D**) Surface accessibility of the heptameric complexes PA28α4β3 (**C**) and PA28γ7 (**D**) viewed from the top (left), front and inside (middle), and bottom (right). Regions that were not detected are represented in black (described in **Fig. 1B,C**). “RDU”: relative deuterium uptake.

The sequence coverages obtained upon digestion were above 90% for all three PA28 subunits across all conditions (**Fig. S1, Fig. S6-7** and **Table S1**). Their deuteration pattern revealed similar dynamics for both PA28αβ and PA28γ with a very strong protection of most residues inside the channel (**Fig. 5C-D**, middle and **Fig. S7**). The “missing” apical loops (**Fig. S7** in red) were very quickly deuterated, a sign of high accessibility/flexibility, that could explain their absence in known structures. The proteasome-facing interface (unstructured N- and C-termini and activation loops) (**Fig. 5C-D**, right) and the entrance of the channel (**Fig. 5C-D**, left) were also strongly deuterated. Remarkably, the regions involved in the binding (C-ter) and activation of the proteasome were very accessible/dynamic when both regulators were alone in solution. However, the activation loop was less deuterated in PA28β.

### Allosteric changes between PA28γ and PA28αβ upon 20S binding

We compared the PA28 regulators alone and in complex with the std20S. As expected, their proteasome-facing interface was highly protected in the complex compared to the regulators alone (**Fig. 6** right, in blue; **Fig. S6**, **Fig. S8-S9**). The activation loops, located at the 20S binding interface, and known to interact with the N-ter of the 20S α subunits to open its central pore, were highly dynamic/accessible in both PA28 analyzed alone (**Fig. 5C-D** right, and **Fig. S6-S7**). Our data show that PA28αβ/std20S binding strongly protected the activation loops as well as their neighboring residues (136-147, 153-159 and 232-249 in PA28α; 123-134, 140-145 and 215-221 in PA28β, **Fig. 6A** and **Fig. S8-S9**). The protection of PA28γ activation loop (region 146-152) upon std20S binding was also statistically significant but notably less pronounced (**Fig. 6B** and **Table S4**), which could indicate a greater affinity of the std20S for PA28αβ than for PA28γ.

**Figure 6.**
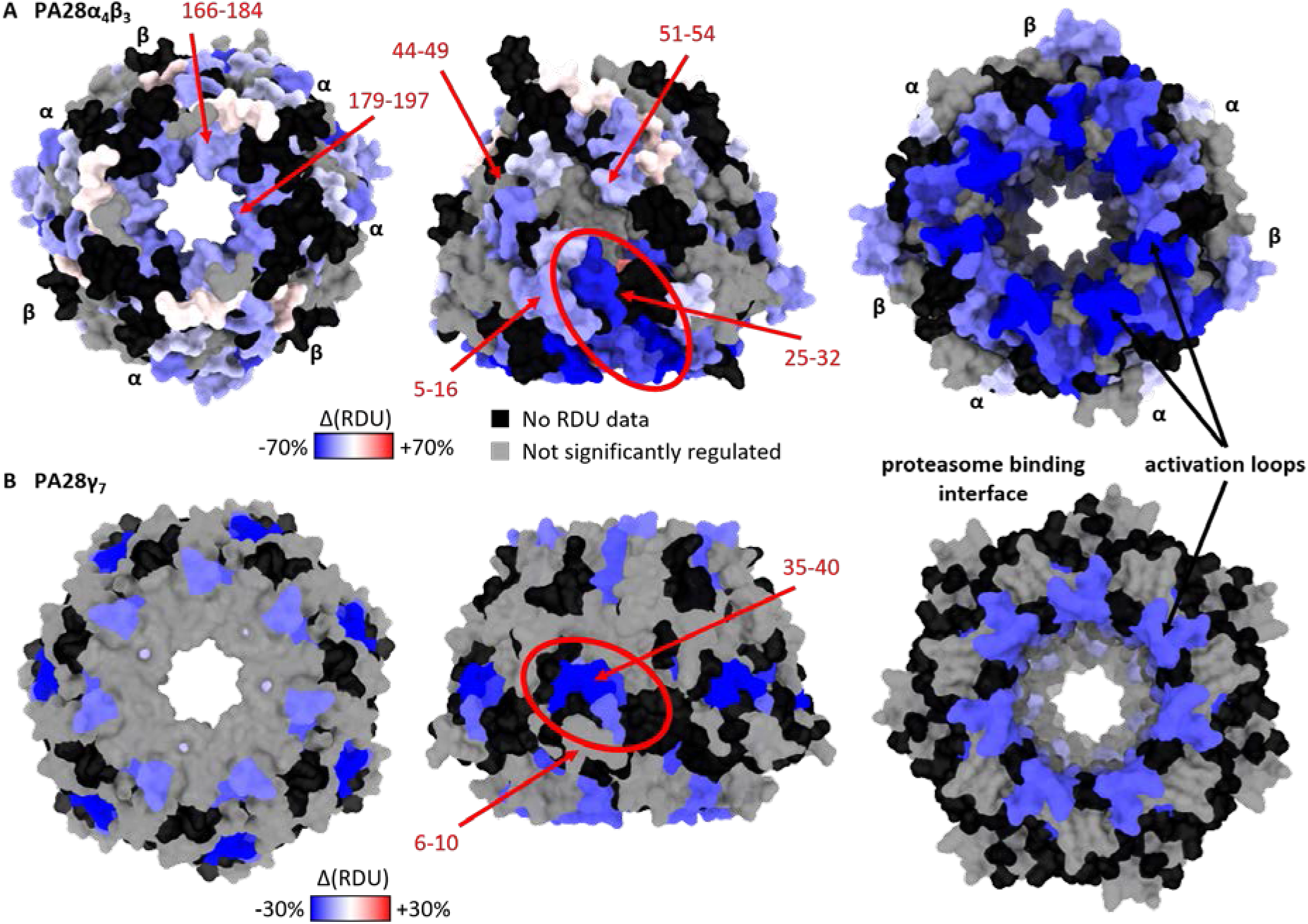
Activation of PA28αβ and PA28γ by the std20S involves a rearrangement of the N-ter kink and an allosteric change. Residues of PA28αβ (**A**) and PA28γ (**B**) that are statistically protected/less dynamic upon binding of the std20S proteasome are located around the N-ter kink (red ovals), the activation loop and 20S interface (arrows) and the entry of the channel (left). The sum of differential relative uptakes accumulated over 30 min deuteration of the PA28 regulators with and without the std20S proteasome is color coded from −70% (blue) to +70% (red) for PA28αβ (**A**) and from −30% (blue) to +30% (red) for PA28γ (**B**). Residues in grey were not considered significantly different according to the statistical thresholds presented in **Fig. 1D**. Residues with no RDU information in one or both conditions are in black. “Δ(RDU)”: sum of differences of relative deuterium uptake.

Interestingly, other regions were less dynamic/accessible upon complex formation. The kinks of the first α-helix of PA28α and PA28γ (**Fig. 5A** and **Fig. 6**, red ovals) were strongly protected upon std20S binding. The PA28β N-ter (5-16) that faces PA28α’s kink, was also significantly protected upon std20S interaction (**Fig. S8**). We thus hypothesize that PA28α/PA28β interaction might be strengthened and/or structurally rearranged upon binding to the std20S. The N-ter (6-10) of PA28γ facing the kink was slightly but not significantly protected (**Fig. 6B** and **Fig. S8**).

The opposite side of the heptamer was also partially protected upon std20S binding: the loops between helices 3 and 4 of PA28α (179-197), PA28β (166-184) and PA28γ (191, 203-208) (**Fig. 6**, K191 of PA28γ not visible in this view) as well as the flexible loop (60-101) of PA28γ (**Fig. 5B**). These regions include the constriction sites at the top of both activators (K190 in PA28α, K177 in PA28β and K191 in PA28γ) that were significantly protected upon std20S binding (**Fig. 6** and **Fig. S8**).

We then compared the impact of std20S *vs*. i20S binding on PA28 solvent accessibility. Most of the reduction in accessibility/flexibility that was observed on the PA28 chains upon binding to the std20S were dampened upon interaction with the i20S, suggesting a stronger interaction of the regulators with the std20S than the i20S (**Fig. S8**).

### A PA28-driven outer-to-inner allosteric change of the 20S proteasome

Atomic force spectroscopy showed that the 20S can alternate between open and closed states, possibly through allosteric regulation^32,33^. Our data indicate that PA28αβ and PA28γ do not present the same conformational rearrangements upon binding to the 20S, so they might allosterically affect the 20S pore entrance and/or its catalytic sites in a regulator-specific outer-to-inner mechanism, as suggested recently^20^.

We compared the std/i20S RDUs in presence or absence of PA28αβ or PA28γ (**Fig. 7, Fig. S10-S12**). Strikingly, very few regions of the α-ring in contact with the regulators were significantly protected in the std20S upon interaction with PA28αβ or PA28γ, and none in the i20S. The same C-ter region of α3 that was found to be more accessible in the std20S *vs*. i20S (**Fig. 4D**) was protected in the std20S upon interaction with PA28αβ, and to a lesser extent with PA28γ (**Fig. 7A-B**). Other regions of the std20S (α4,α5) that were protected when binding PA28αβ form two patches on the same solvent-exposed interface (**Fig. 7A** in blue). These could be the main anchorage sites of PA28αβ at the surface of the std20S. The remaining of the std20S solvent-exposed surface was globally more dynamic upon PA28αβ binding, especially near the pore entrance (N-ter of α1/2/4/5/7), which can be interpreted as an opening of the pore. Another external patch at the α1/α2 interface was also more dynamic upon binding of PA28αβ (**Fig. 7A**).

**Figure 7.**
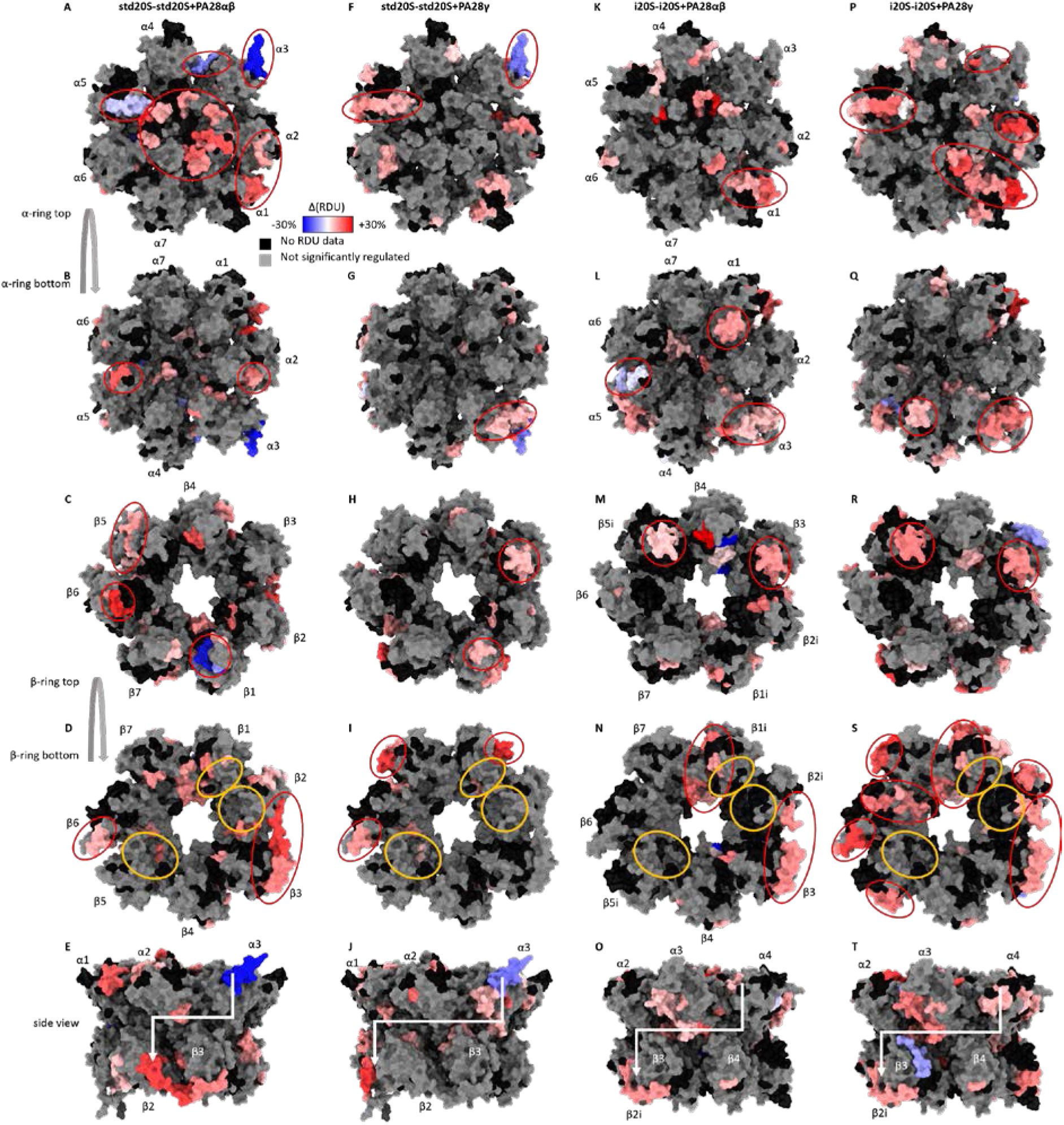
Catalytic-subunit- and regulator-specificity of the outer to inner allosteric changes. Differential relative uptakes accumulated over 30 min deuteration of the 20S proteasomes with and without the PA28 regulators. The regions that were statistically protected or more dynamic upon regulator binding are represented in blue and red, respectively, for the std20S (PDB 5LE5) and PA28αβ (PDB 5MX5) (**A-E**), the std20S and PA28γ (**F-J**), the i20S (PDB: 6E5B) and PA28αβ (**K-O**) and the i20S and PA28γ (**P-T**). Active sites at the β-ring/β-ring interface are circled in orange. The side views of the hemi-proteasome (**J/O/T/E**) show the α-ring and β-ring interface. Residues in grey were not considered significantly different according to the statistical thresholds presented in **Fig. 1D**. Residues with no RDU information in one or both conditions are in black. “Δ(RDU)”: sum of differences of relative deuterium uptake.

On the β-facing α-ring, PA28αβ binding mainly destabilized α2 (135-145) and the α5 bulge (102-107) (**Fig. 7B**, red circles). Interestingly, this bulge faces the 109-135 β6 stretch encompassing 4 peptides more deuterated upon PA28αβ binding and is in close vicinity to β5 interface, also more dynamic (**Fig. 7C**, red circles). Although not close enough to make hydrogen bonds, α2 (135-145) (**Fig. 7B**) faces the β2/β3 interface that was more flexible/accessible upon PA28αβ binding, especially in the 186-202 C-ter region of β2 extending towards β3 (**Fig. 7D**, circled in red). The only peptides of the β-ring that were less dynamic upon PA28αβ binding are located on β1 (**Fig. 7C**, circled in blue).

We compared the effect of PA28αβ binding to the std20S (**Fig. 7A-E**) with the other conditions (**Fig. 7F-T**). Although the destabilization of the α-ring N-ter by PA28αβ was almost complete in the std20S (except α7) (**Fig. 7A**), it was restricted to α1/3/4/5 in the i20S (**Fig. 7K**), suggesting a partial opening of the pore. Binding of PA28γ did not affect any of the std20S α subunit N-ter (**Fig. 7F**) and only the i20S α2 and α3 N-ter (**Fig. 7P**), indicating less PA28γ-driven opening of the i20S than the std20S pore.

Overall, PA28 binding to the std/i20S increased their RDU, especially at the pore entrance, at the α1/α2 interface (**Fig. 7A,F,K,P**), and the β2 C-ter arm extending towards β3 (**Fig. 7D,E,N,O,S,T**). Despite these similarities, our data show subtle proteasome-as well as regulator-specific differences in the allosteric motion triggered upon complex formation, as suggested recently^20^. The std20S α3 C-ter was protected upon binding of PA28αβ and PA28γ (**Fig. 7A,F**). This seemed to induce the PA28γ-specific destabilization of α2 (101-119, **Fig. 7F**), β3 (74-86) and β1 (64-70) bulges (**Fig. 7H**, red circles), down to the outer β-ring/β-ring interface of β1 (178-205), β6 (151-159) and β7 (152-167) (**Fig. 7I**, red circles). According to our statistical thresholds, the α3 C-ter protection upon PA28 binding was std20S-specific (**Fig. 7K,P**), but the remodeling of the α-ring occurred in both 20S (although differently) and was propagated to the β-ring via α1/β3/β5i and α4/β3/β5i bulges for PA28αβ and PA28γ, respectively (**Fig. 7L,M,Q,R**). It resulted in remodeling of the β-ring/β-ring interface eventually affecting β2i and β1i or β2i/1i/5i/6 with PA28αβ (**Fig. 7N**) and PA28γ (**Fig. 7S**), respectively.

The limited sequence coverage around the catalytic sites prevented us from drawing exhaustive conclusions on the differential impact of regulator/20S complex formation on the proteasome active sites. However, β1 (10-25), β2 (16-42) and β5 (29-36) active sites were partially but significantly more dynamic/accessible upon PA28αβ binding (**Fig. 7D**, orange circles), in line with an increased proteolytic activity. Binding of PA28αβ also increased the solvent accessibility of β1i active site (10-17) (**Fig. 7N**), which is in good agreement with previous work showing that the caspase-like activity of the i20S is enhanced to a higher extent by PA28αβ than PA28γ^40^. PA28γ was reported to have little or no effect on the i20S^41^, whereas it is known to increase all three std20S activities^42^. Our data do not indicate any major change in the std20S active site solvent accessibility upon interaction with the regulator (**Fig. 7I**). However, it revealed a significant decrease of β2i N-ter (2-6) solvent accessibility upon PA28γ interaction (**Fig. S11**). Thus, the catalytic activation observed upon PA28γ/std20S binding may be driven by changes occurring further away from the active site.

## Discussion

This work provides the first structural data set on the human i20S compared to the std20S alone or bound to both PA28 regulators. It rationalizes from a mechanistic point of view previous observations that replacement^36^, modification^28^ or ligand binding^36,43^ to the catalytic subunits can allosterically modify the 20S core particle structure and alter its binding to potential PIPs. A main advantage of HDX-MS is that it provides information on very flexible loops, unlike X-ray crystallography or cryo-EM. Also, it is not theoretically limited by the complex size since the proteins are digested into peptides after deuteration: it can be applied to very small proteins (unlike cryo-EM) or very large complexes (unlike NMR). Large oligomeric complexes containing a reduced number of different monomers (limited number of chemical shifts) have already been analyzed by NMR, such as the prokaryotic 20S from *T. acidophilum* constituted of 14 identical α and 14 identical β subunits^28^. With the recent exception of the DNA-PKc analysis^44^ (469 kDa monomer), HDX-MS is usually performed either on rather small and simple systems^45^ or on homo-oligomeric complexes^46,47^, due to some technical limitations in protein digestion, peptide separation and data analysis. In comparison, our study encompasses 16 different ~25 kDa monomers. Despite this analytical challenge, we obtained an average sequence coverage per subunit of 89% for the 20S alone and 81% when in complex with PA28 regulators. The thorough statistical analysis of these data provides a wealth of information on the proteasome dynamics, binding interfaces and allosteric changes upon incorporation of the immuno-subunits or activation by PA28 regulators (**Fig. 8**).

**Figure 8.**
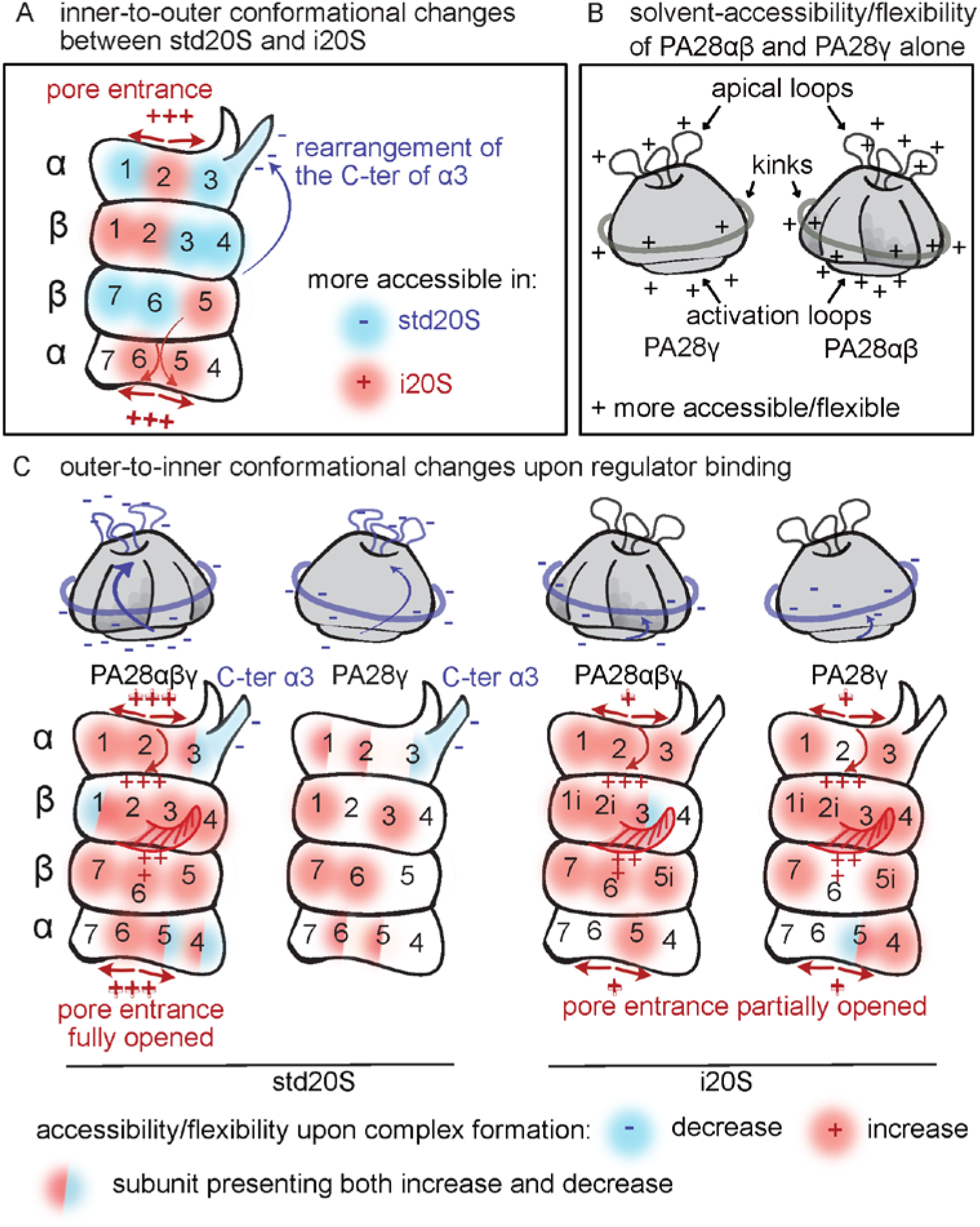
Schematic representation of our major finding. (**A**) Comparison of std20S *vs*. i20S deuteration profiles confirms an inner-to-outer allosteric change. (**B**) PA28 solvent accessibilities alone indicate similarities in their dynamic regions. (**C**) 20S- and PA28-specific outer-to-inner reorganizations occurring upon complex formation.

These conformational maps revealed the subtle rearrangement of the std/i20S α3 C-ter. More precisely, the residues 238-261 were less dynamic/accessible upon incorporation of the immuno-catalytic subunits (in purple **Fig. 8A**). The principal component analysis of the 20S high-resolution structures available in the Protein Data Bank (PDB) in 2014 (46 structures from yeast, 4 murine and 1 bovine) revealed that the 220-230 stretch of the mouse std20S and i20S α3 clustered with the yeast apo and peptide-bound forms, respectively^36^. We believe that the allosteric change occurring on α3 C-ter upon ligand binding to β5 in the yeast 20S could be similar to the change in solvent accessibility/dynamics observed between the std20S and the i20S conformational maps. The highly charged C-ter of α3 points towards the outer surface of the α-ring and has also been suggested to be the Insulin-Degrading-Enzyme binding site^48^. It could be involved in the binding of PA28αβ/γ to the std20S, or in its subsequent activation. Interestingly, the very end of α3 C-ter is absent from most human 20S proteasome structures, which confirms its high flexibility but also impedes any modeling of 20S/PA28 interaction. Unfortunately, this protruding C-ter α helix is not present in the cryo-EM structure of the 20S *Plasmodium falciparum* bound to its PA28 regulator, since it is 9 residues shorter than its human counterpart^30^. It could be characteristic of more evolved 20S proteasomes and tune their binding with different regulators.

The central pore of the 20S, on the other hand, was more flexible in the i20S *vs*. std20S (**Fig. 5D** and in orange **Fig. 8A**), as was the β-facing interface of the α-ring, especially α5 that directly faces the β5 (std20S) or β5i (i20S) catalytic subunit (**Fig. 5C**). We propose that the difference of dynamics between the two proteasomes observed at the catalytic sites could be transduced from the inner β-ring to the outer α-ring via α5 and lead to the conformational differences observed on their α3 C-ter (orange arrows **Fig. 8A**). Although HDX-MS does not allow the direct comparison of the standard and immuno-catalytic subunits (their sequences are different), our data suggests that the substrate pockets S1/S2 of β1 as well as the active site and S1 pocket of β5 are more dynamic than in their immuno-subunit counterparts. It has been shown that β1 PGPH activity is higher than the one of β1i and that β5i chymotrypsin activity is higher than the one of β5^49^, suggesting that we cannot directly correlate the substrate pocket flexibility/accessibility with activity. The slower RDU observed in β5i S1 pocket is in good agreement with the higher number of van der Waals interactions observed in mice^25^ and with the specific hydrogen bond that stabilizes the tetrahedral transition state during catalysis of β5i.

Interestingly, upon binding of any regulator to the 20S, we observed a similar destabilization of the α5 region that faces β5/5i, suggesting that the exact opposite mechanism (from the outer α-ring to the inner β-ring) can also take place upon 20S/PA28 interaction, as suggested earlier^28^.

Overall, our data indicate a general mechanism of PA28 activation by the 20S core proteasome. More precisely, the interaction of PA28 activation loops and C-ter with the 20S seems to trigger a long-range allosteric change at the top of the regulator (loops between helices 1/2 and/or 3/4) via a rearrangement of the N-ter kink of PA28α and PA28γ (**Fig. 8B**). Although present in the three PA28 subtypes, the activation loop protection was stronger in PA28α and PA28γ *vs*. PA28β; most probably explaining why “the affinity of the proteasome towards the PA28αβ complex is about two orders of magnitude higher than towards the homomeric PA28α and PA28β complexes”^50^. Indeed, several studies showed that both PA28α and PA28β were required for an optimum activation of the 20S by PA28αβ^26,27^.

We also provide crucial information on the loops located between helices 1/2, missing in every PA28 high-resolution structure or model. Since these are located at the entrance/exit of the activator, any change in their flexibility could modify the nature of both substrates and/or products of the proteasome complex. Indeed, it has been shown in PA28α to constrain substrates within the catalytic chamber of the core particle, thereby generating shorter peptides^51^. We show that these flexible loops, together with PA28 constriction sites (conserved lysine between alpha helices 3 and 4) are allosterically stabilized upon binding of the std20S but not the i20S (**Fig. 8B**). This could explain how the binding of different PA28 to different 20S subtypes alters the length/nature of the generated peptides. Our data also suggests that PA28 might modify the 20S products by inducing conformational changes in the proteasomal active sites. Indeed, our results show distinctive changes in dynamics detected down to the β-ring and around the catalytic site and/or substrate pockets upon regulator binding (**Fig. 8C**).

We observed a stronger protection of PA28αβ than PA28γ upon std20S binding (**Fig. 6** and **Fig 8B**). Globally, both PA28 were more protected upon interaction with the std20S than with the i20S (**Fig. S8-S9**). This was mirrored by regions on the i20S α-ring that were less protected than in the std20S upon incubation with both PA28, whereas previous studies showed that PA28αβ interaction was stronger with the i20S than with the std20S^52^ or equivalent^16^. We thus think that the protection cannot be directly correlated to relative binding affinities, especially because we monitor deuteration on protein complexes at steady-state (after 30 min incubation).

Although not similarly distributed, the global core proteasome destabilization upon complex formation was similar in all four conditions tested (**Fig. 7**), so we focused on the comparison of the most affected regions. We observed that each pair of PA28/20S complex underwent long-range conformational rearrangements in specific regions, as suggested recently^20^. Different sets of α N-ter were destabilized depending on the PA28/20S pair (in red **Fig.8C**), suggesting a gradual interaction-specific opening of the central pore. The β2 C-ter extending towards β3 was destabilized in all the conditions, except std20S/PA28γ. On the contrary, the region just before the α3 C-ter (217-236) was protected upon PA28 binding in all cases, except std20S/PA28αβ. Interestingly, this stretch is close to β3 74-86, which was also protected in all conditions tested except std20S/PA28αβ. These observations could provide a basis to understand how regulator binding propagates from the α-to the β-ring depending on the activator/20S pair.

Altogether, these data provide a unique resource informing on the allosteric activation of the 20S proteasome complexes that should soon be completed by upcoming structural and functional studies. We think that HDX-MS could be applied to characterize, not only the binding sites of many new proteasome inhibitors, but also to identify their potential allosteric effects on the 20S core complex. The same is true for the many PIPs that regulate proteasome function and are still poorly characterized from a structural point of view^53^. More broadly, this work demonstrates the ability of HDX-MS to investigate dynamic events on megadalton assemblies including ribosomal particles, inflammasomes and nucleosomes, to name a few.

## Methods

### Reagents

Unless stated otherwise, all reagents were purchased from Sigma-Aldrich. The std20S, i20S, PA28γ and PA28αβ were purchased from Enzo Life Science. Deuterium oxide, deuterium chloride and sodium deuteroxide solutions were from Euriso-top.

### Development of a HDX-MS pipeline dedicated to the comparative analysis of 20S proteasome/regulator complexes

In a classical HDX-MS workflow, the proteins of interest are digested online and the generated peptides are separated on a reverse phase column before MS analysis to monitor deuterium incorporation rates. The main challenges encountered when analysing heterogeneous complexes such as the proteasome by HDX-MS are threefold. First, the high number of peptides generated after digestion entailed specific optimization of their chromatographic separation. Second, we optimized the quenching step and injection parameters to reduce dead-volumes, as well as the acquisition method, in order to handle samples at low concentration (<1 μM). Finally, we developed a computational pipeline dedicated to the multi-dimensional analysis of HDX-MS analysis of large complexes. The resulting data were mapped to available 3D structures using our recently developed open-source web application HDX-Viewer^54^.

### Automated Hydrogen-Deuterium eXchange coupled to Mass Spectrometry (HDX-MS)

HDX-MS experiments were performed on a Synapt-G2Si (Waters Scientific, Manchester, UK) coupled to a Twin HTS PAL dispensing and labelling robot (LEAP Technologies, Carborro, NC, USA) via a NanoAcquity system with HDX technology (Waters, Manchester, UK). Each step was optimized to minimize sample loss and work with such a heterogeneous and diluted sample:

- Method in HDxDirector: The method recommended by Waters (HDX System Suitability Test) injects only 25% of the sample that is aspirated from the protein vial. In order to reduce sample loss, we 1) used a sample loop of 100 μl instead of 50 μl, 2) carefully reduced all the dead volumes and 3) used a stronger acid (500 mM glycine pH 2.3 instead of 50 mM K2HPO4, 50 mM KH2PO4, pH 2.3) in order to optimize the ratio of quenching volume (10% instead of 50%). With this workflow, we increased by more than threefold the amount of starting material injected (79%). 20S proteasomes were incubated alone or with a 2-fold molar excess of PA28, with final concentrations of 0.4 μM and 0.8 μM, respectively. 5.7 μL of protein were aspirated and 5.2 μL were diluted in 98.8 μL of protonated (peptide mapping) or deuterated buffer (20 mM Tris pH/pD 7.4, 1 mM EDTA, 1 mM DTT) and incubated at 20 °C for 0, 0.5, 1, 5, 10 and 30 min. 99 μL were then transferred to vials containing 11 μL of pre-cooled quenching solution (500 mM glycine at pH 2.3). For experiments involving PA28αβ, the quenching buffer was supplemented with 250 mM tris-(2-carboxyethyl) phosphine (TCEP) in order to reduce the disulphide bridge between Cys21 of chain α and Cys3 of chain β. After 30 sec. of quenching, 105 μL were injected into a 100 μL loop. Proteins were digested on-line with a 2.1 mm x 30 mm Poros Immobilized Pepsin column (Life Technologies/Applied Biosystems, Carlsbad, CA, USA). The temperature of the digestion room was set at 15°C.
- Chromatographic run In order to cope with the unusual sample heterogeneity, the runtime of the chromatographic separation (12 min) was doubled compared to the one used for smaller protein complexes (6 min). Peptides were desalted for 3 min on a C18 pre-column (Acquity UPLC BEH 1.7 μm, VANGUARD) and separated on a C18 column (Acquity UPLC BEH 1.7 μm, 1.0 x 100 mm) by the following gradient: 5% to 35% buffer B (100% acetonitrile, 0.2% formic acid) for 12 min, 35% to 40% for 1 min, 40% to 95% for 1 min, 2 min at 95% followed by 2 cycles of 5% to 95% for 2 min and a final equilibration at 5% buffer A (5% acetonitrile, 0.2% formic acid) for 2min. The total runtime was 25 min. The temperature of the chromatographic module was set at 4°C. Experiments were run in triplicates and the protonated buffer was injected between each triplicate to wash the column and avoid cross-over contamination.
- MS acquisition The acquisitions were performed in positive and resolution mode in the m/z range 50 to 2000 Th. The sample cone and capillary voltages were set at 30 V and 3 kV, respectively. The analysis cycles for non-deuterated samples alternated between a 0.3 sec low energy scan (Trap and Transfer collision energies set to 4 V and 2 V, respectively), a 0.3 sec high energy scan (Ramp Trap and Transfer collision energies set to 18 V to 40 V and 2 V to 2 V, respectively) and a 0.3 sec lockspray scan (0.1 μM [Glu1]-Fibrinopeptide in 50% acetonitrile, 50% water and 0.2% formic acid infused at 10 μL/min). The lockspray trap collision energy was set at 32 V and a GFP scan of 0.3 sec is acquired every min. In order to double the signal intensity of deuterated peptides, deuterated samples were acquired only with the low energy and lockspray functions.
- Data analysis Peptide identification was performed with ProteinLynx Global SERVER (PLGS, Waters, Manchester, UK) based on the MS^E^ data acquired on the non-deuterated samples. The MSMS spectra were searched against a home-made database containing sequences from the 17 std20S and i20S subunits, PA28α, PA28β, PA28γ and pepsin from *Sus scrofa*. Peptides were filtered in DynamX 3.0 with the following parameters: peptides identified in at least 2 replicates, 0.2 fragments per amino-acid, intensity threshold 1000. The quantitative analysis of deuteration kinetics was performed using the statistical package R (R Development Core Team, 2012; http://www.R-project.org/) on the corresponding MS intensities. The deuterium uptakes of each ion for each time point were calculated based on the theoretical maximum, considering that all amino-acids (except proline residues and the first amino-acid or each peptide) were deuterated, and then averaged (weight = intensity) to get a value of relative deuterium uptake (RDU) per peptide sequence/condition/time point (see **Fig. S2, S6, S10** and **Table S2,S6**). To identify the protein regions that presented conformational changes in complex *vs*. alone, we performed an ANOVA (Anova(), type = “III”, singular.ok = T) followed by Benjamini Hochberg correction of the *P*-value. For each comparison, we considered significantly regulated the peptides with a corrected *P*-value ≤ 0.01 and an absolute difference of RDU above 0.01569 (4 times the mean absolute difference of RDU in the entire data set) for 3 successive time points (**Table S4**). The corresponding volcano plots are presented in **Fig. S5, S9, S12** and all the regions statistically regulated in all the differential HDX-MS analysis are listed in **Table S3**. The RDU and differences of RDU (for protein alones or comparison between conditions, respectively) were consolidated using the values of the smallest peptide to increase the spatial resolution (see consolidation heatmaps in **Fig. S3, S4, S7, S8, S11**). These data (per time point or sum of all time points) were then used for structural data mining with the recently developed open-source web application called HDX-Viewer^54^ that allows to directly plot and visualize the whole data set (14 different subunits per proteasome core particle) on the proteasome 3D structure. The scripts corresponding to the quality control and statistical analysis of this data set are available at zenodo.org with the DOI 10.5281/zenodo.3769174 under the Creative Commons Attribution 4.0 International licence. All molecular representations were generated in UCSF ChimeraX version: 0.9 (2019-06-06)^55^.

### Modeling

The PA28γ model was generated using the structure of human PA28α (PDB:1AV0) as a template with the SWISS-MODEL server^39^. This model misses the first 3 and the last 7 amino-acids, as well as the 64-106 stretch.

### Data availability

The R scripts of the analysis, with the fasta files and output tables and cxc files are freely available at zenodo.org with the DOI 10.5281/zenodo.3769174 under the Creative Commons Attribution 4.0 International licence.

The mass spectrometry proteomics data have been deposited to the ProteomeXchange Consortium via the PRIDE^56^ partner repository with the data set identifier PXD018921.

## Acknowledgments

This work was supported by the French Ministry of Research (ANR-ProteasoRegMS to J.M and Investissements d’Avenir Program, Proteomics French Infrastructure, ANR-10-INBS-08 to O.B.-S.), the Fonds Européens de Développement Régional Toulouse Métropole and the Région Midi-Pyrénées (O.B.-S.) and the Novo Nordisk Foundation (NNF14CC0001).

## Author contributions

All authors contributed to editing the manuscript and figures. J.L. performed the HDX-MS optimization and data acquisition, model building and data analysis. J.P. performed the HDX-MS optimization, data acquisition and data analysis. D.Z. performed experiments. M.L.P. analyzed the data and wrote the paper. Odile Schiltz provided critical input. M.P.B. provided critical expertise on the proteasome. J.M. designed, performed and supervised the experiments, data and structural analysis, and wrote the paper.

## Supplementary Information

**Figure S1. Sequence coverage and number of peptides obtained upon pepsin digestion of the 20S and PA28 subunits**. The bars present the sequence coverage (top) and number of peptides (bottom) obtained for each subunit in the samples with the proteins alone (black) or in complex (grey).

**Figure S2. Deuteration uptake curves obtained for each peptide of each subunit of the std20S and i20S**. Each plot corresponds to a single peptide. Its sequence, mass and location in the protein sequence (start - stop) are indicated in the plot titles. Points are the RDU of each replicate, lines are their mean. These data can be found in **Table S2**.

**Figure S3. Deuteration heatmaps of each α and β subunits of the 20S subunits, alongside their peptide sequence**. Relative deuterium uptakes between 0.5 and 30 min color-coded from 0% (white) to 60% (blue). Non-detected regions are colored in black. The solid lines correspond to the peptides from which the RDU values were reported in the consolidated data, the dashed lines correspond to portions of peptides that were not reported because the same sequence portion was covered by a shorter peptide. This representation is explained in **Fig. 1B**.

**Figure S4. Differential deuteration heatmaps of each α and β subunits of the std20S *vs*. i20S**. Differential relative uptakes between the i20S and the std20S from 0.5 to 30 min color coded from - 10% (blue, more flexible/dynamic in the i20S) to +10% (red, more flexible/dynamic in the std20S). Non-detected regions are colored in black. Peptide coverage is represented by lines above the heatmaps. The solid lines correspond to the peptides from which the RDU values were reported in the consolidated data, the dashed lines correspond to portions of peptides that were not reported because the same sequence portion was covered by a shorter peptide. This representation is explained in **Fig. 1B**. Additionally, the peptides that were considered significantly regulated according to the thresholds indicated in **Fig. 1D** are colored in yellow.

**Figure S5. Volcano plots showing the peptides significantly different when comparing the deuteration of the std20S *vs*. i20S**. Each peptide is plotted in function of its statistical significance (y-axis: −log10(BH-corrected p-value)) and its difference of accessibility in the std20S *vs*. i20s (x-axis: sum of RDU differences across time points). The peptides passing the statistical thresholds presented in **Fid. 1D** are colored: peptides that are statistically more flexible/dynamic in the i20S are represented in blue and those more flexible/dynamic in the std20S in red. These data can be found in **Table S4**.

**Figure S6. Deuteration uptake curves obtained for each peptide of PA28α, PA28β and PA28γ alone and in complex with the std20S and i20S**. Each plot corresponds to a single peptide. Its sequence, mass and location in the protein sequence (start - stop) are indicated in the plot titles. Points are the RDU of each replicate, lines are their mean. These data can be found in **Tables S2,S6**.

**Figure S7. Deuteration heatmaps of PA28α, PA28β and PA28γ alongside their peptide sequence**. Relative deuterium uptakes between 0.5 and 30 min color-coded from 0% (white) to 60% (blue). Non-detected regions are colored in black. The solid lines correspond to the peptides from which the RDU values were reported in the consolidated data, the dashed lines correspond to portions of peptides that were not reported because the same sequence portion was covered by a shorter peptide. This representation is explained in **Fig. 1B**. The residues corresponding to the disordered loops located between helices 1/2 and that are missing in every high-resolution structures are highlighted with red rectangles.

**Figure S8. Differential deuteration heatmaps of each PA28 subunit alone *vs*. in complex with the std20S or i20S**. Differential relative uptakes from 0.5 to 30 min color coded from −30% (blue, less flexible/dynamic in the complex) to +30% (red, more flexible/dynamic in the complex) for PA28αβ and from −10% (blue) to +10% (red) for PA28γ. Peptide coverage is represented by lines above the heatmaps. The solid lines correspond to the peptides from which the RDU values were reported in the consolidated data, the dashed lines correspond to portions of peptides that were not reported because the same sequence portion was covered by a shorter peptide. This representation is explained in **Fig. 1B**. Additionally, the peptides that were considered significantly regulated according to the thresholds indicated in **Fig. 1D** are colored in yellow.

**Figure S9. Volcano plots showing the peptides significantly different when comparing the deuteration of the PA28 subunits alone *vs*. in complex with the std20S or i20S**. Each peptide is plotted in function of its statistical significance (y-axis: −log10(BH-corrected p-value)) and its difference of accessibility in PA alone or in complex with the 20S (x-axis: sum of RDU differences across time points). The peptides passing the statistical thresholds presented in **Fid. 1D** are colored: peptides that are statistically less flexible/dynamic in the PA28/20S complex are represented in blue and those more flexible/dynamic in the PA28/20S complex in red. These data can be found in **Table S4**.

**Figure S10. Deuteration uptake curves obtained for each peptide of each subunit of the std20S and i20S alone or in complex with PA28αβ or PA28γ**. Each plot corresponds to a single peptide. Its sequence, mass and location in the protein sequence (start - stop) are indicated in the plot titles. Points are the RDU of each replicate, lines are their mean. These data can be found in **Table S6**.

**Figure S11. Differential deuteration heatmaps of each α and β subunits of the std20S and i20S alone *vs*. in complex with PA28αβ or PA28γ**. Differential relative uptakes from 0.5 to 30 min color coded from −10% (blue, less flexible/dynamic in the complex) to +10% (red, more flexible/dynamic in the complex). Peptide coverage is represented by lines above the heatmaps. The solid lines correspond to the peptides from which the RDU values were reported in the consolidated data, the dashed lines correspond to portions of peptides that were not reported because the same sequence portion was covered by a shorter peptide. This representation is explained in **Fig. 1B**. Additionally, the peptides that were considered significantly regulated according to the thresholds indicated in **Fig. 1D** are colored in yellow.

**Figure S12. Volcano plots showing the peptides significantly different when comparing the deuteration of the std20S and i20S alone or in complex with PA28αβ or PA28γ**. Each peptide is plotted in function of its statistical significance (y-axis: −log10(BH-corrected p-value)) and its difference of accessibility in PA alone or in complex with the 20S (x-axis: sum of RDU differences across time points). The peptides passing the statistical thresholds presented in **Fid. 1D** are colored: peptides that are statistically less flexible/dynamic in the PA28/20S complex are represented in blue and those more flexible/dynamic in the PA28/20S complex in red. These data can be found in **Table S4**.

**Table S1. Sequence coverage and number of peptides obtained upon pepsin digestion of the 20S and PA28 subunits**.

**Table S2. Relative deuterium uptakes calculated for each peptide in the data set of the proteins alone**.

**Table S3. List of all the amino-acid stretches statistically regulated in all the differential HDX-MS analysis**

**Table S4. Outputs of the statistical analysis**.

**Table S5. Identity matrix generated by Clustal2.1 showing the percentage of identity between *Pf*PA28 and human PA28α, PA28β and PA28γ**.

**Table S6. Relative deuterium uptakes calculated for each peptide in the data set of the proteins alone and in complex**.

**Additional Material (PRIDE repository**)

**.raw** data of each LC-MS(MS) acquisition.

**.fasta** file used for the MSMS interrogation in PLGS.

**IA.csv** output files from PLGS used for the sequence coverage of non-deuterated proteins in DynamX.

**.CSV** output files from our analysis in DanymX used for the kinetic plots (**Fig. S2, S6, S10**).

**.pml** output files from our analysis in DynamX, to be used with the corresponding .pdb files in HDX-Viewer.

**.pdb** files of the std20S (5LE5), the i20S (6E5B), PA28αβ (5MX5) and PA28γ (Swiss-Model) used for the 3D analysis in HDX-Viewer with the corresponding .pml files.

**.csx** sessions for the direct 3D visualization of the deuteration uptakes in ChimeraX.

