## Supplementary figures and images for "Conformational maps of human 20S proteasomes reveal PA28- and immuno-dependent inter-ring crosstalks"

### Fig. S1

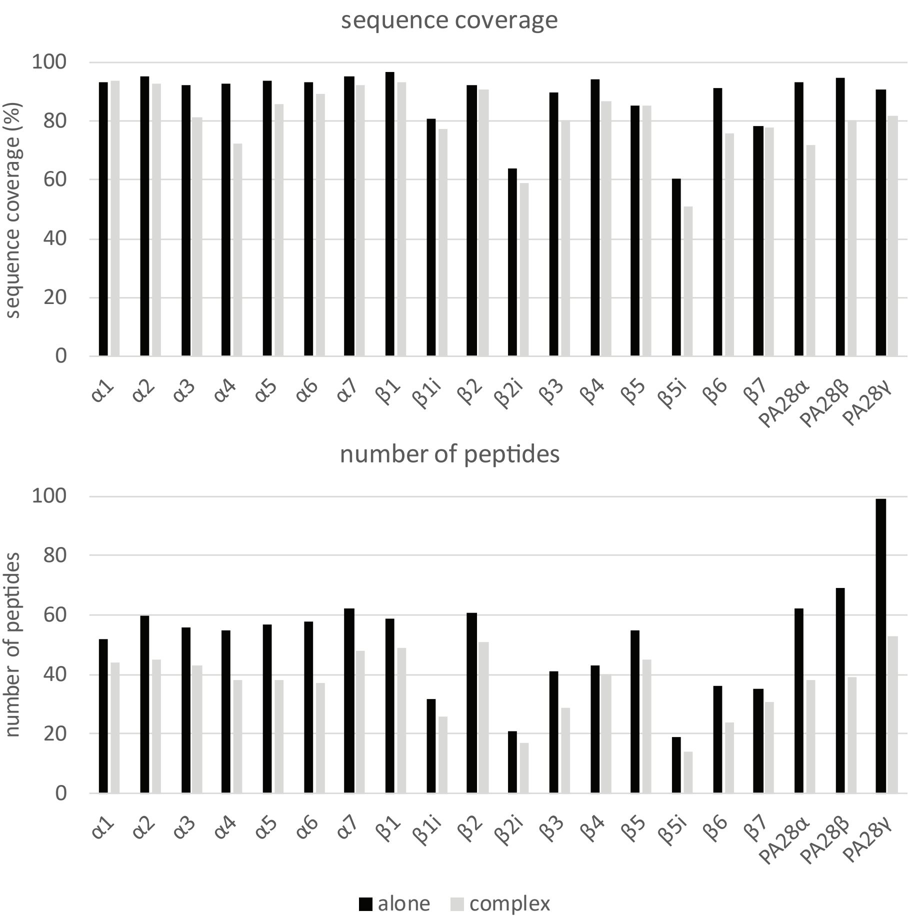

### Fig. S7

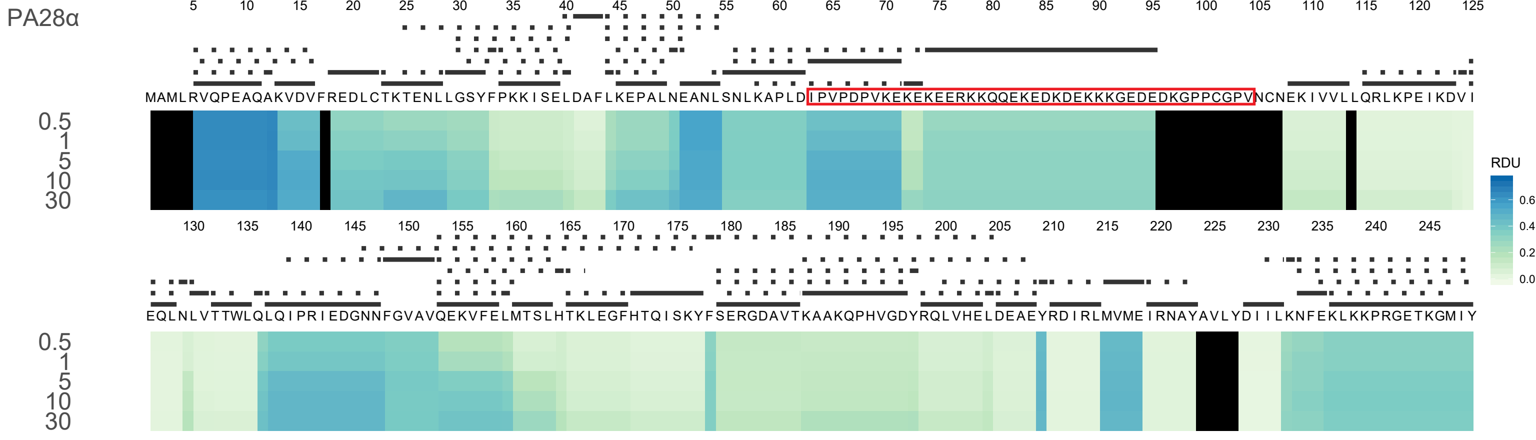

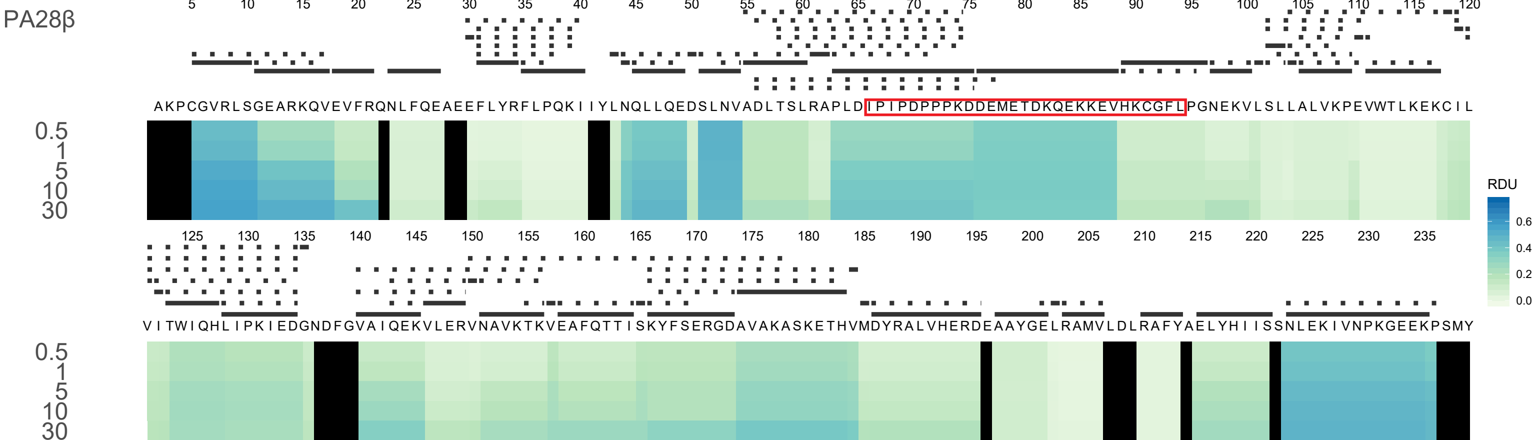

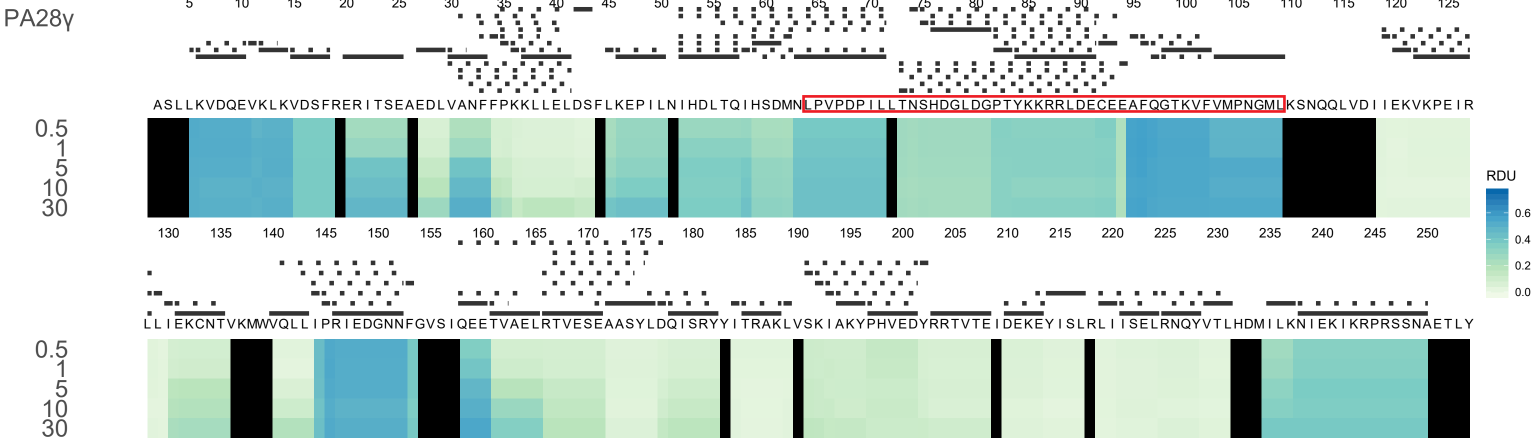

### Fig. S8

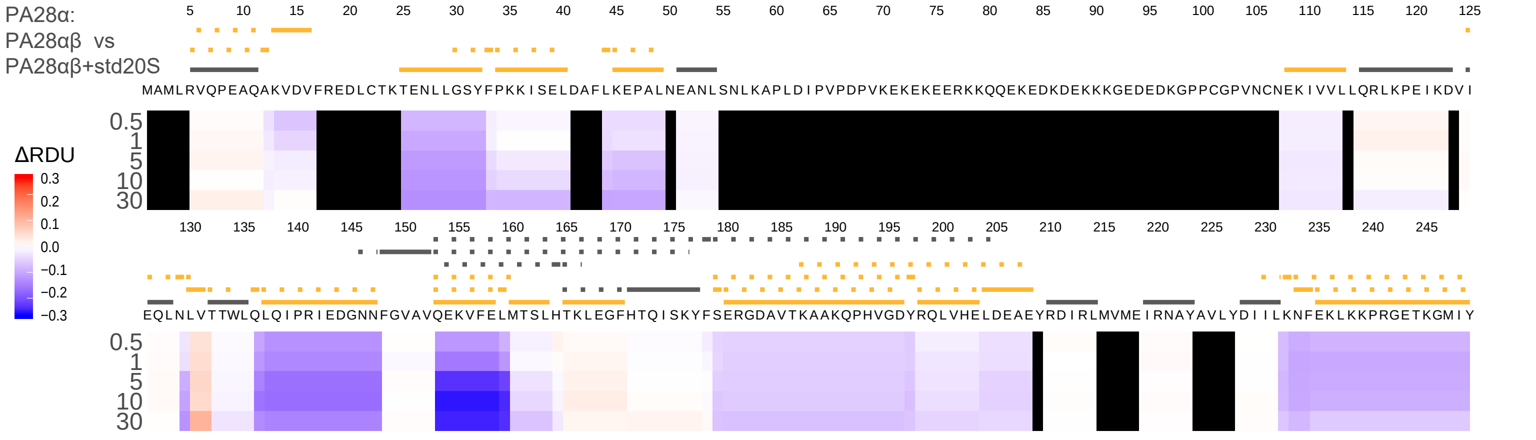

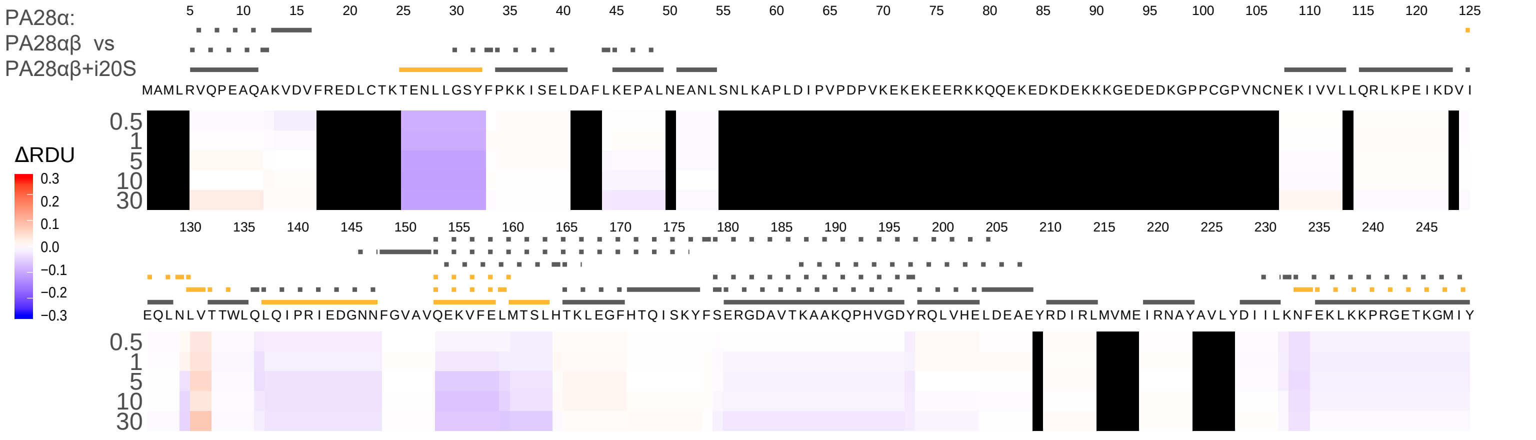

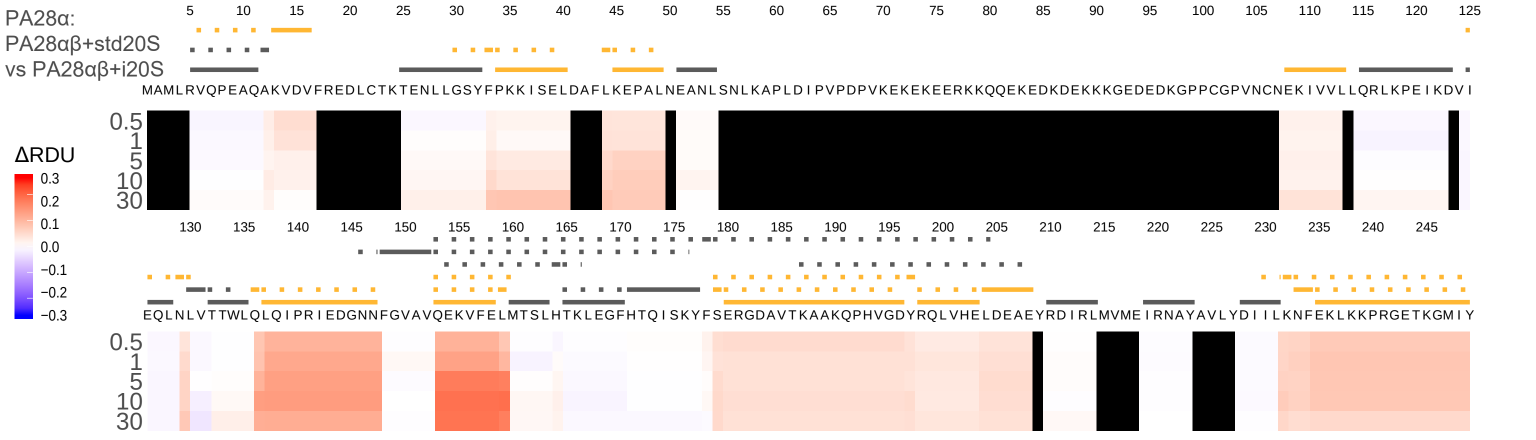

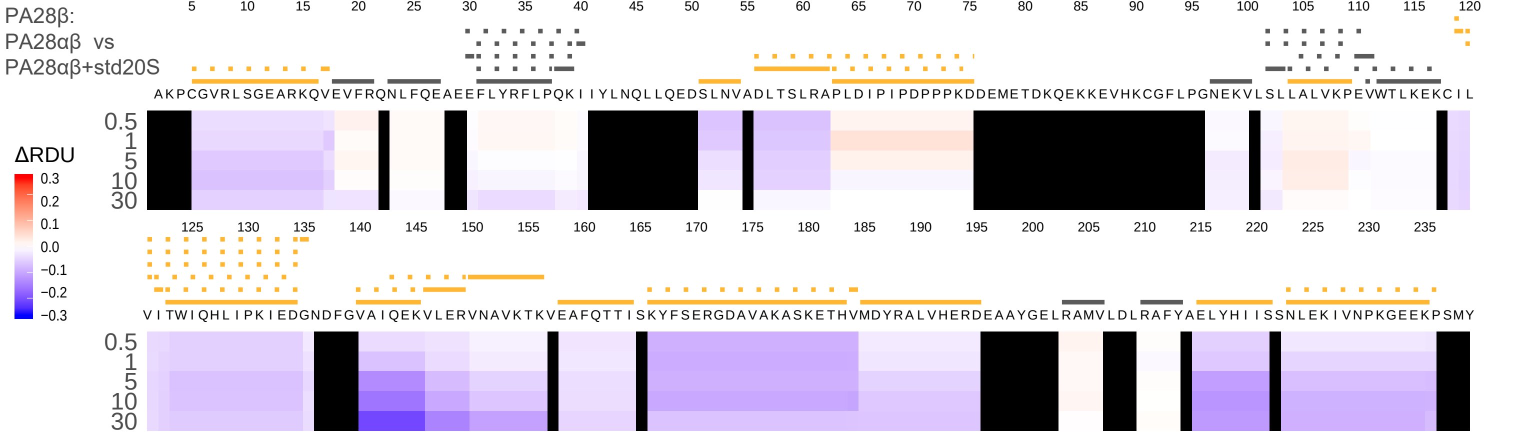

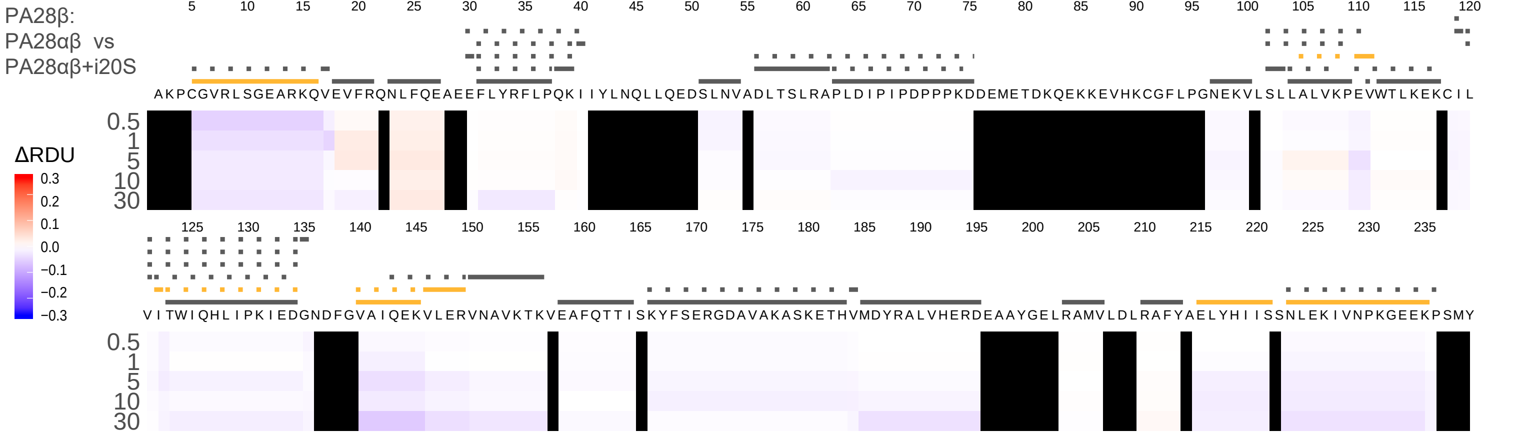

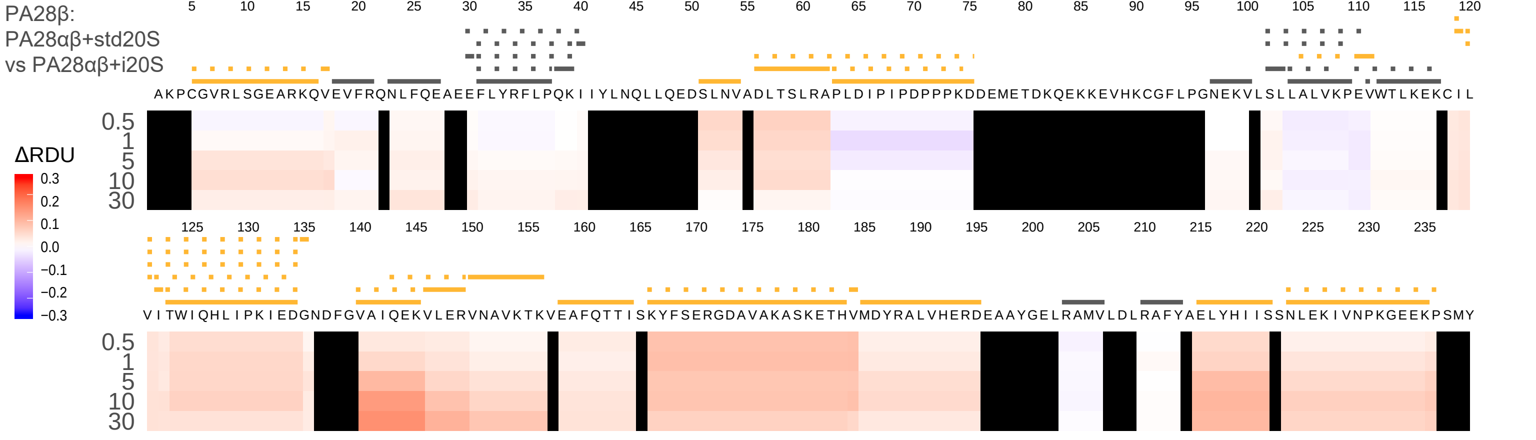

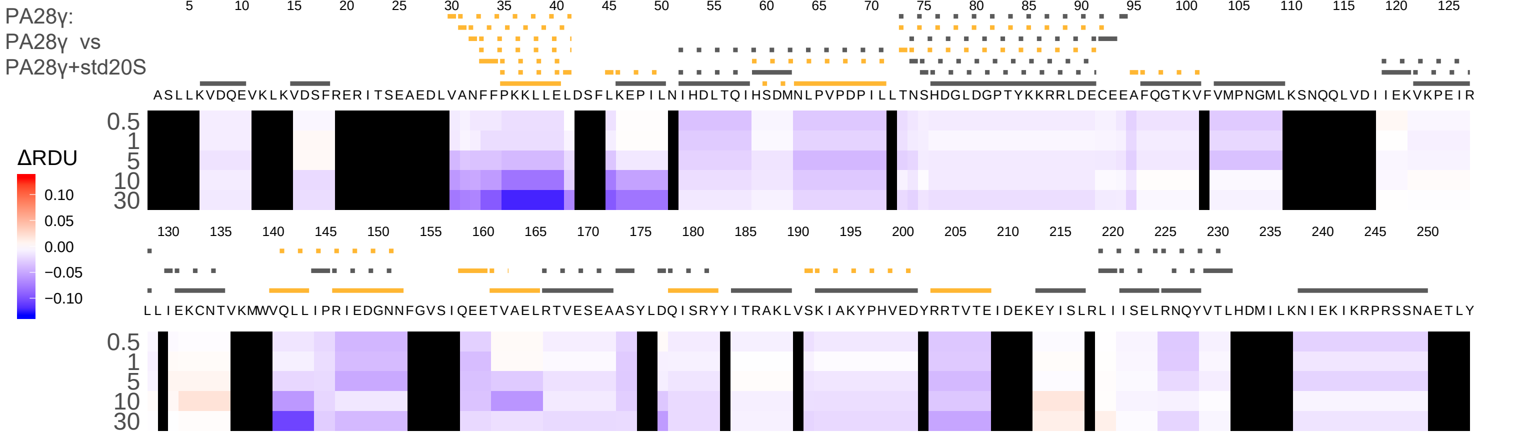

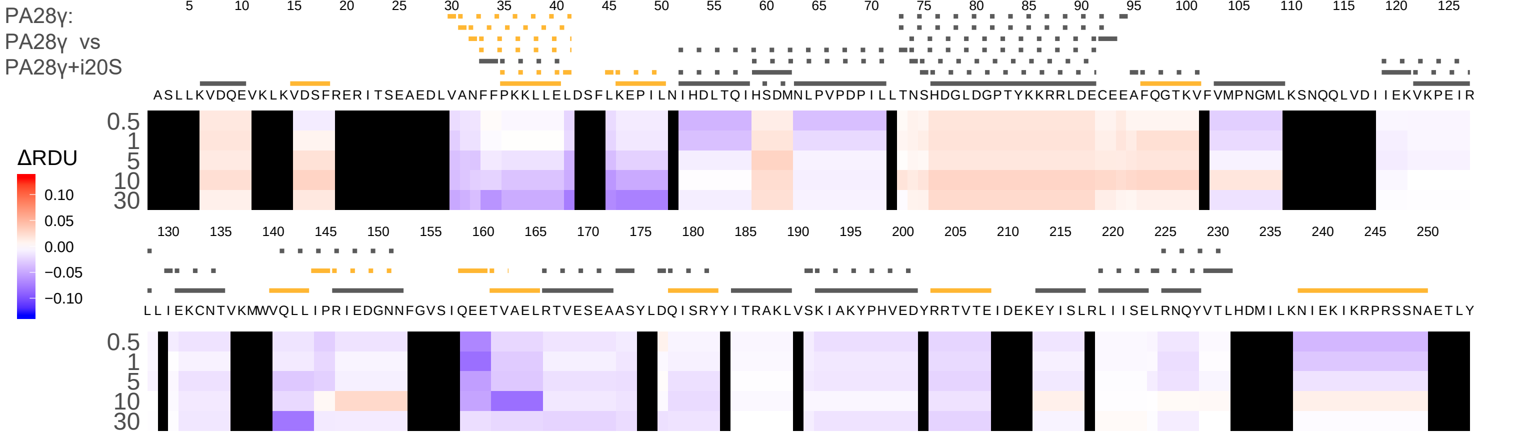

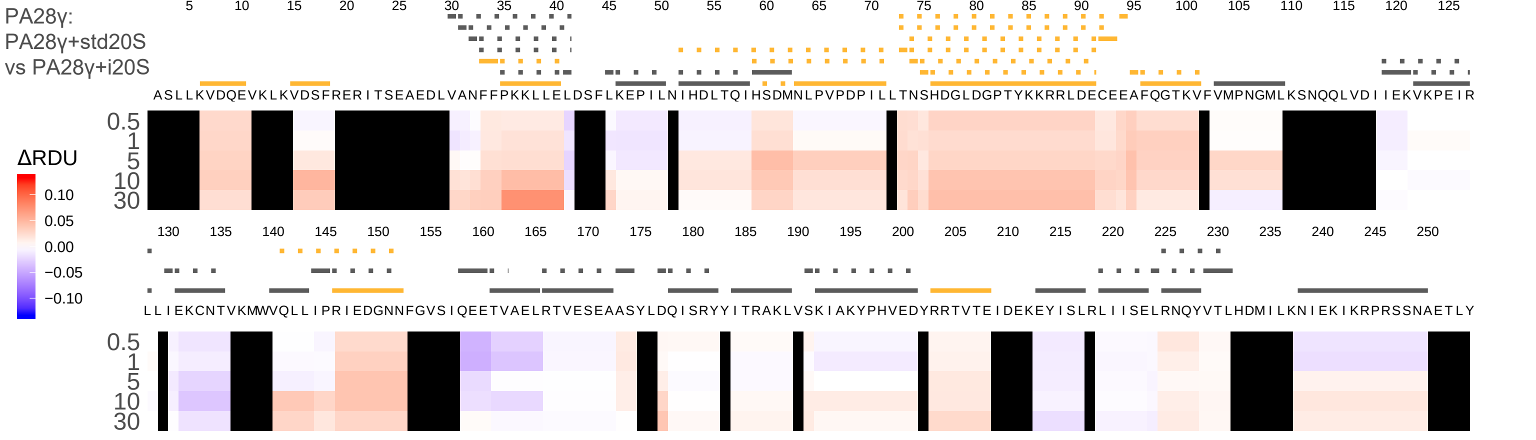

### Fig. S9

PA28α: PA28αβ + std20S Vs PA28αβ

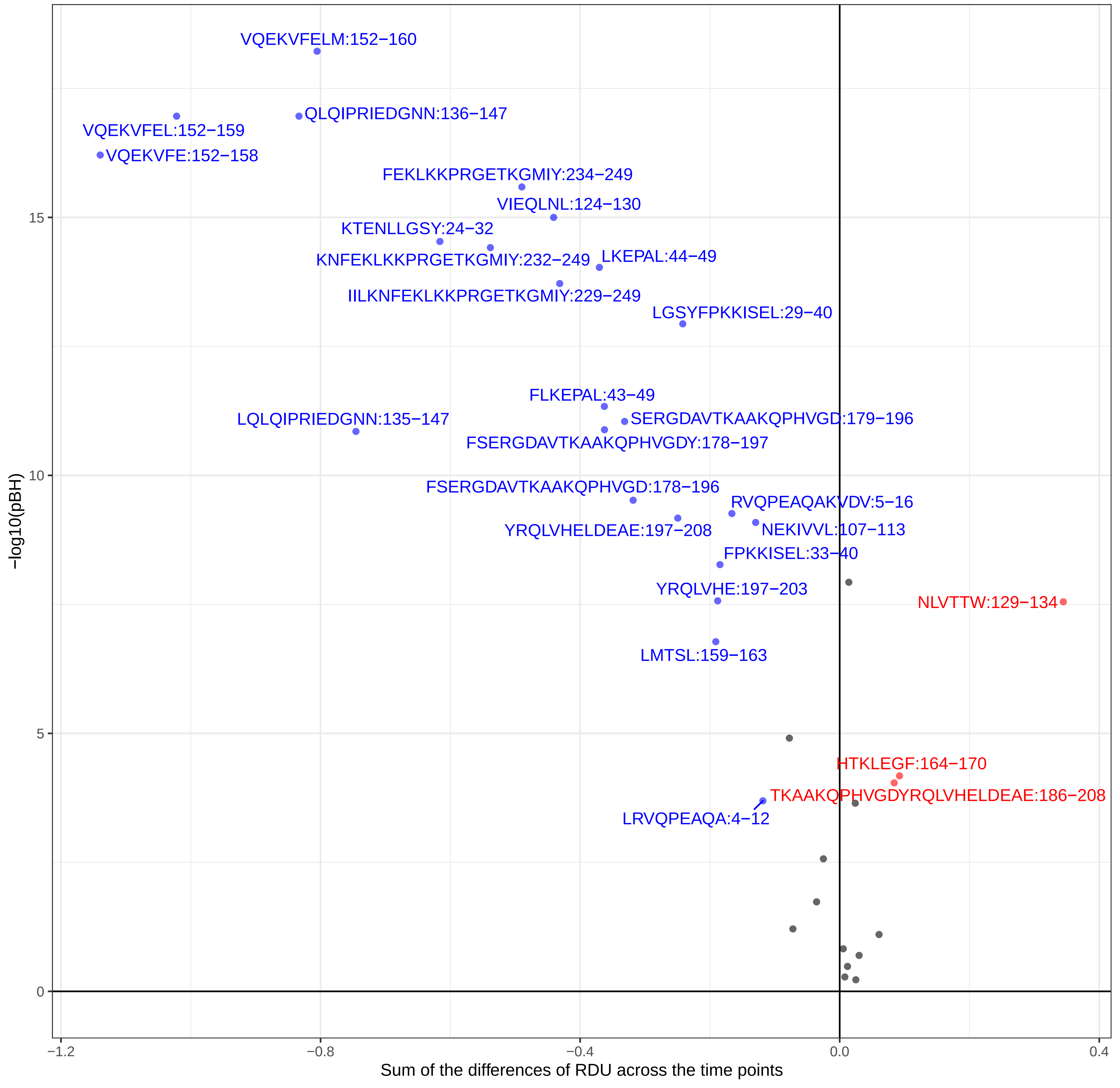

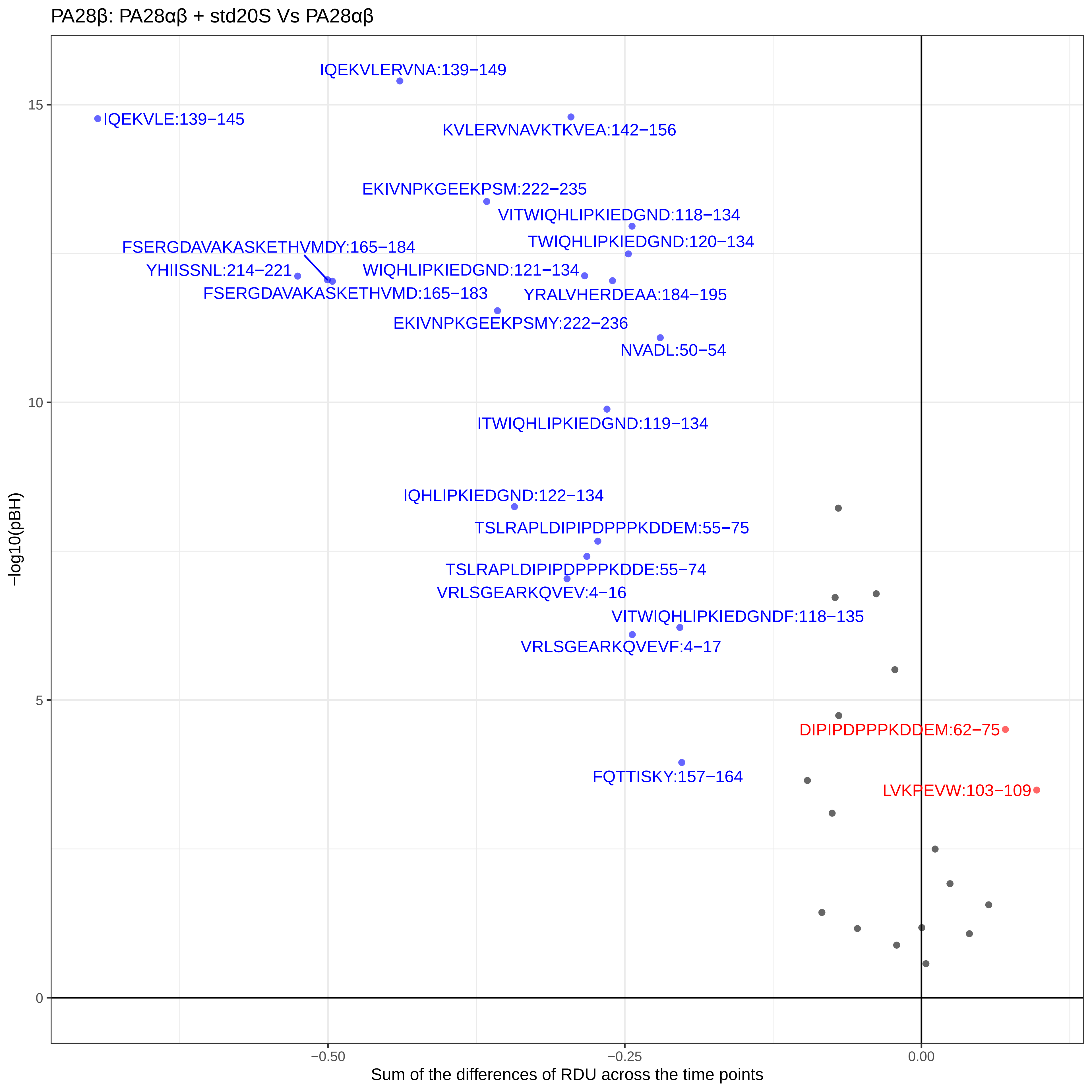

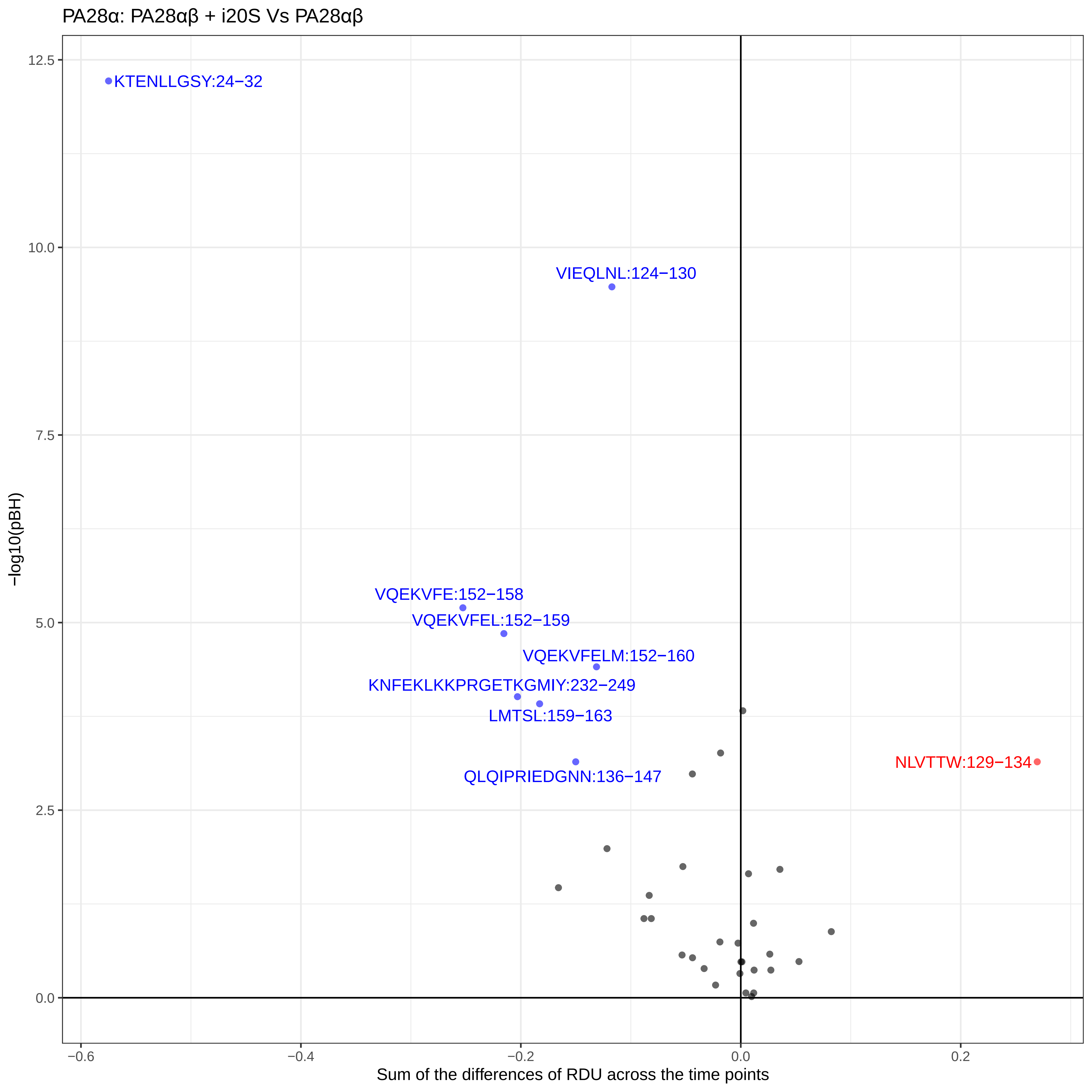

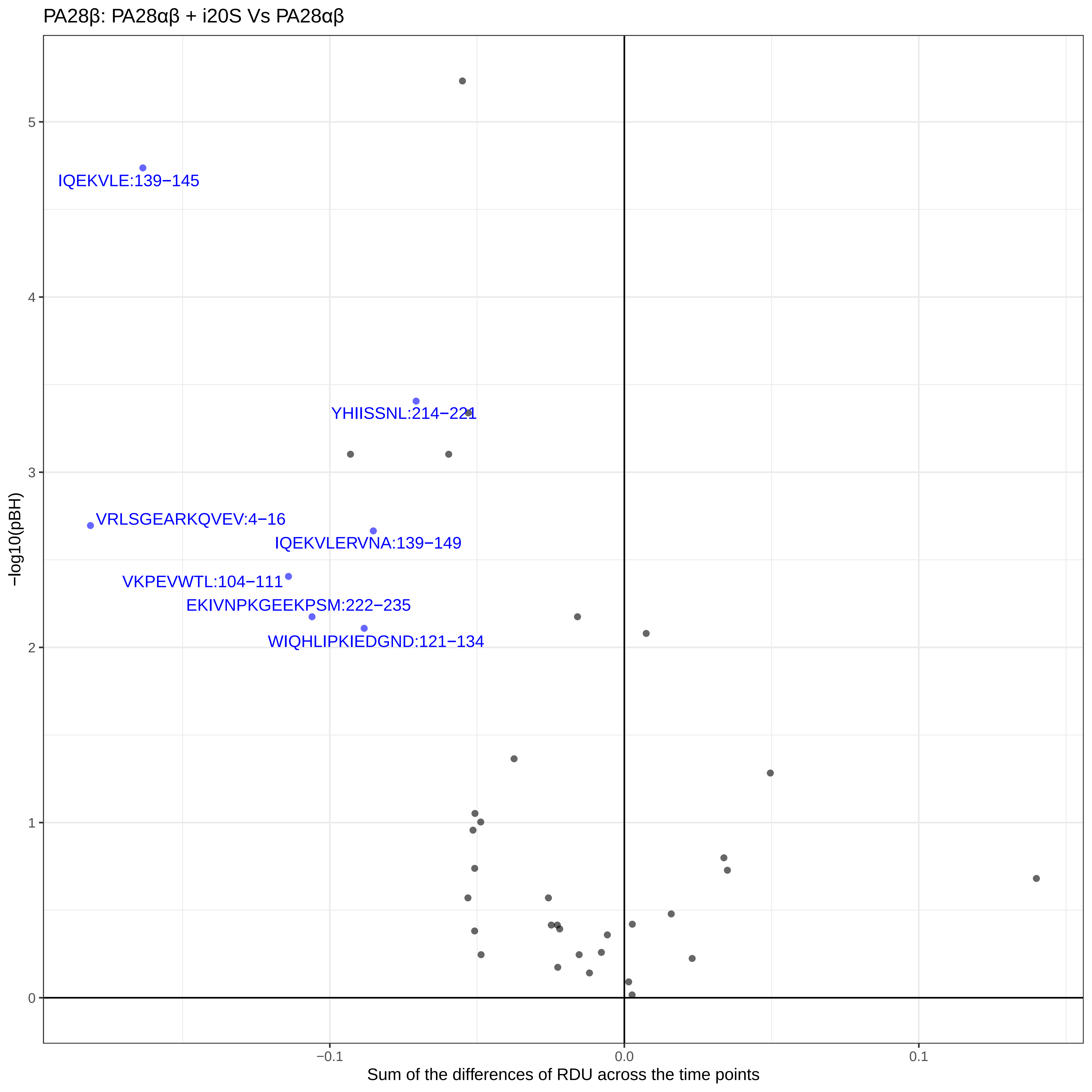

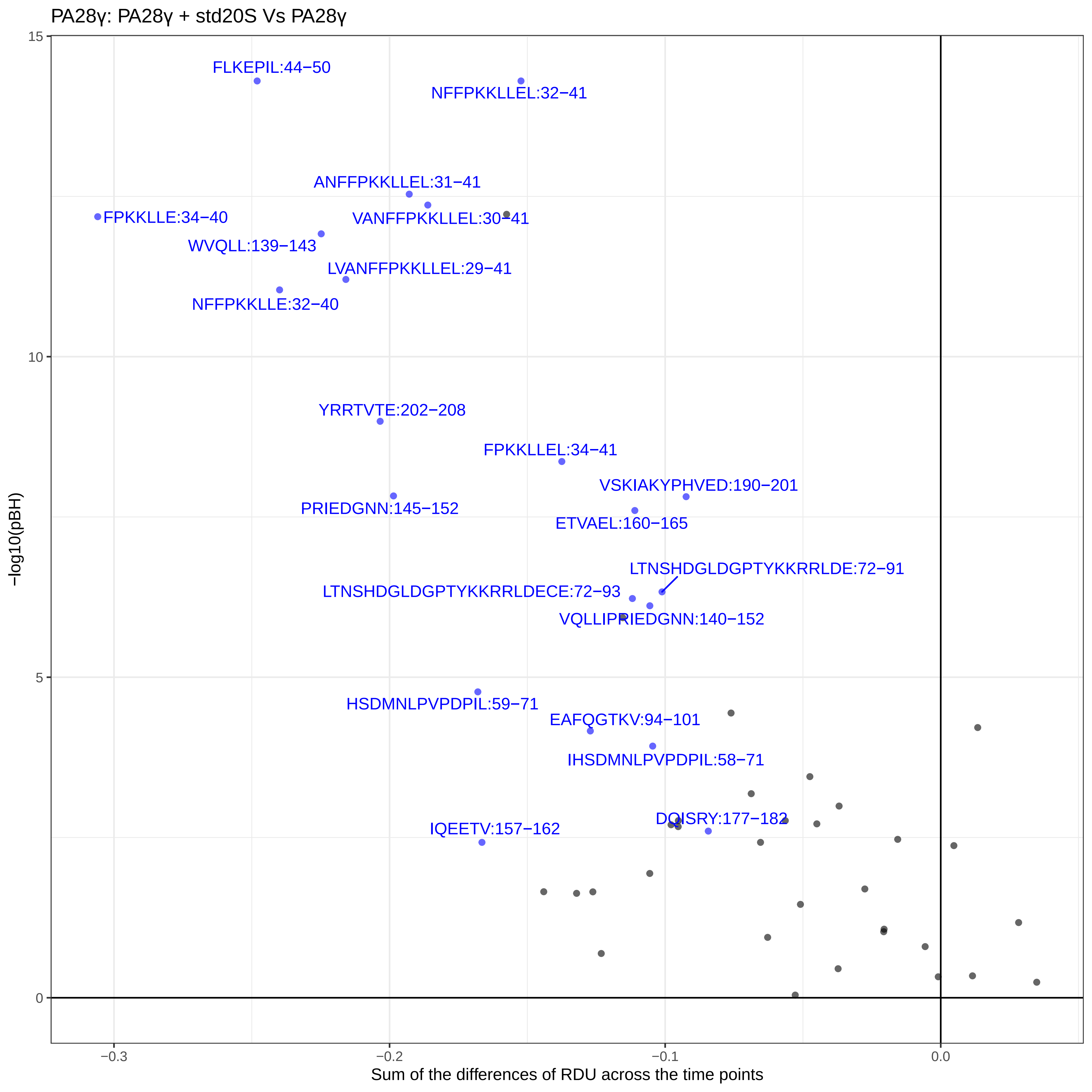

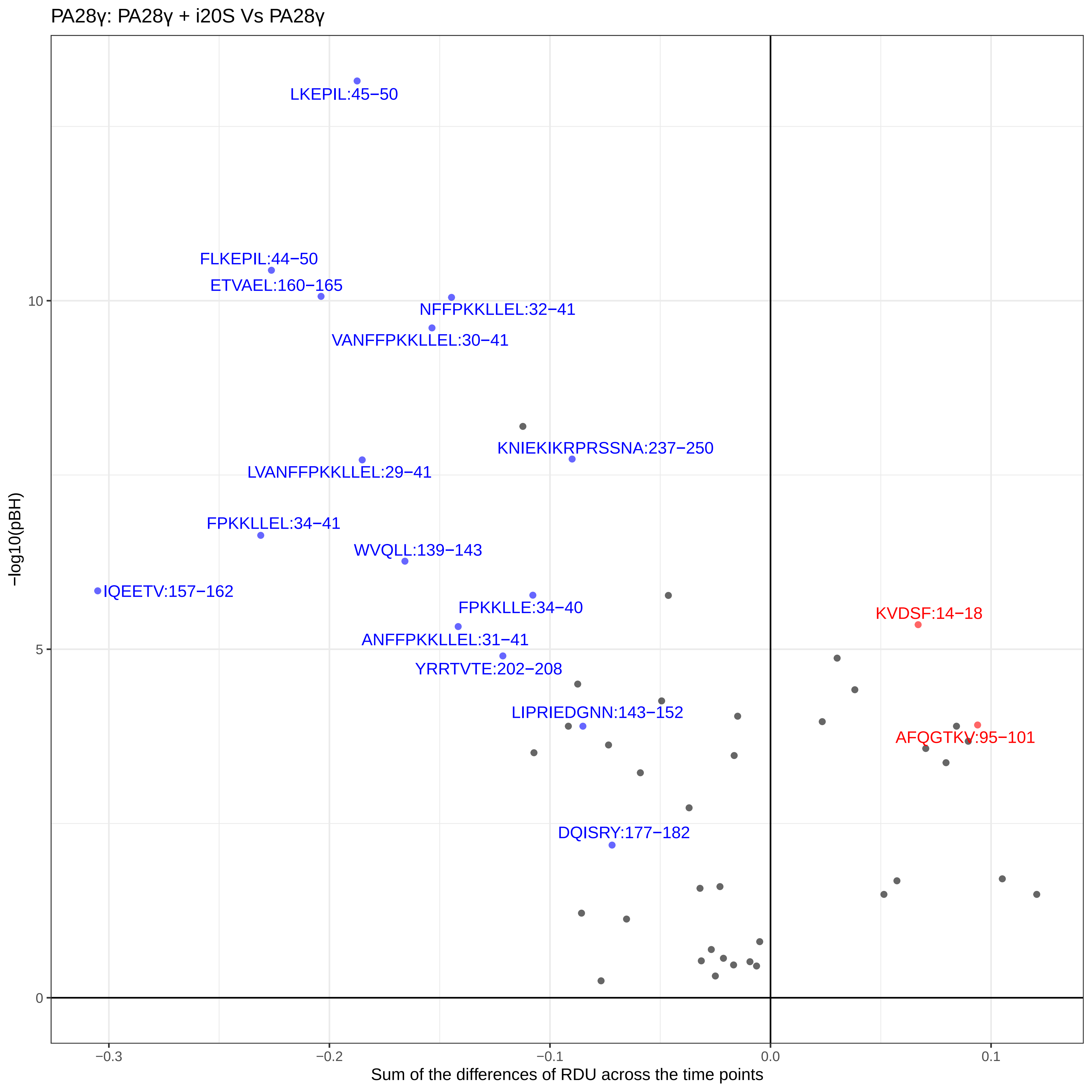
