## Supplementary material for "Conformational maps of human 20S proteasomes reveal PA28- and immuno-dependent inter-ring crosstalks": Fig. S2

α1

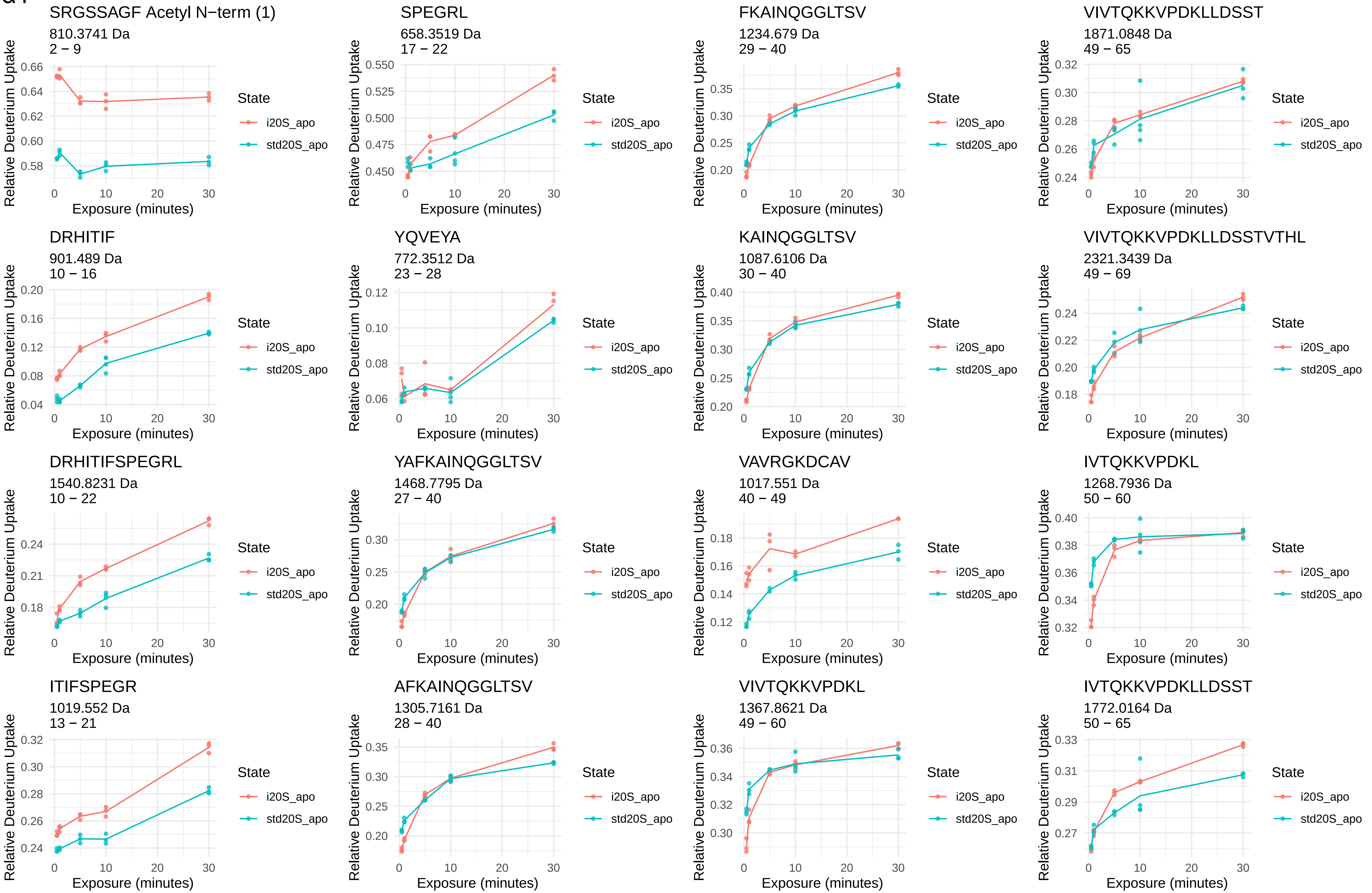
SRGSSAGF Acetyl N−term (1)
810.3741 Da
2 − 9
SPEGRL
658.3519 Da
17 − 22
FKAINQGGLTSV
1234.679 Da
29 − 40
VIVTQKKVPDKLLDSST
1871.0848 Da
49 − 65
DRHITIF
901.489 Da
10 − 16
YQVEYA
772.3512 Da
23 − 28
KAINQGGLTSV
1087.6106 Da
30 − 40
VIVTQKKVPDKLLDSSTVTHL
2321.3439 Da
49 − 69
DRHITIFSPEGRL
1540.8231 Da
10 − 22
YAFKAINQGGLTSV
1468.7795 Da
27 − 40
VAVRGKDCAV
1017.551 Da
40 − 49
IVTQKKVPDKL
1268.7936 Da
50 − 60
ITIFSPEGR
1019.552 Da
13 − 21
AFKAINQGGLTSV
1305.7161 Da
28 − 40
VIVTQKKVPDKL
1367.8621 Da
49 − 60
IVTQKKVPDKLLDSST
1772.0164 Da
50 − 65
Relative Deuterium Uptake
Exposure (minutes)
State
i20S_apo
std20S_apo

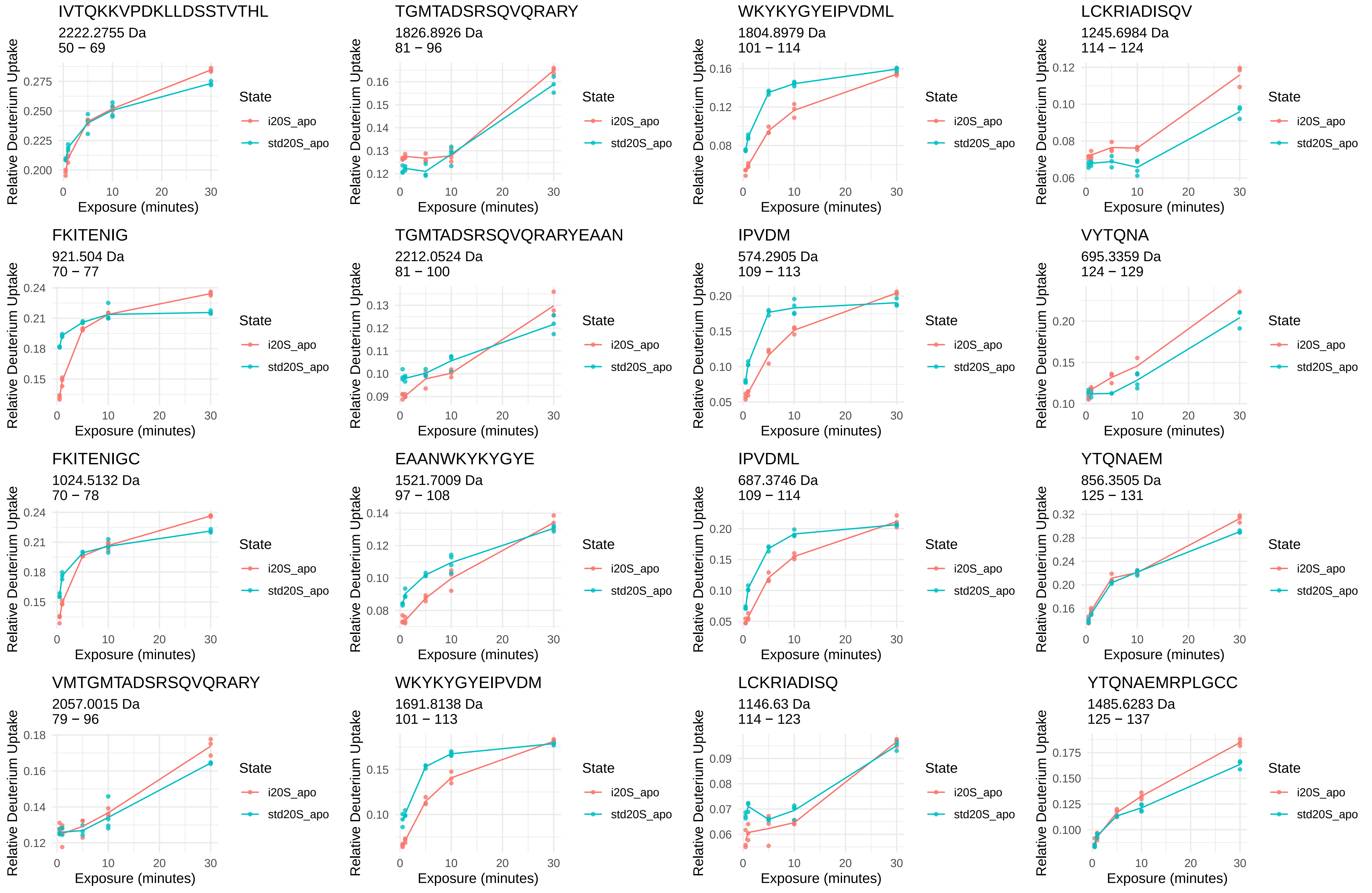

IVTQKKVPDKLLDSSTVTHL
2222.2755 Da
50 − 69
TGMTADSRSQVQRARY
1826.8926 Da
81 − 96
WKYKYGYEIPVDML
1804.8979 Da
101 − 114
LCKRIADISQV
1245.6984 Da
114 − 124
FKITENIG
921.504 Da
70 − 77
TGMTADSRSQVQRARYEAAN
2212.0524 Da
81 − 100
IPVDM
574.2905 Da
109 − 113
VYTQNA
695.3359 Da
124 − 129
FKITENIGC
1024.5132 Da
70 − 78
EAANWKYKYGYE
1521.7009 Da
97 − 108
IPVDML
687.3746 Da
109 − 114
YTQNAEM
856.3505 Da
125 − 131
VMTGMTADSRSQVQRARY
2057.0015 Da
79 − 96
WKYKYGYEIPVDM
1691.8138 Da
101 − 113
LCKRIADISQ
1146.63 Da
114 − 123
YTQNAEMRPLGCC
1485.6283 Da
125 − 137
Relative Deuterium Uptake
Exposure (minutes)
State
i20S_apo
std20S_apo

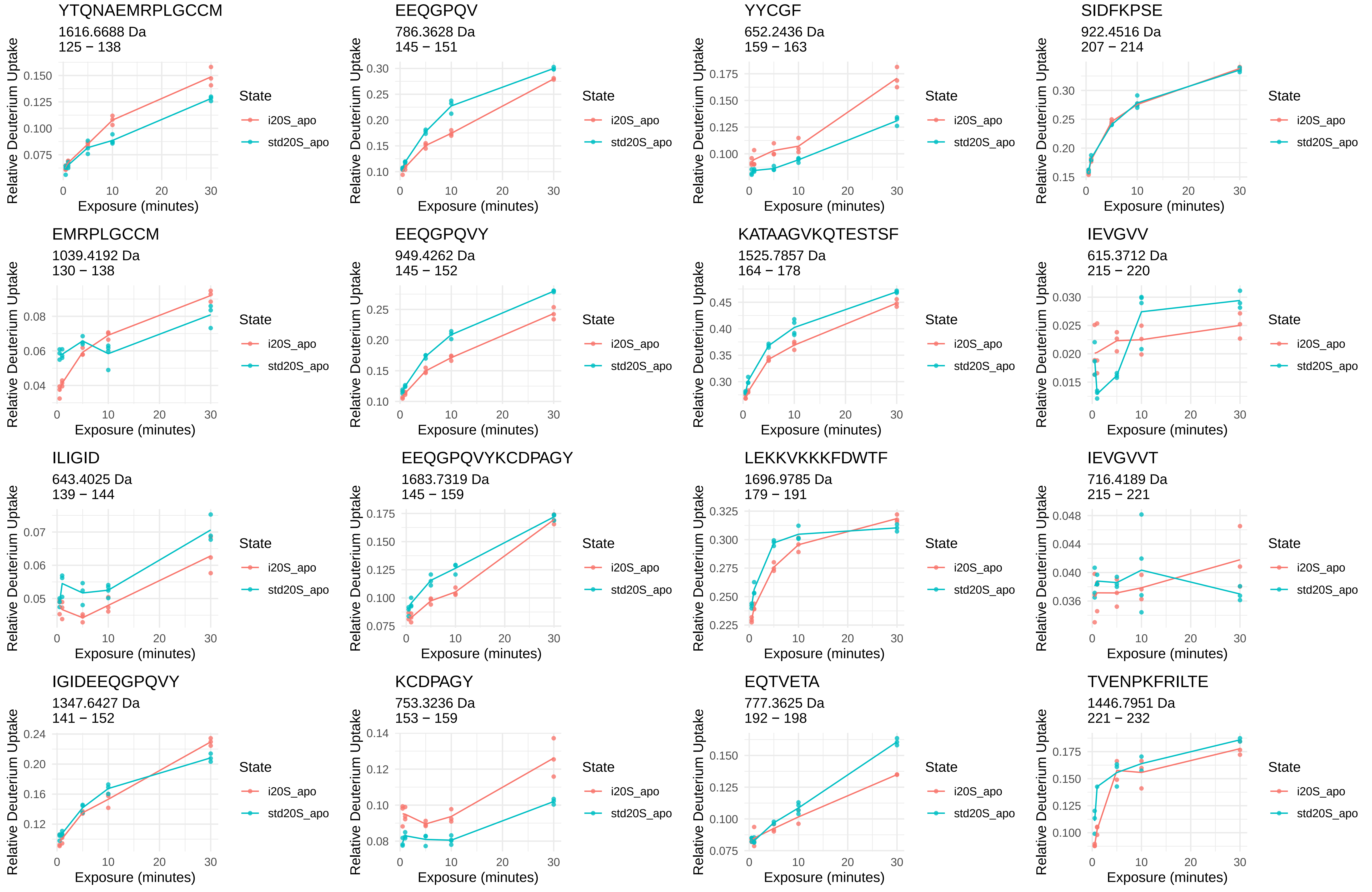
YTQNAEMRPLGCCM
1616.6688 Da
125 − 138
EEQGPQV
786.3628 Da
145 − 151
YYCGF
652.2436 Da
159 − 163
SIDFKPSE
922.4516 Da
207 − 214
EMRPLGCCM
1039.4192 Da
130 − 138
EEQGPQVY
949.4262 Da
145 − 152
KATAAGVKQTESTSF
1525.7857 Da
164 − 178
IEVGVV
615.3712 Da
215 − 220
ILIGID
643.4025 Da
139 − 144
EEQGPQVYKCDPAGY
1683.7319 Da
145 − 159
LEKKVKKKFDWTF
1696.9785 Da
179 − 191
IEVGVVT
716.4189 Da
215 − 221
IGIDEEQGPQVY
1347.6427 Da
141 − 152
KCDPAGY
753.3236 Da
153 − 159
EQTVETA
777.3625 Da
192 − 198
TVENPKFRILTE
1446.7951 Da
221 − 232
Relative Deuterium Uptake
Exposure (minutes)
State
i20S_apo
std20S_apo

VENPKF
733.3879 Da
222 − 227

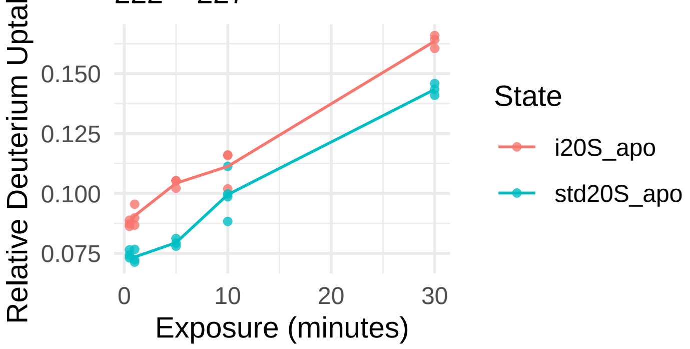


RILTE
631.3774 Da
228 − 232

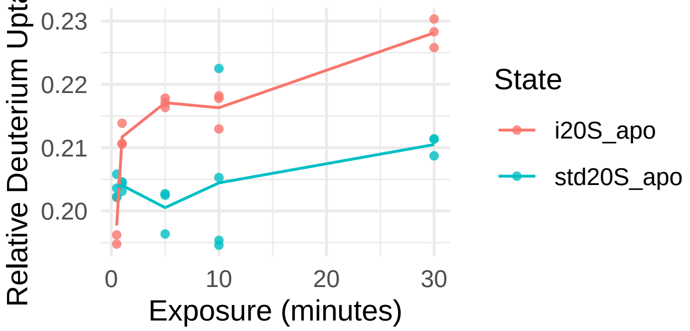


AEIDA
518.2457 Da
233 − 237

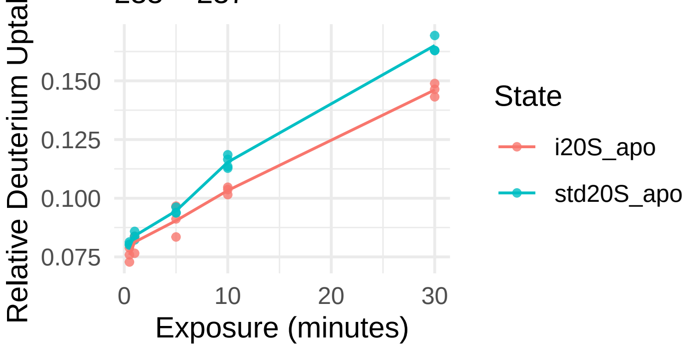


AEIDAHL
768.3886 Da
233 − 239

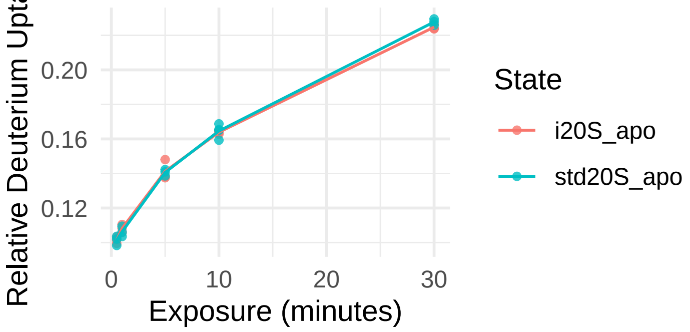

α2

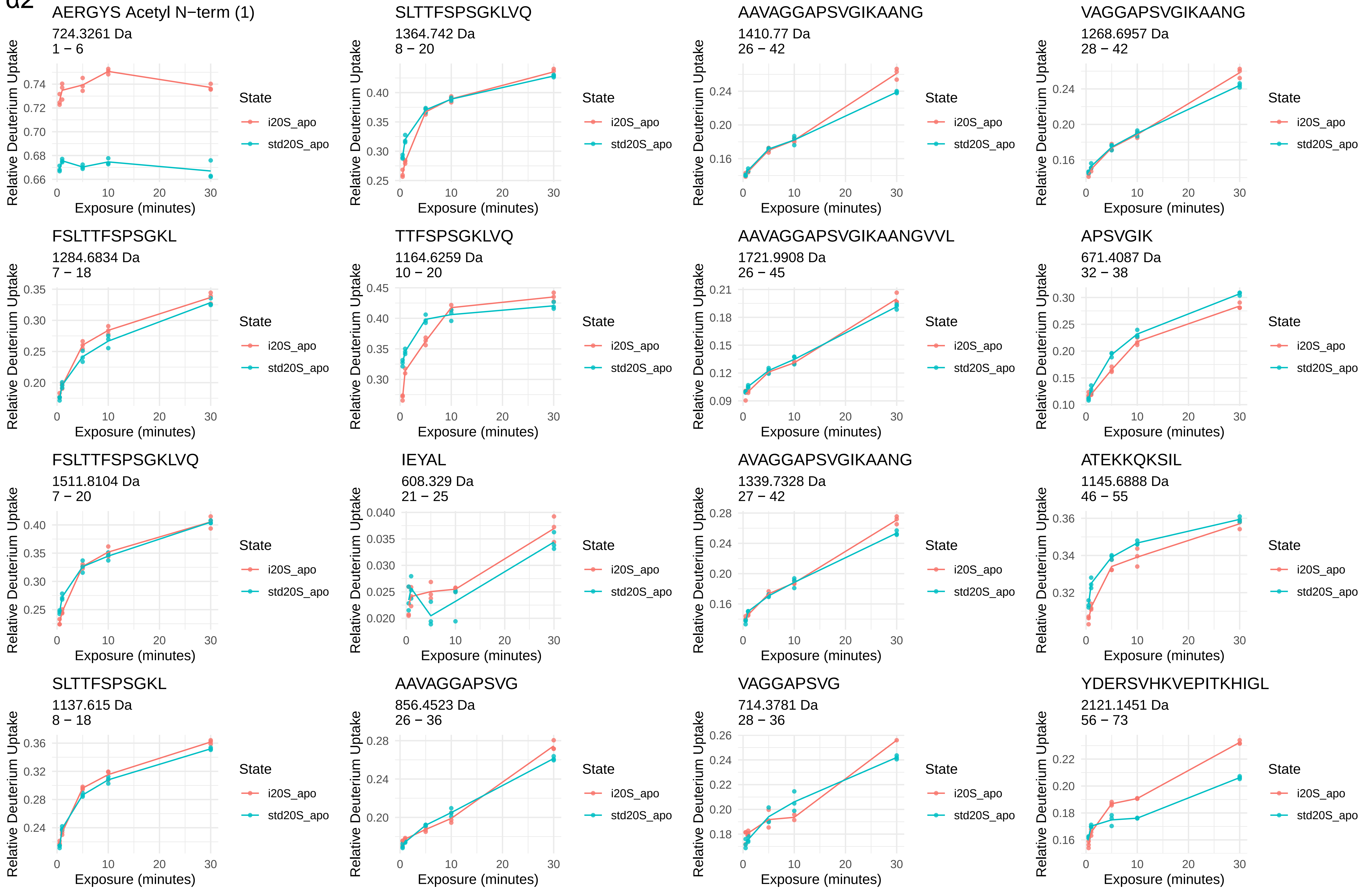
AERGYS Acetyl N−term (1)
724.3261 Da
1 − 6
SLTTFSPSGKLVQ
1364.742 Da
8 − 20
AAVAGGAPSVGIKAANG
1410.77 Da
26 − 42
VAGGAPSVGIKAANG
1268.6957 Da
28 − 42
FSLTTFSPSGKL
1284.6834 Da
7 − 18
TTFSPSGKLVQ
1164.6259 Da
10 − 20
AAVAGGAPSVGIKAANGVVL
1721.9908 Da
26 − 45
APSVGIK
671.4087 Da
32 − 38
FSLTTFSPSGKLVQ
1511.8104 Da
7 − 20
IEYAL
608.329 Da
21 − 25
AVAGGAPSVGIKAANG
1339.7328 Da
27 − 42
ATEKKQKSIL
1145.6888 Da
46 − 55
SLTTFSPSGKL
1137.615 Da
8 − 18
AAVAGGAPSVG
856.4523 Da
26 − 36
VAGGAPSVG
714.3781 Da
28 − 36
YDERSVHKVEPITKHIGL
2121.1451 Da
56 − 73
Relative Deuterium Uptake
Exposure (minutes)
State
i20S_apo
std20S_apo

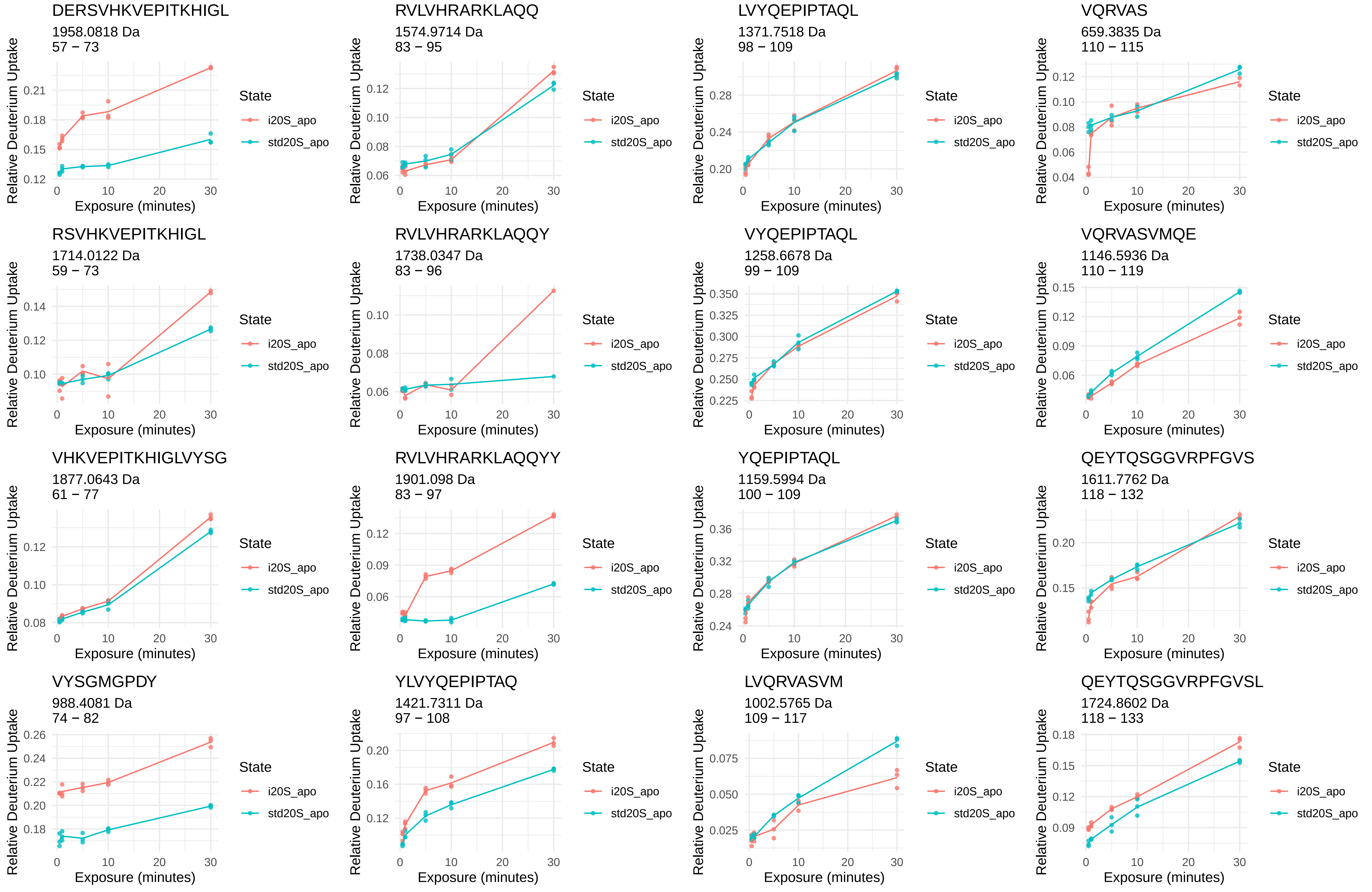
DERSVHKVEPITKHIGL
1958.0818 Da
57 − 73
RVLVHRARKLAQQ
1574.9714 Da
83 − 95
LVYQEPIPTAQL
1371.7518 Da
98 − 109
VQRVAS
659.3835 Da
110 − 115
RSVHKVEPITKHIGL
1714.0122 Da
59 − 73
RVLVHRARKLAQQY
1738.0347 Da
83 − 96
VYQEPIPTAQL
1258.6678 Da
99 − 109
VQRVASVMQE
1146.5936 Da
110 − 119
VHKVEPITKHIGLVYSG
1877.0643 Da
61 − 77
RVLVHRARKLAQQYY
1901.098 Da
83 − 97
YQEPIPTAQL
1159.5994 Da
100 − 109
QEYTQSGGVRPFGVS
1611.7762 Da
118 − 132
VYSGMGPDY
988.4081 Da
74 − 82
YLVYQEPIPTAQ
1421.7311 Da
97 − 108
LVQRVASVM
1002.5765 Da
109 − 117
QEYTQSGGVRPFGVSL
1724.8602 Da
118 − 133
Relative Deuterium Uptake
Exposure (minutes)
State
i20S_apo
std20S_apo

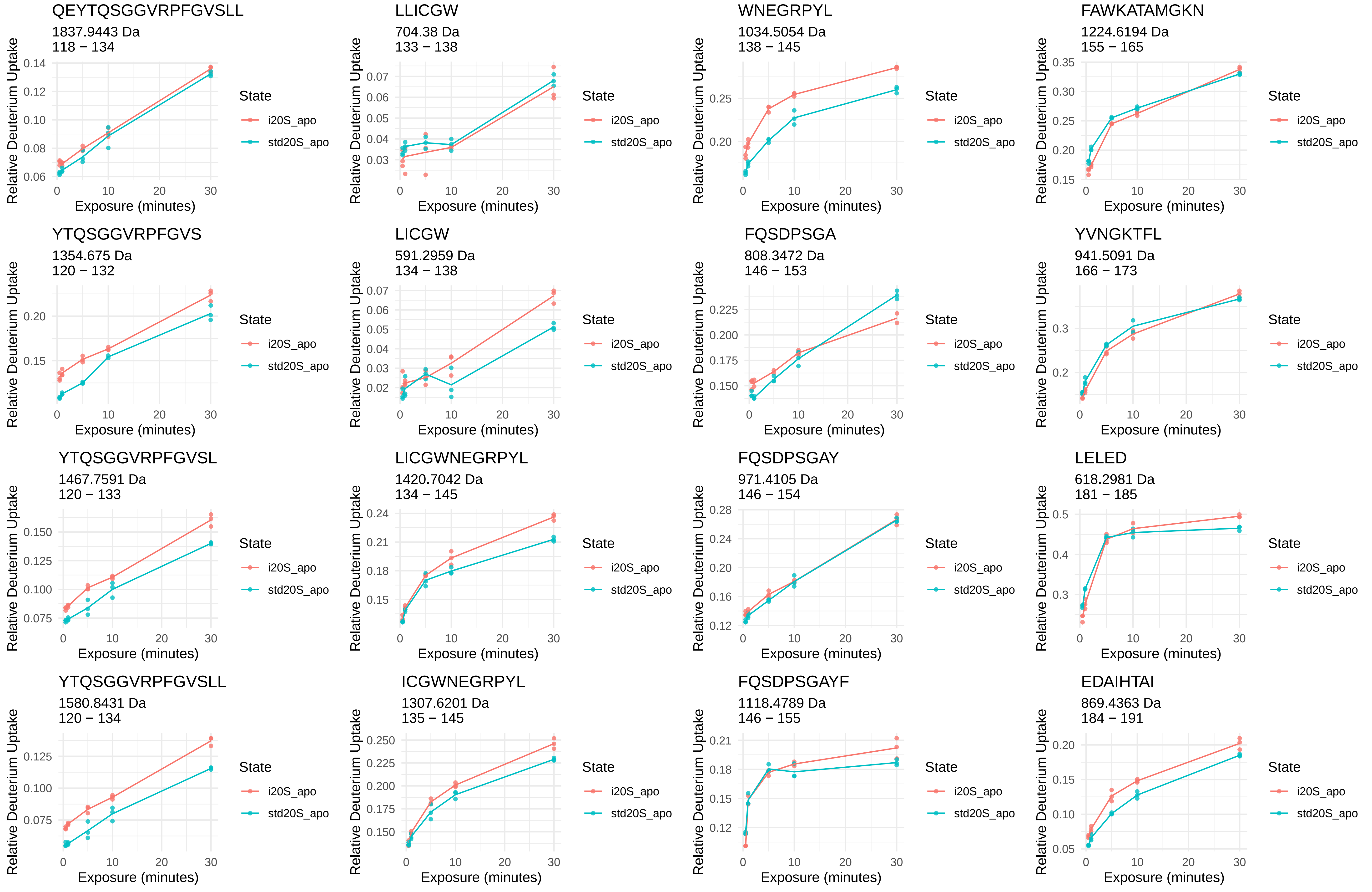
QEYTQSGGVRPFGVSLL
1837.9443 Da
118 − 134
LLICGW
704.38 Da
133 − 138
WNEGRPYL
1034.5054 Da
138 − 145
FAWKATAMGKN
1224.6194 Da
155 − 165
YTQSGGVRPFGVS
1354.675 Da
120 − 132
LICGW
591.2959 Da
134 − 138
FQSDPSGA
808.3472 Da
146 − 153
YVNGKTFL
941.5091 Da
166 − 173
YTQSGGVRPFGVSL
1467.7591 Da
120 − 133
LICGWNEGRPYL
1420.7042 Da
134 − 145
FQSDPSGAY
971.4105 Da
146 − 154
LELED
618.2981 Da
181 − 185
YTQSGGVRPFGVSLL
1580.8431 Da
120 − 134
ICGWNEGRPYL
1307.6201 Da
135 − 145
FQSDPSGAYF
1118.4789 Da
146 − 155
EDAIHTAI
869.4363 Da
184 − 191
Relative Deuterium Uptake
Exposure (minutes)
State
i20S_apo
std20S_apo

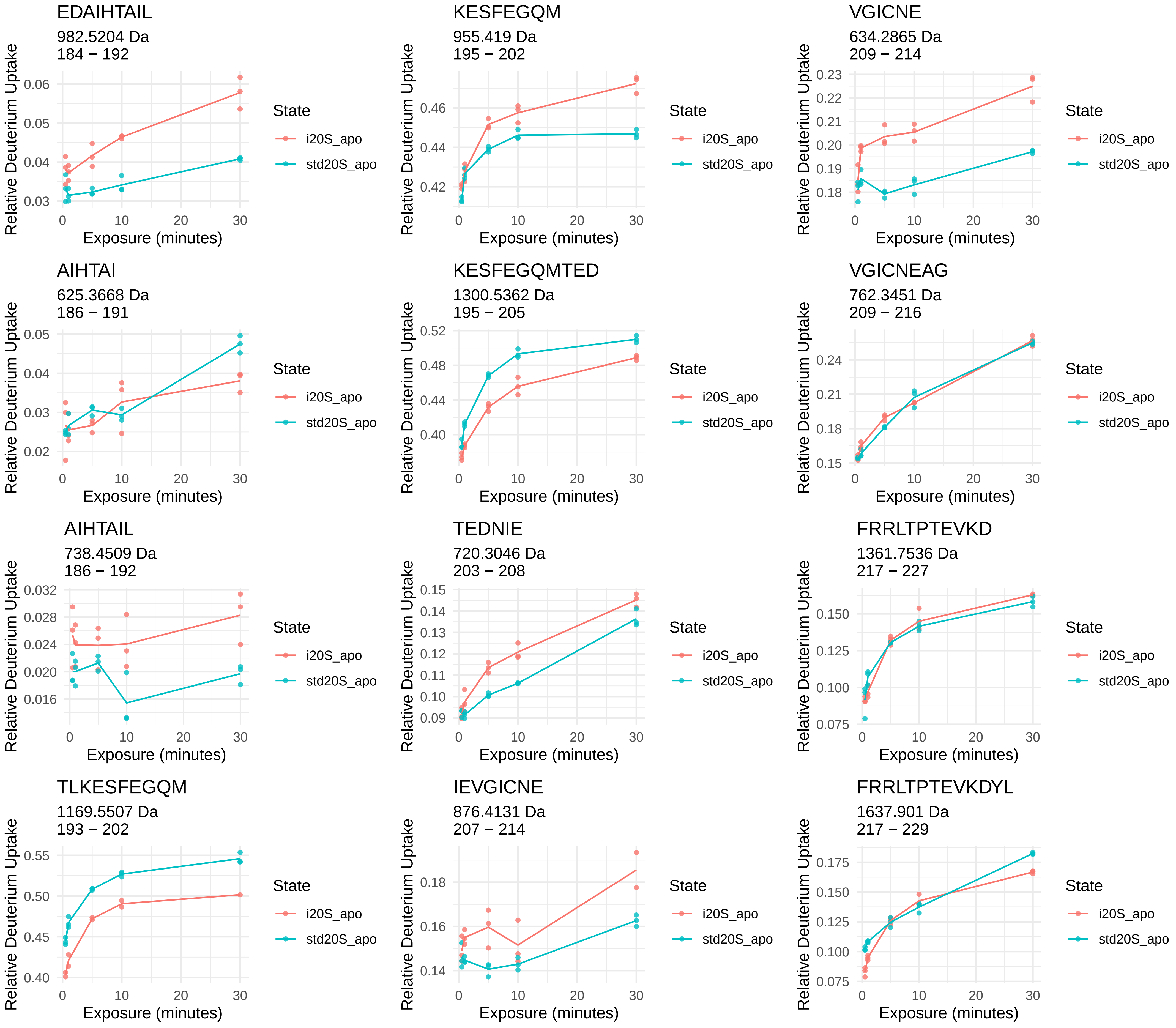
EDAIHTAIL
982.5204 Da
184 − 192
KESFEGQM
955.419 Da
195 − 202
VGICNE
634.2865 Da
209 − 214
AIHTAI
625.3668 Da
186 − 191
KESFEGQMTED
1300.5362 Da
195 − 205
VGICNEAG
762.3451 Da
209 − 216
AIHTAIL
738.4509 Da
186 − 192
TEDNIE
720.3046 Da
203 − 208
FRRLTPTEVKD
1361.7536 Da
217 − 227
TLKESFEGQM
1169.5507 Da
193 − 202
IEVGICNE
876.4131 Da
207 − 214
FRRLTPTEVKDYL
1637.901 Da
217 − 229
Relative Deuterium Uptake
Exposure (minutes)
State
i20S_apo
std20S_apo

α3

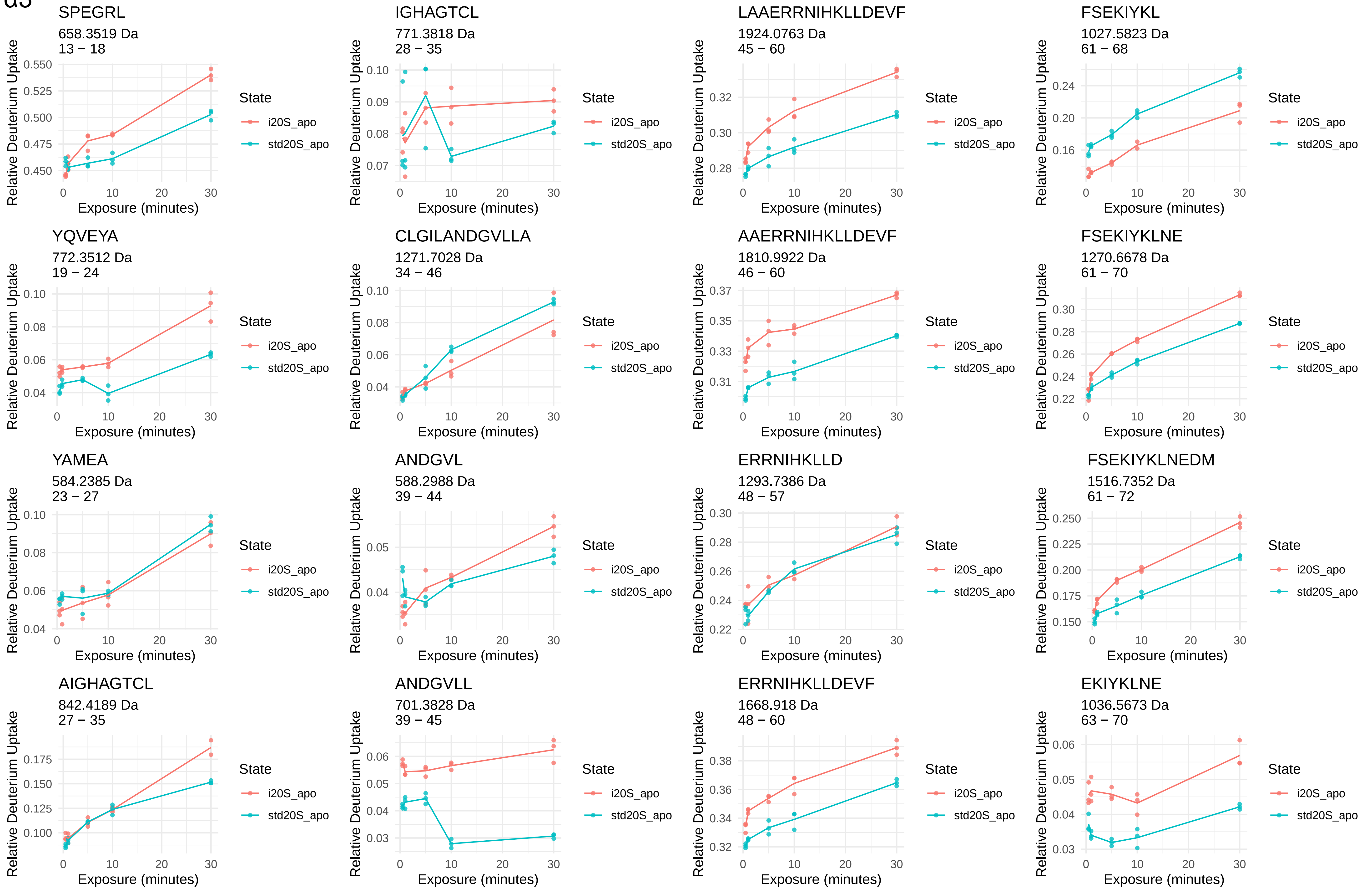
SPEGRL
658.3519 Da
13 − 18
IGHAGTCL
771.3818 Da
28 − 35
LAAERRNIHKLLDEVF
1924.0763 Da
45 − 60
FSEKIYKL
1027.5823 Da
61 − 68
YQVEYA
772.3512 Da
19 − 24
CLGILANDGVLLA
1271.7028 Da
34 − 46
AAERRNIHKLLDEVF
1810.9922 Da
46 − 60
FSEKIYKLNE
1270.6678 Da
61 − 70
YAMEA
584.2385 Da
23 − 27
ANDGVL
588.2988 Da
39 − 44
ERRNIHKLLD
1293.7386 Da
48 − 57
FSEKIYKLNEDM
1516.7352 Da
61 − 72
AIGHAGTCL
842.4189 Da
27 − 35
ANDGVLL
701.3828 Da
39 − 45
ERRNIHKLLDEVF
1668.918 Da
48 − 60
EKIYKLNE
1036.5673 Da
63 − 70
Relative Deuterium Uptake
Exposure (minutes)
State
i20S_apo
std20S_apo

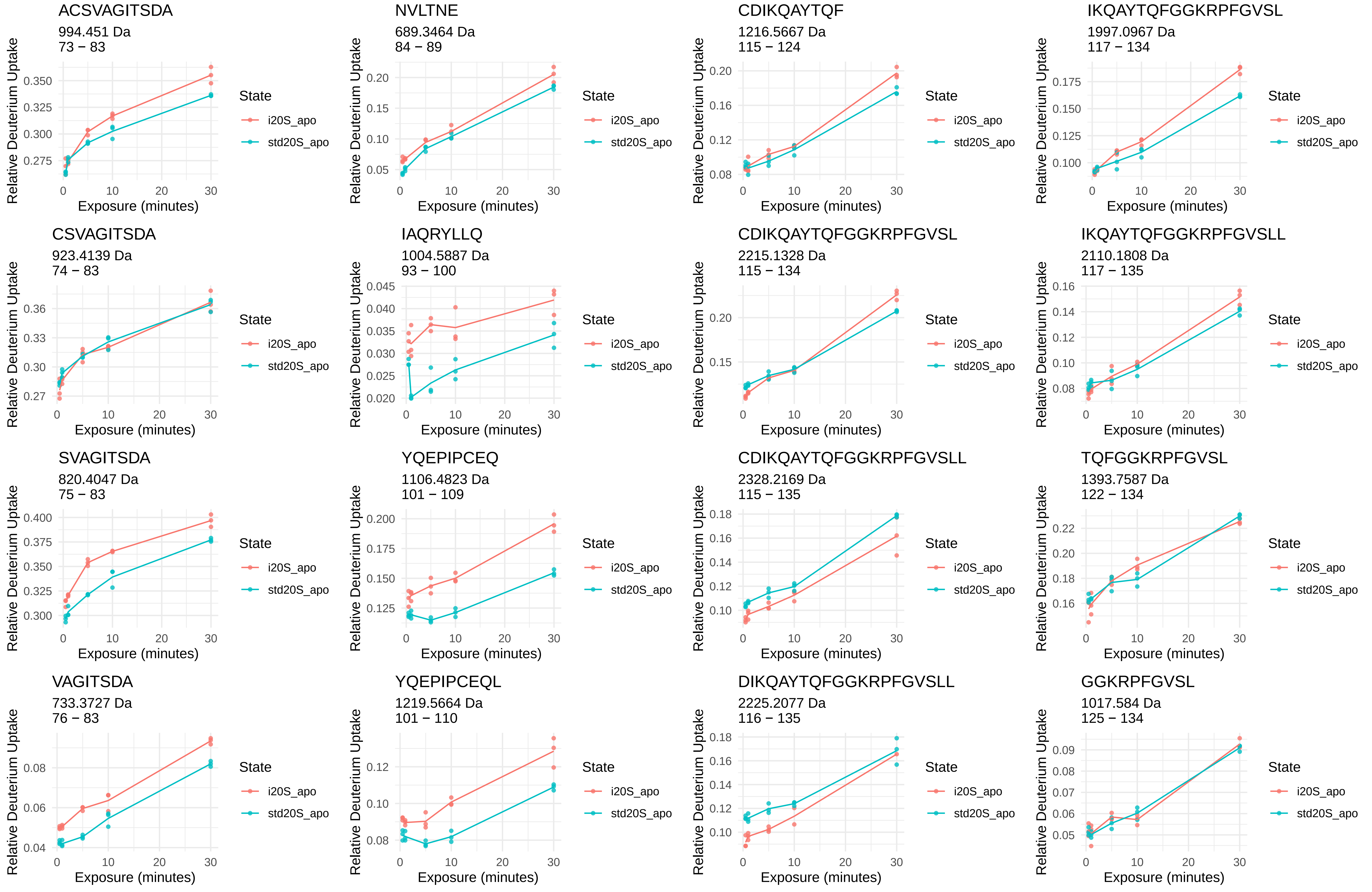
ACSVAGITSDA
994.451 Da
73 – 83
NVLTNE
689.3464 Da
84 – 89
CDIKQAYTQF
1216.5667 Da
115 – 124
IKQAYTQFGGKRPFGVSL
1997.0967 Da
117 – 134
CSVAGITSDA
923.4139 Da
74 – 83
IAQRYLLQ
1004.5887 Da
93 – 100
CDIKQAYTQFGGKRPFGVSL
2215.1328 Da
115 – 134
IKQAYTQFGGKRPFGVSLL
2110.1808 Da
117 – 135
SVAGITSDA
820.4047 Da
75 – 83
YQEPIPCEQ
1106.4823 Da
101 – 109
CDIKQAYTQFGGKRPFGVSLL
2328.2169 Da
115 – 135
TQFGGKRPFGVSL
1393.7587 Da
122 – 134
VAGITSDA
733.3727 Da
76 – 83
YQEPIPCEQL
1219.5664 Da
101 – 110
DIKQAYTQFGGKRPFGVSLL
2225.2077 Da
116 – 135
GGKRPFGVSL
1017.584 Da
125 – 134
Relative Deuterium Uptake
Exposure (minutes)
State
i20S_apo
std20S_apo

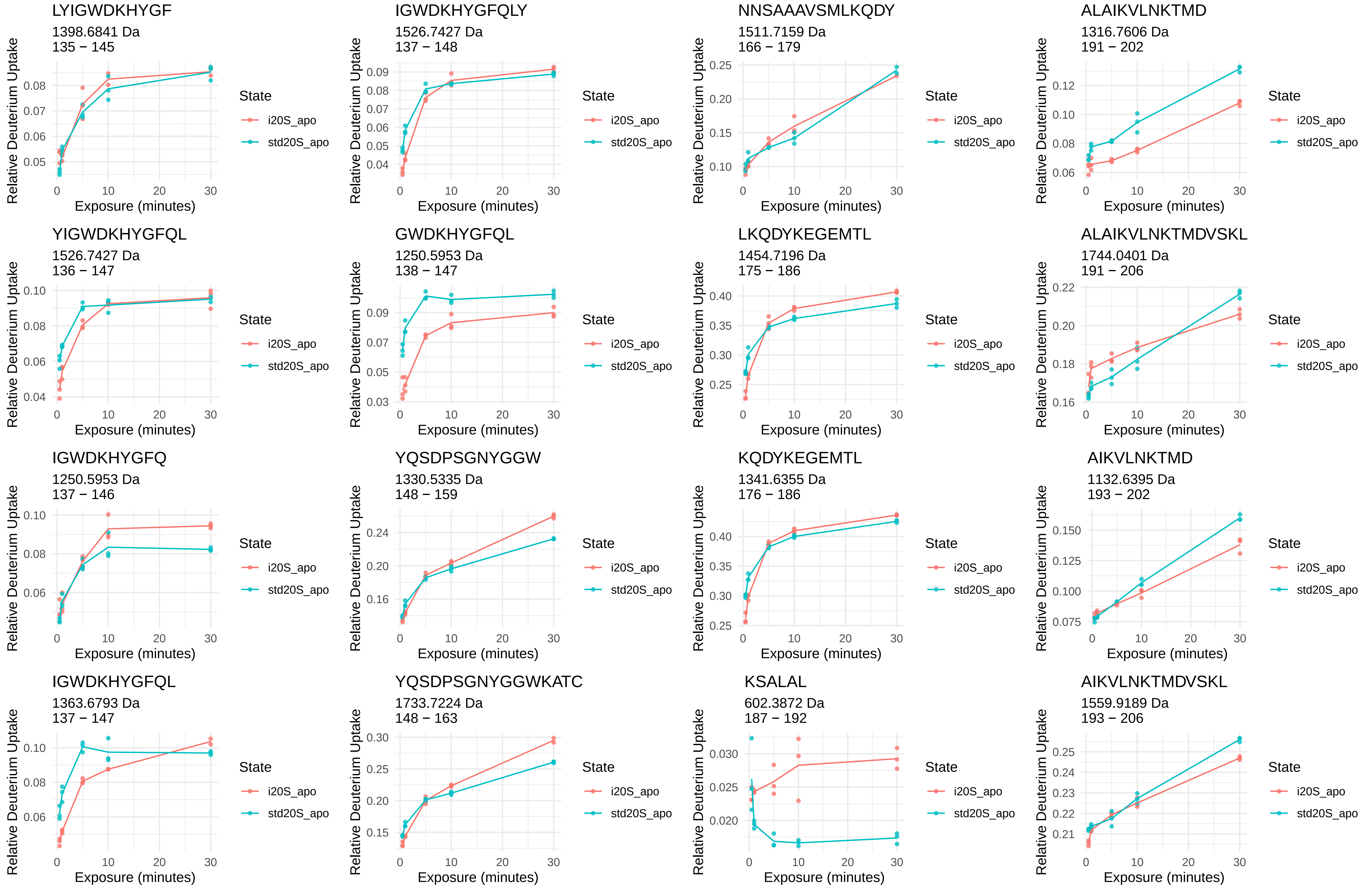
LYIGWDKHYGF
1398.6841 Da
135 − 145
IGWDKHYGFQLY
1526.7427 Da
137 − 148
NNSAAAVSMLKQDY
1511.7159 Da
166 − 179
ALAIKVLNKTMD
1316.7606 Da
191 − 202
YIGWDKHYGFQL
1526.7427 Da
136 − 147
GWDKHYGFQL
1250.5953 Da
138 − 147
LKQDYKEGEMTL
1454.7196 Da
175 − 186
ALAIKVLNKTMDVSKL
1744.0401 Da
191 − 206
IGWDKHYGFQ
1250.5953 Da
137 − 146
YQSDPSGNYGGW
1330.5335 Da
148 − 159
KQDYKEGEMTL
1341.6355 Da
176 − 186
AIKVLNKTMD
1132.6395 Da
193 − 202
IGWDKHYGFQL
1363.6793 Da
137 − 147
YQSDPSGNYGGWKATC
1733.7224 Da
148 − 163
KSALAL
602.3872 Da
187 − 192
AIKVLNKTMDVSKL
1559.9189 Da
193 − 206
Relative Deuterium Uptake
Exposure (minutes)
State
i20S_apo
std20S_apo

AIKVLNKTMDVSKLSA
1717.9881 Da
193 − 208

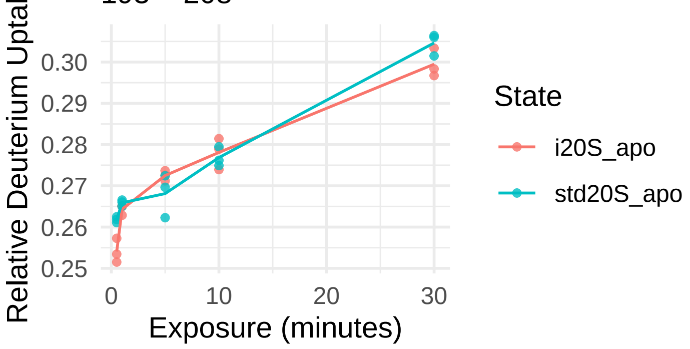


KVEIATL
773.4767 Da
210 − 216

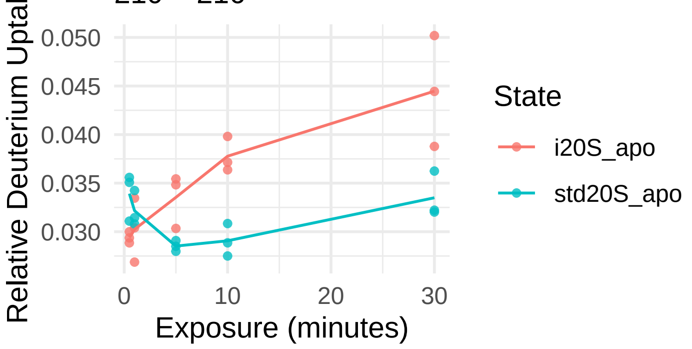


KVLNKTMDVSKL
1375.7977 Da
195 − 206

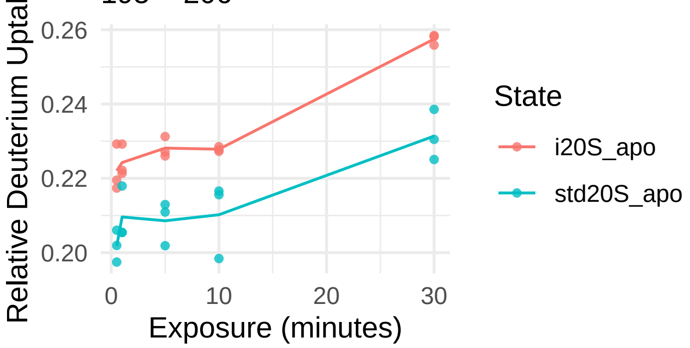


LTRENGKTVIRVLKQKEVEQL
2481.4511 Da
216 − 236

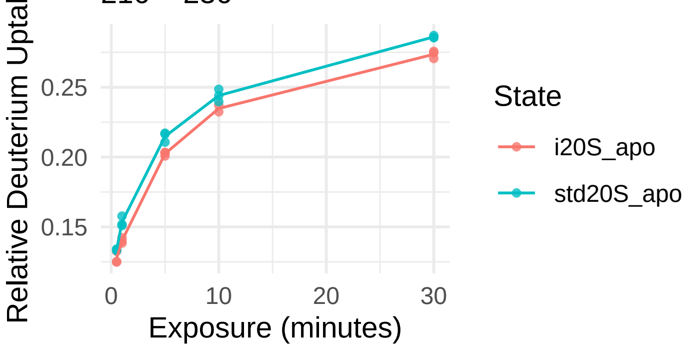


SAEKVEIATL
1060.5885 Da
207 − 216

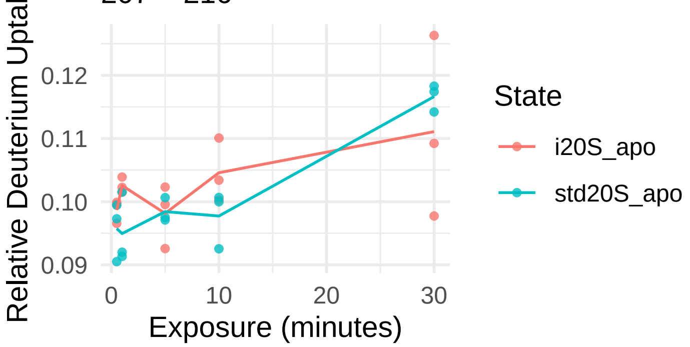


TRENGKTVIRVLKQKEVEQL
2368.3671 Da
217 − 236

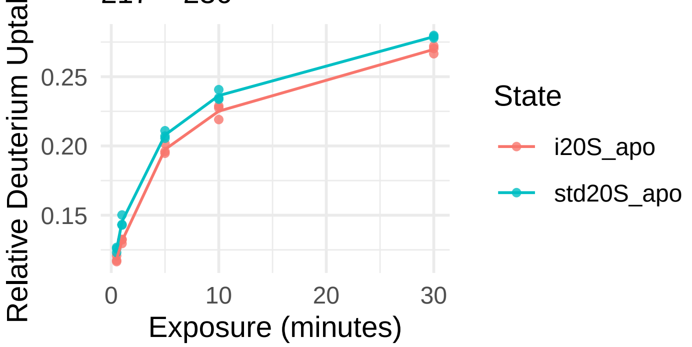


EKVEIATL
902.5193 Da
209 − 216

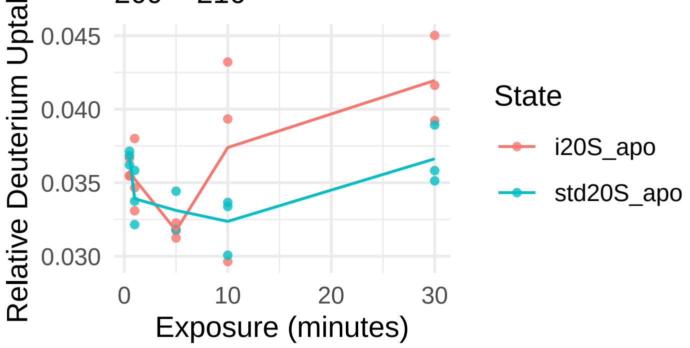


IKKHEEEEAKAEREKKEKEQKEKDK
3124.6597 Da
237 − 261

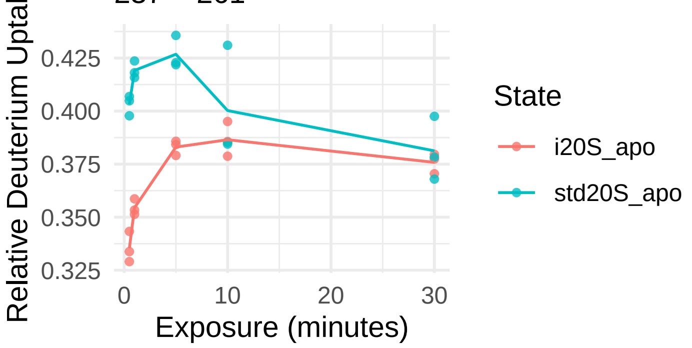

α4

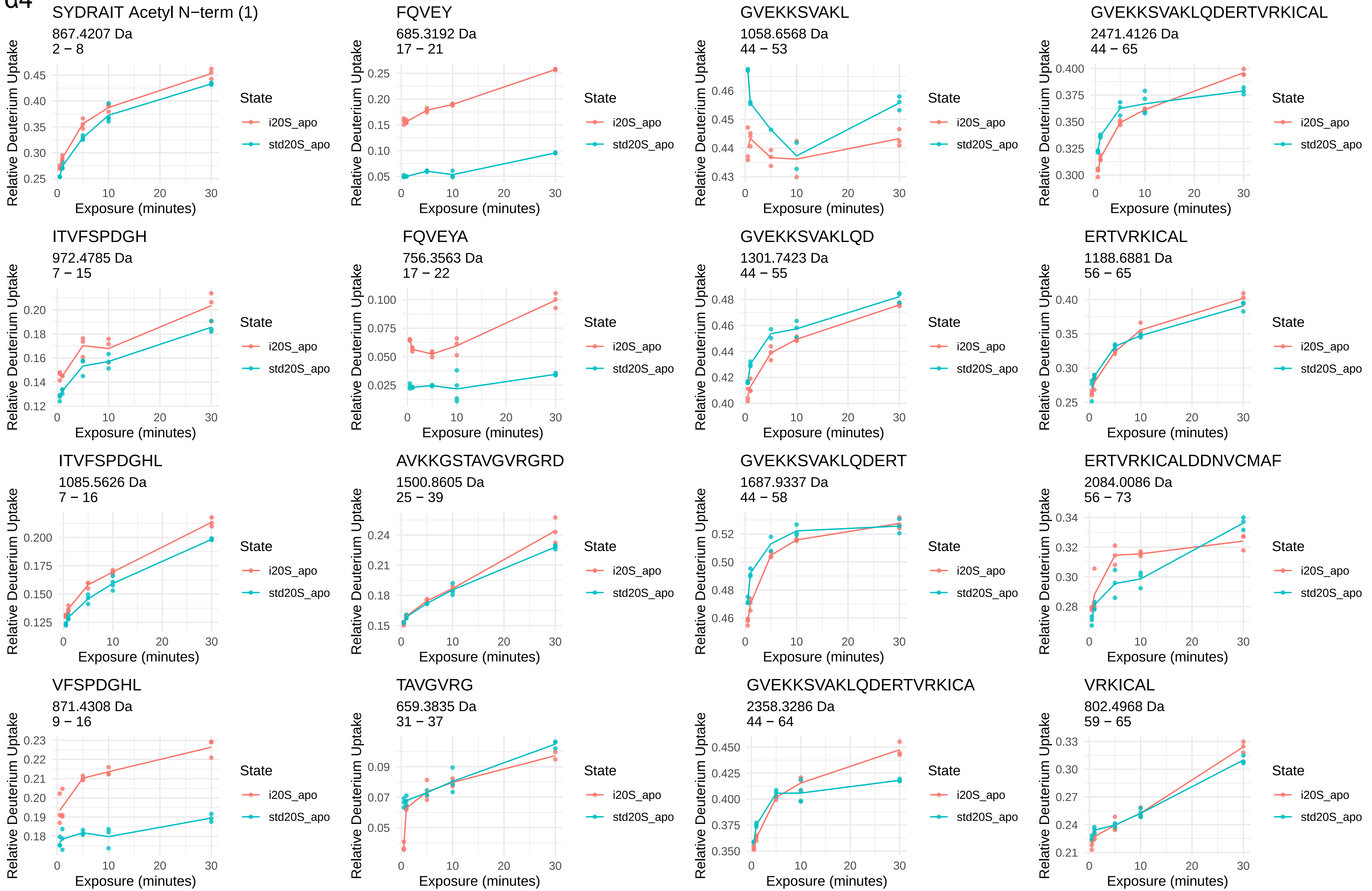
SYDRAIT Acetyl N−term (1)
867.4207 Da
2 − 8
FQVEY
685.3192 Da
17 − 21
GVEKKSVAKL
1058.6568 Da
44 − 53
GVEKKSVAKLQDERTVRKICAL
2471.4126 Da
44 − 65
ITVFSPDGH
972.4785 Da
7 − 15
FQVEYA
756.3563 Da
17 − 22
GVEKKSVAKLQD
1301.7423 Da
44 − 55
ERTVRKICAL
1188.6881 Da
56 − 65
ITVFSPDGHL
1085.5626 Da
7 − 16
AVKKGSTAVGVRGRD
1500.8605 Da
25 − 39
GVEKKSVAKLQDERT
1687.9337 Da
44 − 58
ERTVRKICALDDNVCMAF
2084.0086 Da
56 − 73
VFSPDGHL
871.4308 Da
9 − 16
TAVGVRG
659.3835 Da
31 − 37
GVEKKSVAKLQDERTVRKICA
2358.3286 Da
44 − 64
VRKICAL
802.4968 Da
59 − 65
Relative Deuterium Uptake
Exposure (minutes)
State
i20S_apo
std20S_apo

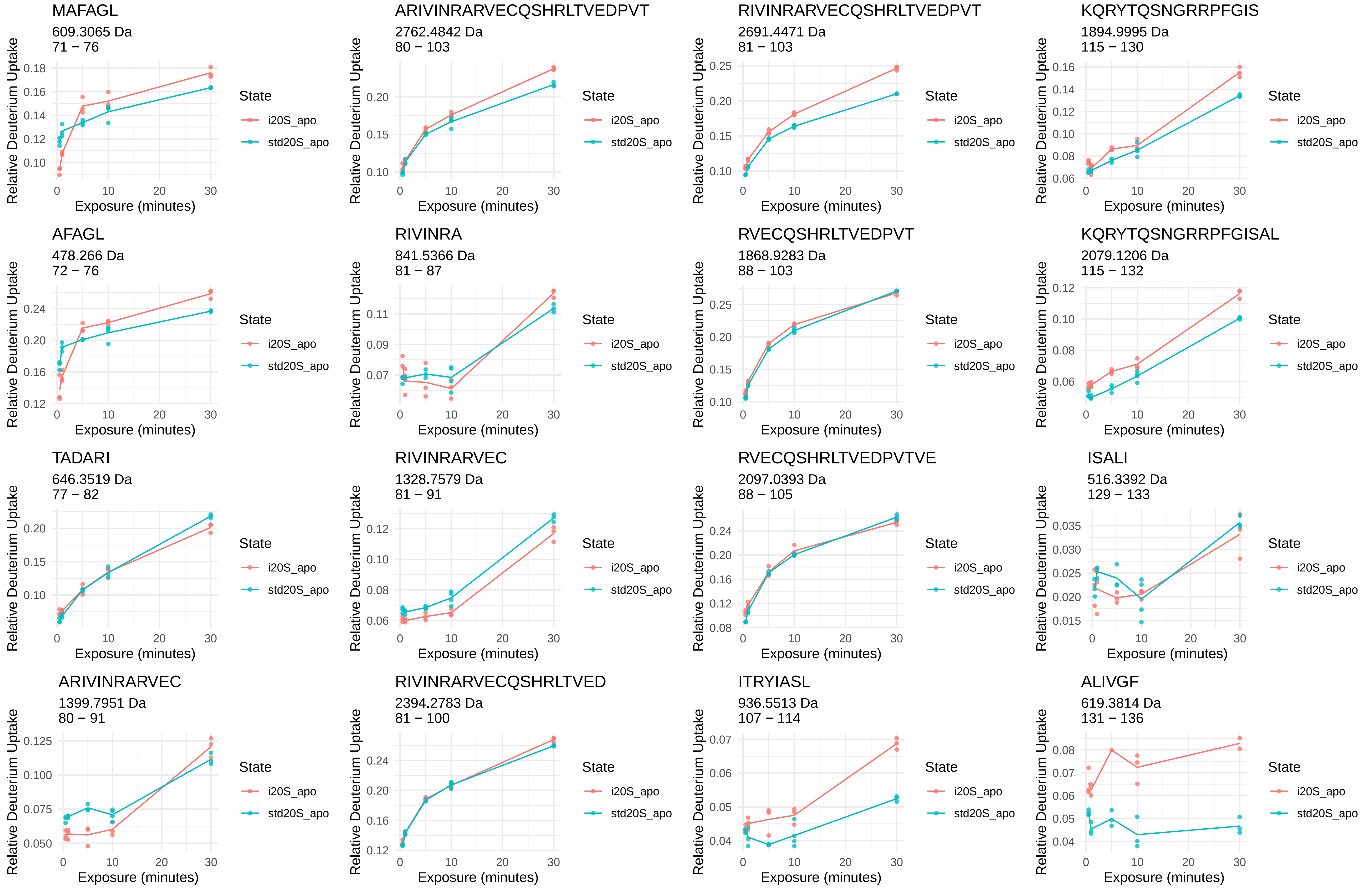
MAFAGL
609.3065 Da
71 − 76
ARIVINRARVECQSHRLTVEDPVT
2762.4842 Da
80 − 103
RIVINRARVECQSHRLTVEDPVT
2691.4471 Da
81 − 103
KQRYTQSNGRRPFGIS
1894.9995 Da
115 − 130
AFAGL
478.266 Da
72 − 76
RIVINRA
841.5366 Da
81 − 87
RVECQSHRLTVEDPVT
1868.9283 Da
88 − 103
KQRYTQSNGRRPFGISAL
2079.1206 Da
115 − 132
TADARI
646.3519 Da
77 − 82
RIVINRARVEC
1328.7579 Da
81 − 91
RVECQSHRLTVEDPVTVE
2097.0393 Da
88 − 105
ISALI
516.3392 Da
129 − 133
ARIVINRARVEC
1399.7951 Da
80 − 91
RIVINRARVECQSHRLTVED
2394.2783 Da
81 − 100
ITRYIASL
936.5513 Da
107 − 114
ALIVGF
619.3814 Da
131 − 136
Relative Deuterium Uptake
Exposure (minutes)
State
i20S_apo
std20S_apo

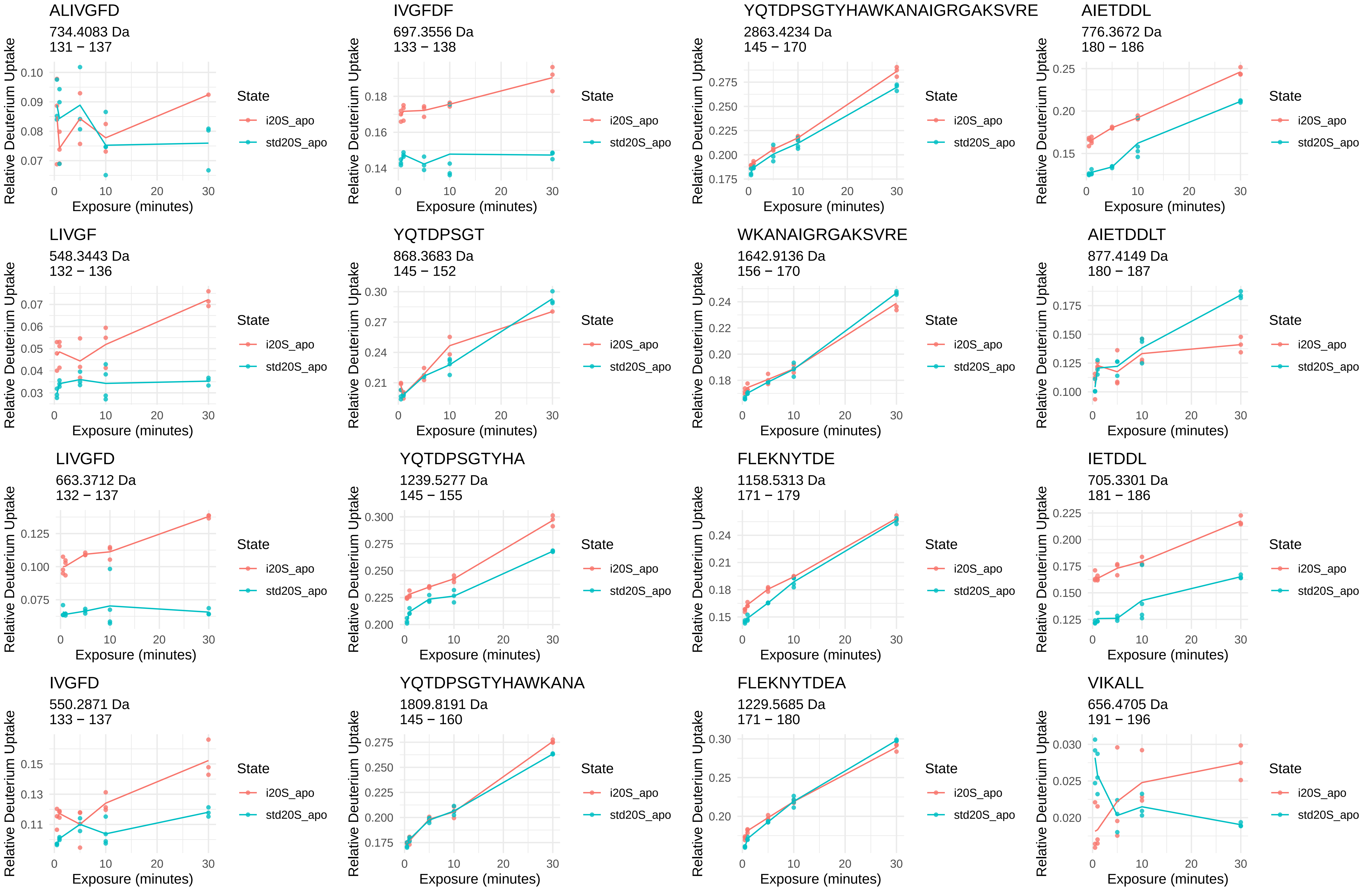
ALIVGFD
734.4083 Da
131 − 137
IVGFDF
697.3556 Da
133 − 138
YQTDPSGTYHAWKANAIGRGAKSVRE
2863.4234 Da
145 − 170
AIETDDL
776.3672 Da
180 − 186
LIVGF
548.3443 Da
132 − 136
YQTDPSGT
868.3683 Da
145 − 152
WKANAIGRGAKSVRE
1642.9136 Da
156 − 170
AIETDDLT
877.4149 Da
180 − 187
LIVGFD
663.3712 Da
132 − 137
YQTDPSGTYHA
1239.5277 Da
145 − 155
FLEKNYTDE
1158.5313 Da
171 − 179
IETDDL
705.3301 Da
181 − 186
IVGFD
550.2871 Da
133 − 137
YQTDPSGTYHAWKANA
1809.8191 Da
145 − 160
FLEKNYTDEA
1229.5685 Da
171 − 180
VIKALL
656.4705 Da
191 − 196
Relative Deuterium Uptake
Exposure (minutes)
State
i20S_apo
std20S_apo

VIKALLE

785.5131 Da

191 − 197

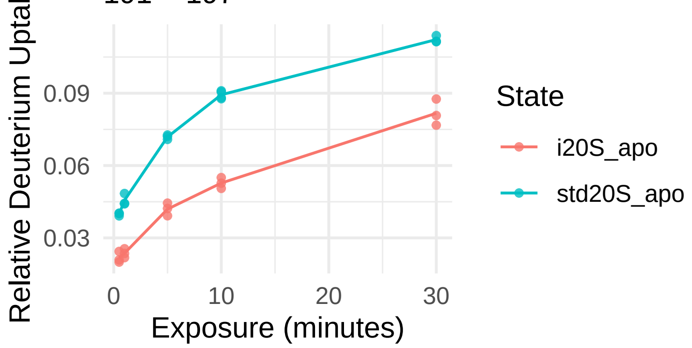


KILNPEE

842.4618 Da

218 − 224

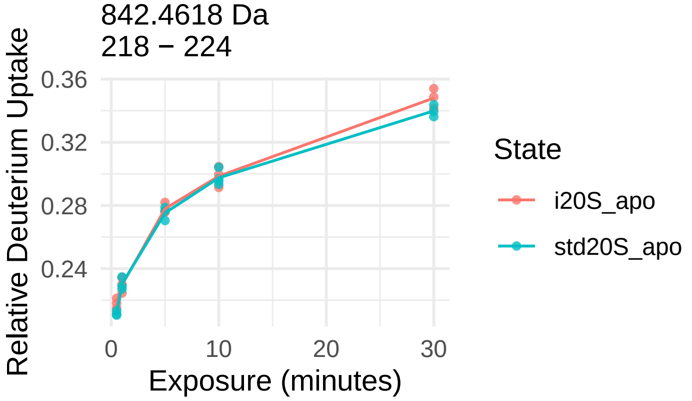


EVVQSGGKNIE

1159.5953 Da

197 − 207

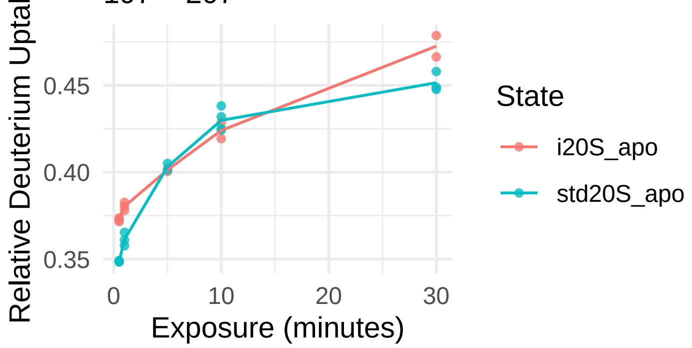


IEKYVAE

851.4509 Da

225 − 231

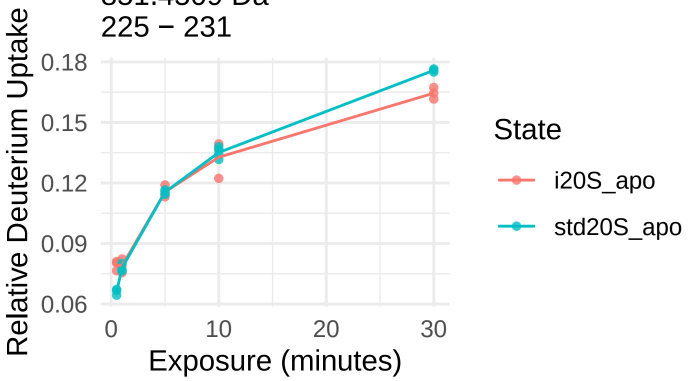


AVMRRDQSL

1075.5677 Da

209 − 217

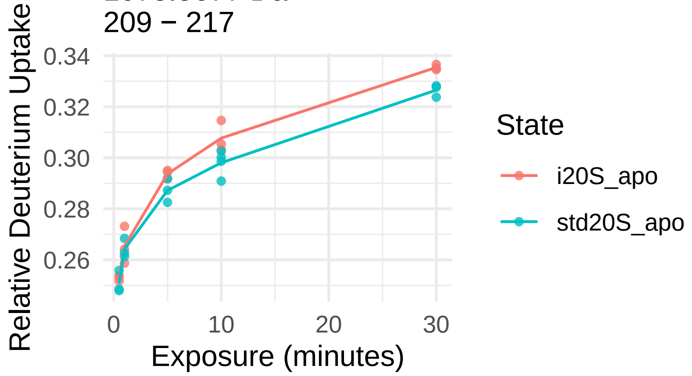


IEKEKEENEKKKQKKAS

2074.1503 Da

232 − 248

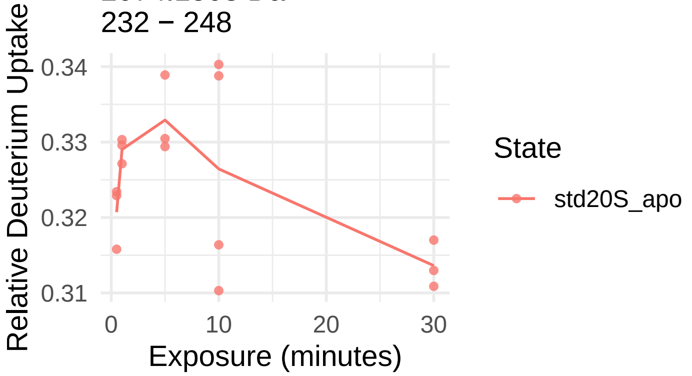


LKILNPEEIEKYV

1587.8992 Da

217 − 229

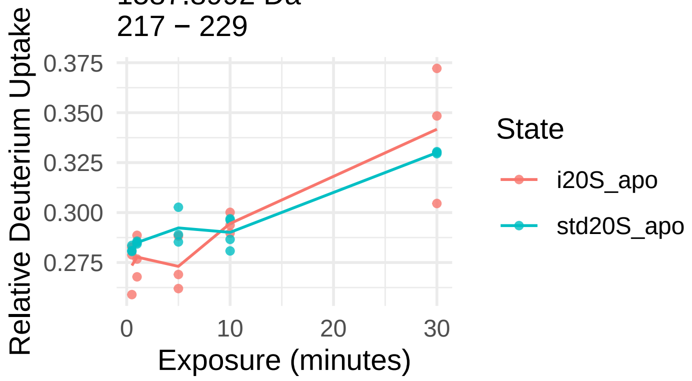

α5

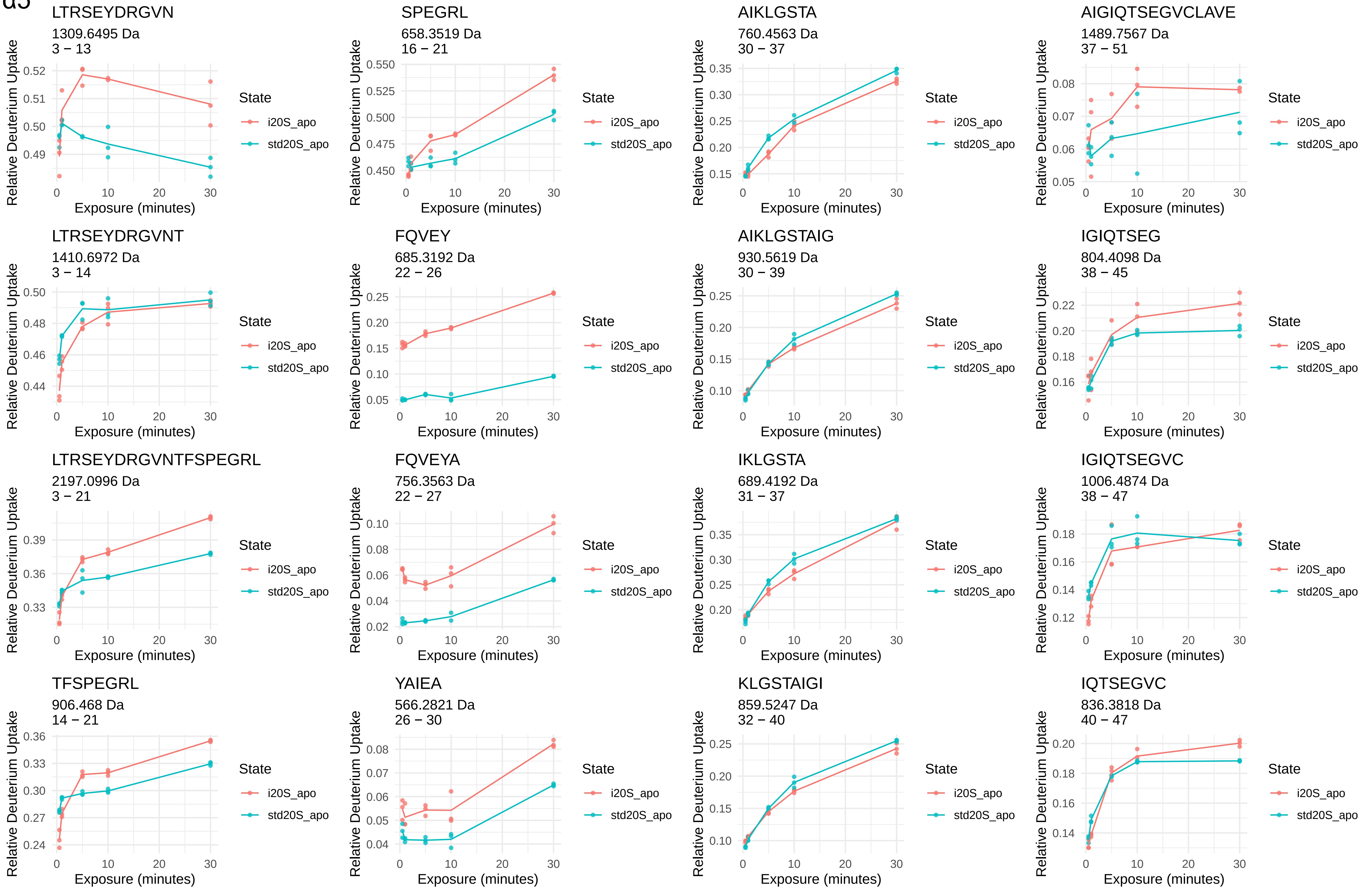
LTRSEYDRGVN
1309.6495 Da
3 – 13
SPEGRL
658.3519 Da
16 – 21
AIKLGSTA
760.4563 Da
30 – 37
AIGIQTSEGVCLAVE
1489.7567 Da
37 – 51
LTRSEYDRGVNT
1410.6972 Da
3 – 14
FQVEY
685.3192 Da
22 – 26
AIKLGSTAIG
930.5619 Da
30 – 39
IGIQTSEG
804.4098 Da
38 – 45
LTRSEYDRGVNTFSPEGRL
2197.0996 Da
3 – 21
FQVEYA
756.3563 Da
22 – 27
IKLGSTA
689.4192 Da
31 – 37
IGIQTSEGVC
1006.4874 Da
38 – 47
TFSPEGRL
906.468 Da
14 – 21
YAIEA
566.2821 Da
26 – 30
KLGSTAIGI
859.5247 Da
32 – 40
IQTSEGVC
836.3818 Da
40 – 47
Relative Deuterium Uptake
Exposure (minutes)
State
i20S_apo
std20S_apo

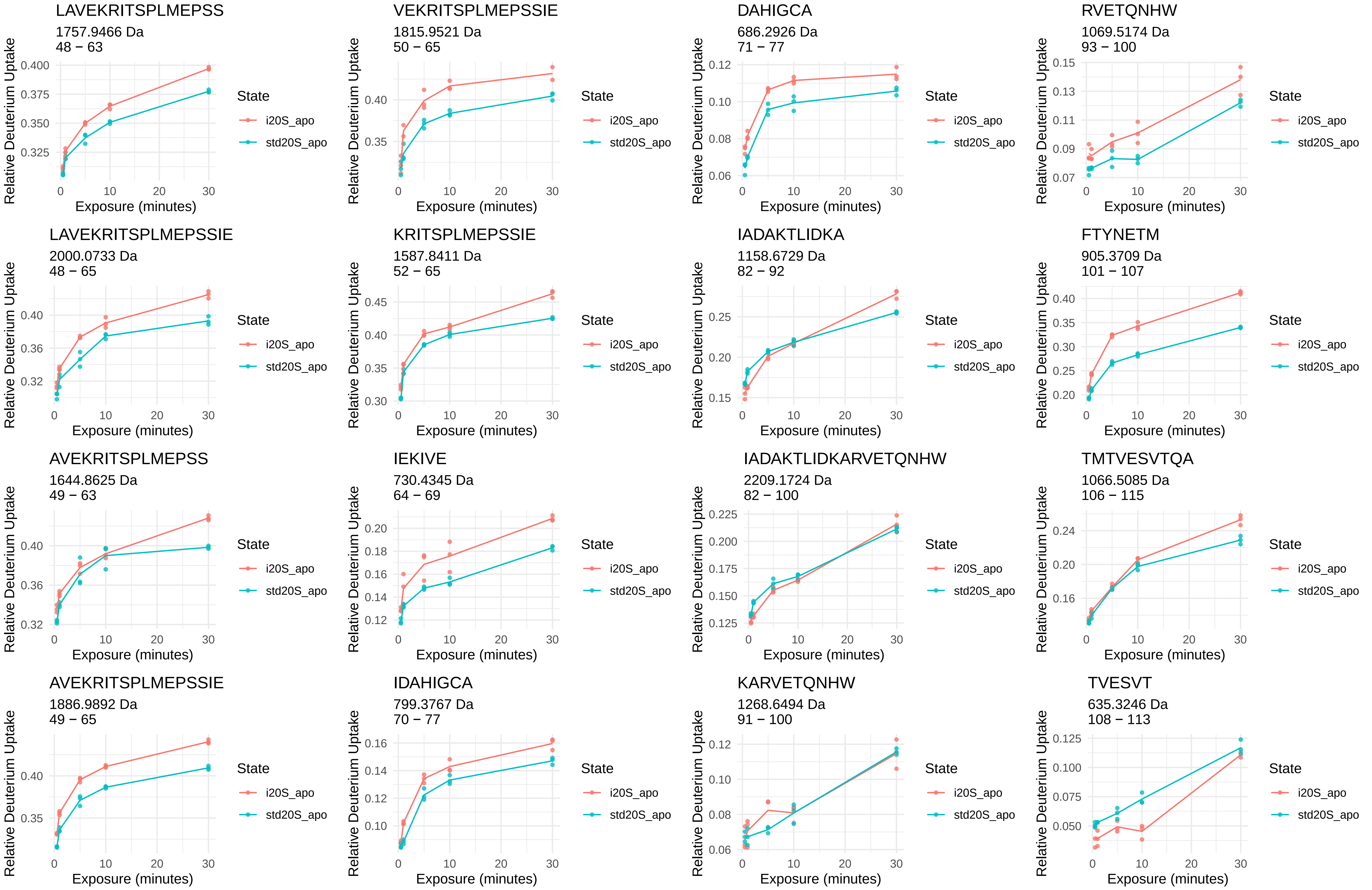
LAVEKRITSPLMEPSS
1757.9466 Da
48 − 63
VEKRITSPLMEPSSIE
1815.9521 Da
50 − 65
DAHIGCA
686.2926 Da
71 − 77
RVETQNHW
1069.5174 Da
93 − 100
LAVEKRITSPLMEPSSIE
2000.0733 Da
48 − 65
KRITSPLMEPSSIE
1587.8411 Da
52 − 65
IADAKTLIDKA
1158.6729 Da
82 − 92
FTYNETM
905.3709 Da
101 − 107
AVEKRITSPLMEPSS
1644.8625 Da
49 − 63
IEKIVE
730.4345 Da
64 − 69
IADAKTLIDKARVETQNHW
2209.1724 Da
82 − 100
TMTVESVTQA
1066.5085 Da
106 − 115
AVEKRITSPLMEPSSIE
1886.9892 Da
49 − 65
IDAHIGCA
799.3767 Da
70 − 77
KARVETQNHW
1268.6494 Da
91 − 100
TVESVT
635.3246 Da
108 − 113
Relative Deuterium Uptake
Exposure (minutes)
State
i20S_apo
std20S_apo

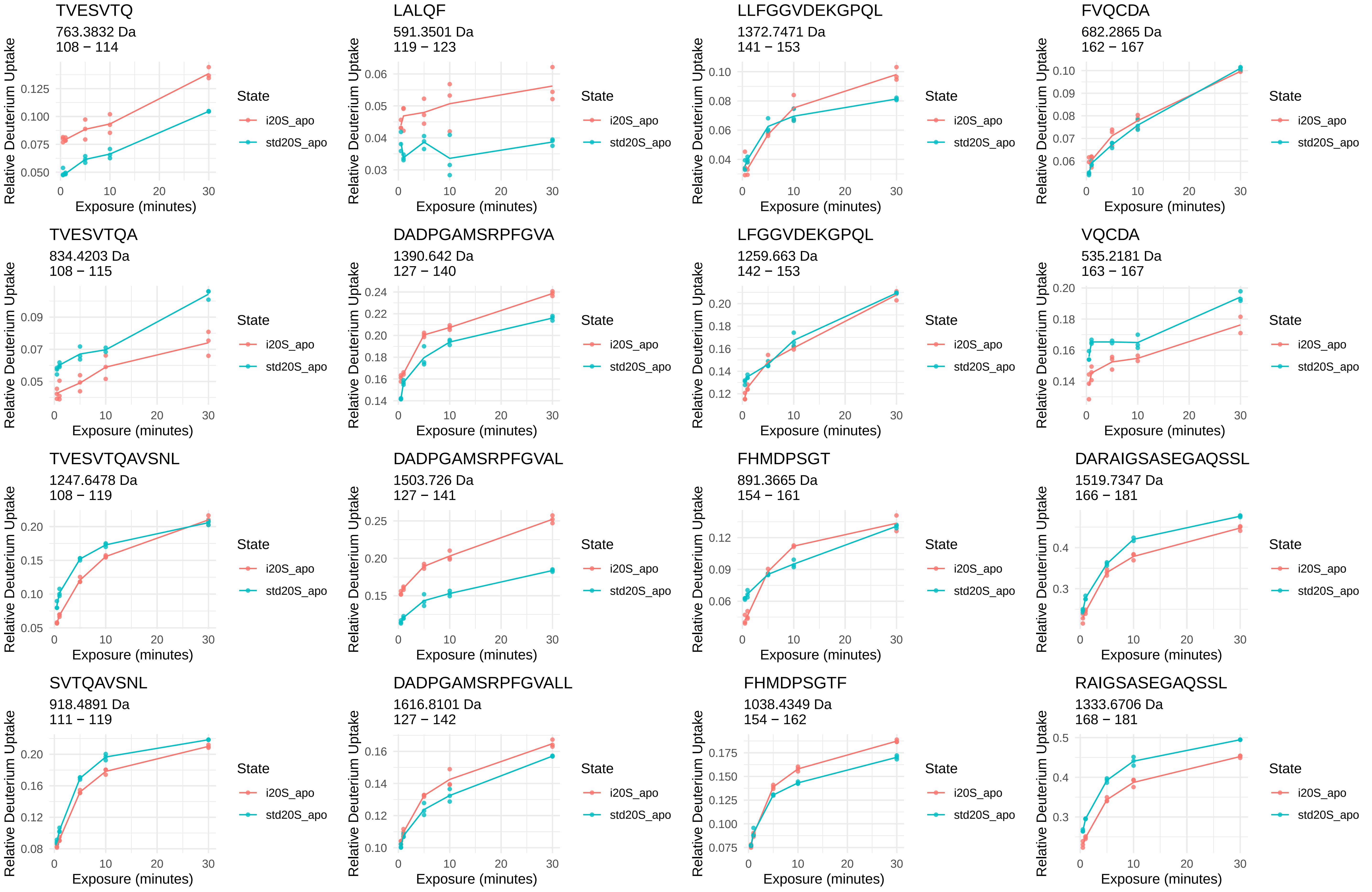
TVESVTQ
763.3832 Da
108 − 114
LALQF
591.3501 Da
119 − 123
LLFGGVDEKGPQL
1372.7471 Da
141 − 153
FVQCDA
682.2865 Da
162 − 167
TVESVTQA
834.4203 Da
108 − 115
DADPGAMSRPFGVA
1390.642 Da
127 − 140
LFGGVDEKGPQL
1259.663 Da
142 − 153
VQCDA
535.2181 Da
163 − 167
TVESVTQAVSNL
1247.6478 Da
108 − 119
DADPGAMSRPFGVAL
1503.726 Da
127 − 141
FHMDPSGT
891.3665 Da
154 − 161
DARAIGSASEGAQSSL
1519.7347 Da
166 − 181
SVTQAVSNL
918.4891 Da
111 − 119
DADPGAMSRPFGVALL
1616.8101 Da
127 − 142
FHMDPSGTF
1038.4349 Da
154 − 162
RAIGSASEGAQSSL
1333.6706 Da
168 − 181
Relative Deuterium Uptake
Exposure (minutes)
State
i20S_apo
std20S_apo

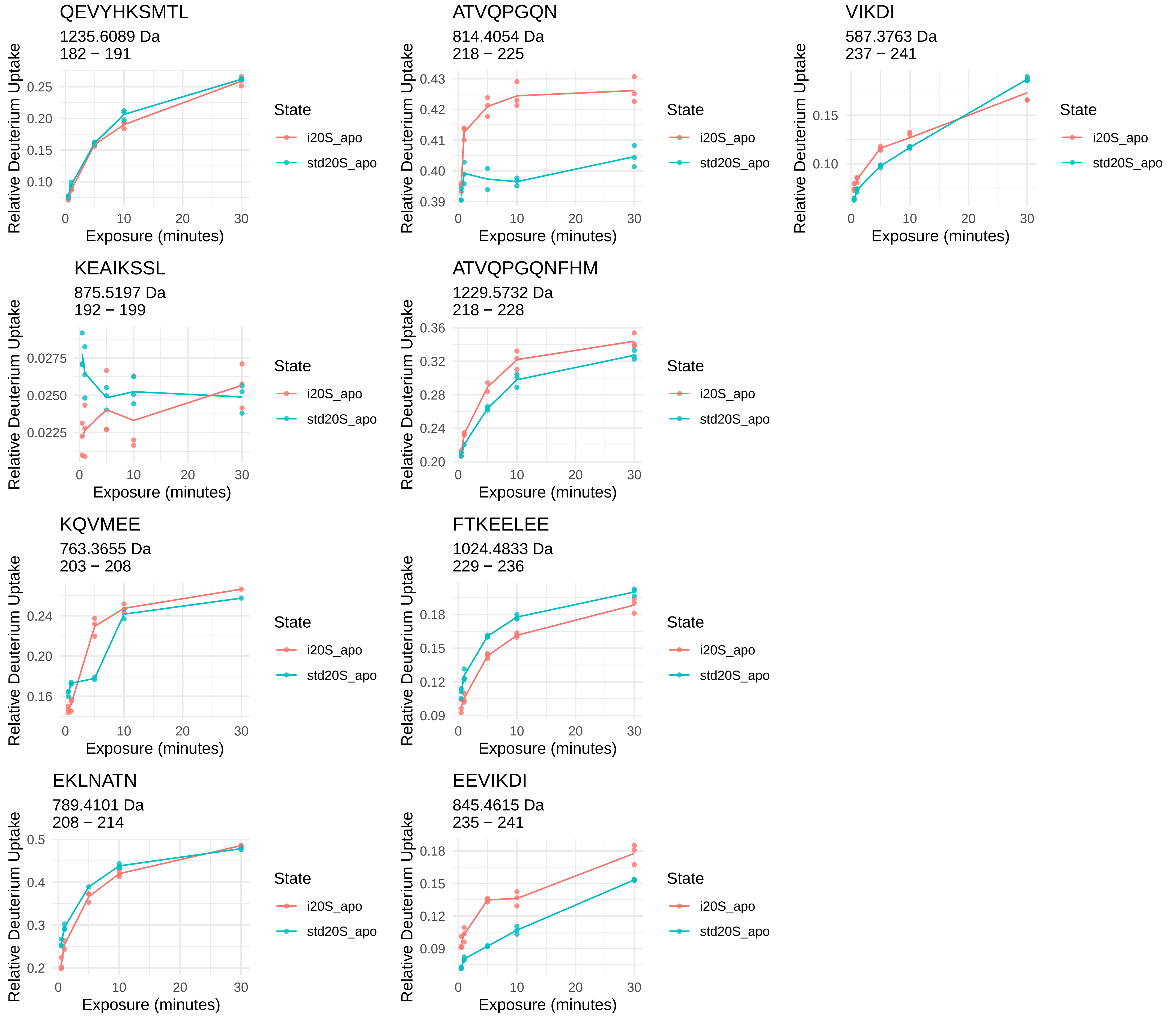
QEVYHKSMTL
1235.6089 Da
182 − 191
ATVQPGQN
814.4054 Da
218 − 225
VIKDI
587.3763 Da
237 − 241
KEAIKSSL
875.5197 Da
192 − 199
ATVQPGQNFHM
1229.5732 Da
218 − 228
KQVMEE
763.3655 Da
203 − 208
FTKEELEE
1024.4833 Da
229 − 236
EKLNATN
789.4101 Da
208 − 214
EEVIKDI
845.4615 Da
235 − 241
Relative Deuterium Uptake
Exposure (minutes)
State
i20S_apo
std20S_apo

α6

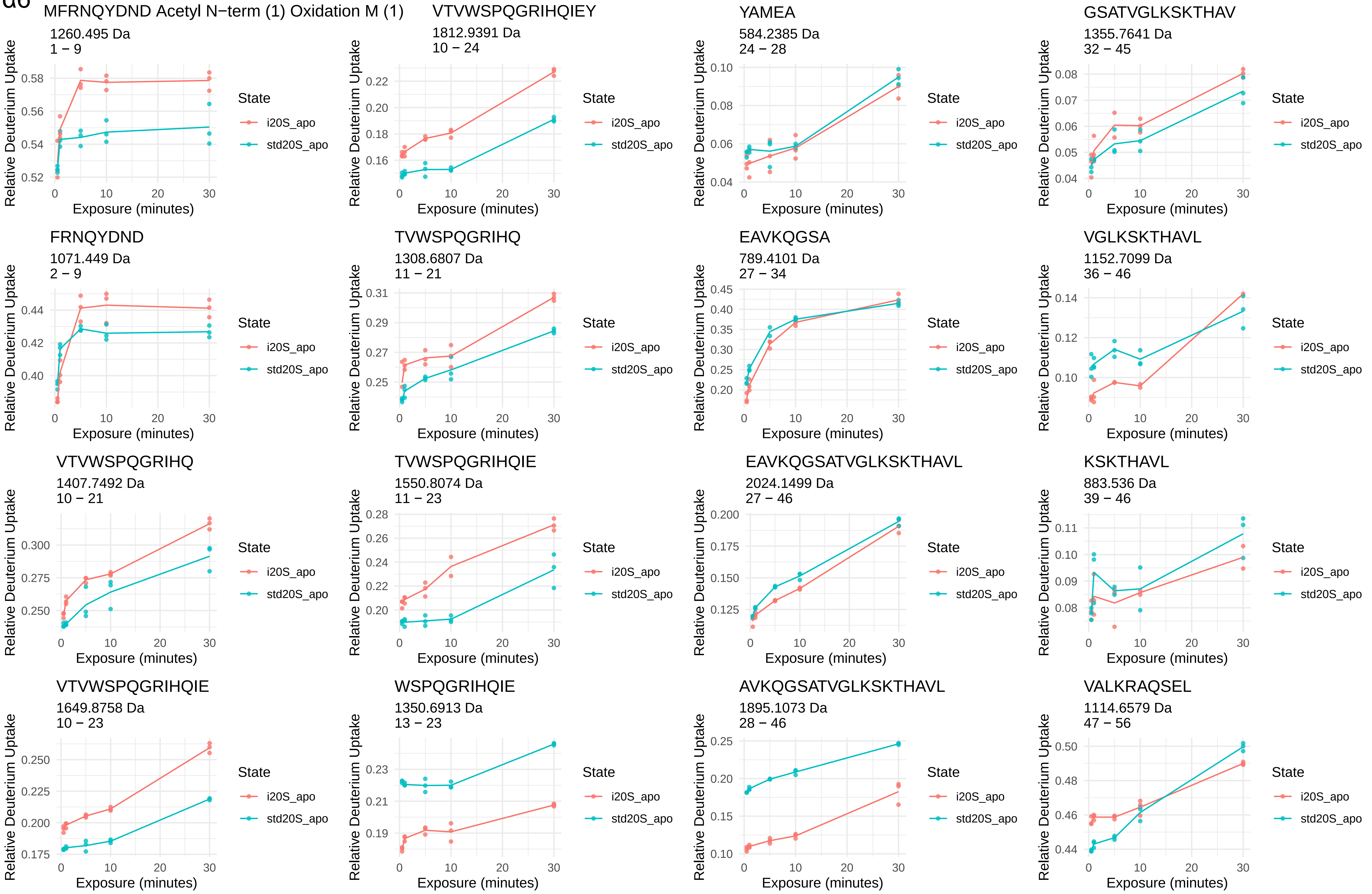
MFRNQYDND Acetyl N−term (1) Oxidation M (1)
1260.495 Da
1 − 9
VTVWSPQGRIHQIEY
1812.9391 Da
10 − 24
YAMEA
584.2385 Da
24 − 28
GSATVGLKSKTHAV
1355.7641 Da
32 − 45
FRNQYDND
1071.449 Da
2 − 9
TVWSPQGRIHQ
1308.6807 Da
11 − 21
EAVKQGSA
789.4101 Da
27 − 34
VGLKSKTHAVL
1152.7099 Da
36 − 46
VTVWSPQGRIHQ
1407.7492 Da
10 − 21
TVWSPQGRIHQIE
1550.8074 Da
11 − 23
EAVKQGSATVGLKSKTHAVL
2024.1499 Da
27 − 46
KSKTHAVL
883.536 Da
39 − 46
VTVWSPQGRIHQIE
1649.8758 Da
10 − 23
WSPQGRIHQIE
1350.6913 Da
13 − 23
AVKQGSATVGLKSKTHAVL
1895.1073 Da
28 − 46
VALKRAQSEL
1114.6579 Da
47 − 56
Relative Deuterium Uptake
Exposure (minutes)
State
i20S_apo
std20S_apo

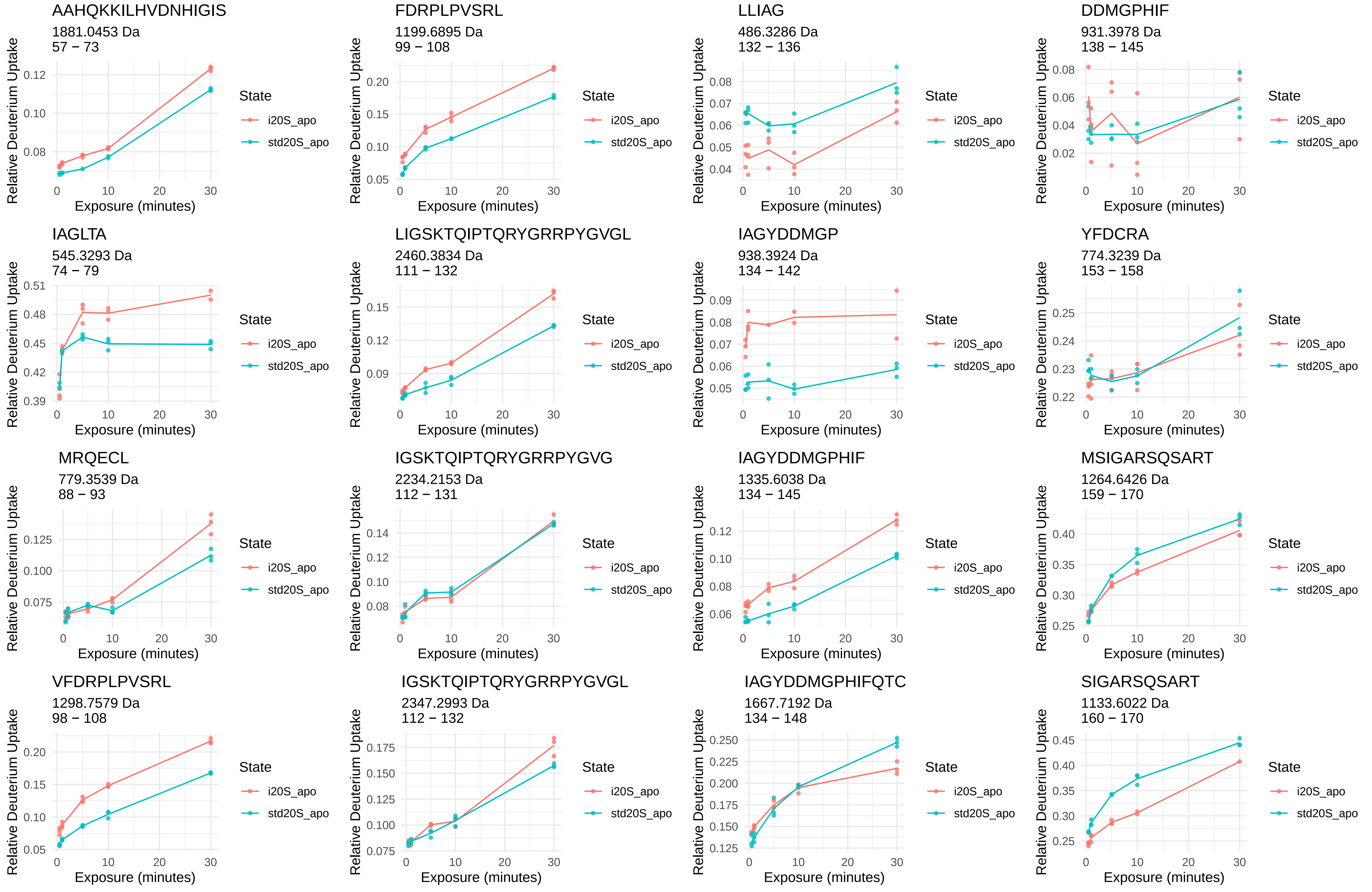
AAHQKKILHVDNHIGIS
1881.0453 Da
57 − 73
FDRPLPVSRL
1199.6895 Da
99 − 108
LLIAG
486.3286 Da
132 − 136
DDMGPHIF
931.3978 Da
138 − 145
IAGLTA
545.3293 Da
74 − 79
LIGSKTQIPTQRYGRRPYGVGL
2460.3834 Da
111 − 132
IAGYDDMGP
938.3924 Da
134 − 142
YFDCRA
774.3239 Da
153 − 158
MRQECL
779.3539 Da
88 − 93
IGSKTQIPTQRYGRRPYGVG
2234.2153 Da
112 − 131
IAGYDDMGPHIF
1335.6038 Da
134 − 145
MSIGARSQSART
1264.6426 Da
159 − 170
VFDRPLPVSRL
1298.7579 Da
98 − 108
IGSKTQIPTQRYGRRPYGVGL
2347.2993 Da
112 − 132
IAGYDDMGPHIFQTC
1667.7192 Da
134 − 148
SIGARSQSART
1133.6022 Da
160 − 170
Relative Deuterium Uptake
Exposure (minutes)
State
i20S_apo
std20S_apo

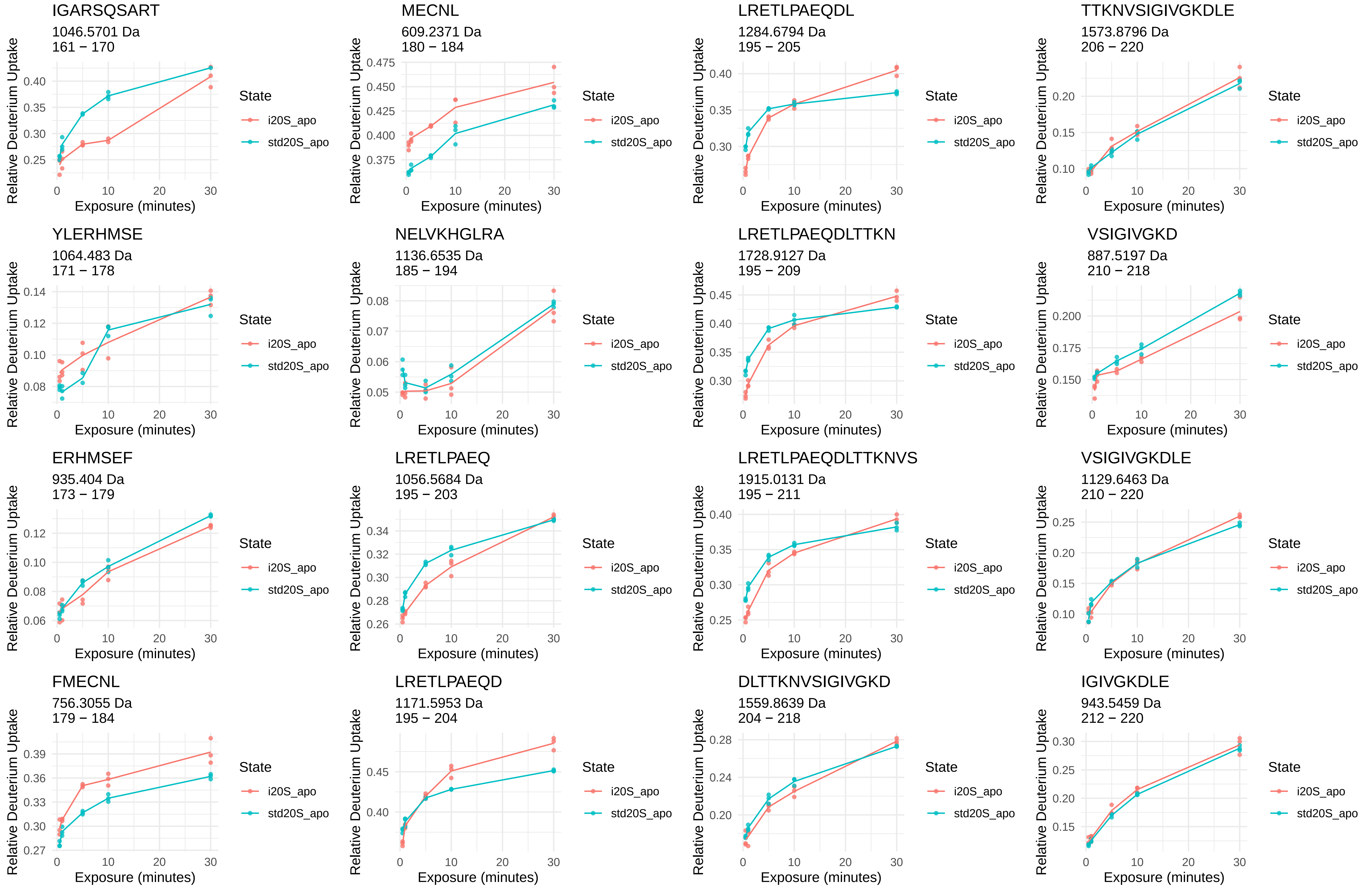
IGARSQSART
1046.5701 Da
161 − 170
MECNL
609.2371 Da
180 − 184
LRETLPAEQDL
1284.6794 Da
195 − 205
TTKNVSIGIVGKDLE
1573.8796 Da
206 − 220
YLERHMSE
1064.483 Da
171 − 178
NELVKHGLRA
1136.6535 Da
185 − 194
LRETLPAEQDLTTKN
1728.9127 Da
195 − 209
VSIGIVGKD
887.5197 Da
210 − 218
ERHMSEF
935.404 Da
173 − 179
LRETLPAEQ
1056.5684 Da
195 − 203
LRETLPAEQDLTTKNVS
1915.0131 Da
195 − 211
VSIGIVGKDLE
1129.6463 Da
210 − 220
FMECNL
756.3055 Da
179 − 184
LRETLPAEQD
1171.5953 Da
195 − 204
DLTTKNVSIGIVGKD
1559.8639 Da
204 − 218
IGIVGKDLE
943.5459 Da
212 − 220
Relative Deuterium Uptake
Exposure (minutes)
State
i20S_apo
std20S_apo

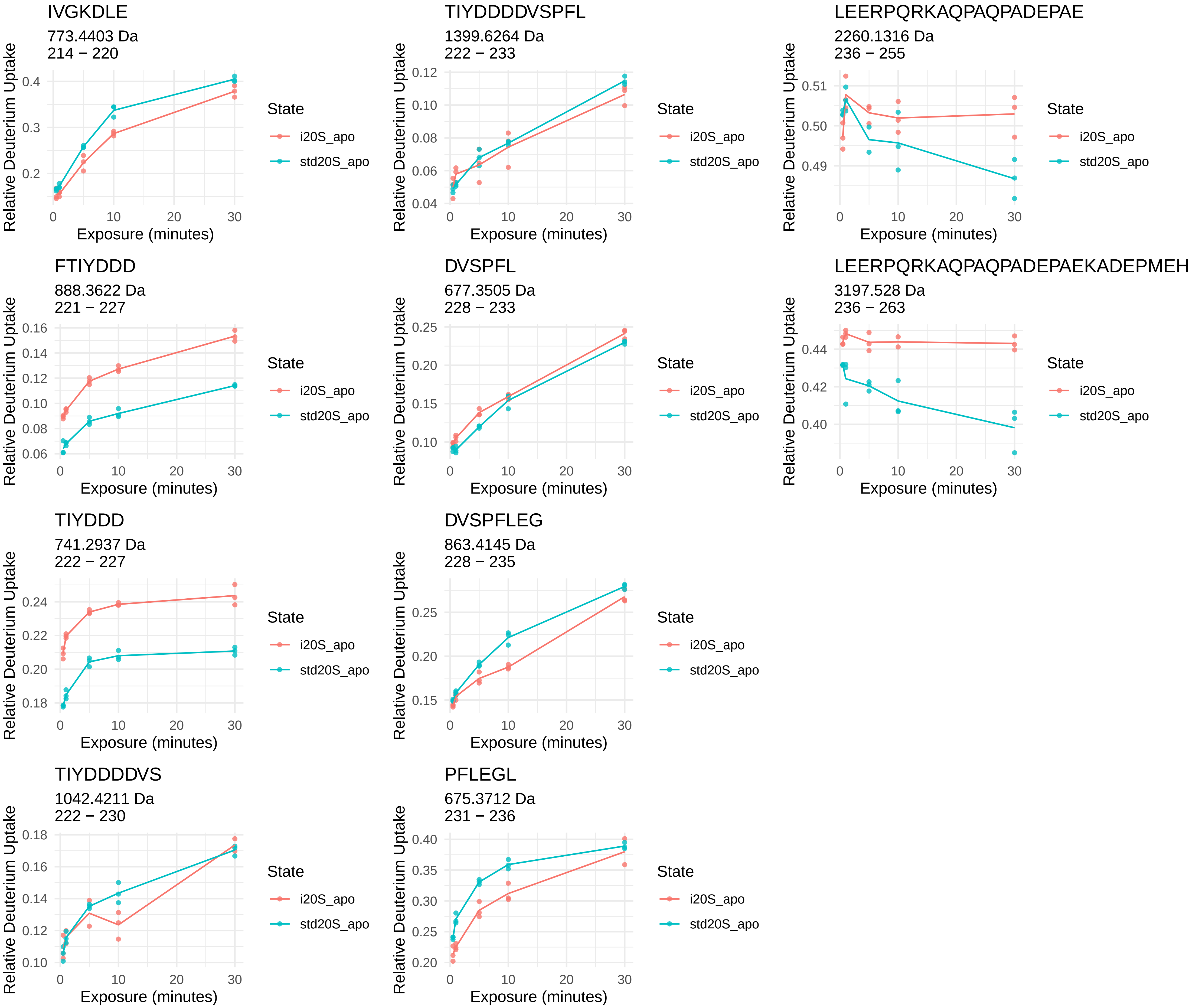
IVGKDLE
773.4403 Da
214 − 220
TIYDDDDVSPFL
1399.6264 Da
222 − 233
LEERPQRKAQPAQPADEPAE
2260.1316 Da
236 − 255
FTIYDDD
888.3622 Da
221 − 227
DVSPFL
677.3505 Da
228 − 233
LEERPQRKAQPAQPADEPAEKADEPMEH
3197.528 Da
236 − 263
TIYDDD
741.2937 Da
222 − 227
DVSPFLEG
863.4145 Da
228 − 235
TIYDDDDVS
1042.4211 Da
222 − 230
PFLEGL
675.3712 Da
231 − 236
Relative Deuterium Uptake
Exposure (minutes)
0
10
20
30
State
i20S_apo
std20S_apo

α7

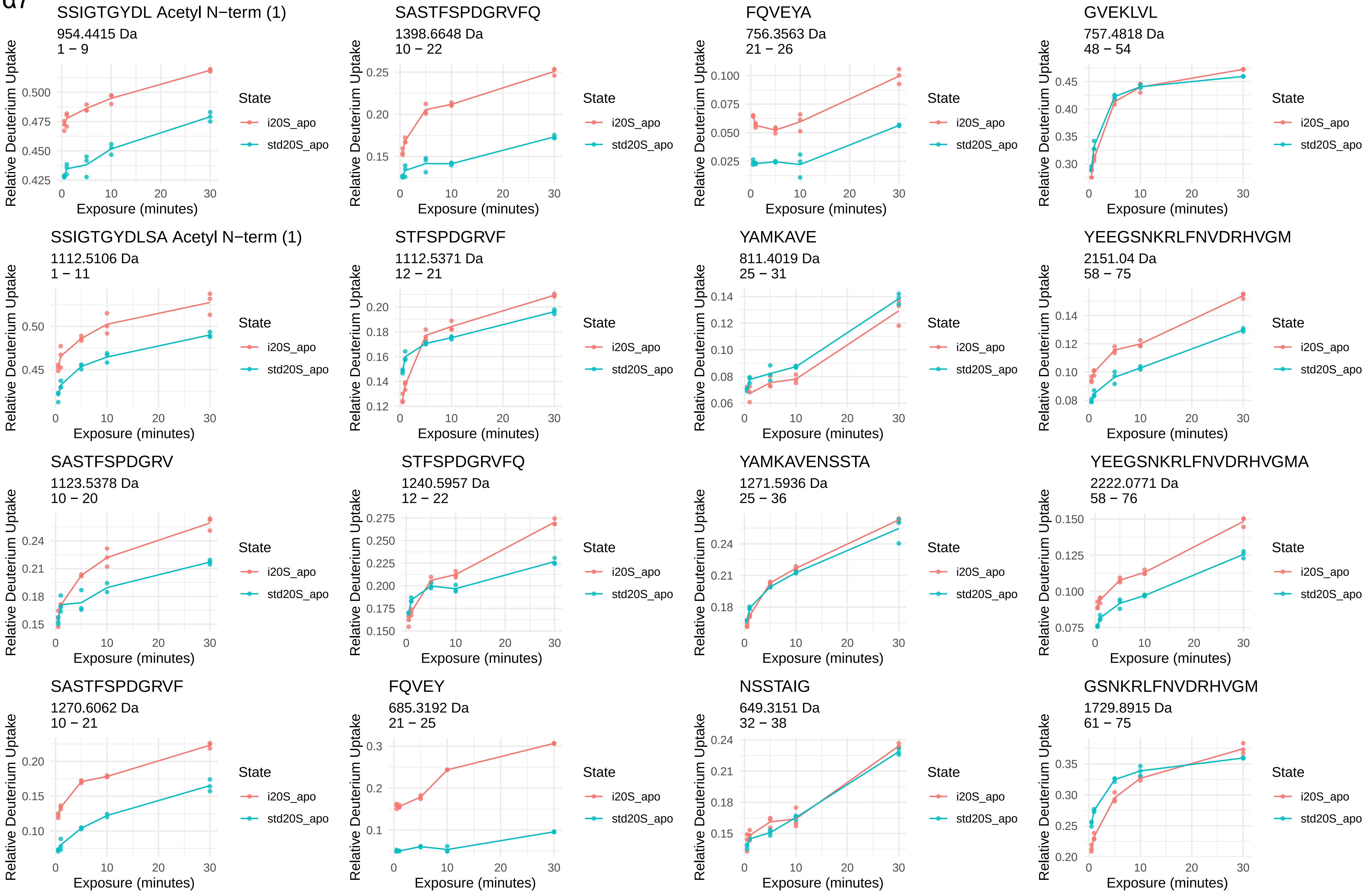
SSIGTGYDL Acetyl N−term (1)
954.4415 Da
1 − 9
SASTFSPDGRVFQ
1398.6648 Da
10 − 22
FQVEYA
756.3563 Da
21 − 26
GVEKLVL
757.4818 Da
48 − 54
SSIGTGYDLSA Acetyl N−term (1)
1112.5106 Da
1 − 11
STFSPDGRVF
1112.5371 Da
12 − 21
YAMKAVE
811.4019 Da
25 − 31
YEEGSNKRLFNVDRHVGM
2151.04 Da
58 − 75
SASTFSPDGRV
1123.5378 Da
10 − 20
STFSPDGRVFQ
1240.5957 Da
12 − 22
YAMKAVENSSTA
1271.5936 Da
25 − 36
YEEGSNKRLFNVDRHVGMA
2222.0771 Da
58 − 76
SASTFSPDGRVF
1270.6062 Da
10 − 21
FQVEY
685.3192 Da
21 − 25
NSSTAIG
649.3151 Da
32 − 38
GSNKRLFNVDRHVGM
1729.8915 Da
61 − 75
Relative Deuterium Uptake
Exposure (minutes)
State
i20S_apo
std20S_apo

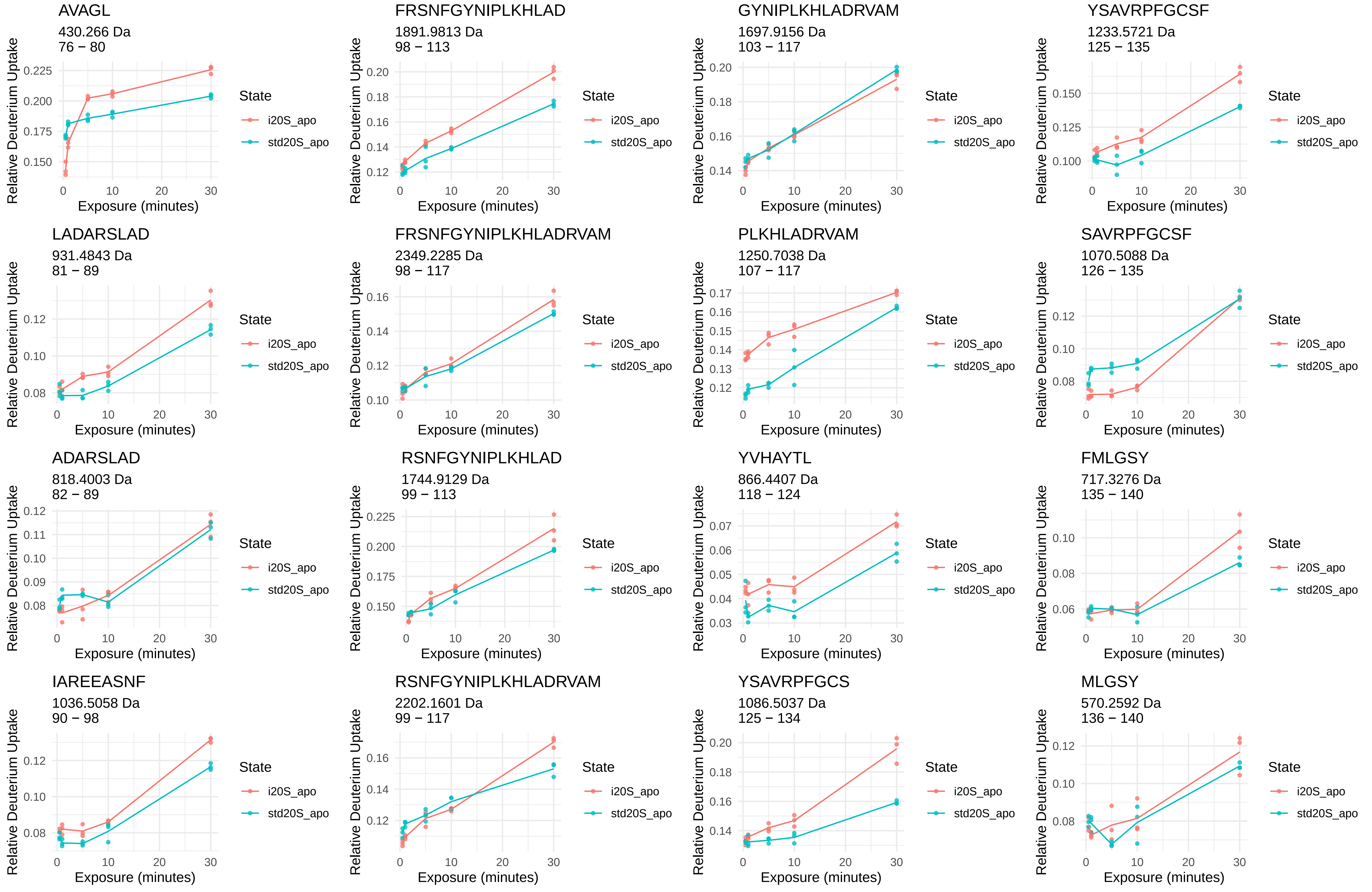
AVAGL
430.266 Da
76 − 80
FRSNFGYNIPLKHLAD
1891.9813 Da
98 − 113
GYNIPLKHLADRVAM
1697.9156 Da
103 − 117
YSAVRPFGCSF
1233.5721 Da
125 − 135
LADARSLAD
931.4843 Da
81 − 89
FRSNFGYNIPLKHLADRVAM
2349.2285 Da
98 − 117
PLKHLADRVAM
1250.7038 Da
107 − 117
SAVRPFGCSF
1070.5088 Da
126 − 135
ADARSLAD
818.4003 Da
82 − 89
RSNFGYNIPLKHLAD
1744.9129 Da
99 − 113
YVHAYTL
866.4407 Da
118 − 124
FMLGSY
717.3276 Da
135 − 140
IAREEASNF
1036.5058 Da
90 − 98
RSNFGYNIPLKHLADRVAM
2202.1601 Da
99 − 117
YSAVRPFGCS
1086.5037 Da
125 − 134
MLGSY
570.2592 Da
136 − 140
Relative Deuterium Uptake
Exposure (minutes)
State
i20S_apo
std20S_apo

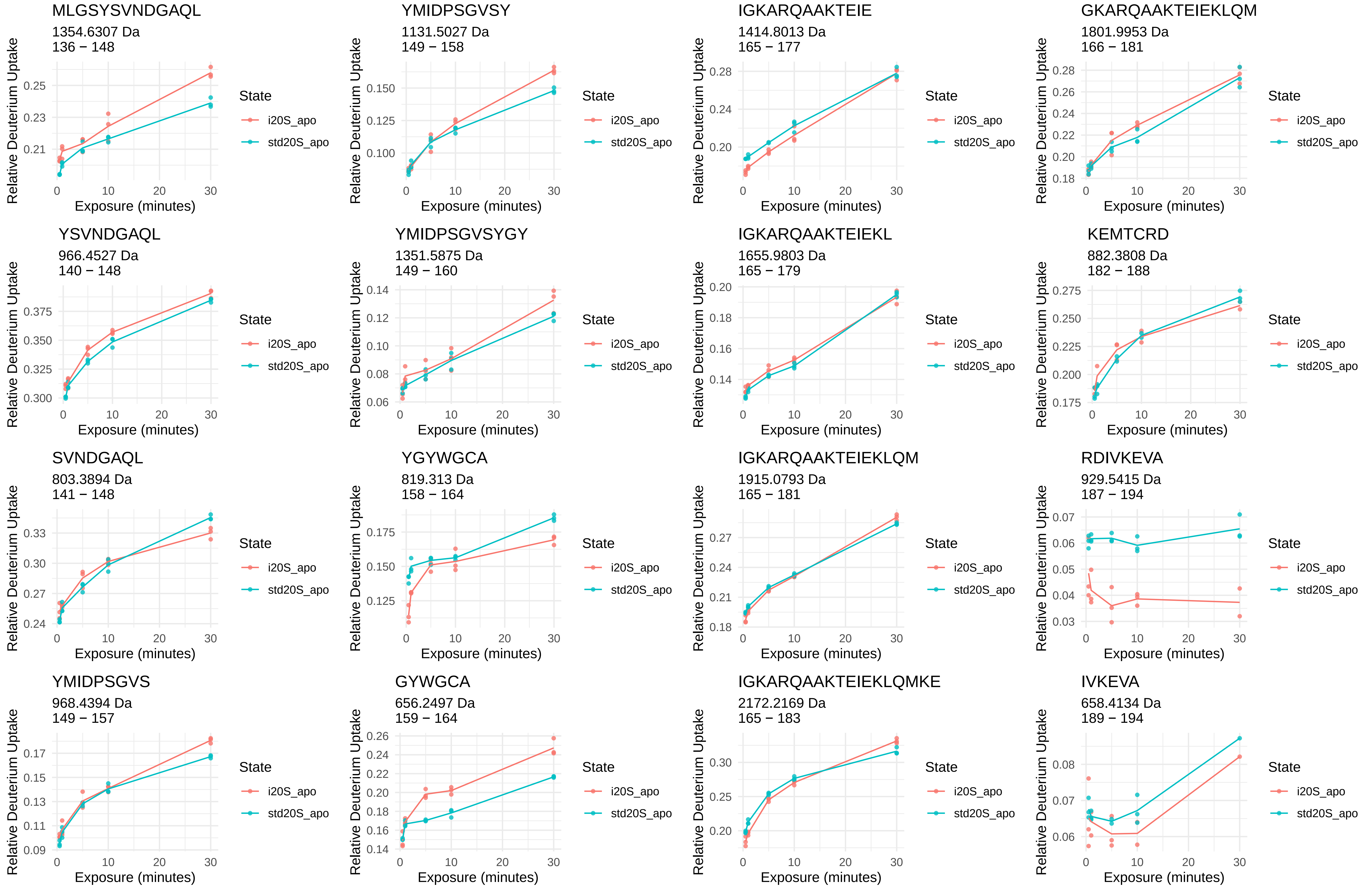
MLGSYSVNDGAQL
1354.6307 Da
136 − 148
0.21 0.23 0.25
YMIDPSGVSY
1131.5027 Da
149 − 158
0.100 0.125 0.150
IGKARQAAKTEIE
1414.8013 Da
165 − 177
0.20 0.24 0.28
GKARQAAKTEIEKLQM
1801.9953 Da
166 − 181
0.18 0.20 0.22 0.24 0.26 0.28
YSVNDGAQL
966.4527 Da
140 − 148
0.300 0.325 0.350 0.375
YMIDPSGVSYGY
1351.5875 Da
149 − 160
0.06 0.08 0.10 0.12 0.14
IGKARQAAKTEIEKL
1655.9803 Da
165 − 179
0.14 0.16 0.18 0.20
KEMTCRD
882.3808 Da
182 − 188
0.175 0.200 0.225 0.250 0.275
SVNDGAQL
803.3894 Da
141 − 148
0.24 0.27 0.30 0.33
YGYWGCA
819.313 Da
158 − 164
0.125 0.150 0.175
IGKARQAAKTEIEKLQM
1915.0793 Da
165 − 181
0.18 0.21 0.24 0.27
RDIVKEVA
929.5415 Da
187 − 194
0.03 0.04 0.05 0.06 0.07
YMIDPSGVS
968.4394 Da
149 − 157
0.09 0.11 0.13 0.15 0.17
GYWGCA
656.2497 Da
159 − 164
0.14 0.16 0.18 0.20 0.22 0.24 0.26
IGKARQAAKTEIEKLQMKE
2172.2169 Da
165 − 183
0.20 0.25 0.30
IVKEVA
658.4134 Da
189 − 194
0.06 0.07 0.08
Relative Deuterium Uptake
Exposure (minutes)
0 10 20 30
State
i20S_apo
std20S_apo

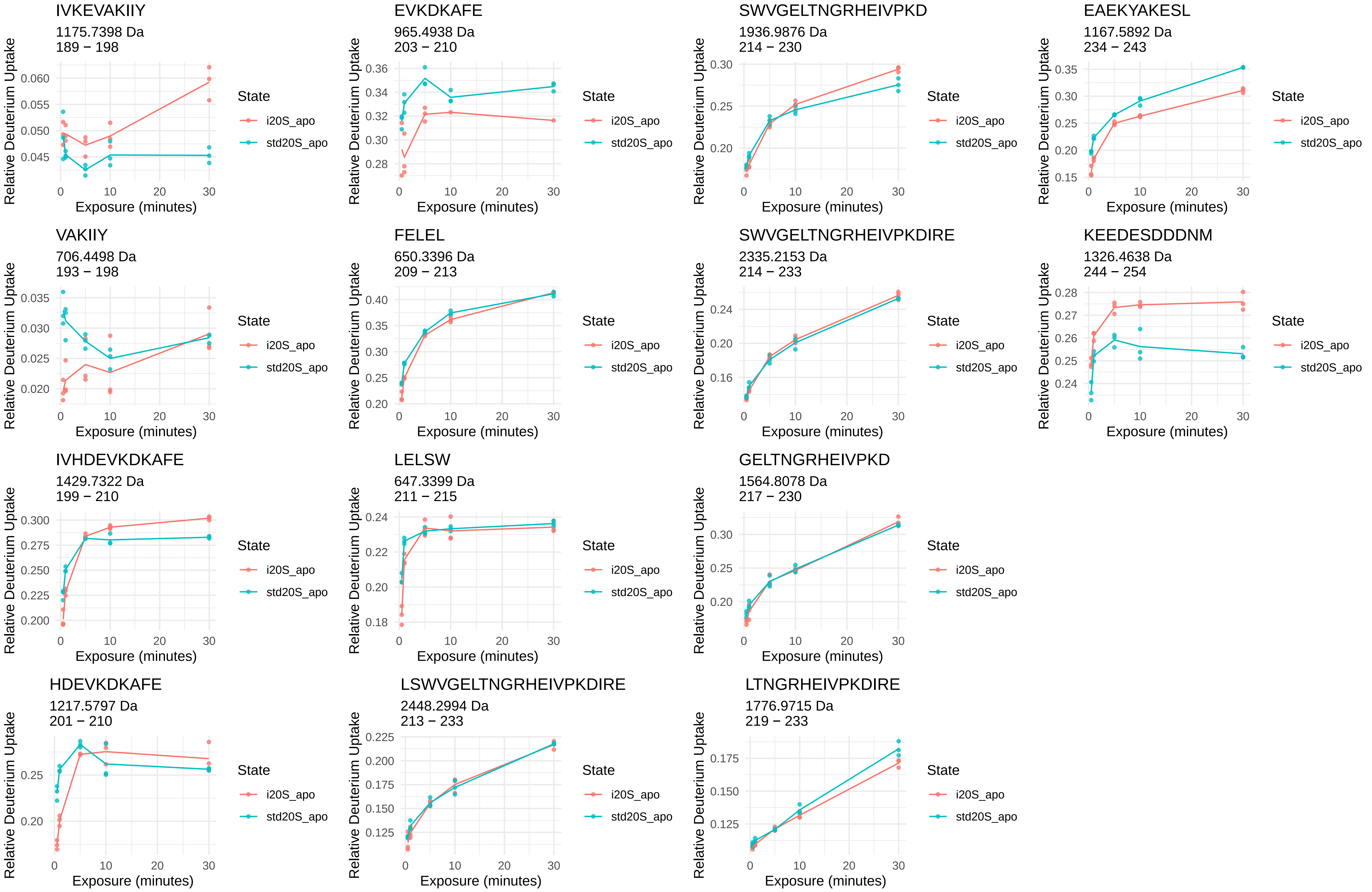
IVKEVAKIIY
1175.7398 Da
189 − 198
EVKDKAFE
965.4938 Da
203 − 210
SWVGELTNGRHEIVPKD
1936.9876 Da
214 − 230
EAEKYAKESL
1167.5892 Da
234 − 243
VAKIIY
706.4498 Da
193 − 198
FELEL
650.3396 Da
209 − 213
SWVGELTNGRHEIVPKDIRE
2335.2153 Da
214 − 233
KEEDESDDDNM
1326.4638 Da
244 − 254
IVHDEVKDKAFE
1429.7322 Da
199 − 210
LELSW
647.3399 Da
211 − 215
GELTNGRHEIVPKD
1564.8078 Da
217 − 230
HDEVKDKAFE
1217.5797 Da
201 − 210
LSWVGELTNGRHEIVPKDIRE
2448.2994 Da
213 − 233
LTNGRHEIVPKDIRE
1776.9715 Da
219 − 233
Relative Deuterium Uptake
Exposure (minutes)
State
i20S_apo
std20S_apo

β1

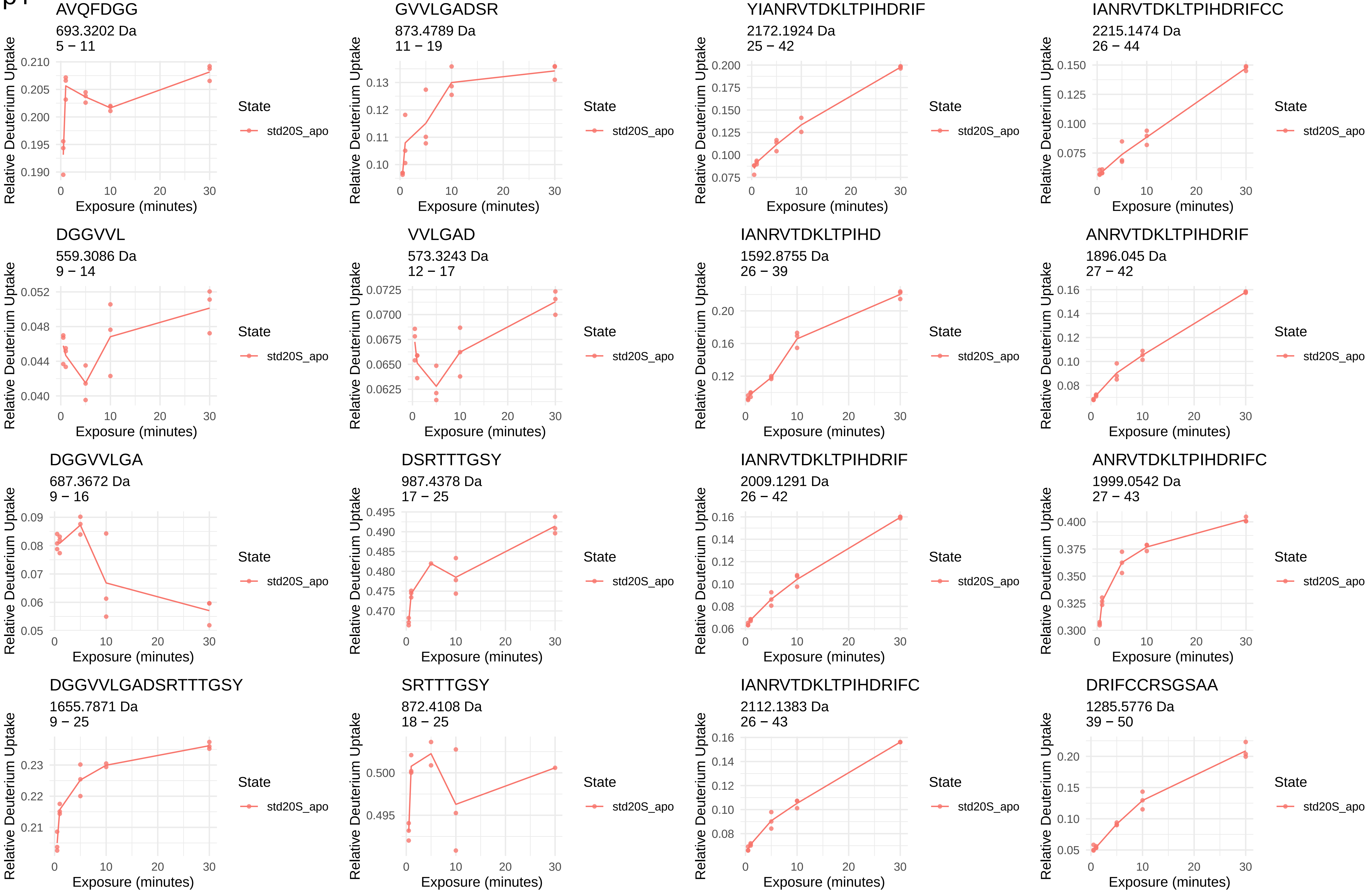
AVQFDGG
693.3202 Da
5 − 11
GVVLGADSR
873.4789 Da
11 − 19
YIANRVTDKLTPIHDRIF
2172.1924 Da
25 − 42
IANRVTDKLTPIHDRIFCC
2215.1474 Da
26 − 44
DGGVVL
559.3086 Da
9 − 14
VVLGAD
573.3243 Da
12 − 17
IANRVTDKLTPIHD
1592.8755 Da
26 − 39
ANRVTDKLTPIHDRIF
1896.045 Da
27 − 42
DGGVVLGA
687.3672 Da
9 − 16
DSRTTTGSY
987.4378 Da
17 − 25
IANRVTDKLTPIHDRIF
2009.1291 Da
26 − 42
ANRVTDKLTPIHDRIFC
1999.0542 Da
27 − 43
DGGVVLGADSRTTTGSY
1655.7871 Da
9 − 25
SRTTTGSY
872.4108 Da
18 − 25
IANRVTDKLTPIHDRIFC
2112.1383 Da
26 − 43
DRIFCCRSGSAA
1285.5776 Da
39 − 50
Relative Deuterium Uptake
Exposure (minutes)
State
std20S_apo

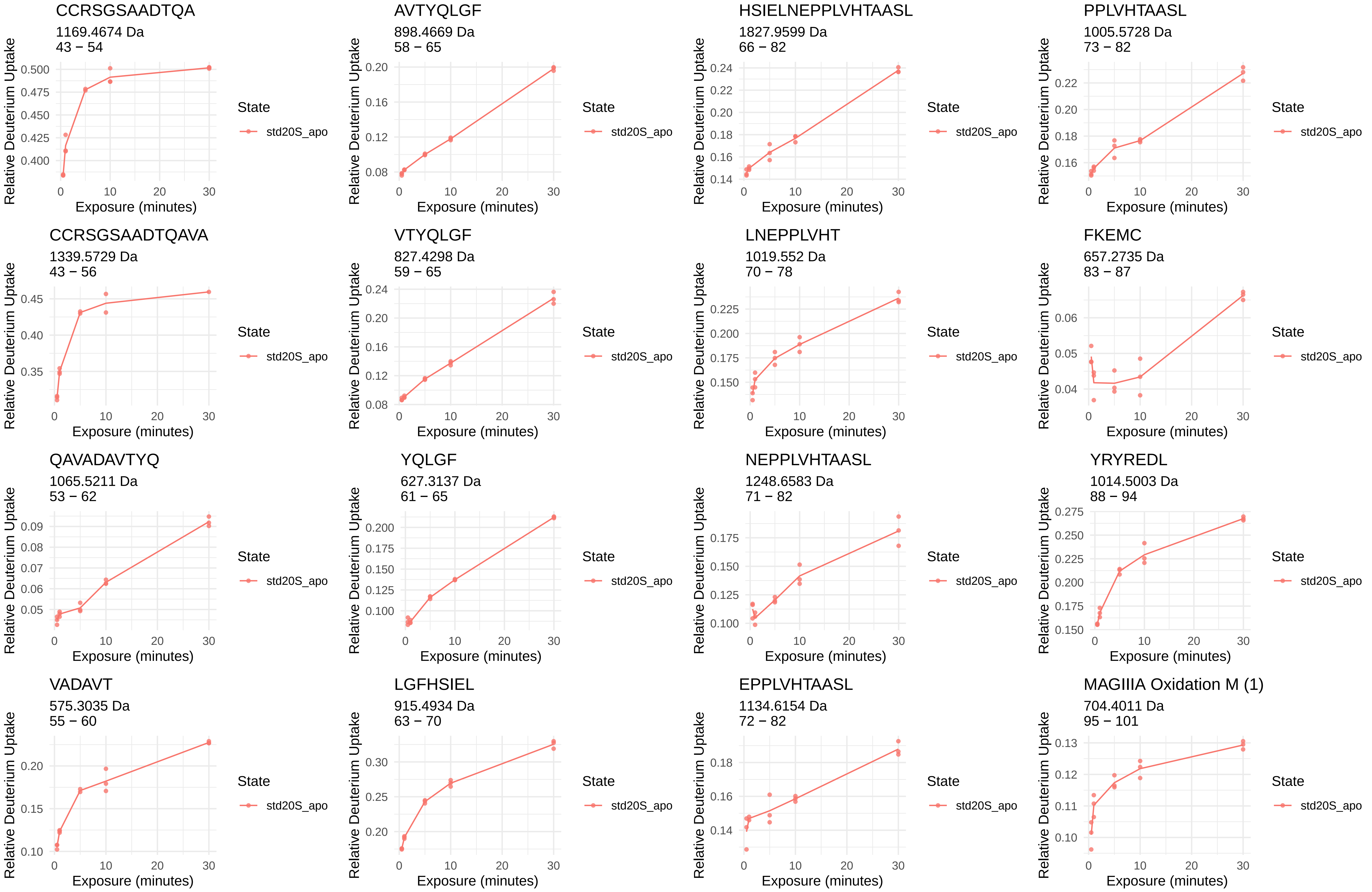
CCRSGSAADTQA
1169.4674 Da
43 − 54
AVTYQLGF
898.4669 Da
58 − 65
HSIELNEPPLVHTAASL
1827.9599 Da
66 − 82
PPLVHTAASL
1005.5728 Da
73 − 82
CCRSGSAADTQAVA
1339.5729 Da
43 − 56
VTYQLGF
827.4298 Da
59 − 65
LNEPPLVHT
1019.552 Da
70 − 78
FKEMC
657.2735 Da
83 − 87
QAVADAVTYQ
1065.5211 Da
53 − 62
YQLGF
627.3137 Da
61 − 65
NEPPLVHTAASL
1248.6583 Da
71 − 82
YRYREDL
1014.5003 Da
88 − 94
VADAVT
575.3035 Da
55 − 60
LGFHSIEL
915.4934 Da
63 − 70
EPPLVHTAASL
1134.6154 Da
72 − 82
MAGIIIA Oxidation M (1)
704.4011 Da
95 − 101
Relative Deuterium Uptake
Exposure (minutes)
State
std20S_apo

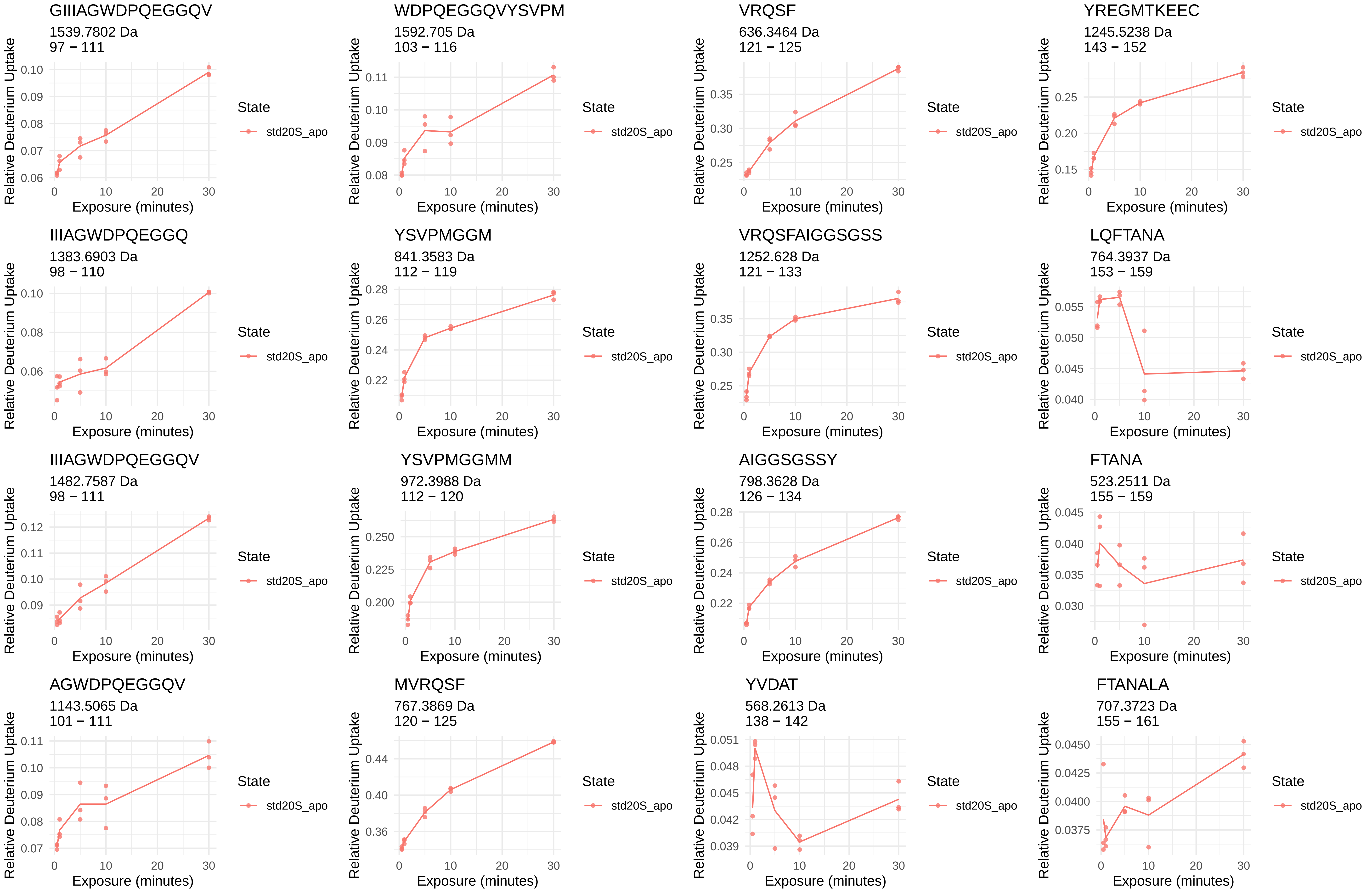
GIIIAGWDPQEGGQV
1539.7802 Da
97 − 111
WDPQEGGQVYSVPM
1592.705 Da
103 − 116
VRQSF
636.3464 Da
121 − 125
YREGMTKEEC
1245.5238 Da
143 − 152
IIIAGWDPQEGGQ
1383.6903 Da
98 − 110
YSVPMGGM
841.3583 Da
112 − 119
VRQSFAIGGSGSS
1252.628 Da
121 − 133
LQFTANA
764.3937 Da
153 − 159
IIIAGWDPQEGGQV
1482.7587 Da
98 − 111
YSVPMGGMM
972.3988 Da
112 − 120
AIGGSGSSY
798.3628 Da
126 − 134
FTANA
523.2511 Da
155 − 159
AGWDPQEGGQV
1143.5065 Da
101 − 111
MVRQSF
767.3869 Da
120 − 125
YVDAT
568.2613 Da
138 − 142
FTANALA
707.3723 Da
155 − 161
Relative Deuterium Uptake
Exposure (minutes)
State
std20S_apo

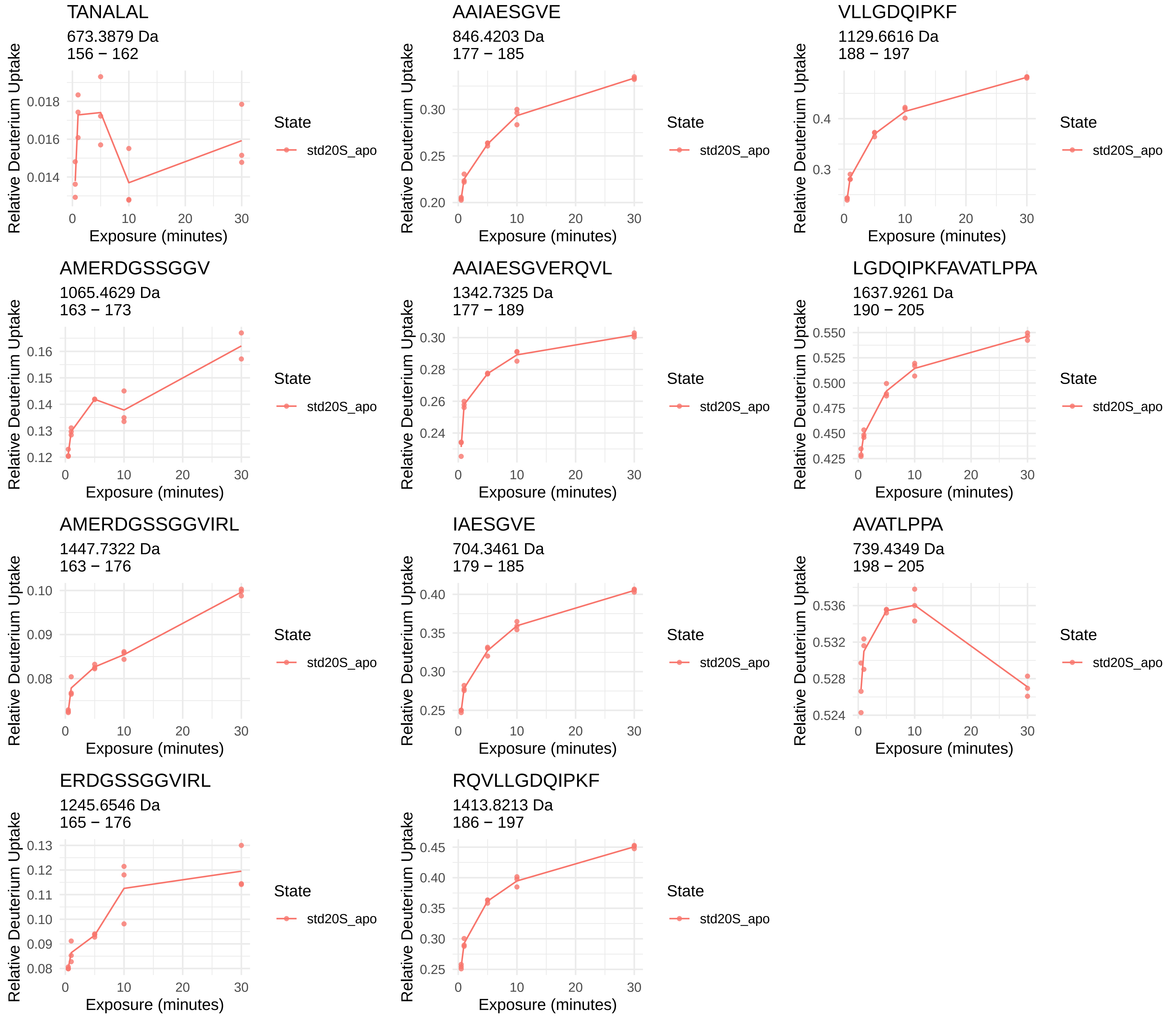
TANALAL
673.3879 Da
156 − 162
AAIAESGVE
846.4203 Da
177 − 185
VLLGDQIPKF
1129.6616 Da
188 − 197
AMERDGSSGGV
1065.4629 Da
163 − 173
AAIAESGVERQVL
1342.7325 Da
177 − 189
LGDQIPKFAVATLPPA
1637.9261 Da
190 − 205
AMERDGSSGGVIRL
1447.7322 Da
163 − 176
IAESGVE
704.3461 Da
179 − 185
AVATLPPA
739.4349 Da
198 − 205
ERDGSSGGVIRL
1245.6546 Da
165 − 176
RQVLLGDQIPKF
1413.8213 Da
186 − 197
Relative Deuterium Uptake
Exposure (minutes)
State
std20S_apo

β1i

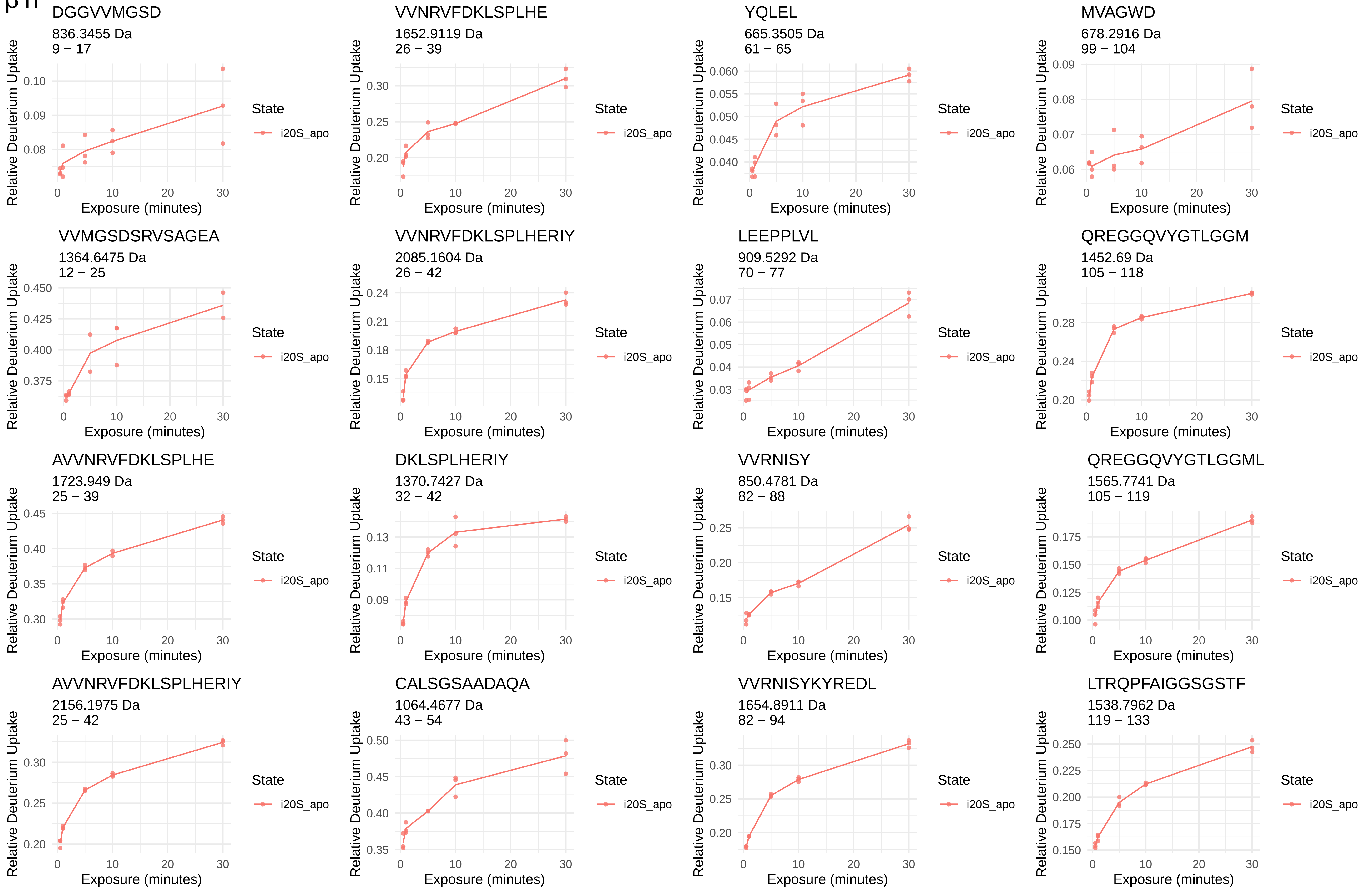
DGGVVMGSD
836.3455 Da
9 − 17
VVNRVFDKLSPLHE
1652.9119 Da
26 − 39
YQLEL
665.3505 Da
61 − 65
MVAGWD
678.2916 Da
99 − 104
VVMGSDSRVSAGEA
1364.6475 Da
12 − 25
VVNRVFDKLSPLHERIY
2085.1604 Da
26 − 42
LEEPPLVL
909.5292 Da
70 − 77
QREGGQVYGTLGGM
1452.69 Da
105 − 118
AVVNRVFDKLSPLHE
1723.949 Da
25 − 39
DKLSPLHERIY
1370.7427 Da
32 − 42
VVRNISY
850.4781 Da
82 − 88
QREGGQVYGTLGGML
1565.7741 Da
105 − 119
AVVNRVFDKLSPLHERIY
2156.1975 Da
25 − 42
CALSGSAADAQA
1064.4677 Da
43 − 54
VVRNISYKYREDL
1654.8911 Da
82 − 94
LTRQPFAIGGSGSTF
1538.7962 Da
119 − 133
Relative Deuterium Uptake
Exposure (minutes)
State
i20S_apo

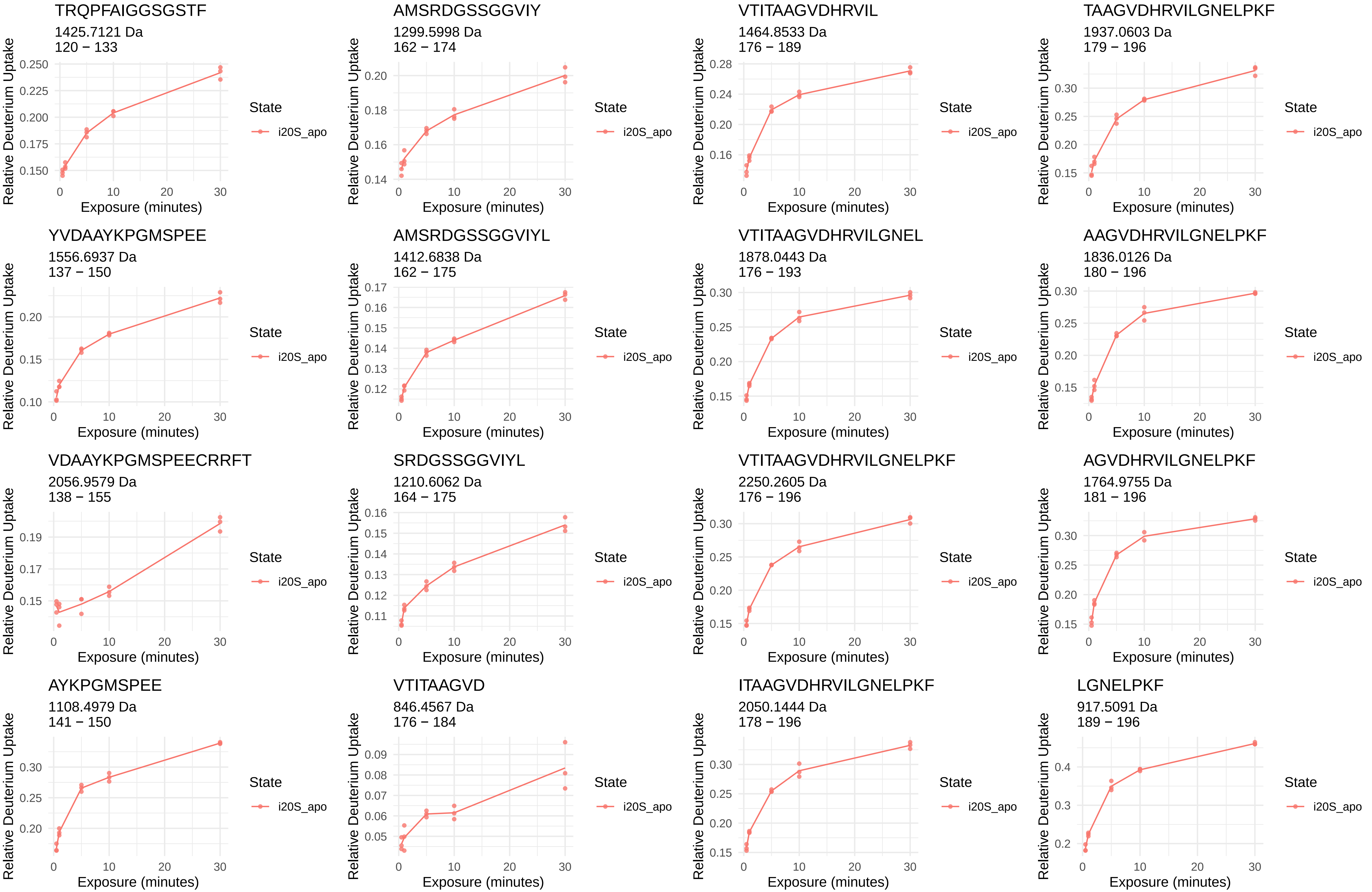
TRQPFAIGGSGSTF
1425.7121 Da
120 − 133
AMSRDGSSGGVIY
1299.5998 Da
162 − 174
VTITAAGVDHRVIL
1464.8533 Da
176 − 189
TAAGVDHRVILGNELPKF
1937.0603 Da
179 − 196
YVDAAYKPGMSPEE
1556.6937 Da
137 − 150
AMSRDGSSGGVIYL
1412.6838 Da
162 − 175
VTITAAGVDHRVILGNEL
1878.0443 Da
176 − 193
AAGVDHRVILGNELPKF
1836.0126 Da
180 − 196
VDAAYKPGMSPEECRRFT
2056.9579 Da
138 − 155
SRDGSSGGVIYL
1210.6062 Da
164 − 175
VTITAAGVDHRVILGNELPKF
2250.2605 Da
176 − 196
AGVDHRVILGNELPKF
1764.9755 Da
181 − 196
AYKPGMSPEE
1108.4979 Da
141 − 150
VTITAAGVD
846.4567 Da
176 − 184
ITAAGVDHRVILGNELPKF
2050.1444 Da
178 − 196
LGNELPKF
917.5091 Da
189 − 196
Relative Deuterium Uptake
Exposure (minutes)
State
i20S_apo

β2

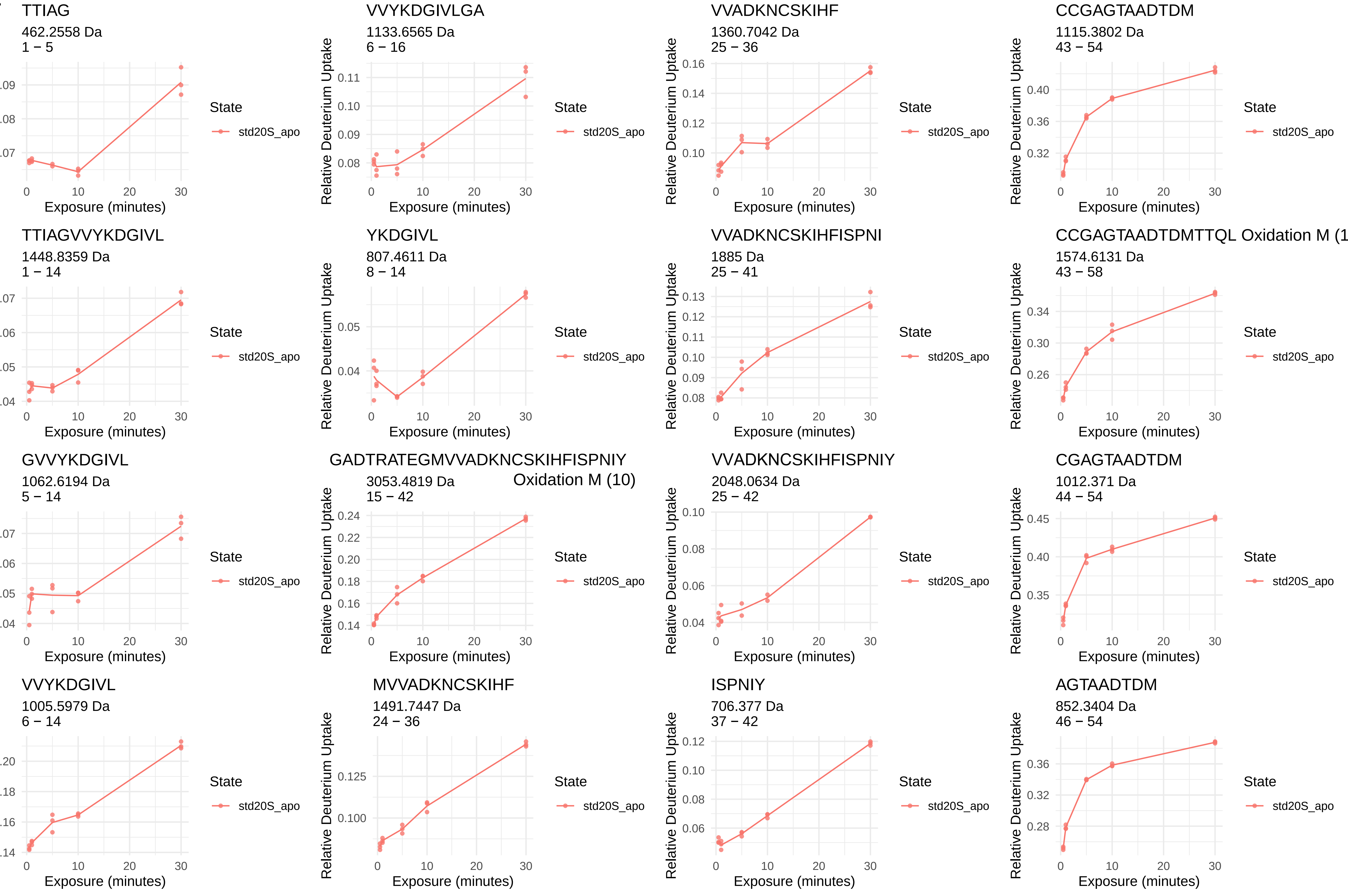
TTIAG
462.2558 Da
1 − 5
VVYKDGIVLGA
1133.6565 Da
6 − 16
VVADKNCSKIHF
1360.7042 Da
25 − 36
CCGAGTAADTDM
1115.3802 Da
43 − 54
TTIAGVVYKDGIVL
1448.8359 Da
1 − 14
YKDGIVL
807.4611 Da
8 − 14
VVADKNCSKIHFISPNI
1885 Da
25 − 41
CCGAGTAADTDMTTQL Oxidation M (1
1574.6131 Da
43 − 58
GVVYKDGIVL
1062.6194 Da
5 − 14
GADTRATEGMVVADKNCSKIHFISPNIY
Oxidation M (10)
3053.4819 Da
15 − 42
VVADKNCSKIHFISPNIY
2048.0634 Da
25 − 42
CGAGTAADTDM
1012.371 Da
44 − 54
VVYKDGIVL
1005.5979 Da
6 − 14
MVVADKNCSKIHF
1491.7447 Da
24 − 36
ISPNIY
706.377 Da
37 − 42
AGTAADTDM
852.3404 Da
46 − 54
Relative Deuterium Uptake
Exposure (minutes)
State
std20S_apo

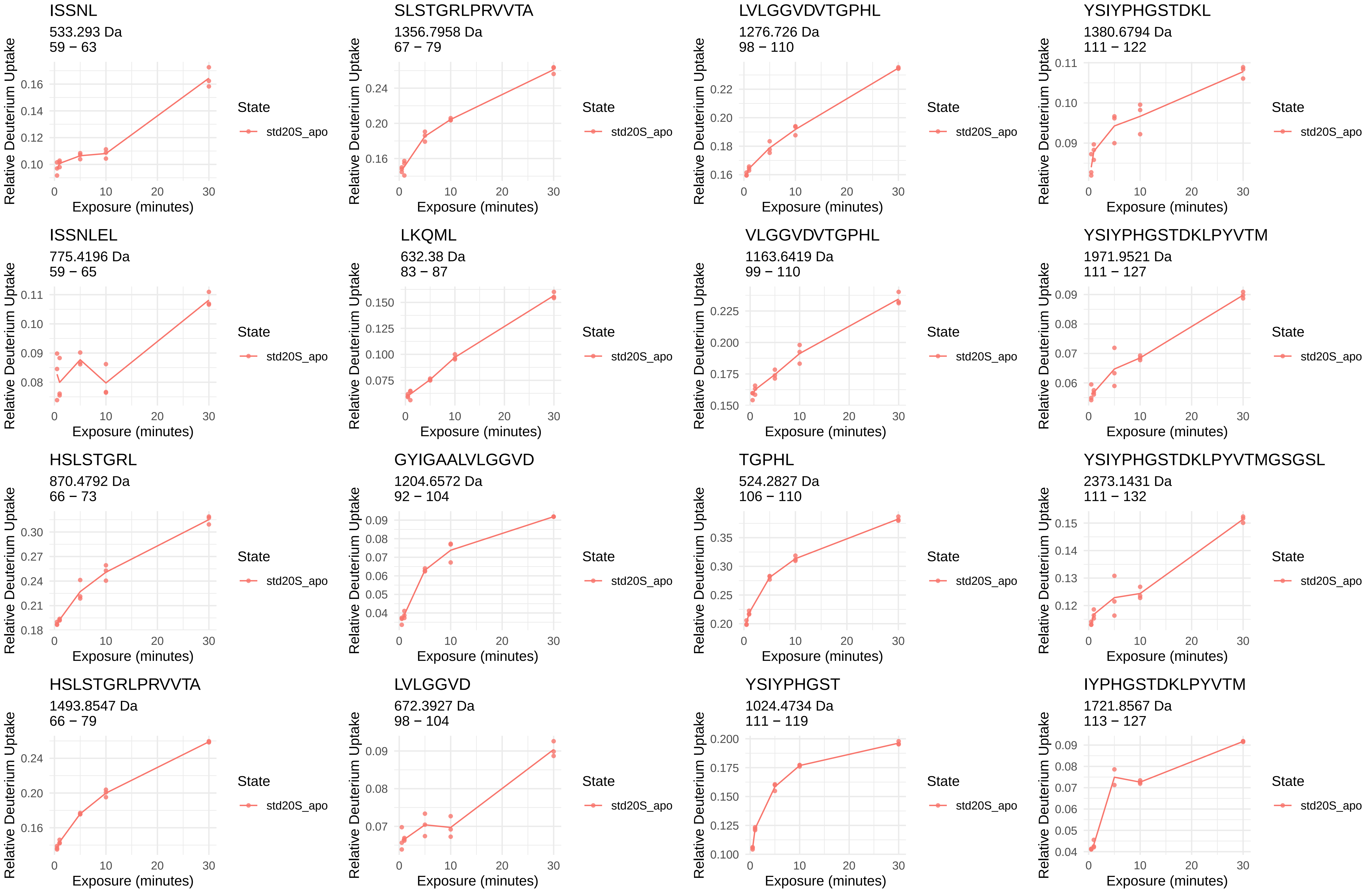
ISSNL
533.293 Da
59 − 63
SLSTGRLPRVVTA
1356.7958 Da
67 − 79
LVLGGVDVTGPHL
1276.726 Da
98 − 110
YSIYPHGSTDKL
1380.6794 Da
111 − 122
ISSNLEL
775.4196 Da
59 − 65
LKQML
632.38 Da
83 − 87
VLGGVDVTGPHL
1163.6419 Da
99 − 110
YSIYPHGSTDKLPYVTM
1971.9521 Da
111 − 127
HSLSTGRL
870.4792 Da
66 − 73
GYIGAALVLGGVD
1204.6572 Da
92 − 104
TGPHL
524.2827 Da
106 − 110
YSIYPHGSTDKLPYVTMGSGSL
2373.1431 Da
111 − 132
HSLSTGRLPRVVTA
1493.8547 Da
66 − 79
LVLGGVD
672.3927 Da
98 − 104
YSIYPHGST
1024.4734 Da
111 − 119
IYPHGSTDKLPYVTM
1721.8567 Da
113 − 127
Relative Deuterium Uptake
Exposure (minutes)
State
std20S_apo

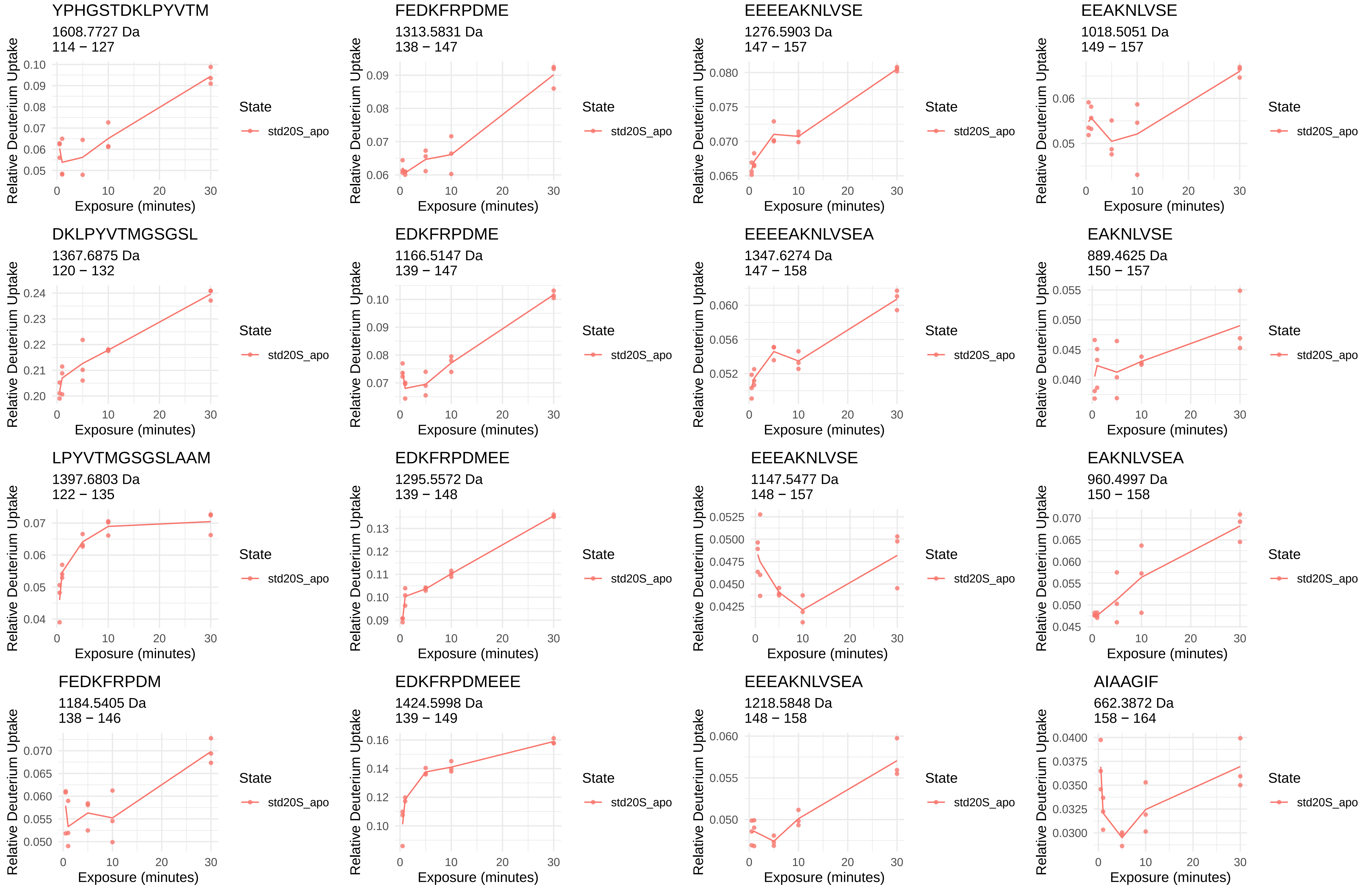
YPHGSTDKLPYVTM
1608.7727 Da
114 − 127
FEDKFRPDME
1313.5831 Da
138 − 147
EEEEAKNLVSE
1276.5903 Da
147 − 157
EEAKNLVSE
1018.5051 Da
149 − 157
DKLPYVTMGSGSL
1367.6875 Da
120 − 132
EDKFRPDME
1166.5147 Da
139 − 147
EEEEAKNLVSEA
1347.6274 Da
147 − 158
EAKNLVSE
889.4625 Da
150 − 157
LPYVTMGSGSLAAM
1397.6803 Da
122 − 135
EDKFRPDMEE
1295.5572 Da
139 − 148
EEEAKNLVSE
1147.5477 Da
148 − 157
EAKNLVSEA
960.4997 Da
150 − 158
FEDKFRPDM
1184.5405 Da
138 − 146
EDKFRPDMEEE
1424.5998 Da
139 − 149
EEEAKNLVSEA
1218.5848 Da
148 − 158
AIAAGIF
662.3872 Da
158 − 164
Relative Deuterium Uptake
Exposure (minutes)
State
std20S_apo

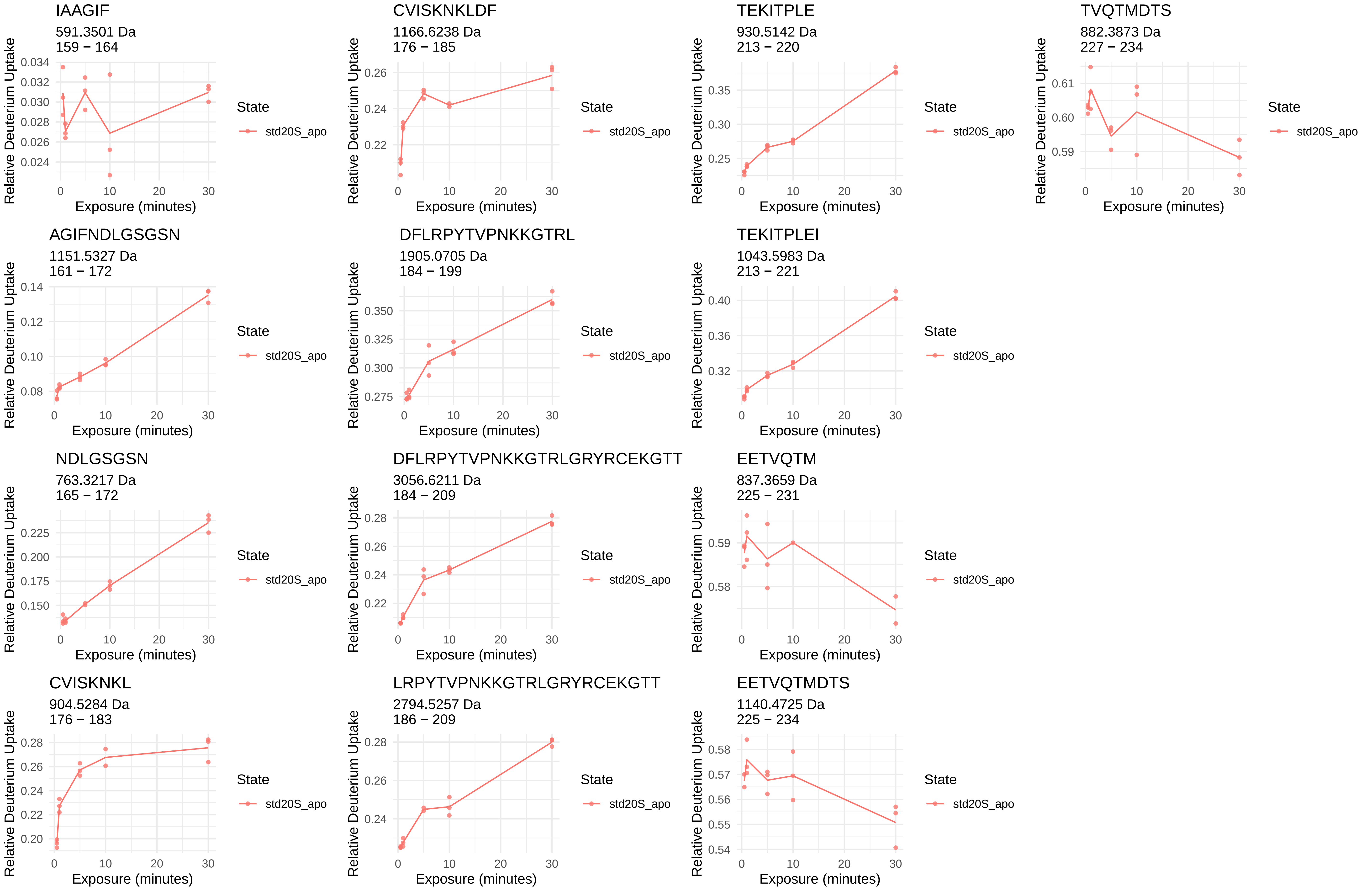
IAAGIF
591.3501 Da
159 − 164
CVISKNKLDF
1166.6238 Da
176 − 185
TEKITPLE
930.5142 Da
213 − 220
TVQTMDTS
882.3873 Da
227 − 234
AGIFNDLGSGSN
1151.5327 Da
161 − 172
DFLRPYTVPNKKGTRL
1905.0705 Da
184 − 199
TEKITPLEI
1043.5983 Da
213 − 221
NDLGSGSN
763.3217 Da
165 − 172
DFLRPYTVPNKKGTRLGRYRCEKGTT
3056.6211 Da
184 − 209
EETVQTM
837.3659 Da
225 − 231
CVISKNKL
904.5284 Da
176 − 183
LRPYTVPNKKGTRLGRYRCEKGTT
2794.5257 Da
186 − 209
EETVQTMDTS
1140.4725 Da
225 − 234
Relative Deuterium Uptake
Exposure (minutes)
State
std20S_apo

β2i

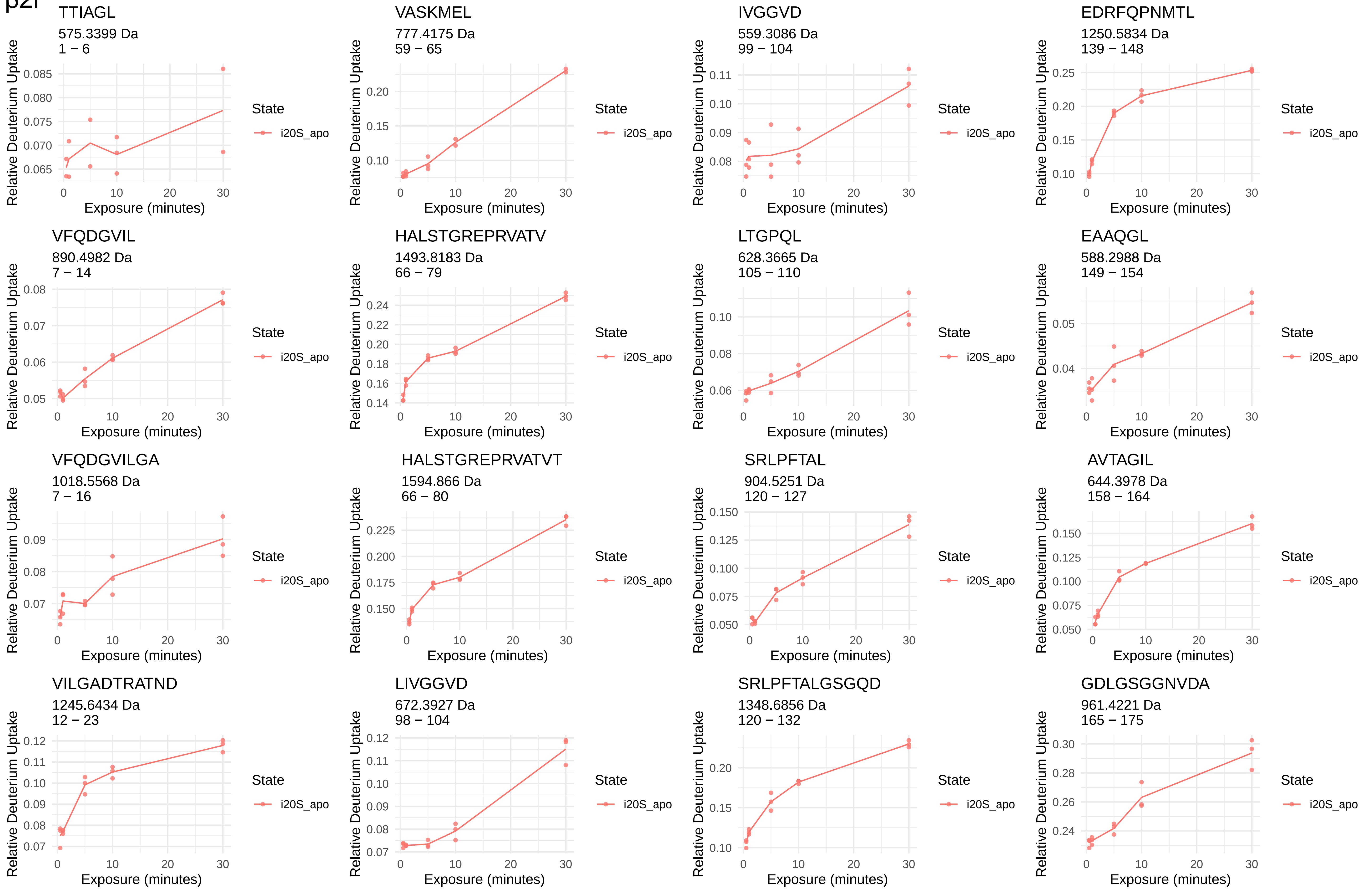
TTIAGL
575.3399 Da
1 − 6
VASKMEL
777.4175 Da
59 − 65
IVGGVD
559.3086 Da
99 − 104
EDRFQPNMTL
1250.5834 Da
139 − 148
VFQDGVIL
890.4982 Da
7 − 14
HALSTGREPRVATV
1493.8183 Da
66 − 79
LTGPQL
628.3665 Da
105 − 110
EAAQGL
588.2988 Da
149 − 154
VFQDGVILGA
1018.5568 Da
7 − 16
HALSTGREPRVATVT
1594.866 Da
66 − 80
SRLPFTAL
904.5251 Da
120 − 127
AVTAGIL
644.3978 Da
158 − 164
VILGADTRATND
1245.6434 Da
12 − 23
LIVGGVD
672.3927 Da
98 − 104
SRLPFTALGSGQD
1348.6856 Da
120 − 132
GDLGSGGNVDA
961.4221 Da
165 − 175
Relative Deuterium Uptake
Exposure (minutes)
State
i20S_apo

CVITKTGAKL
1033.6074 Da
176 − 185

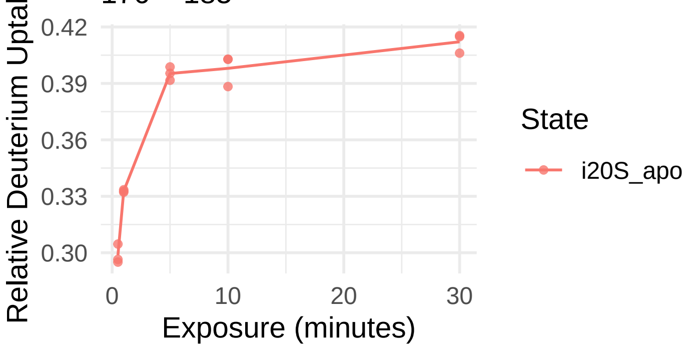


TQTVKPLTLEL
1242.7304 Da
213 − 223

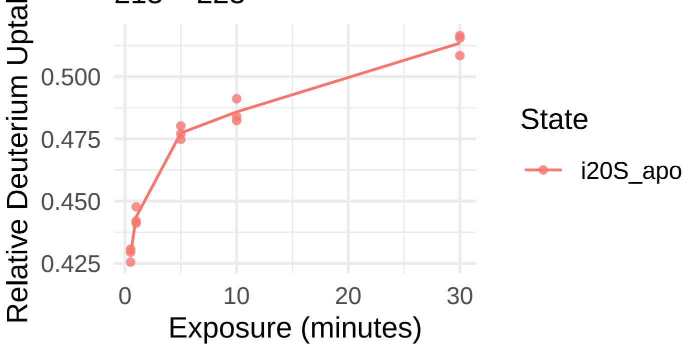


LRTLSSPTEPVKRSGRYHFVPGTT
2686.4424 Da
186 − 209

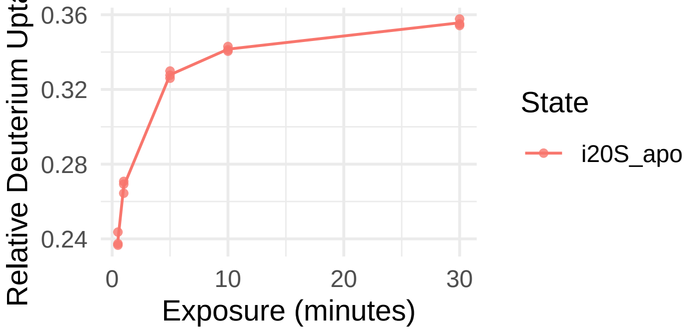


TQTVKPLTL
1000.6037 Da
213 − 221

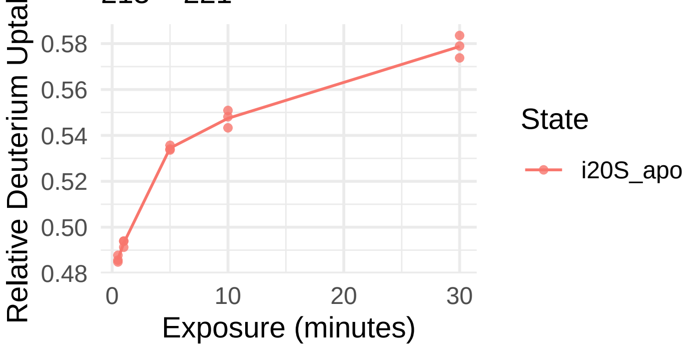


TQTVKPLTLE
1129.6463 Da
213 − 222

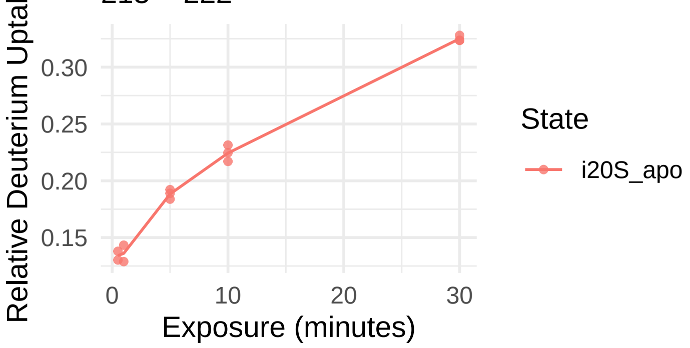

β3

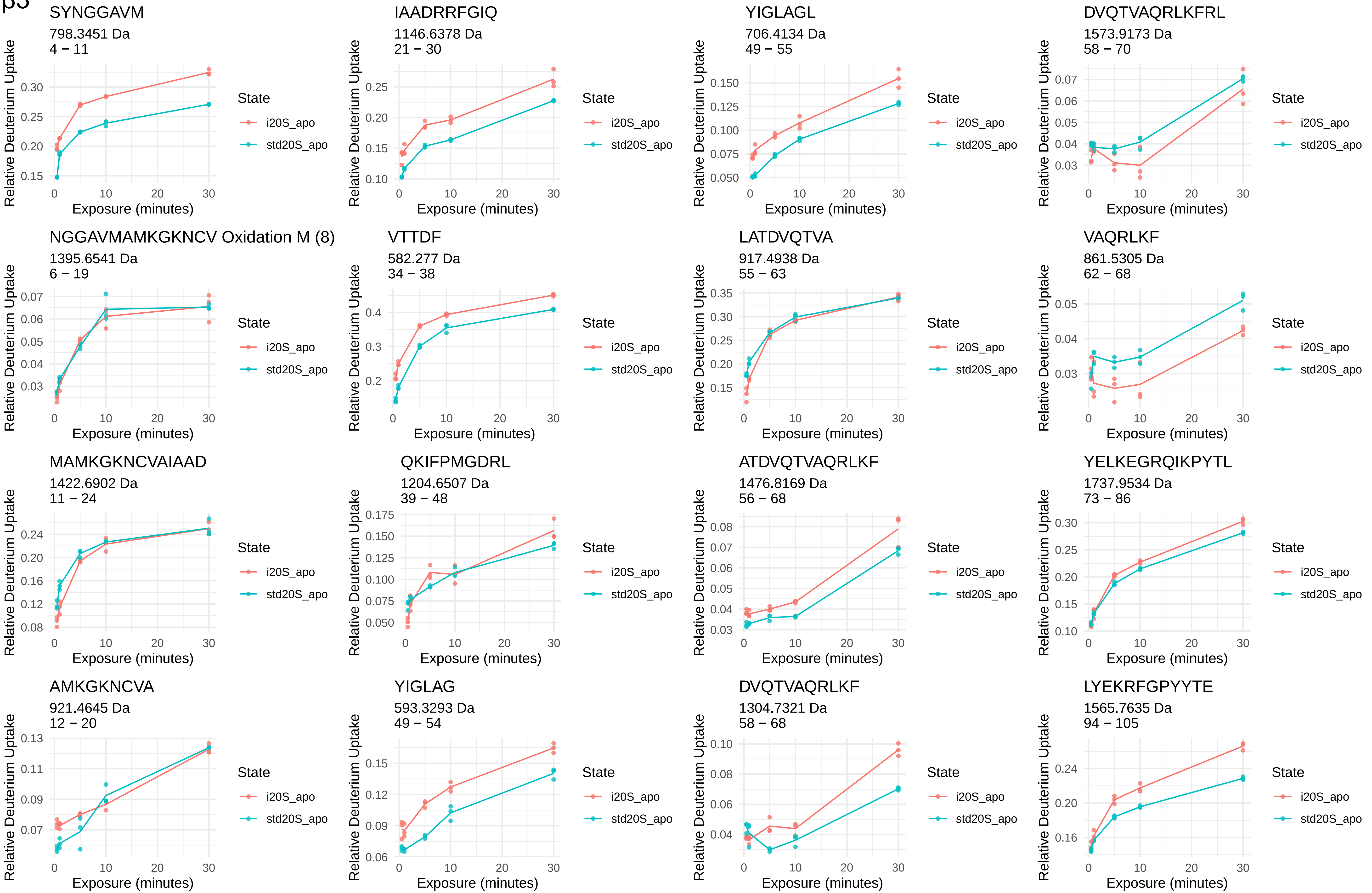
SYNGGAVM
798.3451 Da
4 − 11
IAADRRFGIQ
1146.6378 Da
21 − 30
YIGLAGL
706.4134 Da
49 − 55
DVQTVAQRLKFRL
1573.9173 Da
58 − 70
NGGAVMAMKGKNCV Oxidation M (8)
1395.6541 Da
6 − 19
VTTDF
582.277 Da
34 − 38
LATDVQTVA
917.4938 Da
55 − 63
VAQRLKF
861.5305 Da
62 − 68
MAMKGKNCVAIAAD
1422.6902 Da
11 − 24
QKIFPMGDRL
1204.6507 Da
39 − 48
ATDVQTVAQRLKF
1476.8169 Da
56 − 68
YELKEGRQIKPYTL
1737.9534 Da
73 − 86
AMKGKNCVA
921.4645 Da
12 − 20
YIGLAG
593.3293 Da
49 − 54
DVQTVAQRLKF
1304.7321 Da
58 − 68
LYEKRFGPYYTE
1565.7635 Da
94 − 105
Relative Deuterium Uptake
Exposure (minutes)
State
i20S_apo
std20S_apo

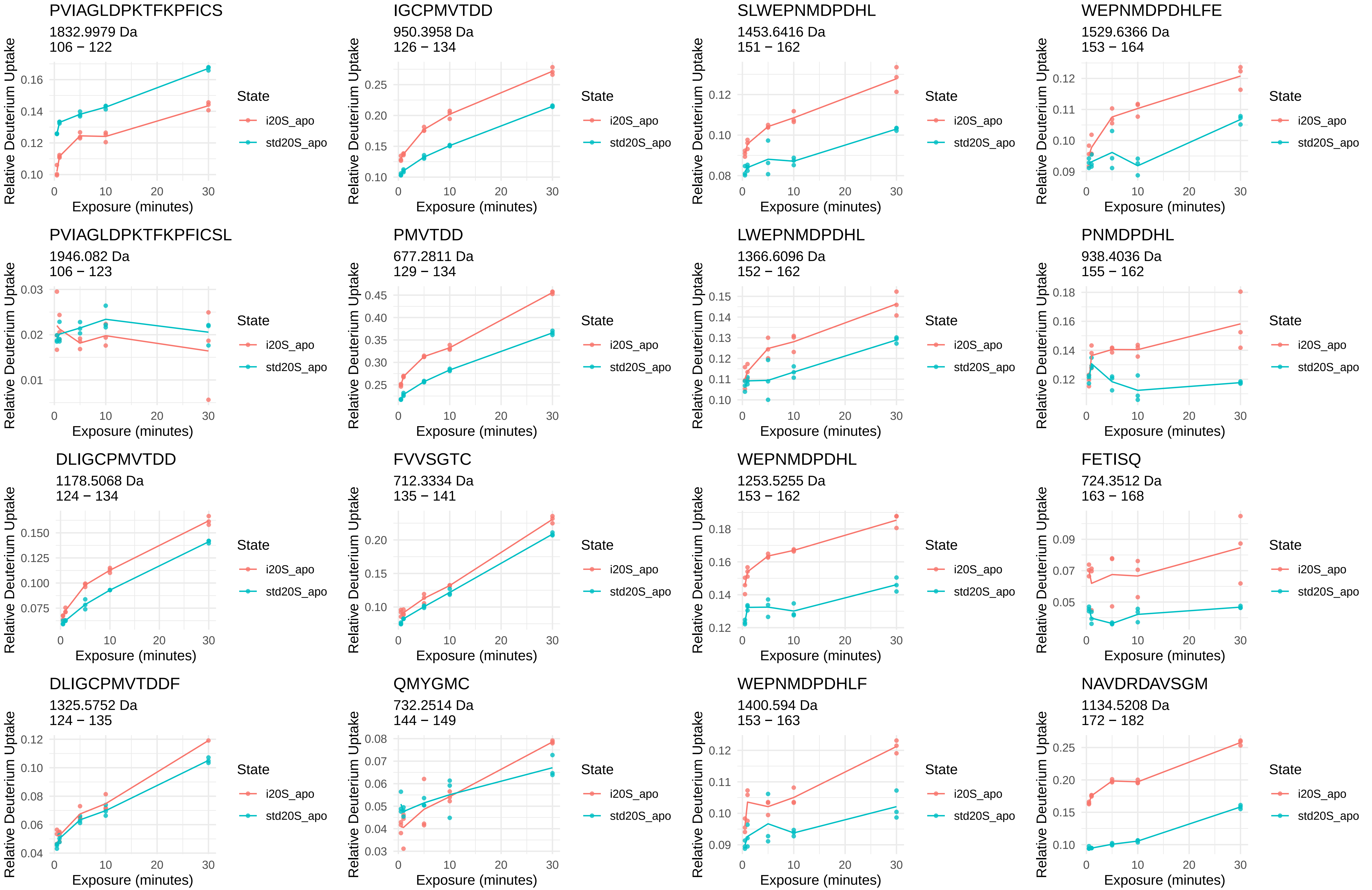
PVIAGLDPKTFKPFICS
1832.9979 Da
106 − 122
IGCPMVTDD
950.3958 Da
126 − 134
SLWEPNMDPDHL
1453.6416 Da
151 − 162
WEPNMDPDHLFE
1529.6366 Da
153 − 164
PVIAGLDPKTFKPFICSL
1946.082 Da
106 − 123
PMVTDD
677.2811 Da
129 − 134
LWEPNMDPDHL
1366.6096 Da
152 − 162
PNMDPDHL
938.4036 Da
155 − 162
DLIGCPMVTDD
1178.5068 Da
124 − 134
FVVSGTC
712.3334 Da
135 − 141
WEPNMDPDHL
1253.5255 Da
153 − 162
FETISQ
724.3512 Da
163 − 168
DLIGCPMVTDDF
1325.5752 Da
124 − 135
QMYGMC
732.2514 Da
144 − 149
WEPNMDPDHLF
1400.594 Da
153 − 163
NAVDRDAVSGM
1134.5208 Da
172 − 182
Relative Deuterium Uptake
Exposure (minutes)
State
i20S_apo
std20S_apo

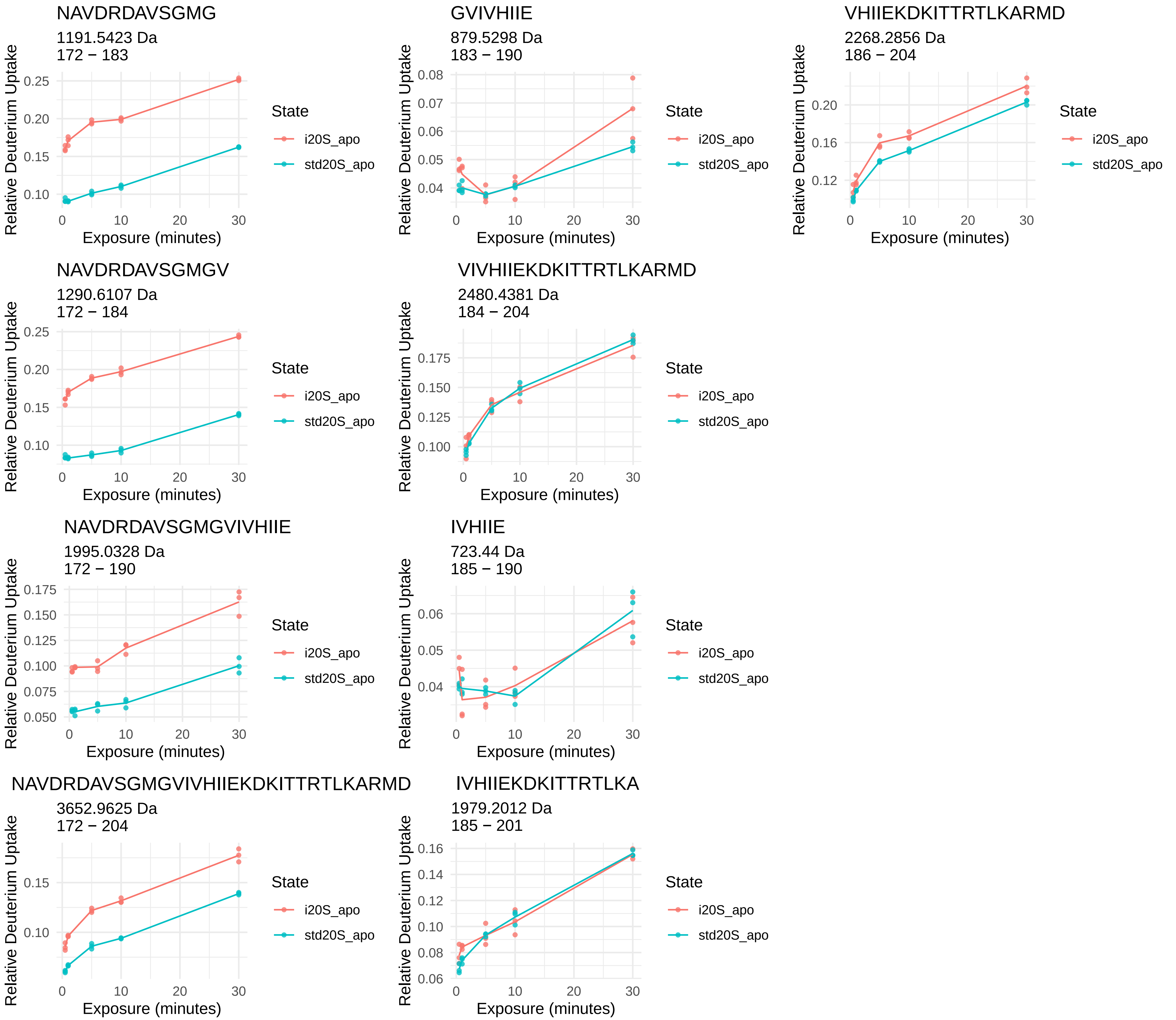
NAVDRDAVSGMG
1191.5423 Da
172 − 183
GVIVHIIE
879.5298 Da
183 − 190
VHIIEKDKITTRTLKARMD
2268.2856 Da
186 − 204
NAVDRDAVSGMGV
1290.6107 Da
172 − 184
VIVHIIEKDKITTRTLKARMD
2480.4381 Da
184 − 204
NAVDRDAVSGMGVIVHIIE
1995.0328 Da
172 − 190
IVHIIE
723.44 Da
185 − 190
NAVDRDAVSGMGVIVHIIEKDKITTRTLKARMD
3652.9625 Da
172 − 204
IVHIIEKDKITTRTLKA
1979.2012 Da
185 − 201
Relative Deuterium Uptake
Exposure (minutes)
State
i20S_apo
std20S_apo

β4

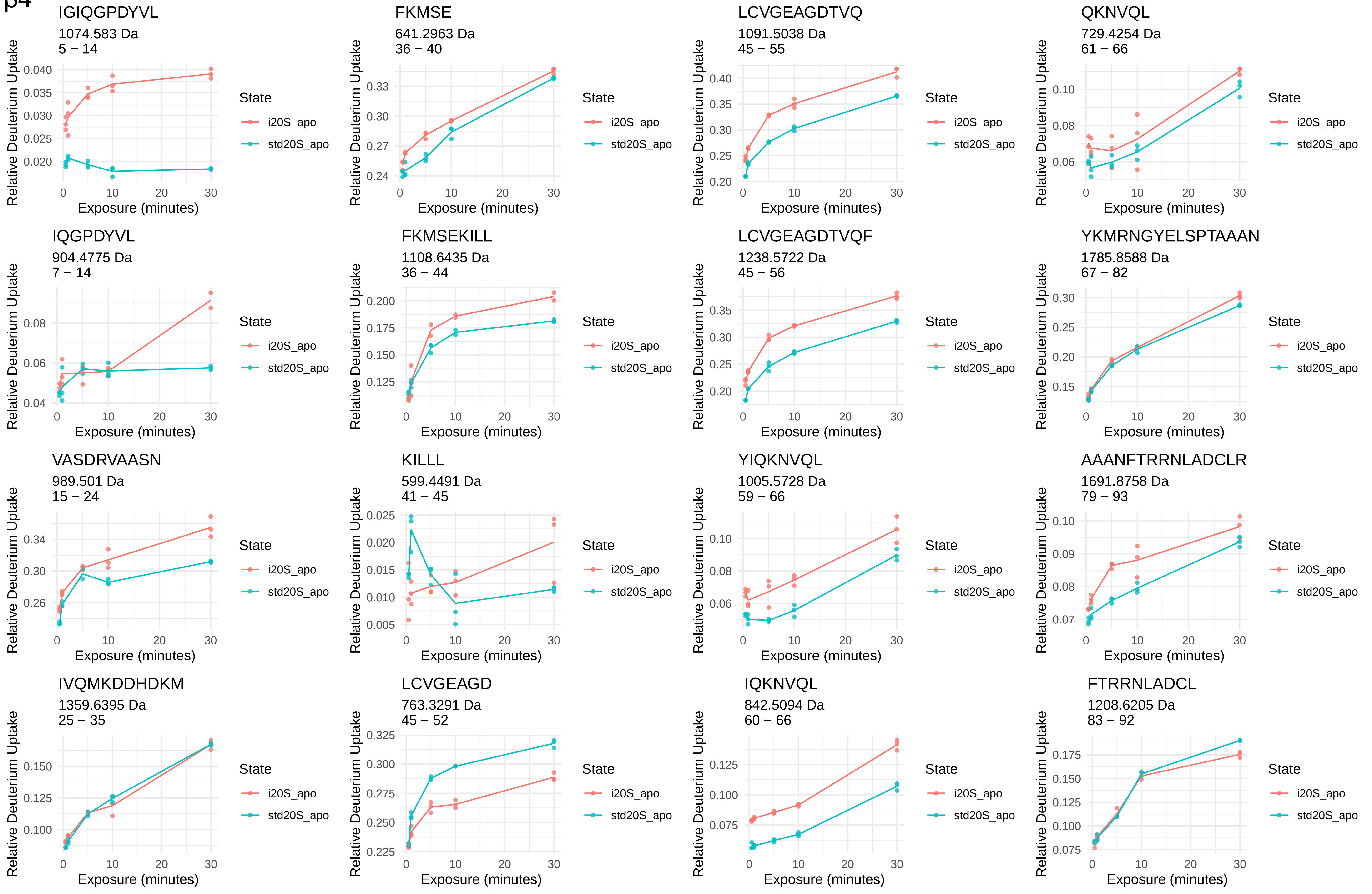
IGIQGPDYVL
1074.583 Da
5 − 14
FKMSE
641.2963 Da
36 − 40
LCVGEAGDTVQ
1091.5038 Da
45 − 55
QKNVQL
729.4254 Da
61 − 66
IQGPDYVL
904.4775 Da
7 − 14
FKMSEKILL
1108.6435 Da
36 − 44
LCVGEAGDTVQF
1238.5722 Da
45 − 56
YKMRNGYELSPTAAAN
1785.8588 Da
67 − 82
VASDRVAASN
989.501 Da
15 − 24
KILLL
599.4491 Da
41 − 45
YIQKNVQL
1005.5728 Da
59 − 66
AAANFTRRNLADCLR
1691.8758 Da
79 − 93
IVQMKDDHDKM
1359.6395 Da
25 − 35
LCVGEAGD
763.3291 Da
45 − 52
IQKNVQL
842.5094 Da
60 − 66
FTRRNLADCL
1208.6205 Da
83 − 92
Relative Deuterium Uptake
Exposure (minutes)
State
i20S_apo
std20S_apo

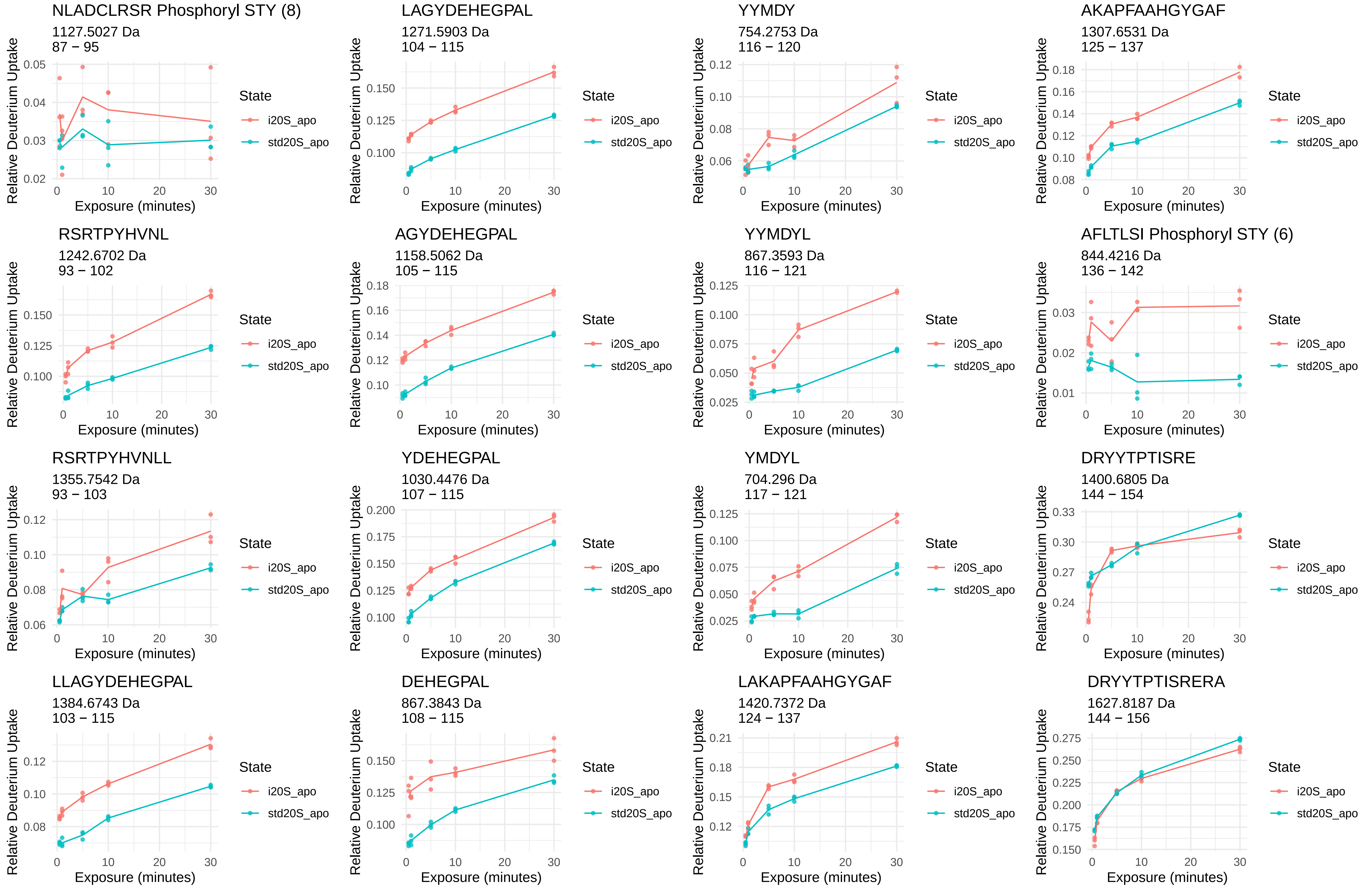
NLADCLRSR Phosphoryl STY (8)
1127.5027 Da
87 − 95
LAGYDEHEGPAL
1271.5903 Da
104 − 115
YYMDY
754.2753 Da
116 − 120
AKAPFAAHGYGAF
1307.6531 Da
125 − 137
RSRTPYHVNL
1242.6702 Da
93 − 102
AGYDEHEGPAL
1158.5062 Da
105 − 115
YYMDYL
867.3593 Da
116 − 121
AFLTLSI Phosphoryl STY (6)
844.4216 Da
136 − 142
RSRTPYHVNLL
1355.7542 Da
93 − 103
YDEHEGPAL
1030.4476 Da
107 − 115
YMDYL
704.296 Da
117 − 121
DRYYTPTISRE
1400.6805 Da
144 − 154
LLAGYDEHEGPAL
1384.6743 Da
103 − 115
DEHEGPAL
867.3843 Da
108 − 115
LAKAPFAAHGYGAF
1420.7372 Da
124 − 137
DRYYTPTISRERA
1627.8187 Da
144 − 156
Relative Deuterium Uptake
Exposure (minutes)
State
i20S_apo
std20S_apo

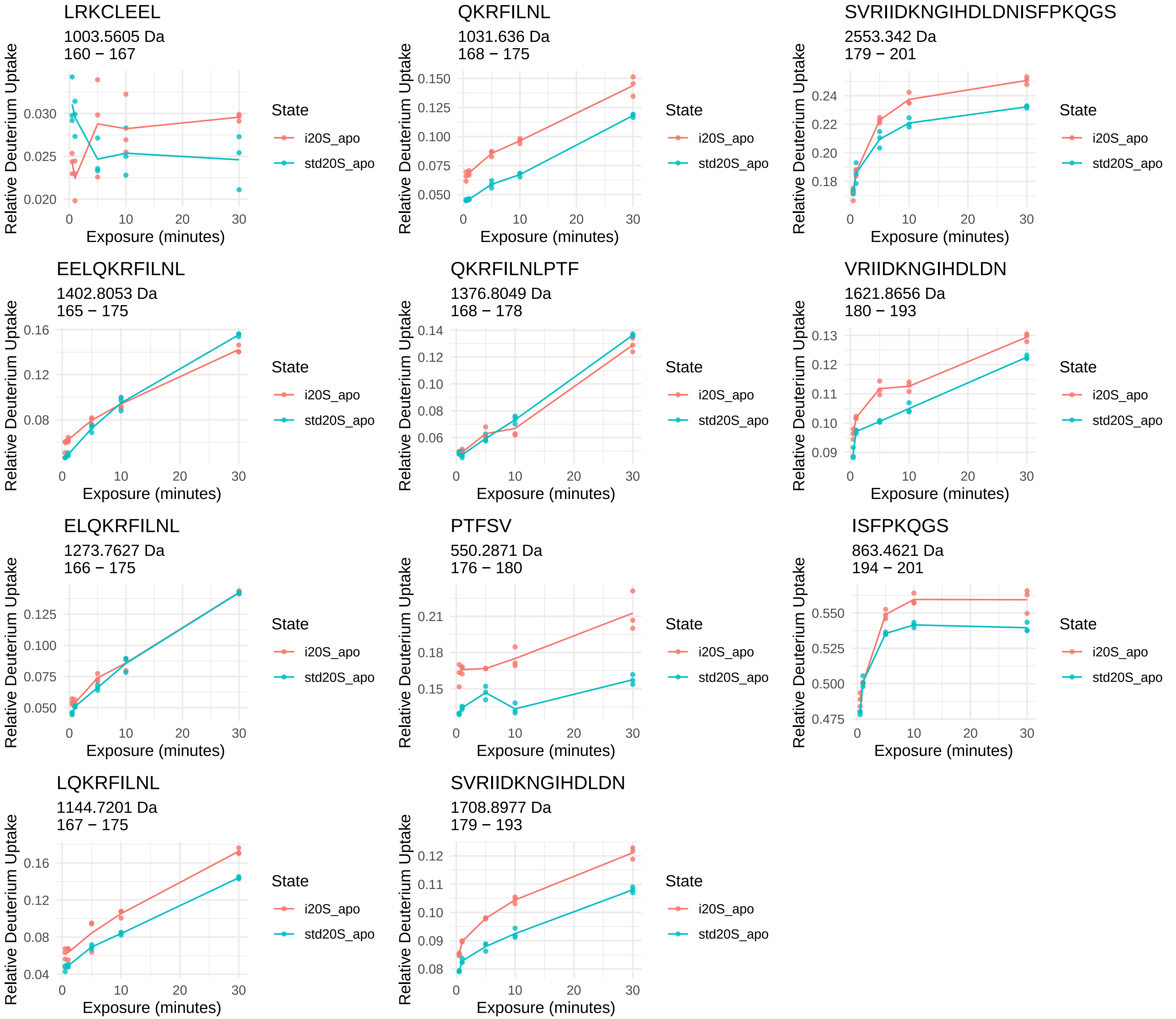
LRKCLEEL
1003.5605 Da
160 − 167
QKRFILNL
1031.636 Da
168 − 175
SVRIIDKNGIHDLDNISFPKQGS
2553.342 Da
179 − 201
EELQKRFILNL
1402.8053 Da
165 − 175
QKRFILNLPTF
1376.8049 Da
168 − 178
VRIIDKNGIHDLDN
1621.8656 Da
180 − 193
ELQKRFILNL
1273.7627 Da
166 − 175
PTFSV
550.2871 Da
176 − 180
ISFPKQGS
863.4621 Da
194 − 201
LQKRFILNL
1144.7201 Da
167 − 175
SVRIIDKNGIHDLDN
1708.8977 Da
179 − 193
Relative Deuterium Uptake
Exposure (minutes)
State
i20S_apo
std20S_apo

β5

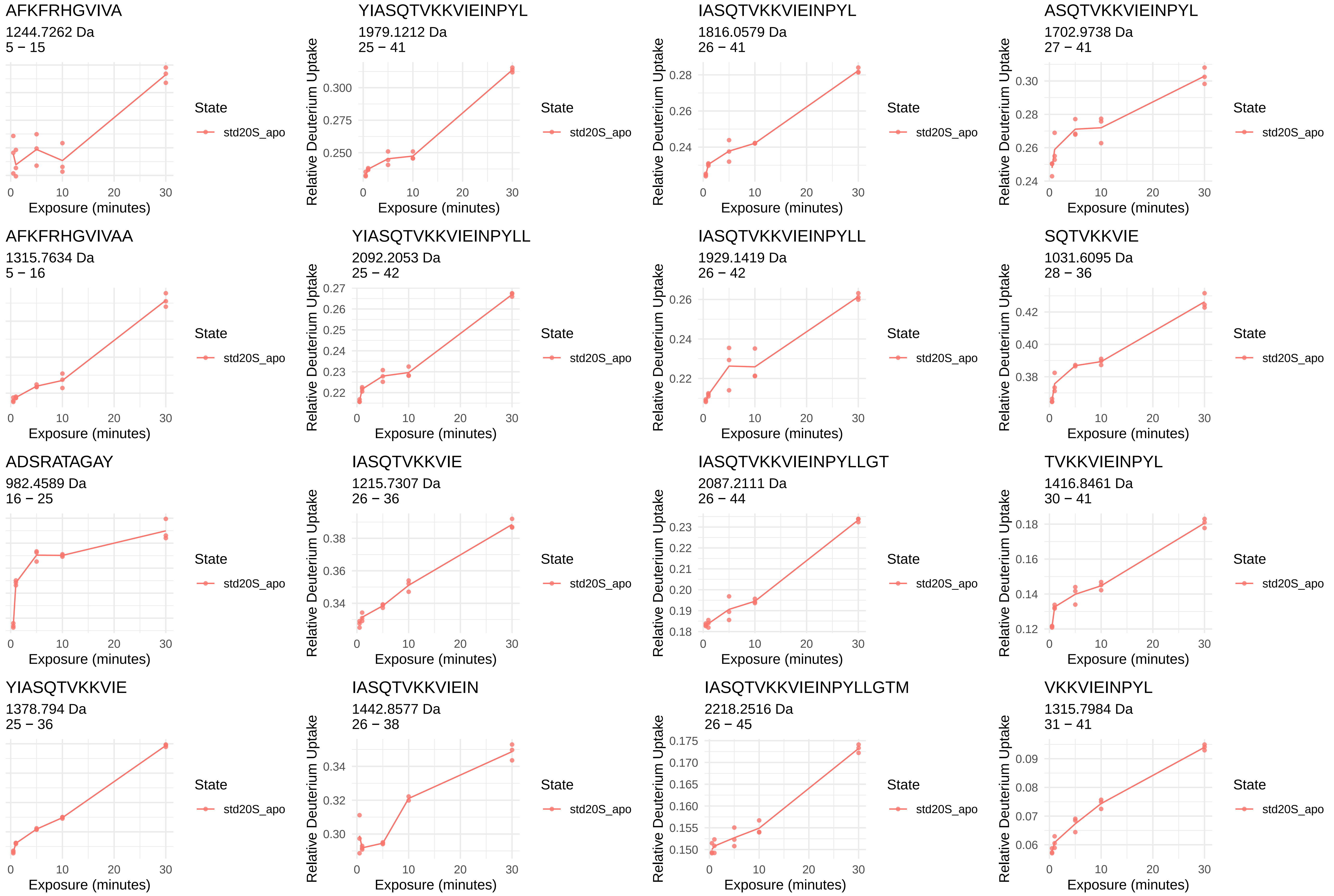
AFKFRHGVIVA
1244.7262 Da
5 − 15
YIASQTVKKVIEINPYL
1979.1212 Da
25 − 41
IASQTVKKVIEINPYL
1816.0579 Da
26 − 41
ASQTVKKVIEINPYL
1702.9738 Da
27 − 41
AFKFRHGVIVAA
1315.7634 Da
5 − 16
YIASQTVKKVIEINPYLL
2092.2053 Da
25 − 42
IASQTVKKVIEINPYLL
1929.1419 Da
26 − 42
SQTVKKVIE
1031.6095 Da
28 − 36
ADSRATAGAY
982.4589 Da
16 − 25
IASQTVKKVIE
1215.7307 Da
26 − 36
IASQTVKKVIEINPYLLGT
2087.2111 Da
26 − 44
TVKKVIEINPYL
1416.8461 Da
30 − 41
YIASQTVKKVIE
1378.794 Da
25 − 36
IASQTVKKVIEIN
1442.8577 Da
26 − 38
IASQTVKKVIEINPYLLGTM
2218.2516 Da
26 − 45
VKKVIEINPYL
1315.7984 Da
31 − 41
Relative Deuterium Uptake
Exposure (minutes)
State
std20S_apo

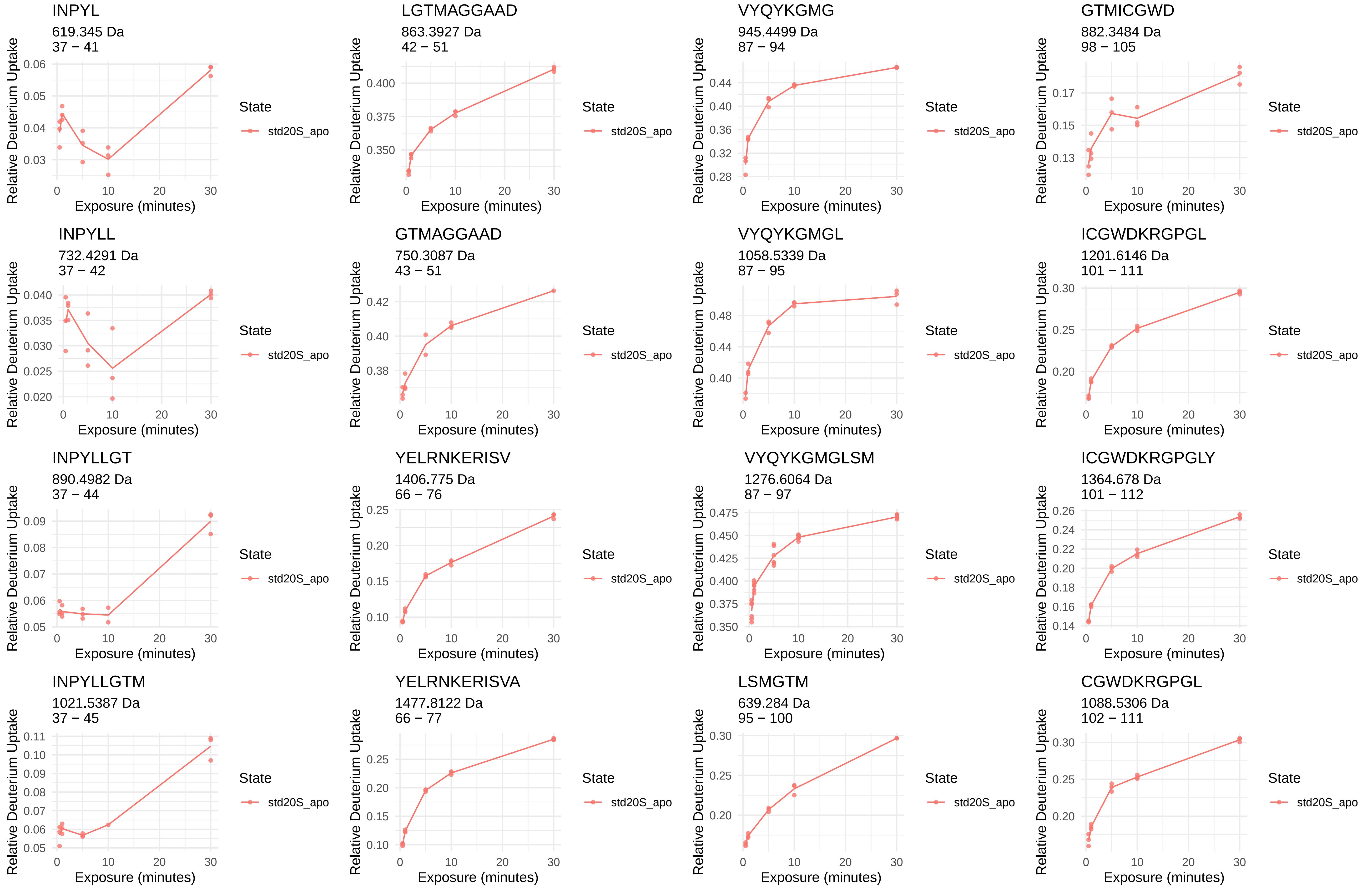
INPYL
619.345 Da
37 − 41
LGTMAGGAAD
863.3927 Da
42 − 51
VYQYKGMG
945.4499 Da
87 − 94
GTMICGWD
882.3484 Da
98 − 105
INPYLL
732.4291 Da
37 − 42
GTMAGGAAD
750.3087 Da
43 − 51
VYQYKGMGL
1058.5339 Da
87 − 95
ICGWDKRGPGL
1201.6146 Da
101 − 111
INPYLLGT
890.4982 Da
37 − 44
YELRNKERISV
1406.775 Da
66 − 76
VYQYKGMGLSM
1276.6064 Da
87 − 97
ICGWDKRGPGLY
1364.678 Da
101 − 112
INPYLLGTM
1021.5387 Da
37 − 45
YELRNKERISVA
1477.8122 Da
66 − 77
LSMGTM
639.284 Da
95 − 100
CGWDKRGPGL
1088.5306 Da
102 − 111
Relative Deuterium Uptake
Exposure (minutes)
State
std20S_apo

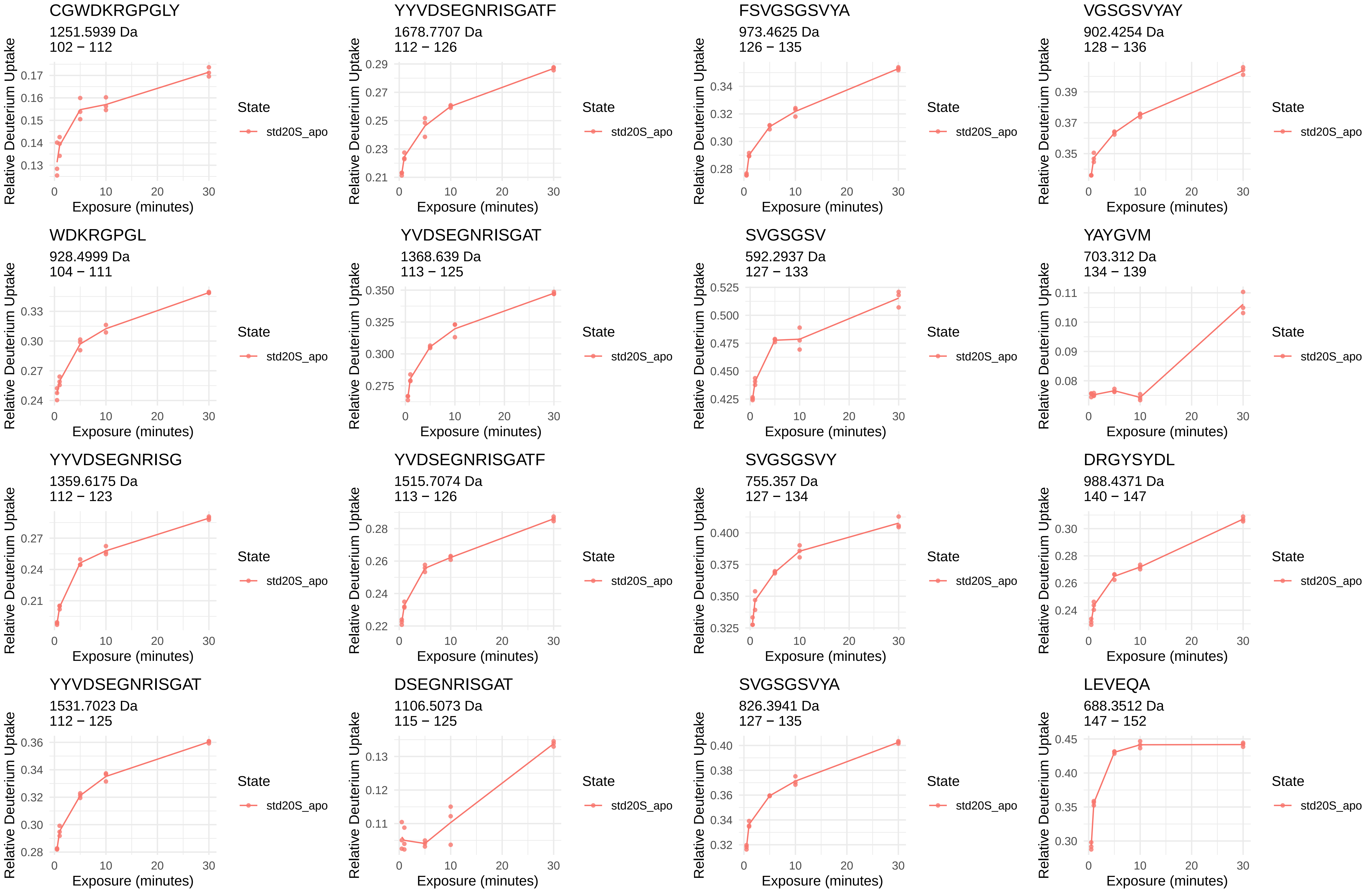
CGWDKRGPGLY
1251.5939 Da
102 − 112
YYVDSEGNRISGATF
1678.7707 Da
112 − 126
FSVGSGSVYA
973.4625 Da
126 − 135
VGSGSVYAY
902.4254 Da
128 − 136
WDKRGPGL
928.4999 Da
104 − 111
YVDSEGNRISGAT
1368.639 Da
113 − 125
SVGSGSV
592.2937 Da
127 − 133
YAYGVM
703.312 Da
134 − 139
YYVDSEGNRISG
1359.6175 Da
112 − 123
YVDSEGNRISGATF
1515.7074 Da
113 − 126
SVGSGSVY
755.357 Da
127 − 134
DRGYSYDL
988.4371 Da
140 − 147
YYVDSEGNRISGAT
1531.7023 Da
112 − 125
DSEGNRISGAT
1106.5073 Da
115 − 125
SVGSGSVYA
826.3941 Da
127 − 135
LEVEQA
688.3512 Da
147 − 152
Relative Deuterium Uptake
Exposure (minutes)
State
std20S_apo

ARRAIYQAT
1049.5851 Da
156 − 164

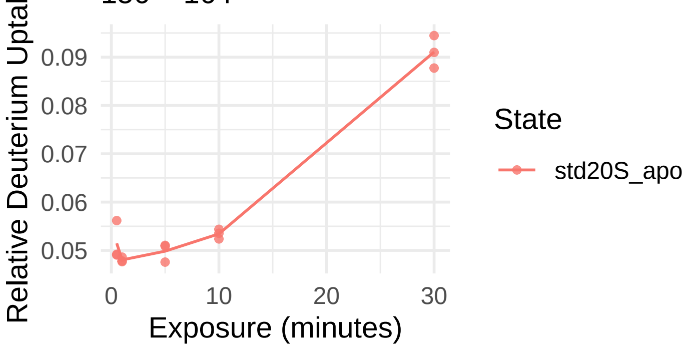


NVADL
531.2773 Da
191 − 195

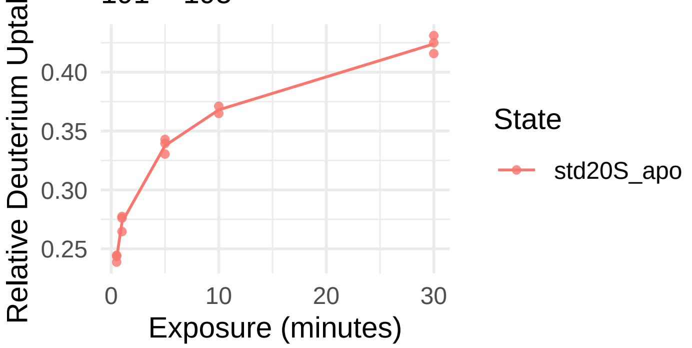


YRDAYSGGAVNL
1285.6171 Da
165 − 176

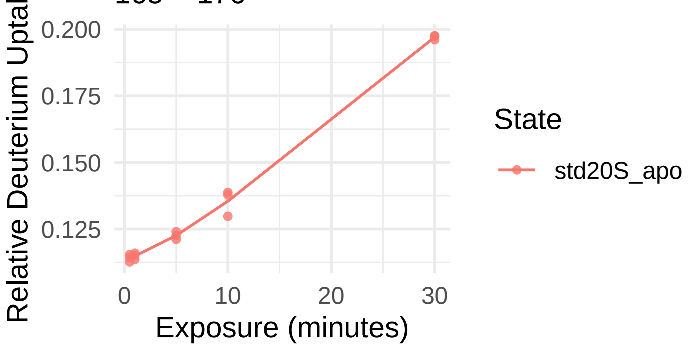


HEKYSGSTP
1005.4636 Da
196 − 204

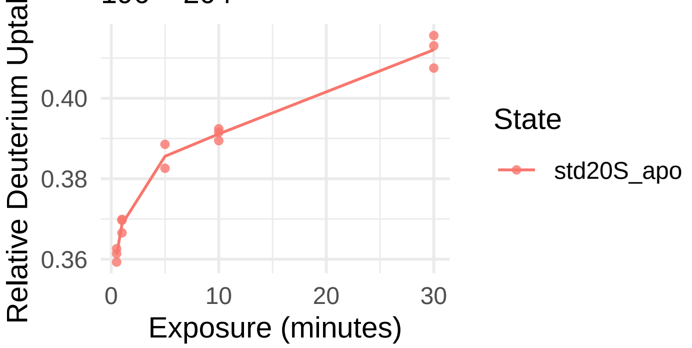


YHVREDGW
1061.4799 Da
177 − 184

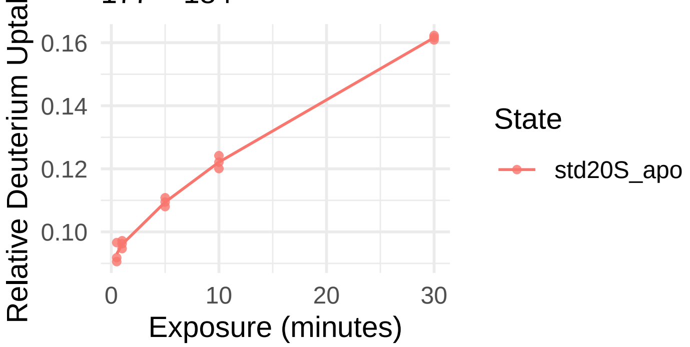


WIRVSSDNVADL
1374.7012 Da
184 − 195

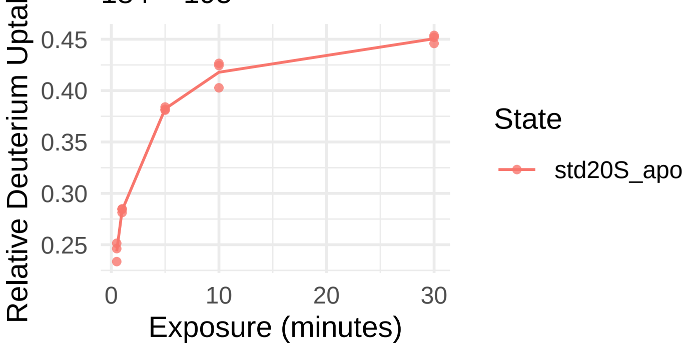

β5i

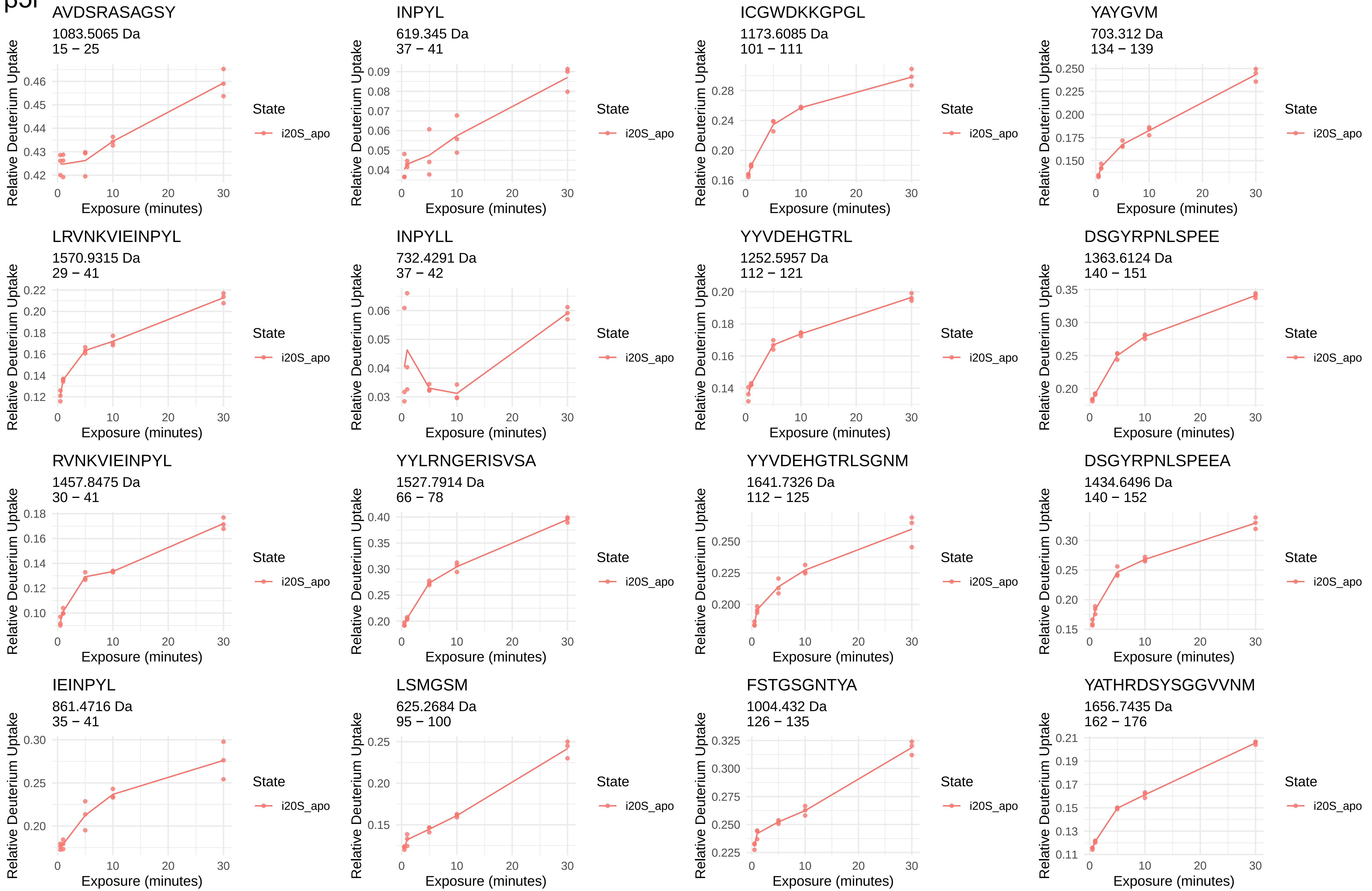
AVDSRASAGSY
1083.5065 Da
15 − 25
INPYL
619.345 Da
37 − 41
ICGWDKKGPGL
1173.6085 Da
101 − 111
YAYGVM
703.312 Da
134 − 139
LRVNKVIEINPYL
1570.9315 Da
29 − 41
INPYLL
732.4291 Da
37 − 42
YYVDEHGTRL
1252.5957 Da
112 − 121
DSGYRPNLSPEE
1363.6124 Da
140 − 151
RVNKVIEINPYL
1457.8475 Da
30 − 41
YYLRNGERISVSA
1527.7914 Da
66 − 78
YYVDEHGTRLSGNM
1641.7326 Da
112 − 125
DSGYRPNLSPEEA
1434.6496 Da
140 − 152
IEINPYL
861.4716 Da
35 − 41
LSMGSM
625.2684 Da
95 − 100
FSTGSGNTYA
1004.432 Da
126 − 135
YATHRDSYSGGVVNM
1656.7435 Da
162 − 176
Relative Deuterium Uptake
Exposure (minutes)
State
i20S_apo

ATHRDSYSGGVVNM

1493.6802 Da
163 − 176

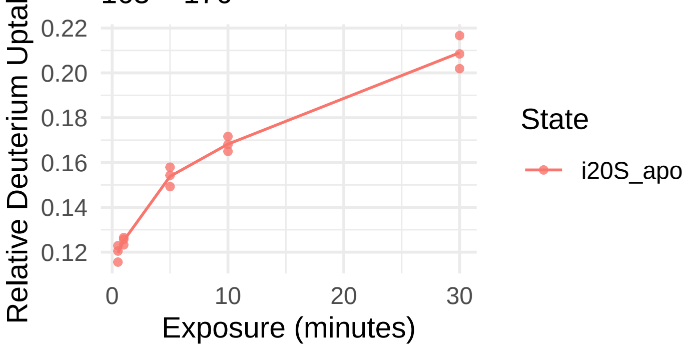


WVKVESTDVSDL

1377.6896 Da
184 − 195

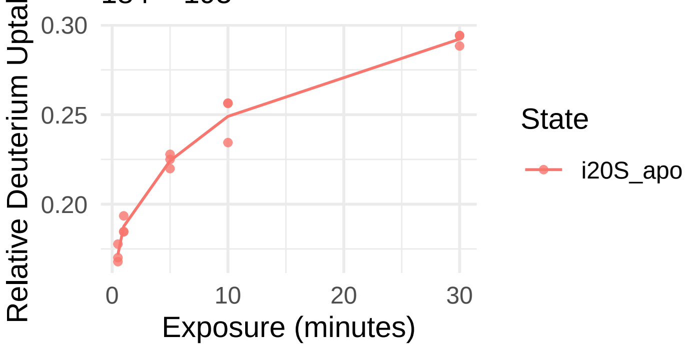


VKVESTDVSDL

1191.6103 Da
185 − 195

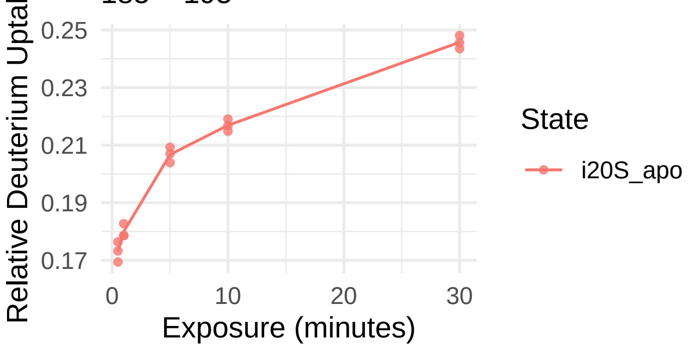

β6

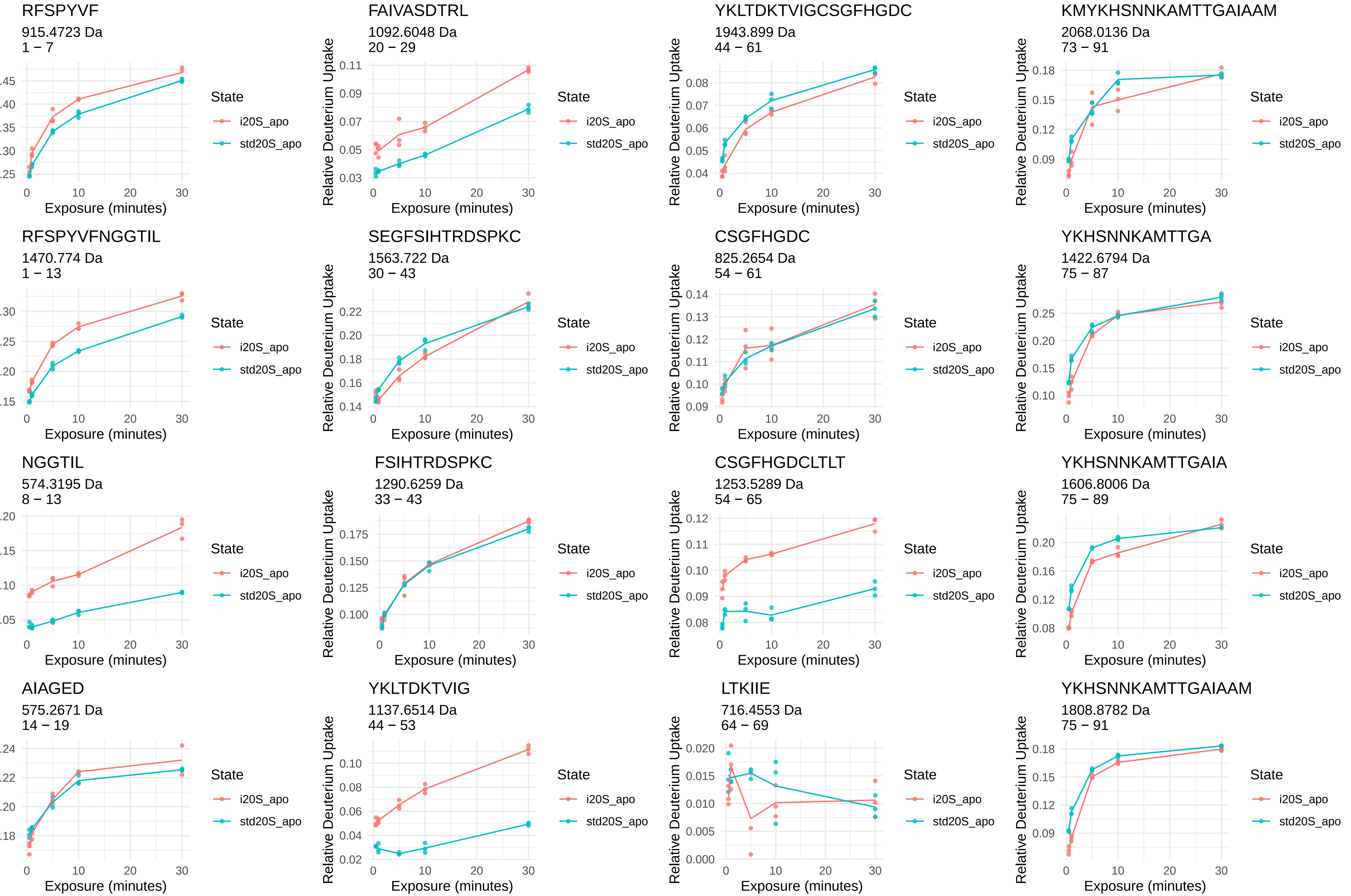
RFSPYVF
915.4723 Da
1 − 7
FAIVASDTRL
1092.6048 Da
20 − 29
YKLTDKTVIGCSGFHGDC
1943.899 Da
44 − 61
KMYKHSNNKAMTTGAIAAM
2068.0136 Da
73 − 91
RFSPYVFNGGTIL
1470.774 Da
1 − 13
SEGFSIHTRDSPKC
1563.722 Da
30 − 43
CSGFHGDC
825.2654 Da
54 − 61
YKHSNNKAMTTGA
1422.6794 Da
75 − 87
NGGTIL
574.3195 Da
8 − 13
FSIHTRDSPKC
1290.6259 Da
33 − 43
CSGFHGDCLTLT
1253.5289 Da
54 − 65
YKHSNNKAMTTGAIA
1606.8006 Da
75 − 89
AIAGED
575.2671 Da
14 − 19
YKLTDKTVIG
1137.6514 Da
44 − 53
LTKIIE
716.4553 Da
64 − 69
YKHSNNKAMTTGAIAAM
1808.8782 Da
75 − 91
Relative Deuterium Uptake
Exposure (minutes)
State
i20S_apo
std20S_apo

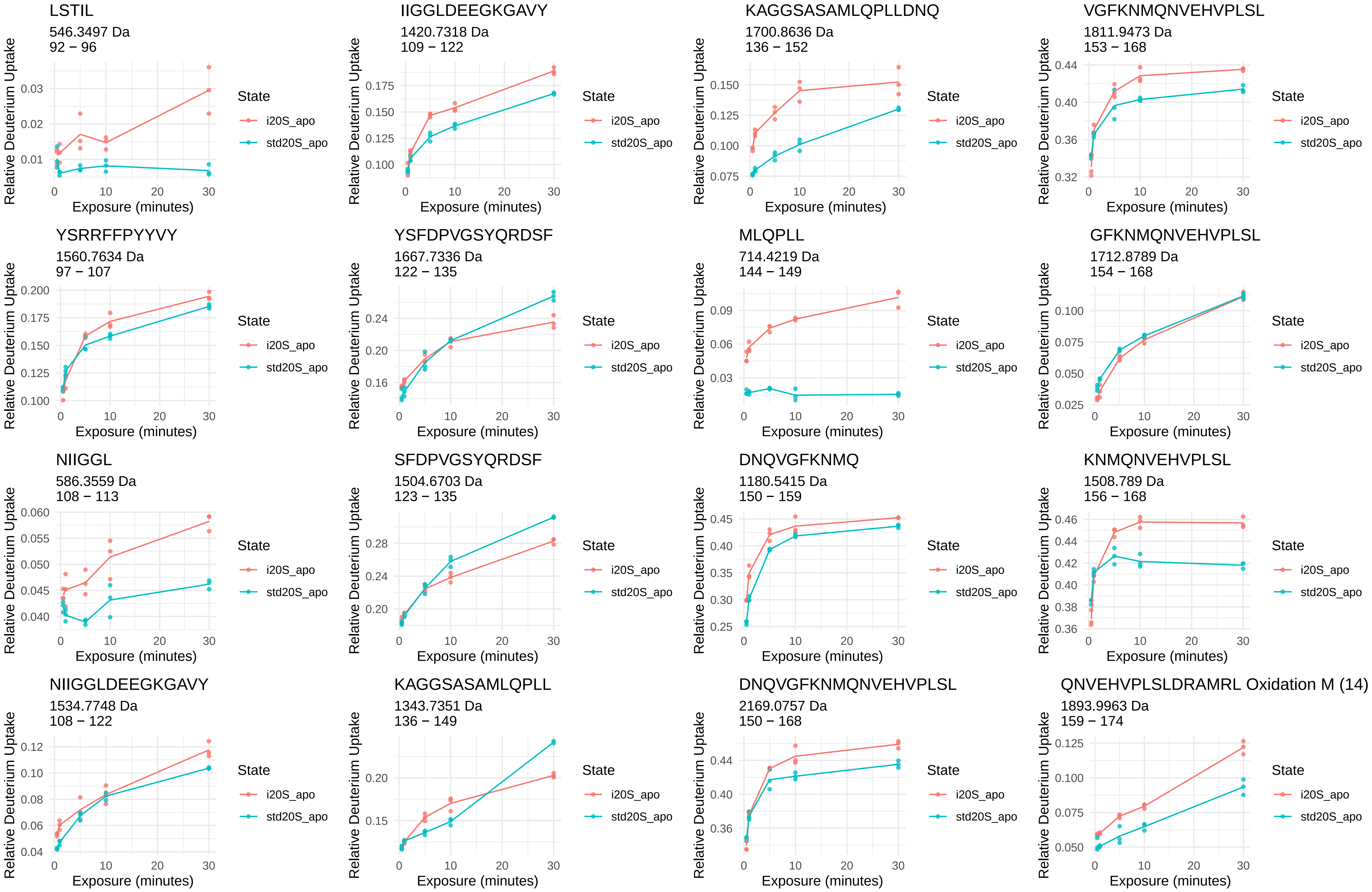
LSTIL
546.3497 Da
92 − 96
IIGGLDEEGKGAVY
1420.7318 Da
109 − 122
KAGGSASAMLQPLLDNQ
1700.8636 Da
136 − 152
VGFKNMQNVEHVPLSL
1811.9473 Da
153 − 168
YSRRFFPYYVY
1560.7634 Da
97 − 107
YSFDPVGSYQRDSF
1667.7336 Da
122 − 135
MLQPLL
714.4219 Da
144 − 149
GFKNMQNVEHVPLSL
1712.8789 Da
154 − 168
NIIGGL
586.3559 Da
108 − 113
SFDPVGSYQRDSF
1504.6703 Da
123 − 135
DNQVGFKNMQ
1180.5415 Da
150 − 159
KNMQNVEHVPLSL
1508.789 Da
156 − 168
NIIGGLDEEGKGAVY
1534.7748 Da
108 − 122
KAGGSASAMLQPLL
1343.7351 Da
136 − 149
DNQVGFKNMQNVEHVPLSL
2169.0757 Da
150 − 168
QNVEHVPLSLDRAMRL Oxidation M (14)
1893.9963 Da
159 − 174
Relative Deuterium Uptake
Exposure (minutes)
State
i20S_apo
std20S_apo

NVEHVPLSL
1007.552 Da
160 − 168

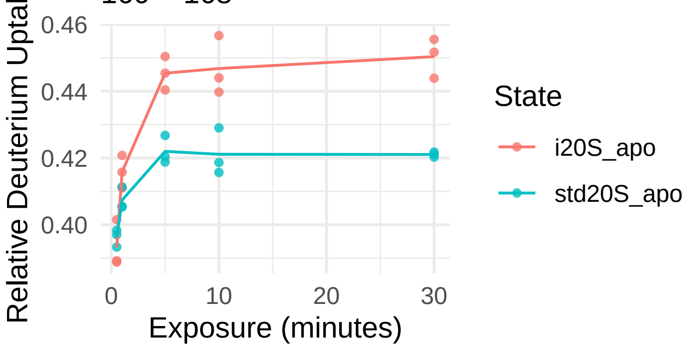


AERDVYTGDAL
1209.5746 Da
183 − 193

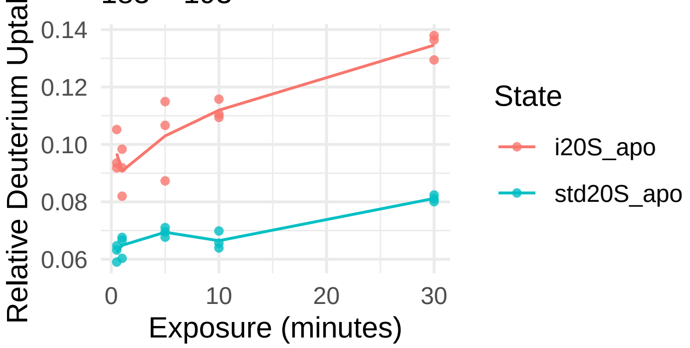


VYTGDA
625.2828 Da
187 − 192

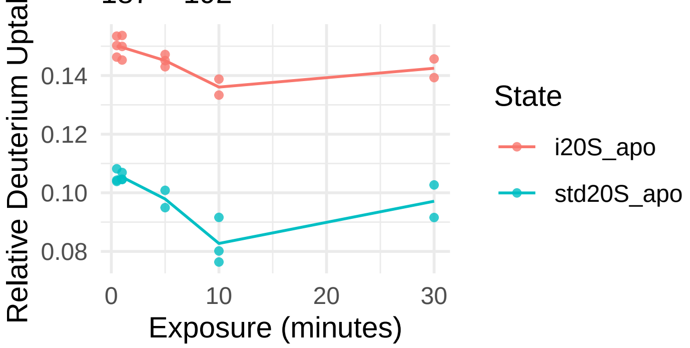


RICIVTKEGIRE
1416.7991 Da
194 − 205

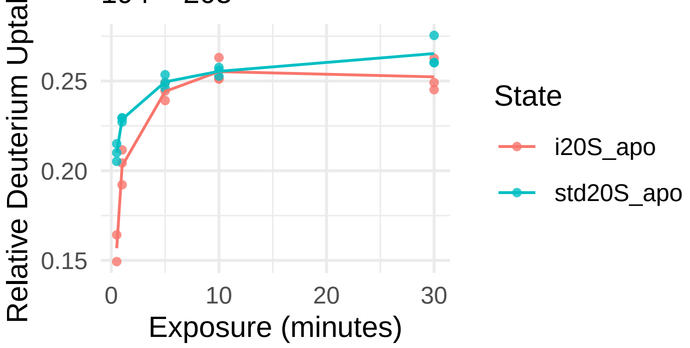

β7

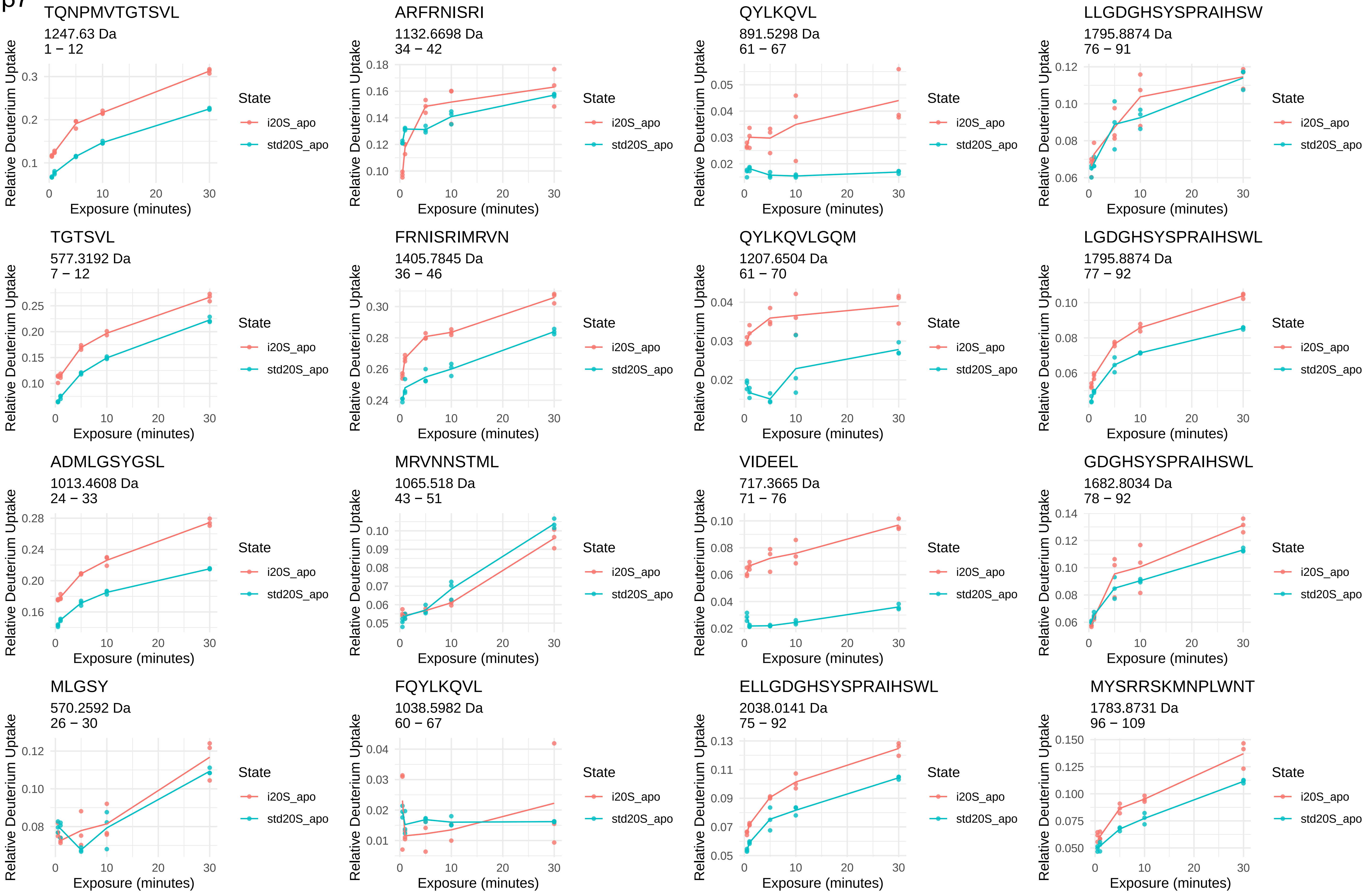
TQNPMVTGTSVL
1247.63 Da
1 − 12
ARFRNISRI
1132.6698 Da
34 − 42
QYLKQVL
891.5298 Da
61 − 67
LLGDGHSYSPRAIHSW
1795.8874 Da
76 − 91
TGTSVL
577.3192 Da
7 − 12
FRNISRIMRVN
1405.7845 Da
36 − 46
QYLKQVLGQM
1207.6504 Da
61 − 70
LGDGHSYSPRAIHSWL
1795.8874 Da
77 − 92
ADMLGSYGSL
1013.4608 Da
24 − 33
MRVNNSTML
1065.518 Da
43 − 51
VIDEEL
717.3665 Da
71 − 76
GDGHSYSPRAIHSWL
1682.8034 Da
78 − 92
MLGSY
570.2592 Da
26 − 30
FQYLKQVL
1038.5982 Da
60 − 67
ELLGDGHSYSPRAIHSWL
2038.0141 Da
75 − 92
MYSRRSKMNPLWNT
1783.8731 Da
96 − 109
Relative Deuterium Uptake
Exposure (minutes)
State
i20S_apo
std20S_apo

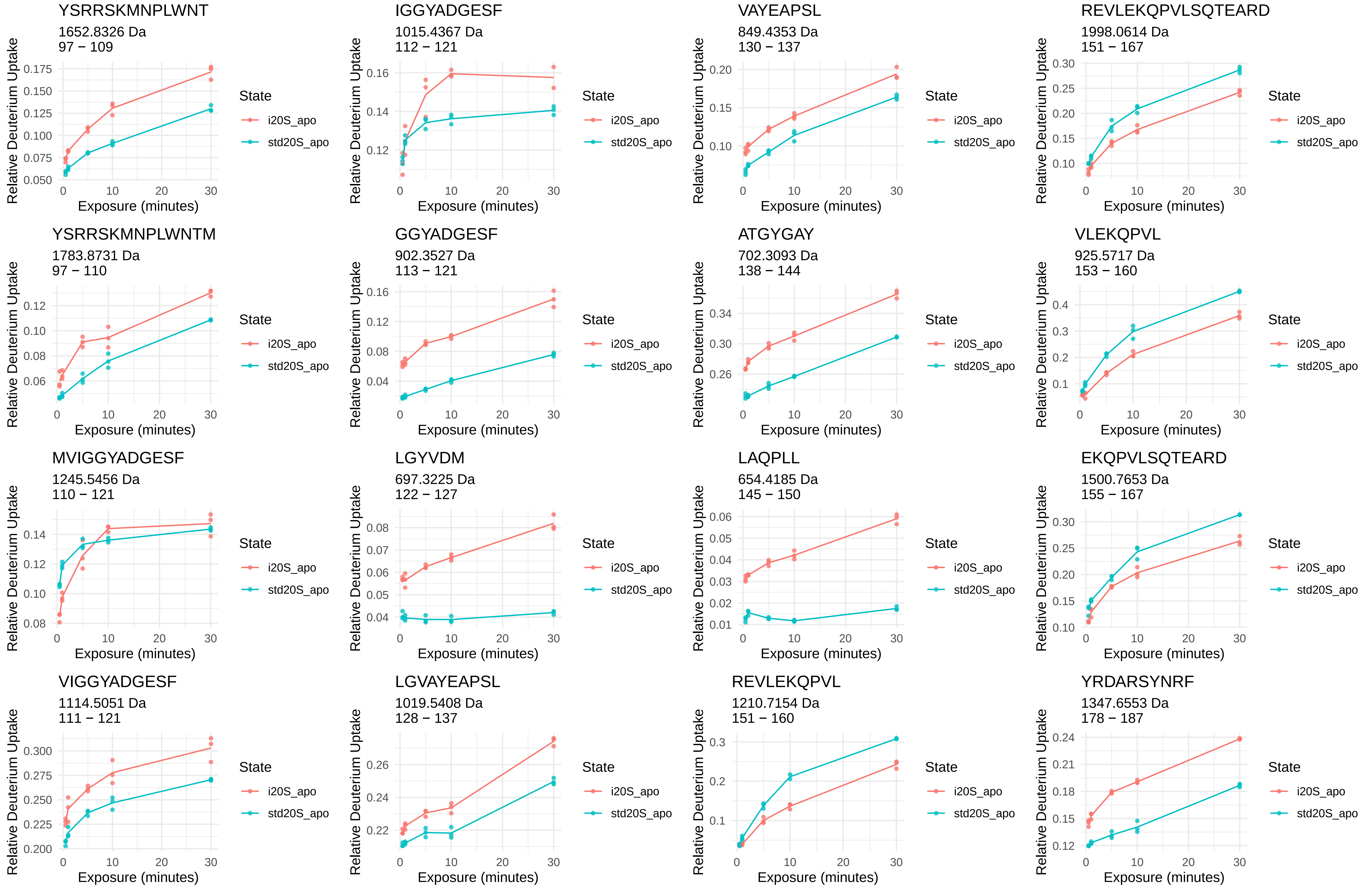
YSRRSKMNPLWNT
1652.8326 Da
97 – 109
IGGYADGESF
1015.4367 Da
112 – 121
VAYEAPSL
849.4353 Da
130 – 137
REVLEKQPVLSQTEARD
1998.0614 Da
151 – 167
YSRRSKMNPLWNTM
1783.8731 Da
97 – 110
GGYADGESF
902.3527 Da
113 – 121
ATGYGAY
702.3093 Da
138 – 144
VLEKQPVL
925.5717 Da
153 – 160
MVIGGYADGESF
1245.5456 Da
110 – 121
LGYVDM
697.3225 Da
122 – 127
LAQPLL
654.4185 Da
145 – 150
EKQPVLSQTEARD
1500.7653 Da
155 – 167
VIGGYADGESF
1114.5051 Da
111 – 121
LGVAYEAPSL
1019.5408 Da
128 – 137
REVLEKQPVL
1210.7154 Da
151 – 160
YRDARSYNRF
1347.6553 Da
178 – 187
Relative Deuterium Uptake
Exposure (minutes)
State
i20S_apo
std20S_apo

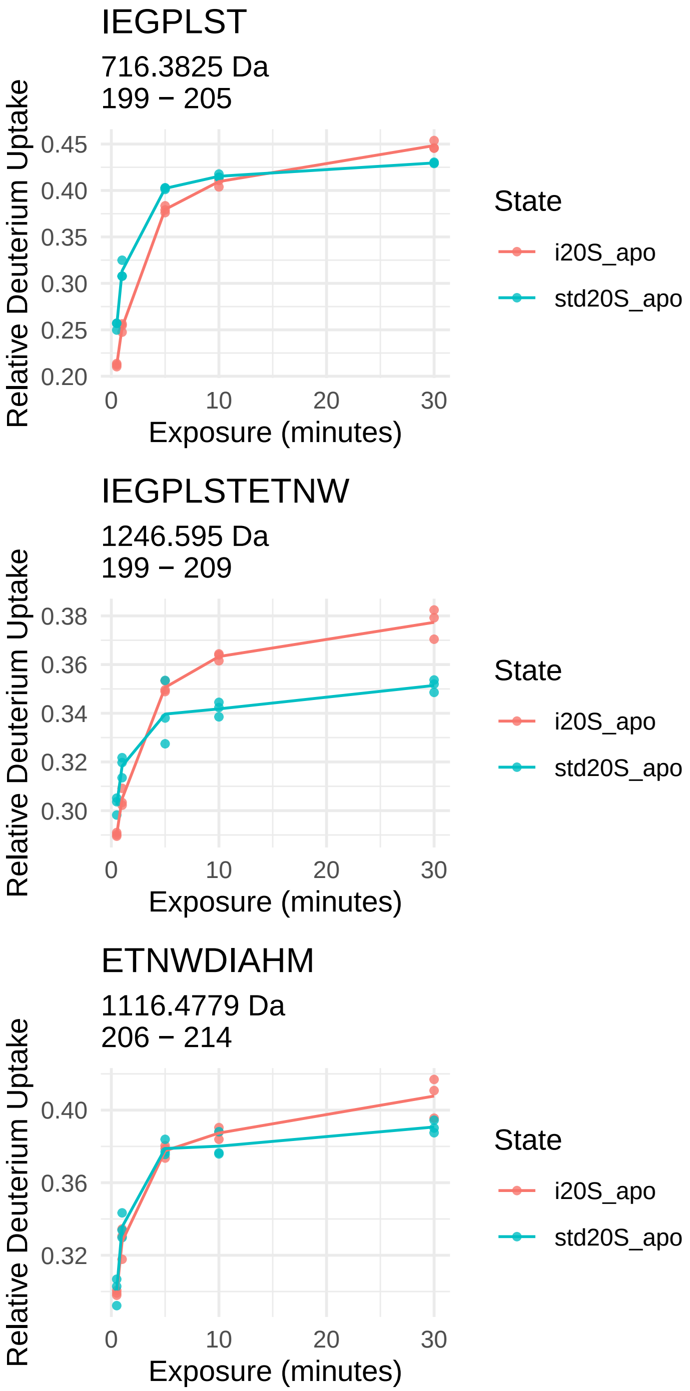
IEGPLST
716.3825 Da
199 − 205
Relative Deuterium Uptake
0.45
0.40
0.35
0.30
0.25
0.20
0
10
20
30
Exposure (minutes)
State
i20S_apo
std20S_apo
IEGPLSTETNW
1246.595 Da
199 − 209
Relative Deuterium Uptake
0.38
0.36
0.34
0.32
0.30
0
10
20
30
Exposure (minutes)
State
i20S_apo
std20S_apo
ETNWDIAHM
1116.4779 Da
206 − 214
Relative Deuterium Uptake
0.40
0.36
0.32
0
10
20
30
Exposure (minutes)
State
i20S_apo
std20S_apo
