## Supplementary material for "Conformational maps of human 20S proteasomes reveal PA28- and immuno-dependent inter-ring crosstalks": Fig. S3

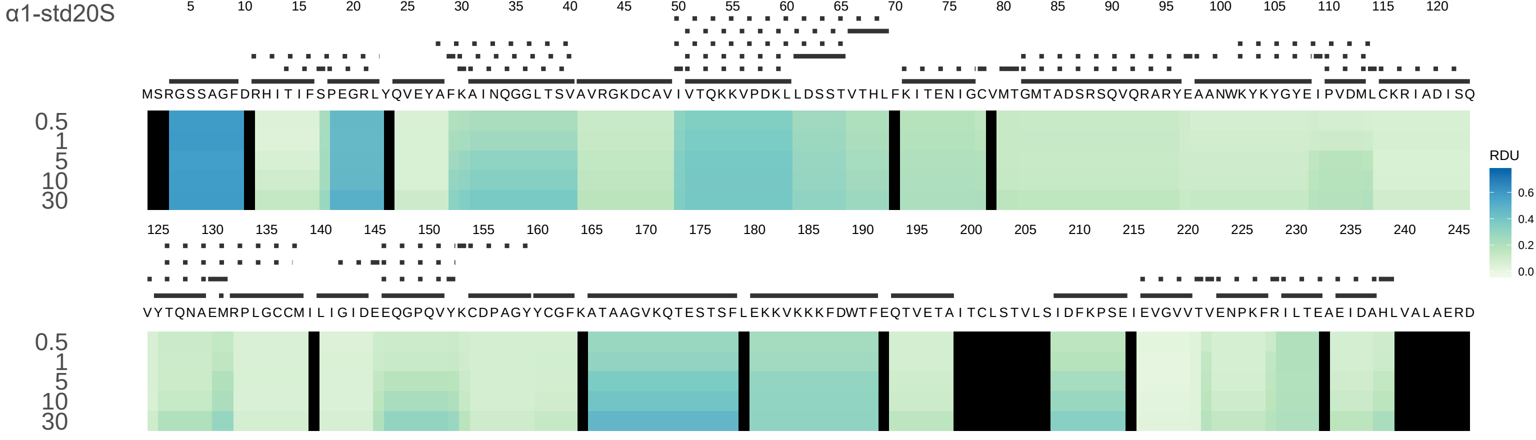
α1-std20S
5 10 15 20 25 30 35 40 45 50 55 60 65 70 75 80 85 90 95 100 105 110 115 120
MSRGSSAGFDRHITIFSPEGRLYQVEYAFKAINQGGLTSVAVRGKDCAVIVTQKKVPDKLLDSSTVTHLFKITENIGCVMTGMTADSRSQVQRARYEAANWKYKYGYEIPVDMLCKRIADISQ
0.5
1
5
10
30
125 130 135 140 145 150 155 160 165 170 175 180 185 190 195 200 205 210 215 220 225 230 235 240 245
VYTQNAEMRPLGCCMILIGIDEEQGPQVYKCDPAGYYCGFKATAAGVKQTESTSFLEKKVKKKFDWTFEQTVETAITCLSTVLSIDFKPSEIEVGVVTVENPKFRILTEAEIDAHLVALAERD
0.5
1
5
10
30
RDU
0.6
0.4
0.2
0.0

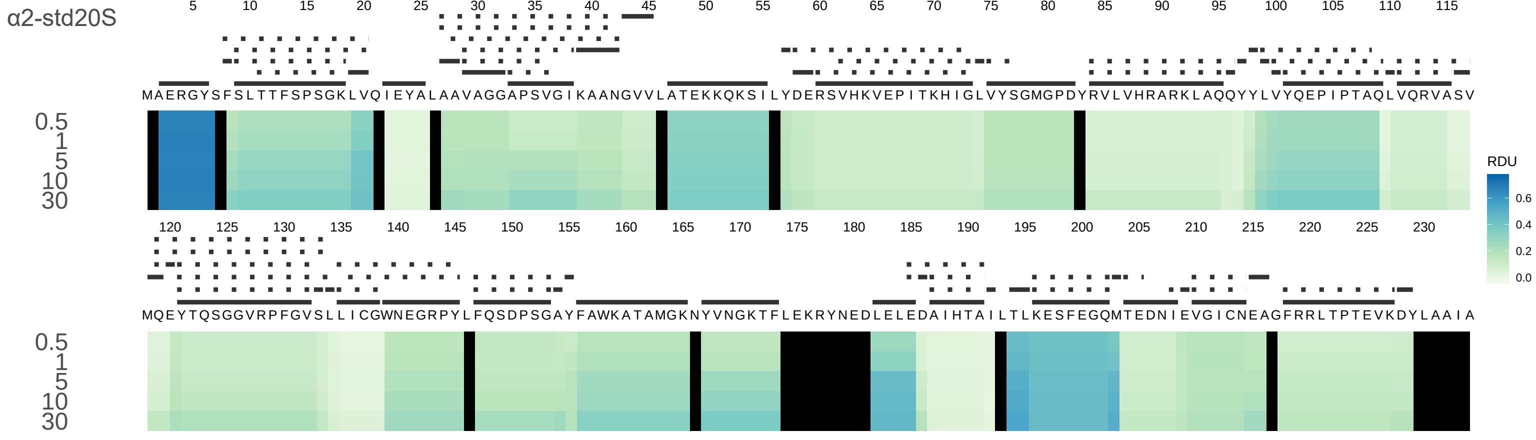

α2-std20S
MAERGYSFSLTTFSPSGKLVQIEYALAAVAGGAPSVGIKAANGVVLATEKKQKSILYDERSVHKVEPITKHIGLVYSGMGPDYRVLVHRARKLAQQYYLVYQEPIPTAQLVQRVASV
MQEYTQSGGVRPFGVSLLICGWNEGRPYLFQSDPSGAYFAWKATAMGKNYVNGKTFLEKRYNEDLELEDAIHTAILTLKESFEGQMTEDNIEVGICNEAGFRRLTPTEVKDYLAAIA
0.5
1
5
10
30
RDU
0.6
0.4
0.2
0.0

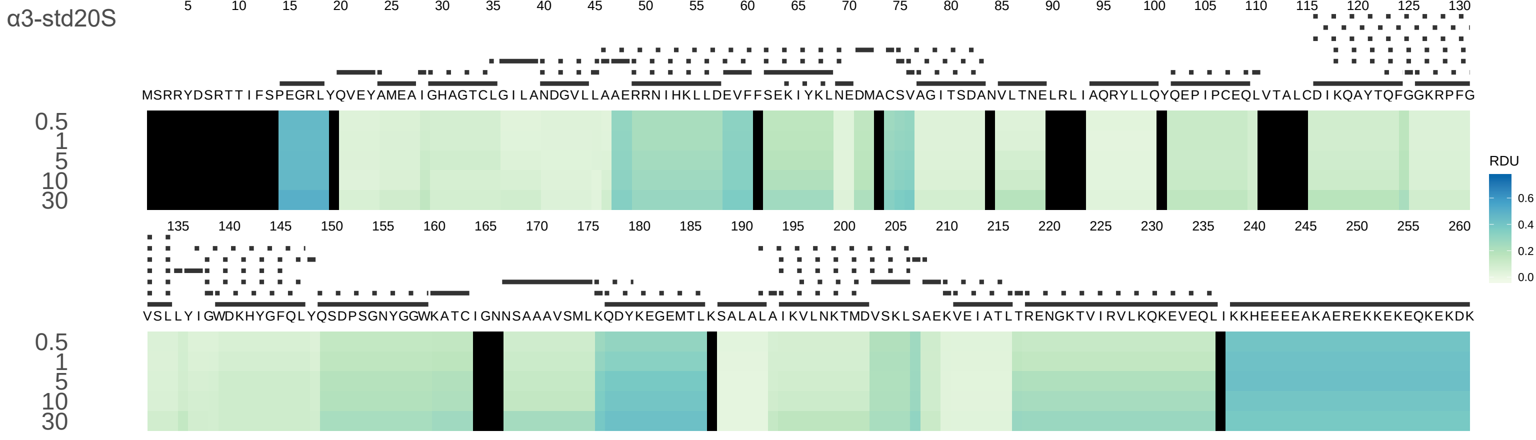
α3-std20S
MSRRYDSRTTIFSPEGRLYQVEYAMEAIGHAGTCLGILANDGVLLAAERRNIHKLLDEVFFSEKIYKLNEDMACSVAGITSDANVLTNELRLIAQRYLLQYQEPIPCEQLVTALCDIKQAYTQFGGKRPFG
VSLLYIGWDKHYGFQLYQSDPSGNYGGWKATCIGNNSAAAVSMLKQDYKEGEMTLKSALALAIKVLNKTMDVSKLSAEKVEIATLTRENGKTVIRVLKQKEVEQLIKKHEEEEAKAEREKKEKEQKEKDK
0.5
1
5
10
30
RDU
0.6
0.4
0.2
0.0

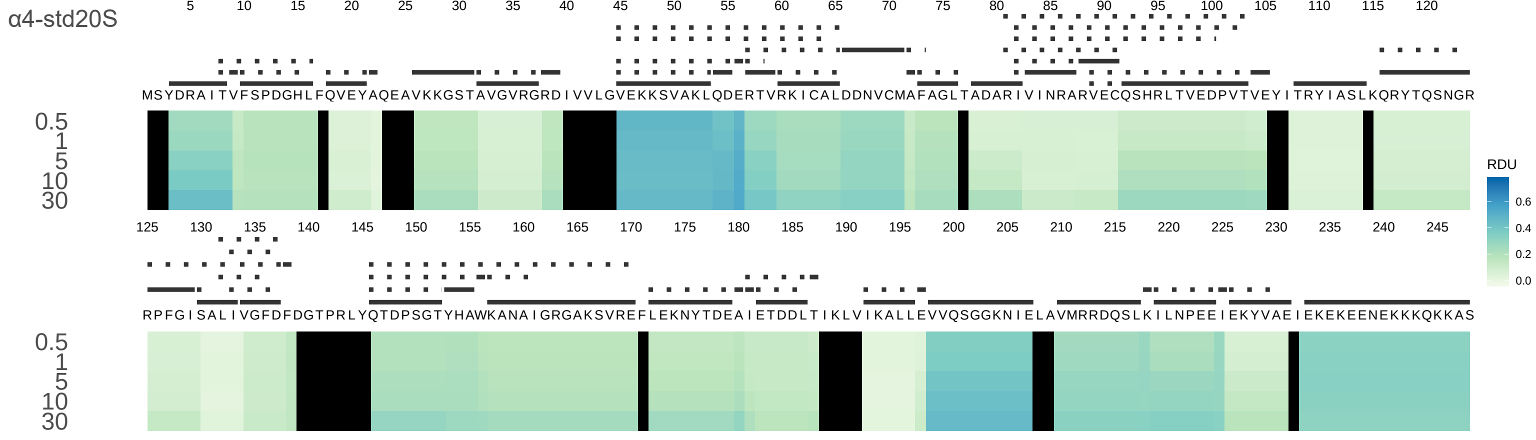
α4-std20S
5 10 15 20 25 30 35 40 45 50 55 60 65 70 75 80 85 90 95 100 105 110 115 120
MSYDRAITVFSPDGHLFQVEYAQEAVKKGSTAVGVRGRDIVVLGVEKKSVAKLQDERTVRKICALDDNVCMAFAGLTADARIVINRARVECQSHRLTVEDPVTVEYITRYIASLKQRYTQSNGR
0.5
1
5
10
30
125 130 135 140 145 150 155 160 165 170 175 180 185 190 195 200 205 210 215 220 225 230 235 240 245
RPFGISALIVGFDFDGTPRLYQTDPSGTYHAWKANAIGRGAKSVREFLEKNYTDEAIETDDLTIKLVIKALLEVVQSGGKNIELAVMRRDQSLKILNPEEIEKYVAEIEKEKEENEKKKQKKAS
0.5
1
5
10
30
RDU
0.6
0.4
0.2
0.0

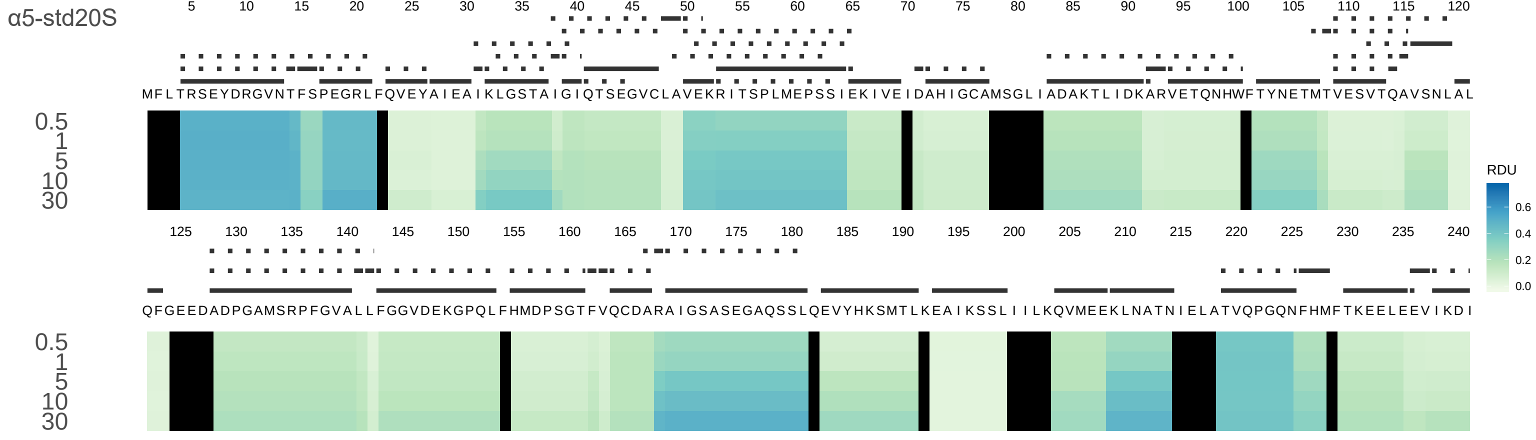

α5-std20S
5 10 15 20 25 30 35 40 45 50 55 60 65 70 75 80 85 90 95 100 105 110 115 120
MFLTRSEYDRGVNTFSPEGRLFQVEYAIEAIKLGSTAIGIQTSEGVCLAVEKRITSPLMEPSSIEKIVEIDAHIGCAMSGLIADAKTLIDKARVETQNHWFTYNETMTVESVTQAVSNLAL
0.5
1
5
10
30
125 130 135 140 145 150 155 160 165 170 175 180 185 190 195 200 205 210 215 220 225 230 235 240
QFGEEDADPGAMSRPFGVALLFGGVDEKGPQLFHMDPSGTFVQCDARAIGSASEGAQSSLQEVYHKSMTLKEAIKSSLIIILKQVMEEKLNATNIELATVQPGQNFHMFTKEELEEVIKDI
0.5
1
5
10
30
RDU
0.6
0.4
0.2
0.0

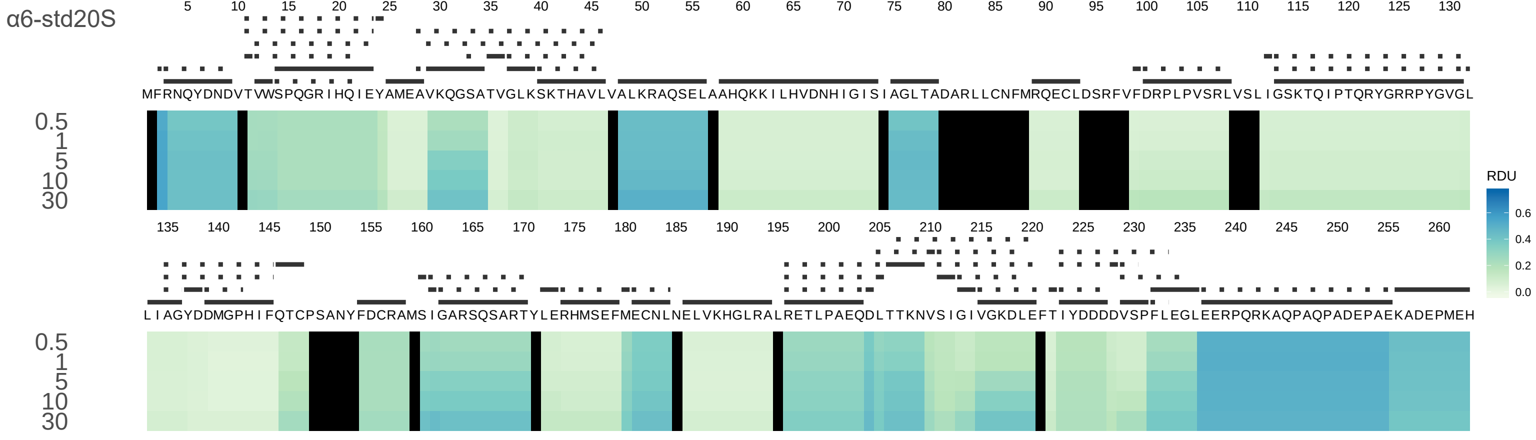
α6-std20S
MFRNQYDNDVTVWSPQGRIHQIEYAMEAVKQGSATVGLKSKTHAVLVALKRAQSELAAHQKKILHVDNHIGISIAGLTADARLLCNFMRQECLDSRFVFDRPLPVSRLVSLIGSKTQIPTQRYGRRPYGVGL
LIAGYDDMGPHIFQTCPSANYFDCRAMSIGARSQSARTYLERHMSEFMECNLNELVKHGLRALRETLPAEQDLTTKNVSIGIVGKDLEFTIYDDDDVSPFLEGLEERPQRKAQPAQPADEPAEKADEPMEH
0.5
1
5
10
30
RDU
0.6
0.4
0.2
0.0

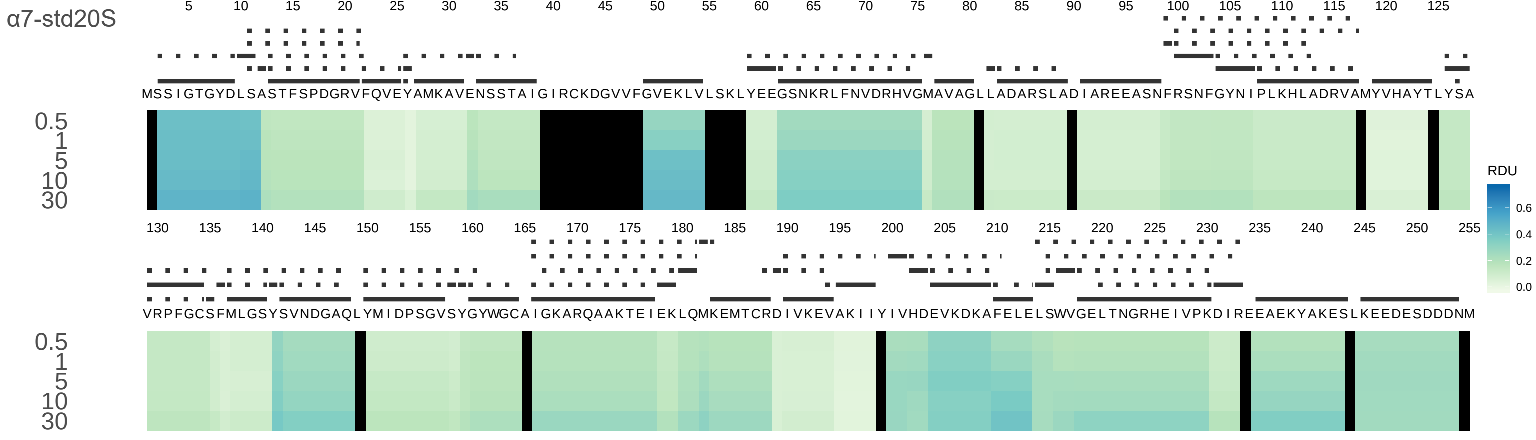
α7-std20S
MSSIGTGYDLSASTFSPDGRVFQVEYAMKAVENSSTAIGIRCKDGVVFGVEKLVLSKLYEEGSNKRLFNVDRHVGMAVAGLLADARSLADIAREEASNFRSNFGYNIPLKHLADRVAMYVHAYTLYSA
VRPFGCSFMLGSYSVNDGAQLYMIDPSGVSYGYWGCAIGKARQAAKTEIEKLQMKEMTCRDIVKEVAKIIYIVHDEVKDKAFELELSWVGELTNGRHEIVPKDIREEAEKYAKESLKEEDESDDDNM
0.5
1
5
10
30
RDU

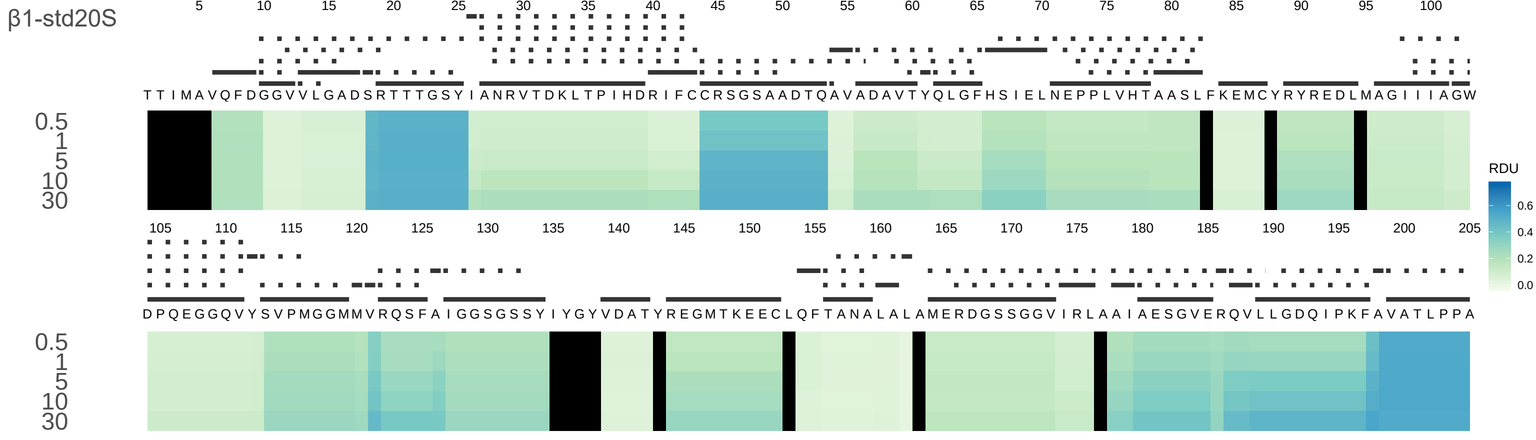
β1-std20S
TTIMAVQFDGGVVLGADSRTTTGSYIANRVTDKLTPIHDRIFCCRSGSAADTQAVADAVTYQLGFHSIELNEPPLVHTAASLFKEMCYRYREDLMAGIIIAGW
DPQEGGQVYSVPMGGMMVRQSFAIGGSGSSYIYGYVDATYREGMTKEECLQFTANALALAMERDGSSGGVIRLAAIAESGVERQVLLGDQIPKFAVATLPPA
0.5
1
5
10
30
RDU
0.6
0.4
0.2
0.0

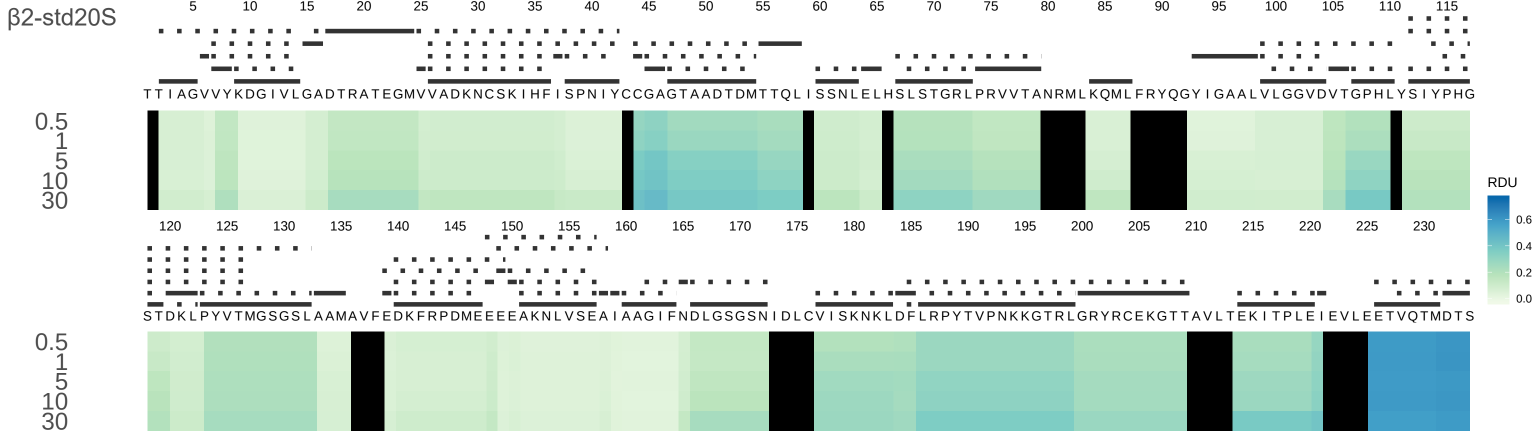
β2-std20S
TTIAGVVYKDGIVLGADTRATEGMVVADKNCSKIHFISPNIYCCGAGTAADTDMTTQLISSNLELHSLSTGRLPRVVTANRMLKQMLFRYQGYIGAALVLGGVDVTGPHLYSIYPHG
STDKLPYYVTMGSGSLAAMAVFEDKFRPDMEEEEAKNLVSEAIAAGIFNDLGSGSNIDLCVISKNKLDFLRPYTVPNKKGTRLGRYRCEKGTTAVLTEKITPLEIEVLEETVQTMDTS
0.5
1
5
10
30
RDU
0.6
0.4
0.2
0.0

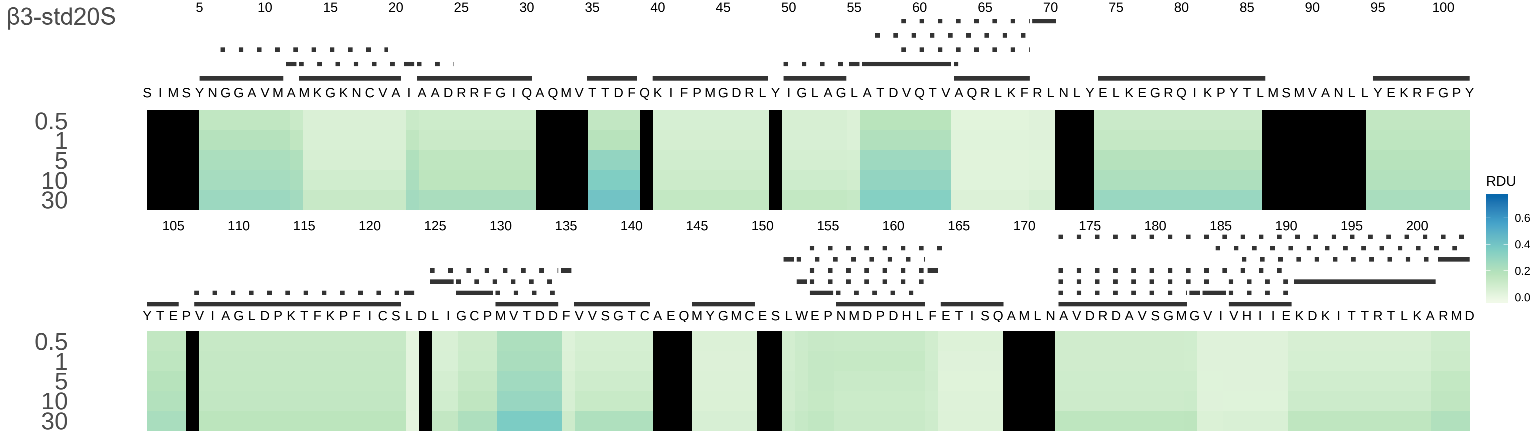
β3-std20S
S I M S Y N G G A V M A M K G K N C V A I A A D R R F G I Q A Q M V T T D F Q K I F P M G D R L Y I G L A G L A T D V Q T T V A Q R L K F R L N L Y E L K E G R Q I K P Y T L M S M V A N L L Y E K R F G P Y
Y T E P V I A G L D P K T F K P F I C S L D L I G C P M V T D D F V V S G T C A E Q M Y G M C E S L W E P N M D P D H L F E T I S Q A M L N A V D R D A V S G M G V I V H I I E K D K I T T R T L K A R M D
0.5
1
5
10
30
RDU

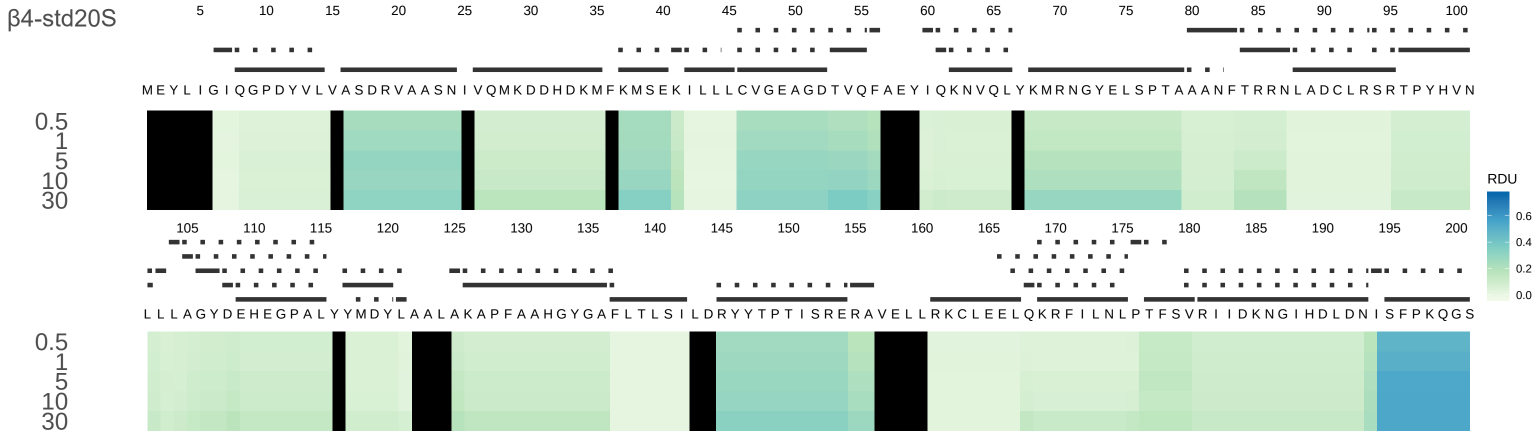

β4-std20S
MEYLIGIQGPDYVLVASDRVAASNIVQMKDDHDKMFKMSEKILLLCVGEAGDTVQFAEYIQKNVQLYKMRNGYELSPTAANFTRRNLADCLRSRTPYHVN
LLLAGYDEHEGPALYYMDYLAALAKAPFAAHGYGAFLTLSILDRYYTPTISRERAVELLRKCLEELQKRFILNLPTFSVRIIDKNGIHDLDNISFPKQGS
0.5
1
5
10
30
RDU

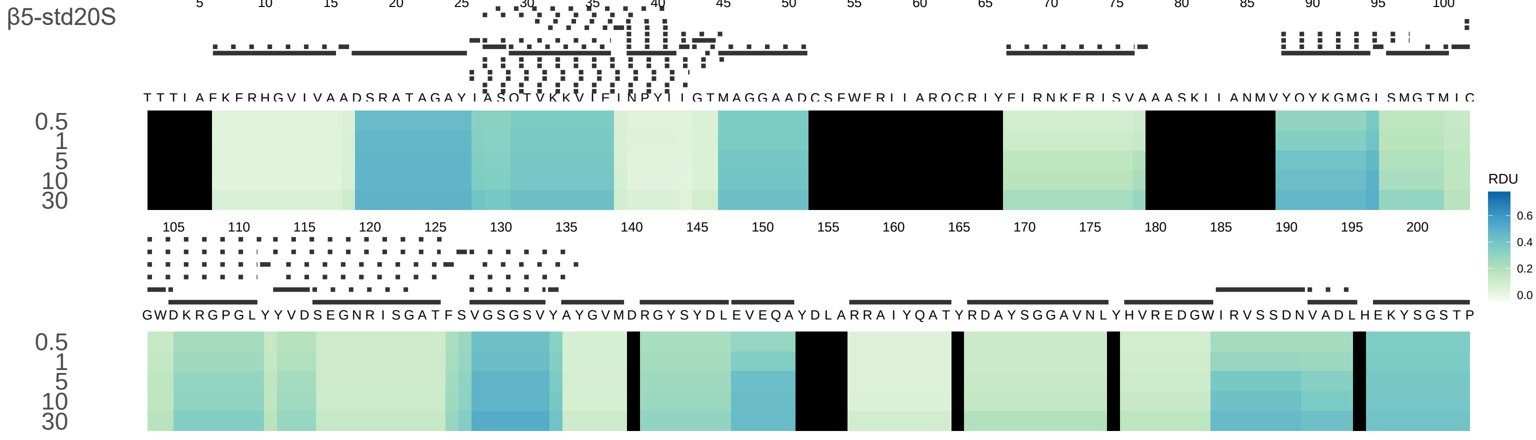
β5-std20S
RDU
0.6
0.4
0.2
0.0
0.5
1
5
10
30
TTTIAFKFRHGVIVAADSRATAGAYIASQTVKKVIEINPYLLGTMAGGAADCSFWERLLARQCRIYELRNKERISVAAASKLLANMVYQYKGMGLSMGTMIC
GWDKRGPGLYYVDSEGNRISGATFSVGSGSVYAYGVMDRGYSYDLEVEQAYDLARRAIYQATYRDAYSGGAVNLYHVREDGWIRVSSDNVADLHEKYSGSTP

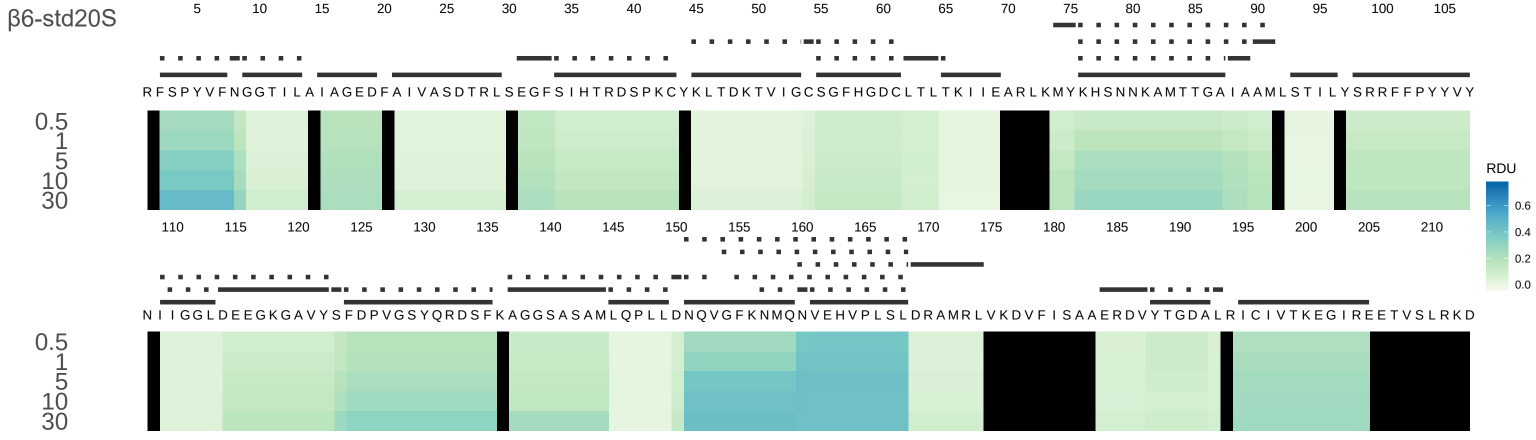
β6-std20S
RDU
0.0
0.2
0.4
0.6
0.5
1
5
10
30

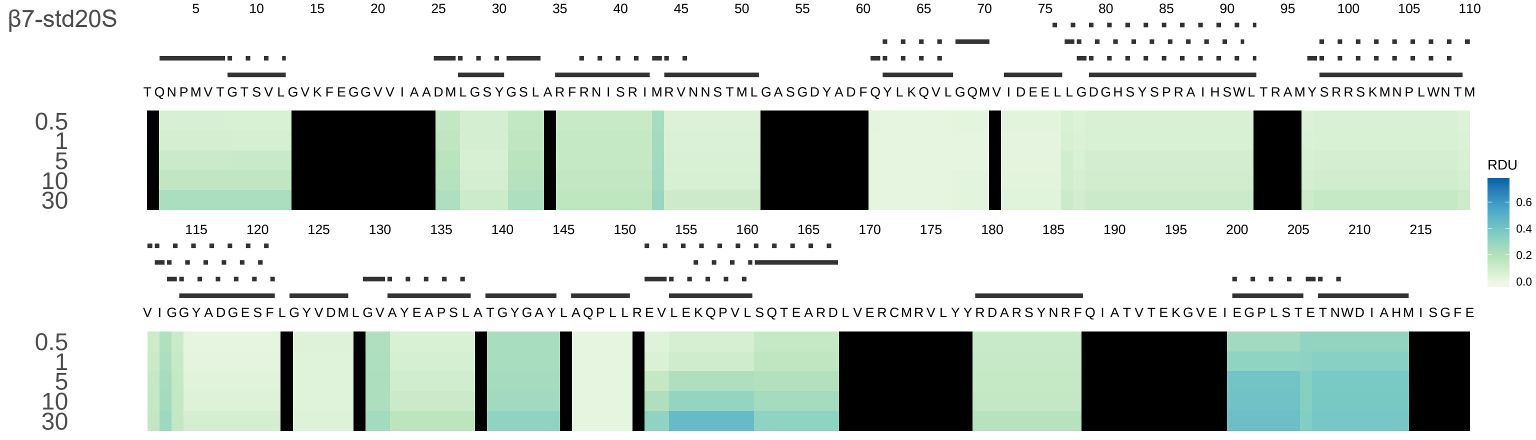
β7-std20S
TQNPMVTGTSVLGVKFEGGVVIAADMLGSYGSLARFRNISRIMRVNNSTMLGASGDYADFQYLKQVLGQMVIDEELLGDGHSYSPRAIHSWLTRAMYSRRSKMNPLWNTM
VIGGYADGESFLGYVDMLGVAYEAPSLATGYGAYLAQPLLREVLEKQPVLSQTEARDLVERCMRVLYYRDARSYNRFQIATVTEKGVEIEGPLSTETNWDIAHMISGFE
0.5
1
5
10
30
RDU
0.6
0.4
0.2
0.0

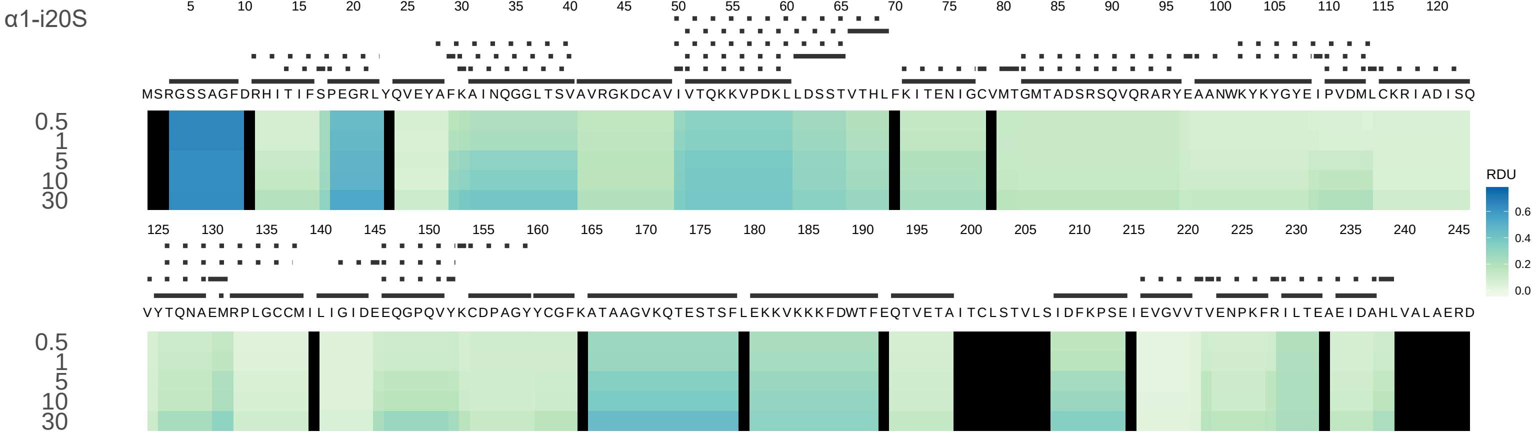
α1-i20S
MSRGSSAGFDRHITIFSPEGRLYQVEYAFKAINQGGLTSVAVRGKDCAVIVTQKKVPDKLLDSSTVTHLFKITENIGCVMTGMTADSRSQVQRARYEAANWKYKYGYEIPVDMLCKRIADISQ
VYTQNAEMRPLGCCMILIGIDEEQGPQVYKCDPAGYYCGFKATAAGVKQTESTSFLEKKVKKKFDWTFEQTVETAITCLSTVLSIDFKPSEIEVGVVTVENPKFRILTEAEIDAHLVALAERD
0.5
1
5
10
30
RDU
0.6
0.4
0.2
0.0

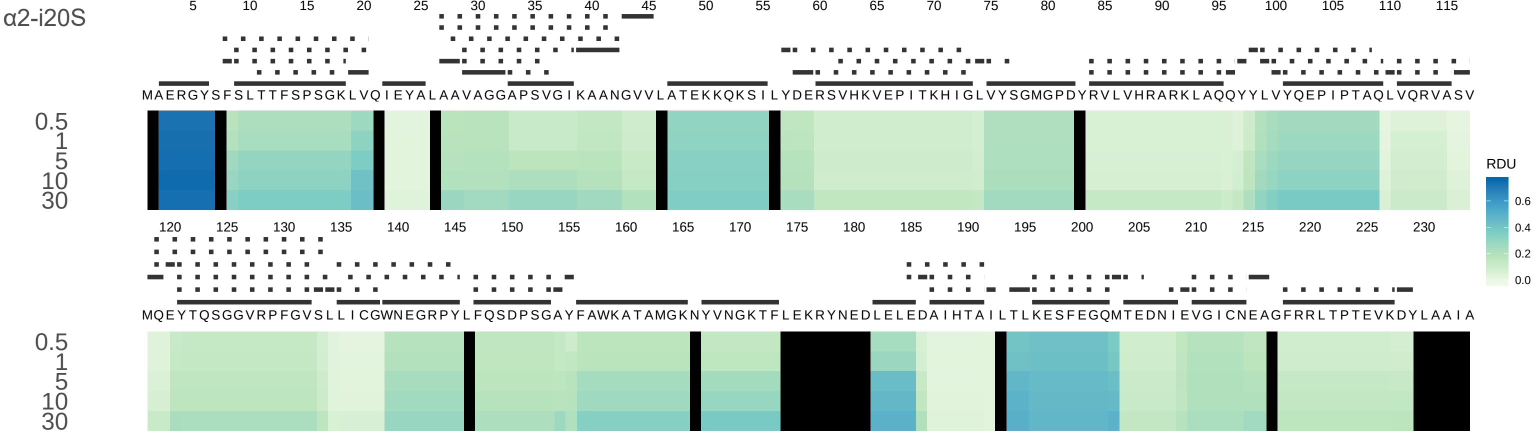

α2-i20S
MAERGYSFSLTTFSPSGKLVQIEYALAAVAGGAPSVGIKAANGVVLATEKKQKSILYDERSVHKVEPITKHIGLVYSGMGPDYRVLVHRARKLAQQYYLVYQEPIPTAQLVQRVASV
MQEYTQSGGVRPFGVSLLICGWNEGRPYLFQSDPSGAYFAWKATAMGKNYVNGKTFLEKRYNEDLELEDAIHTAILTLKESFEGQMTEDNIEVGICNEAGFRRLTPTEVKDYLAAIA
0.5
1
5
10
30
RDU
0.6
0.4
0.2
0.0

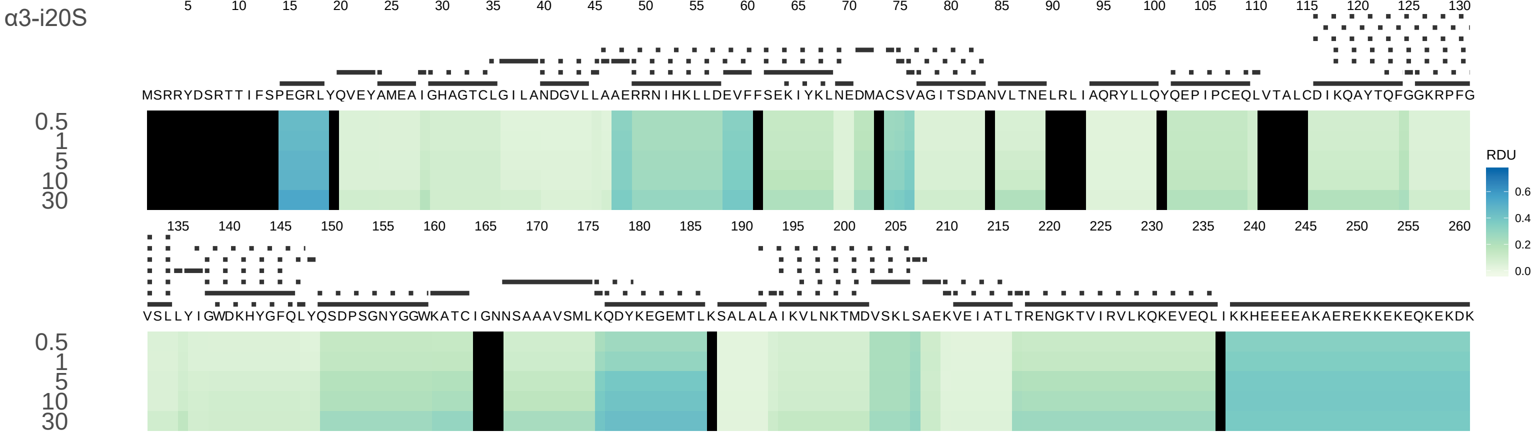
α3-i20S
MSRRYDSRTTIFSPEGRLYQVEYAMEAIGHAGTCLGILANDGVLLAAERRNIHKLLDEVFFSEKIYKLNEDMACSVAGITSDANVLTNELRLIAQRYLLQYQEPIPCEQLVTALCDIKQAYTQFGGKRPFG
VSLLYIGWDKHYGFQLYQSDPSGNYGGWKATCIGNNSAAAVSMLKQDYKEGEMTLKSALALAIKVLNKTMDVSKLSAEKVEIATLTRENGKTVIRVLKQKEVEQLIKKHEEEEAKAEREKKEKEQKEKDK
0.5
1
5
10
30
RDU
0.6
0.4
0.2
0.0

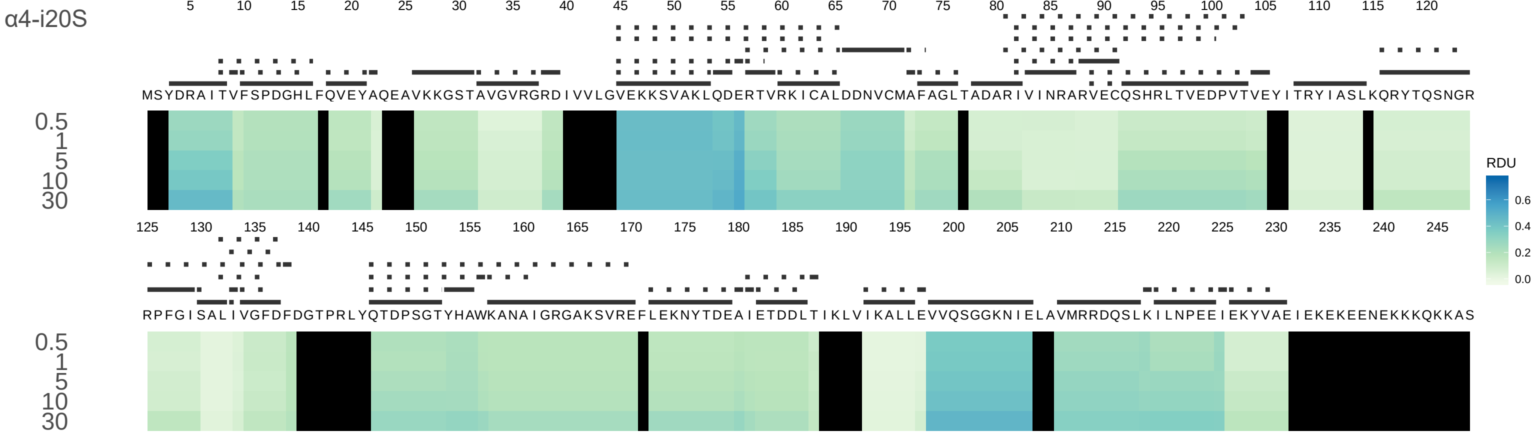

α4-i20S
5 10 15 20 25 30 35 40 45 50 55 60 65 70 75 80 85 90 95 100 105 110 115 120
MSYDRAITVFSPDGHLFQVEYAQEAVKKGSTAVGVRGRDIVVLGVEKKSVAKLQDERTVRKICALDDNVCMAFAGLTADARIVINRARVECQSHRLTVEDPVTVEYITRYIASLKQRYTQSNGR
0.5
1
5
10
30
125 130 135 140 145 150 155 160 165 170 175 180 185 190 195 200 205 210 215 220 225 230 235 240 245
RPFGISALIVGFDFDGTPRLYQTDPSGTYHAWKANAIGRGAKSVREFLEKNYTDEAIETDDLTIKLVIKALLEVVQSGGKNIELAVMRRDQSLKILNPEEIEKYVAEIEKEKEENEKKKQKKAS
0.5
1
5
10
30
RDU
0.6
0.4
0.2
0.0

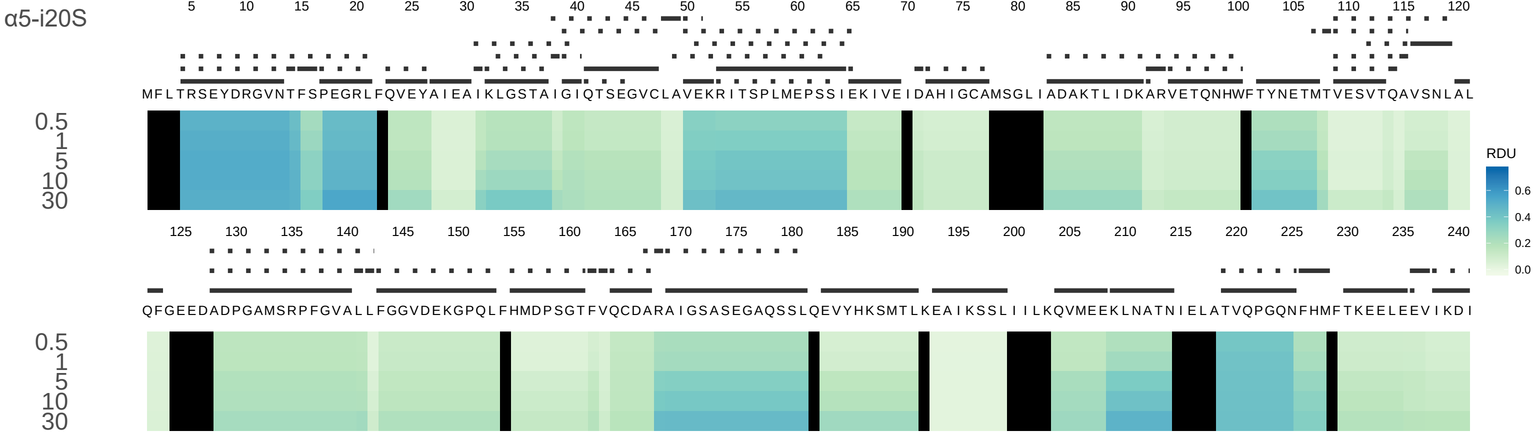
α5-i20S
MFLTRSEYDRGVNTFSPEGRLFQVEYAIEAIKLGSTAIGIQTSEGVCLAVEKRITSPLMEPSSIEKIVEIDAHIGCAMSGLIADAKTLIDKARVETQNHWFTYNETMTVESVTQAVSNLAL
QFGEEDADPGAMSRPFGVALLFGGVDEKGPQLFHMDPSGTFVQCDARAIGSASEGAQSSLQEVYHKSMTLKEAIKSSLIIILKQVMEEKLNATNIELATVQPGQNFHMFTKEELEEVIKDI
0.5
1
5
10
30
RDU
0.6
0.4
0.2
0.0

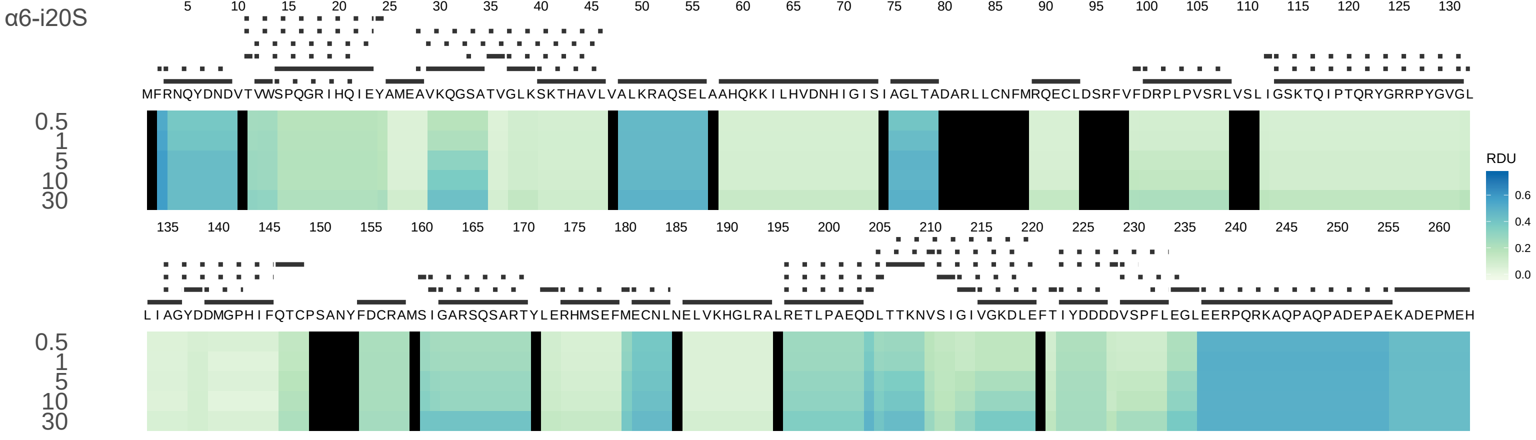
α6-i20S
MFRNQYDNDVTVWSPQGRIHQIEYAMEAVKQGSATVGLKSKTHAVLVALKRAQSELAAHQKKILHVDNHIGISIAGLTADARLLCNFMRQECLDSRFVFDRPLPVSRLVSLIGSKTQIPTQRYGRRPYGVGL
LIAGYDDMGPHIFQTCPSANYFDCRAMSIGARSQSARTYLERHMSEFMECNLNELVKHGLRALRETLPAEQDLTTKNVSIGIVGKDLEFTIYDDDDVSPFLEGLEERPQRKAQPAQPADEPAEKADEPMEH
0.5
1
5
10
30
RDU
0.6
0.4
0.2
0.0

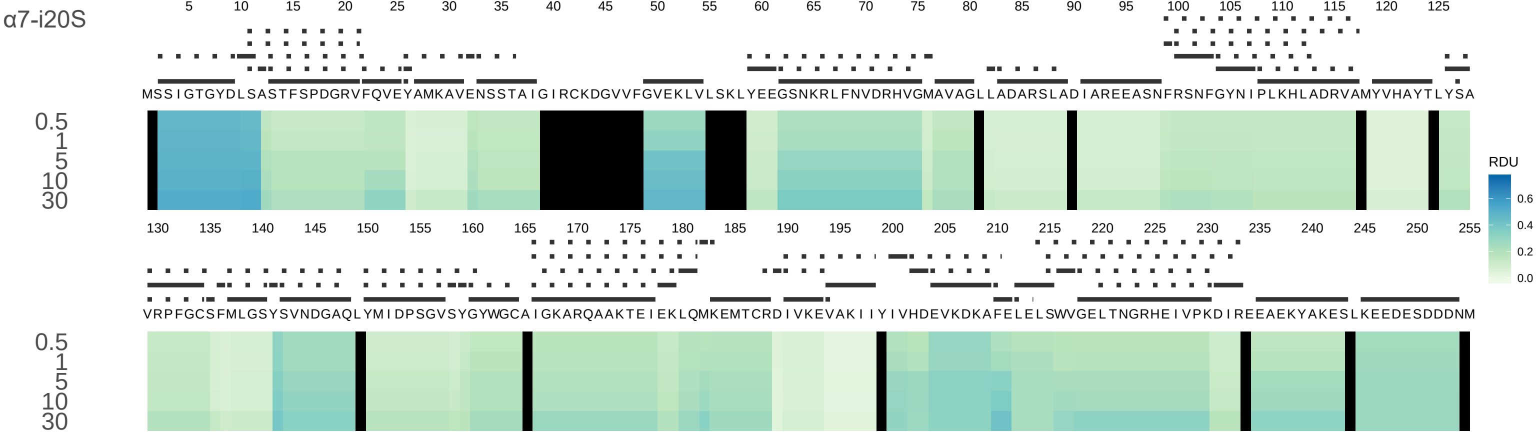

α7-i20S
MSSIGTGYDLSASTFSPDGRVFQVEYAMKAVENSSTAIGIRCKDGVVFGVEKLVLSKLYEEGSNKRLFNVDRHVGMAVAGLLADARSLADIAREEASNFRSNFGYNIPLKHLADRVAMYVHAYTLYSA
VRPFGCSFMLGSYSVNDGAQLYMIDPSGVSYGYWGCAIGKARQAAKTEIEKLQMKEMTCRDIVKEVAKIIYIVHDEVKDKAFELELSWVGELTNGRHEIVPKDIREEAEKYAKESLKEEDESDDDNM
0.5
1
5
10
30
RDU
0.6
0.4
0.2
0.0

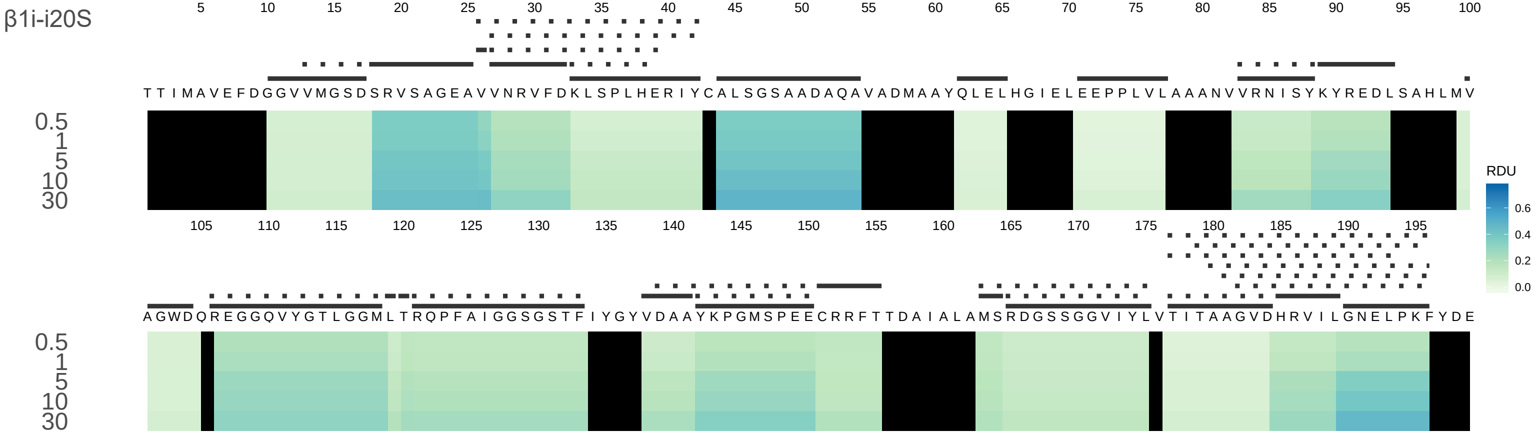
β1i-i20S
TTIMAVEFDGGVVMGSDSRVSAGEAVVNRVFDKLSPLHERIYCALSGSAADAQAVADMAAYQLELHGIELEEPPLVLAAANVVRNISYKYREDLSAHLMV
AGWDQREGGQVYGTLGGMLTRQPFAIGGSGSTFIYGYVDAAYKPGMSPEECRRFTTDAIALAMSRDGSSGGVIYLVTITAAGVDHRVILGNELPKFYDE
0.5
1
5
10
30
RDU
0.6
0.4
0.2
0.0

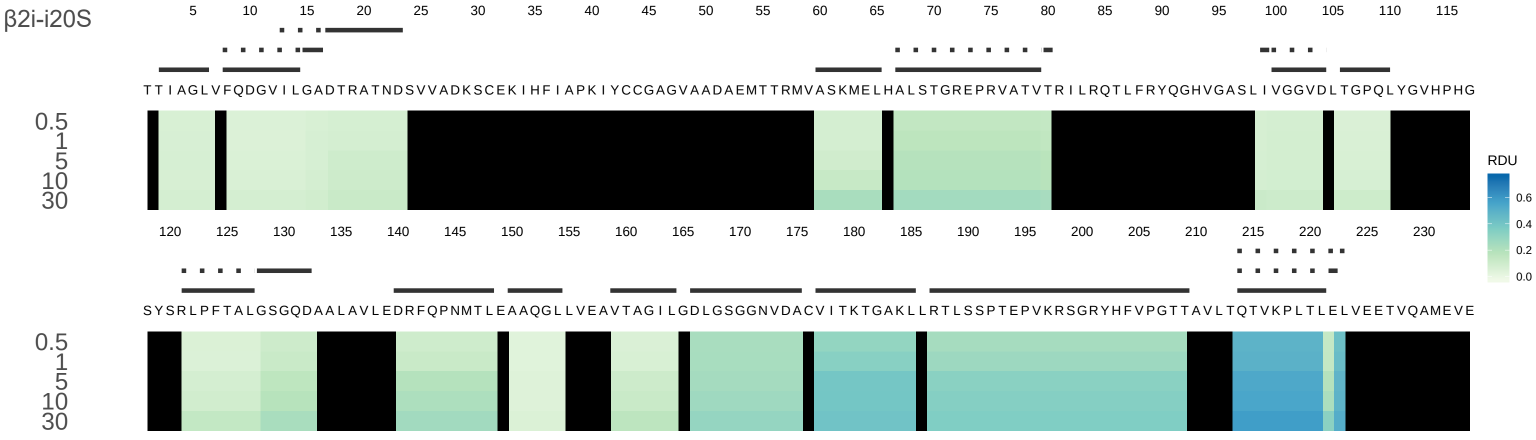
β2i-i20S
TTIAGLVFQDGVILGADTRATNDSVVADKSCEKIHFIAPKIYCCGAGVAADAEMTTRMVASKMELHALSTGREPRVATVTRILRQTLFRYQGHVGASLIVGGVDLTGPQLYGVHPHG
SYSRLPFTALGSGQDAALAVLEDRFQPNMTLEAAQGLLVEAVTAGILGDLGSGGNVDACVITKTGAKLLRTLSSPTEPVKRSGRYHFVPGTTAVLTQTVKPLTLELVEETVQAMEVE
0.5
1
5
10
30
RDU
0.6
0.4
0.2
0.0

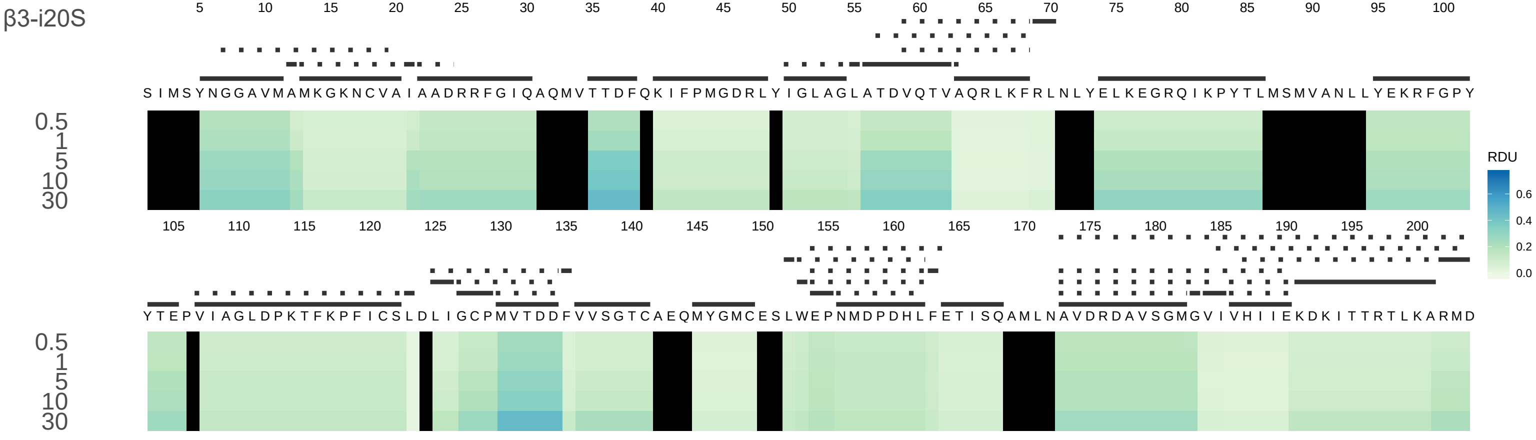
β3-i20S
RDU
0.6
0.4
0.2
0.0
0.5
1
5
10
30
SIMSYNGGAVMAMKGKNCVAIAADRRFGIQAQMVTTDFQKIFPMGDRLYIGLAGLATDVQTVAQRLKFRLNLYELKEGRQIKPYTLMSMVANLLYEKRFGPY
YTEPVIAGLDPKTFKPFICSLDLIGCPMVTDDFVVSGTCAEQMYGMCESLWEPNMDPDHLFETISQAMLNAVDRDAVSGMGVIVHIIEKDKITTRTLKARMD

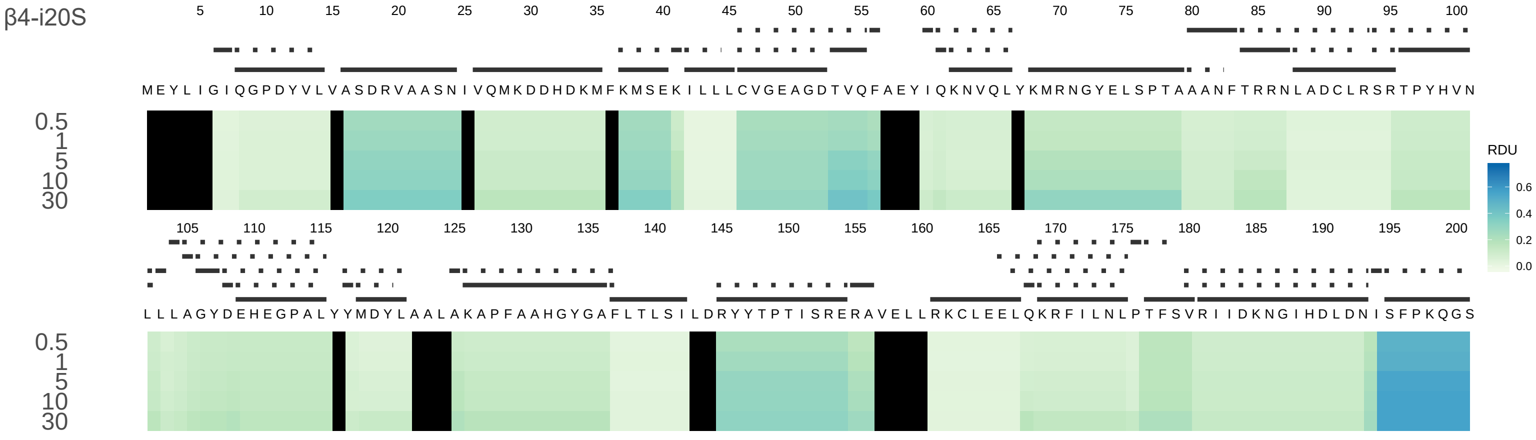

β4-i20S
M E Y L I G I Q G P D Y V L V A S D R V A A S N I V Q M K D D H D K M F K M S E K I L L L C V G E A G D T V Q F A E Y I Q K N V Q L Y K M R N G Y E L S P T A A A N F T R R N L A D C L R S R T P Y H V N
L L L A G Y D E H E G P A L Y Y M D Y L A A L A K A P F A A H G Y G A F L T L S I L D R Y Y T P T I S R E R A V E L L R K C L E E L Q K R F I L N L P T F S V R I I D K N G I H D L D N I S F P K Q G S
0.5
1
5
10
30
RDU
0.6
0.4
0.2
0.0

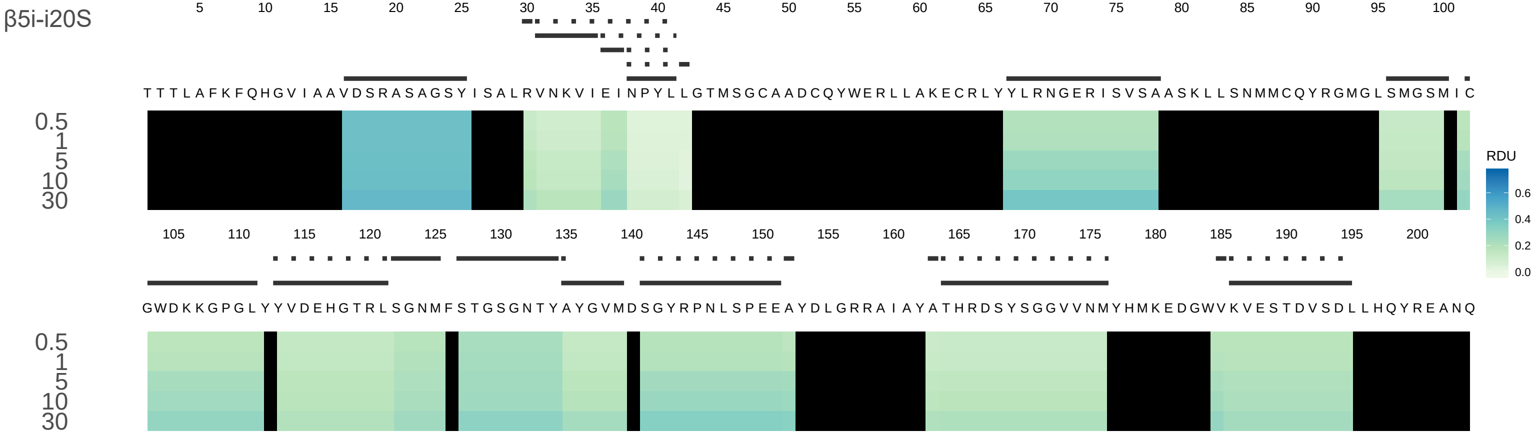
β5i-i20S
TTTLAFKFQHGVIAAVDSRASAGSYISALRVNKVIEINPYLLGTMSGCAADCQYWERLLAKECRLYYLRNGERISVSAASKLLSNMMCQYRGMGLSMGSMIC
GWDKKGPGLYYVDEHGTRLSGNMFSTGSGNTYAYGVMDSGYRPNLSPEEAYDLGRRAIAYATHRDSYSGGVVNMYHMKEDGWVKVESTDVSDLLHQYREANQ
0.5
1
5
10
30
RDU
0.6
0.4
0.2
0.0

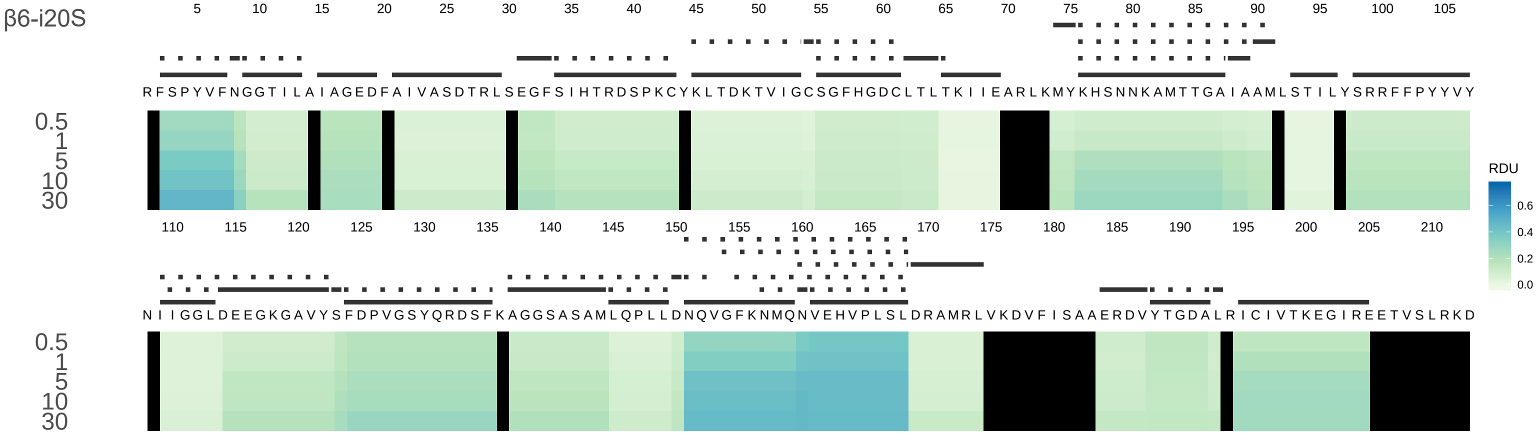
β6-i20S
RDU
0.0
0.2
0.4
0.6
0.5
1
5
10
30
R F S P Y V F N G G T I L A I A G E D F A I V A S D T R L S E G F S I H T R D S P K C Y K L T D K T V I G C S G F H G D C L T L T K I I E A R L K M Y K H S N N K A M T T G A I A A M L S T I L Y S R R F F P Y Y V Y
N I I G G L D E E G K G A V Y S F D P V G S Y Q R D S F K A G G S A S A M L Q P L L D N Q V G F K N M Q N V E H V P L S L D R A M R L V K D V F I S A A E R D V Y T G D A L R I C I V T K E G I R E E T V S L R K D

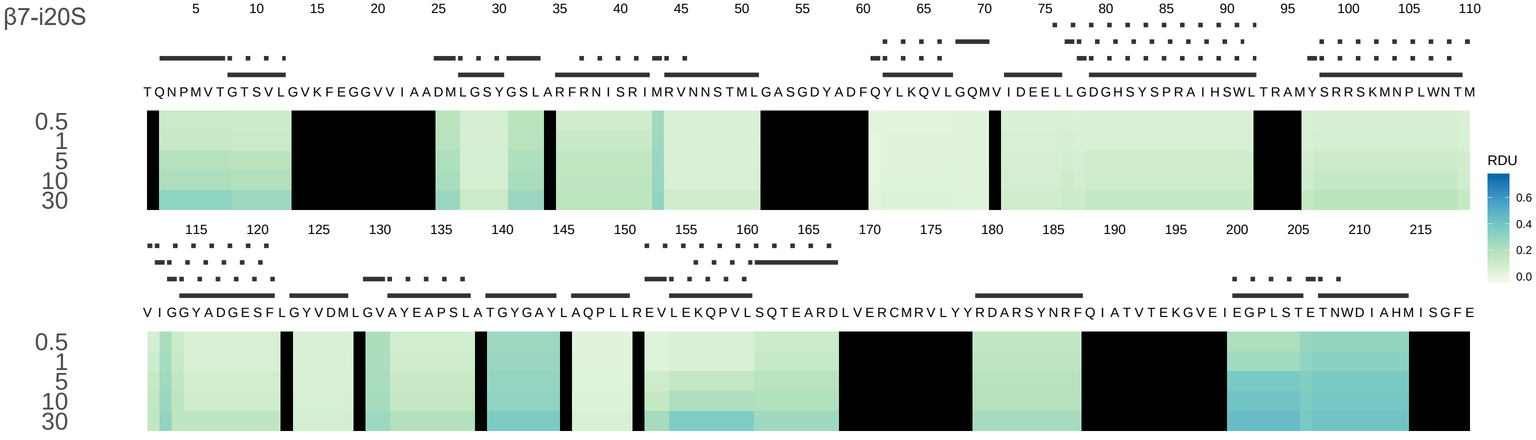
β7-i20S
5 10 15 20 25 30 35 40 45 50 55 60 65 70 75 80 85 90 95 100 105 110
T Q N P M V T G T S V L G V K F E G G V V I A A D M L G S Y G S L A R F R N I S R I M R V N N S T M L G A S G D Y A D F Q Y L K Q V L G Q M V I D E E L L G D G H S Y S P R A I H S W L T R A M Y S R R S K M N P L W N T M
0.5
1
5
10
30
115 120 125 130 135 140 145 150 155 160 165 170 175 180 185 190 195 200 205 210 215
V I G G Y A D G E S F L G Y V D M L G V A Y E A P S L A T G Y G A Y L A Q P L L R E V L E K Q P V L S Q T E A R D L V E R C M R V L Y Y R D A R S Y N R F Q I A T V T E K G V E I E G P L S T E T N W D I A H M I S G F E
0.5
1
5
10
30
RDU
0.6
0.4
0.2
0.0
