## Supplementary material for "Conformational maps of human 20S proteasomes reveal PA28- and immuno-dependent inter-ring crosstalks": Fig. S4

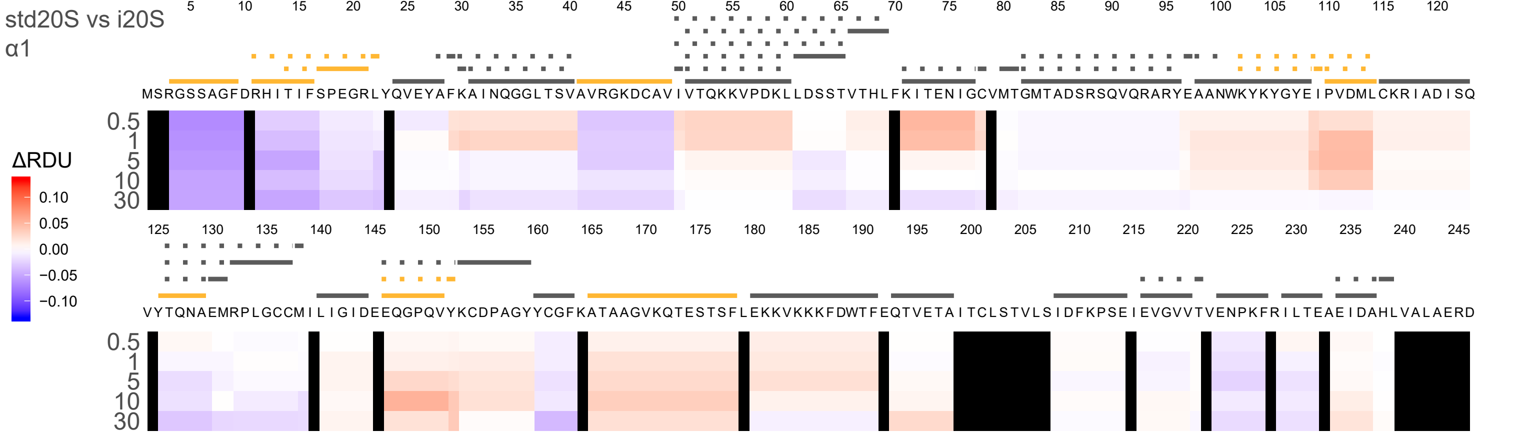
std20S vs i20S
α1
ΔRDU
0.10
0.05
0.00
−0.05
−0.10
MSRGSSAGFDRHITIFSPEGRLYQVEYAFKAINQGGLTSVAVRGKDCAVIVTQKKVPDKLLDSSTVTHLFKITENIGCVMTGMTADSRSQVQRARYEAANWKYKYGYEIPVDMLCKRIADISQ
VYTQNAEMRPLGCCMILIGIDEEQGPQVYKCDPAGYYCGFKATAAGVKQTESTSFLEKKVKKKFDWTFEQTVETAITCLSTVLSIDFKPSEIEVGVVTVENPKFRILTEAEIDAHLVALAERD
0.5
1
5
10
30

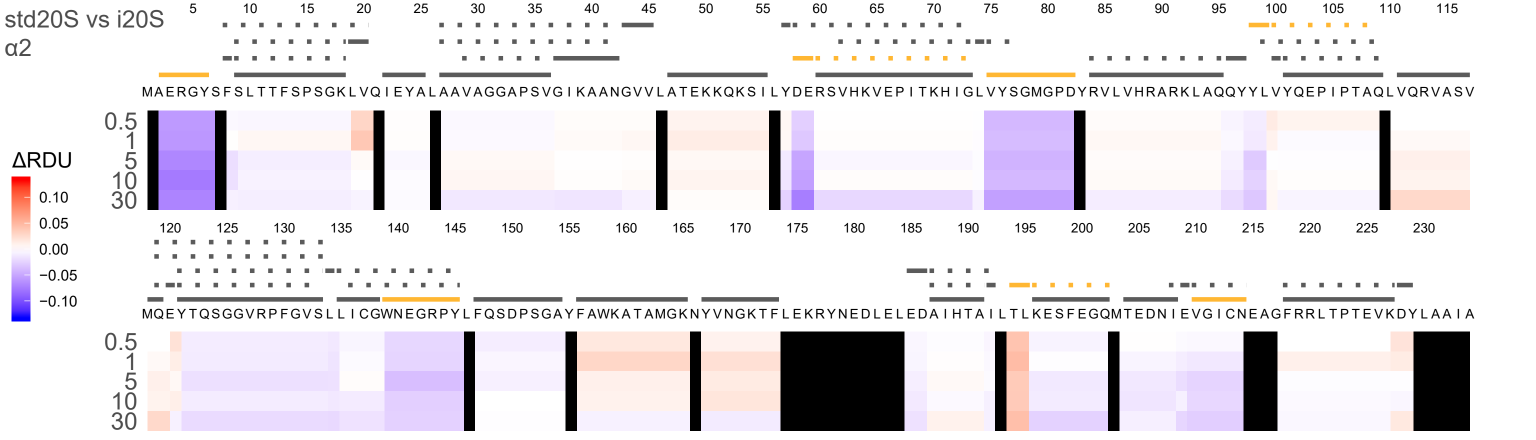
std20S vs i20S
α2
ΔRDU
0.10
0.05
0.00
−0.05
−0.10
5 10 15 20 25 30 35 40 45 50 55 60 65 70 75 80 85 90 95 100 105 110 115
MAERGYSFSLTTFSPSGKLVQIEYALAAVAGGAPSVGIKAANGVVLATEKKQKSILYDERSVHKVEPITKHIGLVYSGMGPDYRVLVHRARKLAQQYYLVYQEPIPTAQLVQRVASV
0.5
1
5
10
30
120 125 130 135 140 145 150 155 160 165 170 175 180 185 190 195 200 205 210 215 220 225 230
MQEYTQSGGVRPFGVSLLICGWNEGRPYLFQSDPSGAYFAWKATAMGKNYVNGKTFLEKRYNEDLELEDAIHTAILTLKESFEGQMTEDNIEVGICNEAGFRRLTPTEVKDYLAAIA
0.5
1
5
10
30

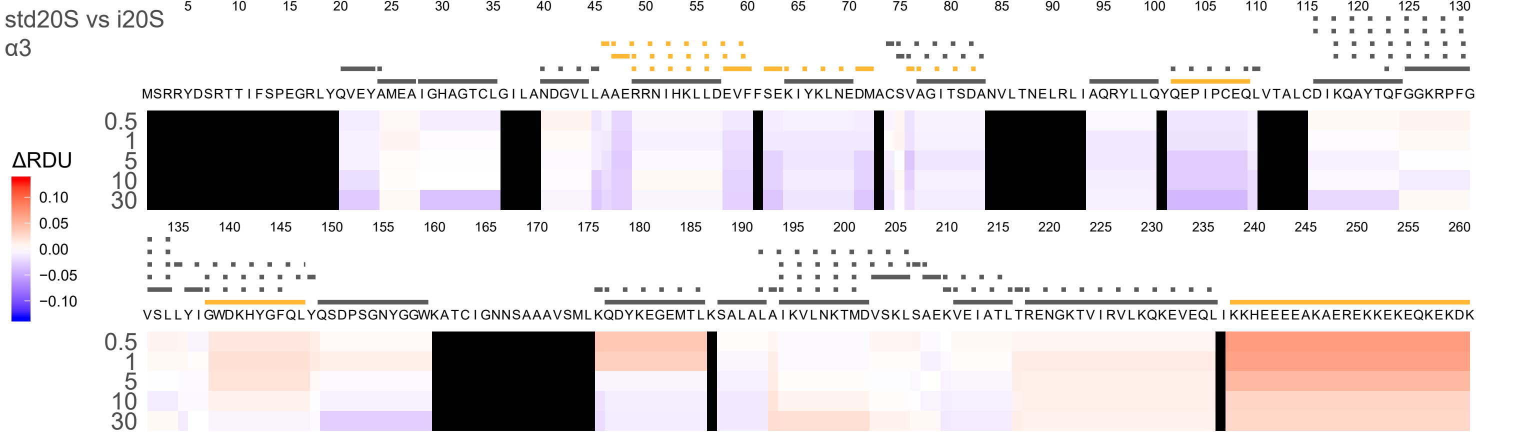
std20S vs i20S
α3
ΔRDU
0.10
0.05
0.00
−0.05
−0.10
5 10 15 20 25 30 35 40 45 50 55 60 65 70 75 80 85 90 95 100 105 110 115 120 125 130
MSRRYDSRTTIFSPEGRLYQVEYAMEAIGHAGTCLGILANDGVLLAAERRNIHKLLDEVFFSEKIYKLNEDMACSVAGITSDANVLTNELRLIAQRYLLQYQEPIPCEQLVTALCDIKQAYTQFGGKRPFG
0.5
1
5
10
30
135 140 145 150 155 160 165 170 175 180 185 190 195 200 205 210 215 220 225 230 235 240 245 250 255 260
VSLLYIGWDKHYGFQLYQSDPSGNYGGWKATCIGNNSAAAVSMLKQDYKEGEMTLKSALALAIKVLNKTMDVSKLSAEKVEIATLTRENGKTVIRVLKQKEVEQLIKKHEEEEAKAEREKKEKEQKEKDK
0.5
1
5
10
30

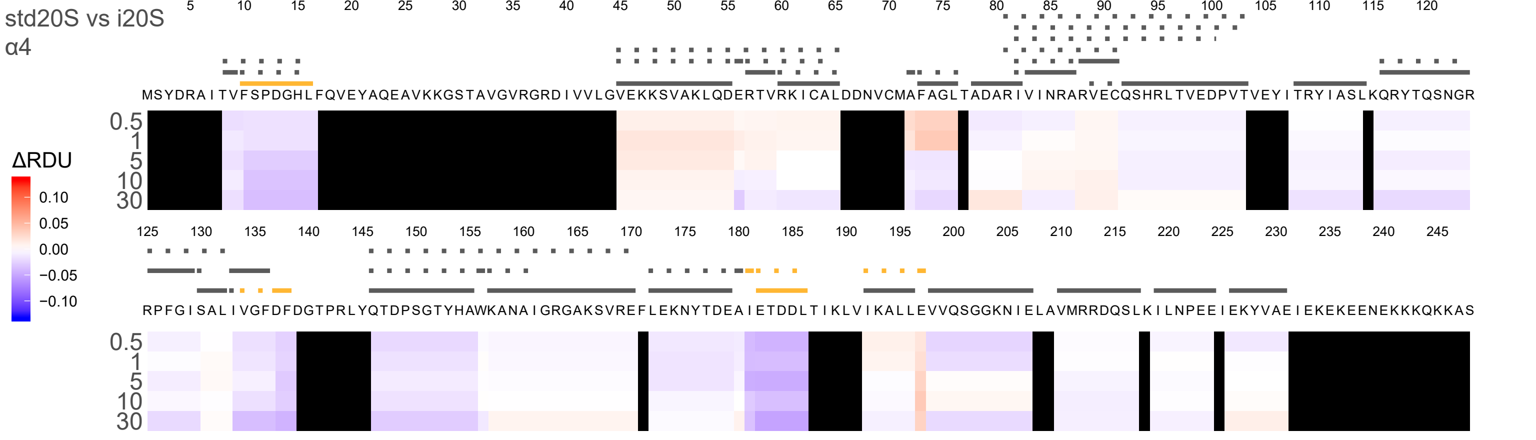

std20S vs i20S
α4
ΔRDU
0.10
0.05
0.00
−0.05
−0.10
5 10 15 20 25 30 35 40 45 50 55 60 65 70 75 80 85 90 95 100 105 110 115 120
MSYDRAITVFSPDGHLFQVEYAQEAVKKGSTAVGVRGRDIVVLGVEKKSVAKLQDERTVRKICALDDNVCMAFAGLTADARIVINRARVECQSHRLTVEDPVTVEYITRYIASLKQRYTQSNGR
0.5
1
5
10
30
125 130 135 140 145 150 155 160 165 170 175 180 185 190 195 200 205 210 215 220 225 230 235 240 245
RPFGISALIVGFDFDGTPRLYQTDPSGTYHAWKANAIGRGAKSVREFLEKNYTDEAIETDDLTIKLVIKALLEVVQSGGKNIELAVMRRDQSLKILNPEEIEKYVAEIEKEKEENEKKKQKKAS
0.5
1
5
10
30

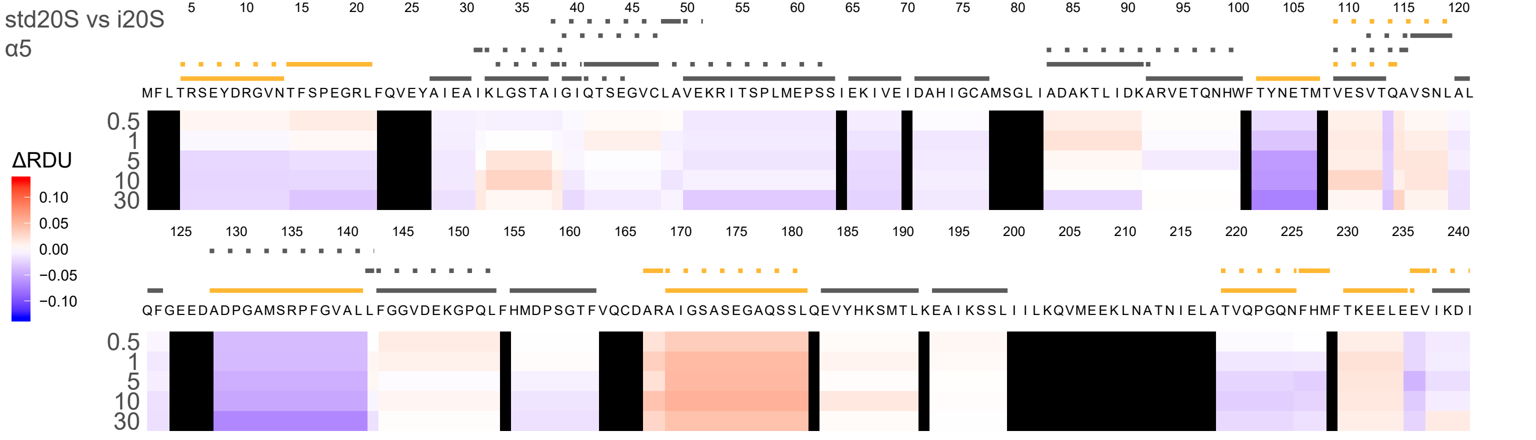
std20S vs i20S
α5
ΔRDU
0.10
0.05
0.00
−0.05
−0.10
MFLTRSEYDRGVNTFSPEGRLFQVEYAIEAIKLGSTAIGIQTSEGVCLAVEKRITSPLMEPSSIEKIVEIDAHIGCAMSGLIADAKTLIDKARVETQNHWFTYNETMTVESVTQAVSNLAL
QFGEEDADPGAMSRPFGVALLFGGVDEKGPQLFHMDPSGTFVQCDARAIGSASEGAQSSLQEVYHKSMTLKEAIKSSLIIILKQVMEEKLNATNIELATVQPGQNFHMFTKEELEEVIKDI
0.5
1
5
10
30

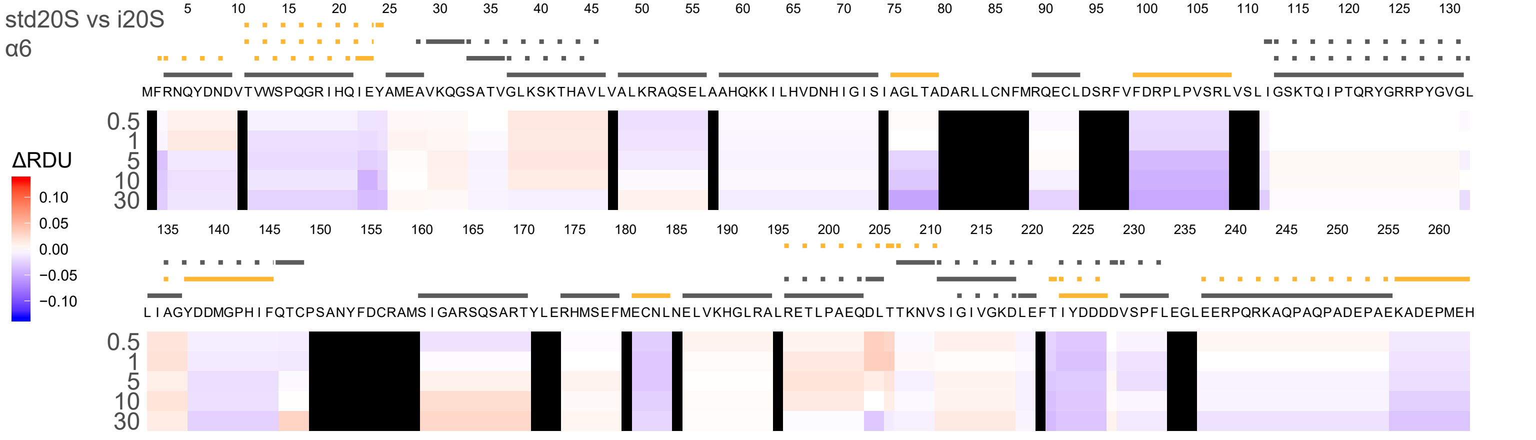

std20S vs i20S
α6
ΔRDU
0.10
0.05
0.00
−0.05
−0.10
0.5
1
5
10
30
MFRNQYDNDVTVWSPQGRIHQIEYAMEAVKQGSATVGLKSKTHAVLVALKRAQSELAAHQKKILHVDNHIGISIAGLTADARLLCNFMRQECLDSRFVFDRPLPVSRLVSLIGSKTQIPTQRYGRRPYGVGL
LIAGYDDMGPHIFQTCPSANYFDCRAMSIGARSQSARTYLERHMSEFMECNLNELVKHGLRALRETLPAEQDLTTKNVSIGIVGKDLEFTIYDDDDVSPFLEGLEERPQRKAQPAQPADEPAEKADEPMEH

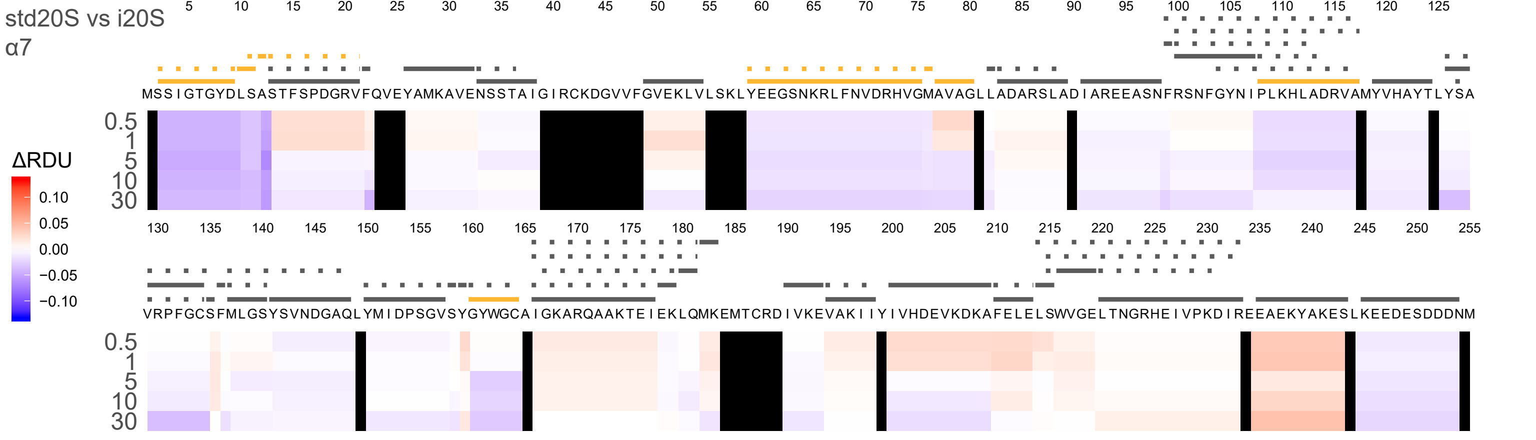
std20S vs i20S
α7
ΔRDU
0.10
0.05
0.00
−0.05
−0.10
0.5
1
5
10
30

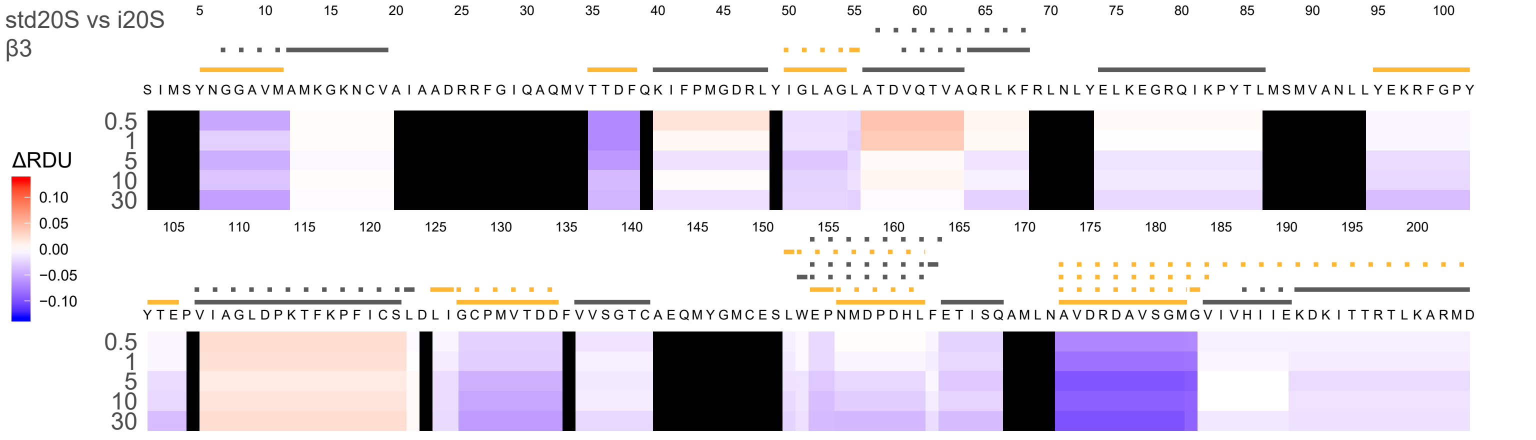

std20S vs i20S
β3
ΔRDU
0.10
0.05
0.00
−0.05
−0.10
5 10 15 20 25 30 35 40 45 50 55 60 65 70 75 80 85 90 95 100
SIMSYNGGAVMAMKGKNCVAIAADRRFGIQAQMVTTDFQKIFPMGDRLYIGLAGLATDVQTVAQRLKFRLNLYELKEGRQIKPYTLMSMVANLLYEKRFGPY
0.5
1
5
10
30
105 110 115 120 125 130 135 140 145 150 155 160 165 170 175 180 185 190 195 200
YTEPVIAGLDPKTFKPFICSLDLIGCPMVTDDFVVSGTCAEQMYGMCESLWEPNMDPDHLFETISQAMLNAVDRDAVSGMGVIVHIIEKDKITTRTLKARMD
0.5
1
5
10
30

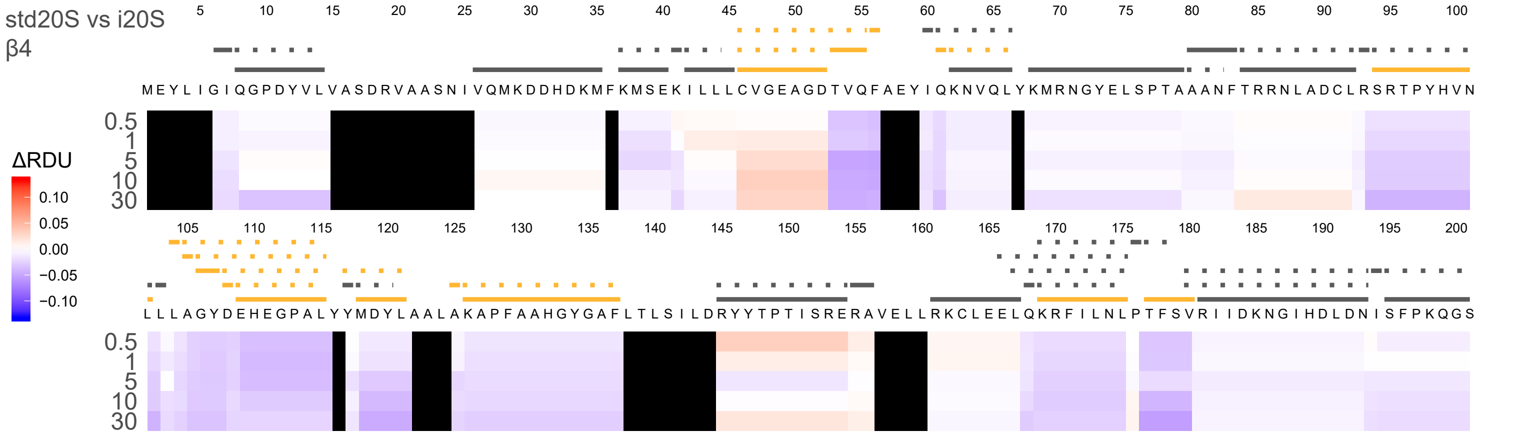
std20S vs i20S
β4
ΔRDU
0.10
0.05
0.00
−0.05
−0.10
0.5
1
5
10
30
MEYLIGIQGPDYVLVASDRVAASNIVQMKDDHDKMFKMSEKILLLCVGEAGDTVQFAEYIQKNVQLYKMRNGYELSPTAAANFTRRNLADCLRSRTPYHVN
LLLAGYDEHEGPALYYMDYLAALAKAPFAAHGYGAFLTLSILDRYYTPTISRERAVELLRKCLEELQKRFILNLPTFSVRIIDKNGIHDLDNISFPKQGS

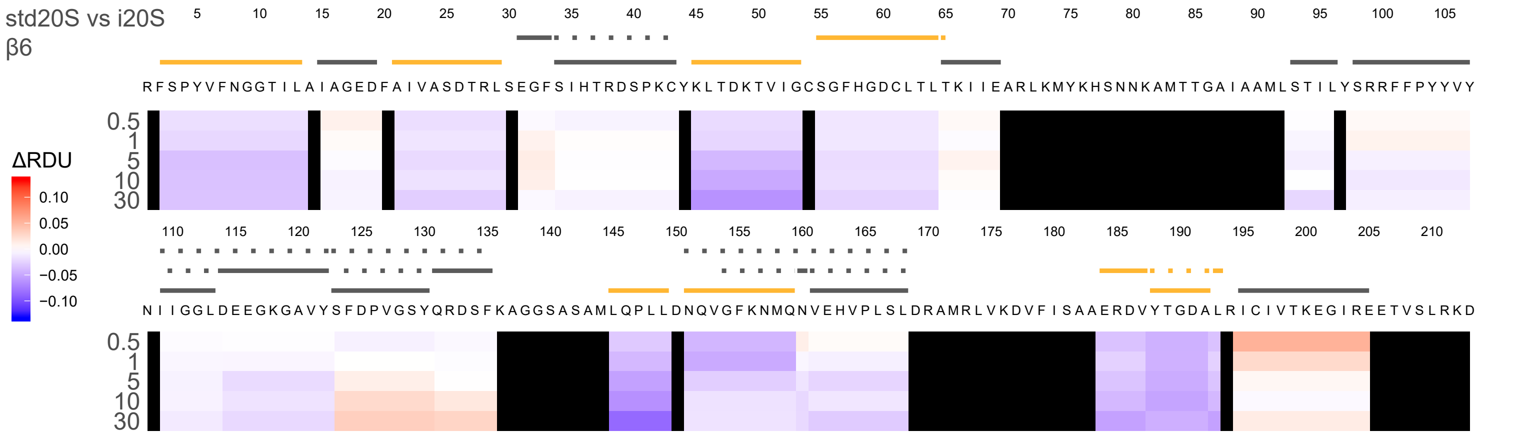
std20S vs i20S
β6
5 10 15 20 25 30 35 40 45 50 55 60 65 70 75 80 85 90 95 100 105
RFSPYVFNGGTILAIAGEDFAIVASDTRLSEGFSIHTRDSPKCYKLTDKTVIGCSGFHGDCLTLTKIIEARLKMYKHSNNKAMTTGAIAAMLSTILYSRRFFPYYVY
0.5
1
5
10
30
ΔRDU
0.10
0.05
0.00
−0.05
−0.10
110 115 120 125 130 135 140 145 150 155 160 165 170 175 180 185 190 195 200 205 210
NIIGGLDEEGKGAVYSFDPVGSYQRDSFKAGGSASAMLQPLLDNQVGFKNMQNVEHVPLSLDRAMRLVKDVFISAAERDVYTGDALRICIVTKEGIREETVSLRKD
0.5
1
5
10
30

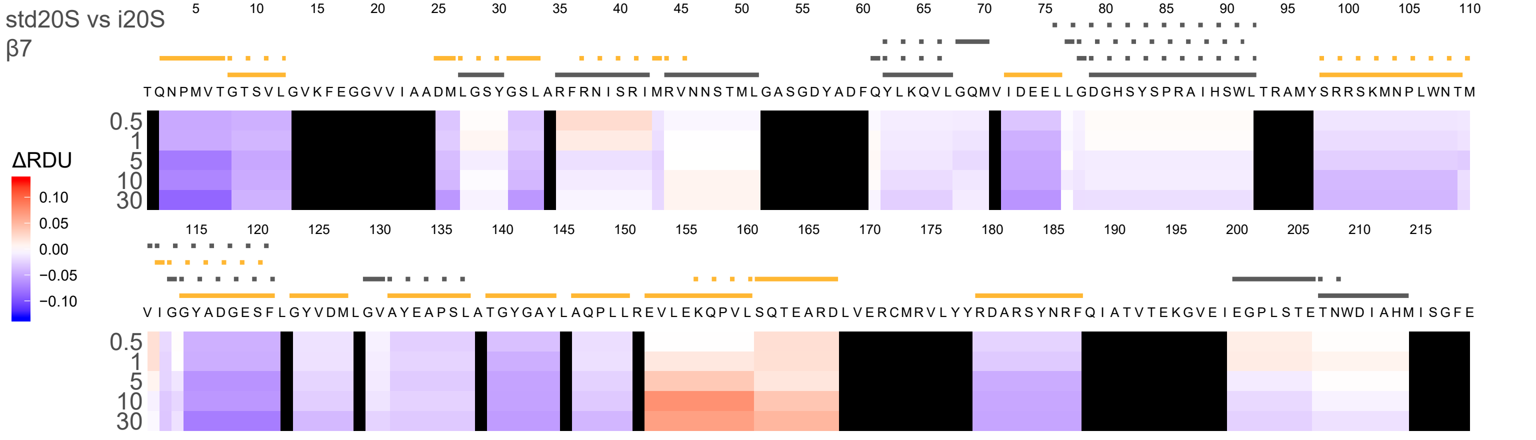

std20S vs i20S
β7
ΔRDU
0.10
0.05
0.00
−0.05
−0.10
0.5
1
5
10
30
TQNPMVTGTSVLGVKFEGGVVIAADMLGSYGSLARFRNISRIMRVNNSTMLGASGDYADFQYLKQVLGQMVIDEELLGDGHSYSPRAIHSWLTRAMYSRRSKMNPLWNTM
VIGGYADGESFLGYVDMLGVAYEAPSLATGYGAYLAQPLLREVLEKQPVLSQTEARDLVERCMRVLYYRDARSYNRFQIATVTEKGVEIEGPLSTETNWDIAHMISGFE
