## Supplementary material for "Conformational maps of human 20S proteasomes reveal PA28- and immuno-dependent inter-ring crosstalks": Fig. S5

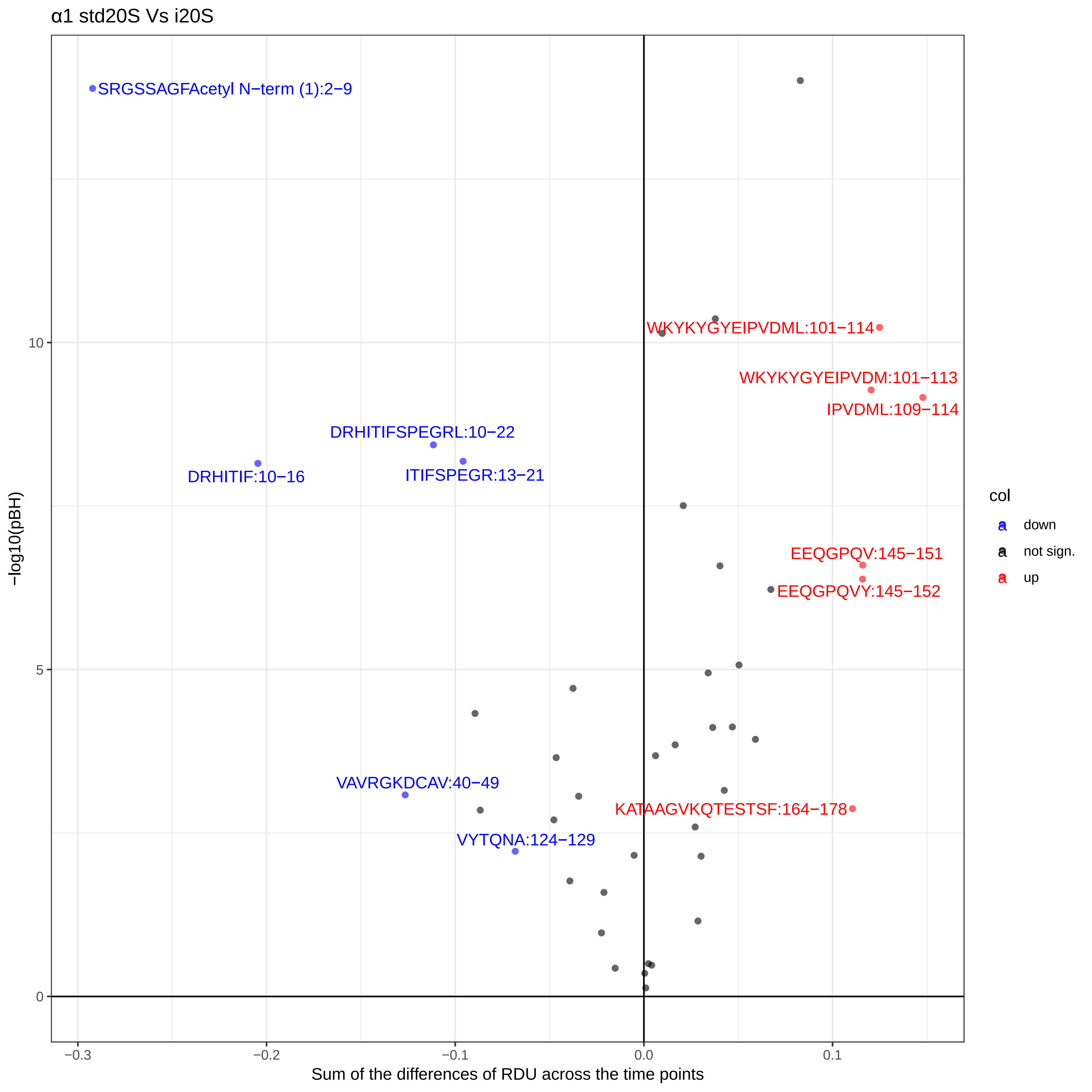

α1 std20S Vs i20S
SRGSSAGFAcetyl N−term (1):2−9
WKYKYGYEIPVDML:101−114
WKYKYGYEIPVDM:101−113
IPVDML:109−114
DRHITIFSPEGRL:10−22
DRHITIF:10−16
ITIFSPEGR:13−21
EEQGPQV:145−151
EEQGPQVY:145−152
VAVRGKDCAV:40−49
KATAAGVKQTESTSF:164−178
VYTQNA:124−129
col
down
not sign.
up
−log10(pBH)
10
5
0
−0.3
−0.2
−0.1
0.0
0.1
Sum of the differences of RDU across the time points

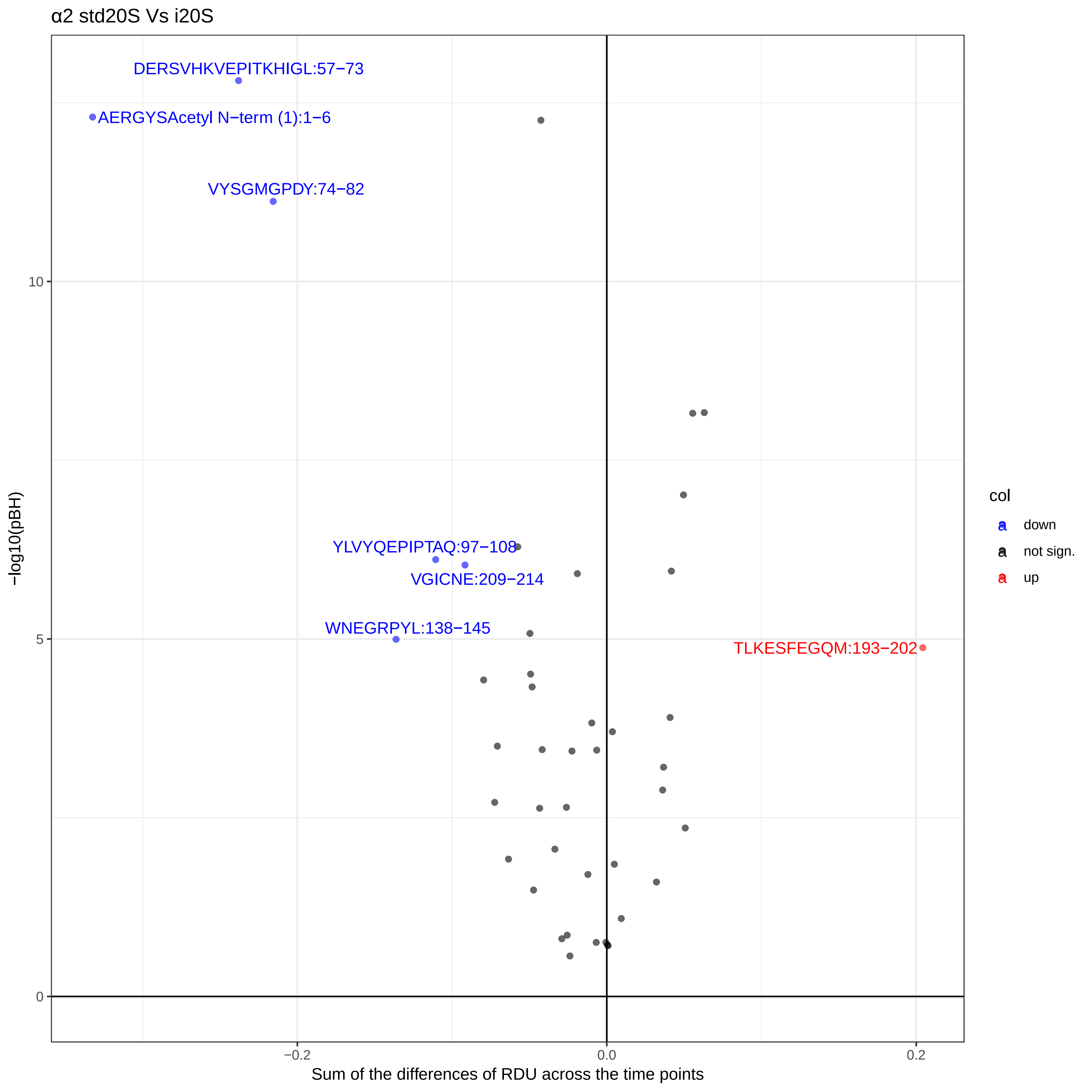

α2 std20S Vs i20S
DERSVHKVEPITKHIGL:57−73
AERGYSAcetyl N−term (1):1−6
VYSGMGPDY:74−82
YLVYQEPIPTAQ:97−108
VGICNE:209−214
WNEGRPYL:138−145
TLKESFEGQM:193−202
−log10(pBH)
10
5
0
−0.2
0.0
0.2
Sum of the differences of RDU across the time points
col
down
not sign.
up

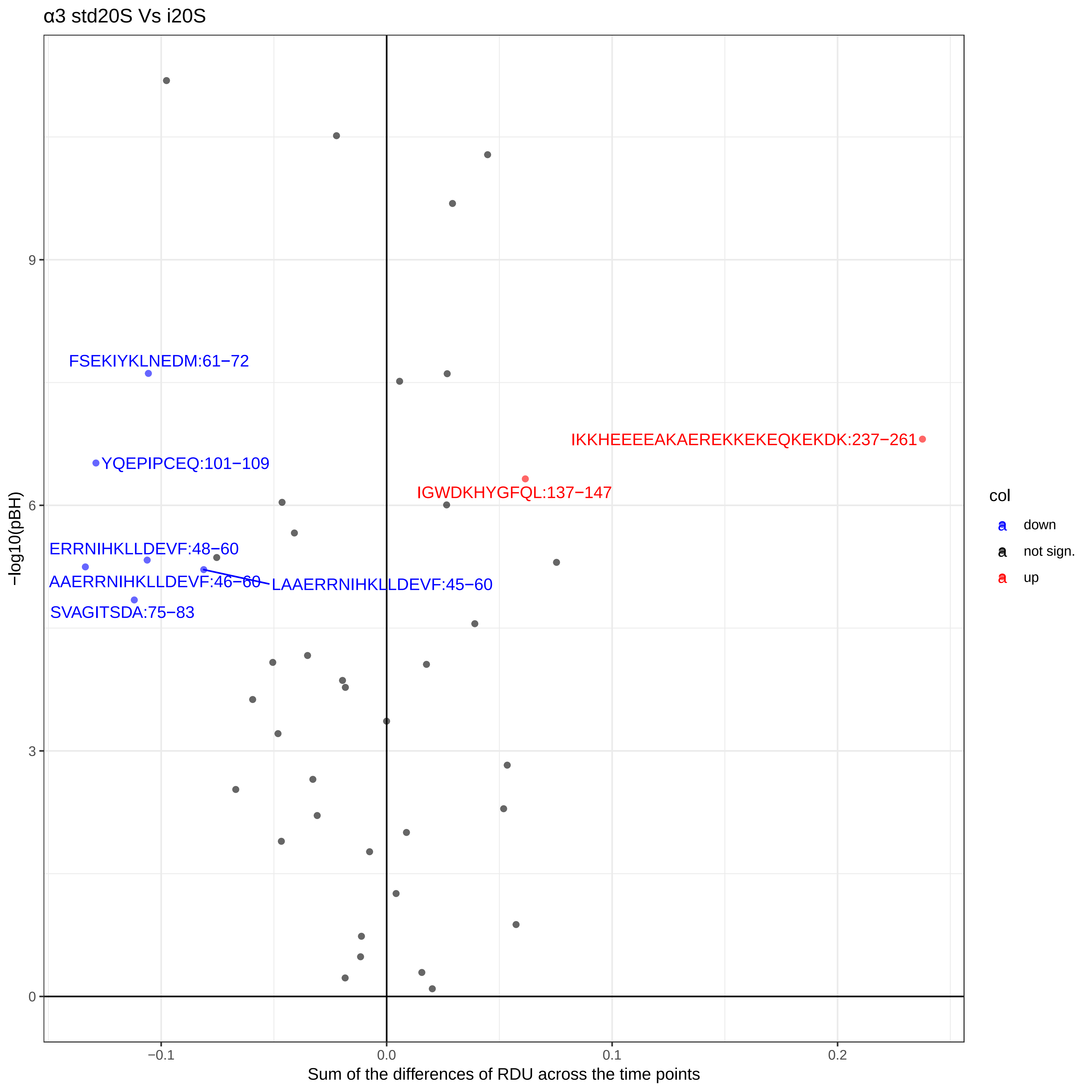
α3 std20S Vs i20S
FSEKIYKLNEDM:61−72
YQEPIPCEQ:101−109
ERRNIHKLLDEVF:48−60
AAERRNIHKLLDEVF:46−60
LAAERRNIHKLLDEVF:45−60
SVAGITSDA:75−83
IKKHEEEEAKAEREKKEKEQKEKDK:237−261
IGWDKHYGFQL:137−147
−log10(pBH)
9
6
3
0
−0.1
0.0
0.1
0.2
Sum of the differences of RDU across the time points
col
down
not sign.
up

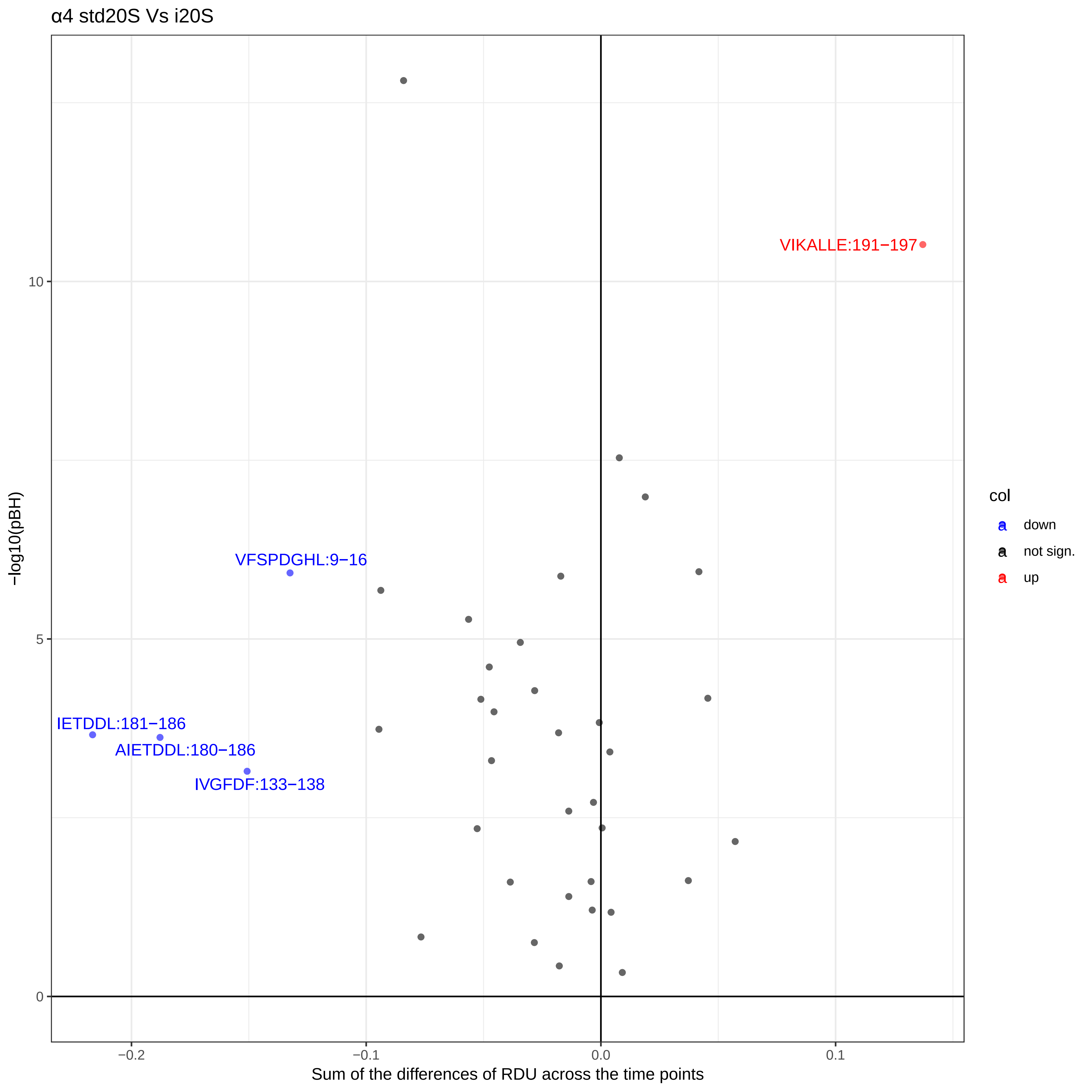
α4 std20S Vs i20S
VIKALLE:191−197
VFSPDGHL:9−16
IETDDL:181−186
AIETDDL:180−186
IVGFDF:133−138
−log10(pBH)
10
5
0
−0.2
−0.1
0.0
0.1
Sum of the differences of RDU across the time points
col
down
not sign.
up

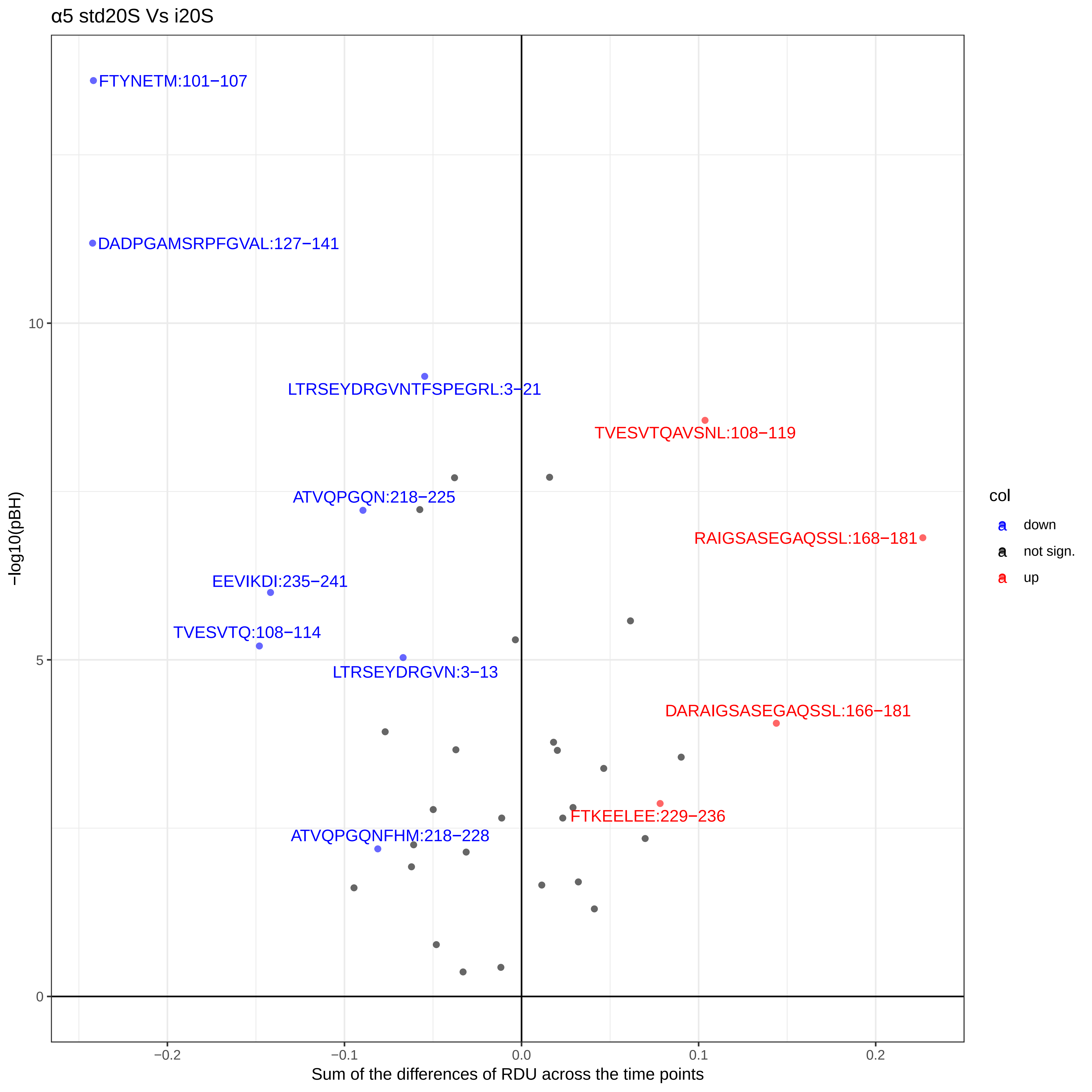

α5 std20S Vs i20S
FTYNETM:101−107
DADPGAMSRPFGVAL:127−141
LTRSEYDRGVNTFSPEGRL:3−21
TVESVTQAVSNL:108−119
ATVQPGQN:218−225
RAIGSASEGAQSSL:168−181
EEVIKDI:235−241
TVESVTQ:108−114
LTRSEYDRGVN:3−13
DARAIGSASEGAQSSL:166−181
FTKEELEE:229−236
ATVQPGQNFHM:218−228
−log10(pBH)
10
5
0
−0.2
−0.1
0.0
0.1
0.2
Sum of the differences of RDU across the time points
col
down
not sign.
up

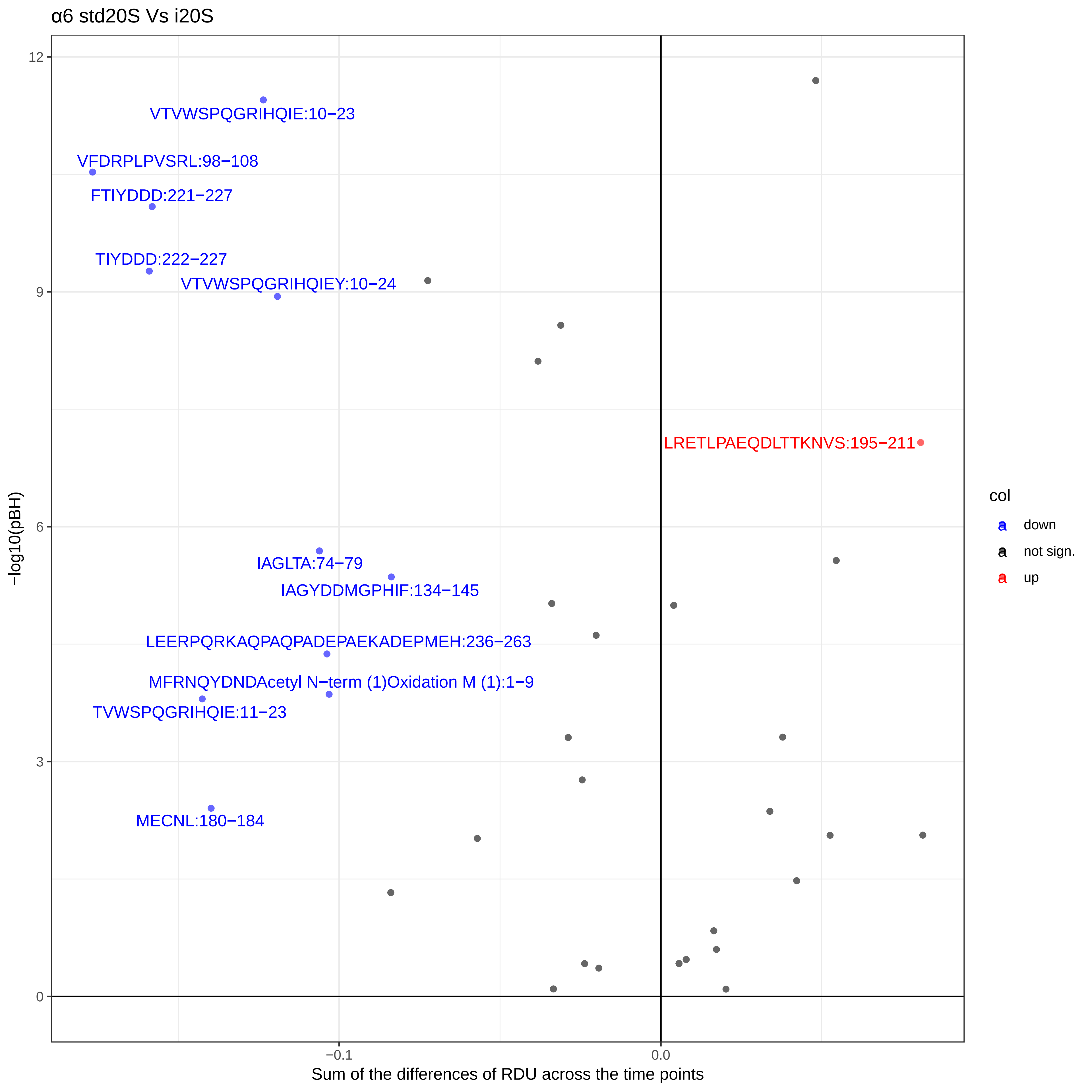
α6 std20S Vs i20S
VTVWSPQGRIHQIE:10−23
VFDRPLPVSRL:98−108
FTIYDDD:221−227
TIYDDD:222−227
VTVWSPQGRIHQIEY:10−24
LRETLPAEQDLTTKNVS:195−211
IAGLTA:74−79
IAGYDDMGPHIF:134−145
LEERPQRKAQPAQPADEPAEKADEPMEH:236−263
MFRNQYDNDAcetyl N−term (1)Oxidation M (1):1−9
TVWSPQGRIHQIE:11−23
MECNL:180−184
−log10(pBH)
12
9
6
3
0
−0.1
0.0
Sum of the differences of RDU across the time points
col
down
not sign.
up

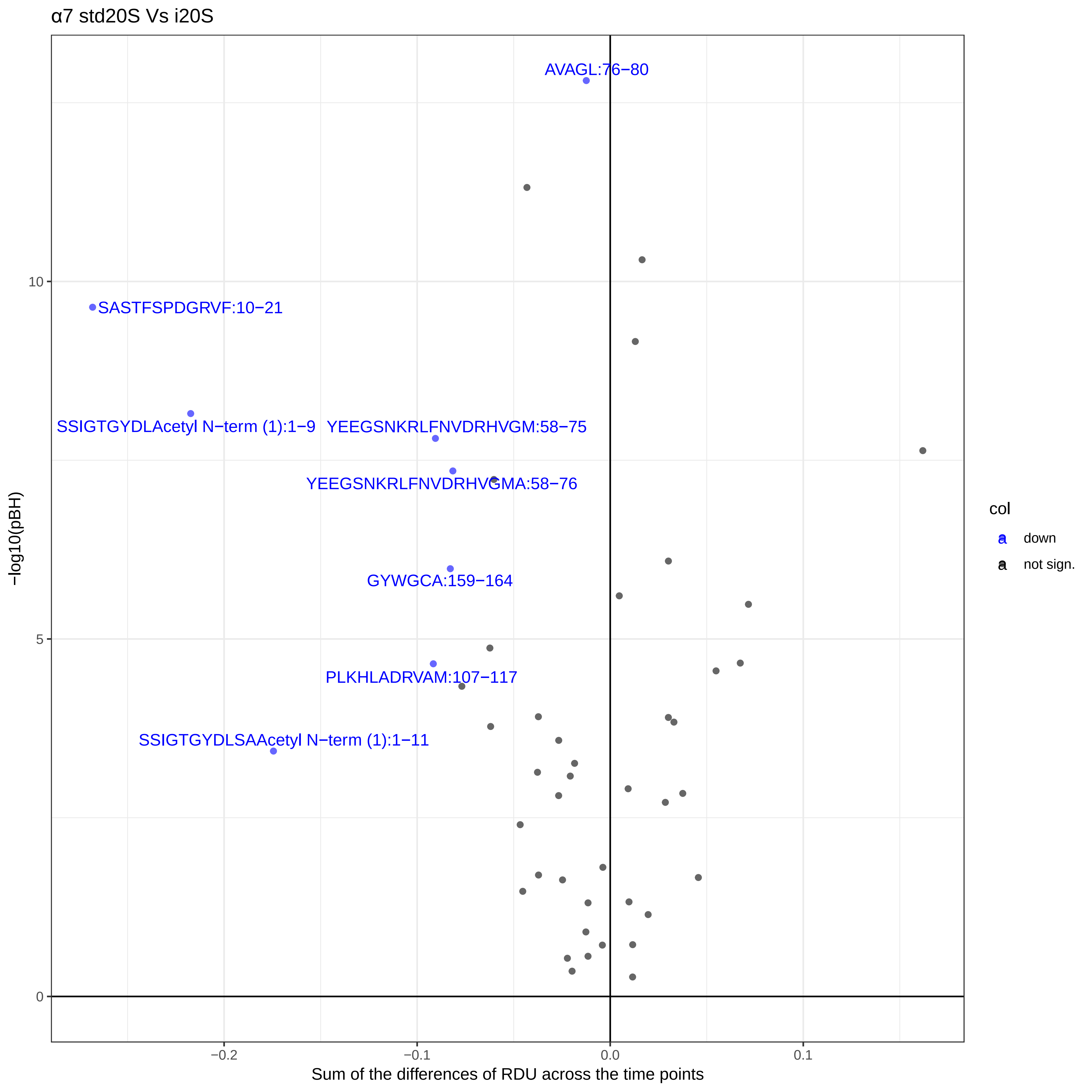
α7 std20S Vs i20S
AVAGL:76−80
SASTFSPDGRVF:10−21
SSIGTGYDLAcetyl N−term (1):1−9
YEEGSNKRLFNVDRHVGM:58−75
YEEGSNKRLFNVDRHVGMA:58−76
GYWGCA:159−164
PLKHLADRVAM:107−117
SSIGTGYDLSAAcetyl N−term (1):1−11
−log10(pBH)
10
5
0
−0.2
−0.1
0.0
0.1
Sum of the differences of RDU across the time points
col
down
not sign.

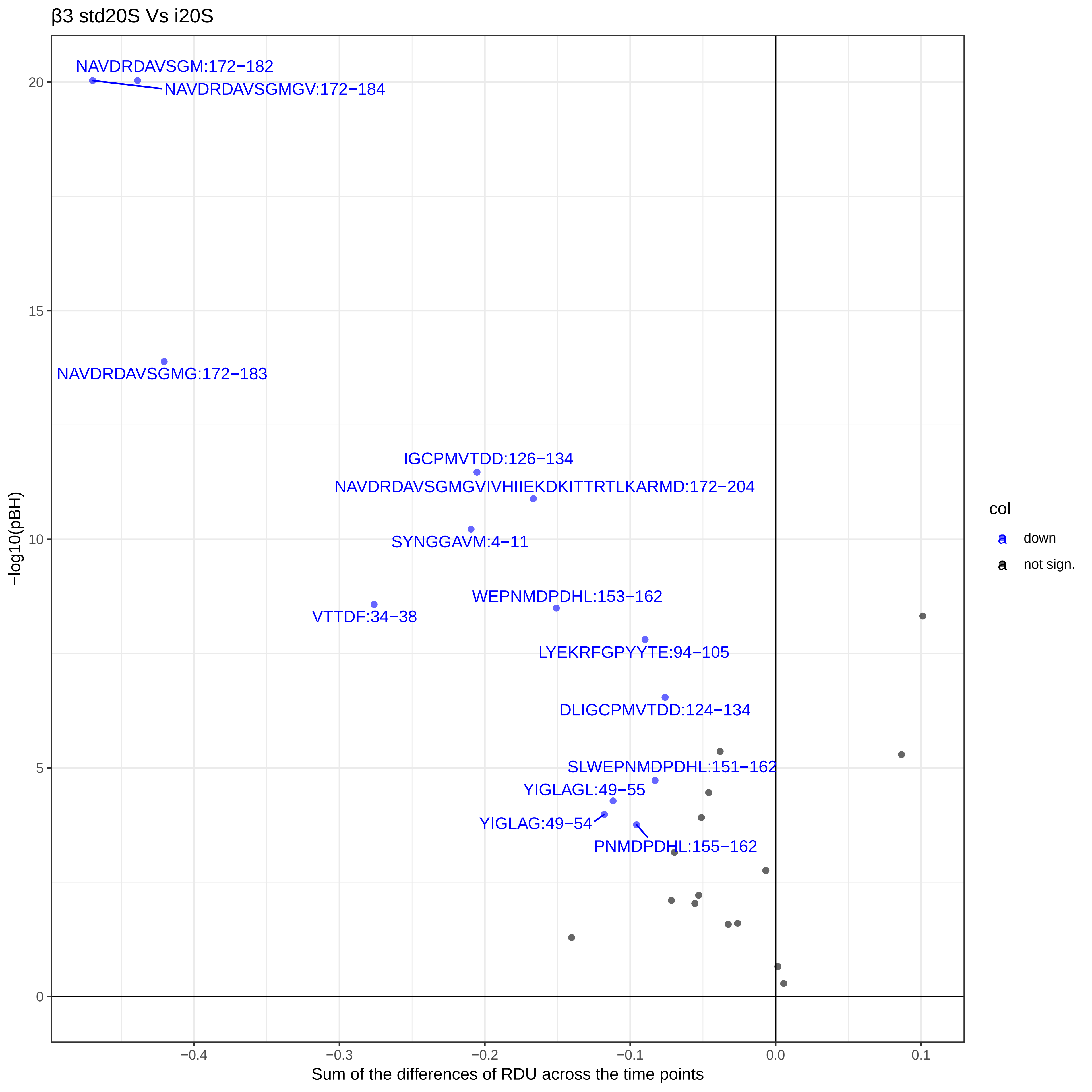
β3 std20S Vs i20S
NAVDRDAVSGM:172−182
NAVDRDAVSGMGV:172−184
NAVDRDAVSGMG:172−183
IGCPMVTDD:126−134
NAVDRDAVSGMGVIVHIIEKDKITTRTLKARMD:172−204
SYNGGAVM:4−11
WEPNMDPDHL:153−162
VTTDF:34−38
LYEKRFGPYYTE:94−105
DLIGCPMVTDD:124−134
SLWEPNMDPDHL:151−162
YIGLAGL:49−55
YIGLAG:49−54
PNMDPDHL:155−162
col
down
not sign.
−log10(pBH)
20
15
10
5
0
−0.4
−0.3
−0.2
−0.1
0.0
0.1
Sum of the differences of RDU across the time points

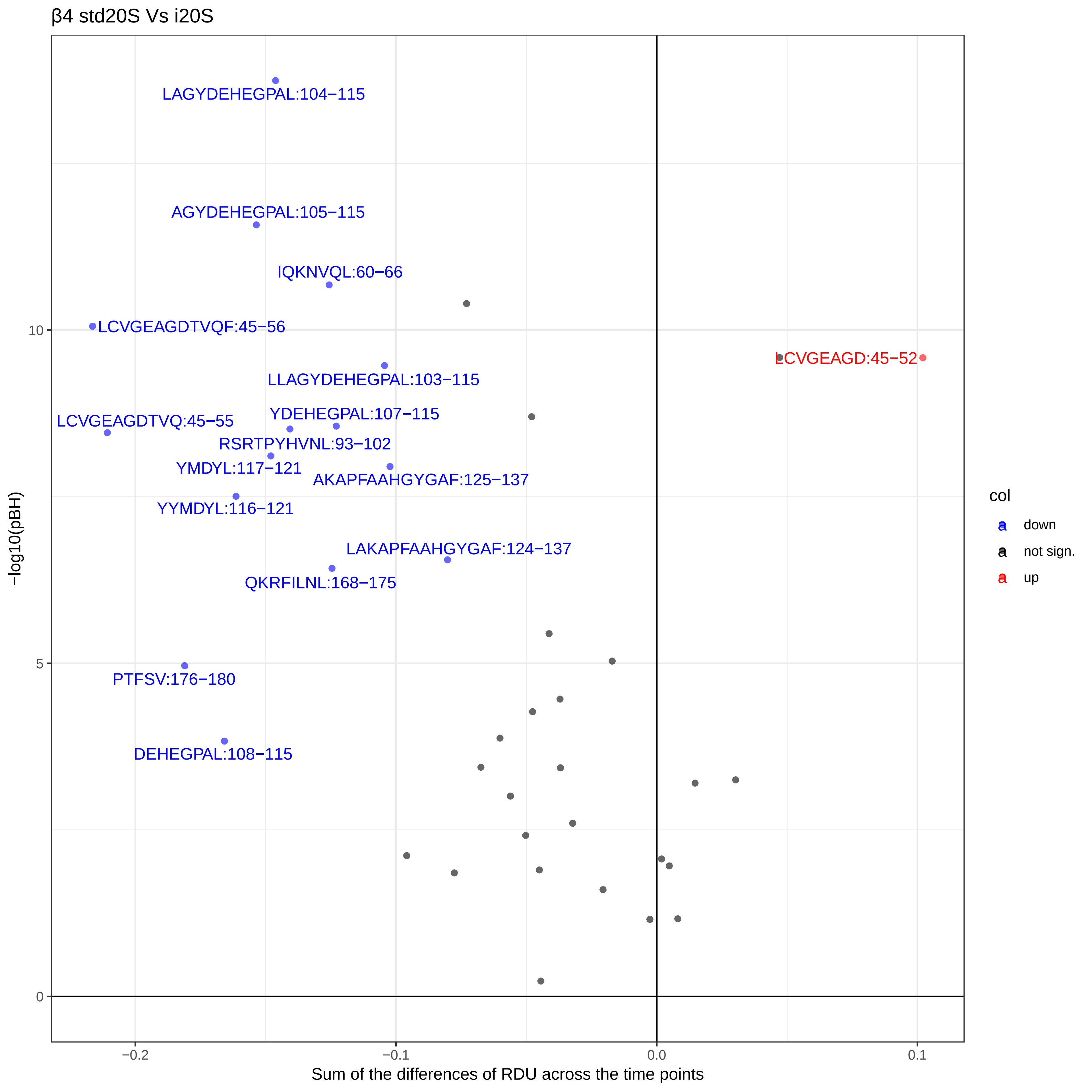

β4 std20S Vs i20S
LAGYDEHEGPAL:104−115
AGYDEHEGPAL:105−115
IQKNVQL:60−66
LCVGEAGDTVQF:45−56
LCVGEAGD:45−52
LLAGYDEHEGPAL:103−115
YDEHEGPAL:107−115
LCVGEAGDTVQ:45−55
RSRTPYHVNL:93−102
YMDYL:117−121
AKAPFAAHGYGAF:125−137
YYMDYL:116−121
LAKAPFAAHGYGAF:124−137
QKRFILNL:168−175
PTFSV:176−180
DEHEGPAL:108−115
−log10(pBH)
Sum of the differences of RDU across the time points
col
down
not sign.
up

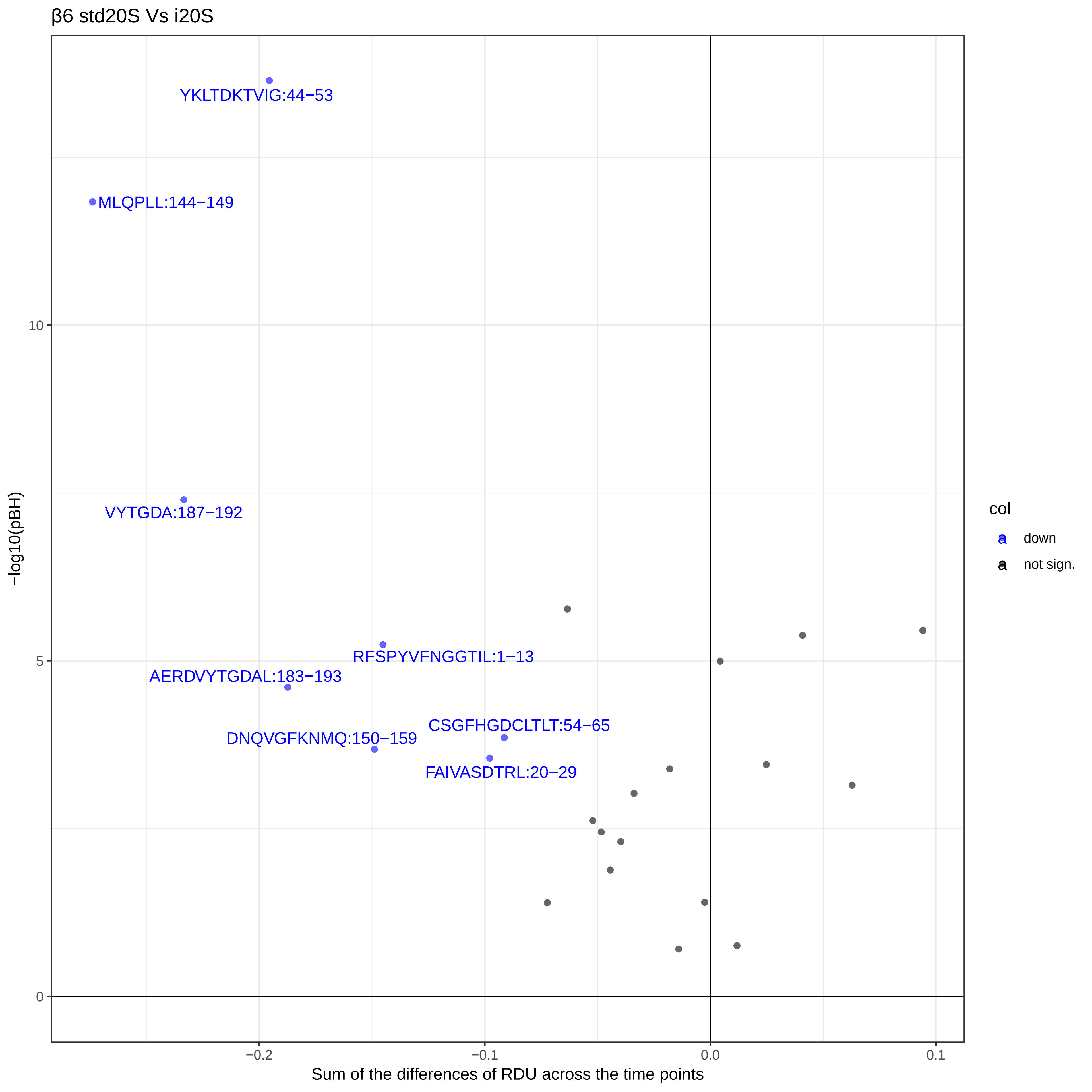

β6 std20S Vs i20S
YKLTDKTVIG:44−53
MLQPLL:144−149
VYTGDA:187−192
RFSPYVFNGGTIL:1−13
AERDVYTGDAL:183−193
CSGFHGDCLTLT:54−65
DNQVGFKNMQ:150−159
FAIVASDTRL:20−29
−log10(pBH)
10
5
0
−0.2
−0.1
0.0
0.1
Sum of the differences of RDU across the time points
col
down
not sign.

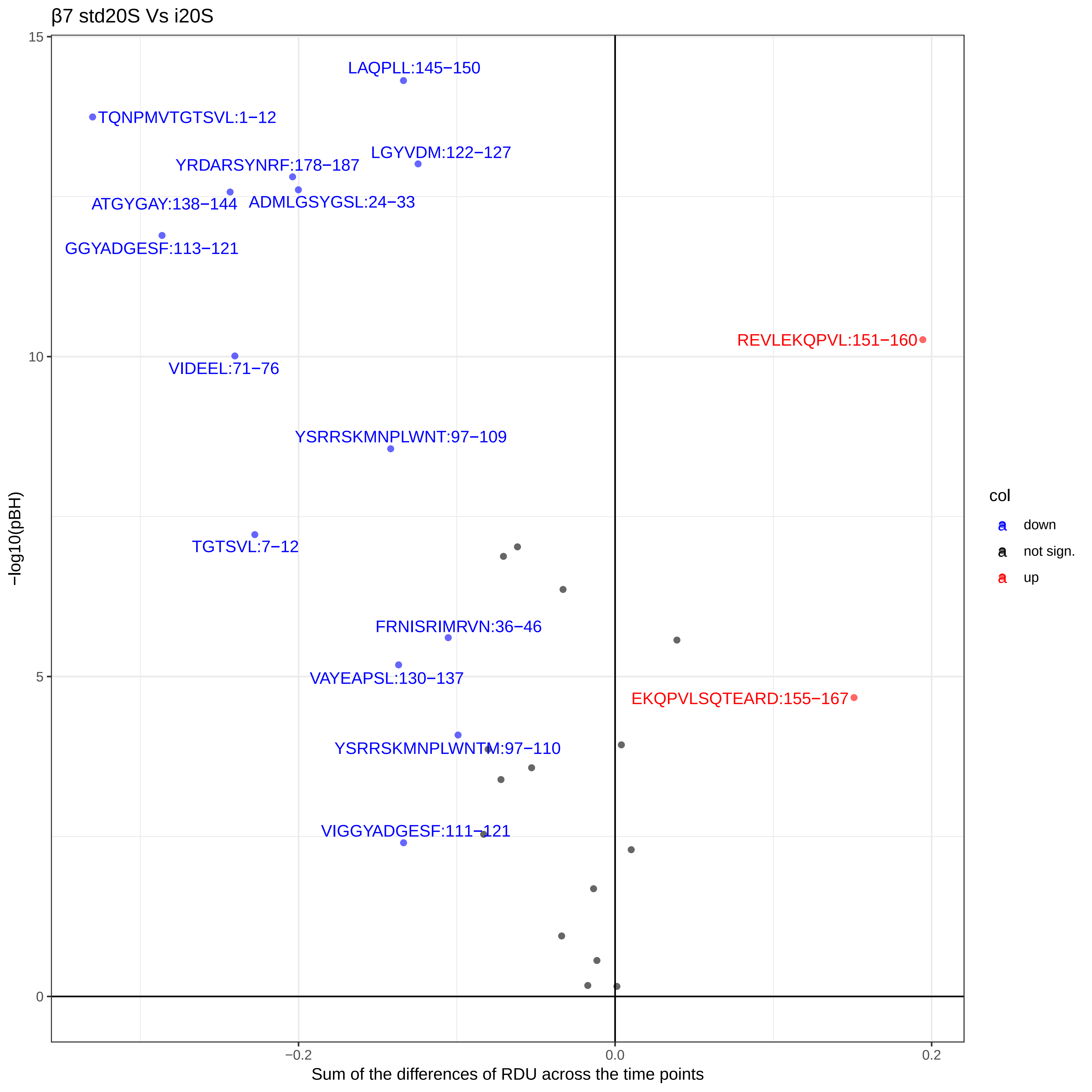

β7 std20S Vs i20S
LAQPLL:145−150
TQNPMVTGTSVL:1−12
LGYVDM:122−127
YRDARSYNRF:178−187
ATGYGAY:138−144
ADMLGSYGSL:24−33
GGYADGESF:113−121
REVLEKQPVL:151−160
VIDEEL:71−76
YSRRSKMNPLWNT:97−109
TGTSVL:7−12
FRNISRIMRVN:36−46
VAYEAPSL:130−137
EKQPVLSQTEARD:155−167
YSRRSKMNPLWNTM:97−110
VIGGYADGESF:111−121
15
10
5
0
−0.2
0.0
0.2
−log10(pBH)
Sum of the differences of RDU across the time points
col
down
not sign.
up
