## Supplementary material for "Conformational maps of human 20S proteasomes reveal PA28- and immuno-dependent inter-ring crosstalks": Fig. S6

# PA28α

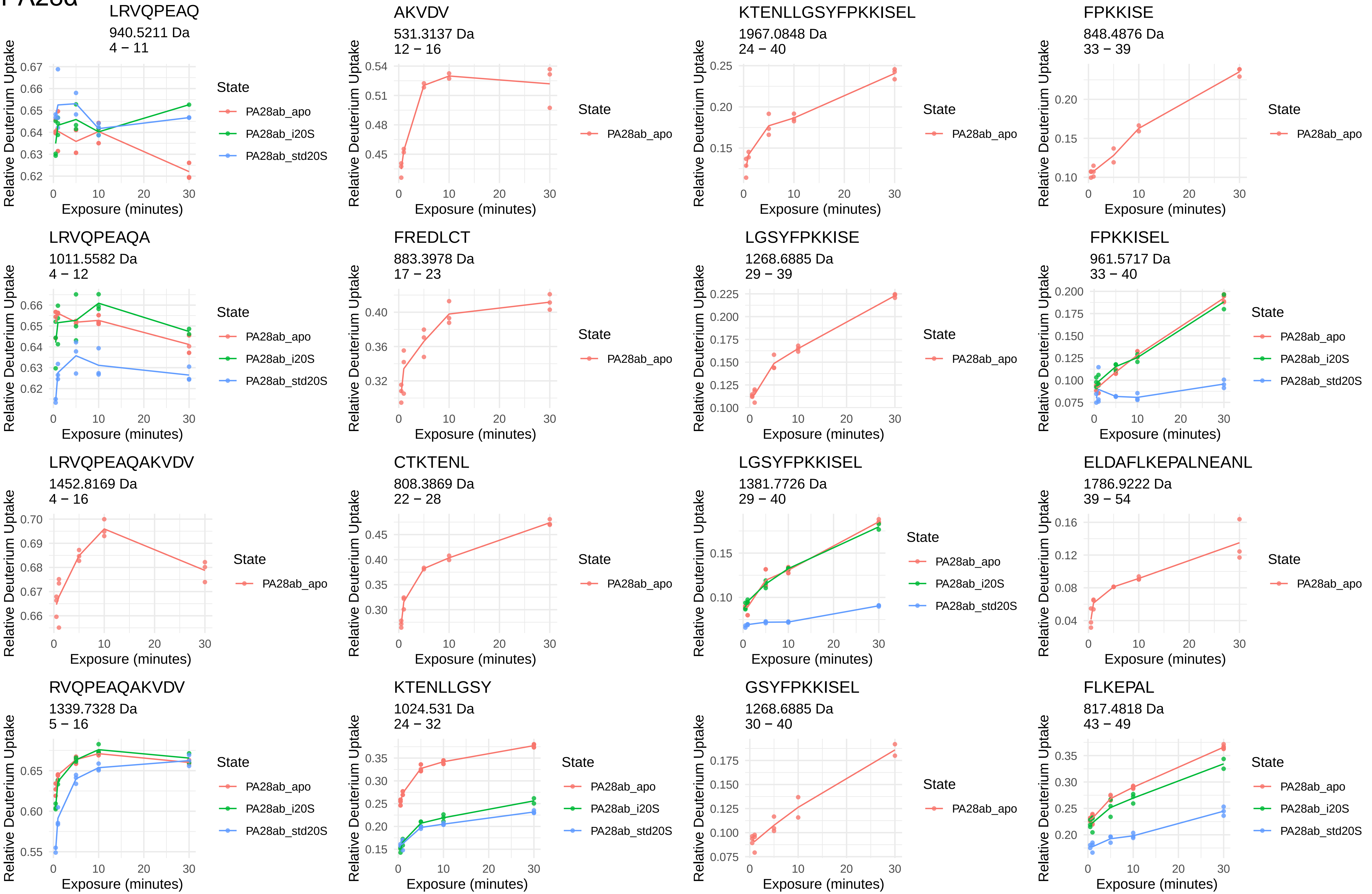
LRVQPEAQ
940.5211 Da
4 − 11
AKVDV
531.3137 Da
12 − 16
KTENLLGSYFPKKISEL
1967.0848 Da
24 − 40
FPKKISE
848.4876 Da
33 − 39
LRVQPEAQA
1011.5582 Da
4 − 12
FREDLCT
883.3978 Da
17 − 23
LGSYFPKKISE
1268.6885 Da
29 − 39
FPKKISEL
961.5717 Da
33 − 40
LRVQPEAQAKVDV
1452.8169 Da
4 − 16
CTKTENL
808.3869 Da
22 − 28
LGSYFPKKISEL
1381.7726 Da
29 − 40
ELDAFLKEPALNEANL
1786.9222 Da
39 − 54
RVQPEAQAKVDV
1339.7328 Da
5 − 16
KTENLLGSY
1024.531 Da
24 − 32
GSYFPKKISEL
1268.6885 Da
30 − 40
FLKEPAL
817.4818 Da
43 − 49
Relative Deuterium Uptake
Exposure (minutes)
State
PA28ab_apo
PA28ab_i20S
PA28ab_std20S

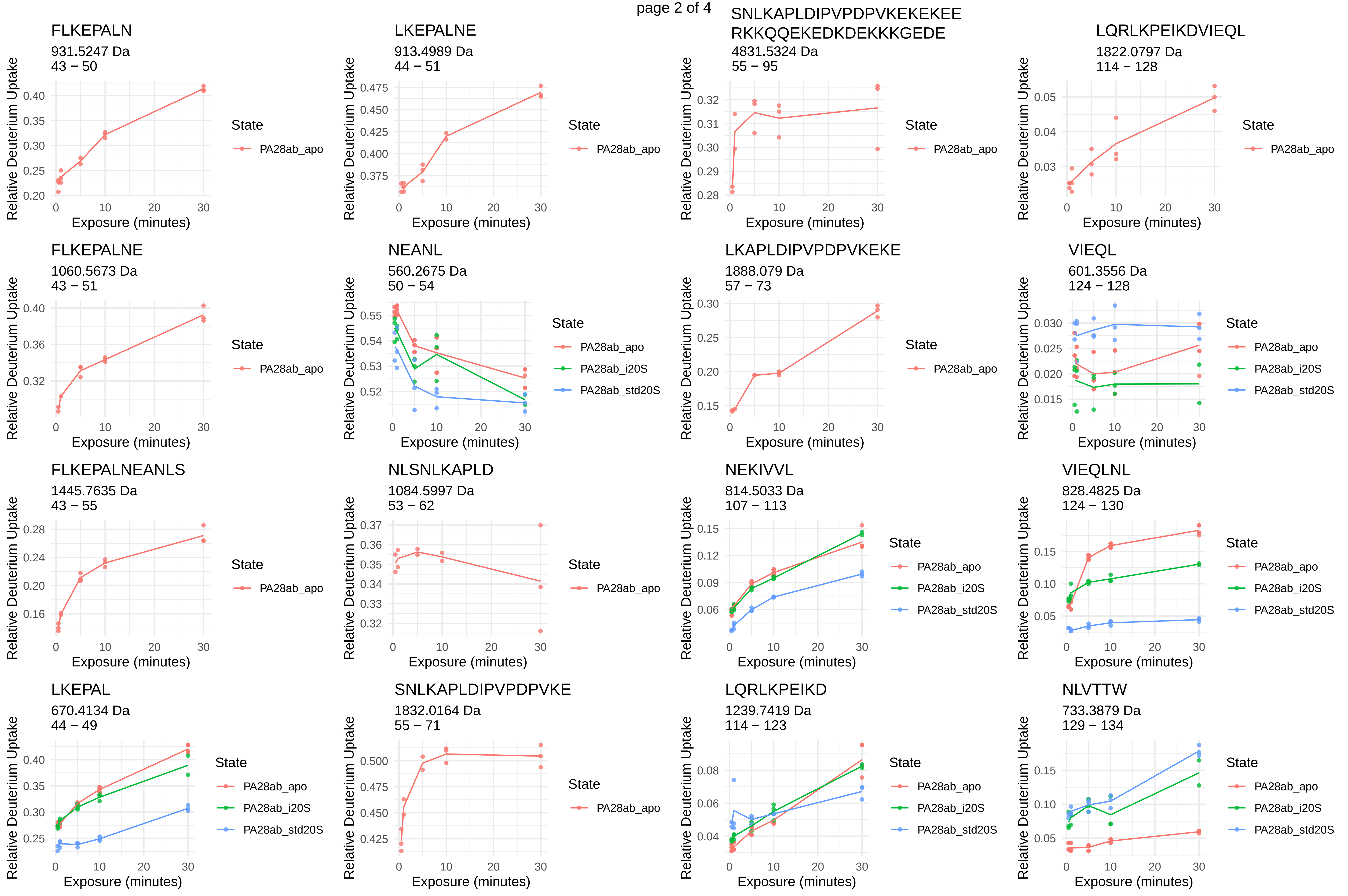
FLKEPALN
931.5247 Da
43 – 50
LKEPALNE
913.4989 Da
44 – 51
SNLKAPLDIPVPDPVKEKEKEE
RKKQQEKEDKDEKKKGEDE
4831.5324 Da
55 – 95
LQRLKPEIKDVIEQL
1822.0797 Da
114 – 128
FLKEPALNE
1060.5673 Da
43 – 51
NEANL
560.2675 Da
50 – 54
LKAPLDIPVPDPVKEKE
1888.079 Da
57 – 73
VIEQL
601.3556 Da
124 – 128
FLKEPALNEANLS
1445.7635 Da
43 – 55
NLSNLKAPLD
1084.5997 Da
53 – 62
NEKIVVL
814.5033 Da
107 – 113
VIEQLNL
828.4825 Da
124 – 130
LKEPAL
670.4134 Da
44 – 49
SNLKAPLDIPVPDPVKE
1832.0164 Da
55 – 71
LQRLKPEIKD
1239.7419 Da
114 – 123
NLVTTW
733.3879 Da
129 – 134
Relative Deuterium Uptake
Exposure (minutes)
State
PA28ab_apo
PA28ab_i20S
PA28ab_std20S

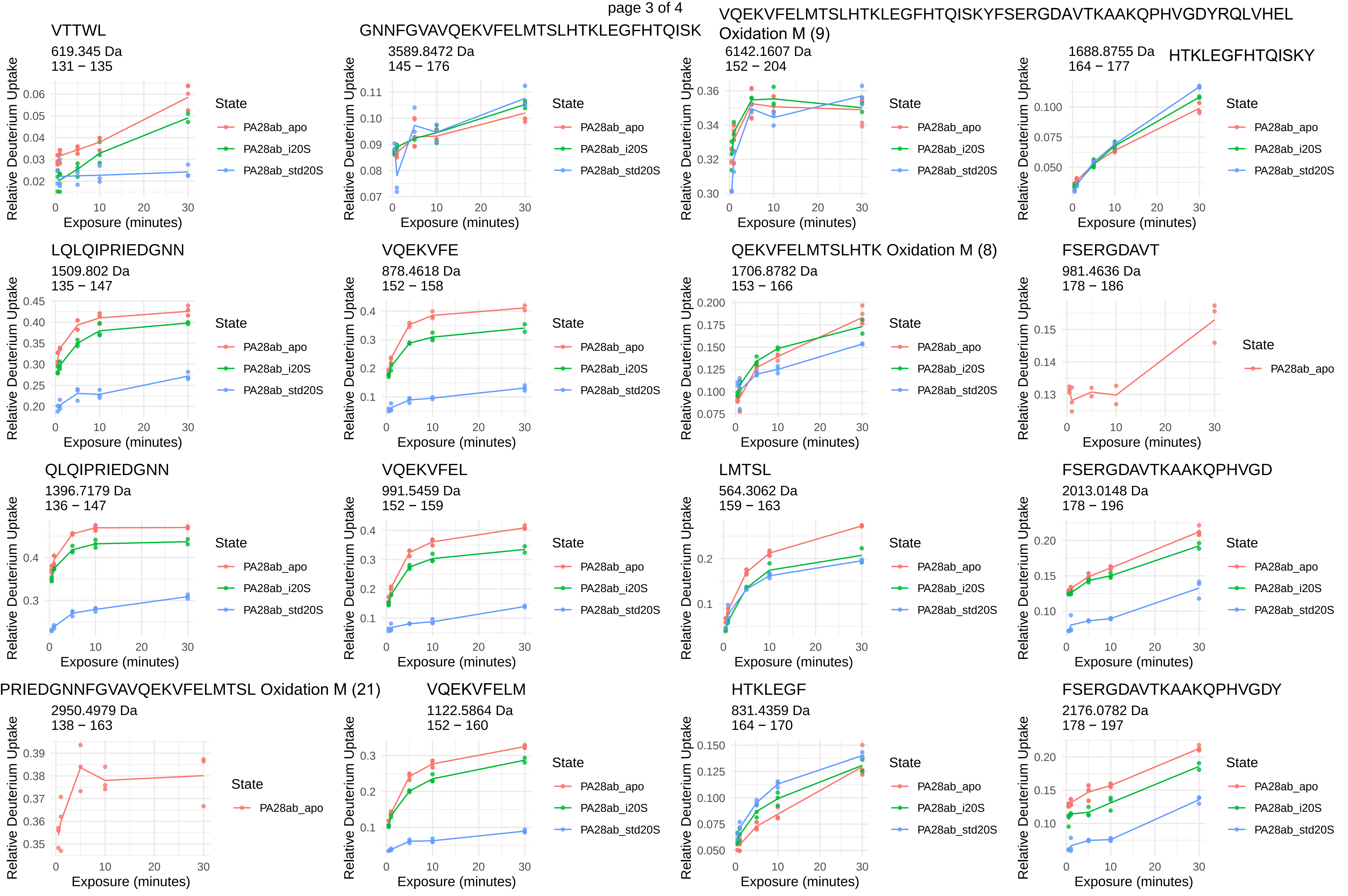
VTTWL
619.345 Da
131 − 135
GNNFGVAVQEKVFELMTSLHTKLEGFHTQISK
3589.8472 Da
145 − 176
VQEKVFELMTSLHTKLEGFHTQISKYFSERGDAVTKAAKQPHVGDYRQLVHEL
Oxidation M (9)
6142.1607 Da
152 − 204
HTKLEGFHTQISKY
1688.8755 Da
164 − 177
LQLQIPRIEDGNN
1509.802 Da
135 − 147
VQEKVFE
878.4618 Da
152 − 158
QEKVFELMTSLHTK Oxidation M (8)
1706.8782 Da
153 − 166
FSERGDAVT
981.4636 Da
178 − 186
QLQIPRIEDGNN
1396.7179 Da
136 − 147
VQEKVFEL
991.5459 Da
152 − 159
LMTSL
564.3062 Da
159 − 163
FSERGDAVTKAAKQPHVGD
2013.0148 Da
178 − 196
PRIEDGNNFGVAVQEKVFELMTSL Oxidation M (21)
2950.4979 Da
138 − 163
VQEKVFELM
1122.5864 Da
152 − 160
HTKLEGF
831.4359 Da
164 − 170
FSERGDAVTKAAKQPHVGDY
2176.0782 Da
178 − 197
Relative Deuterium Uptake
Exposure (minutes)
State
PA28ab_apo
PA28ab_i20S
PA28ab_std20S

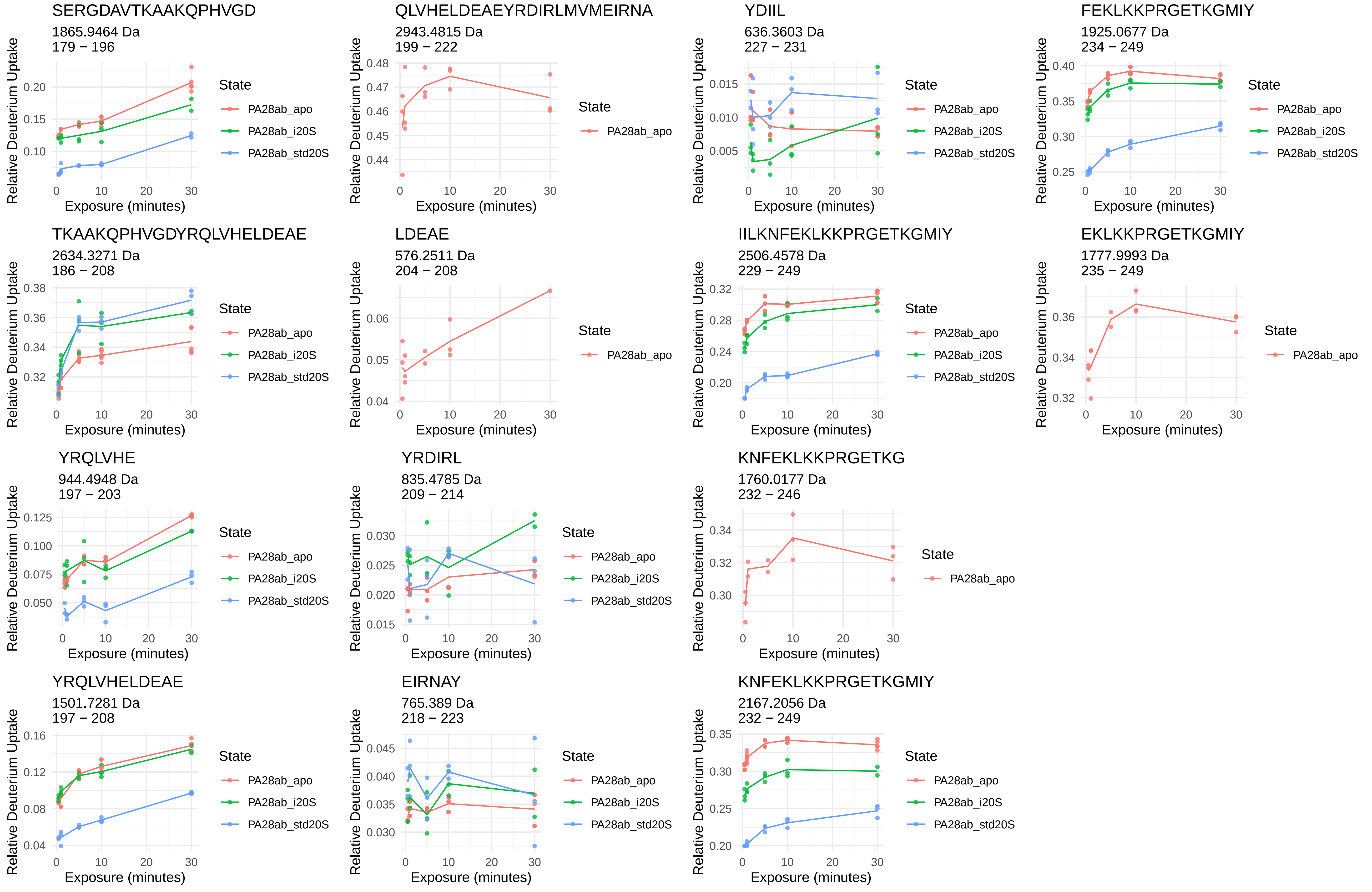
SERGDAVTKAAKQPHVGD
1865.9464 Da
179 − 196
QLVHELDEAEYRDIRLMVMEIRNA
2943.4815 Da
199 − 222
YDIIL
636.3603 Da
227 − 231
FEKLKKPRGETKGMIY
1925.0677 Da
234 − 249
TKAAKQPHVGDYRQLVHELDEAE
2634.3271 Da
186 − 208
LDEAE
576.2511 Da
204 − 208
IILKNFEKLKKPRGETKGMIY
2506.4578 Da
229 − 249
EKLKKPRGETKGMIY
1777.9993 Da
235 − 249
YRQLVHE
944.4948 Da
197 − 203
YRDIRL
835.4785 Da
209 − 214
KNFEKLKKPRGETKG
1760.0177 Da
232 − 246
YRQLVHELDEAE
1501.7281 Da
197 − 208
EIRNAY
765.389 Da
218 − 223
KNFEKLKKPRGETKGMIY
2167.2056 Da
232 − 249
Relative Deuterium Uptake
Exposure (minutes)
State
PA28ab_apo
PA28ab_i20S
PA28ab_std20S

PA28β

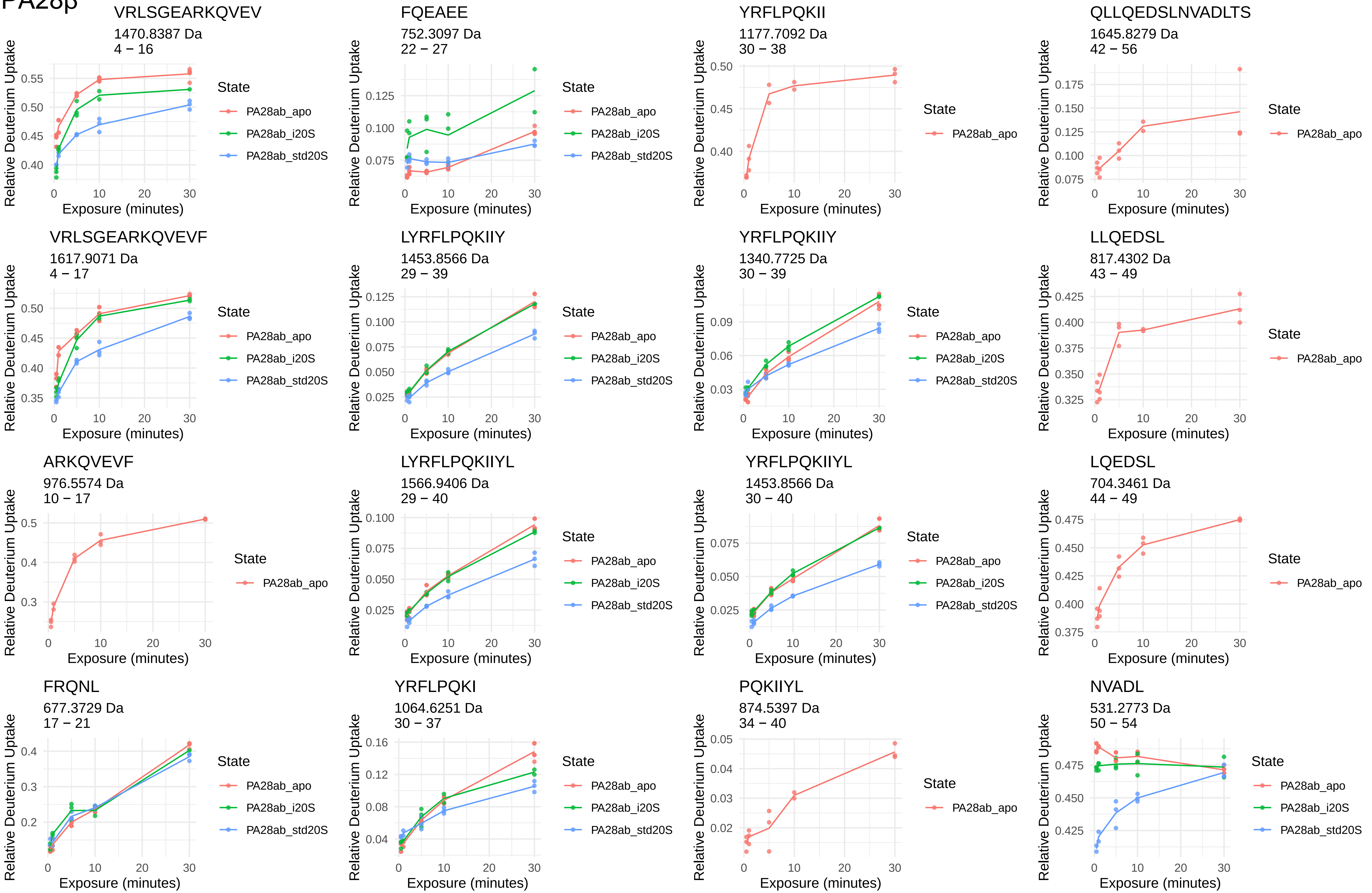
VRLSGEARKQVEV
1470.8387 Da
4 – 16
FQEAEE
752.3097 Da
22 – 27
YRFLPQKII
1177.7092 Da
30 – 38
QLLQEDSLNVADLTS
1645.8279 Da
42 – 56
VRLSGEARKQVEVF
1617.9071 Da
4 – 17
LYRFLPQKIIY
1453.8566 Da
29 – 39
YRFLPQKIIY
1340.7725 Da
30 – 39
LLQEDSL
817.4302 Da
43 – 49
ARKQVEVF
976.5574 Da
10 – 17
LYRFLPQKIIYL
1566.9406 Da
29 – 40
YRFLPQKIIYL
1453.8566 Da
30 – 40
LQEDSL
704.3461 Da
44 – 49
FRQNL
677.3729 Da
17 – 21
YRFLPQKI
1064.6251 Da
30 – 37
PQKIIYL
874.5397 Da
34 – 40
NVADL
531.2773 Da
50 – 54
Relative Deuterium Uptake
Exposure (minutes)
State
PA28ab_apo
PA28ab_i20S
PA28ab_std20S

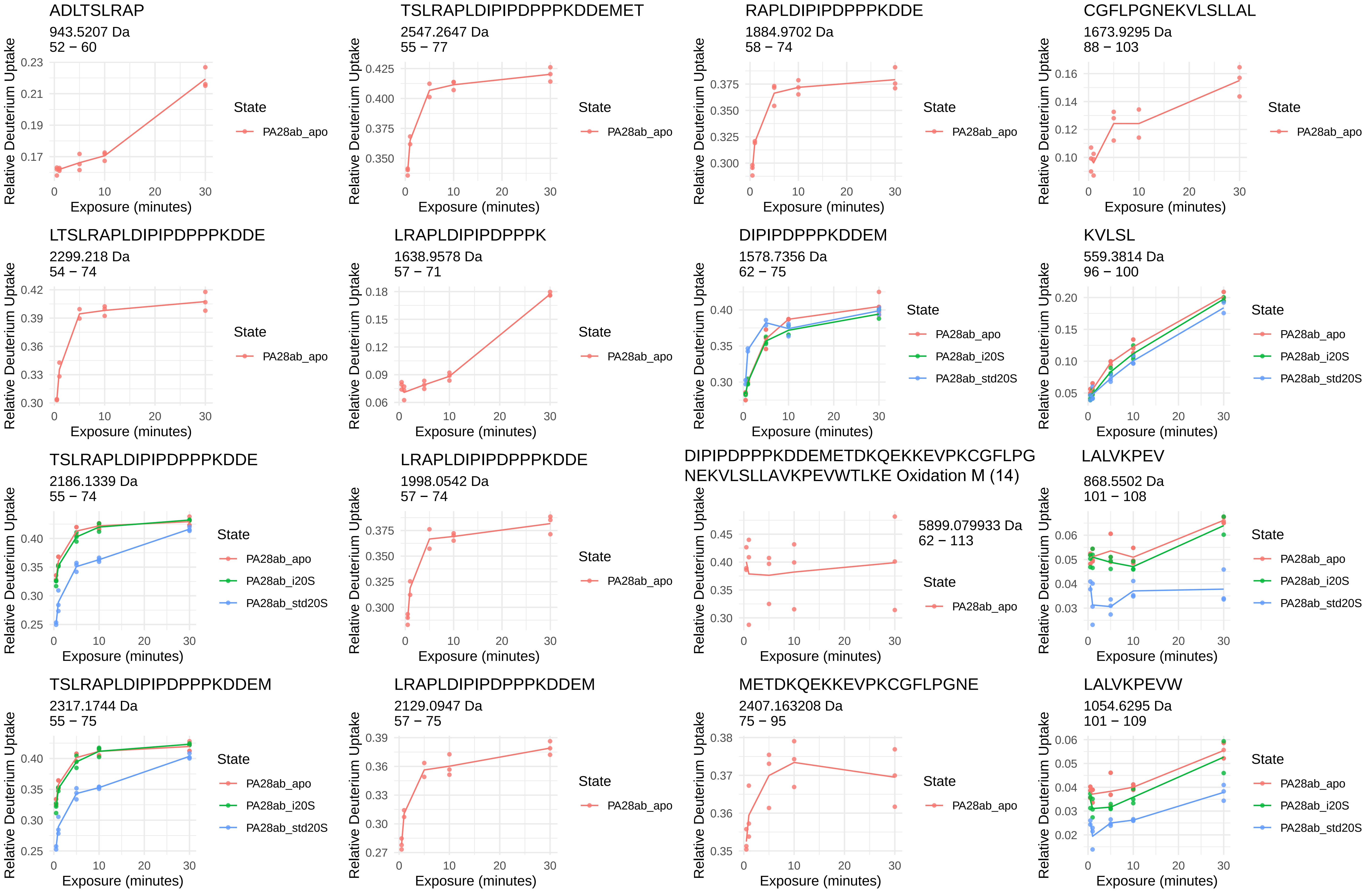

ADLTSLRAP
943.5207 Da
52 – 60
TSLRAPLDIPIPDPPPKDDEMET
2547.2647 Da
55 – 77
RAPLDIPIPDPPPKDDE
1884.9702 Da
58 – 74
CGFLPGNEKVLSLLAL
1673.9295 Da
88 – 103
LTSLRAPLDIPIPDPPPKDDE
2299.218 Da
54 – 74
LRAPLDIPIPDPPPK
1638.9578 Da
57 – 71
DIPIPDPPPKDDEM
1578.7356 Da
62 – 75
KVLSL
559.3814 Da
96 – 100
TSLRAPLDIPIPDPPPKDDE
2186.1339 Da
55 – 74
LRAPLDIPIPDPPPKDDE
1998.0542 Da
57 – 74
DIPIPDPPPKDDEMETDKQEKKEVPKCGFLPG
NEKVLSLLAVKPEVWTLKE Oxidation M (14)
5899.079933 Da
62 – 113
LALVKPEV
868.5502 Da
101 – 108
TSLRAPLDIPIPDPPPKDDEM
2317.1744 Da
55 – 75
LRAPLDIPIPDPPPKDDEM
2129.0947 Da
57 – 75
METDKQEKKEVPKCGFLPGNE
2407.163208 Da
75 – 95
LALVKPEVW
1054.6295 Da
101 – 109
Relative Deuterium Uptake
Exposure (minutes)
State
PA28ab_apo
PA28ab_i20S
PA28ab_std20S

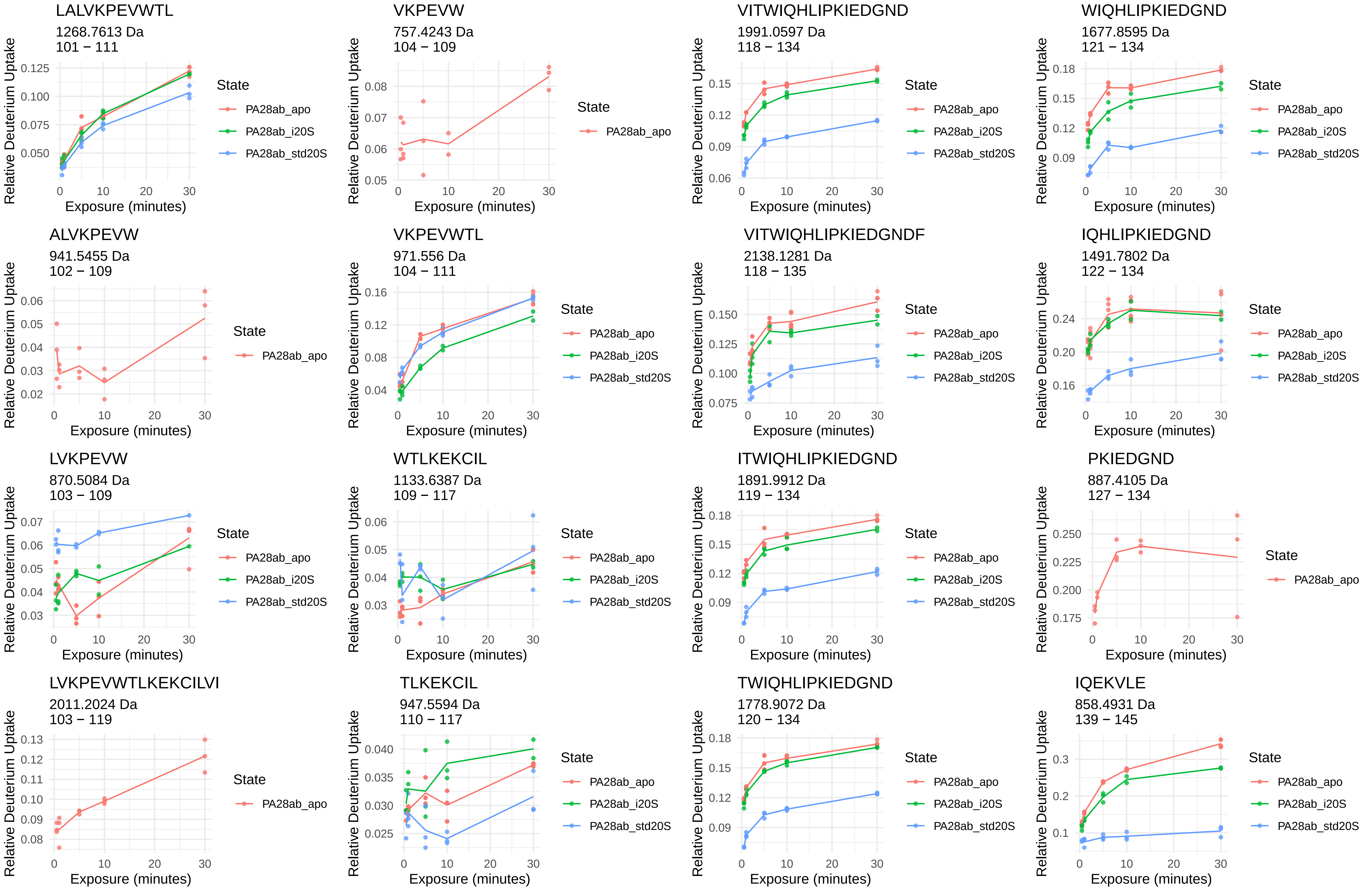
LALVKPEVWTL
1268.7613 Da
101 – 111
VKPEVW
757.4243 Da
104 – 109
VITWIQHLIPKIEDGND
1991.0597 Da
118 – 134
WIQHLIPKIEDGND
1677.8595 Da
121 – 134
ALVKPEVW
941.5455 Da
102 – 109
VKPEVWTL
971.556 Da
104 – 111
VITWIQHLIPKIEDGNDF
2138.1281 Da
118 – 135
IQHLIPKIEDGND
1491.7802 Da
122 – 134
LVKPEVW
870.5084 Da
103 – 109
WTLKEKCIL
1133.6387 Da
109 – 117
ITWIQHLIPKIEDGND
1891.9912 Da
119 – 134
PKIEDGND
887.4105 Da
127 – 134
LVKPEVWTLKEKCILVI
2011.2024 Da
103 – 119
TLKEKCIL
947.5594 Da
110 – 117
TWIQHLIPKIEDGND
1778.9072 Da
120 – 134
IQEKVLE
858.4931 Da
139 – 145
Relative Deuterium Uptake
Exposure (minutes)
State
PA28ab_apo
PA28ab_i20S
PA28ab_std20S

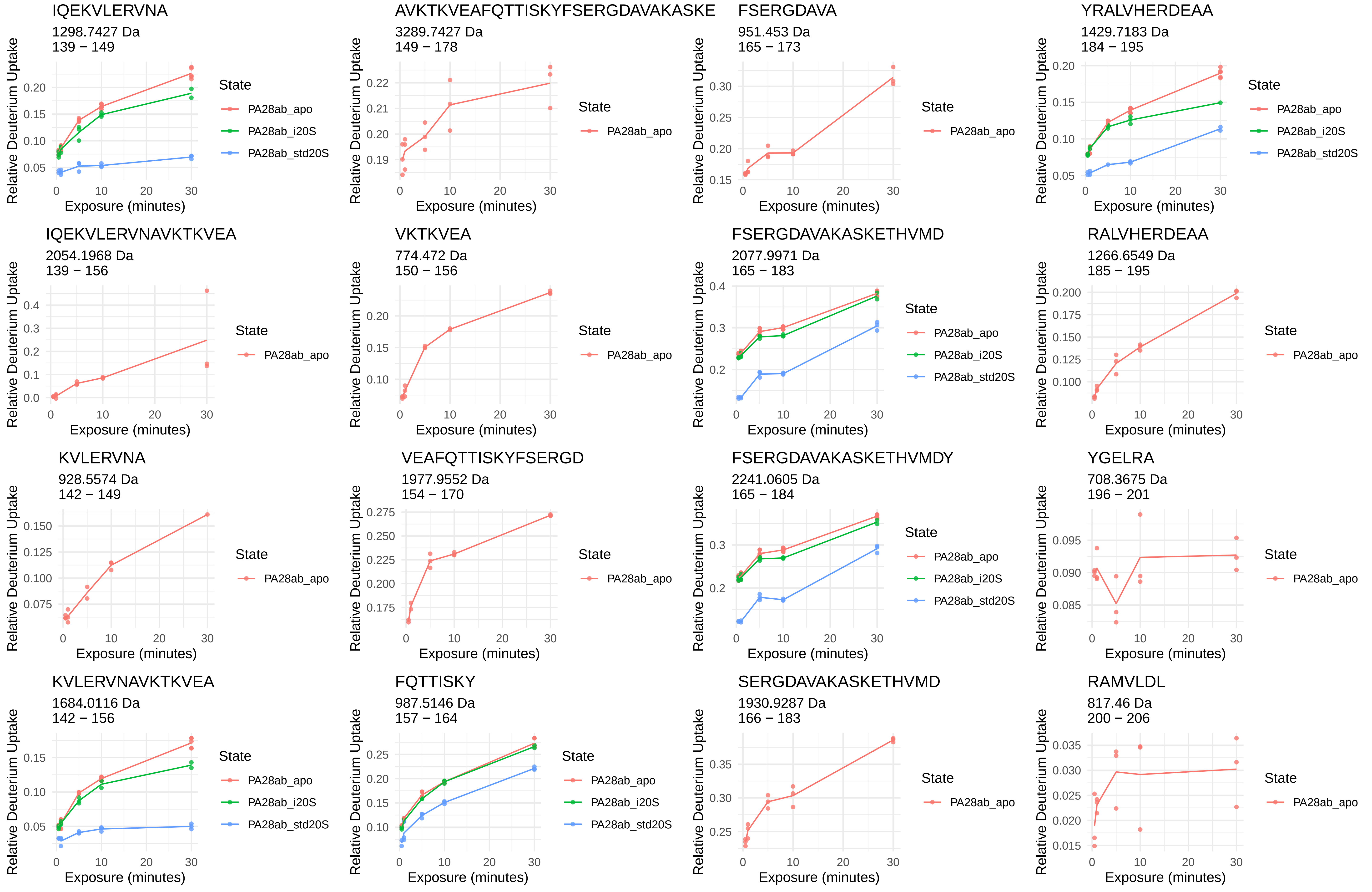
IQEKVLERVNA
1298.7427 Da
139 − 149
AVKTKVEAFQTTISKYFSERGDAVAKASKE
3289.7427 Da
149 − 178
FSERGDAVA
951.453 Da
165 − 173
YRALVHERDEAA
1429.7183 Da
184 − 195
IQEKVLERVNAVKTKVEA
2054.1968 Da
139 − 156
VKTKVEA
774.472 Da
150 − 156
FSERGDAVAKASKETHVMD
2077.9971 Da
165 − 183
RALVHERDEAA
1266.6549 Da
185 − 195
KVLERVNA
928.5574 Da
142 − 149
VEAFQTTISKYFSERGD
1977.9552 Da
154 − 170
FSERGDAVAKASKETHVMDY
2241.0605 Da
165 − 184
YGELRA
708.3675 Da
196 − 201
KVLERVNAVKTKVEA
1684.0116 Da
142 − 156
FQTTISKY
987.5146 Da
157 − 164
SERGDAVAKASKETHVMD
1930.9287 Da
166 − 183
RAMVLDL
817.46 Da
200 − 206
Relative Deuterium Uptake
Exposure (minutes)
State
PA28ab_apo
PA28ab_i20S
PA28ab_std20S

MVLDL

590.3218 Da
202 − 206

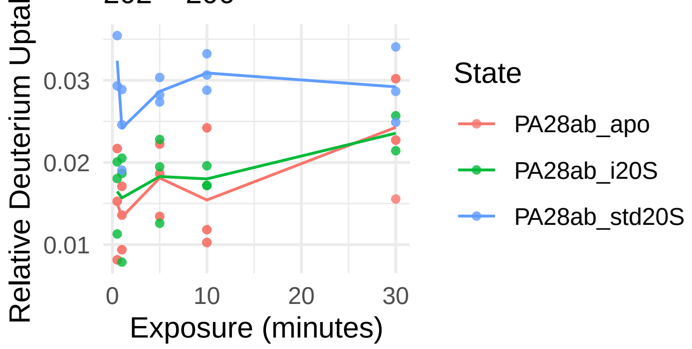


EKIVNPKGEEKPSMY

1748.8887 Da
222 − 236

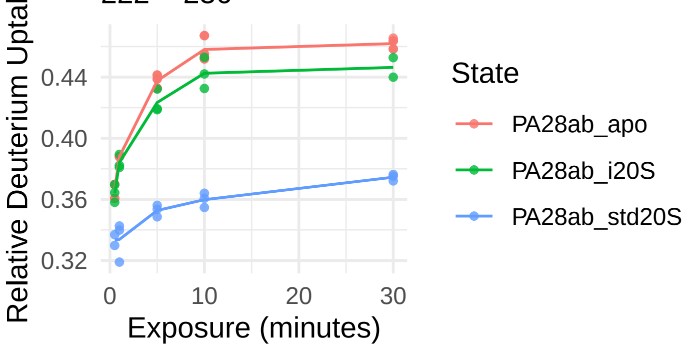


FYAEL

642.3134 Da
209 − 213

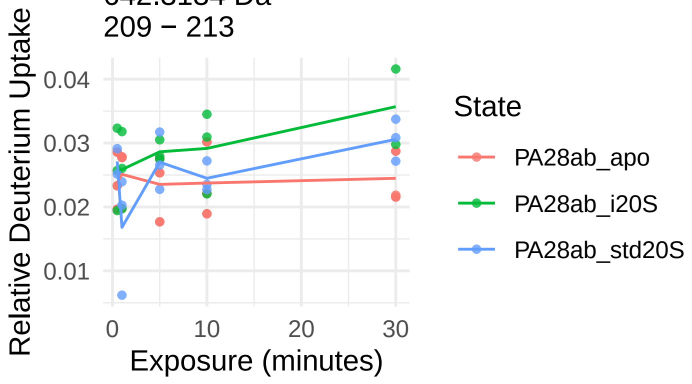


YHIISSNL

946.4993 Da
214 − 221

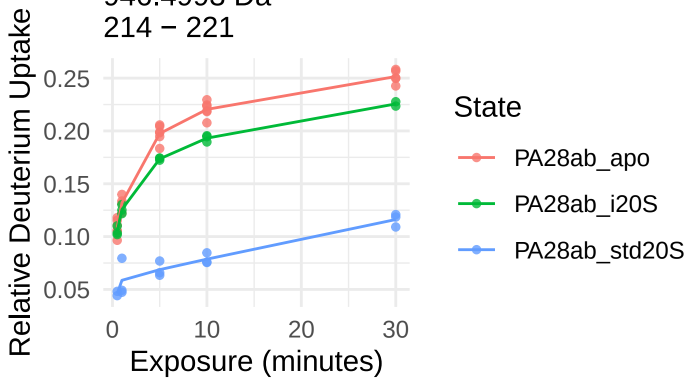


EKIVNPKGEEKPSM

1585.8254 Da
222 − 235

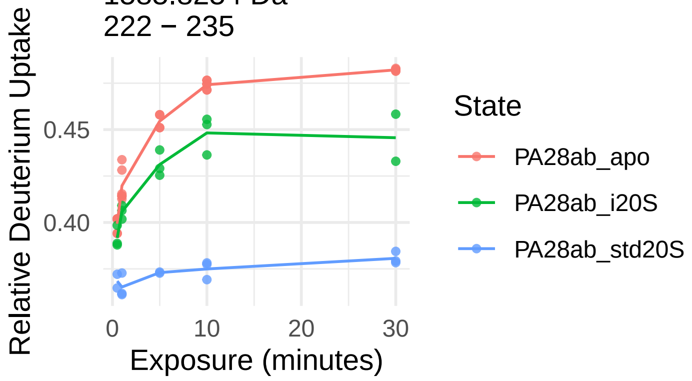

PA28γ

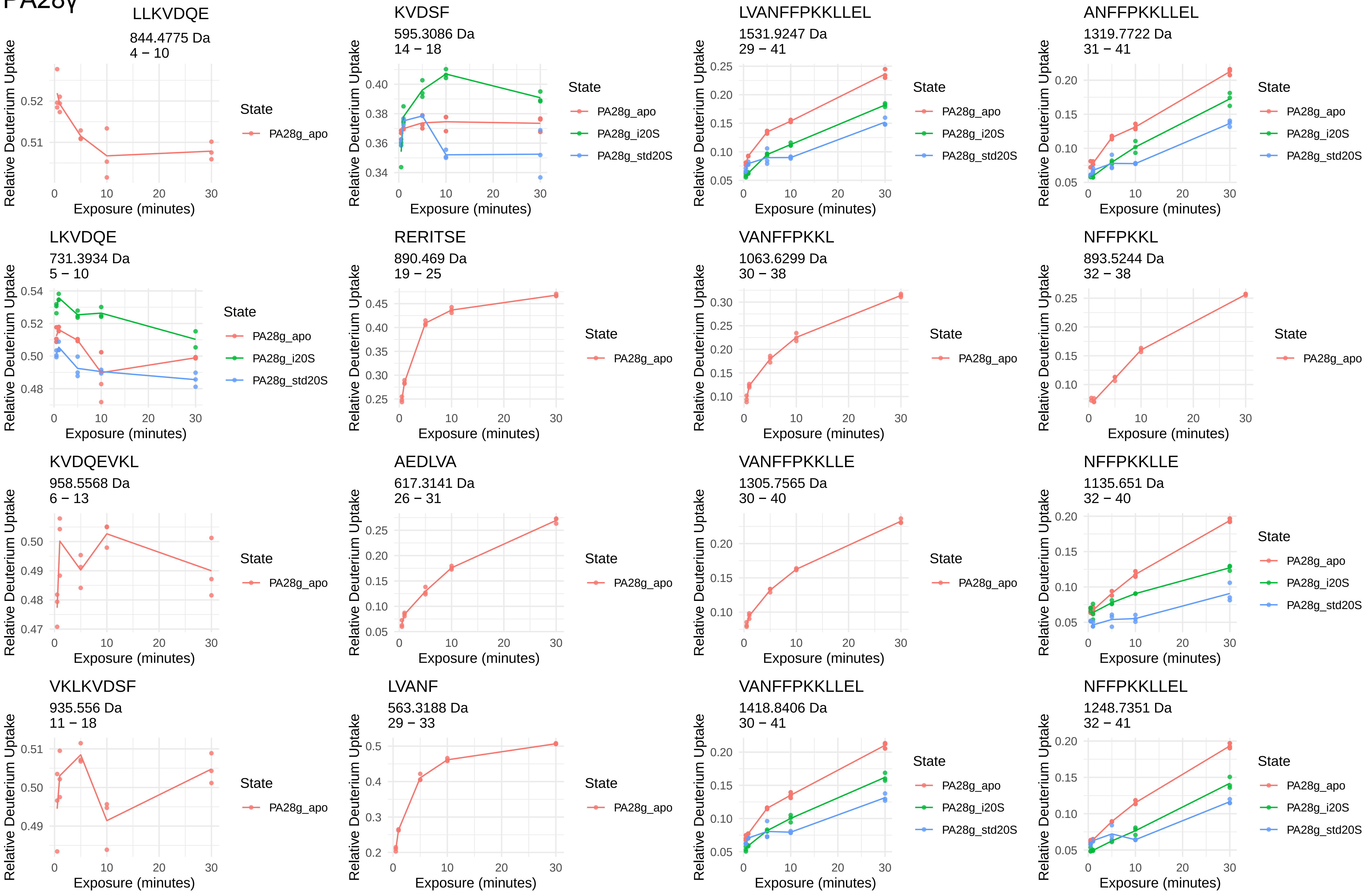
LLKVDQE
844.4775 Da
4 − 10
KVDSF
595.3086 Da
14 − 18
LVANFFPKKLLEL
1531.9247 Da
29 − 41
ANFFPKKLLEL
1319.7722 Da
31 − 41
LKVDQE
731.3934 Da
5 − 10
RERITSE
890.469 Da
19 − 25
VANFFPKKL
1063.6299 Da
30 − 38
NFFPKKL
893.5244 Da
32 − 38
KVDQEVKL
958.5568 Da
6 − 13
AEDLVA
617.3141 Da
26 − 31
VANFFPKKLLE
1305.7565 Da
30 − 40
NFFPKKLLE
1135.651 Da
32 − 40
VKLKVDSF
935.556 Da
11 − 18
LVANF
563.3188 Da
29 − 33
VANFFPKKLLEL
1418.8406 Da
30 − 41
NFFPKKLLEL
1248.7351 Da
32 − 41
Relative Deuterium Uptake
Exposure (minutes)
State
PA28g_apo
PA28g_i20S
PA28g_std20S

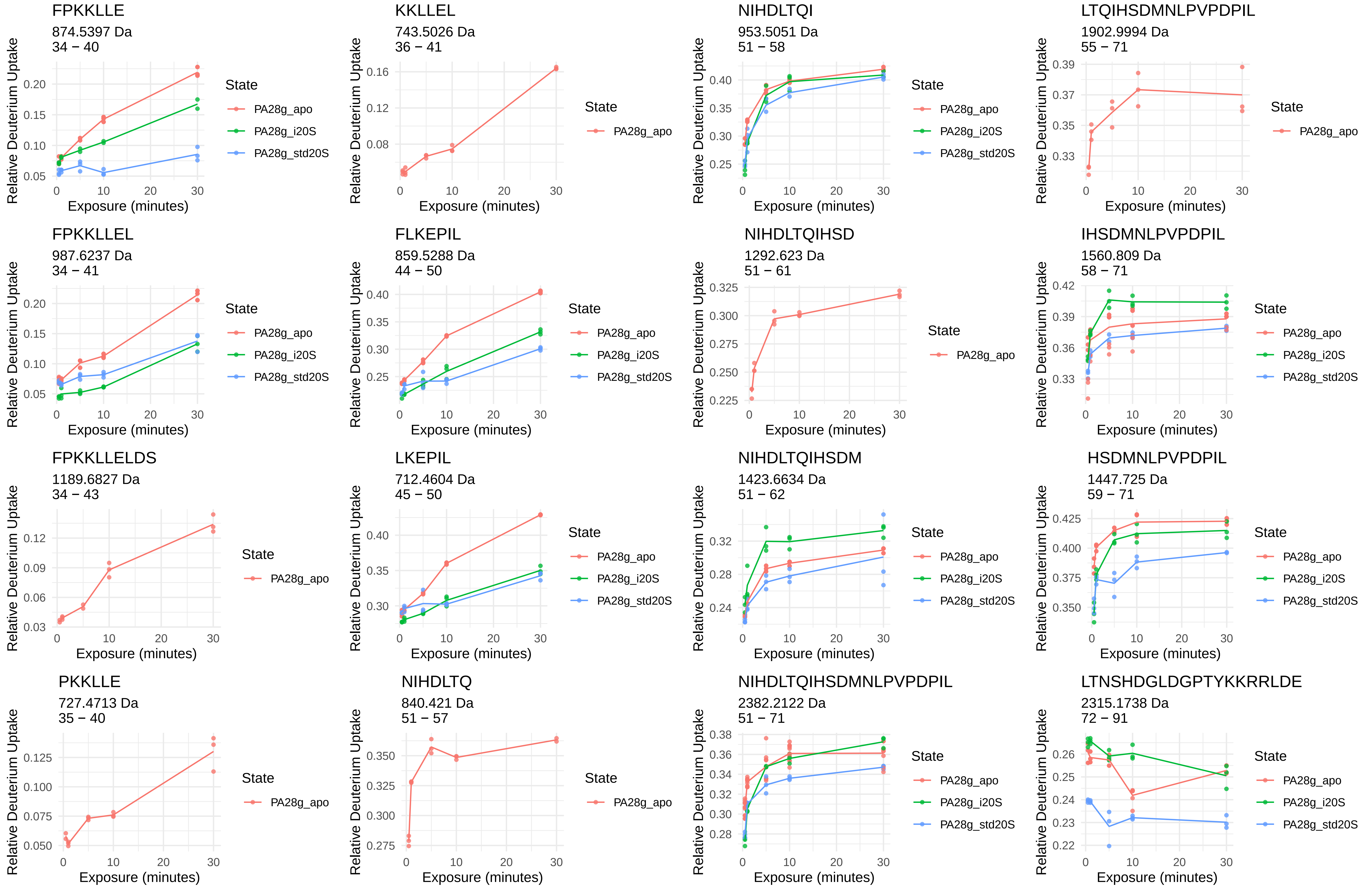
FPKKLLE
874.5397 Da
34 − 40
KKLLEL
743.5026 Da
36 − 41
NIHDLTQI
953.5051 Da
51 − 58
LTQIHSDMNLPVPDPIL
1902.9994 Da
55 − 71
FPKKLLEL
987.6237 Da
34 − 41
FLKEPIL
859.5288 Da
44 − 50
NIHDLTQIHSD
1292.623 Da
51 − 61
IHSDMNLPVPDPIL
1560.809 Da
58 − 71
FPKKLLELDS
1189.6827 Da
34 − 43
LKEPIL
712.4604 Da
45 − 50
NIHDLTQIHSDM
1423.6634 Da
51 − 62
HSDMNLPVPDPIL
1447.725 Da
59 − 71
PKKLLE
727.4713 Da
35 − 40
NIHDLTQ
840.421 Da
51 − 57
NIHDLTQIHSDMNLPVPDPIL
2382.2122 Da
51 − 71
LTNSHDGLDGPTYKKRRLDE
2315.1738 Da
72 − 91
Relative Deuterium Uptake
Exposure (minutes)
State
PA28g_apo
PA28g_i20S
PA28g_std20S

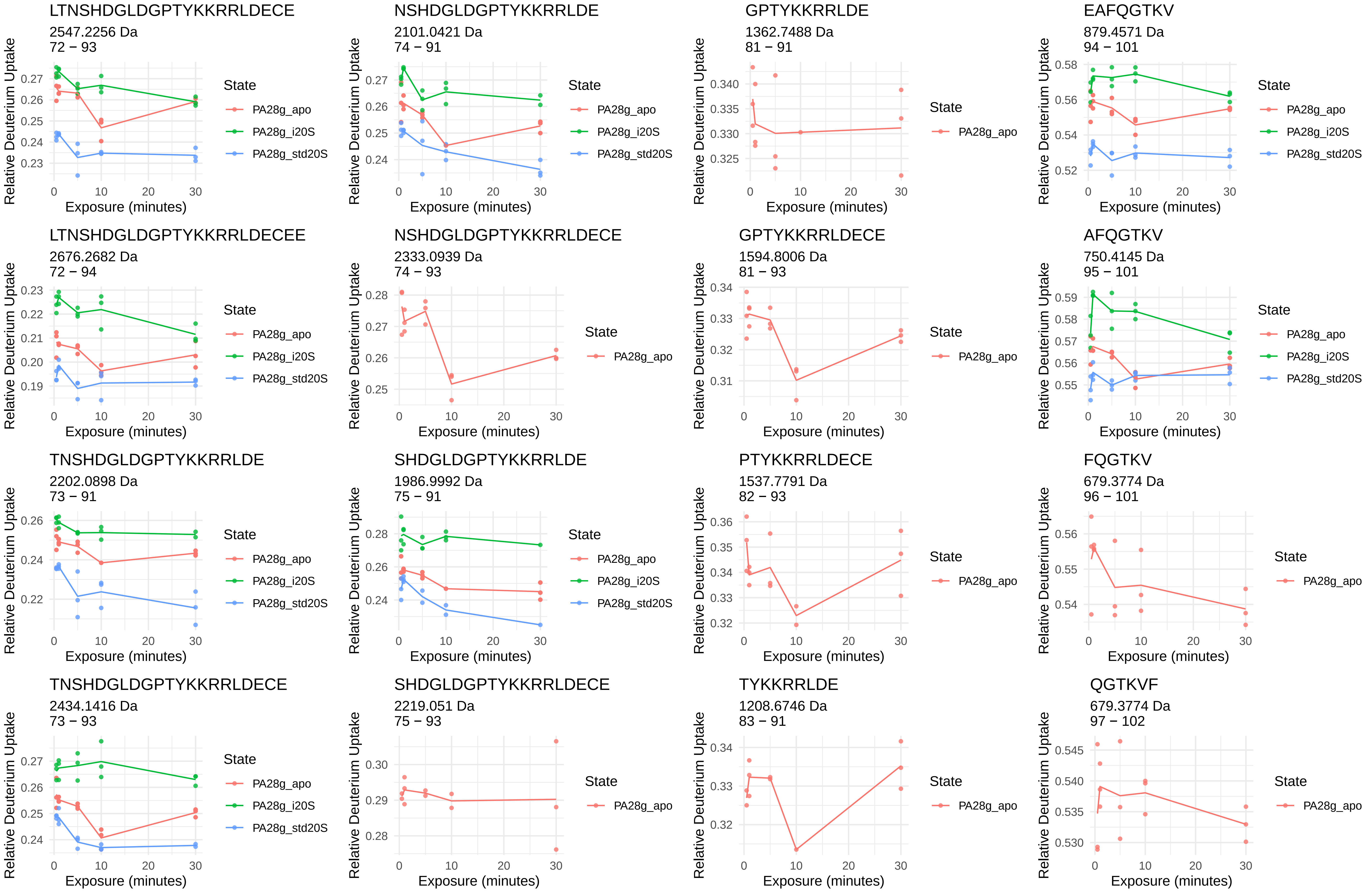
LTNSHDGLDGPTYKKRRLDECE
2547.2256 Da
72 − 93
NSHDGLDGPTYKKRRLDE
2101.0421 Da
74 − 91
GPTYKKRRLDE
1362.7488 Da
81 − 91
EAFQGTKV
879.4571 Da
94 − 101
LTNSHDGLDGPTYKKRRLDECEE
2676.2682 Da
72 − 94
NSHDGLDGPTYKKRRLDECE
2333.0939 Da
74 − 93
GPTYKKRRLDECE
1594.8006 Da
81 − 93
AFQGTKV
750.4145 Da
95 − 101
TNSHDGLDGPTYKKRRLDE
2202.0898 Da
73 − 91
SHDGLDGPTYKKRRLDE
1986.9992 Da
75 − 91
PTYKKRRLDECE
1537.7791 Da
82 − 93
FQGTKV
679.3774 Da
96 − 101
TNSHDGLDGPTYKKRRLDECE
2434.1416 Da
73 − 93
SHDGLDGPTYKKRRLDECE
2219.051 Da
75 − 93
TYKKRRLDE
1208.6746 Da
83 − 91
QGTKVF
679.3774 Da
97 − 102
Relative Deuterium Uptake
Exposure (minutes)
State
PA28g_apo
PA28g_i20S
PA28g_std20S

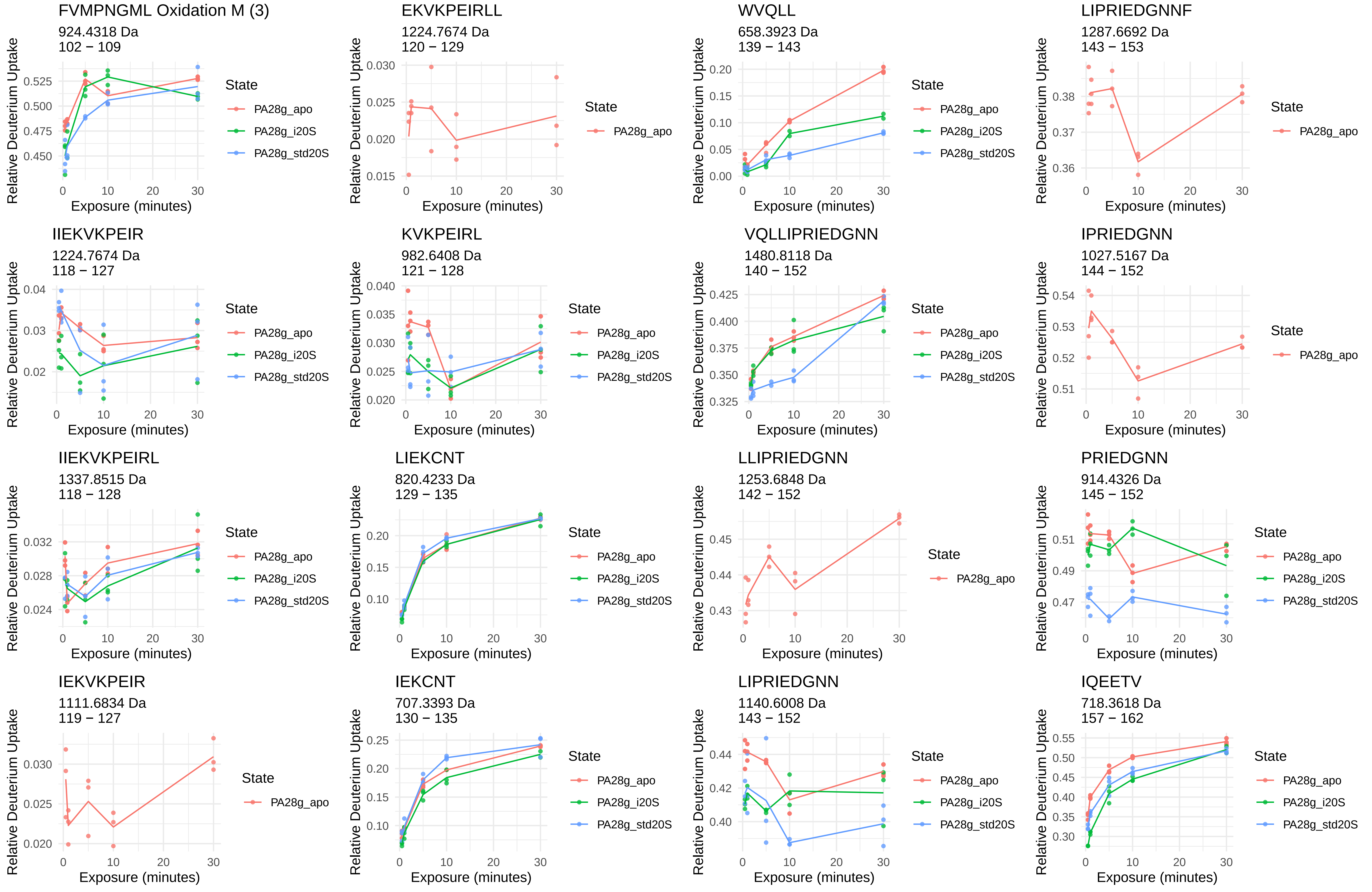
FVMPNGML Oxidation M (3)
924.4318 Da
102 − 109
EKVKPEIRLL
1224.7674 Da
120 − 129
WVQLL
658.3923 Da
139 − 143
LIPRIEDGNNF
1287.6692 Da
143 − 153
IIEKVKPEIR
1224.7674 Da
118 − 127
KVKPEIRL
982.6408 Da
121 − 128
VQLLIPRIEDGNN
1480.8118 Da
140 − 152
IPRIEDGNN
1027.5167 Da
144 − 152
IIEKVKPEIRL
1337.8515 Da
118 − 128
LIEKCNT
820.4233 Da
129 − 135
LLIPRIEDGNN
1253.6848 Da
142 − 152
PRIEDGNN
914.4326 Da
145 − 152
IEKVKPEIR
1111.6834 Da
119 − 127
IEKCNT
707.3393 Da
130 − 135
LIPRIEDGNN
1140.6008 Da
143 − 152
IQEETV
718.3618 Da
157 − 162
Relative Deuterium Uptake
Exposure (minutes)
State
PA28g_apo
PA28g_i20S
PA28g_std20S

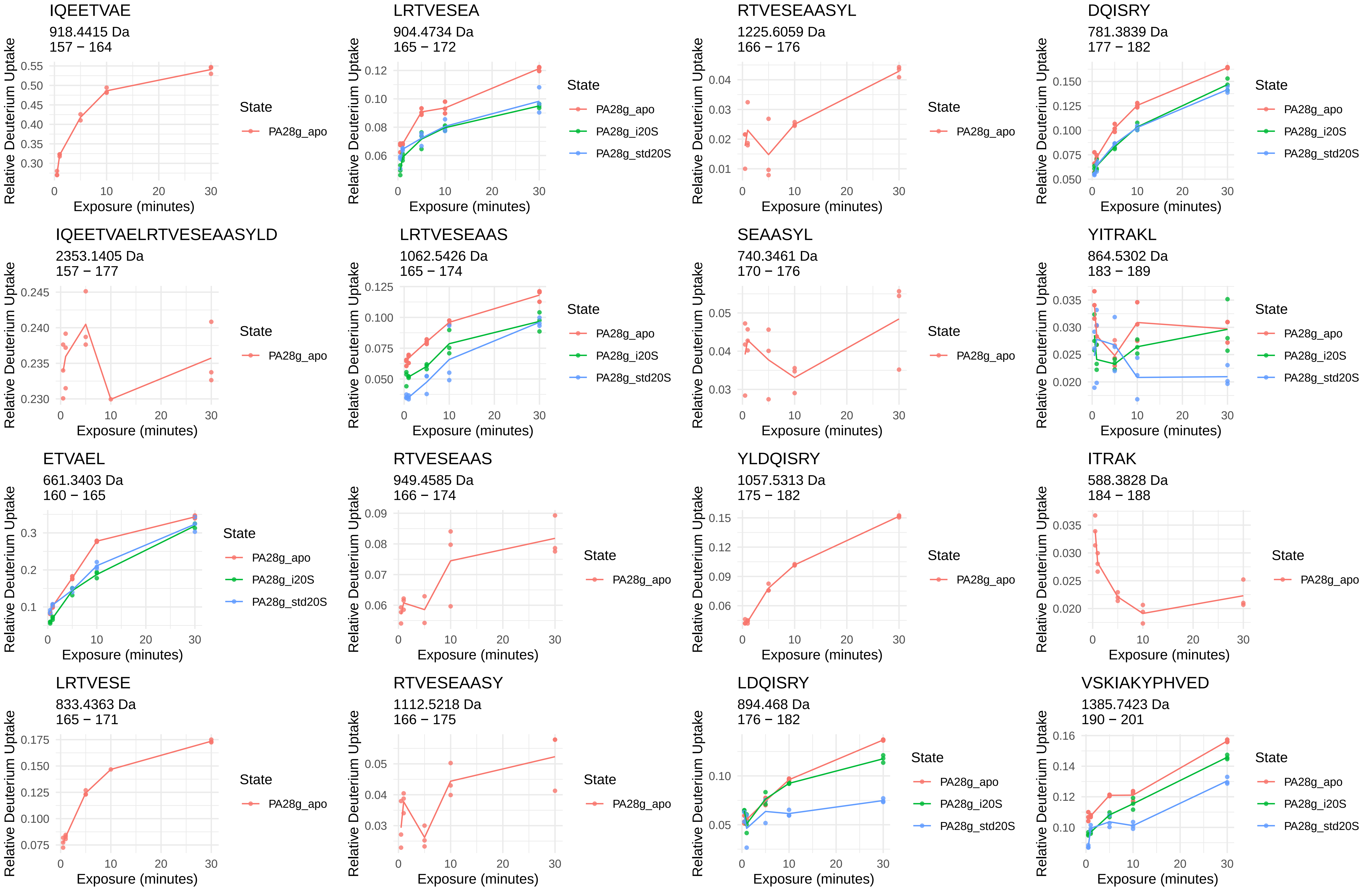
IQEETVAE
918.4415 Da
157 − 164
LRTVESEA
904.4734 Da
165 − 172
RTVESEAASYL
1225.6059 Da
166 − 176
DQISRY
781.3839 Da
177 − 182
IQEETVAELRTVESEAASYLD
2353.1405 Da
157 − 177
LRTVESEAAS
1062.5426 Da
165 − 174
SEAASYL
740.3461 Da
170 − 176
YITRAKL
864.5302 Da
183 − 189
ETVAEL
661.3403 Da
160 − 165
RTVESEAAS
949.4585 Da
166 − 174
YLDQISRY
1057.5313 Da
175 − 182
ITRAK
588.3828 Da
184 − 188
LRTVESE
833.4363 Da
165 − 171
RTVESEAASY
1112.5218 Da
166 − 175
LDQISRY
894.468 Da
176 − 182
VSKIAKYPHVED
1385.7423 Da
190 − 201
Relative Deuterium Uptake
Exposure (minutes)
State
PA28g_apo
PA28g_i20S
PA28g_std20S

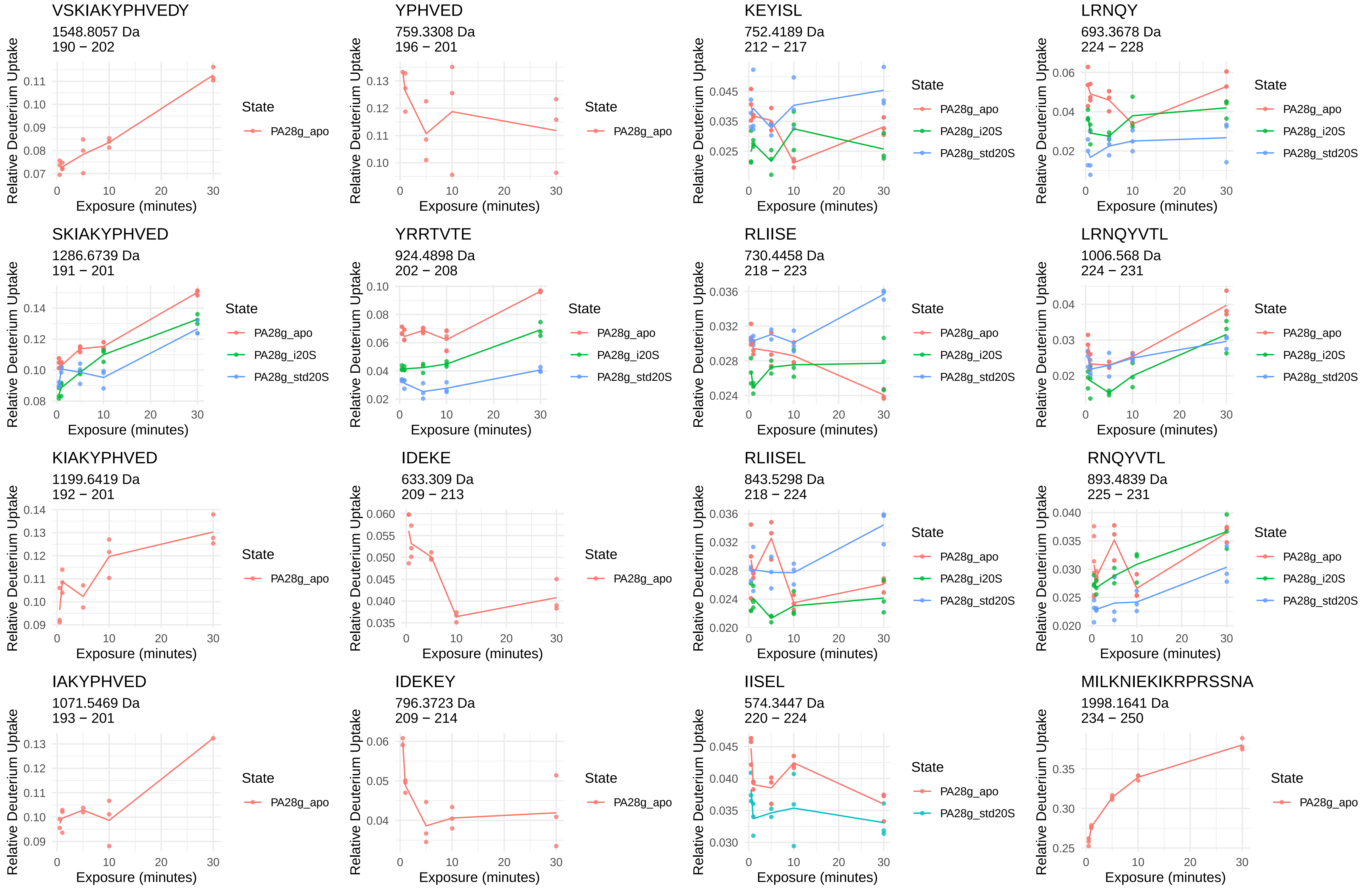
VSKIAKYPHVEDY
1548.8057 Da
190 − 202
YPHVED
759.3308 Da
196 − 201
KEYISL
752.4189 Da
212 − 217
LRNQY
693.3678 Da
224 − 228
SKIAKYPHVED
1286.6739 Da
191 − 201
YRRTVTE
924.4898 Da
202 − 208
RLIISE
730.4458 Da
218 − 223
LRNQYVTL
1006.568 Da
224 − 231
KIAKYPHVED
1199.6419 Da
192 − 201
IDEKE
633.309 Da
209 − 213
RLIISEL
843.5298 Da
218 − 224
RNQYVTL
893.4839 Da
225 − 231
IAKYPHVED
1071.5469 Da
193 − 201
IDEKEY
796.3723 Da
209 − 214
IISEL
574.3447 Da
220 − 224
MILKNIEKIKRPRSSNA
1998.1641 Da
234 − 250
Relative Deuterium Uptake
Exposure (minutes)
State
PA28g_apo
PA28g_i20S
PA28g_std20S

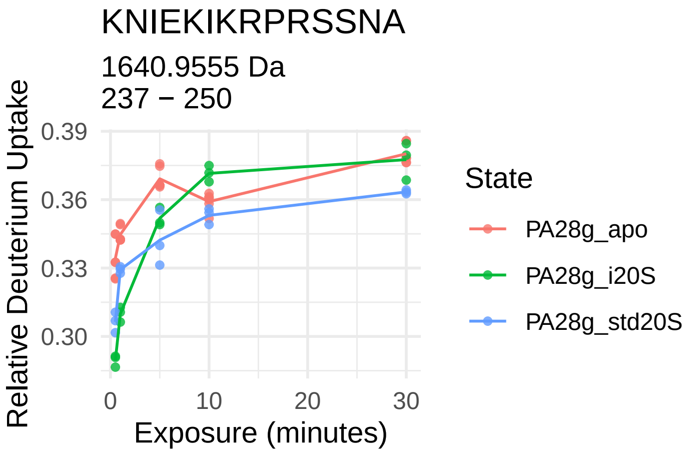
KNIEKIKRPRSSNA
1640.9555 Da
237 − 250
Relative Deuterium Uptake
0.39
0.36
0.33
0.30
0
10
20
30
Exposure (minutes)
State
PA28g_apo
PA28g_i20S
PA28g_std20S
