## Supplementary material for "Conformational maps of human 20S proteasomes reveal PA28- and immuno-dependent inter-ring crosstalks": Fig. S10

# α1

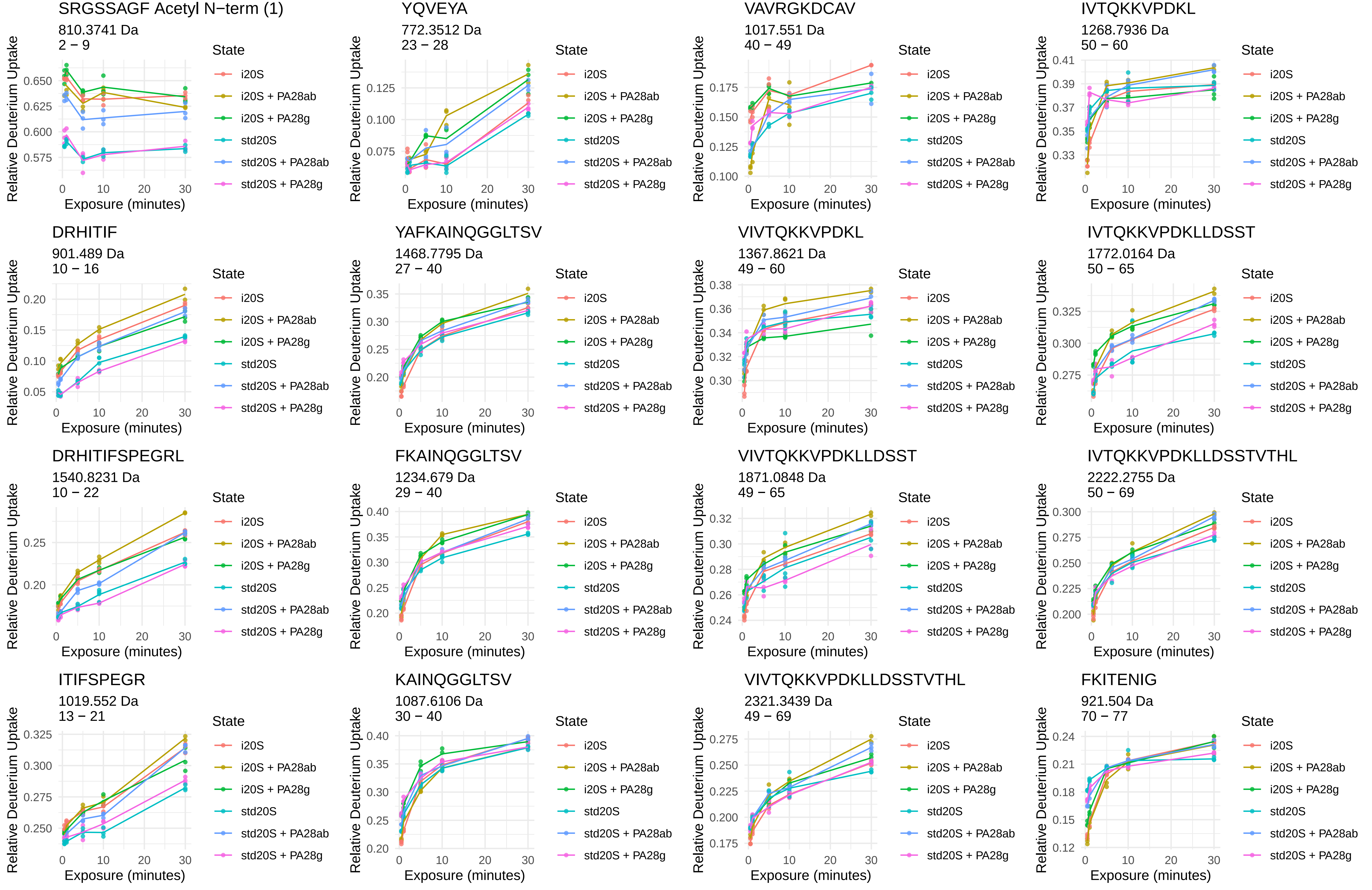
SRGSSAGF Acetyl N−term (1)
810.3741 Da
2 − 9
YQVEYA
772.3512 Da
23 − 28
VAVRGKDCAV
1017.551 Da
40 − 49
IVTQKKVPDKL
1268.7936 Da
50 − 60
DRHITIF
901.489 Da
10 − 16
YAFKAINQGGLTSV
1468.7795 Da
27 − 40
VIVTQKKVPDKL
1367.8621 Da
49 − 60
IVTQKKVPDKLLDSST
1772.0164 Da
50 − 65
DRHITIFSPEGRL
1540.8231 Da
10 − 22
FKAINQGGLTSV
1234.679 Da
29 − 40
VIVTQKKVPDKLLDSST
1871.0848 Da
49 − 65
IVTQKKVPDKLLDSSTVTHL
2222.2755 Da
50 − 69
ITIFSPEGR
1019.552 Da
13 − 21
KAINQGGLTSV
1087.6106 Da
30 − 40
VIVTQKKVPDKLLDSSTVTHL
2321.3439 Da
49 − 69
FKITENIG
921.504 Da
70 − 77
Relative Deuterium Uptake
Exposure (minutes)
State
i20S
i20S + PA28ab
i20S + PA28g
std20S
std20S + PA28ab
std20S + PA28g

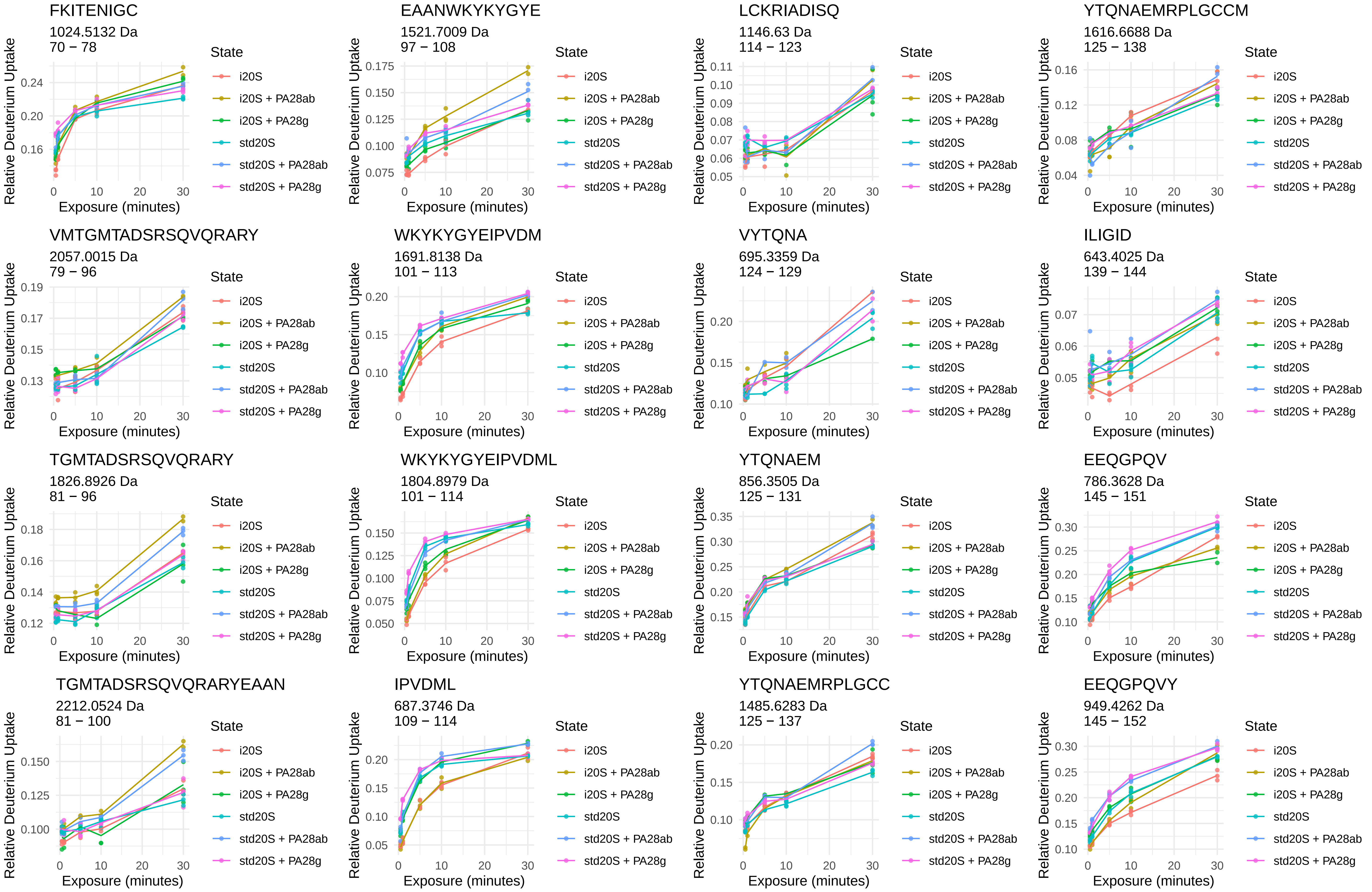
FKITENIGC
1024.5132 Da
70 − 78
EAANWKYKYGYE
1521.7009 Da
97 − 108
LCKRIADISQ
1146.63 Da
114 − 123
YTQNAEMRPLGCCM
1616.6688 Da
125 − 138
VMTGMTADSRSQVQRARY
2057.0015 Da
79 − 96
WKYKYGYEIPVDM
1691.8138 Da
101 − 113
VYTQNA
695.3359 Da
124 − 129
ILIGID
643.4025 Da
139 − 144
TGMTADSRSQVQRARY
1826.8926 Da
81 − 96
WKYKYGYEIPVDML
1804.8979 Da
101 − 114
YTQNAEM
856.3505 Da
125 − 131
EEQGPQV
786.3628 Da
145 − 151
TGMTADSRSQVQRARYEAAN
2212.0524 Da
81 − 100
IPVDML
687.3746 Da
109 − 114
YTQNAEMRPLGCC
1485.6283 Da
125 − 137
EEQGPQVY
949.4262 Da
145 − 152
Relative Deuterium Uptake
Exposure (minutes)
State
i20S
i20S + PA28ab
i20S + PA28g
std20S
std20S + PA28ab
std20S + PA28g

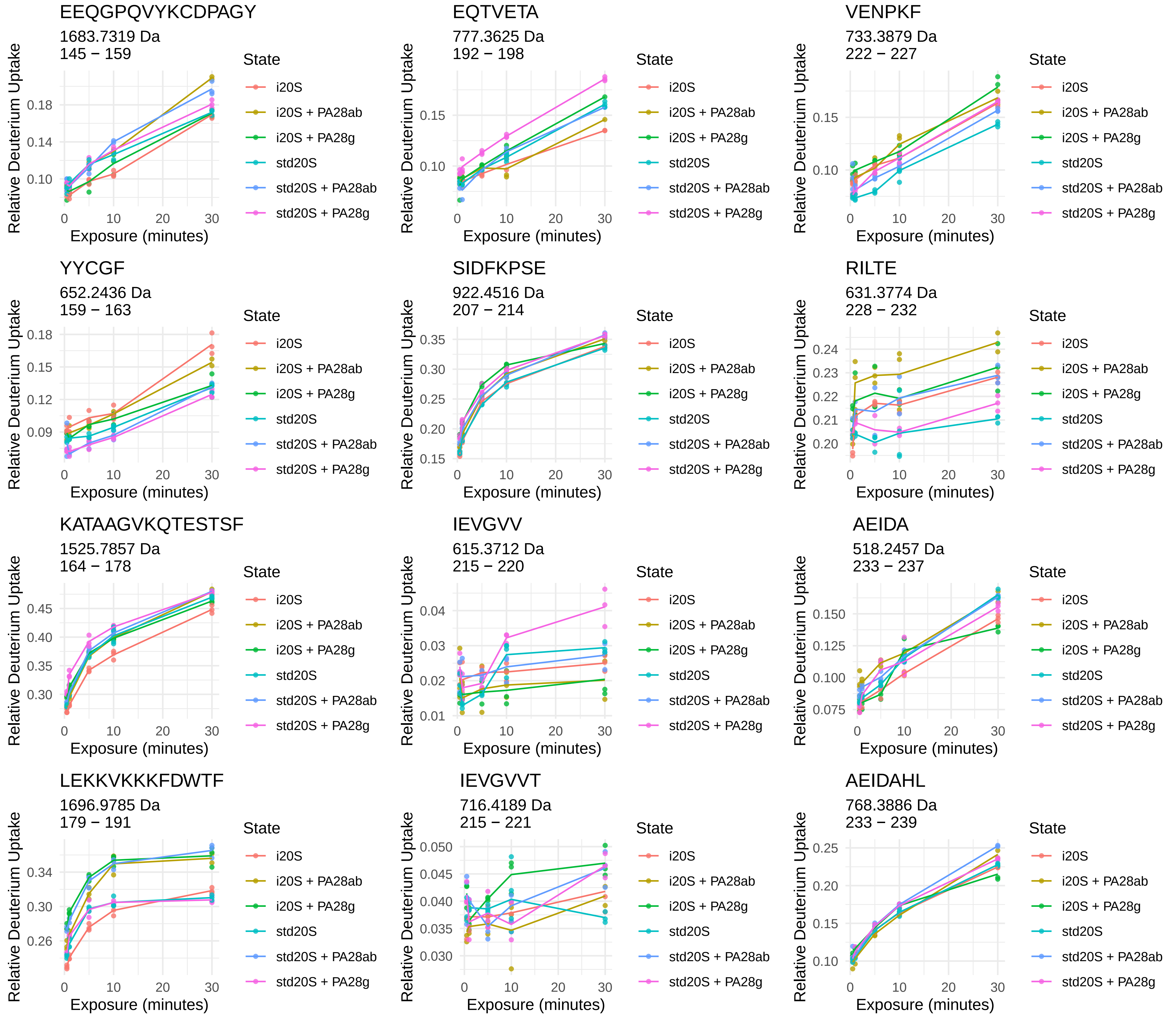
EEQGPQVYKCDPAGY
1683.7319 Da
145 − 159
EQTVETA
777.3625 Da
192 − 198
VENPKF
733.3879 Da
222 − 227
YYCGF
652.2436 Da
159 − 163
SIDFKPSE
922.4516 Da
207 − 214
RILTE
631.3774 Da
228 − 232
KATAAGVKQTESTSF
1525.7857 Da
164 − 178
IEVGVV
615.3712 Da
215 − 220
AEIDA
518.2457 Da
233 − 237
LEKKVKKKFDWTF
1696.9785 Da
179 − 191
IEVGVVT
716.4189 Da
215 − 221
AEIDAHL
768.3886 Da
233 − 239
Relative Deuterium Uptake
Exposure (minutes)
State
i20S
i20S + PA28ab
i20S + PA28g
std20S
std20S + PA28ab
std20S + PA28g

α2



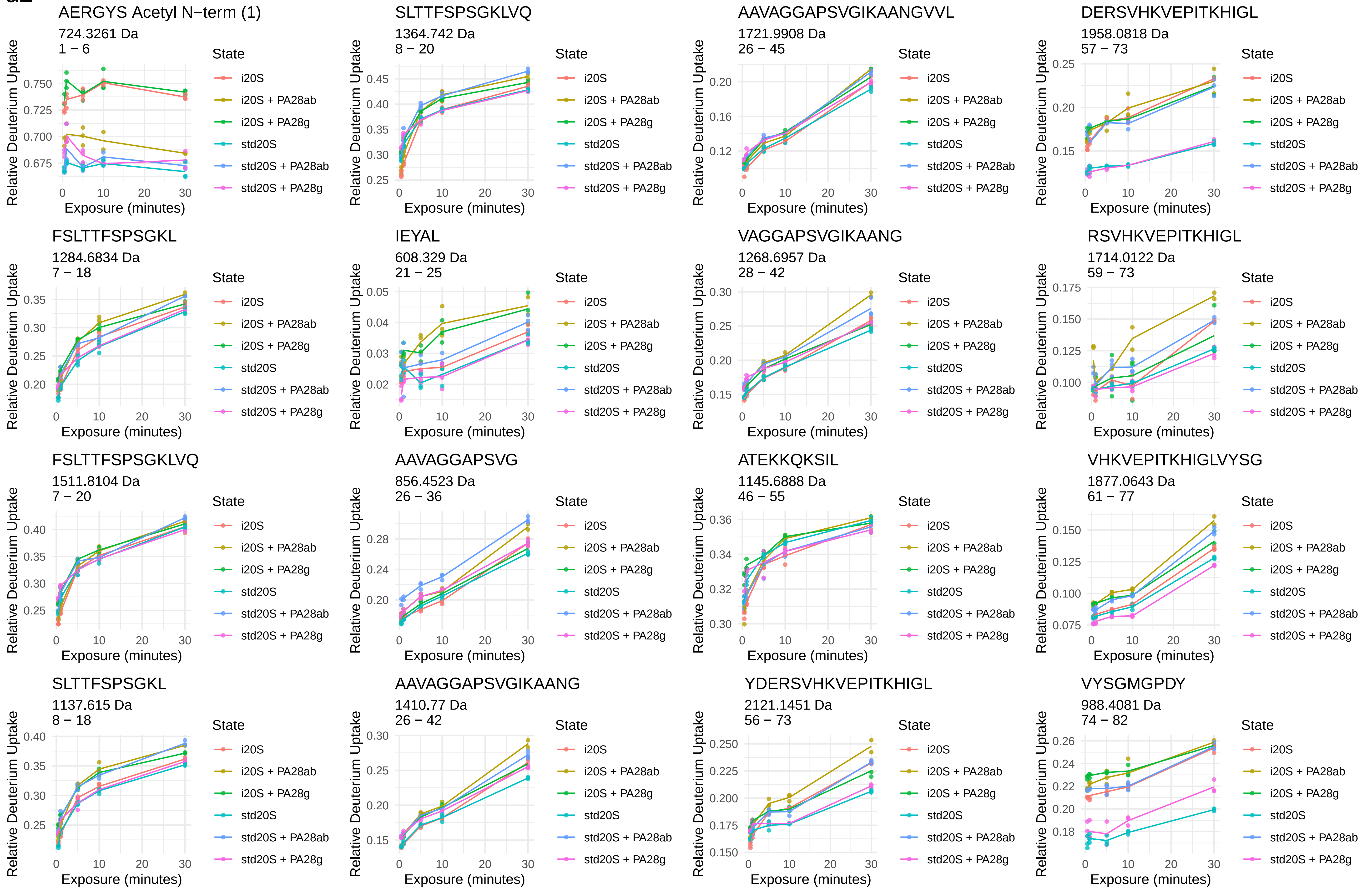
AERGYS Acetyl N−term (1)
724.3261 Da
1 − 6
SLTTFSPSGKLVQ
1364.742 Da
8 − 20
AAVAGGAPSVGIKAANGVVL
1721.9908 Da
26 − 45
DERSVHKVEPITKHIGL
1958.0818 Da
57 − 73
FSLTTFSPSGKL
1284.6834 Da
7 − 18
IEYAL
608.329 Da
21 − 25
VAGGAPSVGIKAANG
1268.6957 Da
28 − 42
RSVHKVEPITKHIGL
1714.0122 Da
59 − 73
FSLTTFSPSGKLVQ
1511.8104 Da
7 − 20
AAVAGGAPSVG
856.4523 Da
26 − 36
ATEKKQKSIL
1145.6888 Da
46 − 55
VHKVEPITKHIGLVYSG
1877.0643 Da
61 − 77
SLTTFSPSGKL
1137.615 Da
8 − 18
AAVAGGAPSVGIKAANG
1410.77 Da
26 − 42
YDERSVHKVEPITKHIGL
2121.1451 Da
56 − 73
VYSGMGPDY
988.4081 Da
74 − 82
Relative Deuterium Uptake
Exposure (minutes)
State
i20S
i20S + PA28ab
i20S + PA28g
std20S
std20S + PA28ab
std20S + PA28g

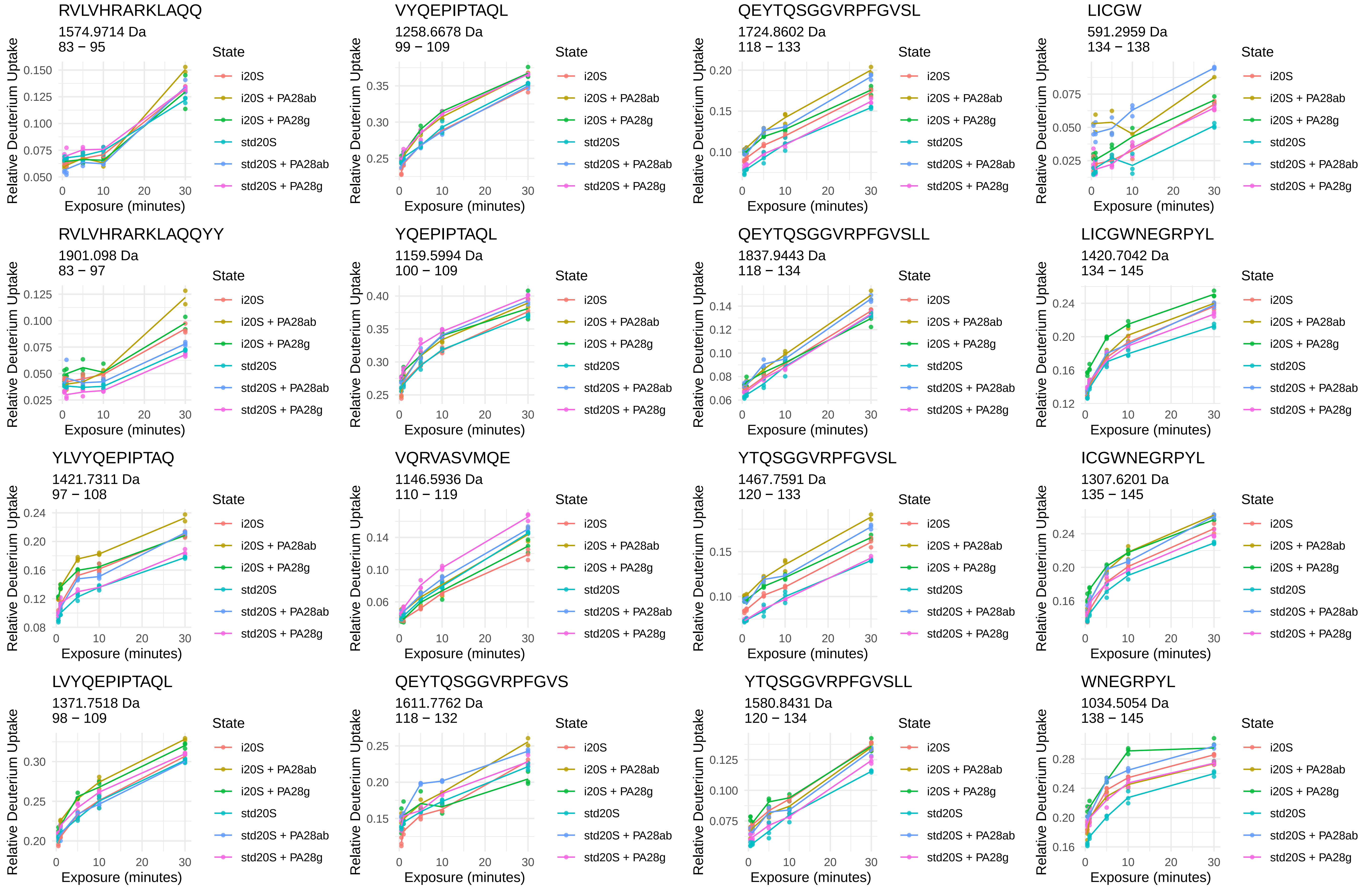
RVLVHRARKLAQQ
1574.9714 Da
83 − 95
VYQEPIPTAQL
1258.6678 Da
99 − 109
QEYTQSGGVRPFGVSL
1724.8602 Da
118 − 133
LICGW
591.2959 Da
134 − 138
RVLVHRARKLAQQYY
1901.098 Da
83 − 97
YQEPIPTAQL
1159.5994 Da
100 − 109
QEYTQSGGVRPFGVSLL
1837.9443 Da
118 − 134
LICGWNEGRPYL
1420.7042 Da
134 − 145
YLVYQEPIPTAQ
1421.7311 Da
97 − 108
VQRVASVMQE
1146.5936 Da
110 − 119
YTQSGGVRPFGVSL
1467.7591 Da
120 − 133
ICGWNEGRPYL
1307.6201 Da
135 − 145
LVYQEPIPTAQL
1371.7518 Da
98 − 109
QEYTQSGGVRPFGVS
1611.7762 Da
118 − 132
YTQSGGVRPFGVSLL
1580.8431 Da
120 − 134
WNEGRPYL
1034.5054 Da
138 − 145
Relative Deuterium Uptake
Exposure (minutes)
State
i20S
i20S + PA28ab
i20S + PA28g
std20S
std20S + PA28ab
std20S + PA28g

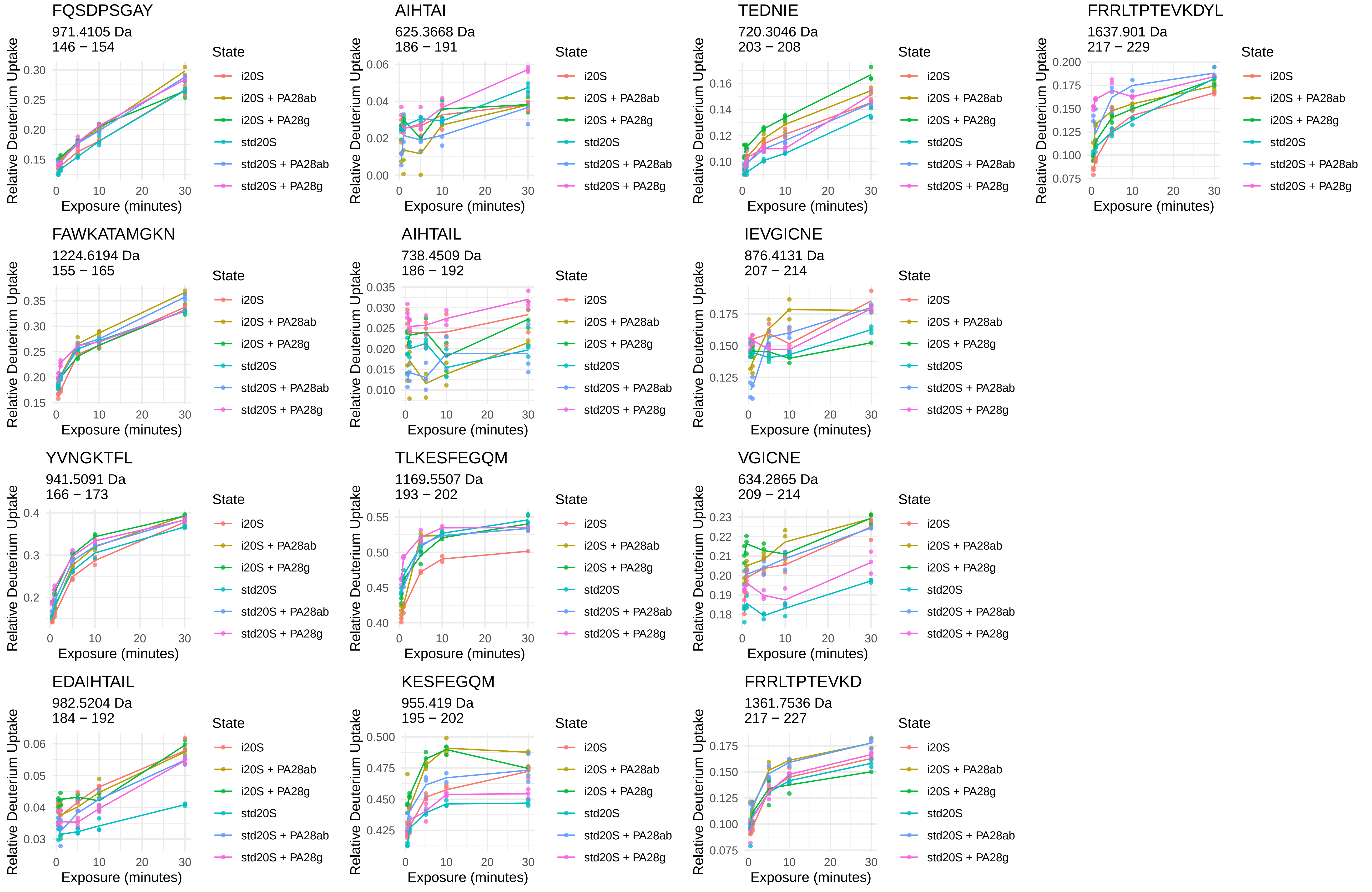
FQSDPSGAY
971.4105 Da
146 − 154
AIHTAI
625.3668 Da
186 − 191
TEDNIE
720.3046 Da
203 − 208
FRRLTPTEVKDYL
1637.901 Da
217 − 229
FAWKATAMGKN
1224.6194 Da
155 − 165
AIHTAIL
738.4509 Da
186 − 192
IEVGICNE
876.4131 Da
207 − 214
YVNGKTFL
941.5091 Da
166 − 173
TLKESFEGQM
1169.5507 Da
193 − 202
VGICNE
634.2865 Da
209 − 214
EDAIHTAIL
982.5204 Da
184 − 192
KESFEGQM
955.419 Da
195 − 202
FRRLTPTEVKD
1361.7536 Da
217 − 227
Relative Deuterium Uptake
Exposure (minutes)
State
i20S
i20S + PA28ab
i20S + PA28g
std20S
std20S + PA28ab
std20S + PA28g

α3



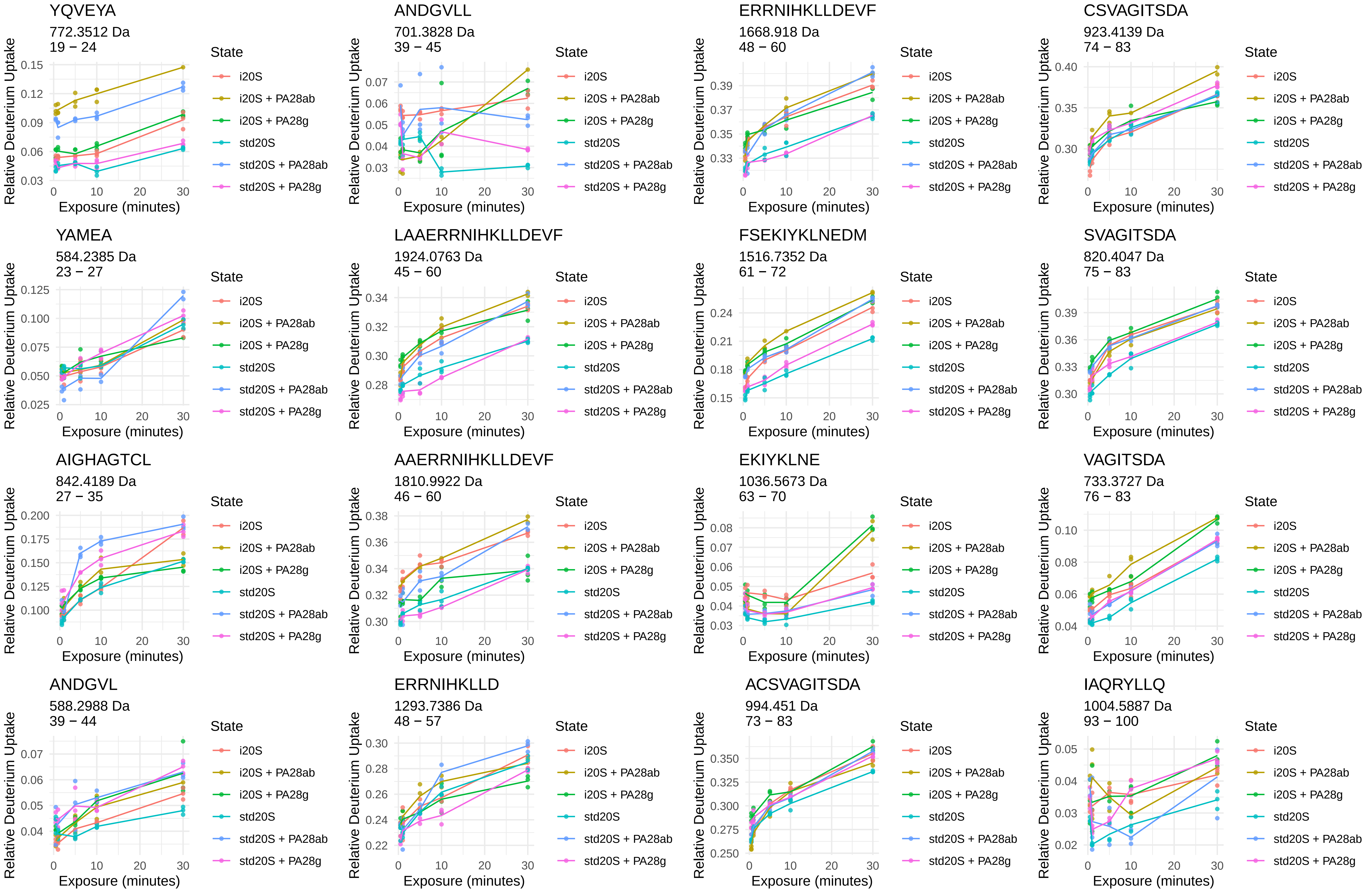
YQVEYA
772.3512 Da
19 − 24
ANDGVLL
701.3828 Da
39 − 45
ERRNIHKLLDEVF
1668.918 Da
48 − 60
CSVAGITSDA
923.4139 Da
74 − 83
YAMEA
584.2385 Da
23 − 27
LAAERRNIHKLLDEVF
1924.0763 Da
45 − 60
FSEKIYKLNEDM
1516.7352 Da
61 − 72
SVAGITSDA
820.4047 Da
75 − 83
AIGHAGTCL
842.4189 Da
27 − 35
AAERRNIHKLLDEVF
1810.9922 Da
46 − 60
EKIYKLNE
1036.5673 Da
63 − 70
VAGITSDA
733.3727 Da
76 − 83
ANDGVL
588.2988 Da
39 − 44
ERRNIHKLLD
1293.7386 Da
48 − 57
ACSVAGITSDA
994.451 Da
73 − 83
IAQRYLLQ
1004.5887 Da
93 − 100
Relative Deuterium Uptake
Exposure (minutes)
State
i20S
i20S + PA28ab
i20S + PA28g
std20S
std20S + PA28ab
std20S + PA28g

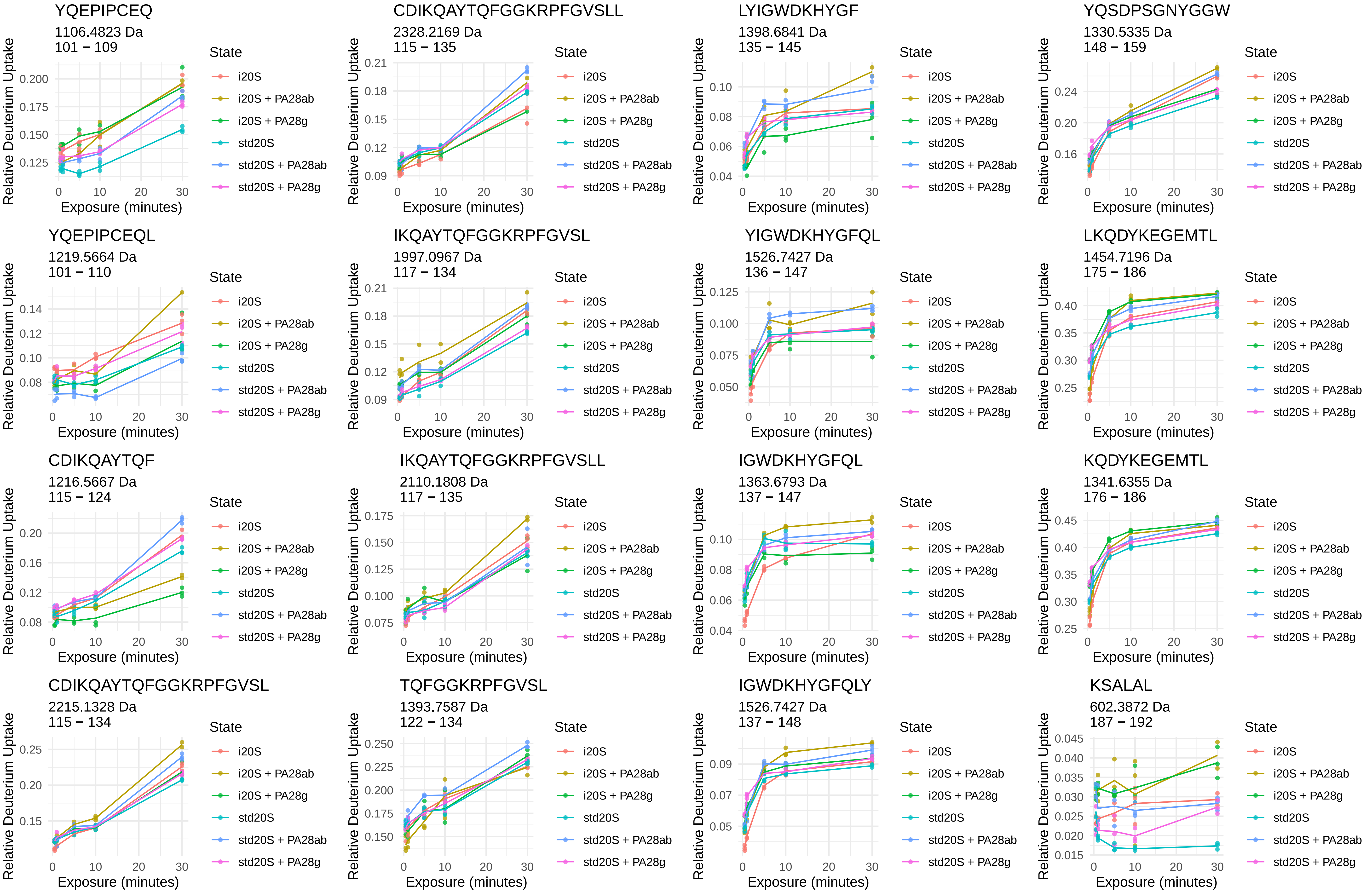

YQEPIPCEQ
1106.4823 Da
101 − 109
CDIKQAYTQFGGKRPFGVSLL
2328.2169 Da
115 − 135
LYIGWDKHYGF
1398.6841 Da
135 − 145
YQSDPSGNYGGW
1330.5335 Da
148 − 159
YQEPIPCEQL
1219.5664 Da
101 − 110
IKQAYTQFGGKRPFGVSL
1997.0967 Da
117 − 134
YIGWDKHYGFQL
1526.7427 Da
136 − 147
LKQDYKEGEMTL
1454.7196 Da
175 − 186
CDIKQAYTQF
1216.5667 Da
115 − 124
IKQAYTQFGGKRPFGVSLL
2110.1808 Da
117 − 135
IGWDKHYGFQL
1363.6793 Da
137 − 147
KQDYKEGEMTL
1341.6355 Da
176 − 186
CDIKQAYTQFGGKRPFGVSL
2215.1328 Da
115 − 134
TQFGGKRPFGVSL
1393.7587 Da
122 − 134
IGWDKHYGFQLY
1526.7427 Da
137 − 148
KSALAL
602.3872 Da
187 − 192
Relative Deuterium Uptake
Exposure (minutes)
State
i20S
i20S + PA28ab
i20S + PA28g
std20S
std20S + PA28ab
std20S + PA28g

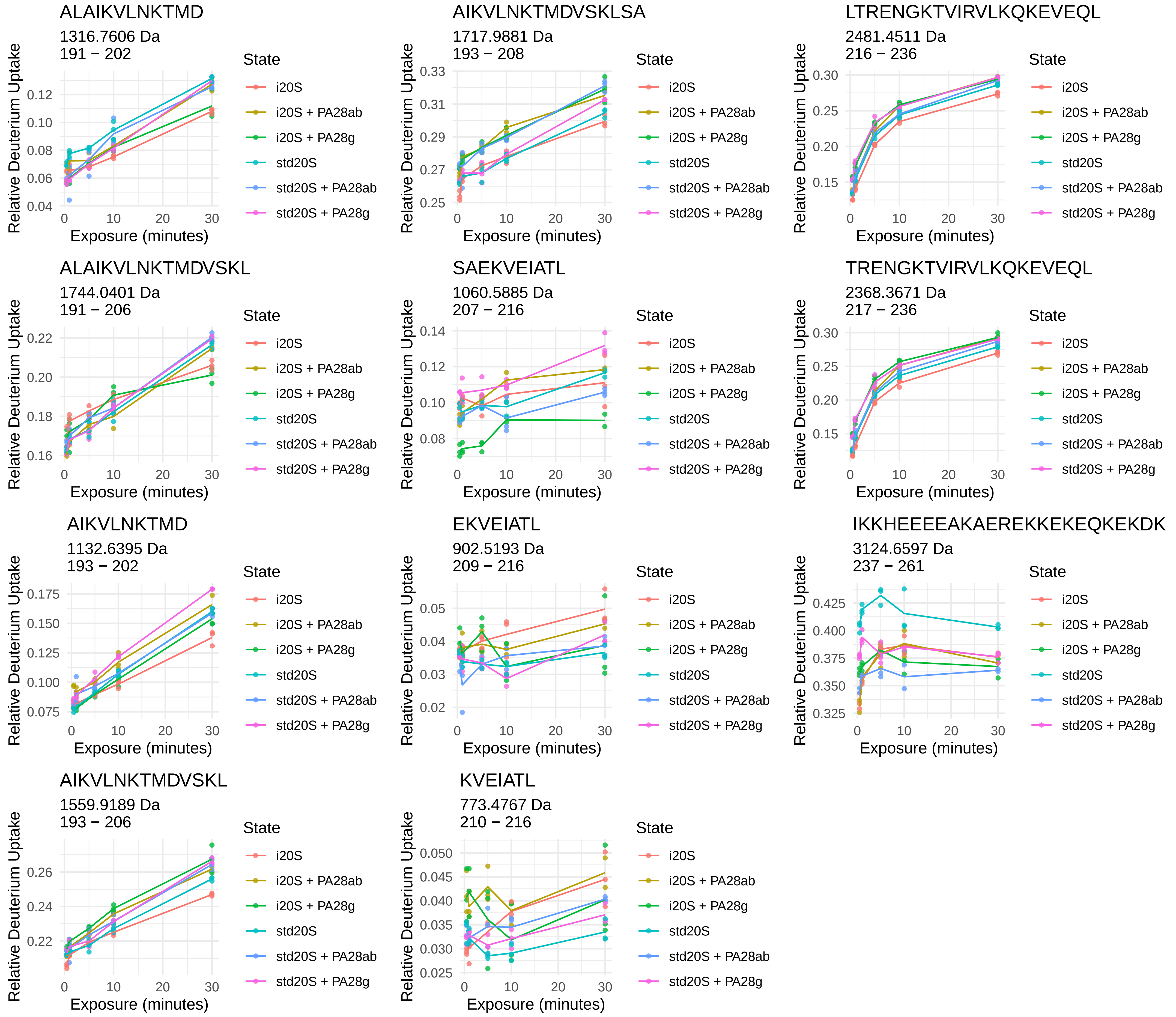
ALAIKVLNKTMD
1316.7606 Da
191 – 202
AIKVLNKTMDVSKLSA
1717.9881 Da
193 – 208
LTRENGKTVIRVLKQKEVEQL
2481.4511 Da
216 – 236
ALAIKVLNKTMDVSKL
1744.0401 Da
191 – 206
SAEKVEIATL
1060.5885 Da
207 – 216
TRENGKTVIRVLKQKEVEQL
2368.3671 Da
217 – 236
AIKVLNKTMD
1132.6395 Da
193 – 202
EKVEIATL
902.5193 Da
209 – 216
IKKHEEEEAKAEREKKEKEQKEKDK
3124.6597 Da
237 – 261
AIKVLNKTMDVSKL
1559.9189 Da
193 – 206
KVEIATL
773.4767 Da
210 – 216
Relative Deuterium Uptake
Exposure (minutes)
State
i20S
i20S + PA28ab
i20S + PA28g
std20S
std20S + PA28ab
std20S + PA28g

α4



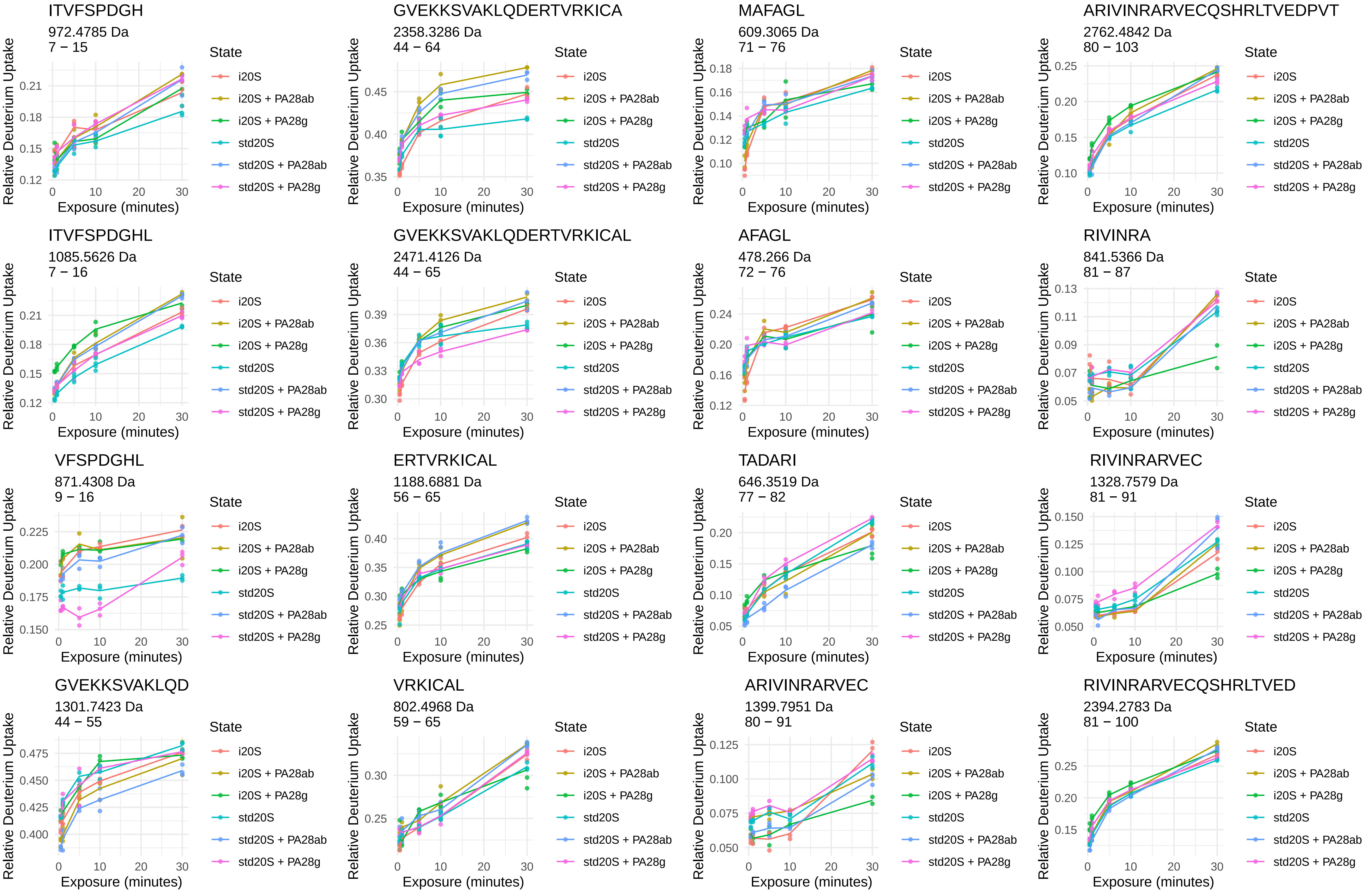
ITVFSPDGH
972.4785 Da
7 − 15
GVEKKSVAKLQDERTVRKICA
2358.3286 Da
44 − 64
MAFAGL
609.3065 Da
71 − 76
ARIVINRARVECQSHRLTVEDPVT
2762.4842 Da
80 − 103
ITVFSPDGHL
1085.5626 Da
7 − 16
GVEKKSVAKLQDERTVRKICAL
2471.4126 Da
44 − 65
AFAGL
478.266 Da
72 − 76
RIVINRA
841.5366 Da
81 − 87
VFSPDGHL
871.4308 Da
9 − 16
ERTVRKICAL
1188.6881 Da
56 − 65
TADARI
646.3519 Da
77 − 82
RIVINRARVEC
1328.7579 Da
81 − 91
GVEKKSVAKLQD
1301.7423 Da
44 − 55
VRKICAL
802.4968 Da
59 − 65
ARIVINRARVEC
1399.7951 Da
80 − 91
RIVINRARVECQSHRLTVED
2394.2783 Da
81 − 100
Relative Deuterium Uptake
Exposure (minutes)
State
i20S
i20S + PA28ab
i20S + PA28g
std20S
std20S + PA28ab
std20S + PA28g

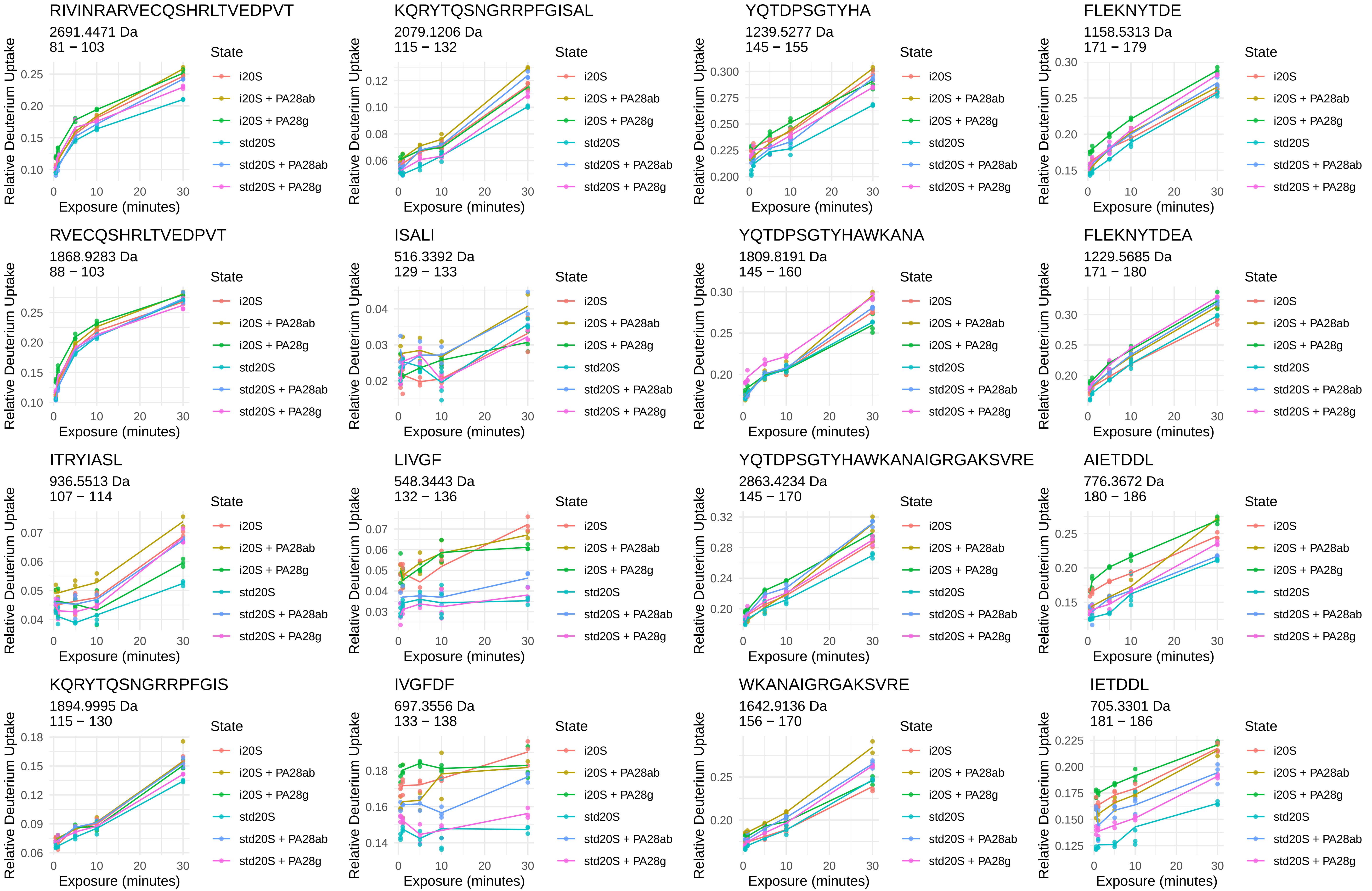
RIVINRARVECQSHRLTVEDPVT
2691.4471 Da
81 − 103
KQRYTQSNGRRPFGISAL
2079.1206 Da
115 − 132
YQTDPSGTYHA
1239.5277 Da
145 − 155
FLEKNYTDE
1158.5313 Da
171 − 179
RVECQSHRLTVEDPVT
1868.9283 Da
88 − 103
ISALI
516.3392 Da
129 − 133
YQTDPSGTYHAWKANA
1809.8191 Da
145 − 160
FLEKNYTDEA
1229.5685 Da
171 − 180
ITRYIASL
936.5513 Da
107 − 114
LIVGF
548.3443 Da
132 − 136
YQTDPSGTYHAWKANAIGRGAKSVRE
2863.4234 Da
145 − 170
AIETDDL
776.3672 Da
180 − 186
KQRYTQSNGRRPFGIS
1894.9995 Da
115 − 130
IVGFDF
697.3556 Da
133 − 138
WKANAIGRGAKSVRE
1642.9136 Da
156 − 170
IETDDL
705.3301 Da
181 − 186
Relative Deuterium Uptake
Exposure (minutes)
State
i20S
i20S + PA28ab
i20S + PA28g
std20S
std20S + PA28ab
std20S + PA28g

VIKALL
656.4705 Da
191 − 196

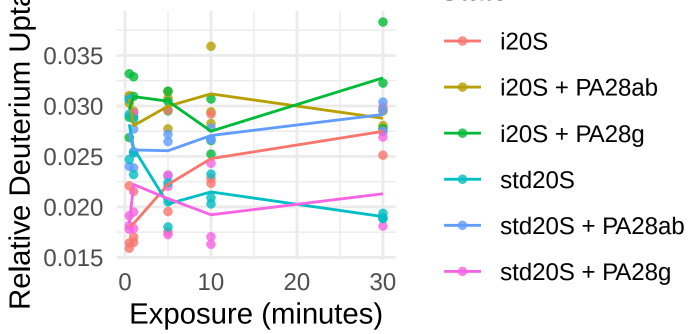


KILNPEE
842.4618 Da
218 − 224

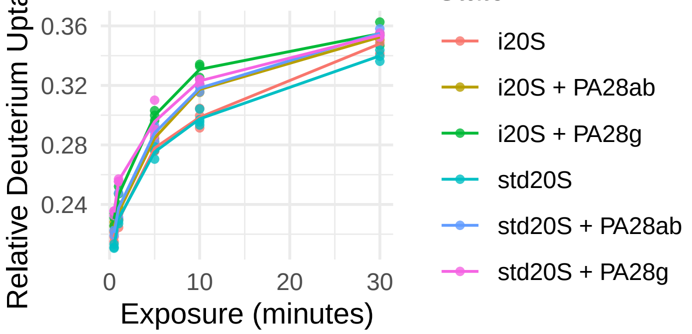


VIKALLE
785.5131 Da
191 − 197

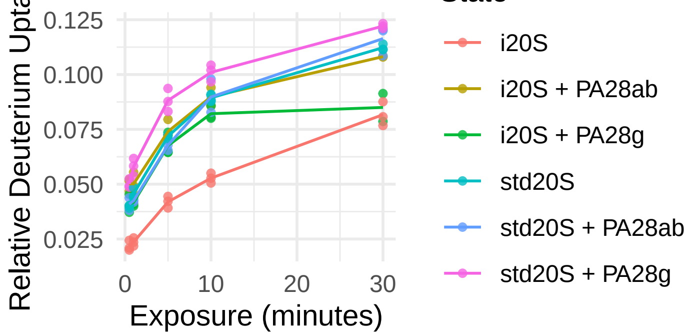


IEKYVAE
851.4509 Da
225 − 231

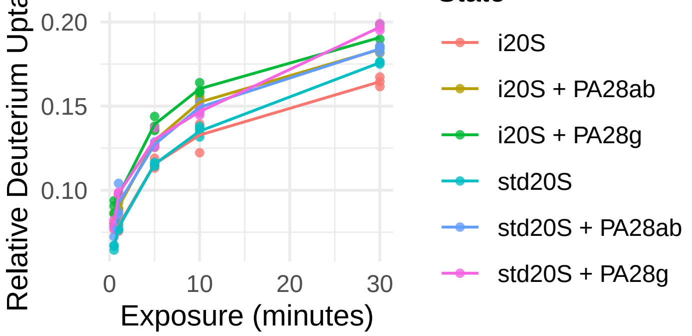


EVVQSGGKNIE
1159.5953 Da
197 − 207

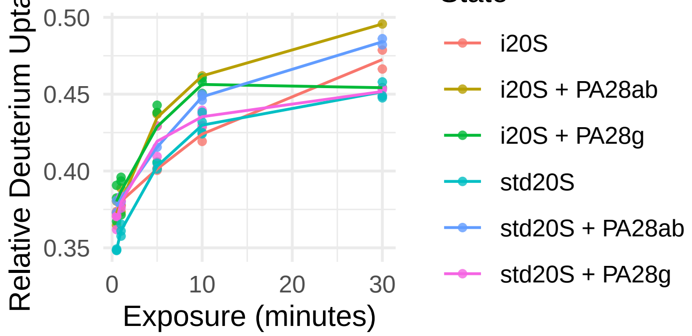


AVMRRDQSL
1075.5677 Da
209 − 217

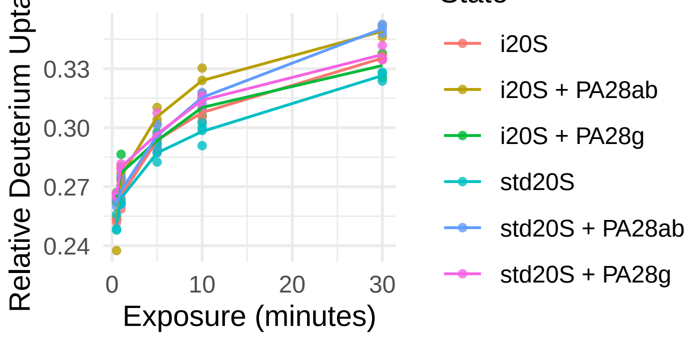

α5

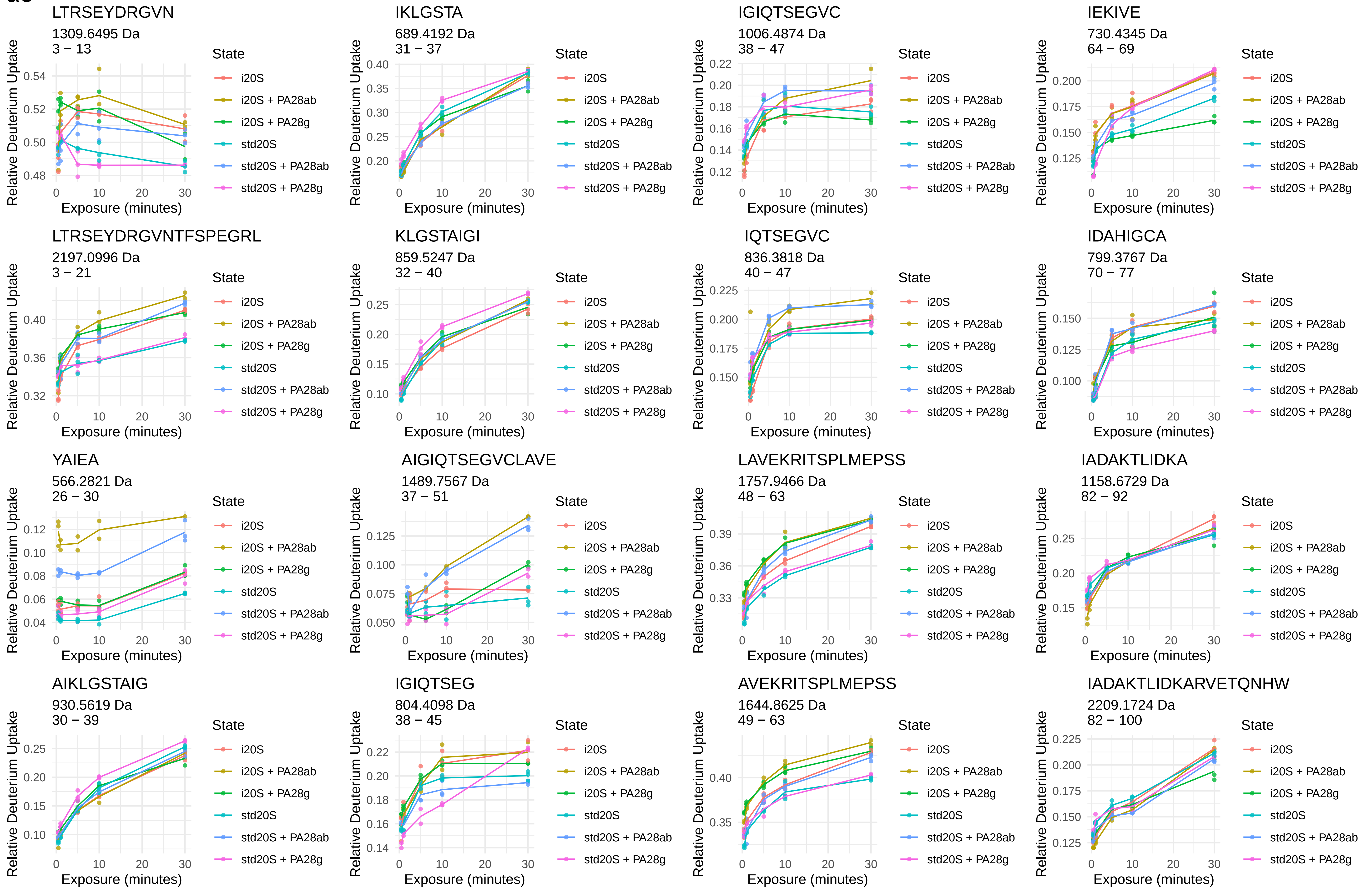
LTRSEYDRGVN
1309.6495 Da
3 − 13
IKLGSTA
689.4192 Da
31 − 37
IGIQTSEGVC
1006.4874 Da
38 − 47
IEKIVE
730.4345 Da
64 − 69
LTRSEYDRGVNTFSPEGRL
2197.0996 Da
3 − 21
KLGSTAIGI
859.5247 Da
32 − 40
IQTSEGVC
836.3818 Da
40 − 47
IDAHIGCA
799.3767 Da
70 − 77
YAIEA
566.2821 Da
26 − 30
AIGIQTSEGVCLAVE
1489.7567 Da
37 − 51
LAVEKRITSPLMEPSS
1757.9466 Da
48 − 63
IADAKTLIDKA
1158.6729 Da
82 − 92
AIKLGSTAIG
930.5619 Da
30 − 39
IGIQTSEG
804.4098 Da
38 − 45
AVEKRITSPLMEPSS
1644.8625 Da
49 − 63
IADAKTLIDKARVETQNHW
2209.1724 Da
82 − 100
Relative Deuterium Uptake
Exposure (minutes)
State
i20S
i20S + PA28ab
i20S + PA28g
std20S
std20S + PA28ab
std20S + PA28g

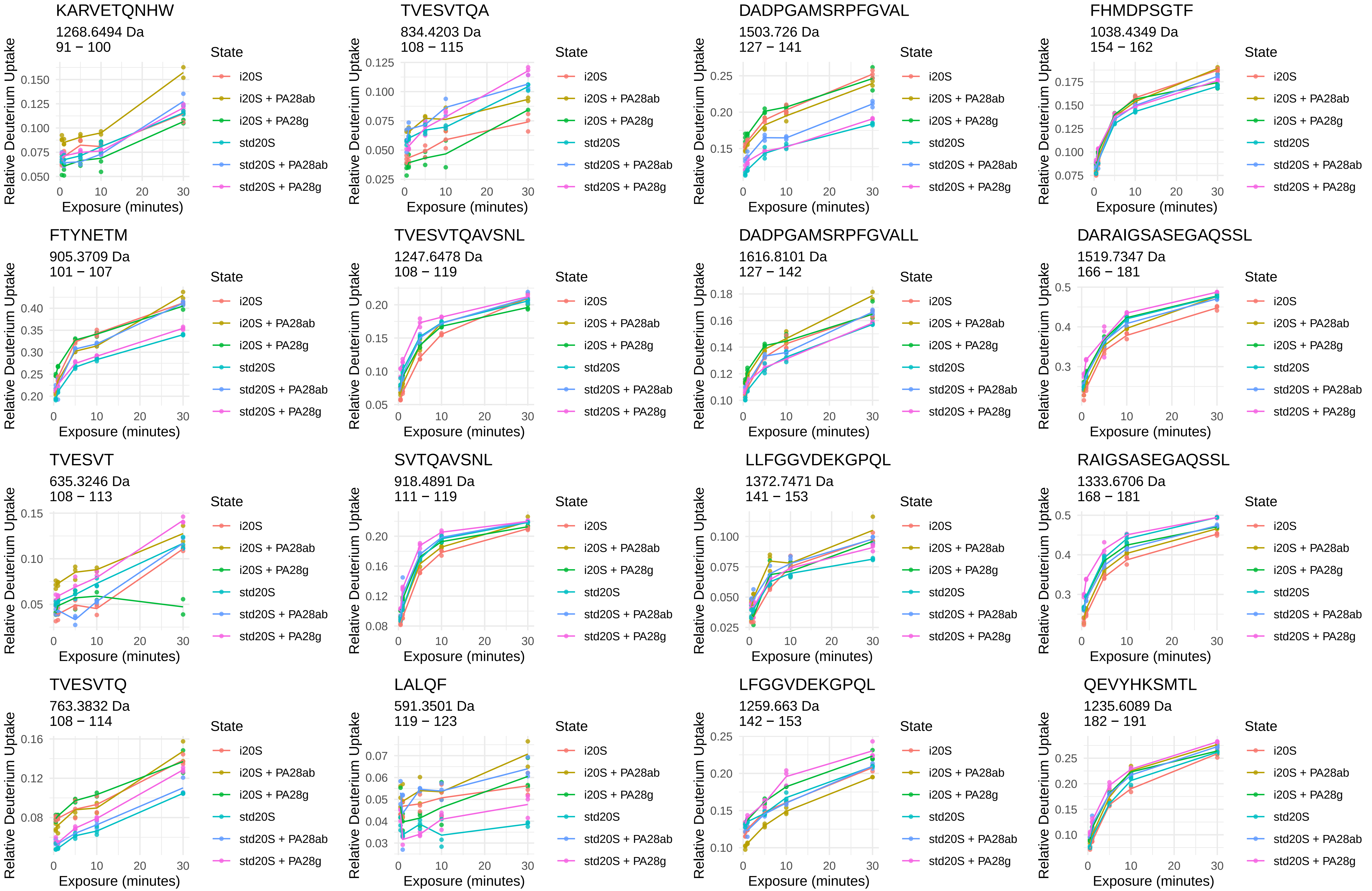
KARVETQNHW
1268.6494 Da
91 − 100
TVESVTQA
834.4203 Da
108 − 115
DADPGAMSRPFGVAL
1503.726 Da
127 − 141
FHMDPSGTF
1038.4349 Da
154 − 162
FTYNETM
905.3709 Da
101 − 107
TVESVTQAVSNL
1247.6478 Da
108 − 119
DADPGAMSRPFGVALL
1616.8101 Da
127 − 142
DARAIGSASEGAQSSL
1519.7347 Da
166 − 181
TVESVT
635.3246 Da
108 − 113
SVTQAVSNL
918.4891 Da
111 − 119
LLFGGVDEKGPQL
1372.7471 Da
141 − 153
RAIGSASEGAQSSL
1333.6706 Da
168 − 181
TVESVTQ
763.3832 Da
108 − 114
LALQF
591.3501 Da
119 − 123
LFGGVDEKGPQL
1259.663 Da
142 − 153
QEVYHKSMTL
1235.6089 Da
182 − 191
Relative Deuterium Uptake
Exposure (minutes)
State
i20S
i20S + PA28ab
i20S + PA28g
std20S
std20S + PA28ab
std20S + PA28g

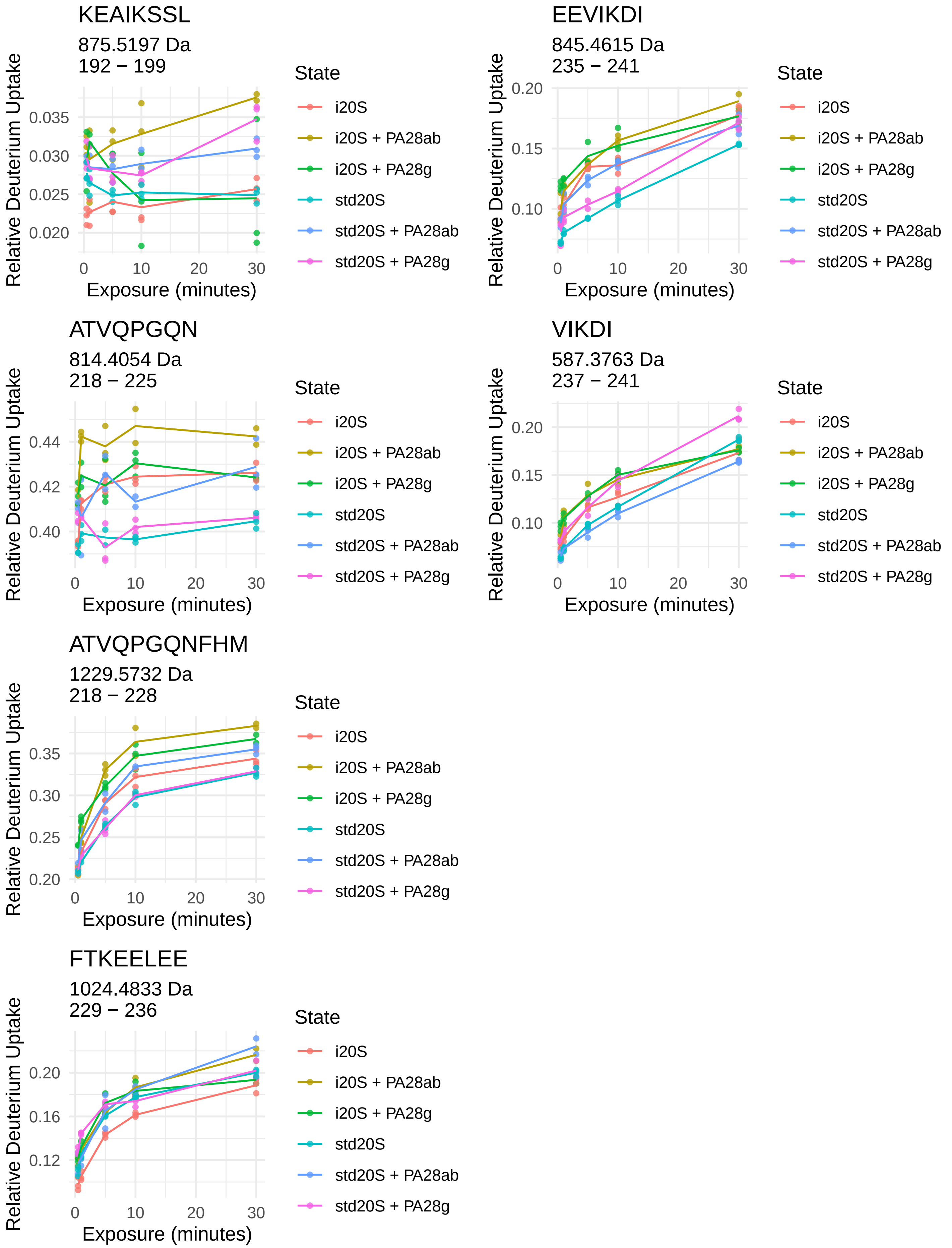
KEAIKSSL
875.5197 Da
192 − 199
EEVIKDI
845.4615 Da
235 − 241
ATVQPGQN
814.4054 Da
218 − 225
VIKDI
587.3763 Da
237 − 241
ATVQPGQNFHM
1229.5732 Da
218 − 228
FTKEELEE
1024.4833 Da
229 − 236
Relative Deuterium Uptake
Exposure (minutes)
State
i20S
i20S + PA28ab
i20S + PA28g
std20S
std20S + PA28ab
std20S + PA28g

# α6



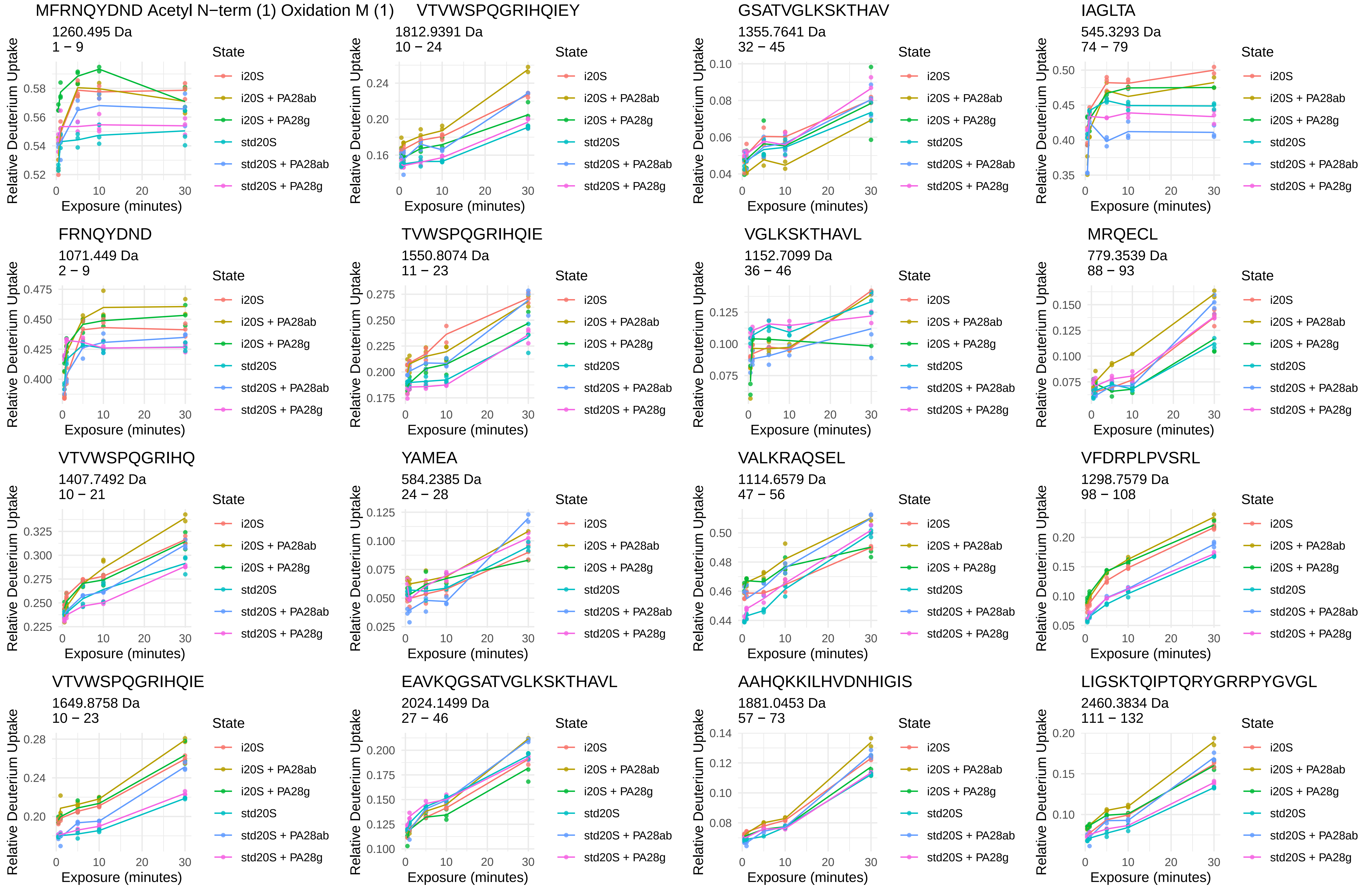
MFRNQYDND Acetyl N−term (1) Oxidation M (1)
1260.495 Da
1 − 9
VTVWSPQGRIHQIEY
1812.9391 Da
10 − 24
GSATVGLKSKTHAV
1355.7641 Da
32 − 45
IAGLTA
545.3293 Da
74 − 79
FRNQYDND
1071.449 Da
2 − 9
TVWSPQGRIHQIE
1550.8074 Da
11 − 23
VGLKSKTHAVL
1152.7099 Da
36 − 46
MRQECL
779.3539 Da
88 − 93
VTVWSPQGRIHQ
1407.7492 Da
10 − 21
YAMEA
584.2385 Da
24 − 28
VALKRAQSEL
1114.6579 Da
47 − 56
VFDRPLPVSRL
1298.7579 Da
98 − 108
VTVWSPQGRIHQIE
1649.8758 Da
10 − 23
EAVKQGSATVGLKSKTHAVL
2024.1499 Da
27 − 46
AAHQKKILHVDNHIGIS
1881.0453 Da
57 − 73
LIGSKTQIPTQRYGRRPYGVGL
2460.3834 Da
111 − 132
Relative Deuterium Uptake
Exposure (minutes)
State
i20S
i20S + PA28ab
i20S + PA28g
std20S
std20S + PA28ab
std20S + PA28g

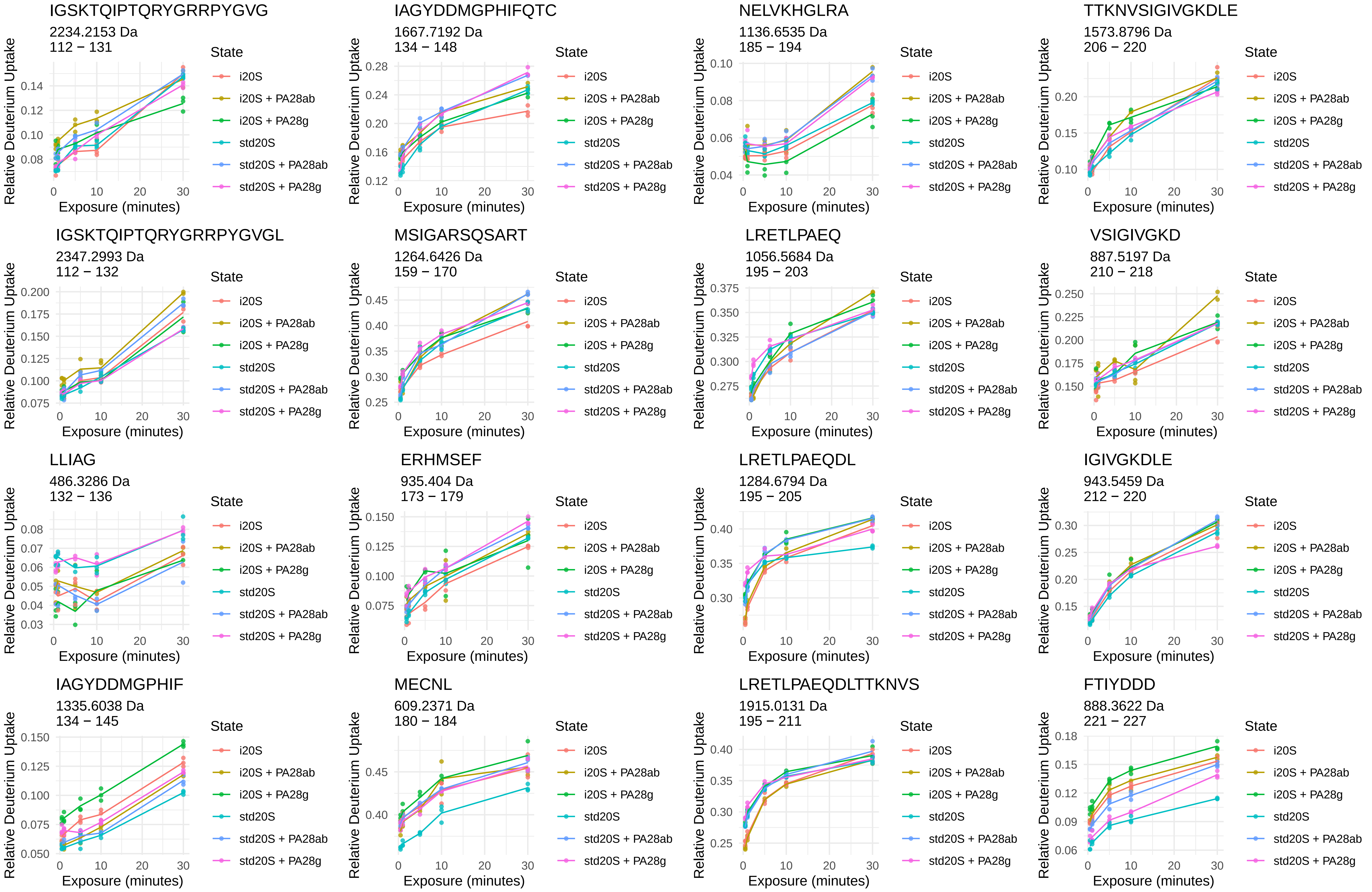
IGSKTQIPTQRYGRRPYGVG
2234.2153 Da
112 − 131
IAGYDDMGPHIFQTC
1667.7192 Da
134 − 148
NELVKHGLRA
1136.6535 Da
185 − 194
TTKNVSIGIVGKDLE
1573.8796 Da
206 − 220
IGSKTQIPTQRYGRRPYGVGL
2347.2993 Da
112 − 132
MSIGARSQSART
1264.6426 Da
159 − 170
LRETLPAEQ
1056.5684 Da
195 − 203
VSIGIVGKD
887.5197 Da
210 − 218
LLIAG
486.3286 Da
132 − 136
ERHMSEF
935.404 Da
173 − 179
LRETLPAEQDL
1284.6794 Da
195 − 205
IGIVGKDLE
943.5459 Da
212 − 220
IAGYDDMGPHIF
1335.6038 Da
134 − 145
MECNL
609.2371 Da
180 − 184
LRETLPAEQDLTTKNVS
1915.0131 Da
195 − 211
FTIYDDD
888.3622 Da
221 − 227
Relative Deuterium Uptake
Exposure (minutes)
State
i20S
i20S + PA28ab
i20S + PA28g
std20S
std20S + PA28ab
std20S + PA28g

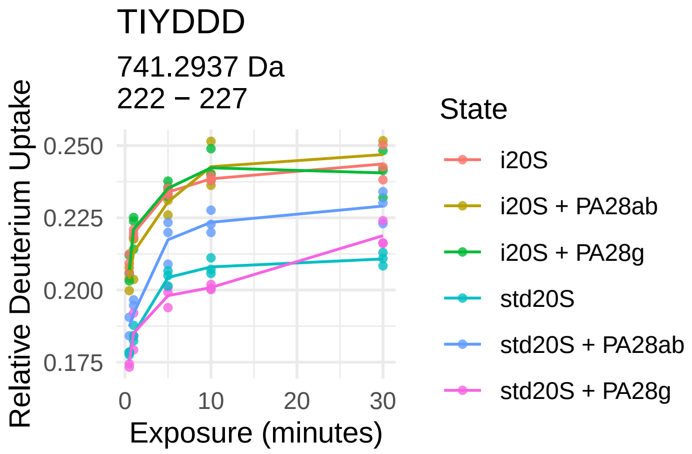
TIYDDD
741.2937 Da
222 − 227
Relative Deuterium Uptake
0.250
0.225
0.200
0.175
0
10
20
30
Exposure (minutes)
State
i20S
i20S + PA28ab
i20S + PA28g
std20S
std20S + PA28ab
std20S + PA28g

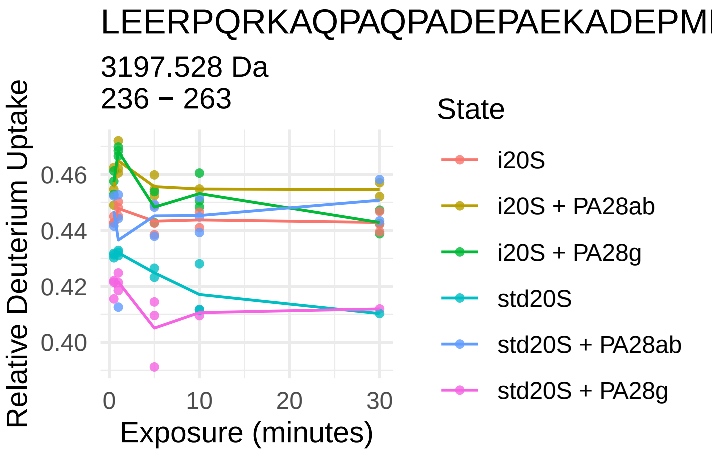
LEERPQRKAQPAQPADEPAEKADEPMEH
3197.528 Da
236 − 263
Relative Deuterium Uptake
0.46
0.44
0.42
0.40
0
10
20
30
Exposure (minutes)
State
i20S
i20S + PA28ab
i20S + PA28g
std20S
std20S + PA28ab
std20S + PA28g

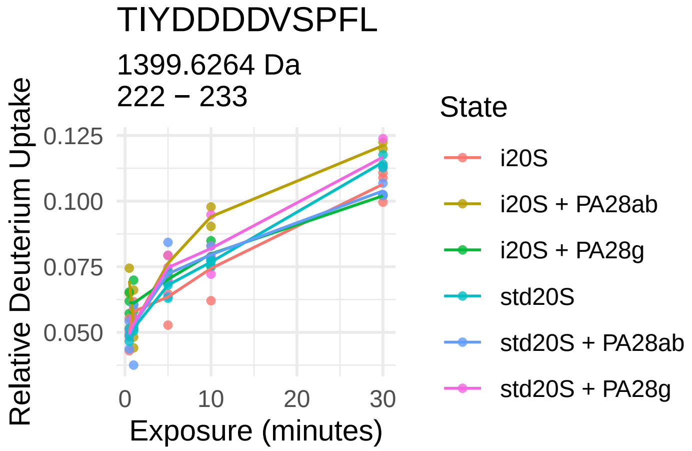
TIYDDDDVSPFL
1399.6264 Da
222 − 233
Relative Deuterium Uptake
0.125
0.100
0.075
0.050
0
10
20
30
Exposure (minutes)
State
i20S
i20S + PA28ab
i20S + PA28g
std20S
std20S + PA28ab
std20S + PA28g

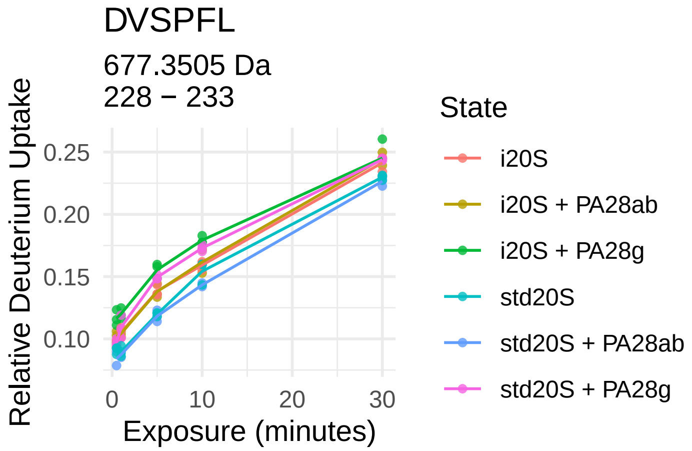
DVSPFL
677.3505 Da
228 − 233
Relative Deuterium Uptake
0.25
0.20
0.15
0.10
0
10
20
30
Exposure (minutes)
State
i20S
i20S + PA28ab
i20S + PA28g
std20S
std20S + PA28ab
std20S + PA28g

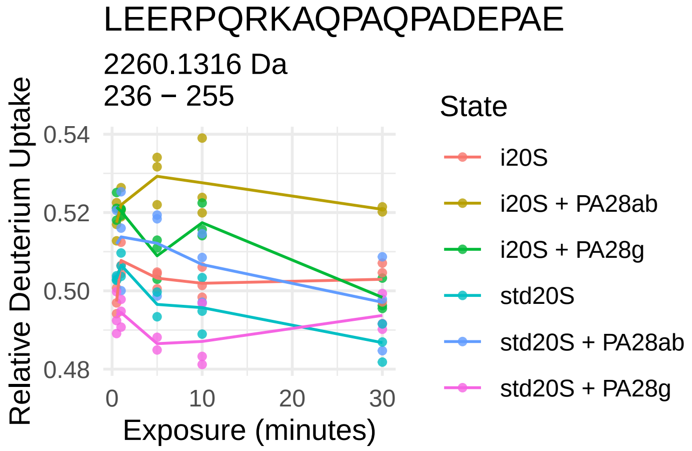
LEERPQRKAQPAQPADEPAE
2260.1316 Da
236 − 255
Relative Deuterium Uptake
0.54
0.52
0.50
0.48
0
10
20
30
Exposure (minutes)
State
i20S
i20S + PA28ab
i20S + PA28g
std20S
std20S + PA28ab
std20S + PA28g

α7

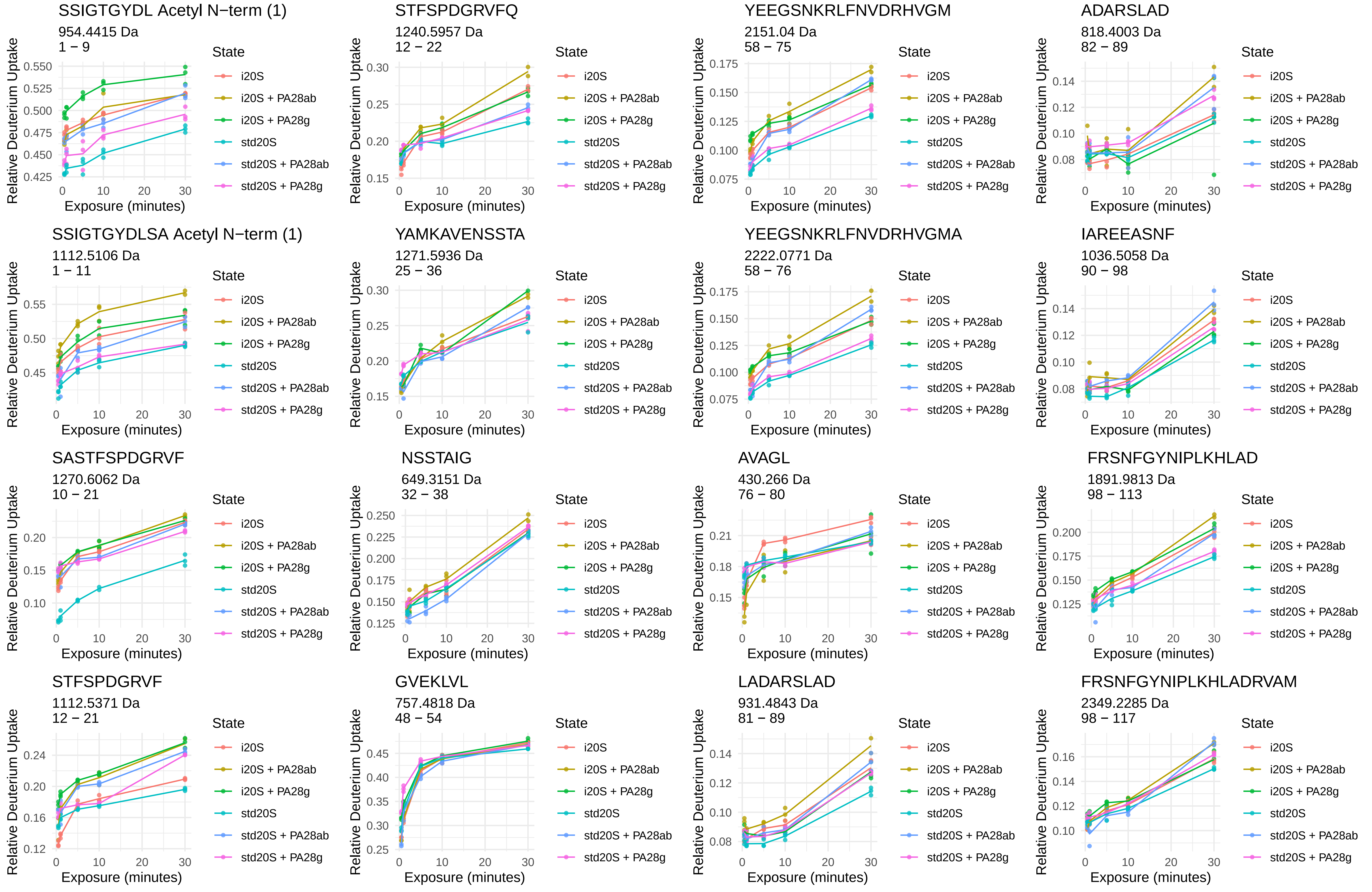
SSIGTGYDL Acetyl N−term (1)
954.4415 Da
1 − 9
STFSPDGRVFQ
1240.5957 Da
12 − 22
YEEGSNKRLFNVDRHVGM
2151.04 Da
58 − 75
ADARSLAD
818.4003 Da
82 − 89
SSIGTGYDLSA Acetyl N−term (1)
1112.5106 Da
1 − 11
YAMKAVENSSTA
1271.5936 Da
25 − 36
YEEGSNKRLFNVDRHVGMA
2222.0771 Da
58 − 76
IAREEASNF
1036.5058 Da
90 − 98
SASTFSPDGRVF
1270.6062 Da
10 − 21
NSSTAIG
649.3151 Da
32 − 38
AVAGL
430.266 Da
76 − 80
FRSNFGYNIPLKHLAD
1891.9813 Da
98 − 113
STFSPDGRVF
1112.5371 Da
12 − 21
GVEKLVL
757.4818 Da
48 − 54
LADARSLAD
931.4843 Da
81 − 89
FRSNFGYNIPLKHLADRVAM
2349.2285 Da
98 − 117
Relative Deuterium Uptake
Exposure (minutes)
State
i20S
i20S + PA28ab
i20S + PA28g
std20S
std20S + PA28ab
std20S + PA28g

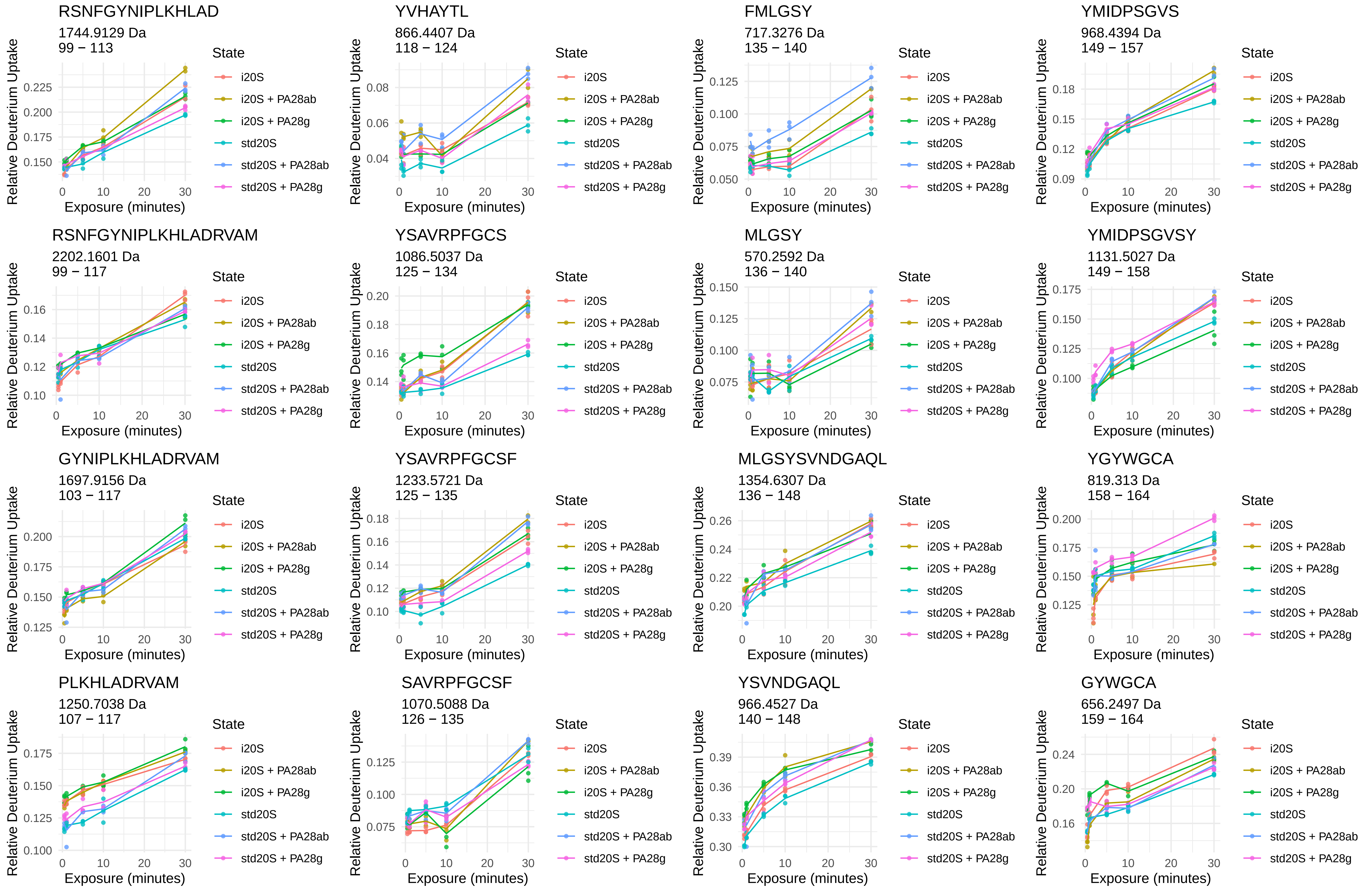

RSNFGYNIPLKHLAD
1744.9129 Da
99 − 113
YVHAYTL
866.4407 Da
118 − 124
FMLGSY
717.3276 Da
135 − 140
YMIDPSGVS
968.4394 Da
149 − 157
RSNFGYNIPLKHLADRVAM
2202.1601 Da
99 − 117
YSAVRPFGCS
1086.5037 Da
125 − 134
MLGSY
570.2592 Da
136 − 140
YMIDPSGVSY
1131.5027 Da
149 − 158
GYNIPLKHLADRVAM
1697.9156 Da
103 − 117
YSAVRPFGCSF
1233.5721 Da
125 − 135
MLGSYSVNDGAQL
1354.6307 Da
136 − 148
YGYWGCA
819.313 Da
158 − 164
PLKHLADRVAM
1250.7038 Da
107 − 117
SAVRPFGCSF
1070.5088 Da
126 − 135
YSVNDGAQL
966.4527 Da
140 − 148
GYWGCA
656.2497 Da
159 − 164
Relative Deuterium Uptake
Exposure (minutes)
State
i20S
i20S + PA28ab
i20S + PA28g
std20S
std20S + PA28ab
std20S + PA28g

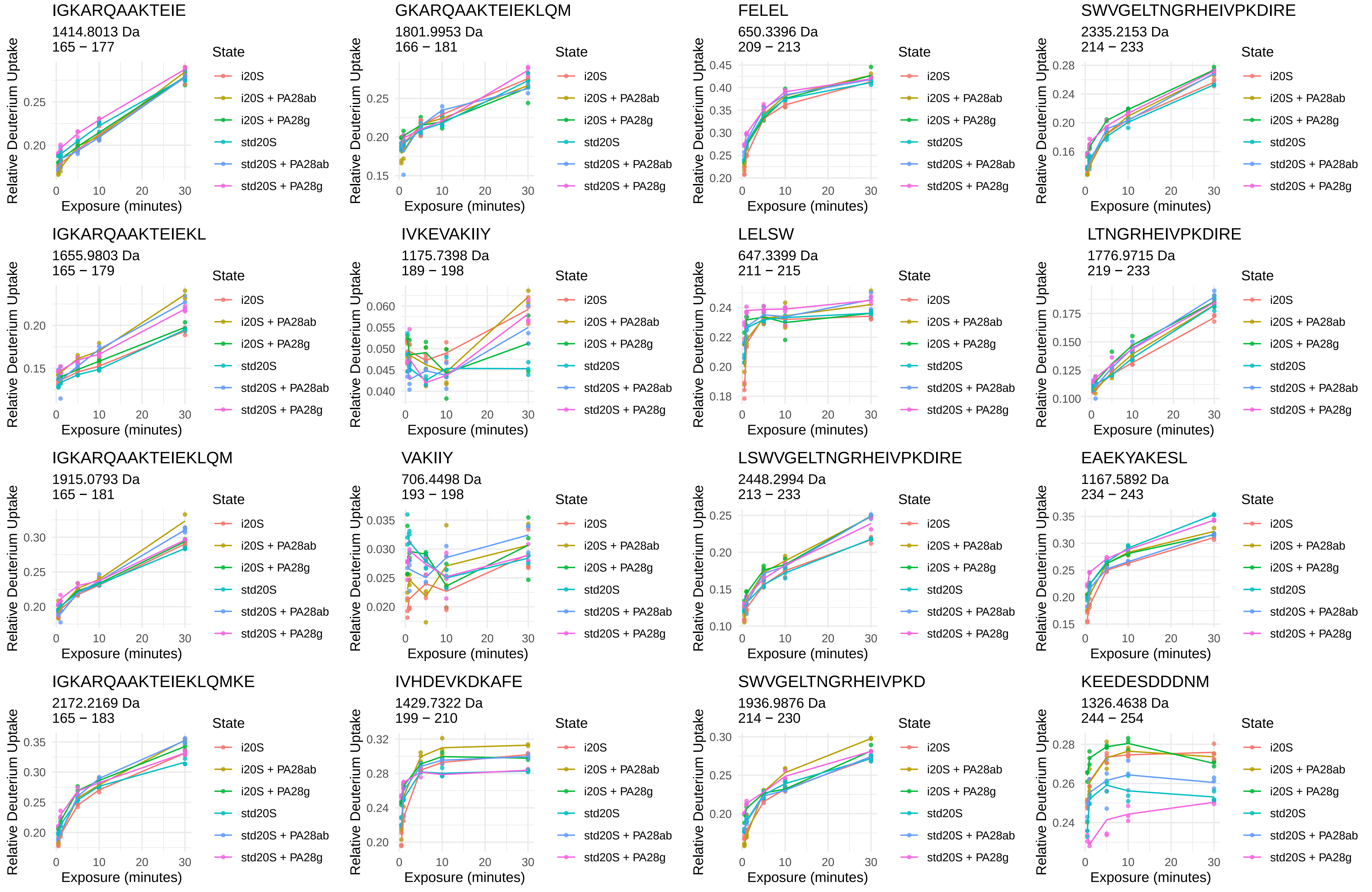
IGKARQAAKTEIE
1414.8013 Da
165 − 177
GKARQAAKTEIEKLQM
1801.9953 Da
166 − 181
FELEL
650.3396 Da
209 − 213
SWVGELTNGRHEIVPKDIRE
2335.2153 Da
214 − 233
IGKARQAAKTEIEKL
1655.9803 Da
165 − 179
IVKEVAKIIY
1175.7398 Da
189 − 198
LELSW
647.3399 Da
211 − 215
LTNGRHEIVPKDIRE
1776.9715 Da
219 − 233
IGKARQAAKTEIEKLQM
1915.0793 Da
165 − 181
VAKIIY
706.4498 Da
193 − 198
LSWVGELTNGRHEIVPKDIRE
2448.2994 Da
213 − 233
EAEKYAKESL
1167.5892 Da
234 − 243
IGKARQAAKTEIEKLQMKE
2172.2169 Da
165 − 183
IVHDEVKDKAFE
1429.7322 Da
199 − 210
SWVGELTNGRHEIVPKD
1936.9876 Da
214 − 230
KEEDESDDDNM
1326.4638 Da
244 − 254
Relative Deuterium Uptake
Exposure (minutes)
State
i20S
i20S + PA28ab
i20S + PA28g
std20S
std20S + PA28ab
std20S + PA28g

β1

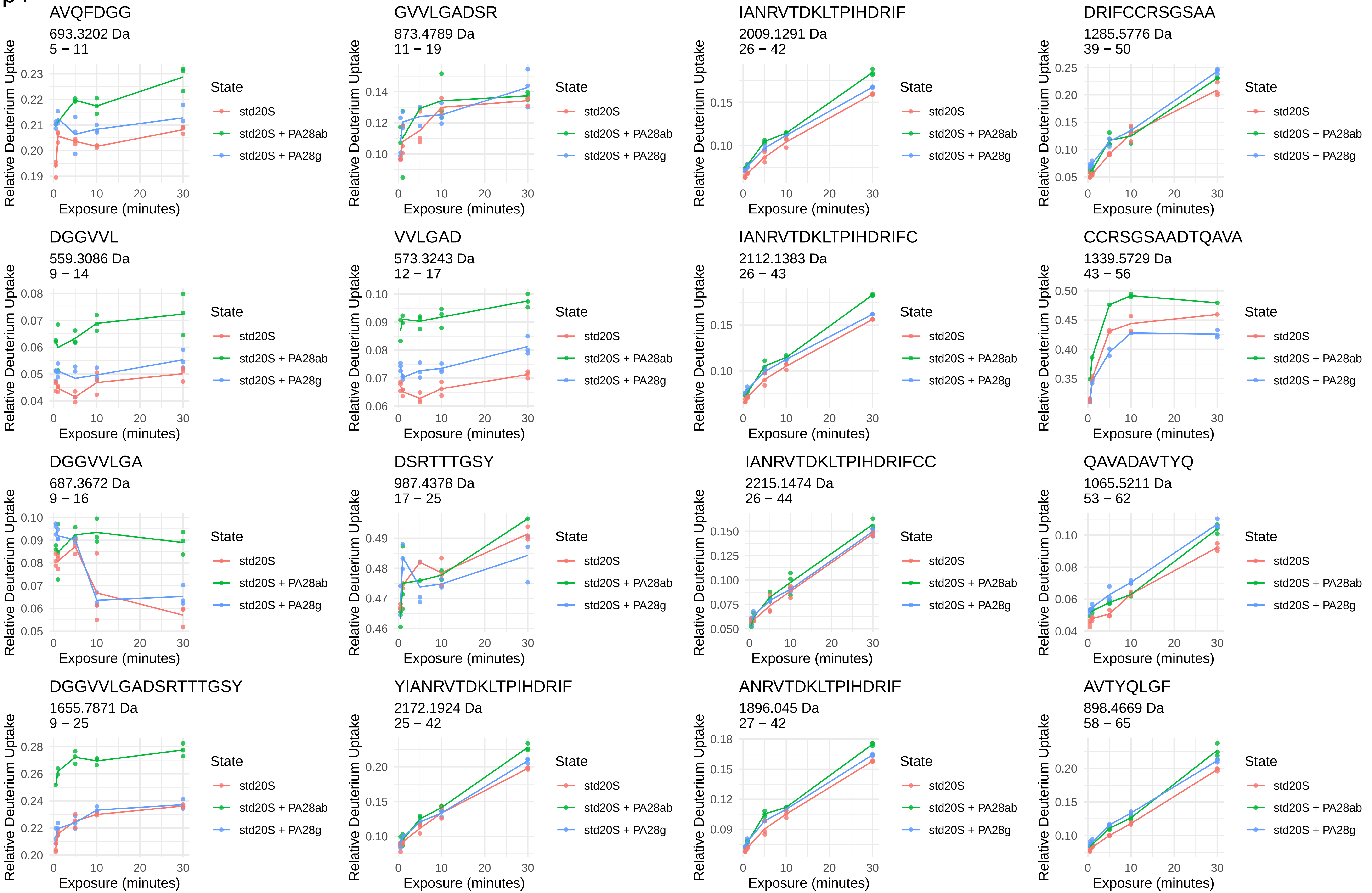
AVQFDGG
693.3202 Da
5 − 11
GVVLGADSR
873.4789 Da
11 − 19
IANRVTDKLTPIHDRIF
2009.1291 Da
26 − 42
DRIFCCRSGSAA
1285.5776 Da
39 − 50
DGGVVL
559.3086 Da
9 − 14
VVLGAD
573.3243 Da
12 − 17
IANRVTDKLTPIHDRIFC
2112.1383 Da
26 − 43
CCRSGSAADTQAVA
1339.5729 Da
43 − 56
DGGVVLGA
687.3672 Da
9 − 16
DSRTTTGSY
987.4378 Da
17 − 25
IANRVTDKLTPIHDRIFCC
2215.1474 Da
26 − 44
QAVADAVTYQ
1065.5211 Da
53 − 62
DGGVVLGADSRTTTGSY
1655.7871 Da
9 − 25
YIANRVTDKLTPIHDRIF
2172.1924 Da
25 − 42
ANRVTDKLTPIHDRIF
1896.045 Da
27 − 42
AVTYQLGF
898.4669 Da
58 − 65
Relative Deuterium Uptake
Exposure (minutes)
State
std20S
std20S + PA28ab
std20S + PA28g

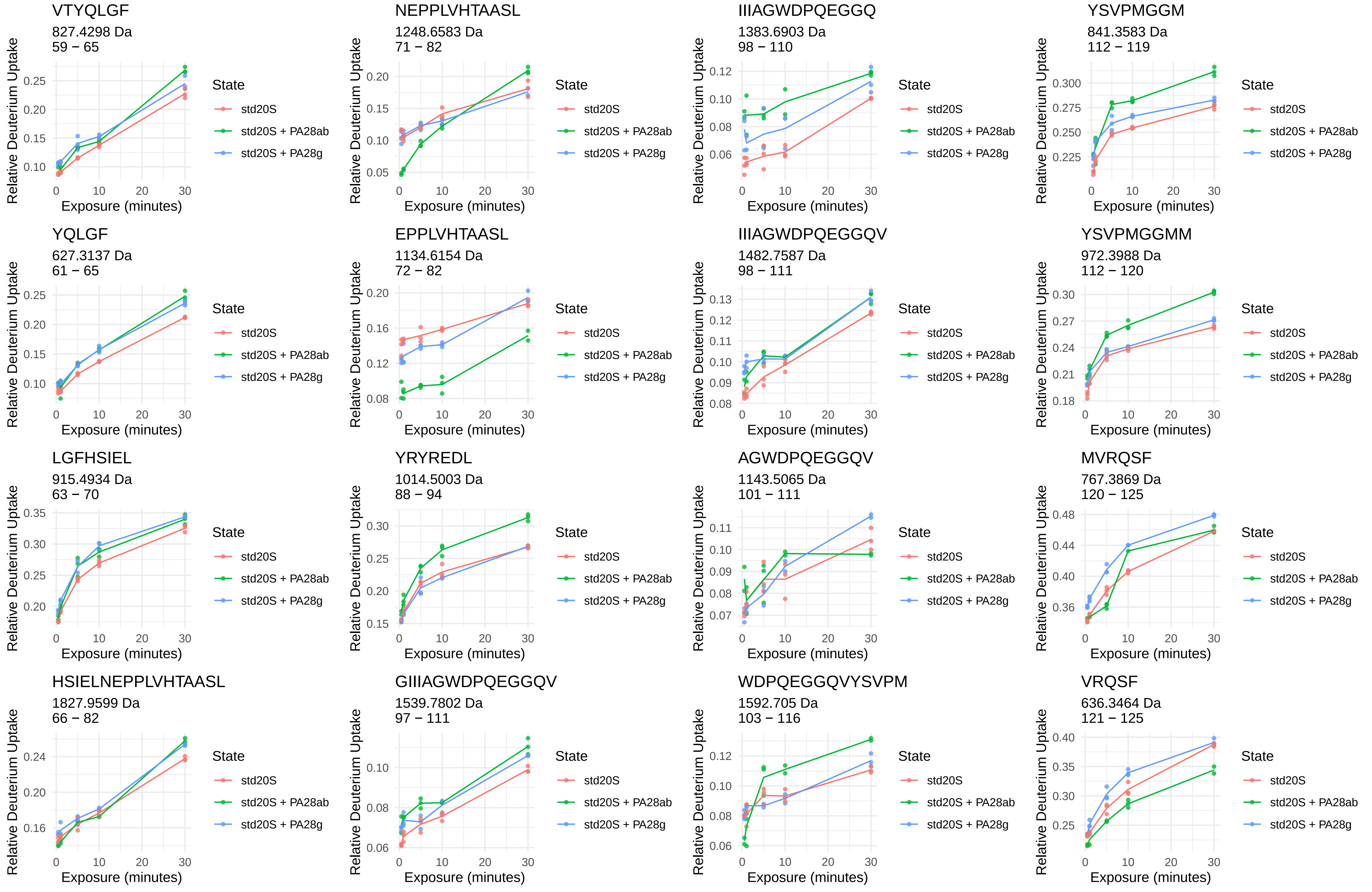
VTYQLGF
827.4298 Da
59 − 65
NEPPLVHTAASL
1248.6583 Da
71 − 82
IIIAGWDPQEGGQ
1383.6903 Da
98 − 110
YSVPMGGM
841.3583 Da
112 − 119
YQLGF
627.3137 Da
61 − 65
EPPLVHTAASL
1134.6154 Da
72 − 82
IIIAGWDPQEGGQV
1482.7587 Da
98 − 111
YSVPMGGMM
972.3988 Da
112 − 120
LGFHSIEL
915.4934 Da
63 − 70
YRYREDL
1014.5003 Da
88 − 94
AGWDPQEGGQV
1143.5065 Da
101 − 111
MVRQSF
767.3869 Da
120 − 125
HSIELNEPPLVHTAASL
1827.9599 Da
66 − 82
GIIIAGWDPQEGGQV
1539.7802 Da
97 − 111
WDPQEGGQVYSVPM
1592.705 Da
103 − 116
VRQSF
636.3464 Da
121 − 125
Relative Deuterium Uptake
Exposure (minutes)
State
std20S
std20S + PA28ab
std20S + PA28g

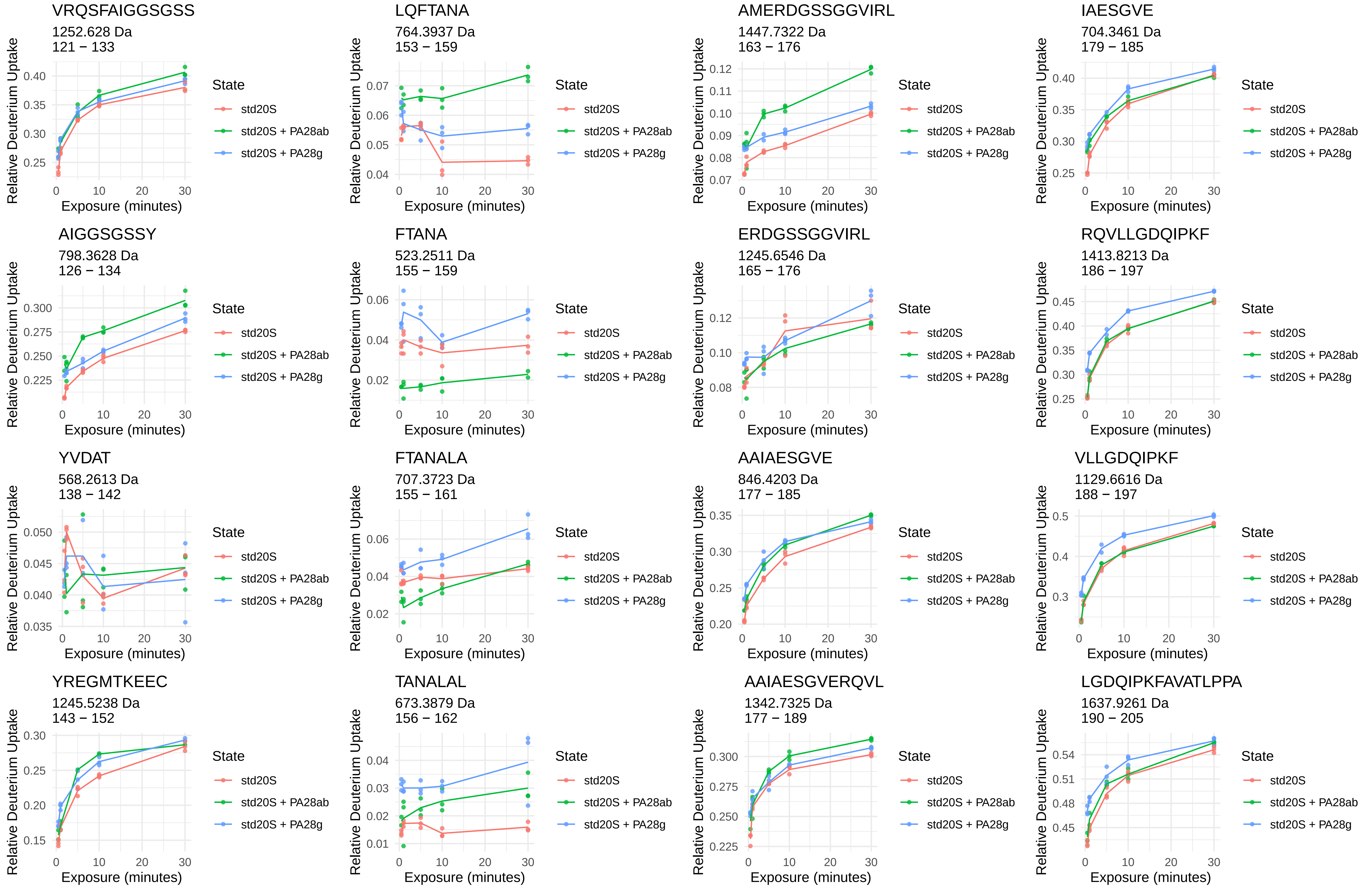
VRQSFAIGGSGSS
1252.628 Da
121 − 133
LQFTANA
764.3937 Da
153 − 159
AMERDGSSGGVIRL
1447.7322 Da
163 − 176
IAESGVE
704.3461 Da
179 − 185
AIGGSGSSY
798.3628 Da
126 − 134
FTANA
523.2511 Da
155 − 159
ERDGSSGGVIRL
1245.6546 Da
165 − 176
RQVLLGDQIPKF
1413.8213 Da
186 − 197
YVDAT
568.2613 Da
138 − 142
FTANALA
707.3723 Da
155 − 161
AAIAESGVE
846.4203 Da
177 − 185
VLLGDQIPKF
1129.6616 Da
188 − 197
YREGMTKEEC
1245.5238 Da
143 − 152
TANALAL
673.3879 Da
156 − 162
AAIAESGVERQVL
1342.7325 Da
177 − 189
LGDQIPKFAVATLPPA
1637.9261 Da
190 − 205
Relative Deuterium Uptake
Exposure (minutes)
State
std20S
std20S + PA28ab
std20S + PA28g

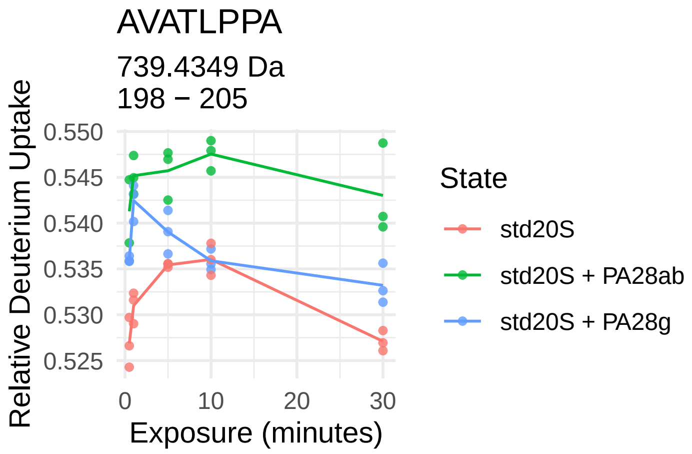
AVATLPPA
739.4349 Da
198 − 205
Relative Deuterium Uptake
0.550
0.545
0.540
0.535
0.530
0.525
0
10
20
30
Exposure (minutes)
State
std20S
std20S + PA28ab
std20S + PA28g

β1i

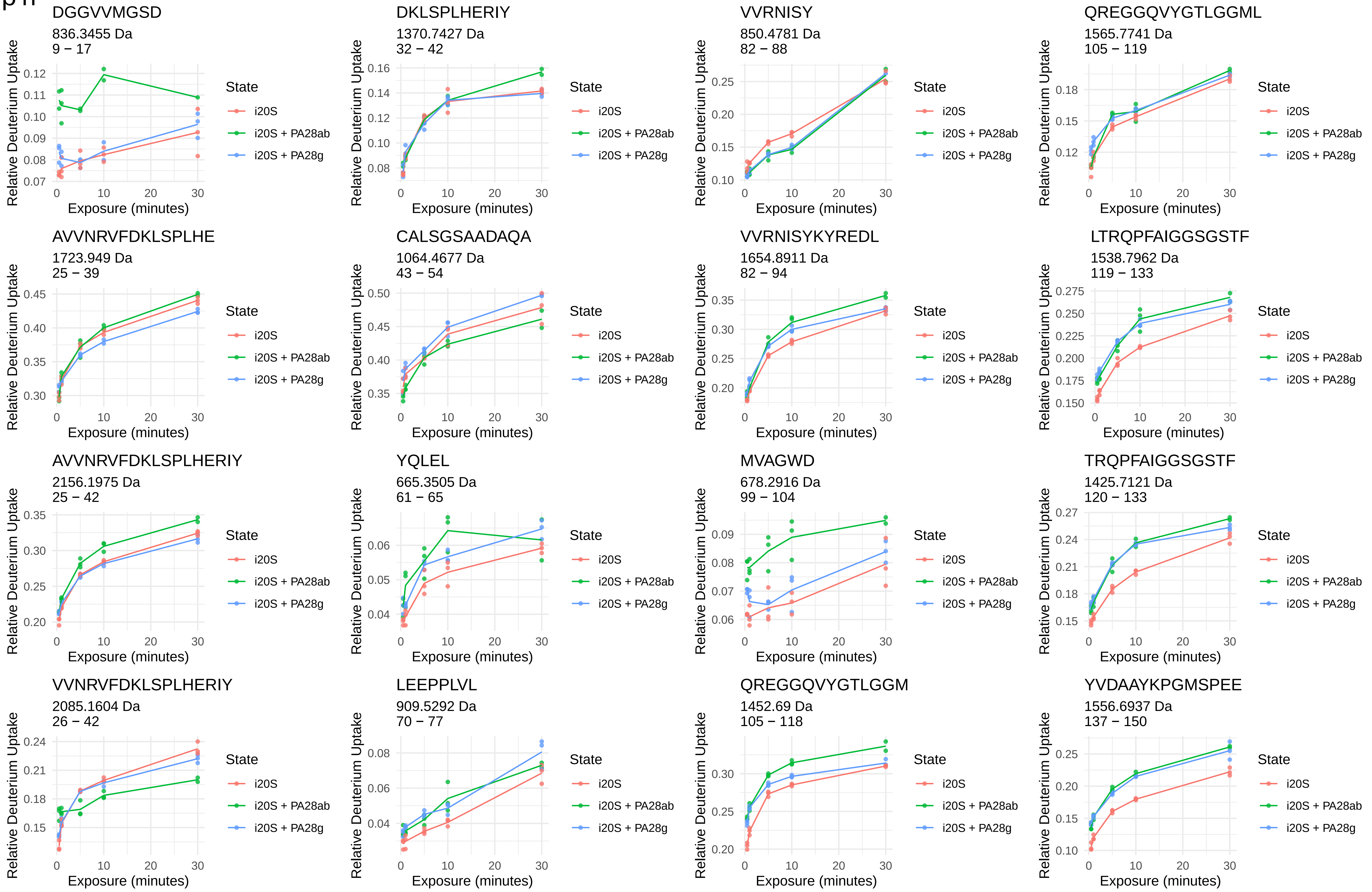
DGGVVMGSD
836.3455 Da
9 − 17
DKLSPLHERIY
1370.7427 Da
32 − 42
VVRNISY
850.4781 Da
82 − 88
QREGGQVYGTLGGML
1565.7741 Da
105 − 119
AVVNRVFDKLSPLHE
1723.949 Da
25 − 39
CALSGSAADAQA
1064.4677 Da
43 − 54
VVRNISYKYREDL
1654.8911 Da
82 − 94
LTRQPFAIGGSGSTF
1538.7962 Da
119 − 133
AVVNRVFDKLSPLHERIY
2156.1975 Da
25 − 42
YQLEL
665.3505 Da
61 − 65
MVAGWD
678.2916 Da
99 − 104
TRQPFAIGGSGSTF
1425.7121 Da
120 − 133
VVNRVFDKLSPLHERIY
2085.1604 Da
26 − 42
LEEPPLVL
909.5292 Da
70 − 77
QREGGQVYGTLGGM
1452.69 Da
105 − 118
YVDAAYKPGMSPEE
1556.6937 Da
137 − 150
Relative Deuterium Uptake
Exposure (minutes)
State
i20S
i20S + PA28ab
i20S + PA28g

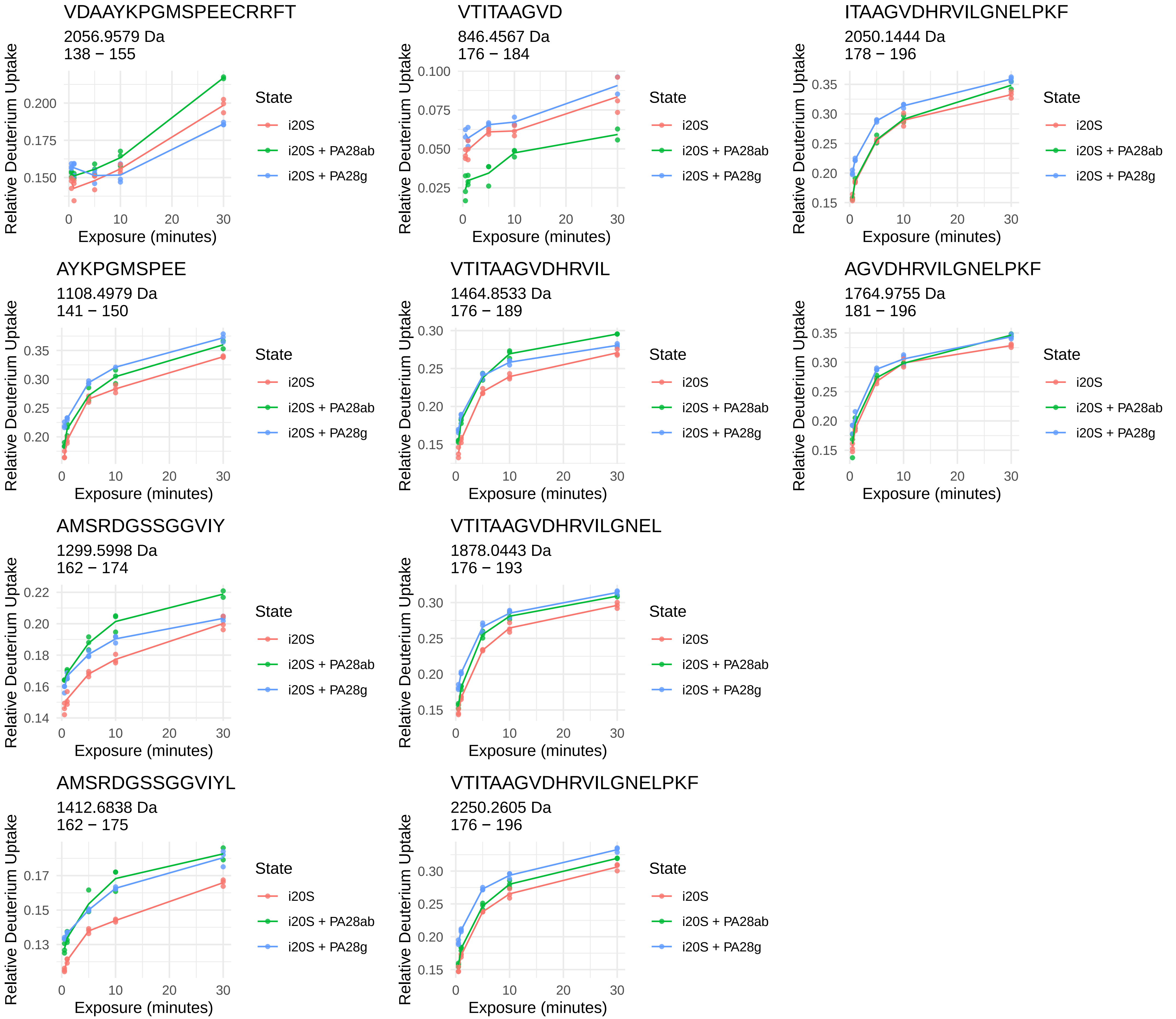
VDAAYKPGMSPEECRRFT
2056.9579 Da
138 − 155
VTITAAGVD
846.4567 Da
176 − 184
ITAAGVDHRVILGNELPKF
2050.1444 Da
178 − 196
AYKPGMSPEE
1108.4979 Da
141 − 150
VTITAAGVDHRVIL
1464.8533 Da
176 − 189
AGVDHRVILGNELPKF
1764.9755 Da
181 − 196
AMSRDGSSGGVIY
1299.5998 Da
162 − 174
VTITAAGVDHRVILGNEL
1878.0443 Da
176 − 193
AMSRDGSSGGVIYL
1412.6838 Da
162 − 175
VTITAAGVDHRVILGNELPKF
2250.2605 Da
176 − 196
Relative Deuterium Uptake
Exposure (minutes)
State
i20S
i20S + PA28ab
i20S + PA28g

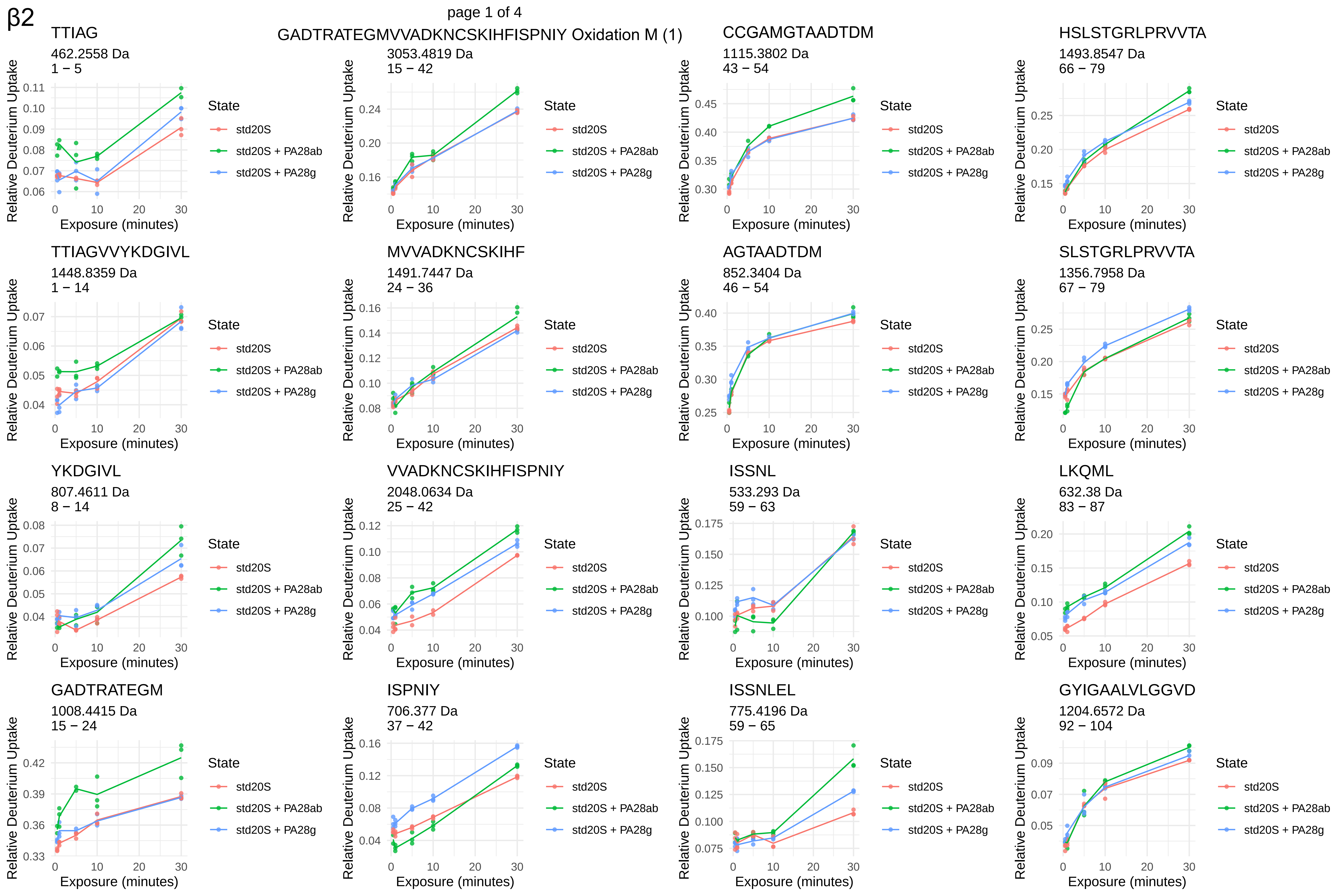
β2
page 1 of 4
TTIAG
462.2558 Da
1 − 5
GADTRATEGMVVADKNCSKIHFISPNIY Oxidation M (1)
3053.4819 Da
15 − 42
CCGAMGTAADTDM
1115.3802 Da
43 − 54
HSLSTGRLPRVVTA
1493.8547 Da
66 − 79
TTIAGVVYKDGIVL
1448.8359 Da
1 − 14
MVVADKNCSKIHF
1491.7447 Da
24 − 36
AGTAADTDM
852.3404 Da
46 − 54
SLSTGRLPRVVTA
1356.7958 Da
67 − 79
YKDGIVL
807.4611 Da
8 − 14
VVADKNCSKIHFISPNIY
2048.0634 Da
25 − 42
ISSNL
533.293 Da
59 − 63
LKQML
632.38 Da
83 − 87
GADTRATEGM
1008.4415 Da
15 − 24
ISPNIY
706.377 Da
37 − 42
ISSNLEL
775.4196 Da
59 − 65
GYIGAALVLGGVD
1204.6572 Da
92 − 104
Relative Deuterium Uptake
Exposure (minutes)
State
std20S
std20S + PA28ab
std20S + PA28g

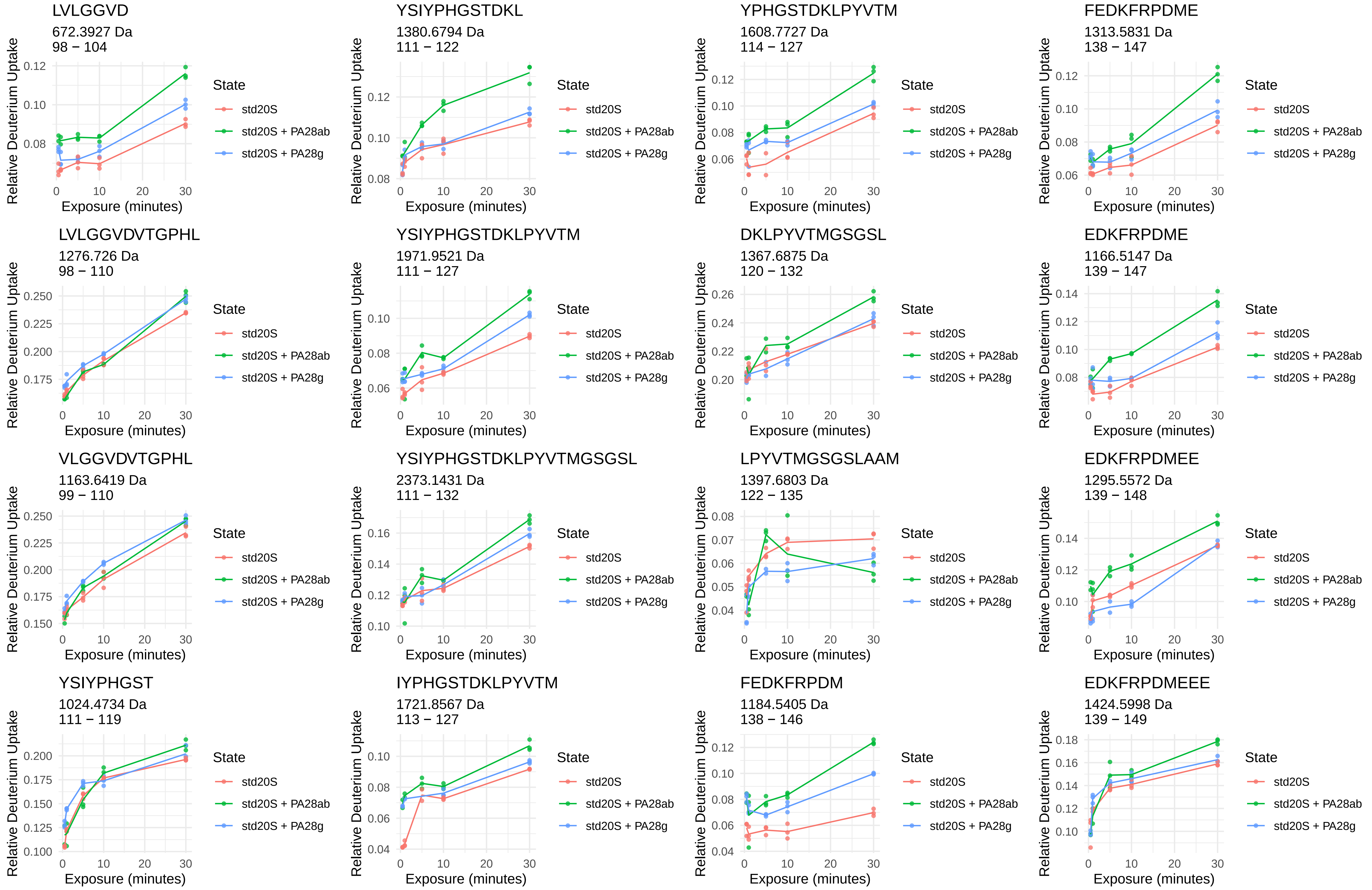
LVLGGVD
672.3927 Da
98 − 104
YSIYPHGSTDKL
1380.6794 Da
111 − 122
YPHGSTDKLPYVTM
1608.7727 Da
114 − 127
FEDKFRPDME
1313.5831 Da
138 − 147
LVLGGVDVTGPHL
1276.726 Da
98 − 110
YSIYPHGSTDKLPYVTM
1971.9521 Da
111 − 127
DKLPYVTMGSGSL
1367.6875 Da
120 − 132
EDKFRPDME
1166.5147 Da
139 − 147
VLGGVDVTGPHL
1163.6419 Da
99 − 110
YSIYPHGSTDKLPYVTMGSGSL
2373.1431 Da
111 − 132
LPYVTMGSGSLAAM
1397.6803 Da
122 − 135
EDKFRPDMEE
1295.5572 Da
139 − 148
YSIYPHGST
1024.4734 Da
111 − 119
IYPHGSTDKLPYVTM
1721.8567 Da
113 − 127
FEDKFRPDM
1184.5405 Da
138 − 146
EDKFRPDMEEE
1424.5998 Da
139 − 149
Relative Deuterium Uptake
Exposure (minutes)
State
std20S
std20S + PA28ab
std20S + PA28g

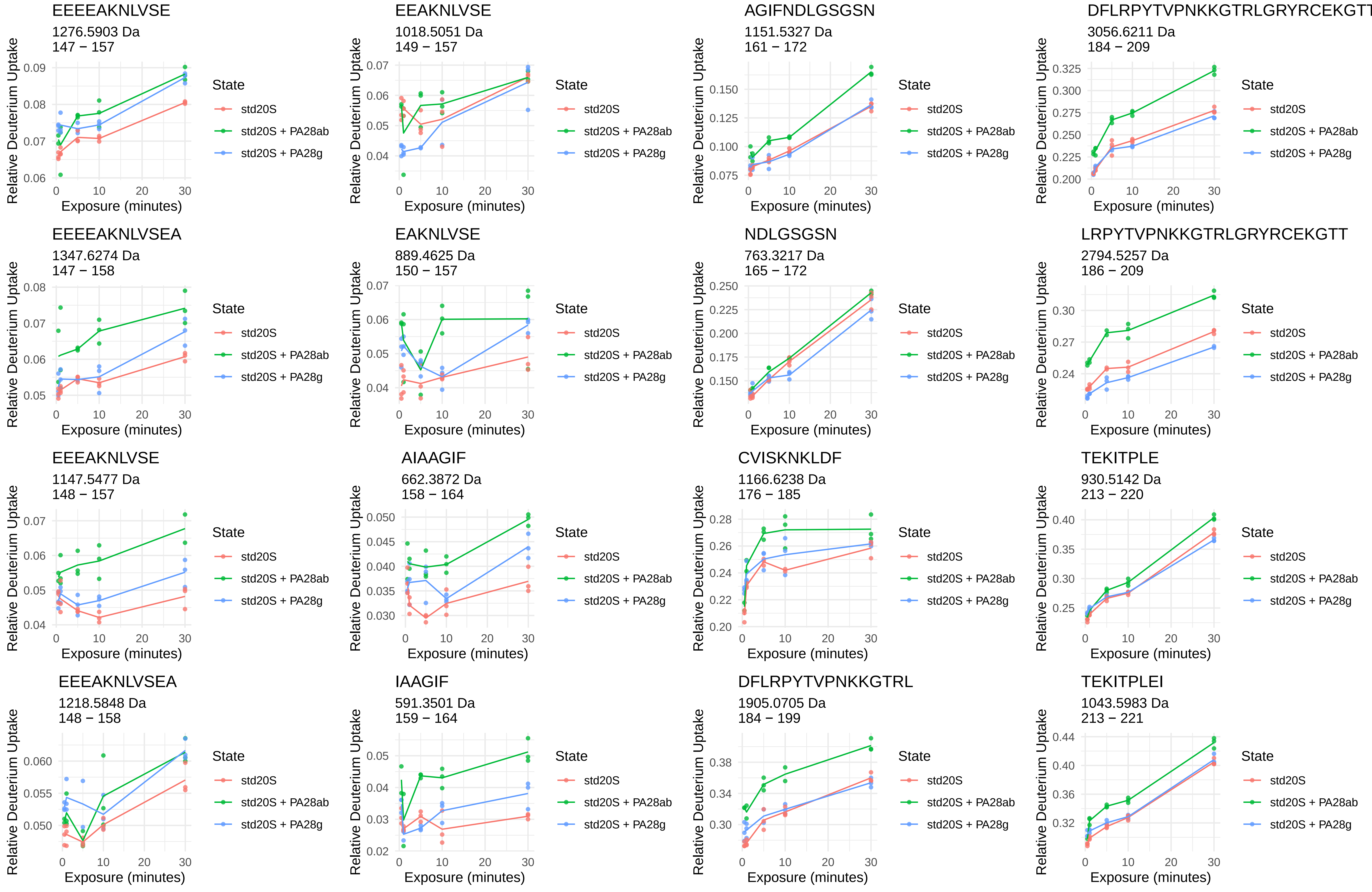
EEEEAKNLVSE
1276.5903 Da
147 − 157
EEAKNLVSE
1018.5051 Da
149 − 157
AGIFNDLGSGSN
1151.5327 Da
161 − 172
DFLRPYTVPNKKGTRLGRYRCEKGTT
3056.6211 Da
184 − 209
EEEEAKNLVSEA
1347.6274 Da
147 − 158
EAKNLVSE
889.4625 Da
150 − 157
NDLGSGSN
763.3217 Da
165 − 172
LRPYTVPNKKGTRLGRYRCEKGTT
2794.5257 Da
186 − 209
EEEAKNLVSE
1147.5477 Da
148 − 157
AIAAGIF
662.3872 Da
158 − 164
CVISKNKLDF
1166.6238 Da
176 − 185
TEKITPLE
930.5142 Da
213 − 220
EEEAKNLVSEA
1218.5848 Da
148 − 158
IAAGIF
591.3501 Da
159 − 164
DFLRPYTVPNKKGTRL
1905.0705 Da
184 − 199
TEKITPLEI
1043.5983 Da
213 − 221
Relative Deuterium Uptake
Exposure (minutes)
State
std20S
std20S + PA28ab
std20S + PA28g

EETVQTM
837.3659 Da
225 − 231

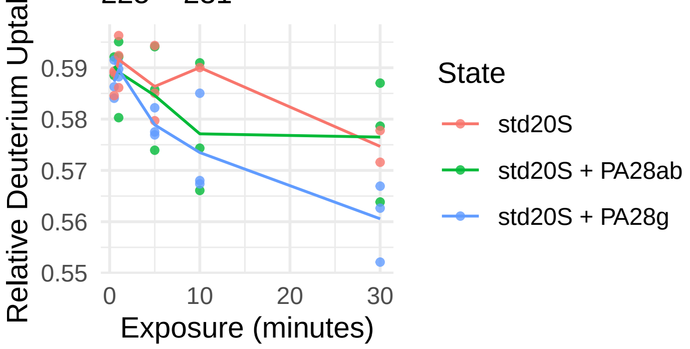


EETVQTMDTS
1140.4725 Da
225 − 234

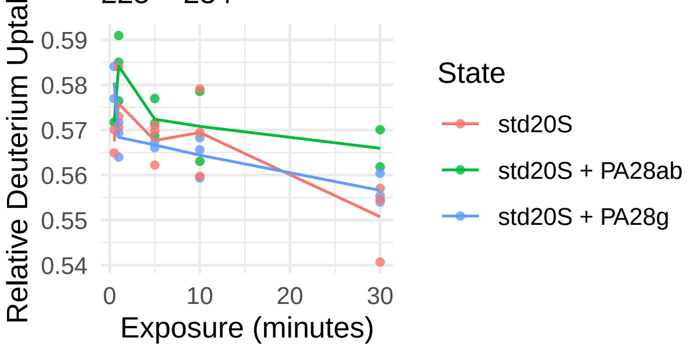


TVQTMDTS
882.3873 Da
227 − 234

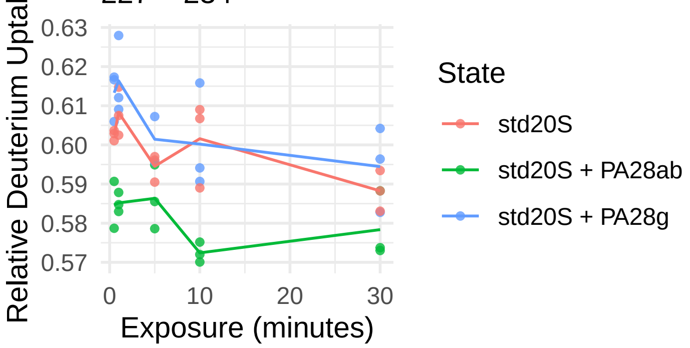

β2i

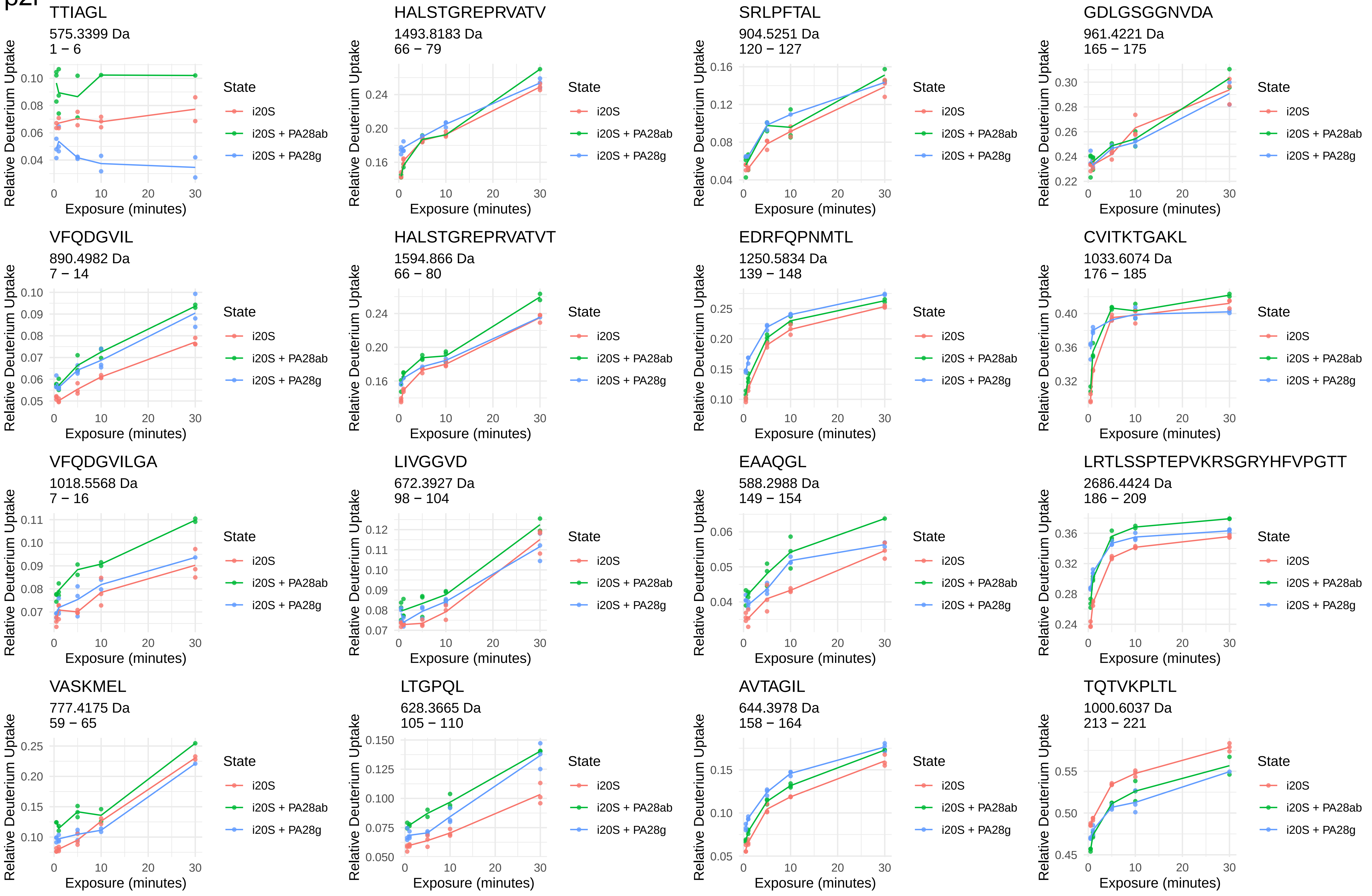
TTIAGL
575.3399 Da
1 − 6
HALSTGREPRVATV
1493.8183 Da
66 − 79
SRLPFTAL
904.5251 Da
120 − 127
GDLGSGGNVDA
961.4221 Da
165 − 175
VFQDGVIL
890.4982 Da
7 − 14
HALSTGREPRVATVT
1594.866 Da
66 − 80
EDRFQPNMTL
1250.5834 Da
139 − 148
CVITKTGAKL
1033.6074 Da
176 − 185
VFQDGVILGA
1018.5568 Da
7 − 16
LIVGGVD
672.3927 Da
98 − 104
EAAQGL
588.2988 Da
149 − 154
LRTLSSPTEPVKRSGRYHFVPGTT
2686.4424 Da
186 − 209
VASKMEL
777.4175 Da
59 − 65
LTGPQL
628.3665 Da
105 − 110
AVTAGIL
644.3978 Da
158 − 164
TQTVKPLTL
1000.6037 Da
213 − 221
Relative Deuterium Uptake
Exposure (minutes)
State
i20S
i20S + PA28ab
i20S + PA28g

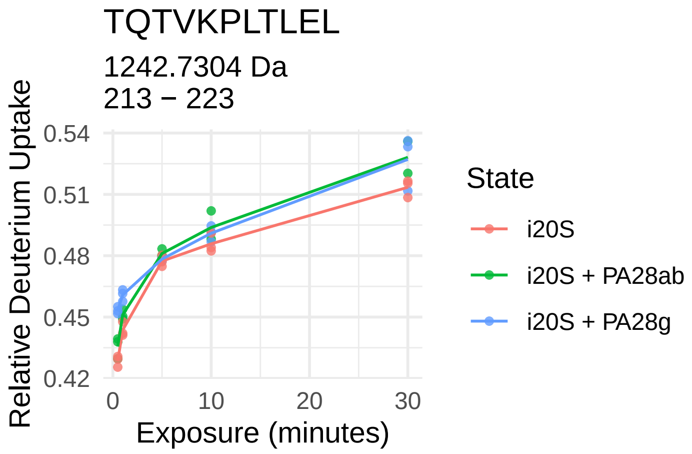

TQTVKPLTLEL
1242.7304 Da
213 − 223
Relative Deuterium Uptake
0.54
0.51
0.48
0.45
0.42
0
10
20
30
Exposure (minutes)
State
i20S
i20S + PA28ab
i20S + PA28g

β3

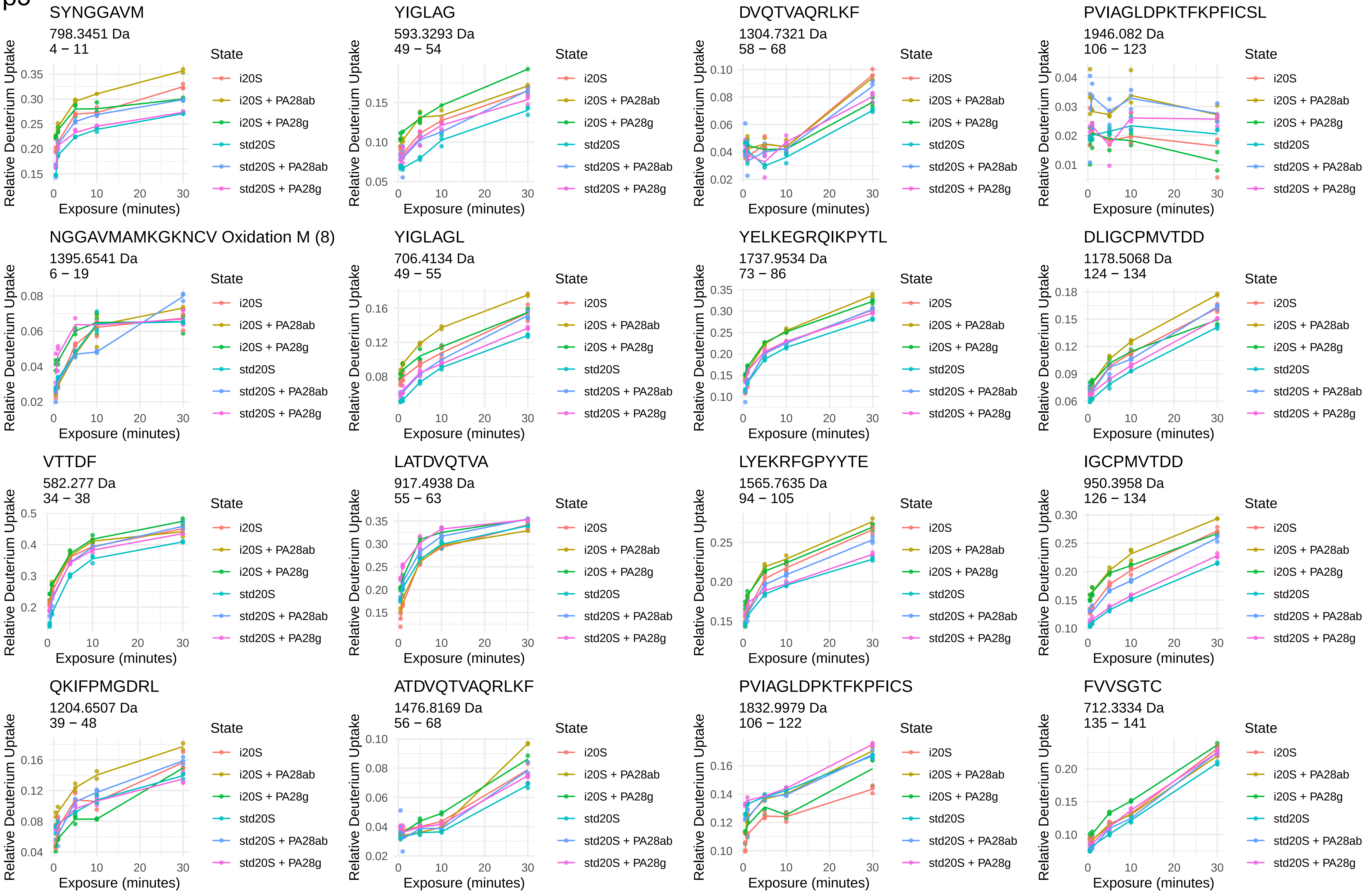
SYNGGAVM
798.3451 Da
4 − 11
YIGLAG
593.3293 Da
49 − 54
DVQTVAQRLKF
1304.7321 Da
58 − 68
PVIAGLDPKTFKPFICSL
1946.082 Da
106 − 123
NGGAVMAMKGKNCV Oxidation M (8)
1395.6541 Da
6 − 19
YIGLAGL
706.4134 Da
49 − 55
YELKEGRQIKPYTL
1737.9534 Da
73 − 86
DLIGCPMVTDD
1178.5068 Da
124 − 134
VTTDF
582.277 Da
34 − 38
LATDVQTVA
917.4938 Da
55 − 63
LYEKRFGPYYTE
1565.7635 Da
94 − 105
IGCPMVTDD
950.3958 Da
126 − 134
QKIFPMGDRL
1204.6507 Da
39 − 48
ATDVQTVAQRLKF
1476.8169 Da
56 − 68
PVIAGLDPKTFKPFICS
1832.9979 Da
106 − 122
FVVSGTC
712.3334 Da
135 − 141
Relative Deuterium Uptake
Exposure (minutes)
State
i20S
i20S + PA28ab
i20S + PA28g
std20S
std20S + PA28ab
std20S + PA28g

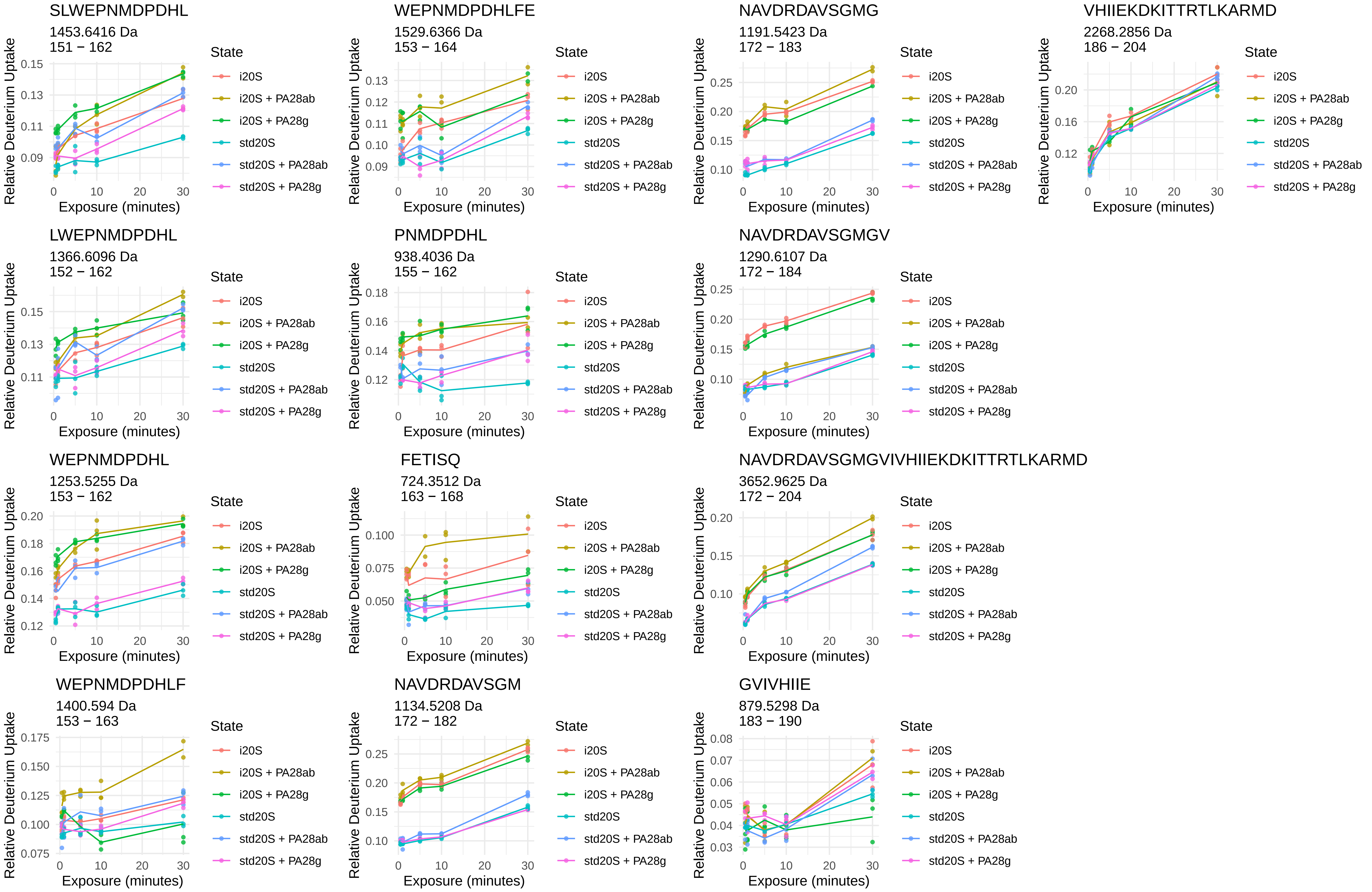

SLWEPNMDPDHL
1453.6416 Da
151 − 162
WEPNMDPDHLFE
1529.6366 Da
153 − 164
NAVDRDAVSGMG
1191.5423 Da
172 − 183
VHIIEKDKITTRTLKARMD
2268.2856 Da
186 − 204
LWEPNMDPDHL
1366.6096 Da
152 − 162
PNMDPDHL
938.4036 Da
155 − 162
NAVDRDAVSGMGV
1290.6107 Da
172 − 184
WEPNMDPDHL
1253.5255 Da
153 − 162
FETISQ
724.3512 Da
163 − 168
NAVDRDAVSGMGVIVHIIEKDKITTRTLKARMD
3652.9625 Da
172 − 204
WEPNMDPDHLF
1400.594 Da
153 − 163
NAVDRDAVSGM
1134.5208 Da
172 − 182
GVIVHIIE
879.5298 Da
183 − 190
Relative Deuterium Uptake
Exposure (minutes)
State
i20S
i20S + PA28ab
i20S + PA28g
std20S
std20S + PA28ab
std20S + PA28g

β4

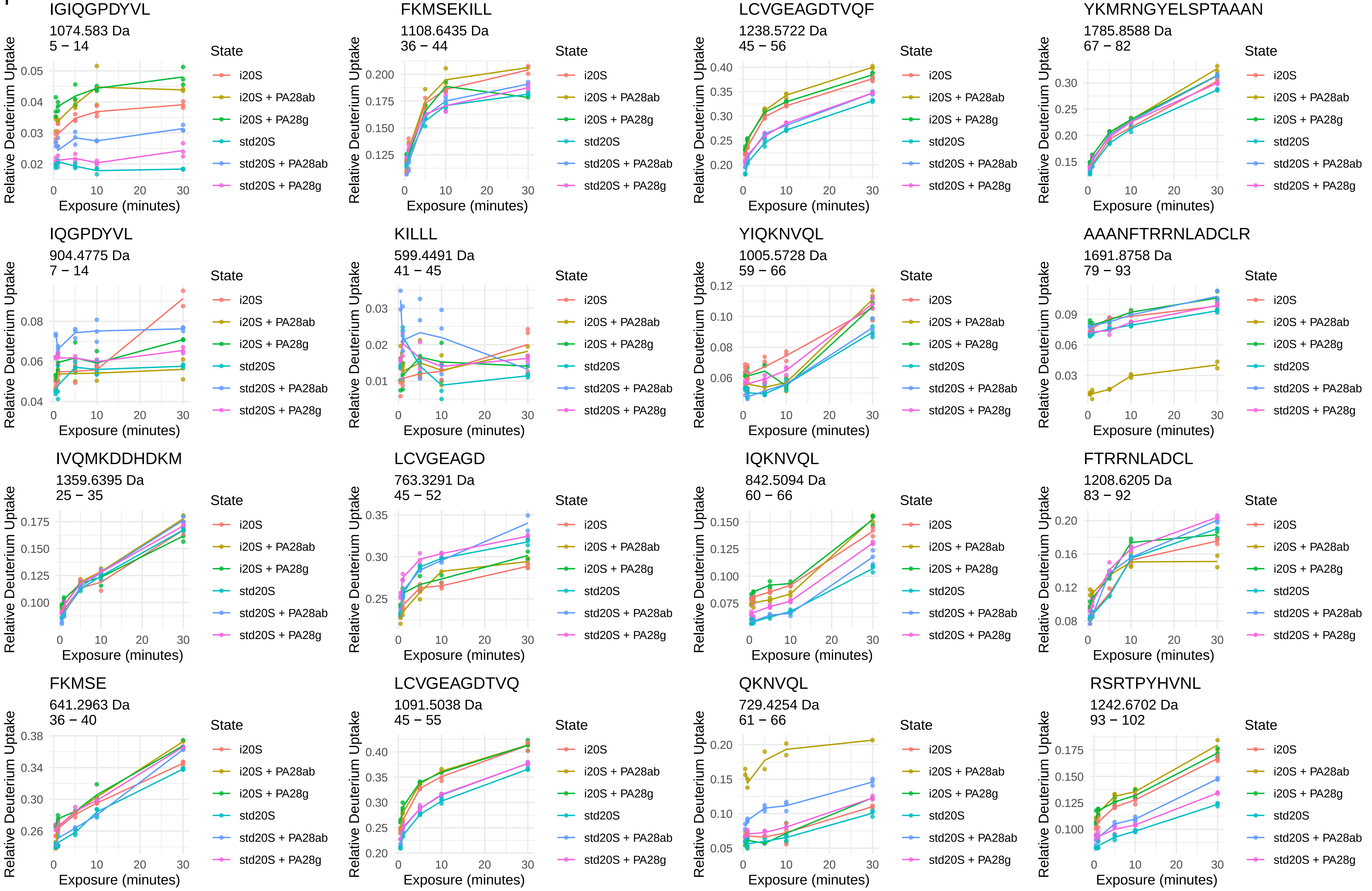
IGIQGPDYVL
1074.583 Da
5 − 14
FKMSEKILL
1108.6435 Da
36 − 44
LCVGEAGDTVQF
1238.5722 Da
45 − 56
YKMRNGYELSPTAAAN
1785.8588 Da
67 − 82
IQGPDYVL
904.4775 Da
7 − 14
KILLL
599.4491 Da
41 − 45
YIQKNVQL
1005.5728 Da
59 − 66
AAANFTRRNLADCLR
1691.8758 Da
79 − 93
IVQMKDDHDKM
1359.6395 Da
25 − 35
LCVGEAGD
763.3291 Da
45 − 52
IQKNVQL
842.5094 Da
60 − 66
FTRRNLADCL
1208.6205 Da
83 − 92
FKMSE
641.2963 Da
36 − 40
LCVGEAGDTVQ
1091.5038 Da
45 − 55
QKNVQL
729.4254 Da
61 − 66
RSRTPYHVNL
1242.6702 Da
93 − 102
Relative Deuterium Uptake
Exposure (minutes)
State
i20S
i20S + PA28ab
i20S + PA28g
std20S
std20S + PA28ab
std20S + PA28g

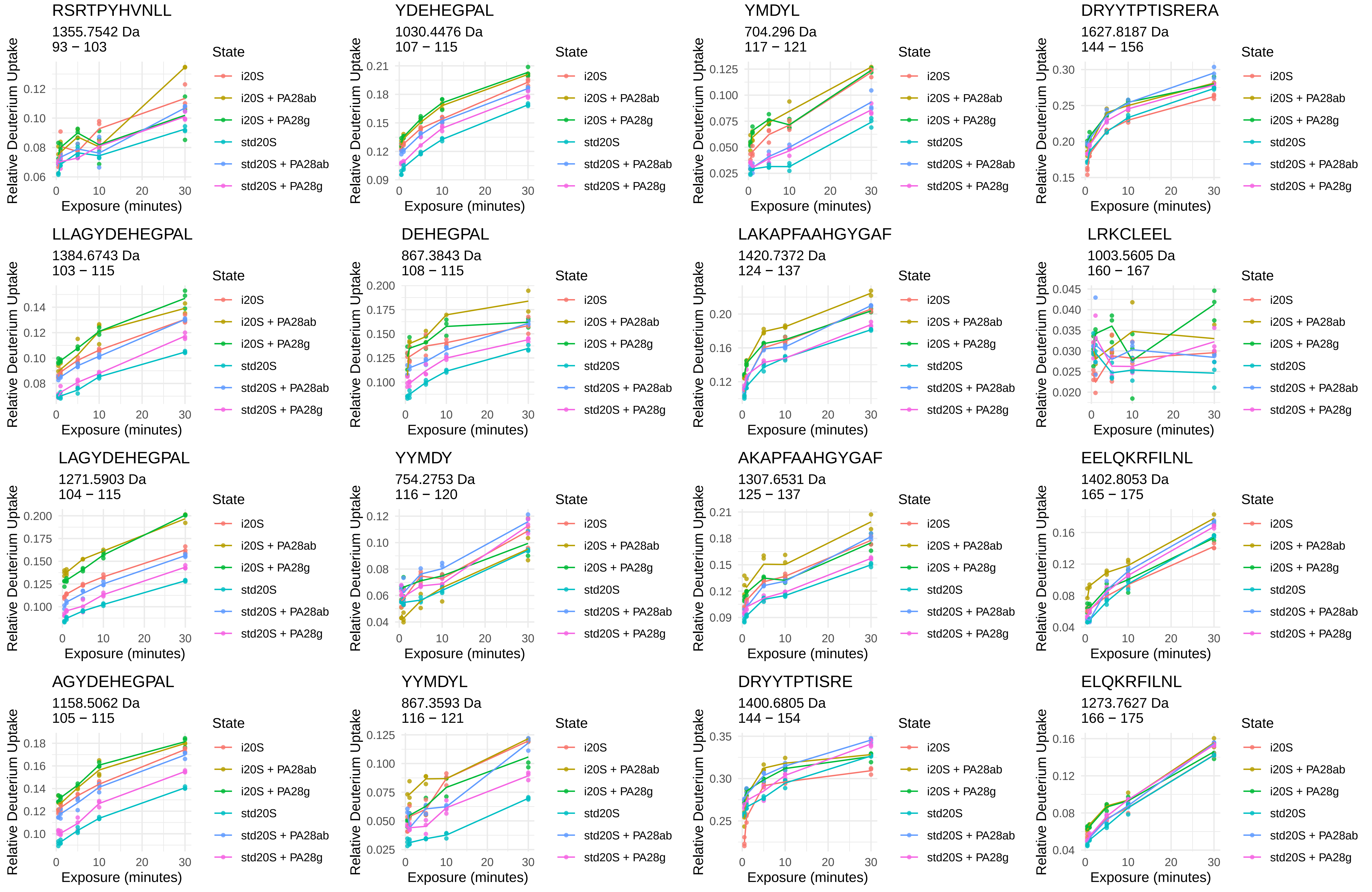
RSRTPYHVNLL
1355.7542 Da
93 − 103
YDEHEGPAL
1030.4476 Da
107 − 115
YMDYL
704.296 Da
117 − 121
DRYYTPTISRERA
1627.8187 Da
144 − 156
LLAGYDEHEGPAL
1384.6743 Da
103 − 115
DEHEGPAL
867.3843 Da
108 − 115
LAKAPFAAHGYGAF
1420.7372 Da
124 − 137
LRKCLEEL
1003.5605 Da
160 − 167
LAGYDEHEGPAL
1271.5903 Da
104 − 115
YYMDY
754.2753 Da
116 − 120
AKAPFAAHGYGAF
1307.6531 Da
125 − 137
EELQKRFILNL
1402.8053 Da
165 − 175
AGYDEHEGPAL
1158.5062 Da
105 − 115
YYMDYL
867.3593 Da
116 − 121
DRYYTPTISRE
1400.6805 Da
144 − 154
ELQKRFILNL
1273.7627 Da
166 − 175
Relative Deuterium Uptake
Exposure (minutes)
State
i20S
i20S + PA28ab
i20S + PA28g
std20S
std20S + PA28ab
std20S + PA28g

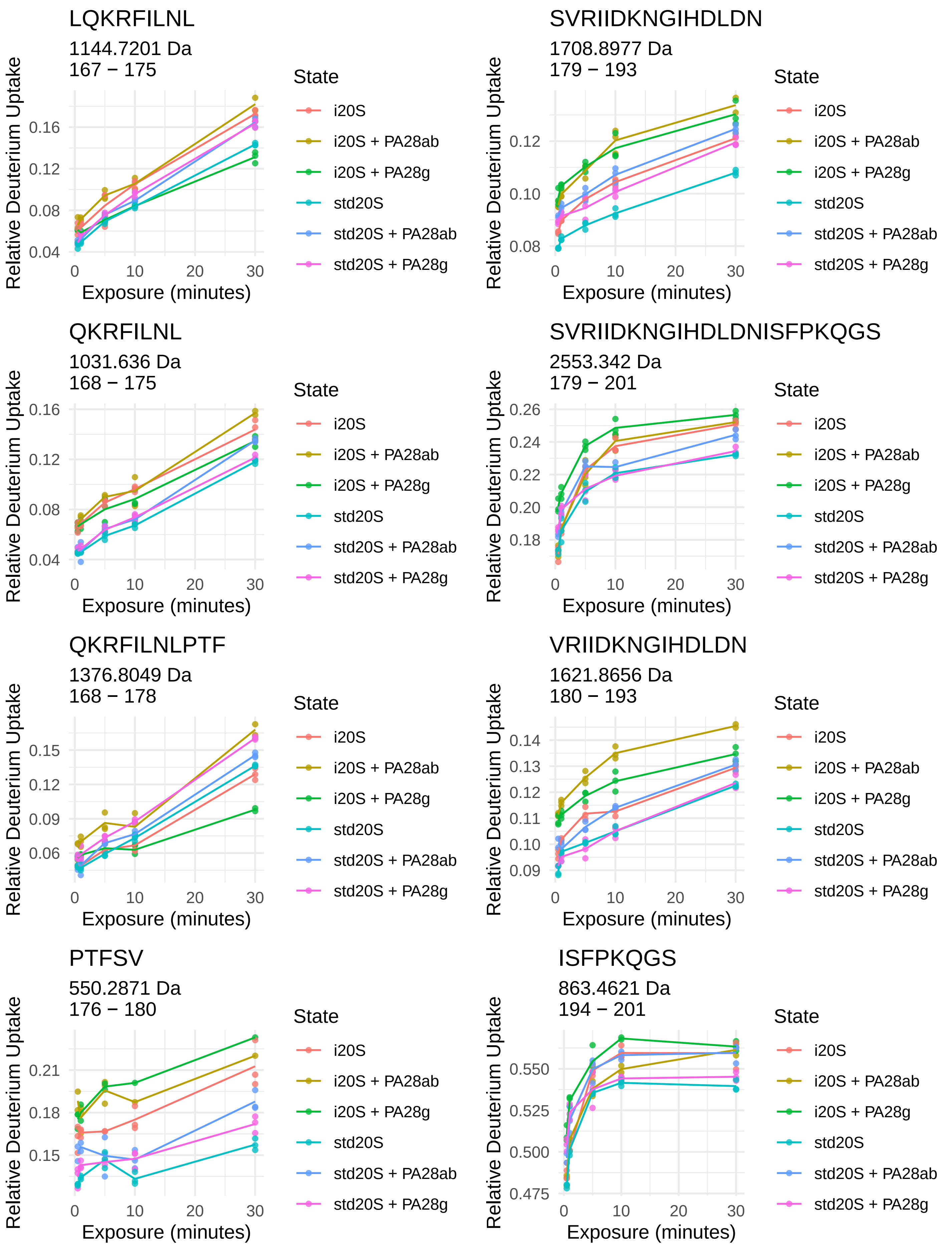
LQKRFILNL
1144.7201 Da
167 − 175
SVRIIDKNGIHDLDN
1708.8977 Da
179 − 193
QKRFILNL
1031.636 Da
168 − 175
SVRIIDKNGIHDLDNISFPKQGS
2553.342 Da
179 − 201
QKRFILNLPTF
1376.8049 Da
168 − 178
VRIIDKNGIHDLDN
1621.8656 Da
180 − 193
PTFSV
550.2871 Da
176 − 180
ISFPKQGS
863.4621 Da
194 − 201
Relative Deuterium Uptake
Exposure (minutes)
State
i20S
i20S + PA28ab
i20S + PA28g
std20S
std20S + PA28ab
std20S + PA28g

β5

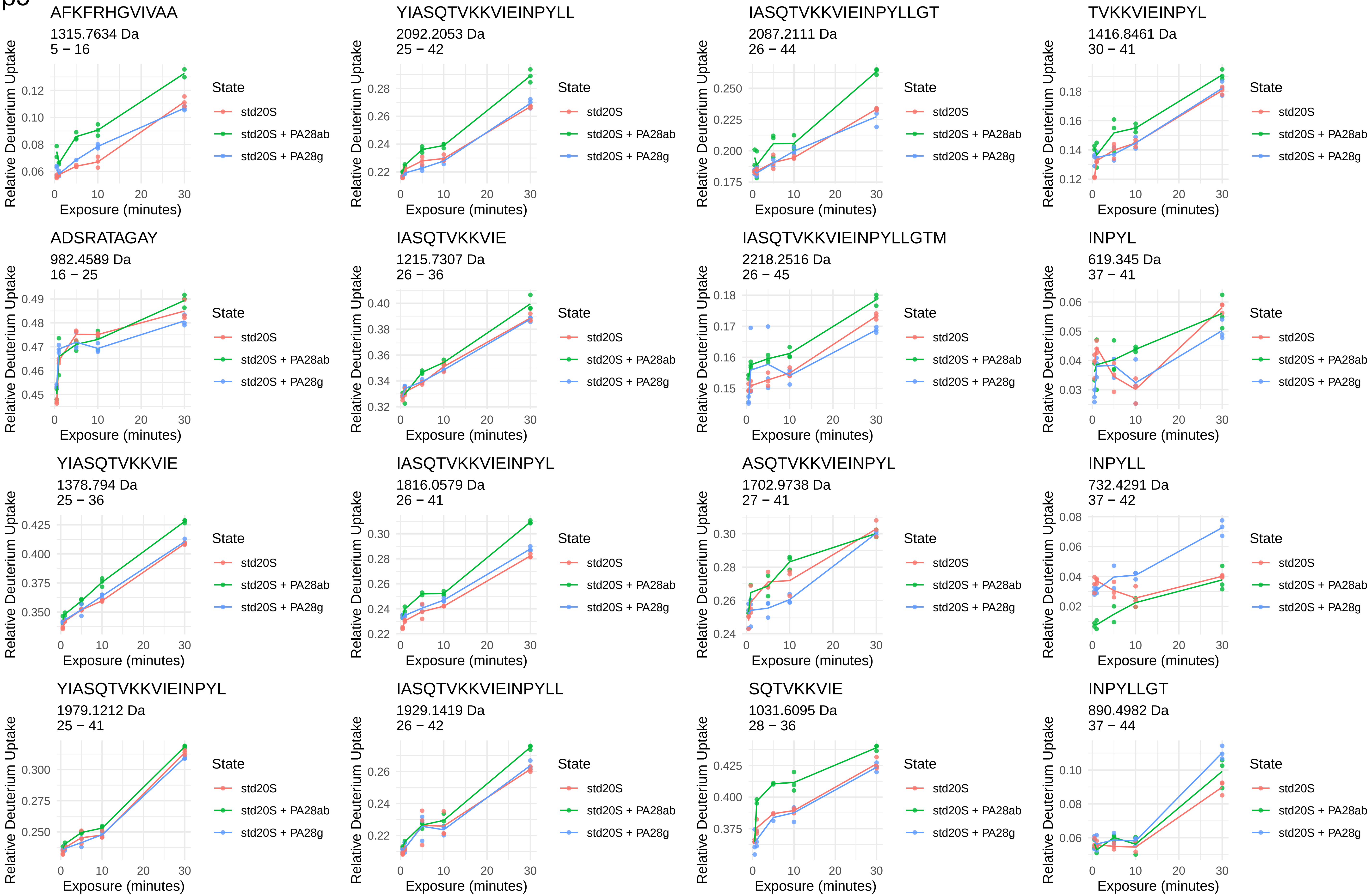
AFKFRHGVIVAA
1315.7634 Da
5 − 16
YIASQTVKKVIEINPYLL
2092.2053 Da
25 − 42
IASQTVKKVIEINPYLLGT
2087.2111 Da
26 − 44
TVKKVIEINPYL
1416.8461 Da
30 − 41
ADSRATAGAY
982.4589 Da
16 − 25
IASQTVKKVIE
1215.7307 Da
26 − 36
IASQTVKKVIEINPYLLGTM
2218.2516 Da
26 − 45
INPYL
619.345 Da
37 − 41
YIASQTVKKVIE
1378.794 Da
25 − 36
IASQTVKKVIEINPYL
1816.0579 Da
26 − 41
ASQTVKKVIEINPYL
1702.9738 Da
27 − 41
INPYLL
732.4291 Da
37 − 42
YIASQTVKKVIEINPYL
1979.1212 Da
25 − 41
IASQTVKKVIEINPYLL
1929.1419 Da
26 − 42
SQTVKKVIE
1031.6095 Da
28 − 36
INPYLLGT
890.4982 Da
37 − 44
Relative Deuterium Uptake
Exposure (minutes)
State
std20S
std20S + PA28ab
std20S + PA28g

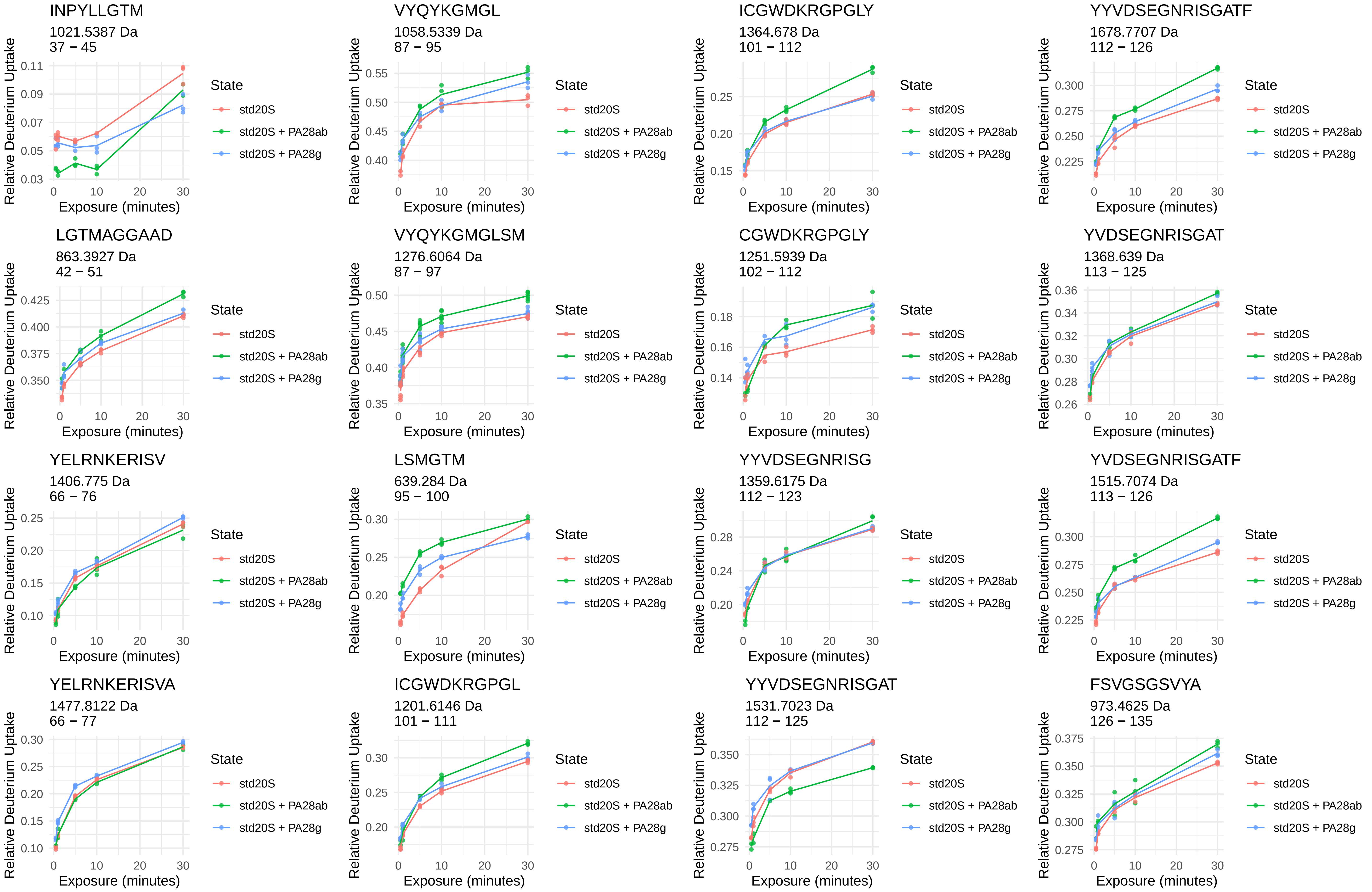
INPYLLGTM
1021.5387 Da
37 – 45
VYQYKGMGL
1058.5339 Da
87 – 95
ICGWDKRGPGLY
1364.678 Da
101 – 112
YYVDSEGNRISGATF
1678.7707 Da
112 – 126
LGTMAGGAAD
863.3927 Da
42 – 51
VYQYKGMGLSM
1276.6064 Da
87 – 97
CGWDKRGPGLY
1251.5939 Da
102 – 112
YVDSEGNRISGAT
1368.639 Da
113 – 125
YELRNKERISV
1406.775 Da
66 – 76
LSMGTM
639.284 Da
95 – 100
YYVDSEGNRISG
1359.6175 Da
112 – 123
YVDSEGNRISGATF
1515.7074 Da
113 – 126
YELRNKERISVA
1477.8122 Da
66 – 77
ICGWDKRGPGL
1201.6146 Da
101 – 111
YYVDSEGNRISGAT
1531.7023 Da
112 – 125
FSVGSGSVYA
973.4625 Da
126 – 135
Relative Deuterium Uptake
Exposure (minutes)
State
std20S
std20S + PA28ab
std20S + PA28g

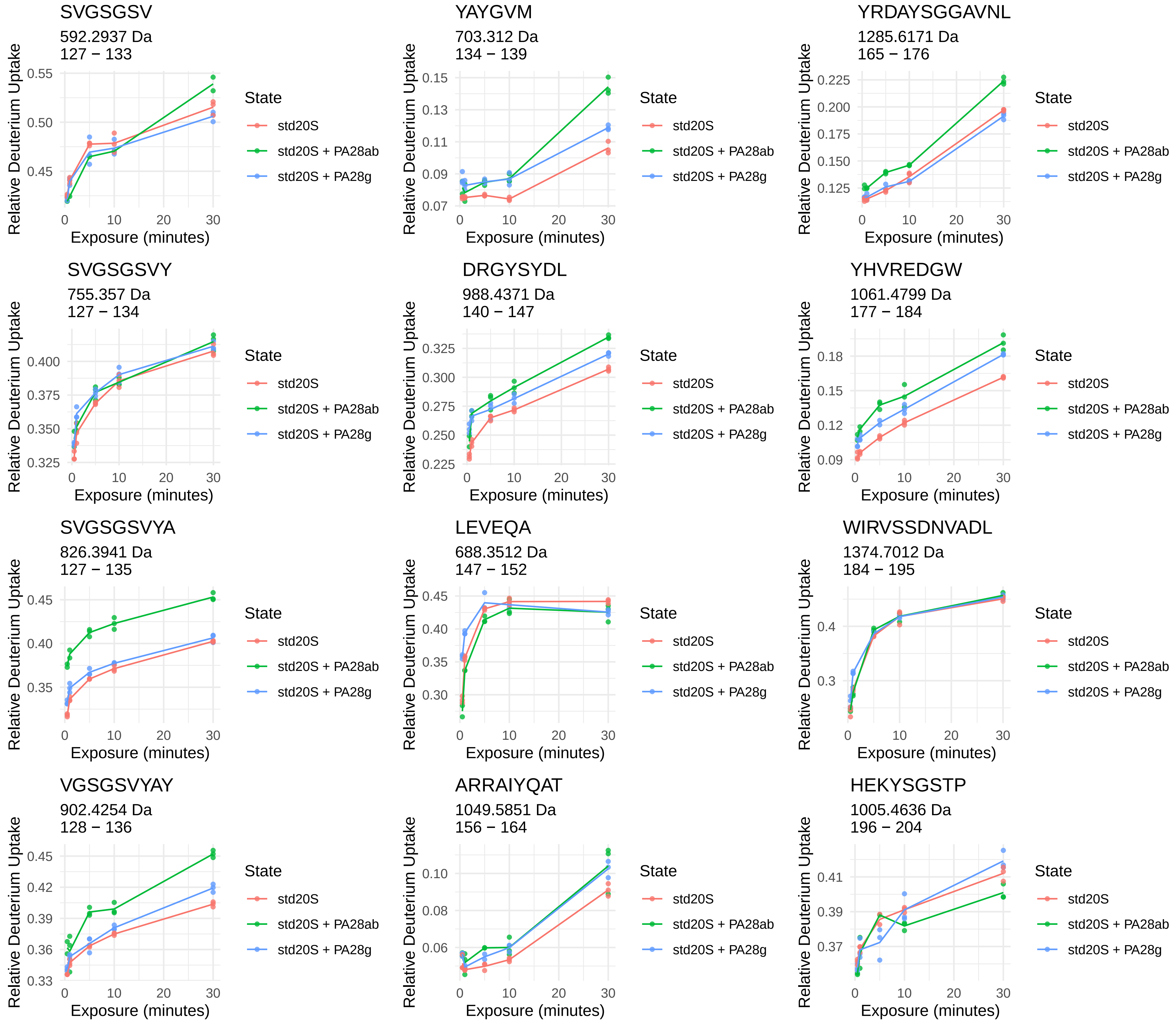
SVGSGSV
592.2937 Da
127 − 133
YAYGVM
703.312 Da
134 − 139
YRDAYSGGAVNL
1285.6171 Da
165 − 176
SVGSGSVY
755.357 Da
127 − 134
DRGYSYDL
988.4371 Da
140 − 147
YHVREDGW
1061.4799 Da
177 − 184
SVGSGSVYA
826.3941 Da
127 − 135
LEVEQA
688.3512 Da
147 − 152
WIRVSSDNVADL
1374.7012 Da
184 − 195
VGSGSVYAY
902.4254 Da
128 − 136
ARRAIYQAT
1049.5851 Da
156 − 164
HEKYSGSTP
1005.4636 Da
196 − 204
Relative Deuterium Uptake
Exposure (minutes)
State
std20S
std20S + PA28ab
std20S + PA28g

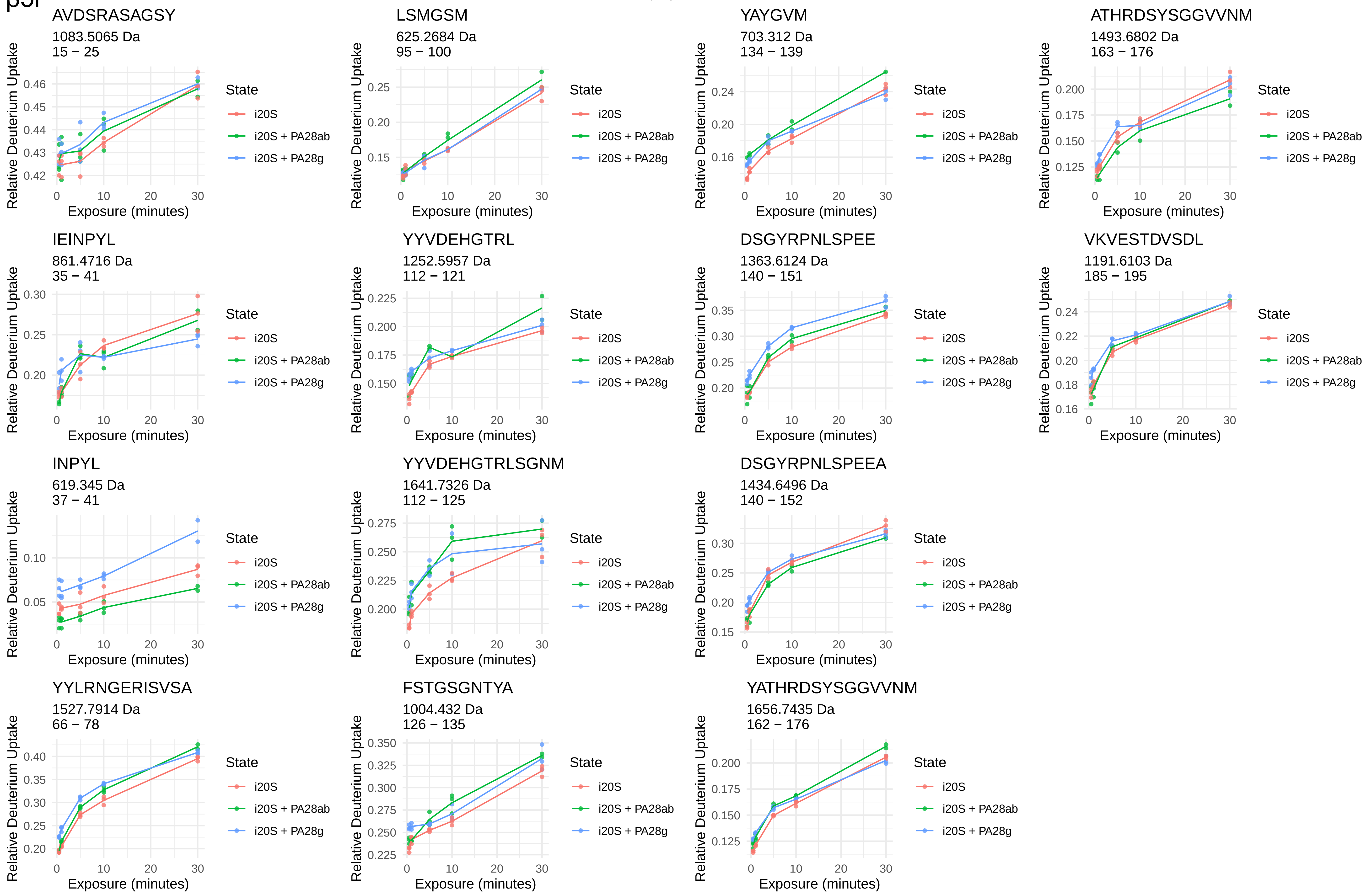
β5i
AVDSRASAGSY
1083.5065 Da
15 − 25
LSMGSM
625.2684 Da
95 − 100
YAYGVM
703.312 Da
134 − 139
ATHRDSYSGGVVNM
1493.6802 Da
163 − 176
IEINPYL
861.4716 Da
35 − 41
YYVDEHGTRL
1252.5957 Da
112 − 121
DSGYRPNLSPEE
1363.6124 Da
140 − 151
VKVESTDVSDL
1191.6103 Da
185 − 195
INPYL
619.345 Da
37 − 41
YYVDEHGTRLSGNM
1641.7326 Da
112 − 125
DSGYRPNLSPEEA
1434.6496 Da
140 − 152
YYLRNGERISVSA
1527.7914 Da
66 − 78
FSTGSGNTYA
1004.432 Da
126 − 135
YATHRDSYSGGVVNM
1656.7435 Da
162 − 176
Relative Deuterium Uptake
Exposure (minutes)
State
i20S
i20S + PA28ab
i20S + PA28g

β6

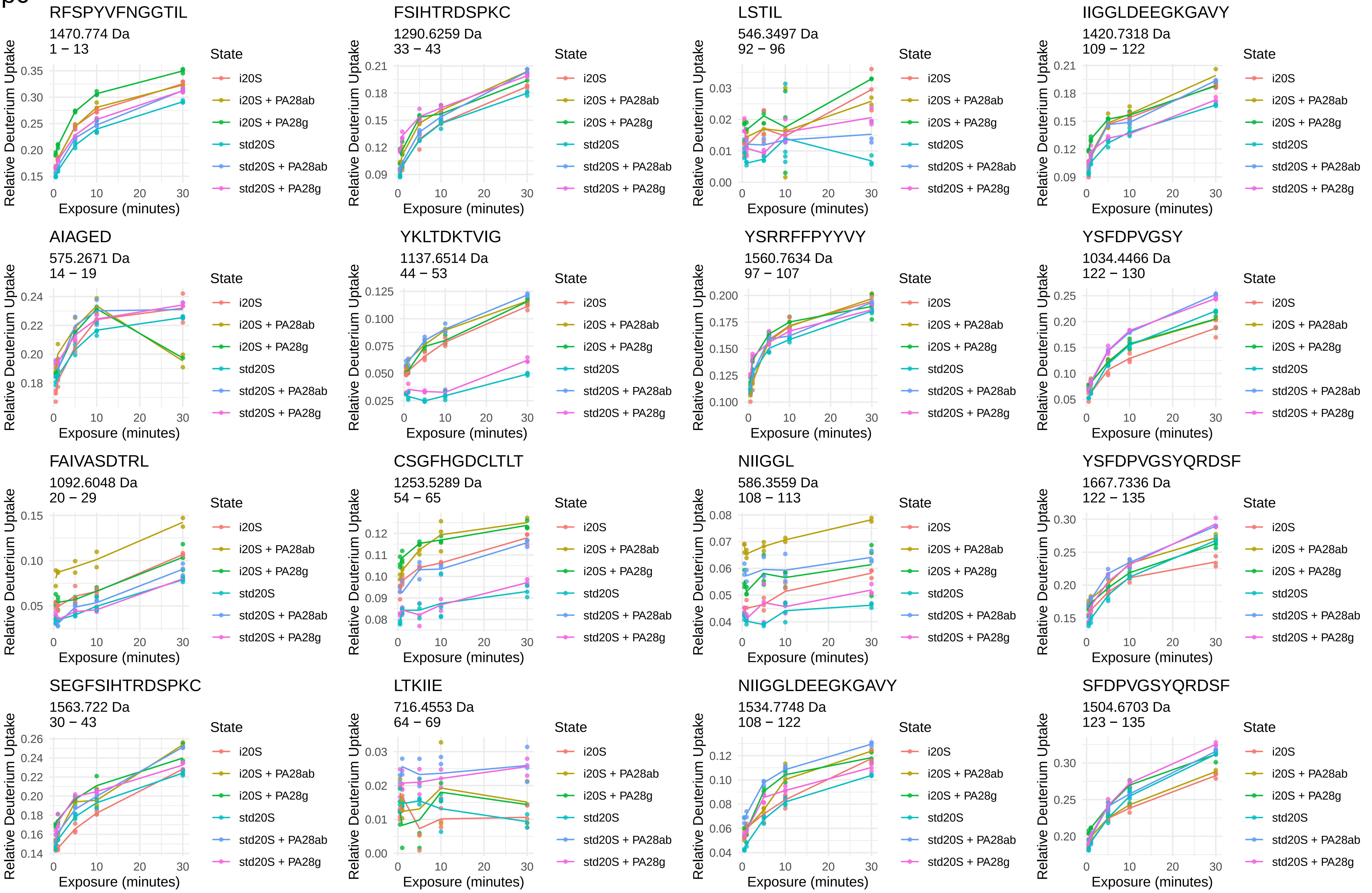
RFSPYVFNGGTIL
1470.774 Da
1 − 13
FSIHTRDSPKC
1290.6259 Da
33 − 43
LSTIL
546.3497 Da
92 − 96
IIGGLDEEGKGAVY
1420.7318 Da
109 − 122
AIAGED
575.2671 Da
14 − 19
YKLTDKTVIG
1137.6514 Da
44 − 53
YSRRFFPYYVY
1560.7634 Da
97 − 107
YSFDPVGSY
1034.4466 Da
122 − 130
FAIVASDTRL
1092.6048 Da
20 − 29
CSGFHGDCLTLT
1253.5289 Da
54 − 65
NIIGGL
586.3559 Da
108 − 113
YSFDPVGSYQRDSF
1667.7336 Da
122 − 135
SEGFSIHTRDSPKC
1563.722 Da
30 − 43
LTKIIE
716.4553 Da
64 − 69
NIIGGLDEEGKGAVY
1534.7748 Da
108 − 122
SFDPVGSYQRDSF
1504.6703 Da
123 − 135
Relative Deuterium Uptake
Exposure (minutes)
State
i20S
i20S + PA28ab
i20S + PA28g
std20S
std20S + PA28ab
std20S + PA28g

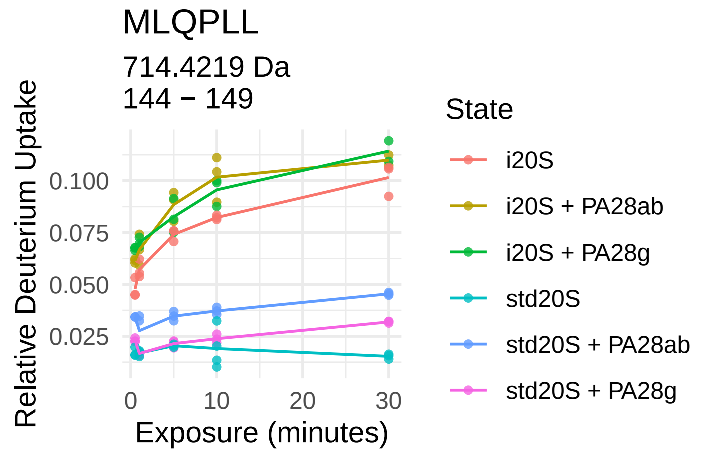
MLQPLL
714.4219 Da
144 − 149
Relative Deuterium Uptake
0.025
0.050
0.075
0.100
0
10
20
30
Exposure (minutes)
State
i20S
i20S + PA28ab
i20S + PA28g
std20S
std20S + PA28ab
std20S + PA28g

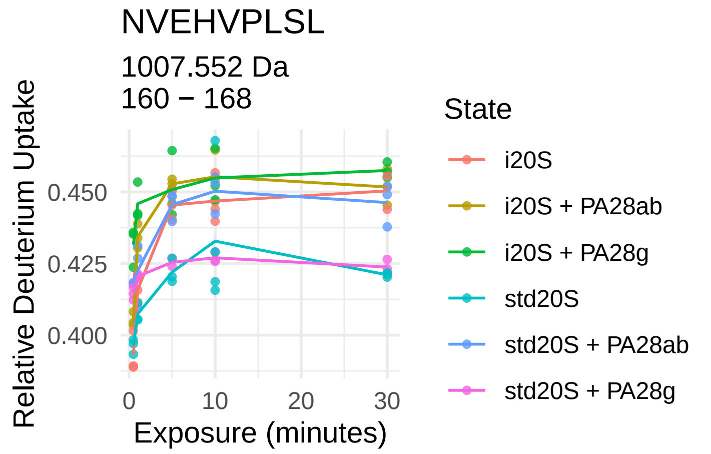
NVEHVPLSL
1007.552 Da
160 − 168
Relative Deuterium Uptake
0.400
0.425
0.450
0
10
20
30
Exposure (minutes)
State
i20S
i20S + PA28ab
i20S + PA28g
std20S
std20S + PA28ab
std20S + PA28g

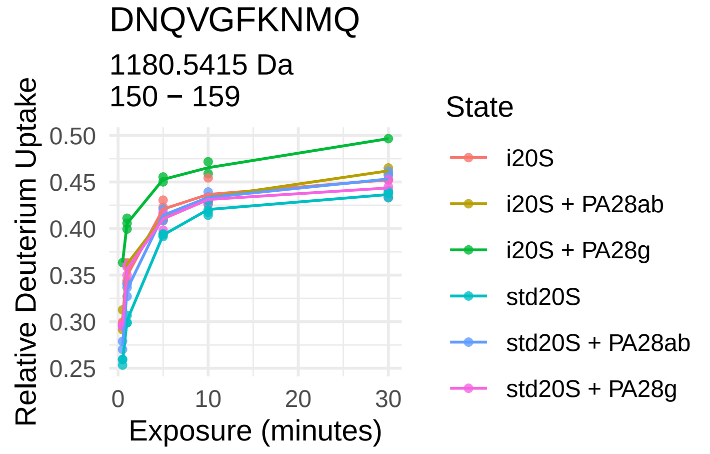
DNQVGFKNMQ
1180.5415 Da
150 − 159
Relative Deuterium Uptake
0.25
0.30
0.35
0.40
0.45
0.50
0
10
20
30
Exposure (minutes)
State
i20S
i20S + PA28ab
i20S + PA28g
std20S
std20S + PA28ab
std20S + PA28g

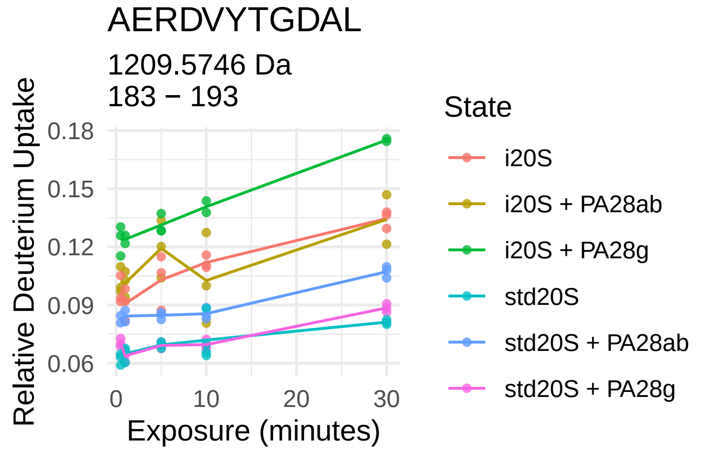
AERDVYTGDAL
1209.5746 Da
183 − 193
Relative Deuterium Uptake
0.06
0.09
0.12
0.15
0.18
0
10
20
30
Exposure (minutes)
State
i20S
i20S + PA28ab
i20S + PA28g
std20S
std20S + PA28ab
std20S + PA28g

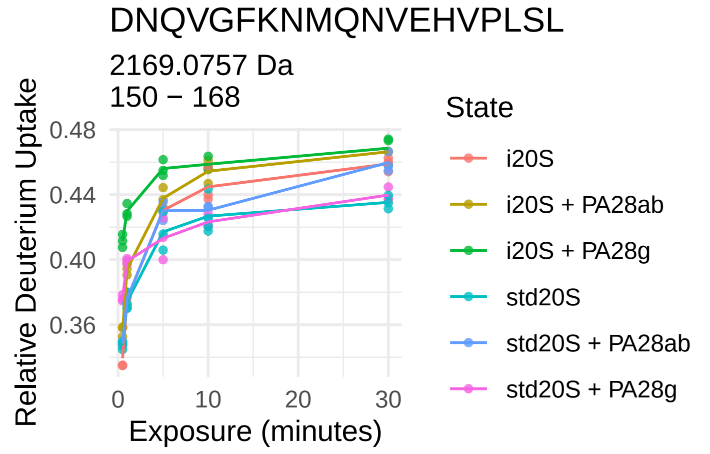
DNQVGFKNMQNVEHVPLSL
2169.0757 Da
150 − 168
Relative Deuterium Uptake
0.36
0.40
0.44
0.48
0
10
20
30
Exposure (minutes)
State
i20S
i20S + PA28ab
i20S + PA28g
std20S
std20S + PA28ab
std20S + PA28g

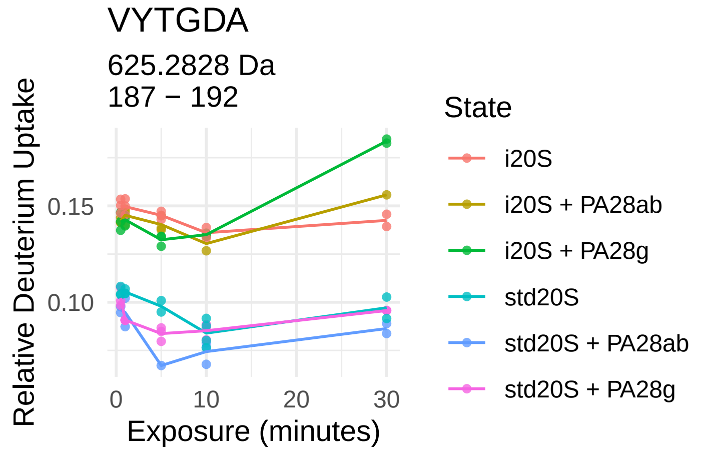
VYTGDA
625.2828 Da
187 − 192
Relative Deuterium Uptake
0.10
0.15
0
10
20
30
Exposure (minutes)
State
i20S
i20S + PA28ab
i20S + PA28g
std20S
std20S + PA28ab
std20S + PA28g

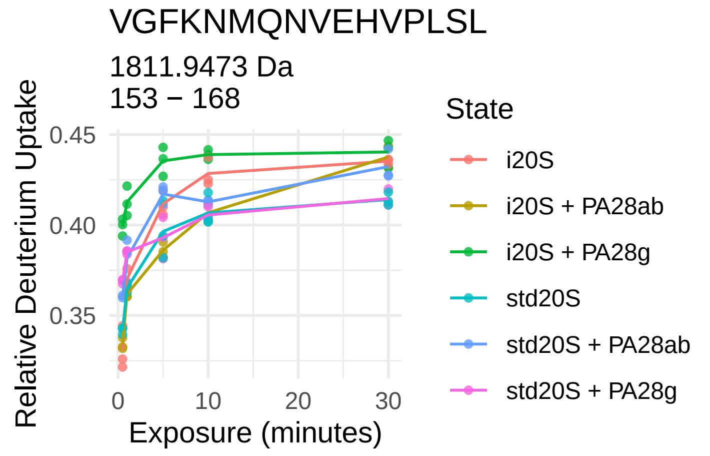
VGFKNMQNVEHVPLSL
1811.9473 Da
153 − 168
Relative Deuterium Uptake
0.35
0.40
0.45
0
10
20
30
Exposure (minutes)
State
i20S
i20S + PA28ab
i20S + PA28g
std20S
std20S + PA28ab
std20S + PA28g

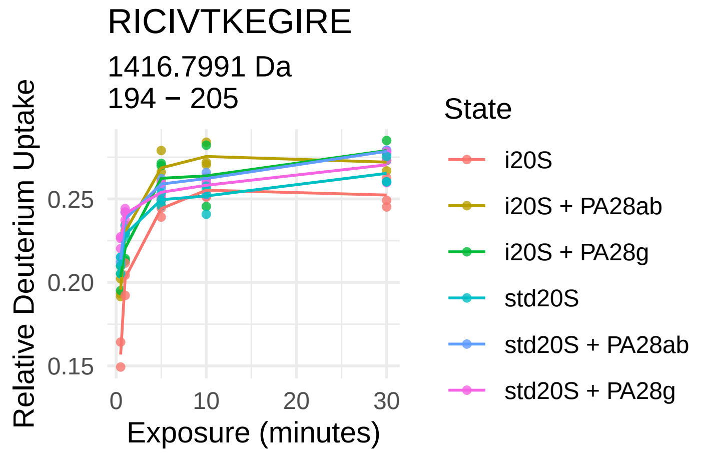
RICIVTKEGIRE
1416.7991 Da
194 − 205
Relative Deuterium Uptake
0.15
0.20
0.25
0
10
20
30
Exposure (minutes)
State
i20S
i20S + PA28ab
i20S + PA28g
std20S
std20S + PA28ab
std20S + PA28g

β7

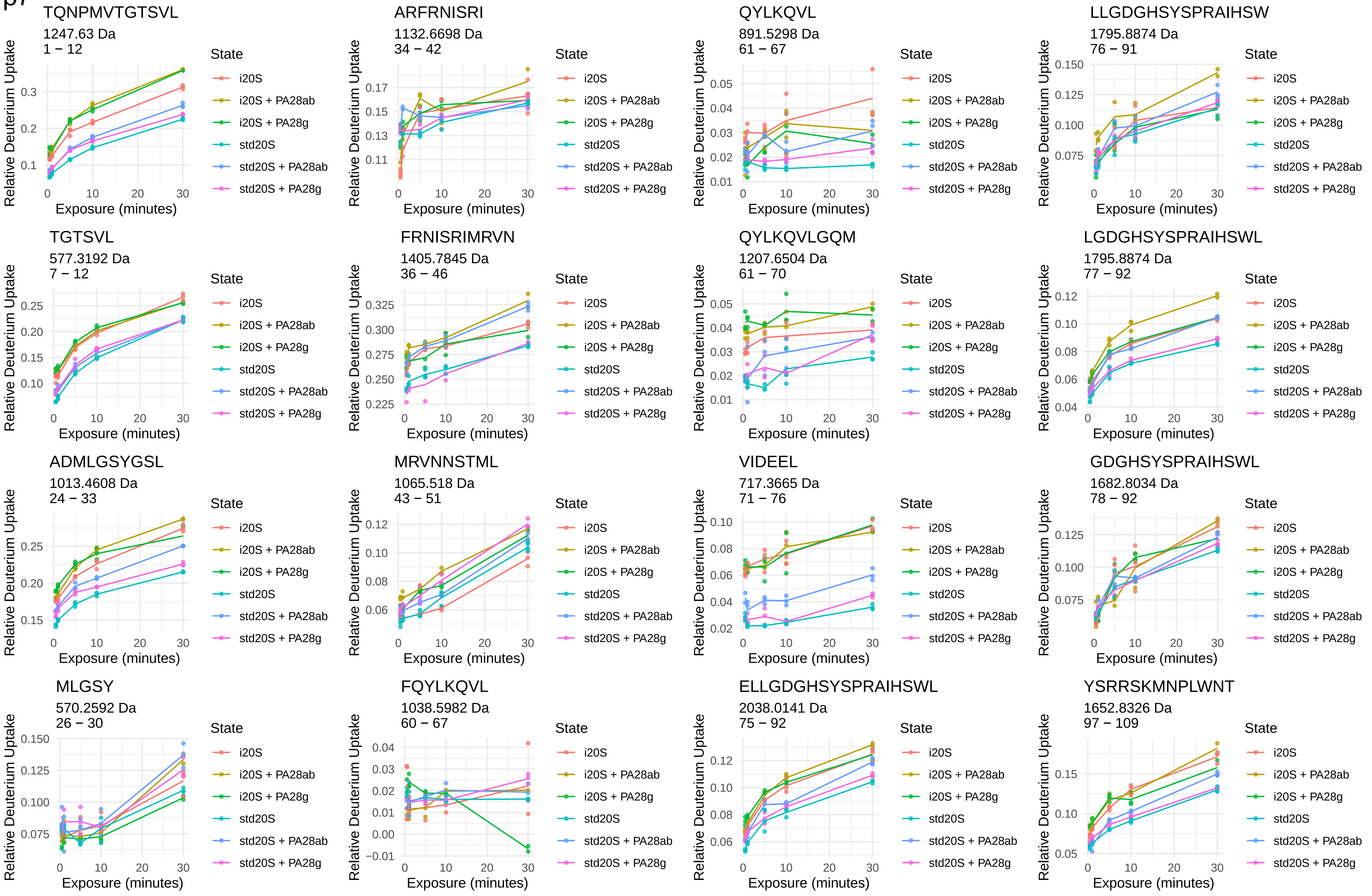
TQNPMVTGTSVL
1247.63 Da
1 − 12
ARFRNISRI
1132.6698 Da
34 − 42
QYLKQVL
891.5298 Da
61 − 67
LLGDGHSYSPRAIHSW
1795.8874 Da
76 − 91
TGTSVL
577.3192 Da
7 − 12
FRNISRIMRVN
1405.7845 Da
36 − 46
QYLKQVLGQM
1207.6504 Da
61 − 70
LGDGHSYSPRAIHSWL
1795.8874 Da
77 − 92
ADMLGSYGSL
1013.4608 Da
24 − 33
MRVNNSTML
1065.518 Da
43 − 51
VIDEEL
717.3665 Da
71 − 76
GDGHSYSPRAIHSWL
1682.8034 Da
78 − 92
MLGSY
570.2592 Da
26 − 30
FQYLKQVL
1038.5982 Da
60 − 67
ELLGDGHSYSPRAIHSWL
2038.0141 Da
75 − 92
YSRRSKMNPLWNT
1652.8326 Da
97 − 109
Relative Deuterium Uptake
Exposure (minutes)
State
i20S
i20S + PA28ab
i20S + PA28g
std20S
std20S + PA28ab
std20S + PA28g

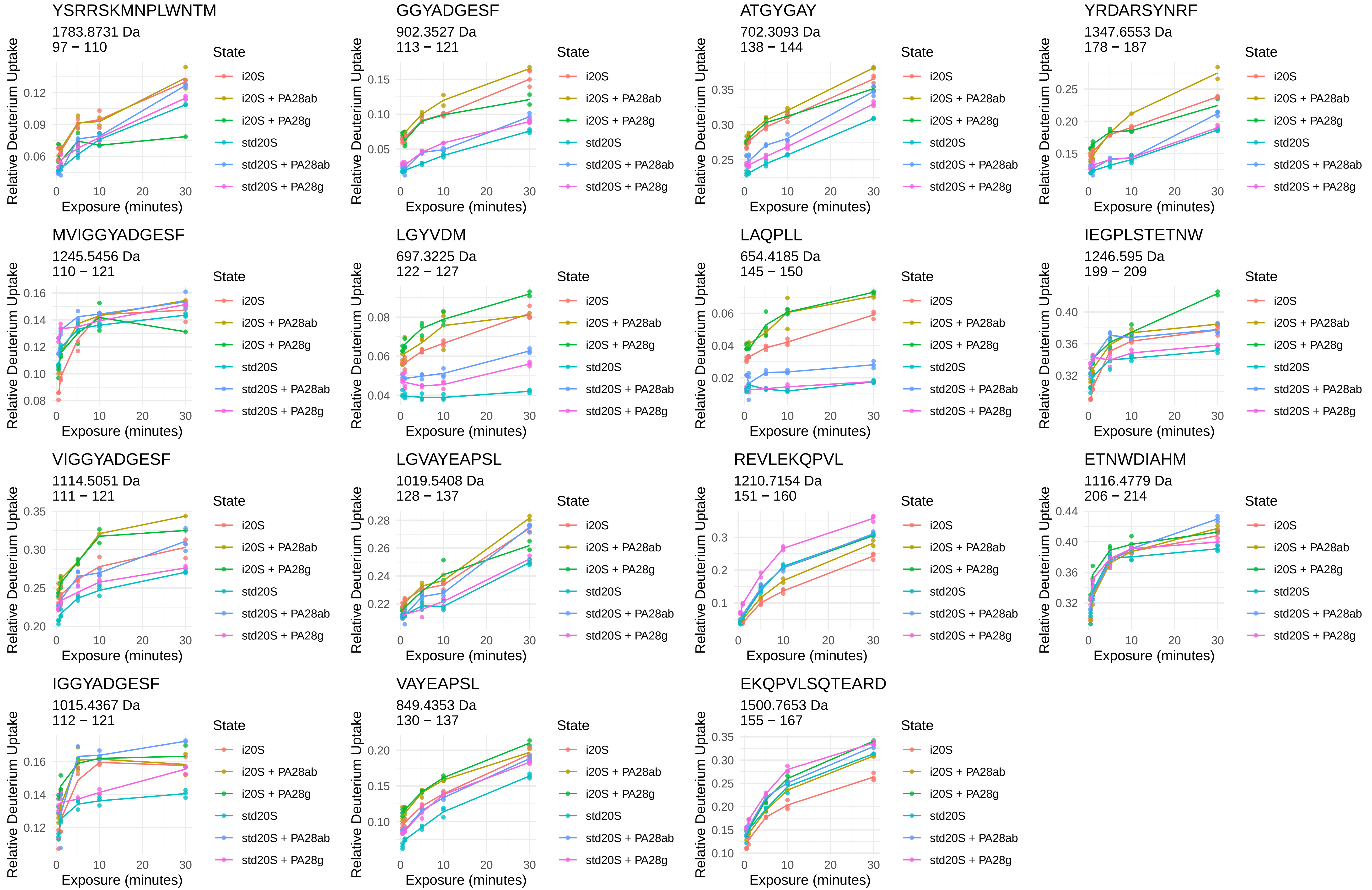
YSRRSKMNPLWNTM
1783.8731 Da
97 – 110
GGYADGESF
902.3527 Da
113 – 121
ATGYGAY
702.3093 Da
138 – 144
YRDARSYNRF
1347.6553 Da
178 – 187
MVIGGYADGESF
1245.5456 Da
110 – 121
LGYVDM
697.3225 Da
122 – 127
LAQPLL
654.4185 Da
145 – 150
IEGPLSTETNW
1246.595 Da
199 – 209
VIGGYADGESF
1114.5051 Da
111 – 121
LGVAYEAPSL
1019.5408 Da
128 – 137
REVLEKQPVL
1210.7154 Da
151 – 160
ETNWDIAHM
1116.4779 Da
206 – 214
IGGYADGESF
1015.4367 Da
112 – 121
VAYEAPSL
849.4353 Da
130 – 137
EKQPVLSQTEARD
1500.7653 Da
155 – 167
Relative Deuterium Uptake
Exposure (minutes)
State
i20S
i20S + PA28ab
i20S + PA28g
std20S
std20S + PA28ab
std20S + PA28g

PA28α



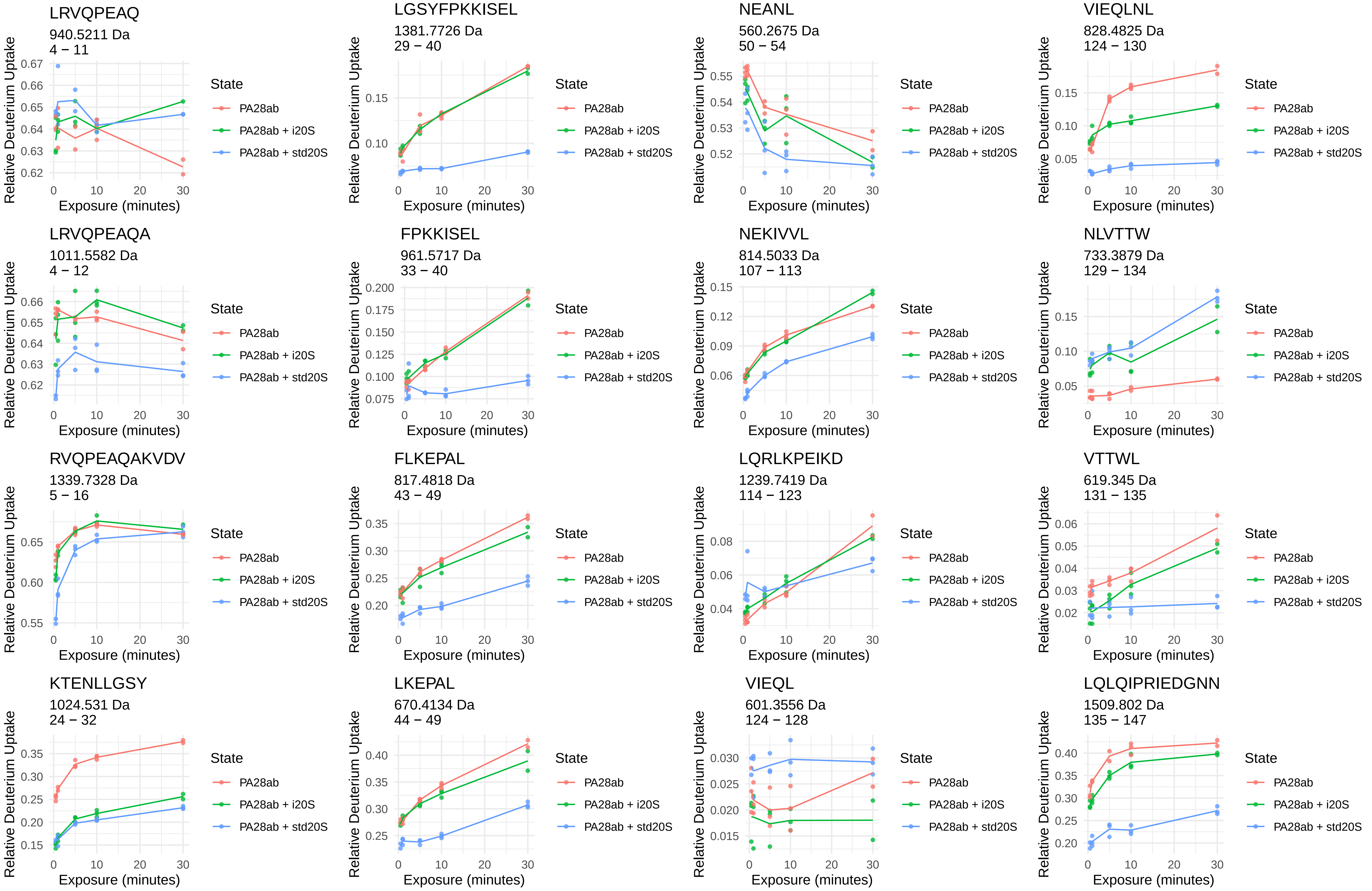
LRVQPEAQ
940.5211 Da
4 − 11
LGSYFPKKISEL
1381.7726 Da
29 − 40
NEANL
560.2675 Da
50 − 54
VIEQLNL
828.4825 Da
124 − 130
LRVQPEAQA
1011.5582 Da
4 − 12
FPKKISEL
961.5717 Da
33 − 40
NEKIVVL
814.5033 Da
107 − 113
NLVTTW
733.3879 Da
129 − 134
RVQPEAQAKVDV
1339.7328 Da
5 − 16
FLKEPAL
817.4818 Da
43 − 49
LQRLKPEIKD
1239.7419 Da
114 − 123
VTTWL
619.345 Da
131 − 135
KTENLLGSY
1024.531 Da
24 − 32
LKEPAL
670.4134 Da
44 − 49
VIEQL
601.3556 Da
124 − 128
LQLQIPRIEDGNN
1509.802 Da
135 − 147
Relative Deuterium Uptake
Exposure (minutes)
State
PA28ab
PA28ab + i20S
PA28ab + std20S

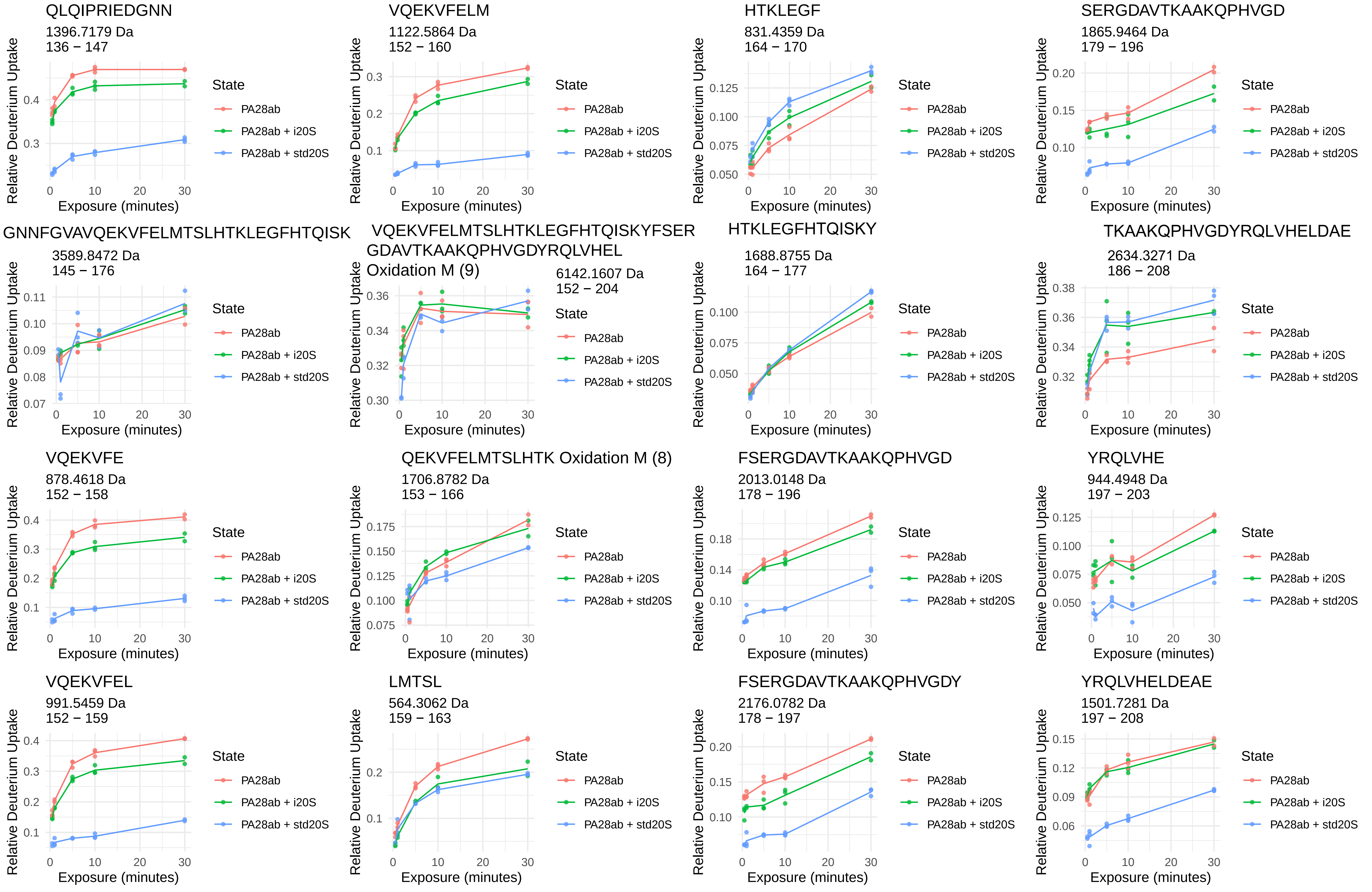
QLQIPRIEDGNN
1396.7179 Da
136 − 147
VQEKVFELM
1122.5864 Da
152 − 160
HTKLEGF
831.4359 Da
164 − 170
SERGDAVTKAAKQPHVGD
1865.9464 Da
179 − 196
GNNFGVAVQEKVFELMTSLHTKLEGFHTQISK
3589.8472 Da
145 − 176
VQEKVFELMTSLHTKLEGFHTQISKYFSER GDAVTKAAKQPHVGDYRQLVHEL Oxidation M (9)
6142.1607 Da
152 − 204
HTKLEGFHTQISKY
1688.8755 Da
164 − 177
TKAAKQPHVGDYRQLVHELDAE
2634.3271 Da
186 − 208
VQEKVFE
878.4618 Da
152 − 158
QEKVFELMTSLHTK Oxidation M (8)
1706.8782 Da
153 − 166
FSERGDAVTKAAKQPHVGD
2013.0148 Da
178 − 196
YRQLVHE
944.4948 Da
197 − 203
VQEKVFEL
991.5459 Da
152 − 159
LMTSL
564.3062 Da
159 − 163
FSERGDAVTKAAKQPHVGDY
2176.0782 Da
178 − 197
YRQLVHELDEAE
1501.7281 Da
197 − 208
Relative Deuterium Uptake
Exposure (minutes)
State
PA28ab
PA28ab + i20S
PA28ab + std20S

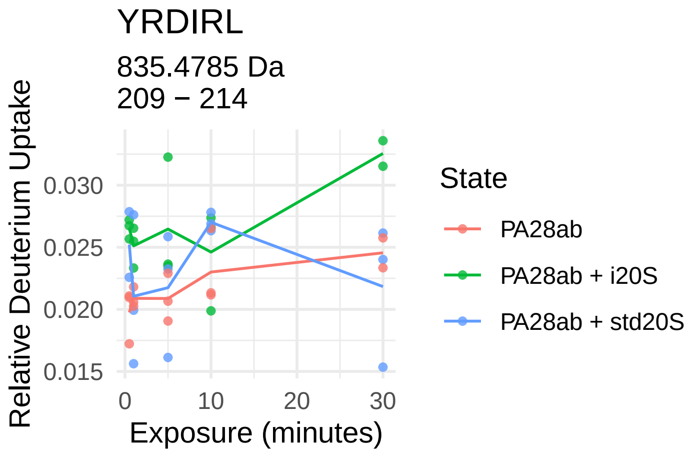
YRDIRL
835.4785 Da
209 − 214
Relative Deuterium Uptake
0.030
0.025
0.020
0.015
0
10
20
30
Exposure (minutes)
State
PA28ab
PA28ab + i20S
PA28ab + std20S

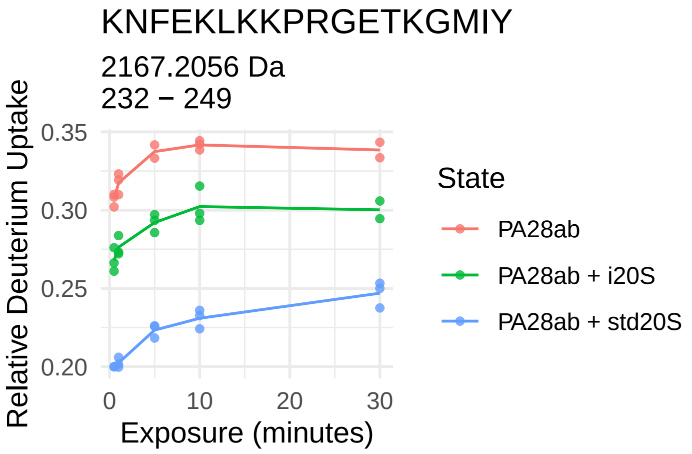
KNFEKLKKPRGETKGMIY
2167.2056 Da
232 − 249
Relative Deuterium Uptake
0.35
0.30
0.25
0.20
0
10
20
30
Exposure (minutes)
State
PA28ab
PA28ab + i20S
PA28ab + std20S

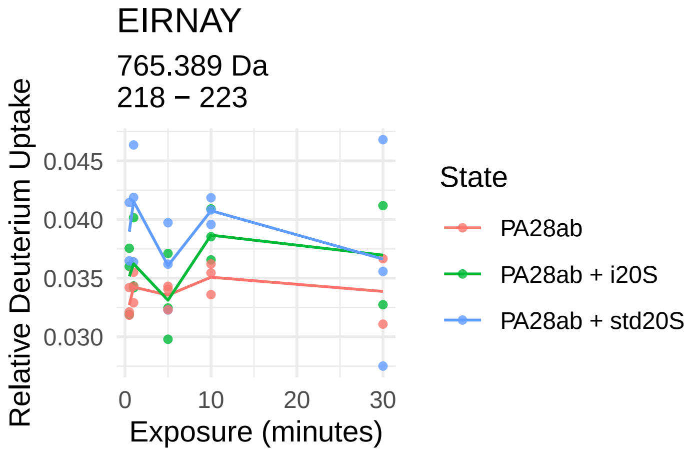
EIRNAY
765.389 Da
218 − 223
Relative Deuterium Uptake
0.045
0.040
0.035
0.030
0
10
20
30
Exposure (minutes)
State
PA28ab
PA28ab + i20S
PA28ab + std20S

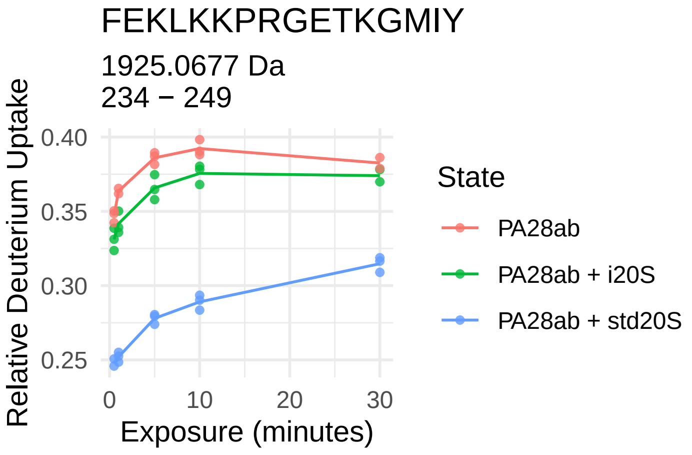
FEKLKKPRGETKGMIY
1925.0677 Da
234 − 249
Relative Deuterium Uptake
0.40
0.35
0.30
0.25
0
10
20
30
Exposure (minutes)
State
PA28ab
PA28ab + i20S
PA28ab + std20S

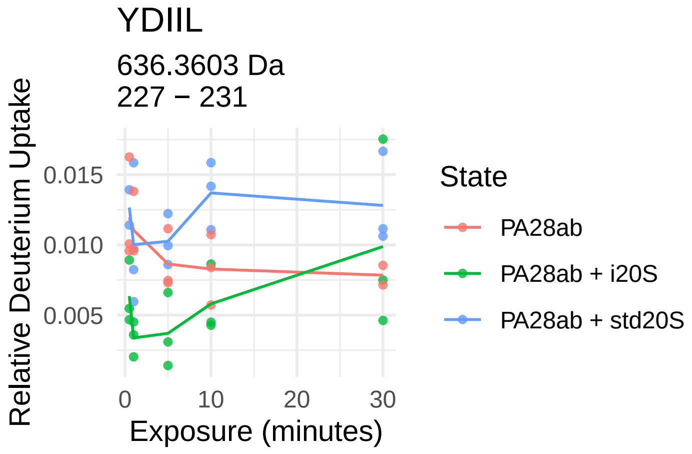
YDIIL
636.3603 Da
227 − 231
Relative Deuterium Uptake
0.015
0.010
0.005
0
10
20
30
Exposure (minutes)
State
PA28ab
PA28ab + i20S
PA28ab + std20S

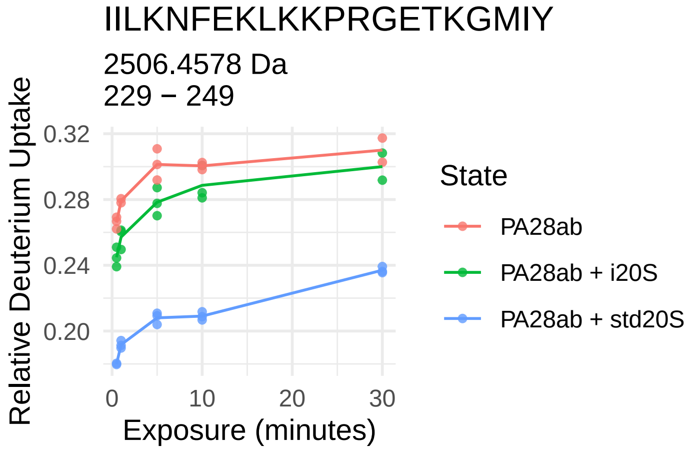
IILKNFEKLKKPRGETKGMIY
2506.4578 Da
229 − 249
Relative Deuterium Uptake
0.32
0.28
0.24
0.20
0
10
20
30
Exposure (minutes)
State
PA28ab
PA28ab + i20S
PA28ab + std20S

# PA28β

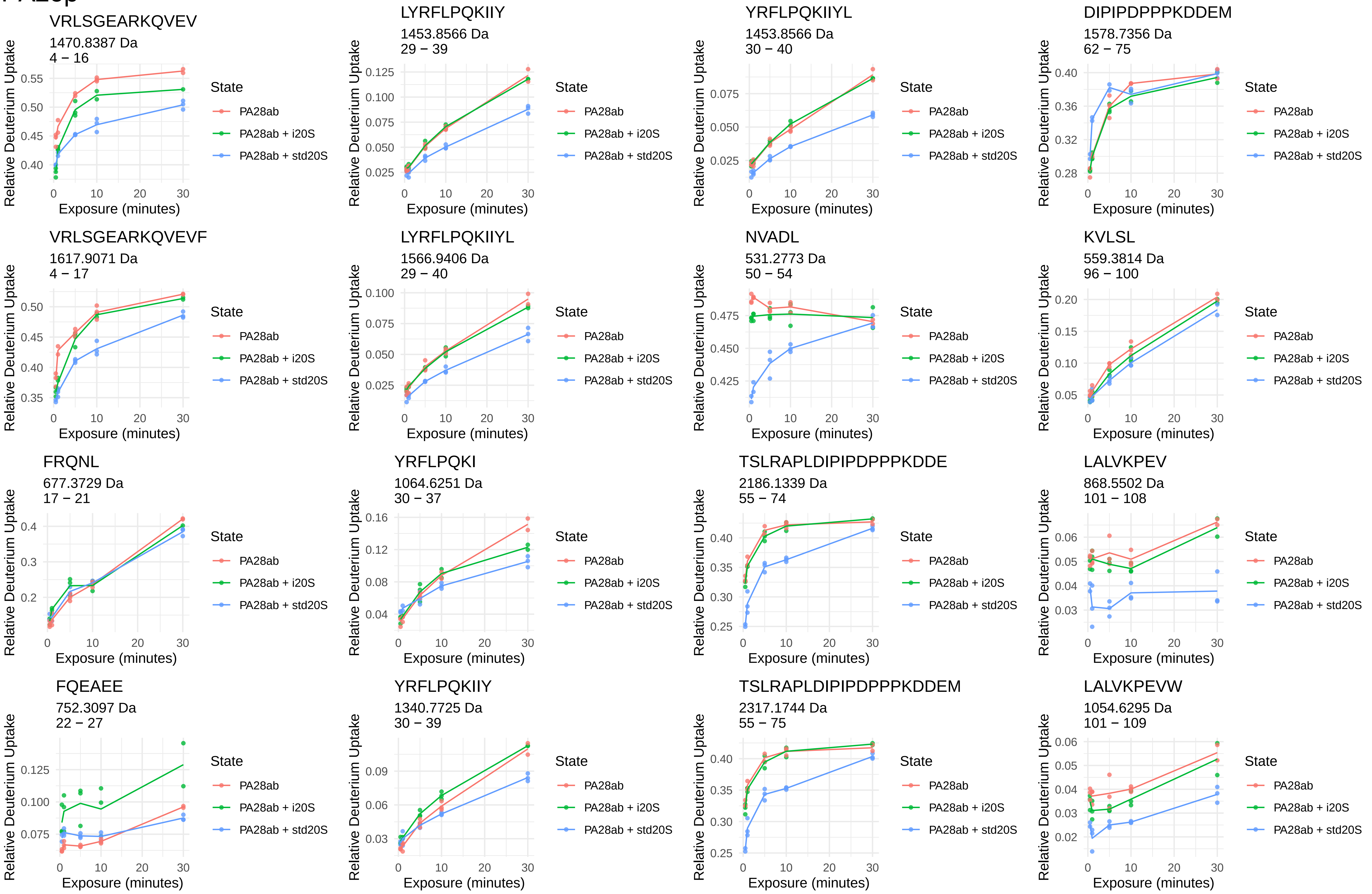
VRLSGEARKQVEV
1470.8387 Da
4 – 16
LYRFLPQKIIY
1453.8566 Da
29 – 39
YRFLPQKIIYL
1453.8566 Da
30 – 40
DIPIPDPPPKDDEM
1578.7356 Da
62 – 75
VRLSGEARKQVEVF
1617.9071 Da
4 – 17
LYRFLPQKIIYL
1566.9406 Da
29 – 40
NVADL
531.2773 Da
50 – 54
KVLSL
559.3814 Da
96 – 100
FRQNL
677.3729 Da
17 – 21
YRFLPQKI
1064.6251 Da
30 – 37
TSLRAPLDIPIPDPPPKDDE
2186.1339 Da
55 – 74
LALVKPEV
868.5502 Da
101 – 108
FQEAEE
752.3097 Da
22 – 27
YRFLPQKIIY
1340.7725 Da
30 – 39
TSLRAPLDIPIPDPPPKDDEM
2317.1744 Da
55 – 75
LALVKPEVW
1054.6295 Da
101 – 109
Relative Deuterium Uptake
Exposure (minutes)
State
PA28ab
PA28ab + i20S
PA28ab + std20S

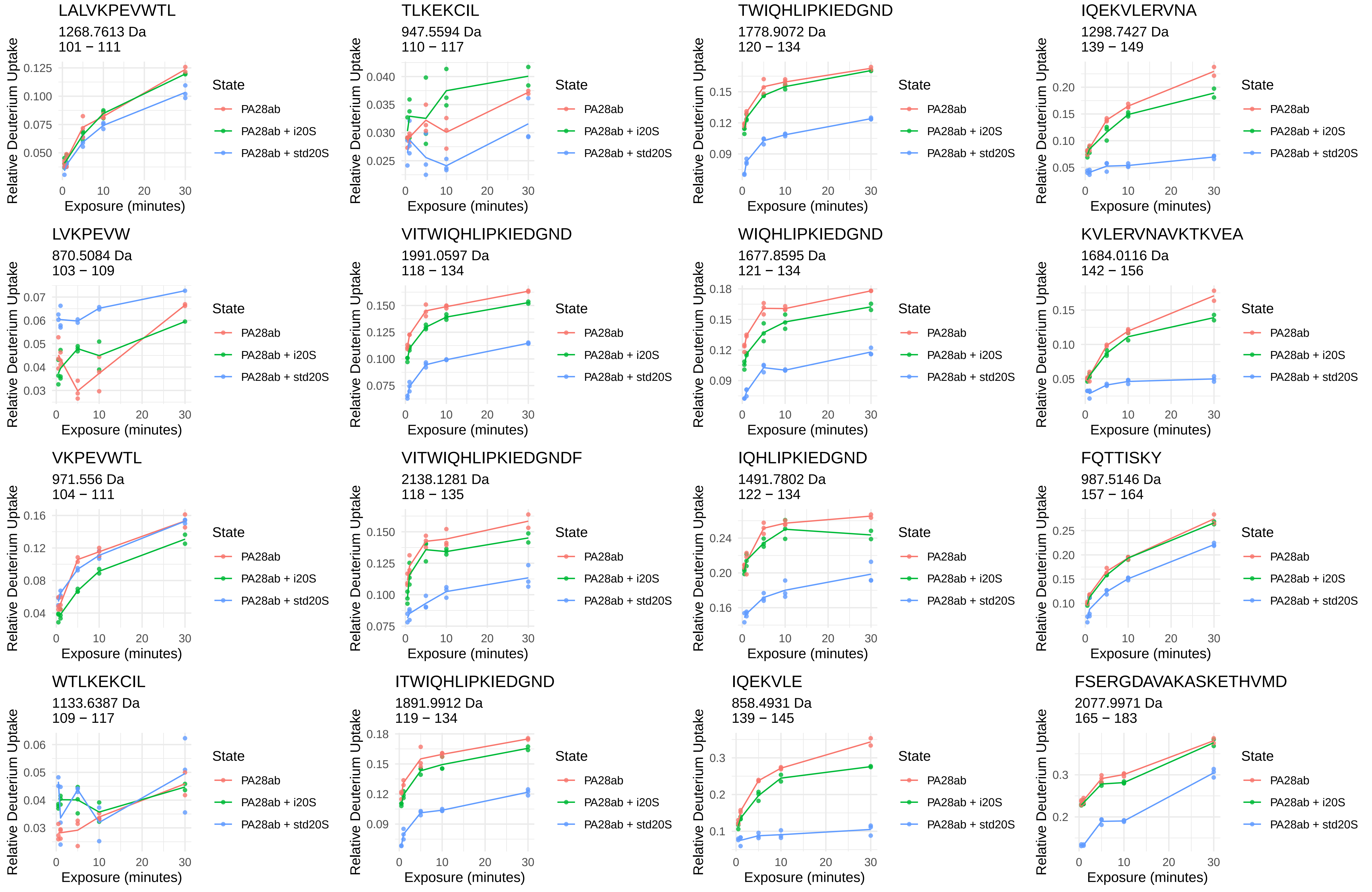
LALVKPEVWTL
1268.7613 Da
101 − 111
TLKEKCIL
947.5594 Da
110 − 117
TWIQHLIPKIEDGND
1778.9072 Da
120 − 134
IQEKVLERVNA
1298.7427 Da
139 − 149
LVKPEVW
870.5084 Da
103 − 109
VITWIQHLIPKIEDGND
1991.0597 Da
118 − 134
WIQHLIPKIEDGND
1677.8595 Da
121 − 134
KVLERVNAVKTKVEA
1684.0116 Da
142 − 156
VKPEVWTL
971.556 Da
104 − 111
VITWIQHLIPKIEDGNDF
2138.1281 Da
118 − 135
IQHLIPKIEDGND
1491.7802 Da
122 − 134
FQTTISKY
987.5146 Da
157 − 164
WTLKEKCIL
1133.6387 Da
109 − 117
ITWIQHLIPKIEDGND
1891.9912 Da
119 − 134
IQEKVLE
858.4931 Da
139 − 145
FSERGDAVAKASKETHVMD
2077.9971 Da
165 − 183
Relative Deuterium Uptake
Exposure (minutes)
State
PA28ab
PA28ab + i20S
PA28ab + std20S

FSERGDAVAKASKETHVMDY
2241.0605 Da
165 − 184

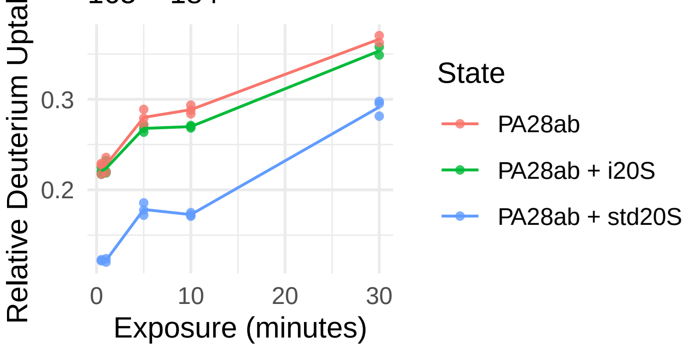


YHIISSNL
946.4993 Da
214 − 221

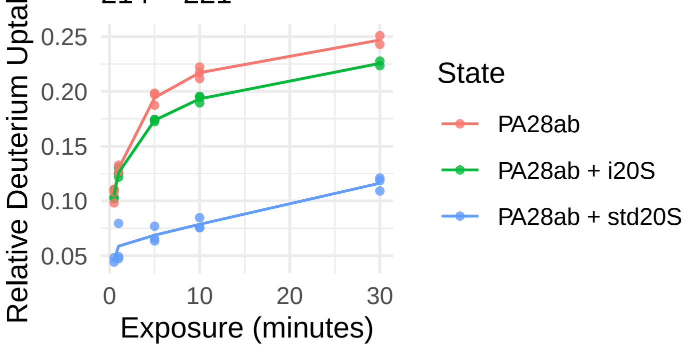


YRALVHERDEAA
1429.7183 Da
184 − 195

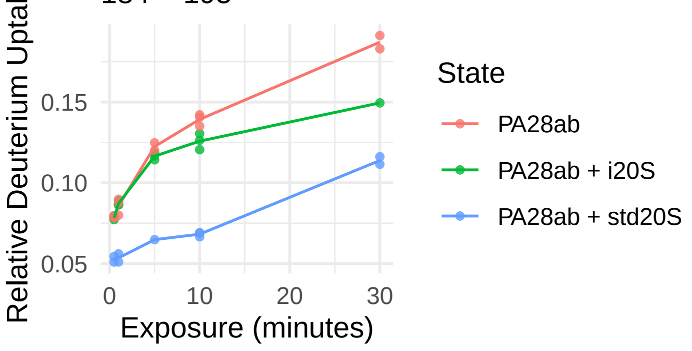


EKIVNPKGEEKPSM
1585.8254 Da
222 − 235

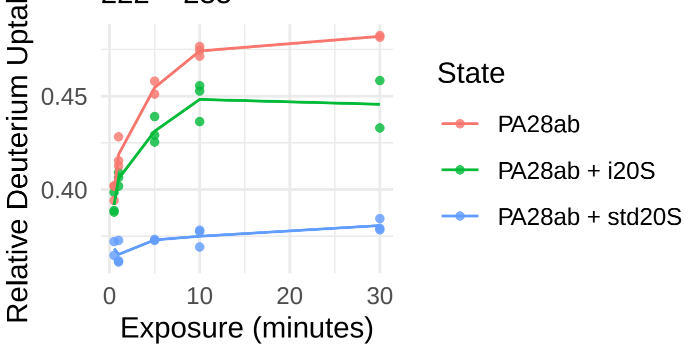


MVLDL
590.3218 Da
202 − 206

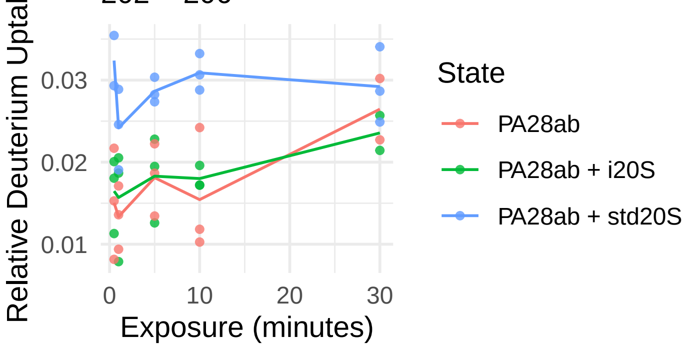


EKIVNPKGEEKPSMY
1748.8887 Da
222 − 236

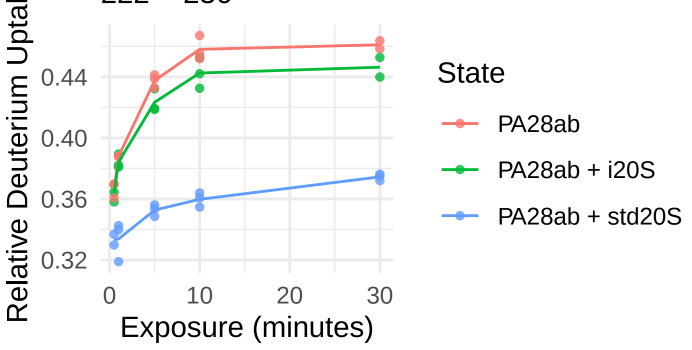


FYAEL
642.3134 Da
209 − 213

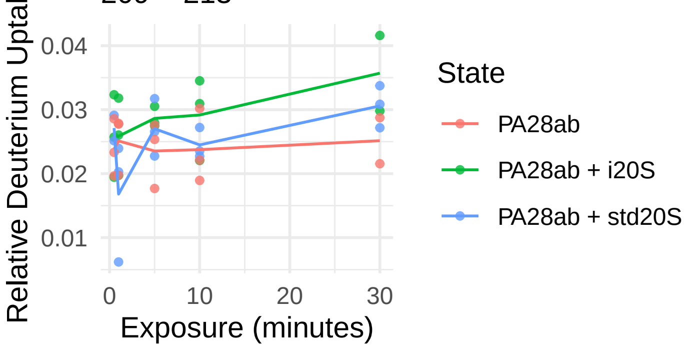

PA28γ

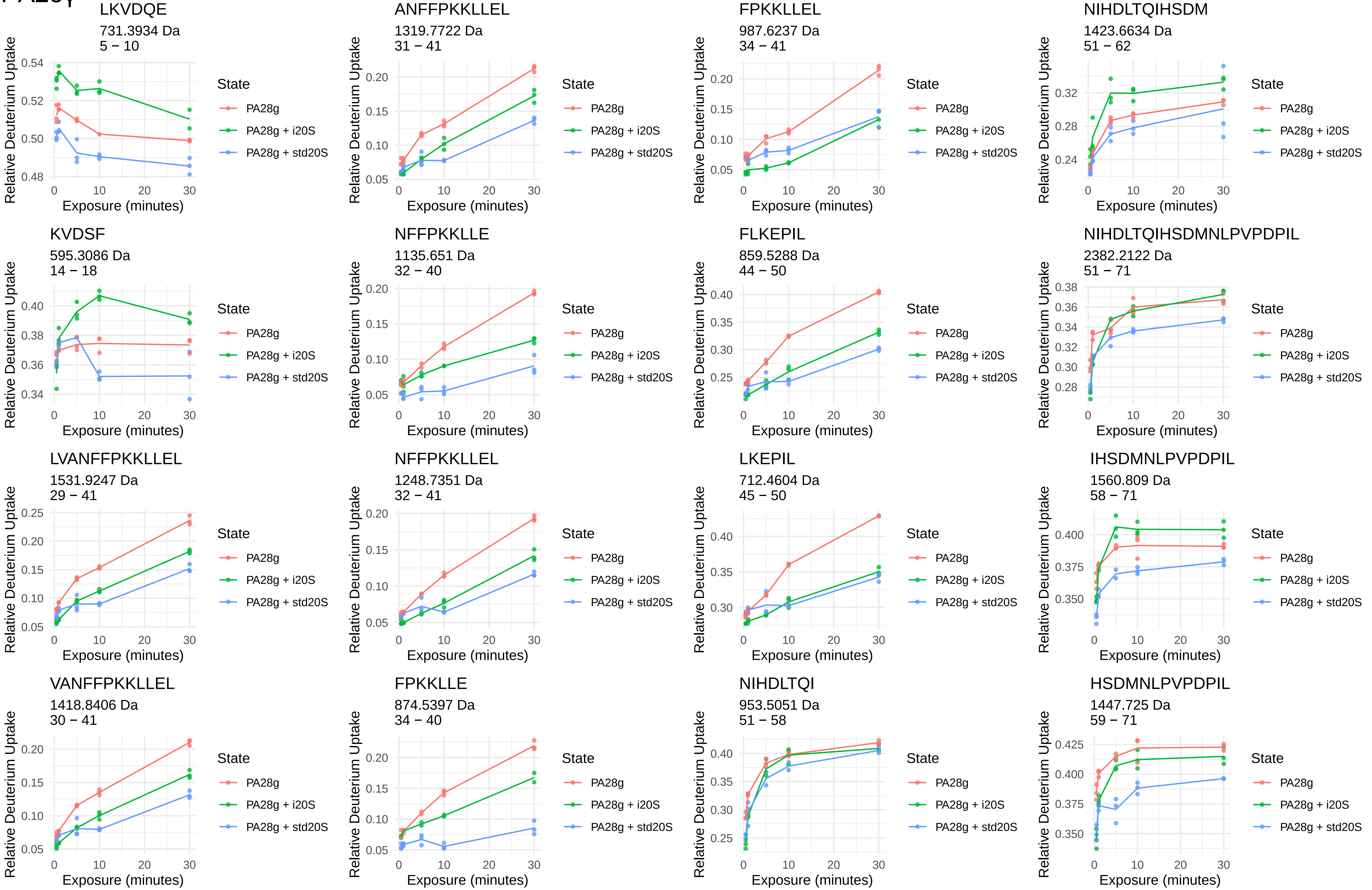
LKVDQE
731.3934 Da
5 − 10
ANFFPKKLLEL
1319.7722 Da
31 − 41
FPKKLLEL
987.6237 Da
34 − 41
NIHDLTQIHSDM
1423.6634 Da
51 − 62
KVDSF
595.3086 Da
14 − 18
NFFPKKLLE
1135.651 Da
32 − 40
FLKEPIL
859.5288 Da
44 − 50
NIHDLTQIHSDMNLPVPDPIL
2382.2122 Da
51 − 71
LVANFFPKKLLEL
1531.9247 Da
29 − 41
NFFPKKLLEL
1248.7351 Da
32 − 41
LKEPIL
712.4604 Da
45 − 50
IHSDMNLPVPDPIL
1560.809 Da
58 − 71
VANFFPKKLLEL
1418.8406 Da
30 − 41
FPKKLLE
874.5397 Da
34 − 40
NIHDLTQI
953.5051 Da
51 − 58
HSDMNLPVPDPIL
1447.725 Da
59 − 71
Relative Deuterium Uptake
Exposure (minutes)
State
PA28g
PA28g + i20S
PA28g + std20S

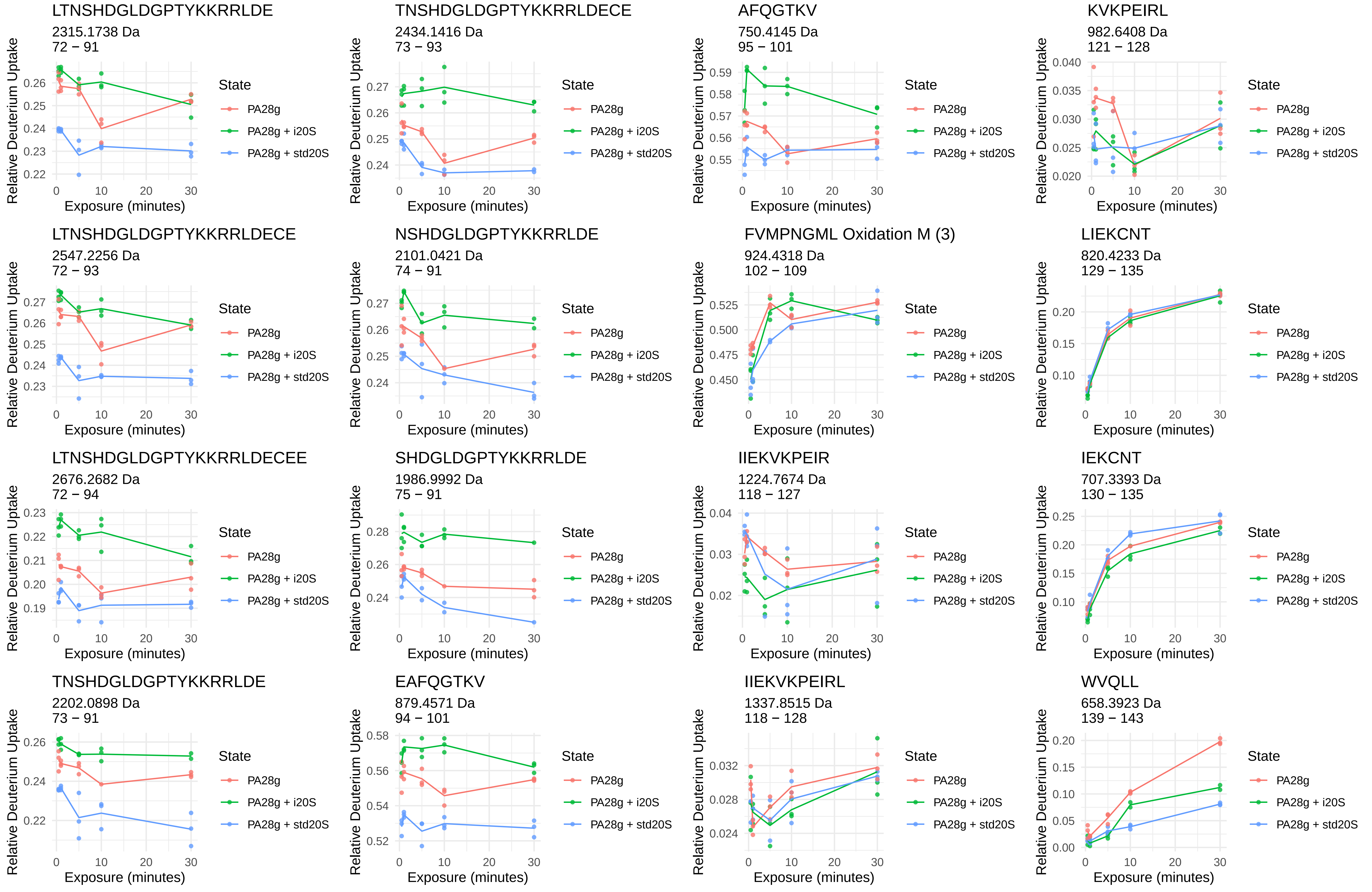
LTNSHDGLDGPTYKKRRLDE
2315.1738 Da
72 – 91
TNSHDGLDGPTYKKRRLDECE
2434.1416 Da
73 – 93
AFQGTKV
750.4145 Da
95 – 101
KVKPEIRL
982.6408 Da
121 – 128
LTNSHDGLDGPTYKKRRLDECE
2547.2256 Da
72 – 93
NSHDGLDGPTYKKRRLDE
2101.0421 Da
74 – 91
FVMPNGML Oxidation M (3)
924.4318 Da
102 – 109
LIEKCNT
820.4233 Da
129 – 135
LTNSHDGLDGPTYKKRRLDECEE
2676.2682 Da
72 – 94
SHDGLDGPTYKKRRLDE
1986.9992 Da
75 – 91
IIEKVKPEIR
1224.7674 Da
118 – 127
IEKCNT
707.3393 Da
130 – 135
TNSHDGLDGPTYKKRRLDE
2202.0898 Da
73 – 91
EAFQGTKV
879.4571 Da
94 – 101
IIEKVKPEIRL
1337.8515 Da
118 – 128
WVQLL
658.3923 Da
139 – 143
Relative Deuterium Uptake
Exposure (minutes)
State
PA28g
PA28g + i20S
PA28g + std20S

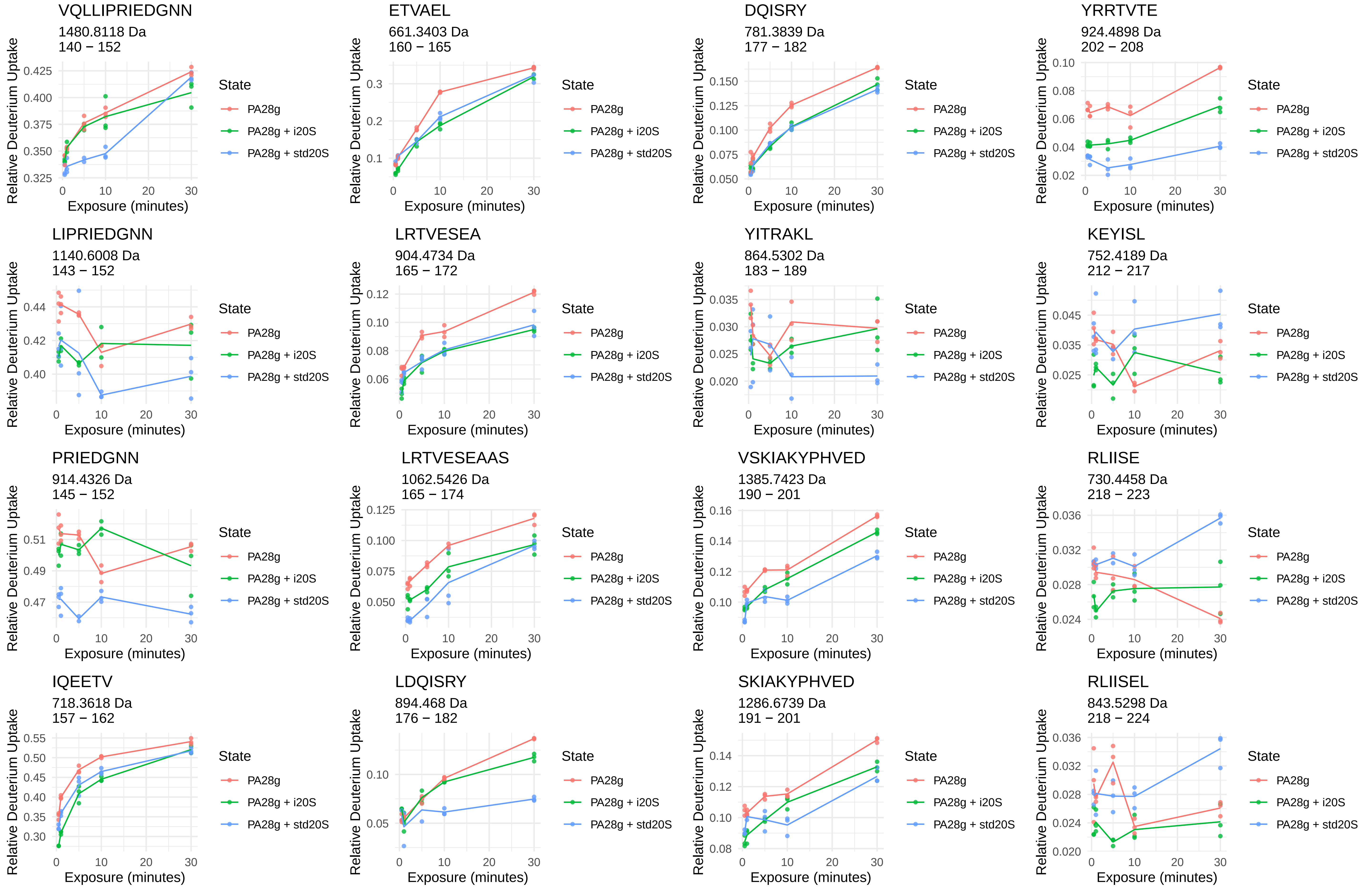
VQLLIPRIEDGNN
1480.8118 Da
140 − 152
ETVAEL
661.3403 Da
160 − 165
DQISRY
781.3839 Da
177 − 182
YRRTVTE
924.4898 Da
202 − 208
LIPRIEDGNN
1140.6008 Da
143 − 152
LRTVESEA
904.4734 Da
165 − 172
YITRAKL
864.5302 Da
183 − 189
KEYISL
752.4189 Da
212 − 217
PRIEDGNN
914.4326 Da
145 − 152
LRTVESEAAS
1062.5426 Da
165 − 174
VSKIAKYPHVED
1385.7423 Da
190 − 201
RLIISE
730.4458 Da
218 − 223
IQEETV
718.3618 Da
157 − 162
LDQISRY
894.468 Da
176 − 182
SKIAKYPHVED
1286.6739 Da
191 − 201
RLIISEL
843.5298 Da
218 − 224
Relative Deuterium Uptake
Exposure (minutes)
State
PA28g
PA28g + i20S
PA28g + std20S

IISEL
574.3447 Da
220 – 224

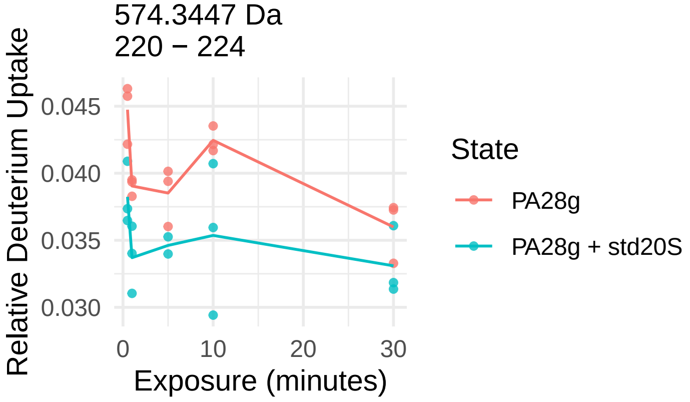


KNIEKIKRPRSSNA
1640.9555 Da
237 – 250

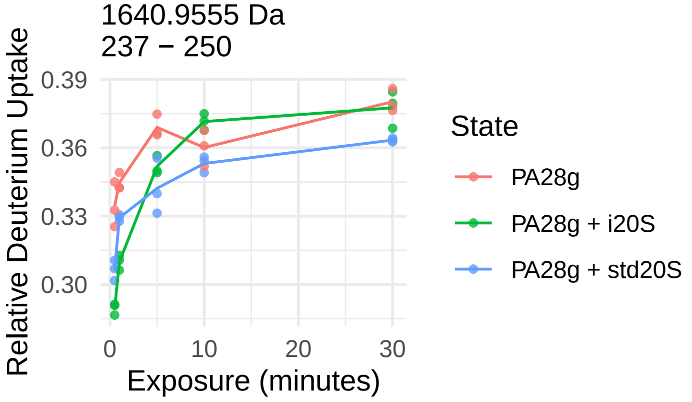


LRNQY
693.3678 Da
224 – 228

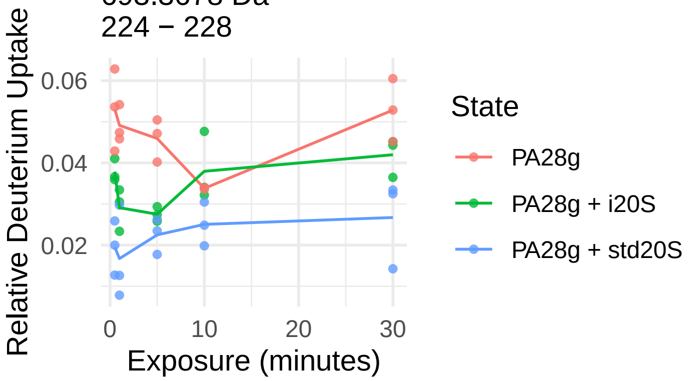


LRNQYVTL
1006.568 Da
224 – 231

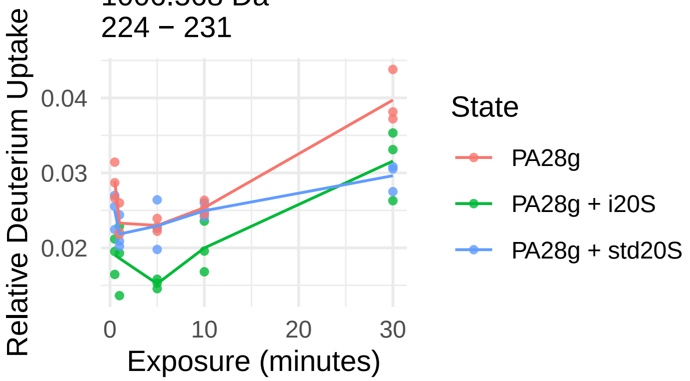


RNQYVTL
893.4839 Da
225 – 231

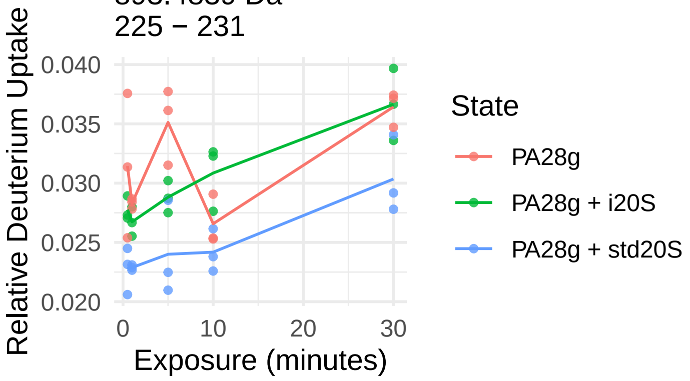
