## Supplementary material for "Conformational maps of human 20S proteasomes reveal PA28- and immuno-dependent inter-ring crosstalks": Fig. S11

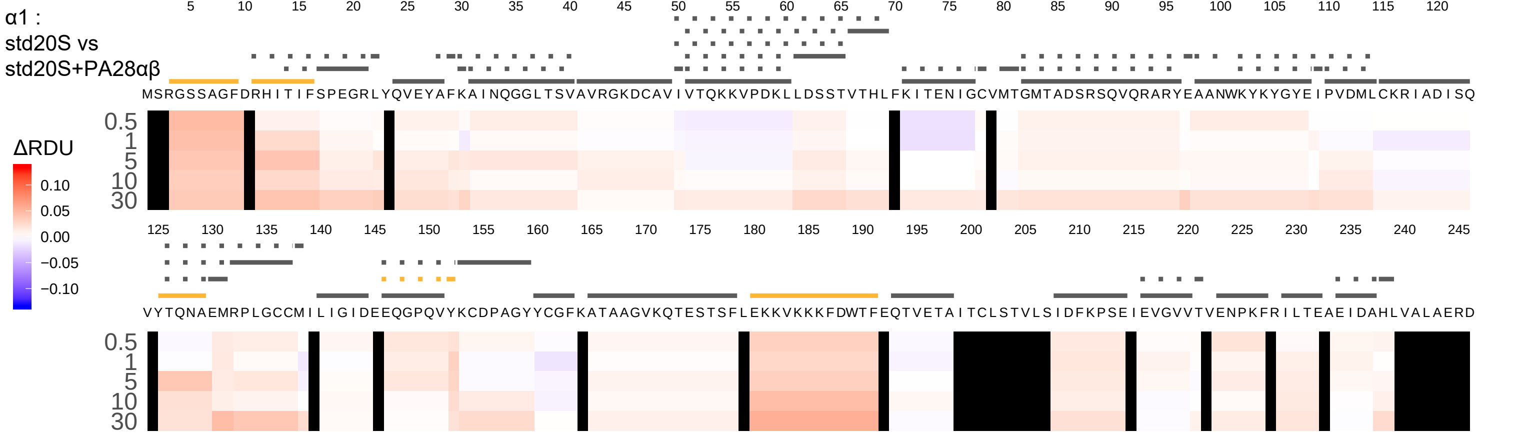

α1 :
std20S vs
std20S+PA28αβ
ΔRDU
0.10
0.05
0.00
−0.05
−0.10
5 10 15 20 25 30 35 40 45 50 55 60 65 70 75 80 85 90 95 100 105 110 115 120
MSRGSSAGFDRHITIFSPEGRLYQVEYAFKAINQGGLTSVAVRGKDCAVIVTQKKVPDKLLDSSTVTHLFKITENIGCVMTGMTADSRSQVQRARYEAANWKYKYGYEIPVDMLCKRIADISQ
0.5
1
5
10
30
125 130 135 140 145 150 155 160 165 170 175 180 185 190 195 200 205 210 215 220 225 230 235 240 245
VYTQNAEMRPLGCCMILIGIDEEQGPQVYKCDPAGYYCGFKATAAGVKQTESTSFLEKKVKKKFDWTFEQTVETAITCLSTVLSIDFKPSEIEVGVVTVENPKFRILTEAEIDAHLVALAERD
0.5
1
5
10
30

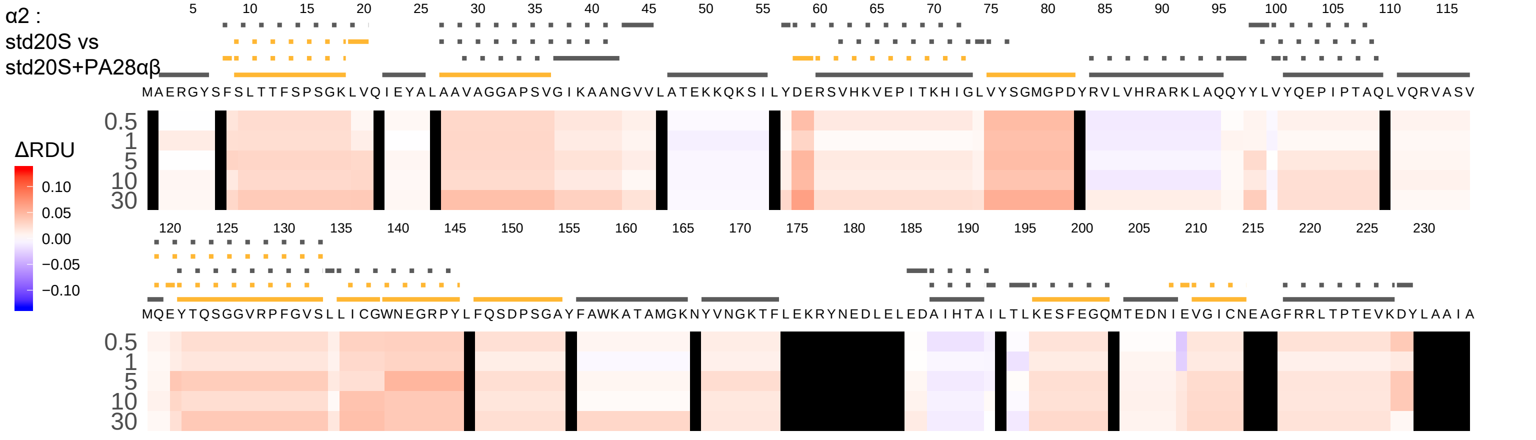
α2 :
std20S vs
std20S+PA28αβ
5 10 15 20 25 30 35 40 45 50 55 60 65 70 75 80 85 90 95 100 105 110 115
MAERGYSFSLTTFSPSGKLVQIEYALAAVAGGAPSVGIKAANGVVLATEKKQKSILYDERSVHKVEPITKHIGLVYSGMGPDYRVLVHRARKLAQQYYLVYQEPIPTAQLVQRVASV
0.5
1
5
10
30
ΔRDU
0.10
0.05
0.00
−0.05
−0.10
120 125 130 135 140 145 150 155 160 165 170 175 180 185 190 195 200 205 210 215 220 225 230
MQEYTQSGGVRPFGVSLLICGWNEGRPYLFQSDPSGAYFAWKATAMGKNYVNGKTFLEKRYNEDLELEDAIHTAILTLKESFEGQMTEDNIEVGICNEAGFRRLTPTEVKDYLAAIA
0.5
1
5
10
30

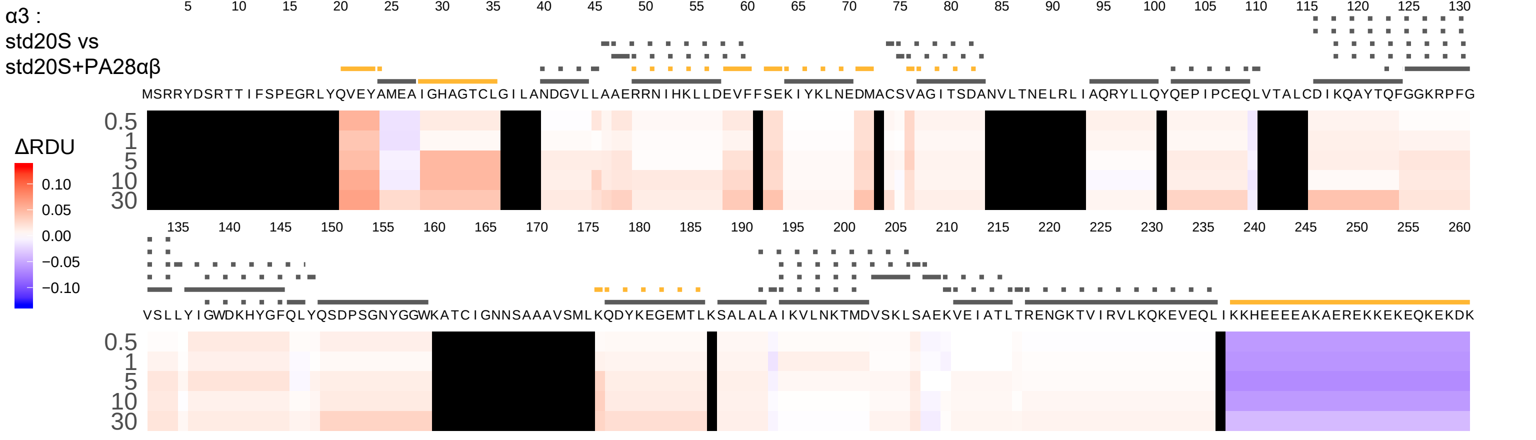
α3 :
std20S vs
std20S+PA28αβ
MSRRYDSRTTIFSPEGRLYQVEYAMEAIGHAGTCLGILANDGVLLAAERRNIHKLLDEVFFSEKIYKLNEDMACSVAGITSDANVLTNELRLIAQRYLLQYQEPIPCEQLVTALCDIKQAYTQFGGKRPFG
VSLLYIGWDKHYGFQLYQSDPSGNYGGWKATCIGNNSAAAVSMLKQDYKEGEMTLKSALALAIKVLNKTMDVSKLSAEKVEIATLTRENGKTVIRVLKQKEVEQLIKKHEEEEAKAEREKKEKEQKEKDK
0.5
1
5
10
30
ΔRDU
0.10
0.05
0.00
−0.05
−0.10

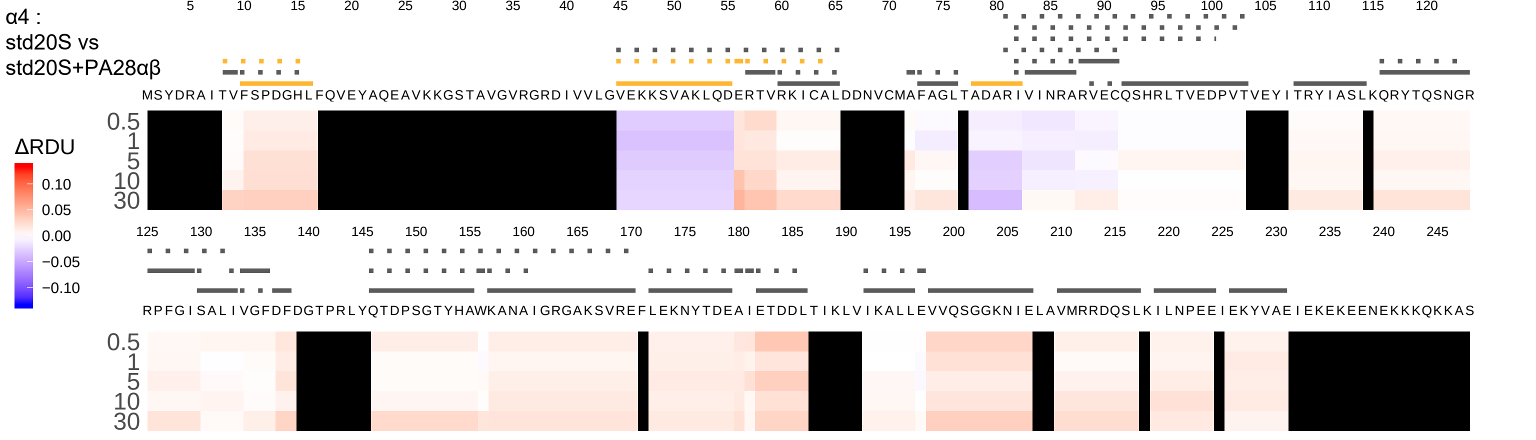

α4 :
std20S vs
std20S+PA28αβ
ΔRDU
0.10
0.05
0.00
−0.05
−0.10
0.5
1
5
10
30
MSYDRAITVFSPDGHLFQVEYAQEAVKKGSTAVGVRGRDIVVLGVEKKSVAKLQDERTVRKICALDDNVCMAFAGLTADARIVINRARVECQSHRLTVEDPVTVEYITRYIASLKQRYTQSNGR
RPFGISALIVGFDFDGTPRLYQTDPSGTYHAWKANAIGRGAKSVREFLEKNYTDEAIETDDLTIKLVIKALLEVVQSGGKNIELAVMRRDQSLKILNPEEIEKYVAEIEKEKEENEKKKQKKAS

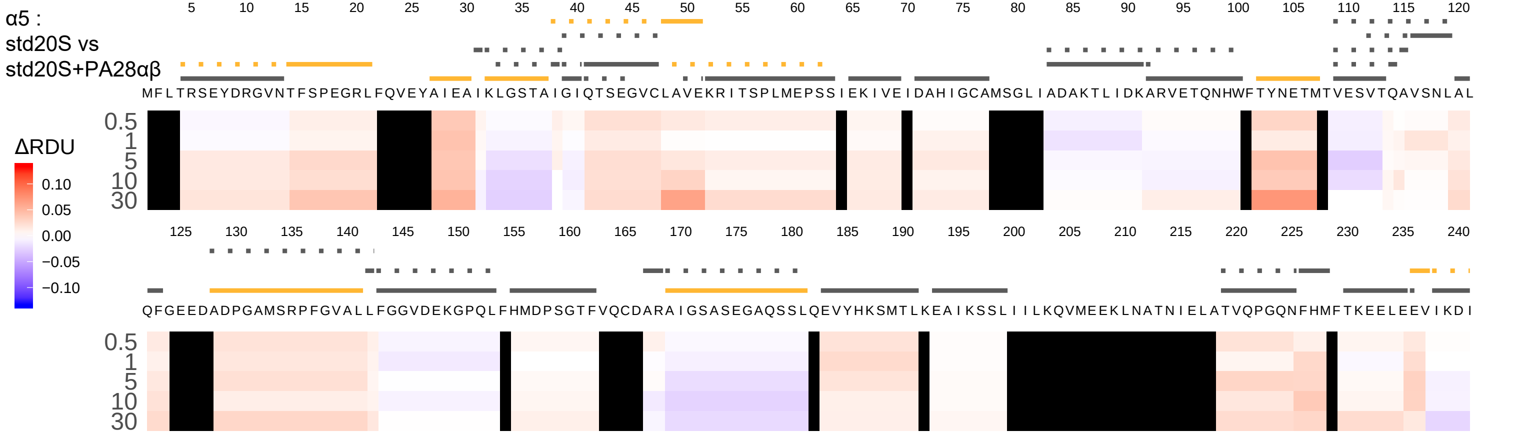

α5 :
std20S vs
std20S+PA28αβ
MFLTRSEYDRGVNTFSPEGRLFQVEYAIEAIKLGSTAIGIQTSEGVCLAVEKRITSPLMEPSSIEKIVEIDAHIGCAMSGLIADAKTLIDKARVETQNHWFTYNETMTVESVTQAVSNLAL
QFGEEDADPGAMSRPFGVALLFGGVDEKGPQLFHMDPSGTFVQCDARAIGSASEGAQSSLQEVYHKSMTLKEAIKSSLIIILKQVMEEKLNATNIELATVQPGQNFHMFTKEELEEVIKDI
ΔRDU
0.10
0.05
0.00
−0.05
−0.10
0.5
1
5
10
30

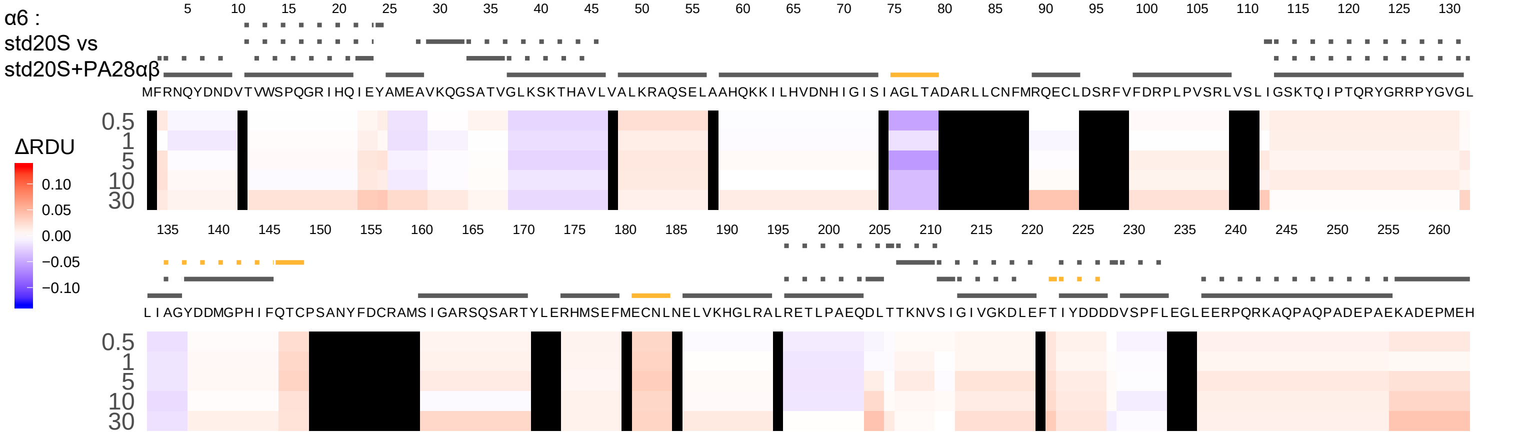
α6 :
std20S vs
std20S+PA28αβ
ΔRDU
0.10
0.05
0.00
−0.05
−0.10
MFRNQYDNDVTVWSPQGRIHQIEYAMEAVKQGSATVGLKSKTHAVLVALKRAQSELAAHQKKILHVDNHIGISIAGLTADARLLCNFMRQECLDSRFVFDRPLPVSRLVSLIGSKTQIPTQRYGRRPYGVGL
LIAGYDDMGPHIFQTCPSANYFDCRAMSIGARSQSARTYLERHMSEFMECNLNELVKHGLRALRETLPAEQDLTTKNVSIGIVGKDLEFTIYDDDDVSPFLEGLEERPQRKAQPAQPADEPAEKADEPMEH
0.5
1
5
10
30

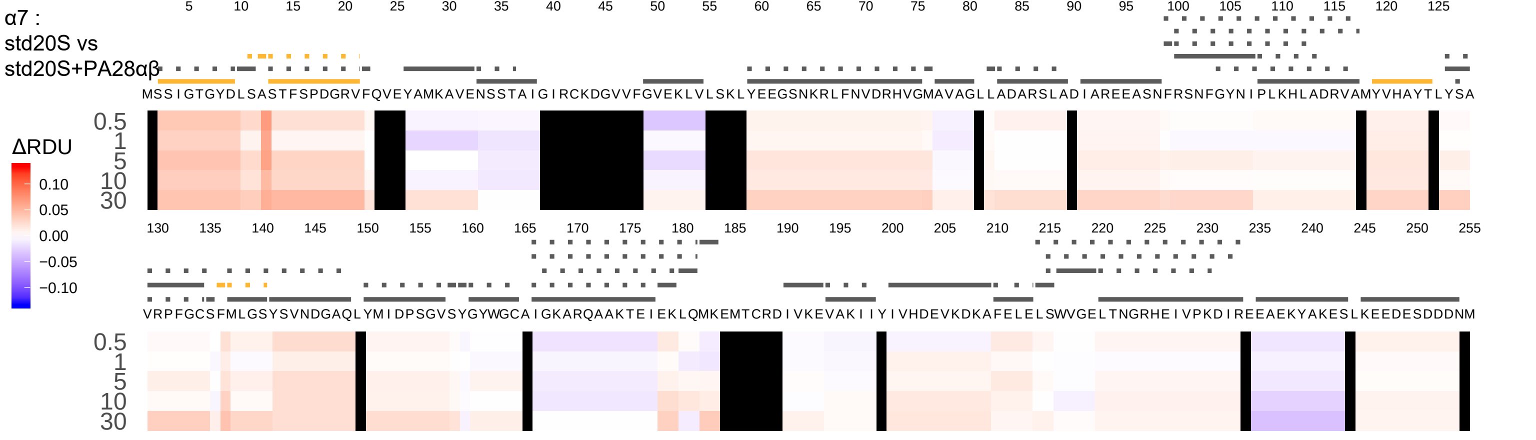

α7 :
std20S vs
std20S+PA28αβ
ΔRDU
0.10
0.05
0.00
−0.05
−0.10
0.5
1
5
10
30
MSSIGTGYDLSASTFSPDGRVFQVEYAMKAVENSSTAIGIRCKDGVVFGVEKLVLSKLYEEGSNKRLFNVDRHVGMAVAGLLADARSLADIAREEASNFRSNFGYNIPLKHLADRVAMYVHAYTLYSA
VRPFGCSFMLGSYSVNDGAQLYMIDPSGVSYGYWGCAIGKARQAAKTEIEKLQMKEMTCRDIVKEVAKIIYIVHDEVKDKAFELELSWVGELTNGRHEIVPKDIREEAEKYAKESLKEEDESDDDNM

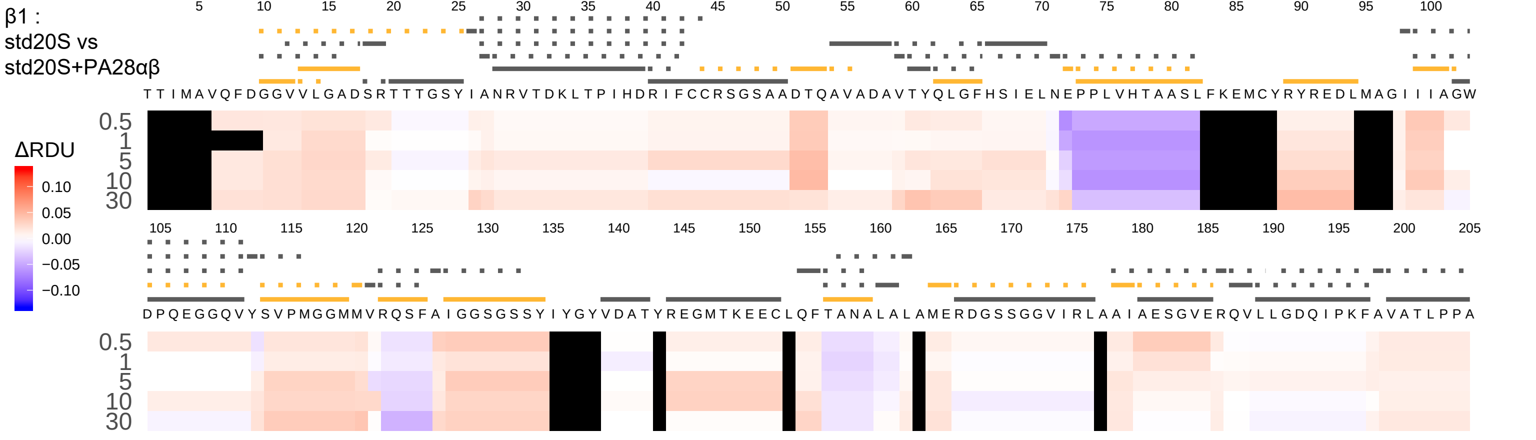

β1 :
std20S vs
std20S+PA28αβ
5 10 15 20 25 30 35 40 45 50 55 60 65 70 75 80 85 90 95 100
T T I M A V Q F D G G V V L G A D S R T T T G S Y I A N R V T D K L T P I H D R I F C C R S G S A A D T Q A V A D A V T Y Q L G F H S I E L N E P P L V H T A A S L F K E M C Y R Y R E D L M A G I I I A G W
0.5
1
5
10
30
ΔRDU
0.10
0.05
0.00
−0.05
−0.10
105 110 115 120 125 130 135 140 145 150 155 160 165 170 175 180 185 190 195 200 205
D P Q E G G Q V Y S V P M G G M M V R Q S F A I G G S G S S Y I Y G Y V D A T Y R E G M T K E E C L Q F T A N A L A L A M E R D G S S G G V I R L A A I A E S G V E R Q V L L G D Q I P K F A V A T L P P A
0.5
1
5
10
30

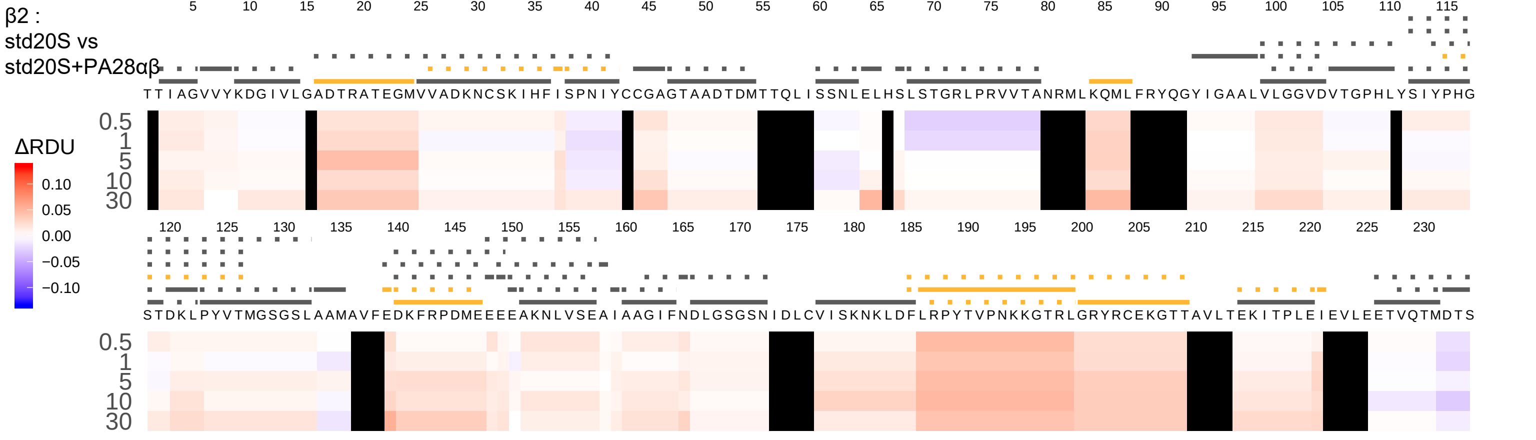

β2 :
std20S vs
std20S+PA28αβ
ΔRDU
0.10
0.05
0.00
−0.05
−0.10
0.5
1
5
10
30
TTIAGVVYKDGIVLGADTRATEGMVVADKNCSKIHFISPNIYCCGAGTAADTDMTTQLISSNLELHSLSTGRLPRVVTANRMLKQMLFRYQGYIGAALVLGGVDVTGPHLYSIYPHG
STDKLPYVTMGSGSLAAMAVFEDKFRPDMEEEEAKNLVSEAIAAGIFNDLGSGSNIDLCVISKNKLDFLRPYTVPNKKGTRLGRYRCEKGTTAVLTEKITPLEIEVLEETVQTMDTS

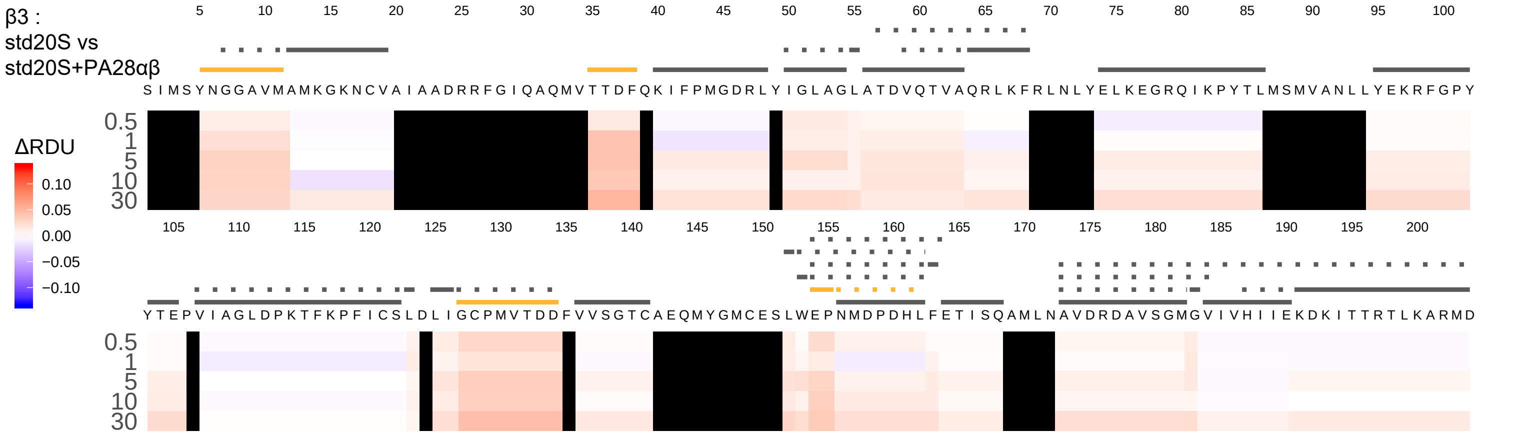
β3 :
std20S vs
std20S+PA28αβ
ΔRDU
0.10
0.05
0.00
−0.05
−0.10
0.5
1
5
10
30
S I M S Y N G G A V M A M K G K N C V A I A A D R R F G I Q A Q M V T T D F Q K I F P M G D R L Y I G L A G L A T D V Q T V A Q R L K F R L N L Y E L K E G R Q I K P Y T L M S M V A N L L Y E K R F G P Y
Y T E P V I A G L D P K T F K P F I C S L D L I G C P M V T D D F V V S G T C A E Q M Y G M C E S L W E P N M D P D H L F E T I S Q A M L N A V D R D A V S G M G V I V H I I E K D K I T T R T L K A R M D

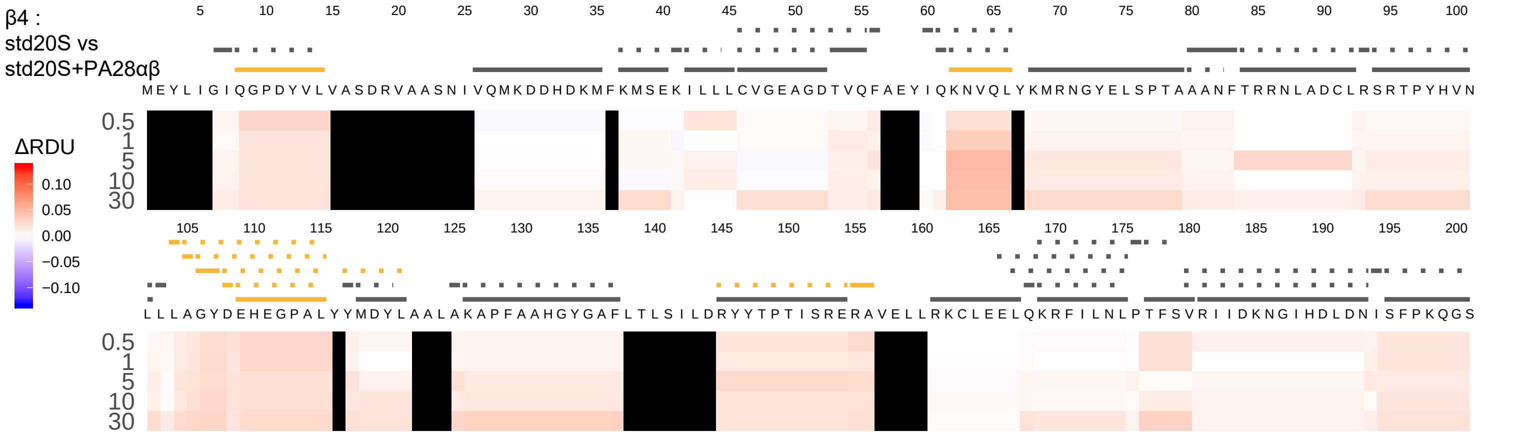
β4 :
std20S vs
std20S+PA28αβ
5 10 15 20 25 30 35 40 45 50 55 60 65 70 75 80 85 90 95 100
M E Y L I G I Q G P D Y V L V A S D R V A A S N I V Q M K D D H D K M F K M S E K I L L L C V G E A G D T V Q F A E Y I Q K N V Q L Y K M R N G Y E L S P T A A A N F T R R N L A D C L R S R T P Y H V N
0.5
1
5
10
30
ΔRDU
0.10
0.05
0.00
−0.05
−0.10
105 110 115 120 125 130 135 140 145 150 155 160 165 170 175 180 185 190 195 200
L L L A G Y D E H E G P A L Y Y M D Y L A A L A K A P F A A H G Y G A F L T L S I L D R Y Y T P T I S R E R A V E L L R K C L E E L Q K R F I L N L P T F S V R I I D K N G I H D L D N I S F P K Q G S
0.5
1
5
10
30

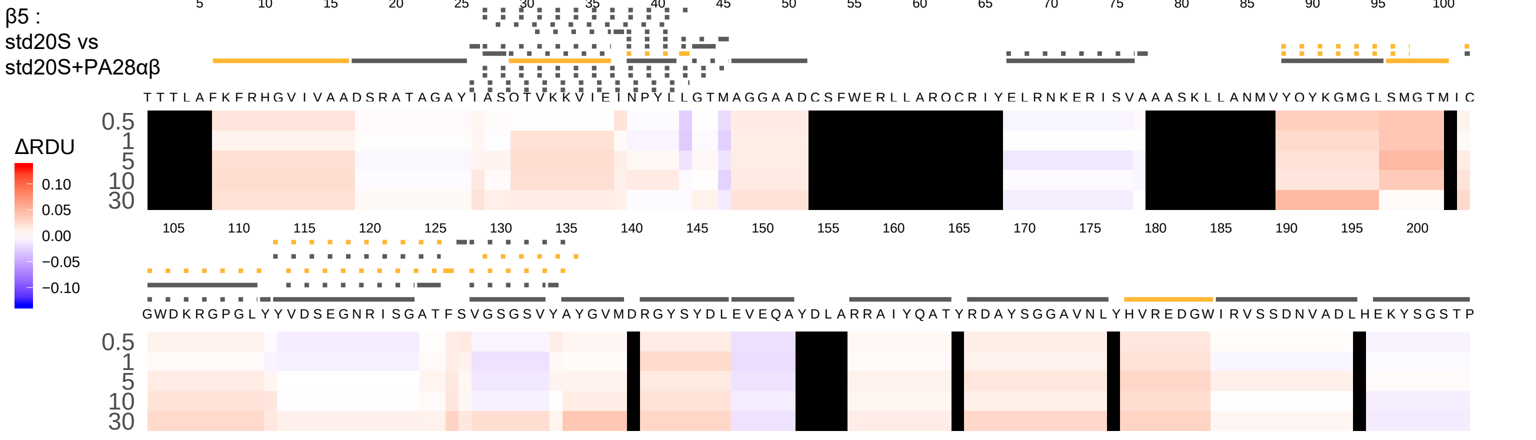

β5 :
std20S vs
std20S+PA28αβ
ΔRDU
0.10
0.05
0.00
−0.05
−0.10
0.5
1
5
10
30
5 10 15 20 25 30 35 40 45 50 55 60 65 70 75 80 85 90 95 100
105 110 115 120 125 130 135 140 145 150 155 160 165 170 175 180 185 190 195 200

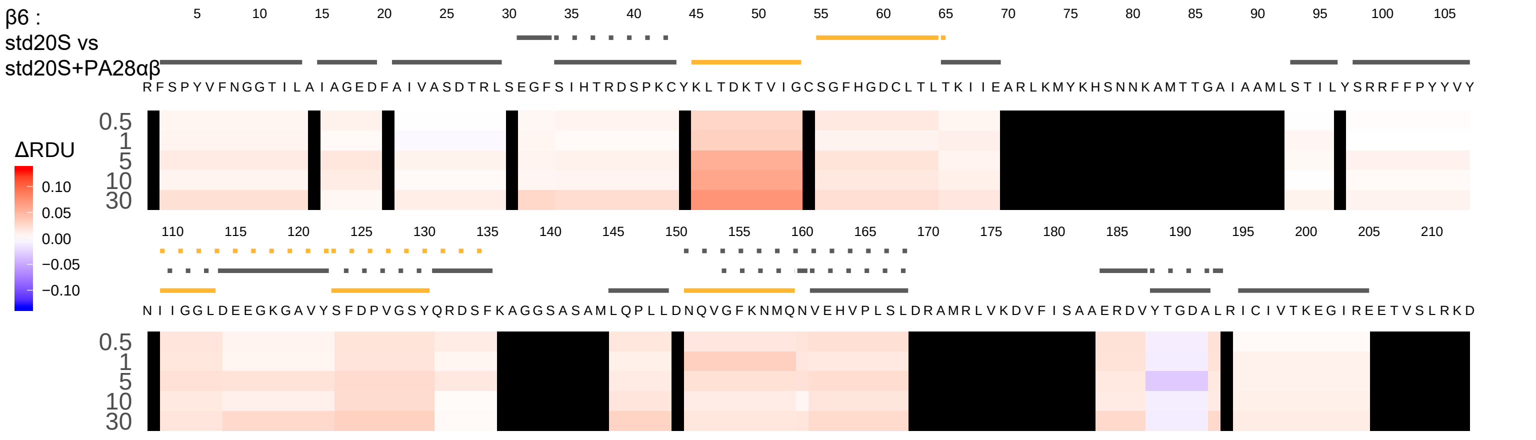

β6 :
std20S vs
std20S+PA28αβ
ΔRDU
0.10
0.05
0.00
−0.05
−0.10
0.5
1
5
10
30
RFSPYVFNGGTILAIAGEDFAIVASDTRLSEGFSIHTRDSPKCYKLTDKTVIGCSGFHGDCLTLTKIIEARLKMYKHSNNKAMTTGAIAAMLSTILYSRRFFPYYVY
NIIGGLDEEGKGAVYSFDPVGSYQRDSFKAGGSASAMLQPLLDNQVGFKNMQNVEHVPLSLDRAMRLVKDVFISAAERDVYTGDALRICIVTKEGIREETVSLRKD

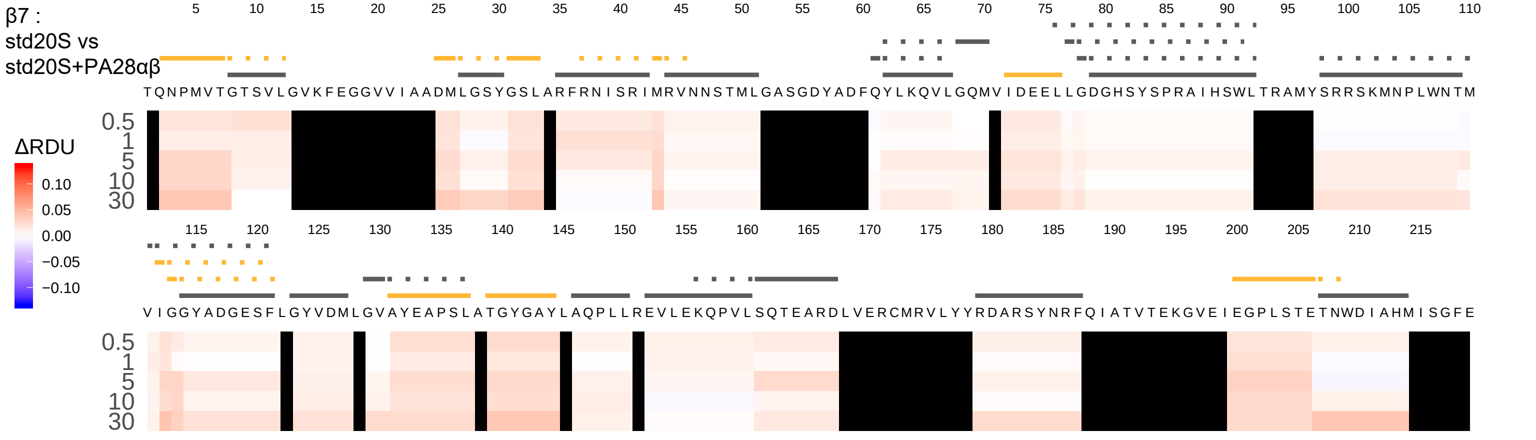
β7 :
std20S vs
std20S+PA28αβ
5 10 15 20 25 30 35 40 45 50 55 60 65 70 75 80 85 90 95 100 105 110
T Q N P M V T G T S V L G V K F E G G V V I A A D M L G S Y G S L A R F R N I S R I M R V N N S T M L G A S G D Y A D F Q Y L K Q V L G Q M V I D E E L L G D G H S Y S P R A I H S W L T R A M Y S R R S K M N P L W N T M
0.5
1
5
10
30
ΔRDU
0.10
0.05
0.00
−0.05
−0.10
115 120 125 130 135 140 145 150 155 160 165 170 175 180 185 190 195 200 205 210 215
V I G G Y A D G E S F L G Y V D M L G V A Y E A P S L A T G Y G A Y L A Q P L L R E V L E K Q P V L S Q T E A R D L V E R C M R V L Y Y R D A R S Y N R F Q I A T V T E K G V E I E G P L S T E T N W D I A H M I S G F E
0.5
1
5
10
30

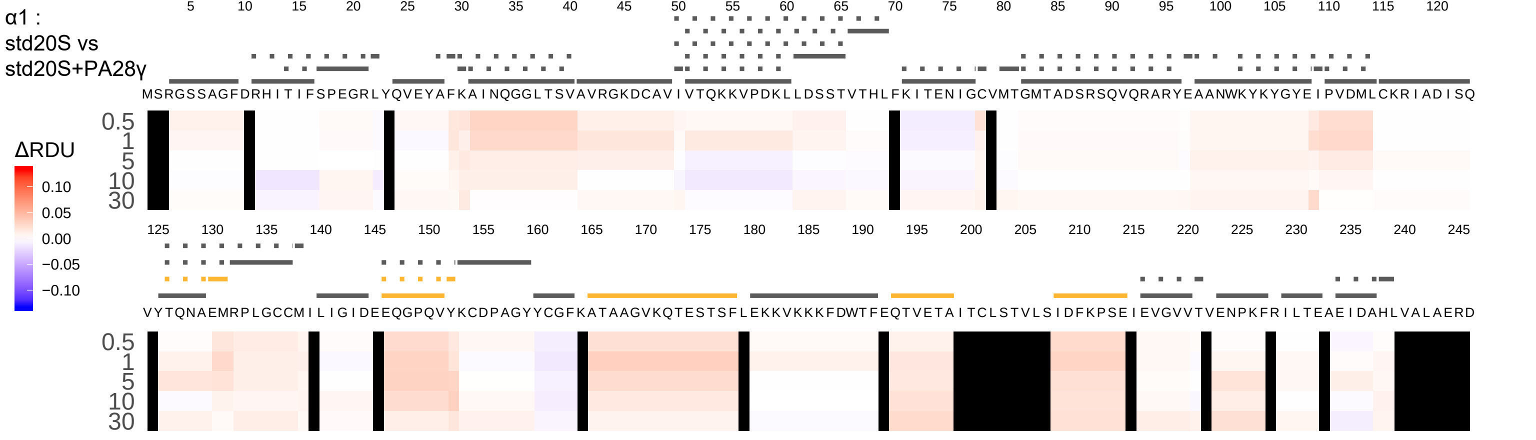
α1 :
std20S vs
std20S+PA28γ
ΔRDU
0.10
0.05
0.00
−0.05
−0.10
0.5
1
5
10
30
MSRGSSAGFDRHITIFSPEGRLYQVEYAFKAINQGGLTSVAVRGKDCAVIVTQKKVPDKLLDSSTVTHLFKITENIGCVMTGMTADSRSQVQRARYEAANWKYKYGYEIPVDMLCKRIADISQ
VYTQNAEMRPLGCCMILIGIDEEQGPQVYKCDPAGYYCGFKATAAGVKQTESTSFLEKKVKKKFDWTFEQTVETAITCLSTVLSIDFKPSEIEVGVVTVENPKFRILTEAEIDAHLVALAERD

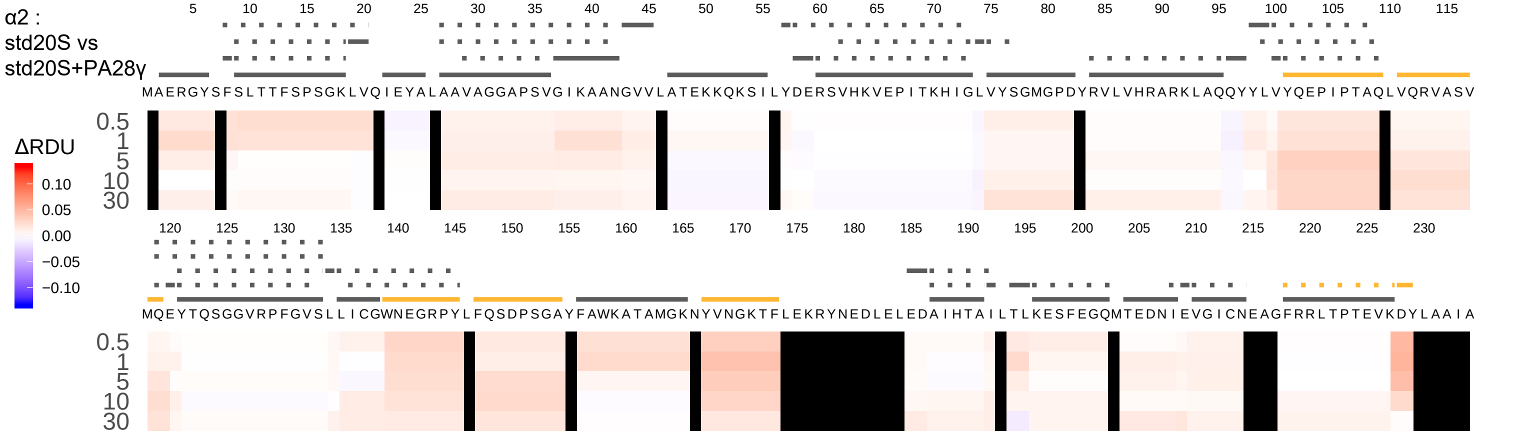

α2 :
std20S vs
std20S+PA28γ
ΔRDU
0.10
0.05
0.00
−0.05
−0.10
5 10 15 20 25 30 35 40 45 50 55 60 65 70 75 80 85 90 95 100 105 110 115
MAERGYSFSLTTFSPSGKLVQIEYALAAVAGGAPSVGIKAANGVVLATEKKQKSILYDERSVHKVEPITKHIGLVYSGMGPDYRVLVHRARKLAQQYYLVYQEPIPTAQLVQRVASV
0.5
1
5
10
30
120 125 130 135 140 145 150 155 160 165 170 175 180 185 190 195 200 205 210 215 220 225 230
MQEYTQSGGVRPFGVSLLICGWNEGRPYLFQSDPSGAYFAWKATAMGKNYVNGKTFLEKRYNEDLELEDAIHTAILTLKESFEGQMTEDNIEVGICNEAGFRRLTPTEVKDYLAAIA
0.5
1
5
10
30

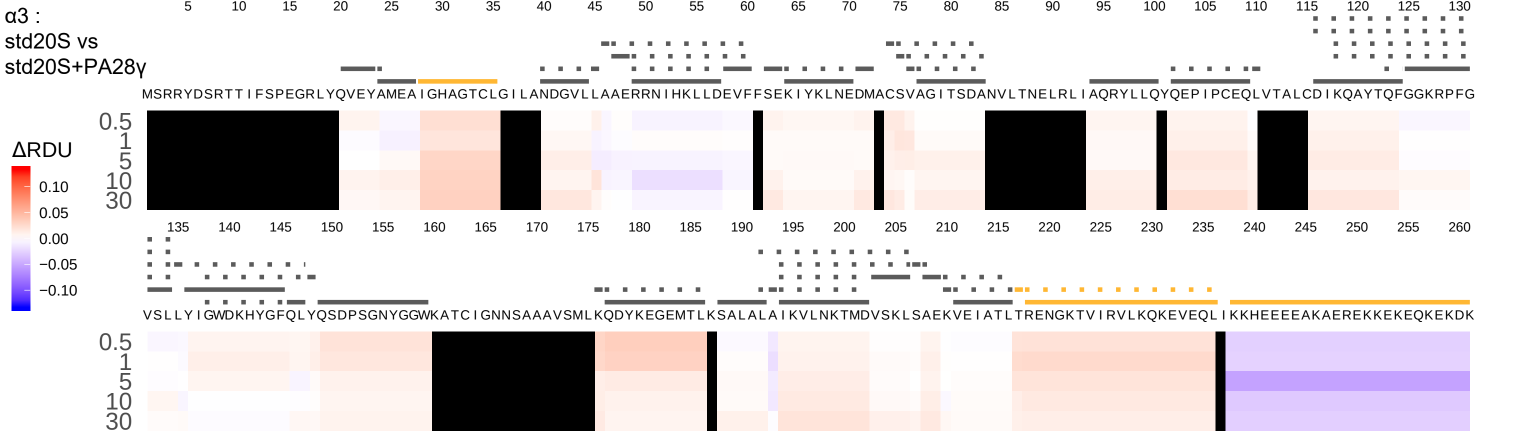
α3 :
std20S vs
std20S+PA28γ
5 10 15 20 25 30 35 40 45 50 55 60 65 70 75 80 85 90 95 100 105 110 115 120 125 130
MSRRYDSRTTIFSPEGRLYQVEYAMEAIGHAGTCLGILANDGVLLAAERRNIHKLLDEVFFSEKIYKLNEDMACSVAGITSDANVLTNELRLIAQRYLLQYQEPIPCEQLVTALCDIKQAYTQFGGKRPFG
0.5
1
5
10
30
135 140 145 150 155 160 165 170 175 180 185 190 195 200 205 210 215 220 225 230 235 240 245 250 255 260
VSLLYIGWDKHYGFQLYQSDPSGNYGGWKATCIGNNSAAAVSMLKQDYKEGEMTLKSALALAIKVLNKTMDVSKLSAEKVEIATLTRENGKTVIRVLKQKEVEQLIKKHEEEEAKAEREKKEKEQKEKDK
0.5
1
5
10
30
ΔRDU
0.10
0.05
0.00
−0.05
−0.10

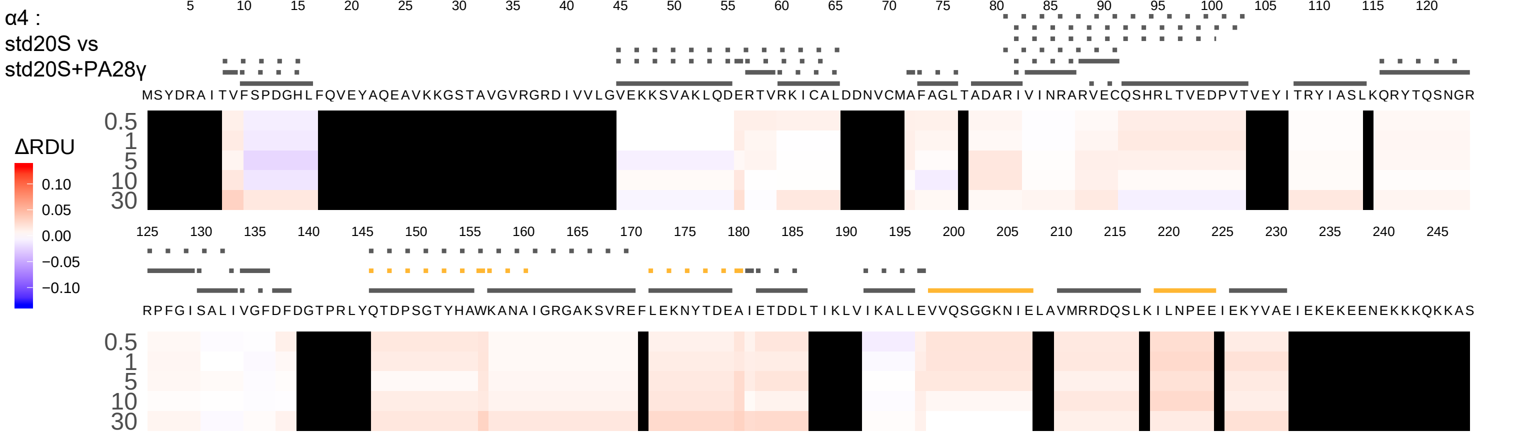
α4 :
std20S vs
std20S+PA28γ
ΔRDU
0.10
0.05
0.00
−0.05
−0.10
MSYDRAITVFSPDGHLFQVEYAQEAVKKGSTAVGVRGRDIVVLGVEKKSVAKLQDERTVRKICALDDNVCMAFAGLTADARIVINRARVECQSHRLTVEDPVTVEYITRYIASLKQRYTQSNGR
RPFGISALIVGFDFDGTPRLYQTDPSGTYHAWKANAIGRGAKSVREFLEKNYTDEAIETDDLTIKLVIKALLEVVQSGGKNIELAVMRRDQSLKILNPEEIEKYVAEIEKEKEENEKKKQKKAS
0.5
1
5
10
30

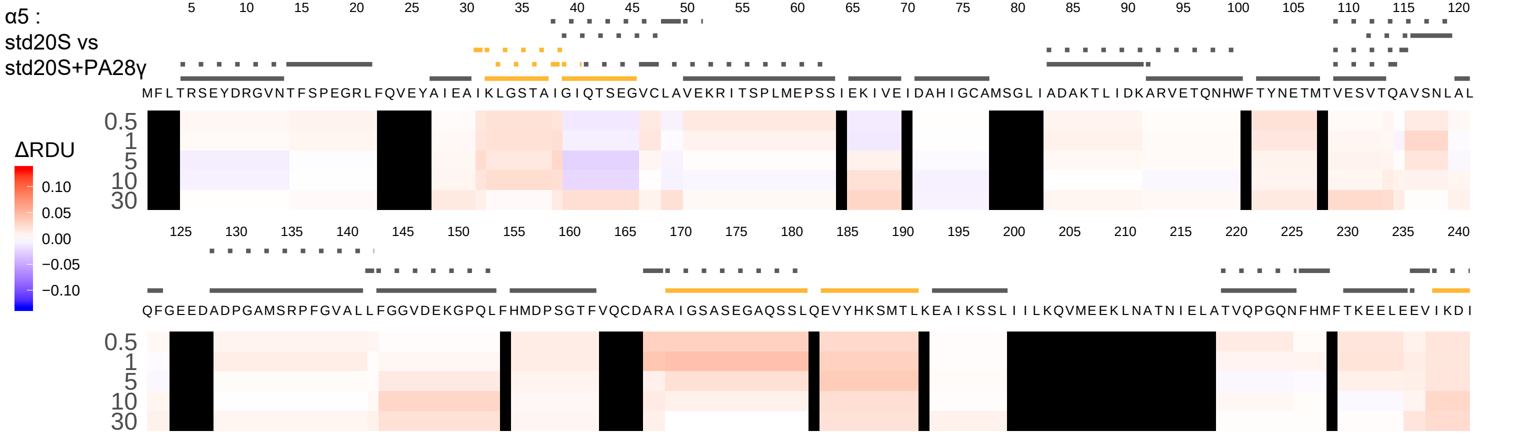
α5 :
std20S vs
std20S+PA28γ
ΔRDU
0.10
0.05
0.00
−0.05
−0.10
0.5
1
5
10
30
5 10 15 20 25 30 35 40 45 50 55 60 65 70 75 80 85 90 95 100 105 110 115 120
M F L T R S E Y D R G V N T F S P E G R L F Q V E Y A I E A I K L G S T A I G I Q T S E G V C L A V E K R I T S P L M E P S S I E K I V E I D A H I G C A M S G L I A D A K T L I D K A R V E T Q N H W F T Y N E T M T V E S V T Q A V S N L A L
125 130 135 140 145 150 155 160 165 170 175 180 185 190 195 200 205 210 215 220 225 230 235 240
Q F G E E D A D P G A M S R P F G V A L L F G G V D E K G P Q L F H M D P S G T F V Q C D A R A I G S A S E G A Q S S L Q E V Y H K S M T L K E A I K S S L I I I L K Q V M E E K L N A T N I E L A T V Q P G Q N F H M F T K E E L E E V I K D I

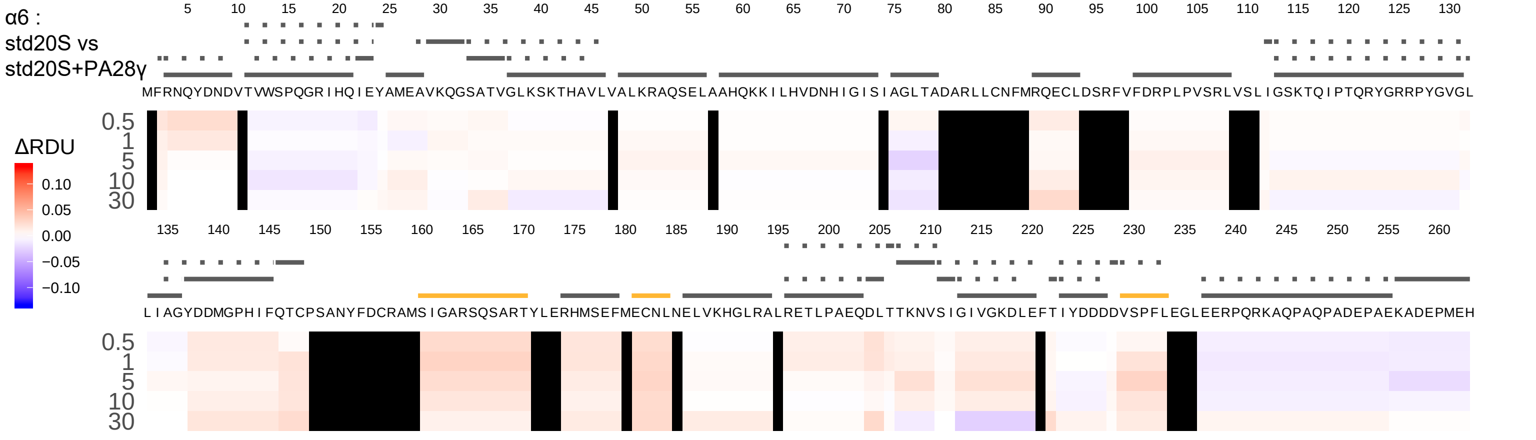

α6 :
std20S vs
std20S+PA28γ
ΔRDU
0.10
0.05
0.00
−0.05
−0.10
0.5
1
5
10
30
5 10 15 20 25 30 35 40 45 50 55 60 65 70 75 80 85 90 95 100 105 110 115 120 125 130
MFRNQYDNDVTVWSPQGRIHQIEYAMEAVKQGSATVGLKSKTHAVLVALKRAQSELAAHQKKILHVDNHIGISIAGLTADARLLCNFMRQECLDSRFVFDRPLPVSRLVSLIGSKTQIPTQRYGRRPYGVGL
135 140 145 150 155 160 165 170 175 180 185 190 195 200 205 210 215 220 225 230 235 240 245 250 255 260
LIAGYDDMGPHIFQTCPSANYFDCRAMSIGARSQSARTYLERHMSEFMECNLNELVKHGLRALRETLPAEQDLTTKNVSIGIVGKDLEFTIYDDDDVSPFLEGLEERPQRKAQPAQPADEPAEKADEPMEH

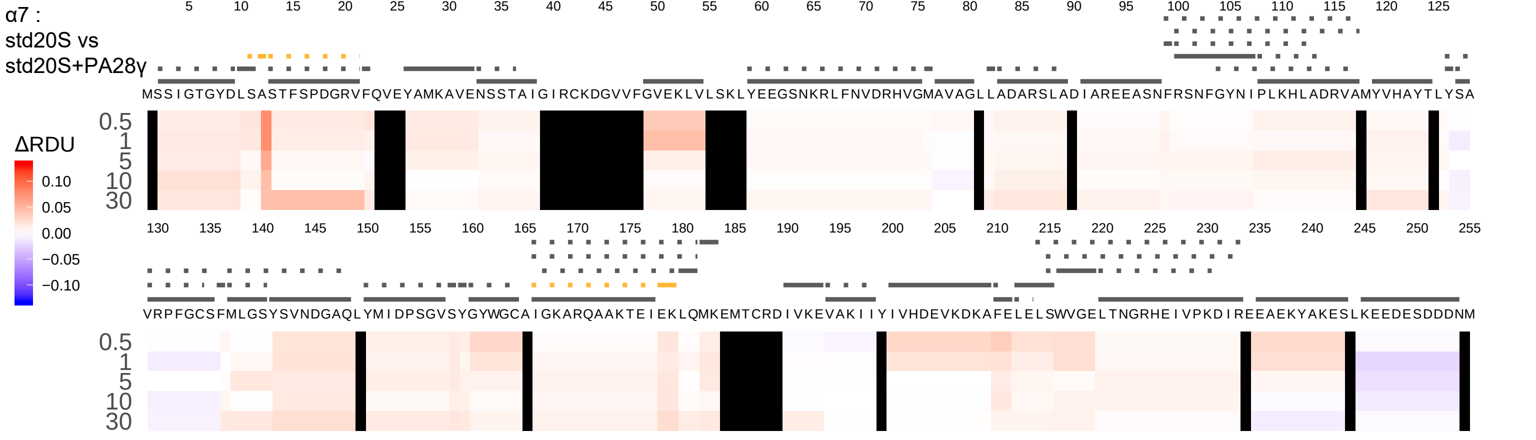
α7 :
std20S vs
std20S+PA28γ
ΔRDU
0.10
0.05
0.00
−0.05
−0.10
0.5
1
5
10
30
MSSIGTGYDLSASTFSPDGRVFQVEYAMKAVENSSTAIGIRCKDGVVFGVEKLVLSKLYEEGSNKRLFNVDRHVGMAVAGLLADARSLADIAREEASNFRSNFGYNIPLKHLADRVAMYVHAYTLYSA
VRPFGCSFMLGSYSVNDGAQLYMIDPSGVSYGYWGCAIGKARQAAKTEIEKLQMKEMTCRDIVKEVAKIIYIVHDEVKDKAFELELSWVGELTNGRHEIVPKDIREEAEKYAKESLKEEDESDDDNM

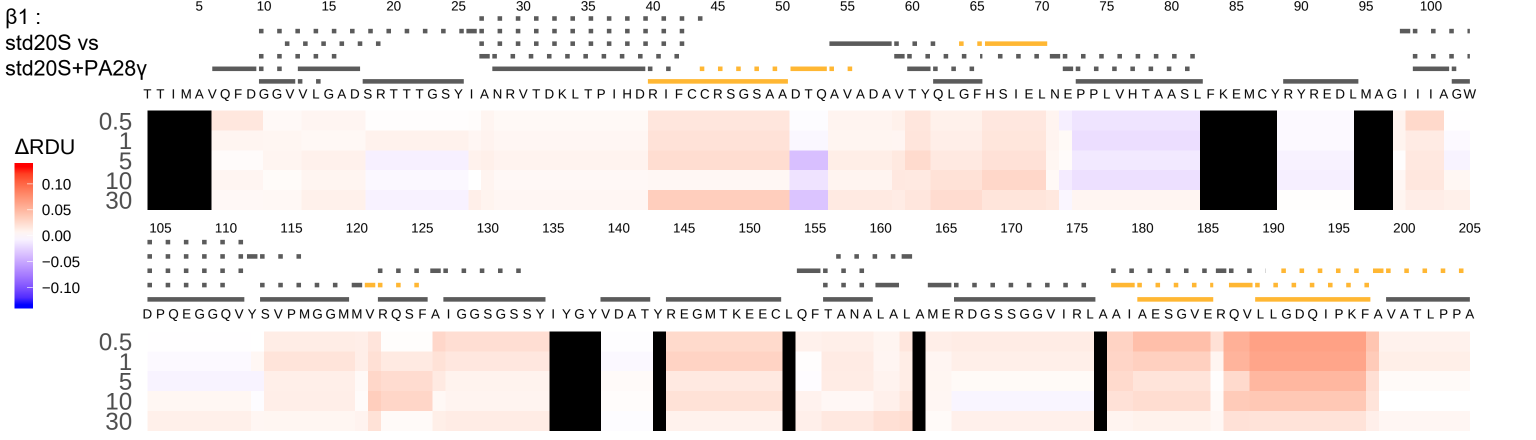
β1 :
std20S vs
std20S+PA28γ
ΔRDU
0.10
0.05
0.00
−0.05
−0.10
5 10 15 20 25 30 35 40 45 50 55 60 65 70 75 80 85 90 95 100
T T I M A V Q F D G G V V L G A D S R T T T G S Y I A N R V T D K L T P I H D R I F C C R S G S A A D T Q A V A D A V T Y Q L G F H S I E L N E P P L V H T A A S L F K E M C Y R Y R E D L M A G I I I A G W
0.5
1
5
10
30
105 110 115 120 125 130 135 140 145 150 155 160 165 170 175 180 185 190 195 200 205
D P Q E G G Q V Y S V P M G G M M V R Q S F A I G G S G S S Y I Y G Y V D A T Y R E G M T K E E C L Q F T A N A L A L A M E R D G S S G G V I R L A A I A E S G V E R Q V L L G D Q I P K F A V A T L P P A
0.5
1
5
10
30

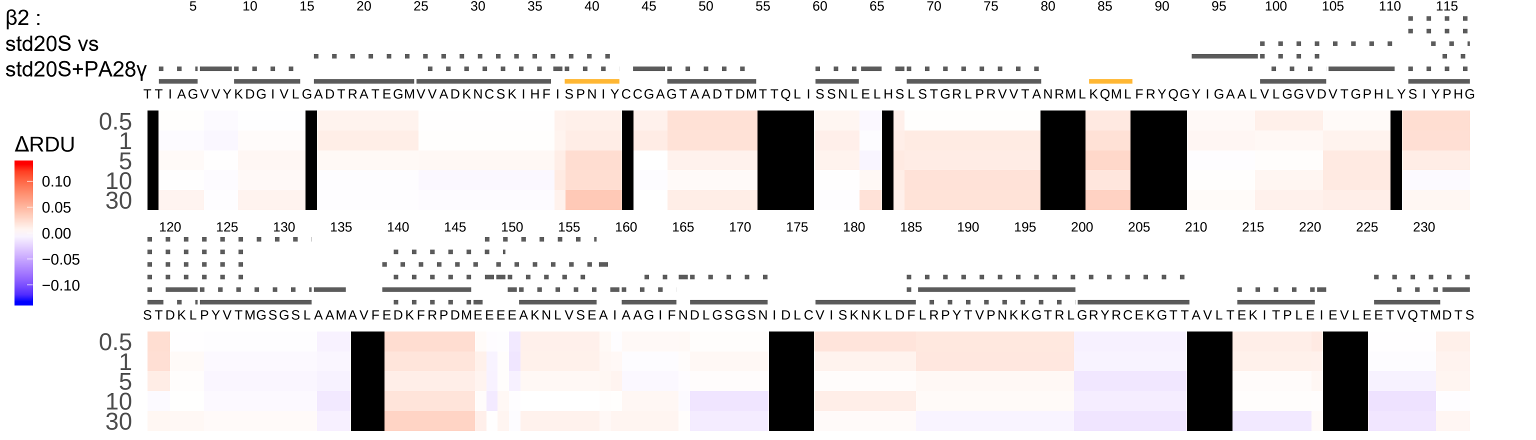

β2 :
std20S vs
std20S+PA28γ
ΔRDU
0.10
0.05
0.00
−0.05
−0.10
0.5
1
5
10
30
TTIAGVVYKDGIVLGADTRATEGMVVADKNCSKIHFISPNIYCCGAGTAADTDMTTQLISSNLELHSLSTGRLPRVVTANRMLKQMLFRYQGYIGAALVLGGVDVTGPHLYSIYPHG
STDKLPYVTMGSGSLAAMAVFEDKFRPDMEEEEAKNLVSEAIAAGIFNDLGSGSNIDLCVISKNKLDFLRPYTVPNKKGTRLGRYRCEKGTTAVLTEKITPLEIEVLEETVQTMDTS

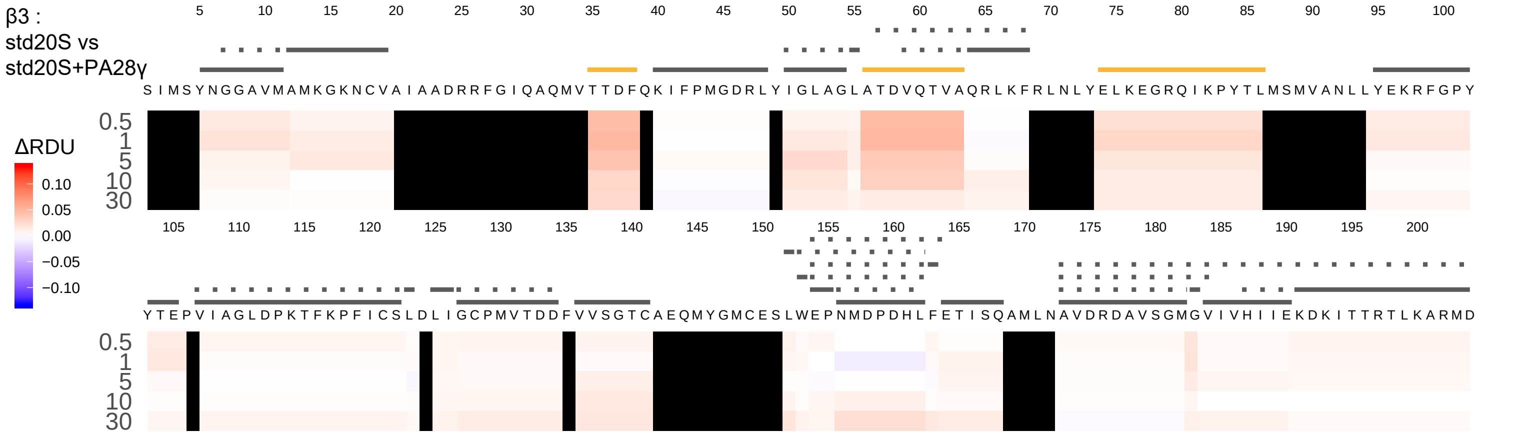
β3 :
std20S vs
std20S+PA28γ
ΔRDU
0.10
0.05
0.00
−0.05
−0.10
0.5
1
5
10
30
SIMSYNGGAVMAMKGKNCVAIAADRRFGIQAQMVTTDFQKIFPMGDRLYIGLAGLATDVQTVAQRLKFRLNLYELKEGRQIKPYTLMSMVANLLYEKRFGPY
YTEPVIAGLDPKTFKPFICSLDLIGCPMVTDDFVVSGTCAEQMYGMCESLWEPNMDPDHLFETISQAMLNAVDRDAVSGMGVIVHIIEKDKITTRTLKARMD

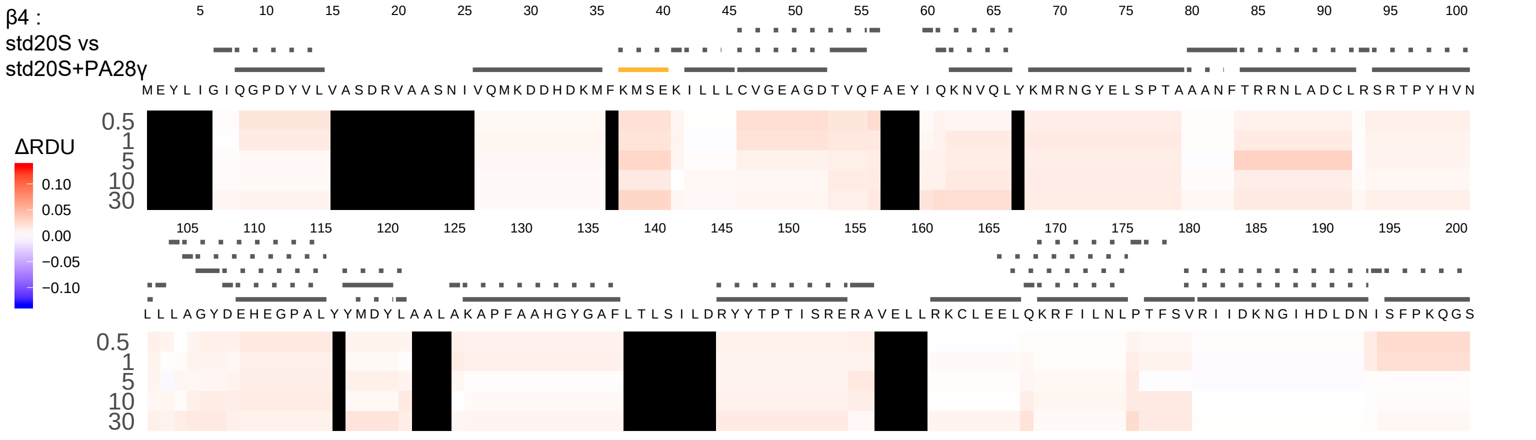

β4 :
std20S vs
std20S+PA28γ
ΔRDU
0.10
0.05
0.00
−0.05
−0.10
0.5
1
5
10
30
5 10 15 20 25 30 35 40 45 50 55 60 65 70 75 80 85 90 95 100
MEYLIGIQGPDYVLVASDRVAASNIVQMKDDHDKMFKMSEKILLLCVGEAGDTVQFAEYIQKNVQLYKMRNGYELSPTAAANFTRRNLADCLRSRTPYHVN
105 110 115 120 125 130 135 140 145 150 155 160 165 170 175 180 185 190 195 200
LLLAGYDEHEGPALYYMDYLAALAKAPFAAHGYGAFLTLSILDRYYTPTISRERAVELLRKCLEELQKRFILNLPTFSVRIIDKNGIHDLDNISFPKQGS

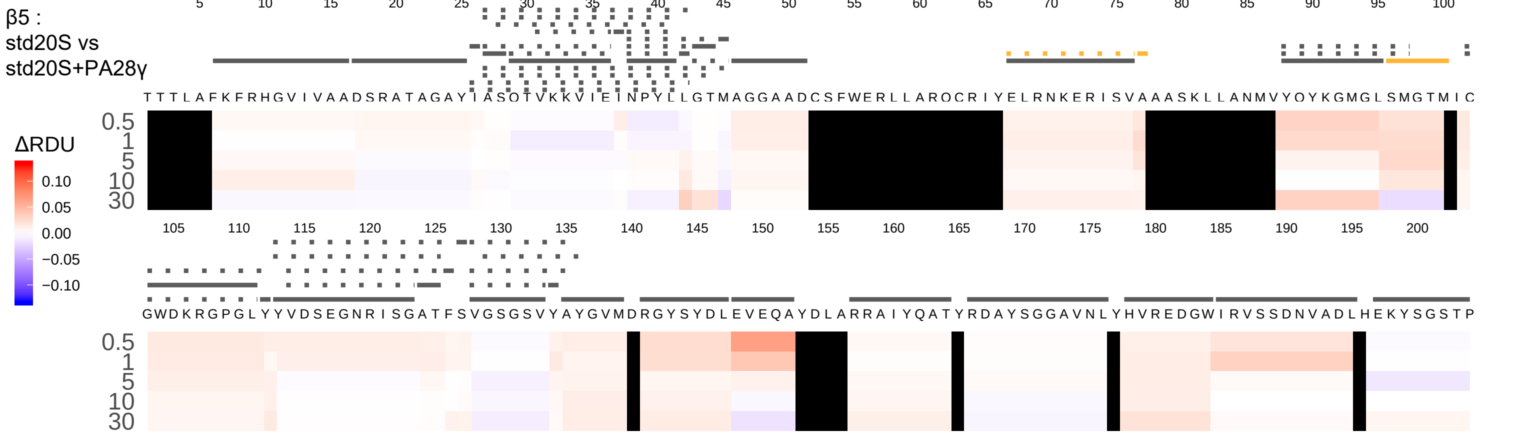
β5 :
std20S vs
std20S+PA28γ
ΔRDU
0.10
0.05
0.00
−0.05
−0.10
0.5
1
5
10
30
TTTLAFKFRHGVIVAADSRATAGAYIASQTVKKVIEINPYLLGTMAGGAADCSFWERLLARQCRIYELRNKERISVAAASKLLANMVYQYKGMGLSMGTMIC
GWDKRGPGLYYVDSEGNRISGATFSVGSGSVYAYGVMDRGYSYDLEVEQAYDLARRAIYQATYRDAYSGGAVNLYHVREDGWIRVSSDNVADLHEKYSGSTP

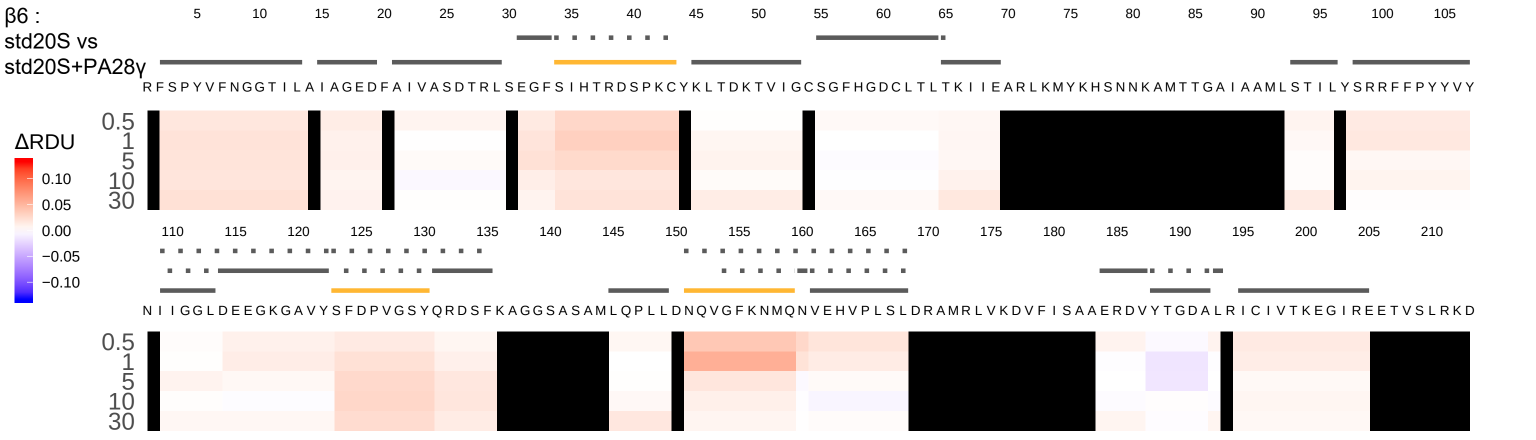

β6 :
std20S vs
std20S+PA28γ
ΔRDU
0.10
0.05
0.00
−0.05
−0.10
0.5
1
5
10
30
RFSPYVFNGGTILAIAGEDFAIVASDTRLSEGFSIHTRDSPKCYKLTDKTVIGCSGFHGDCLTLTKIIEARLKMYKHSNNKAMTTGAIAAMLSTILYSRRFFPYYVY
NIIGGLDEEGKGAVYSFDPVGSYQRDSFKAGGSASAMLQPLLDNQVGFKNMQNVEHVPLSLDRAMRLVKDVFISAAERDVYTGDALRICIVTKEGIREETVSLRKD

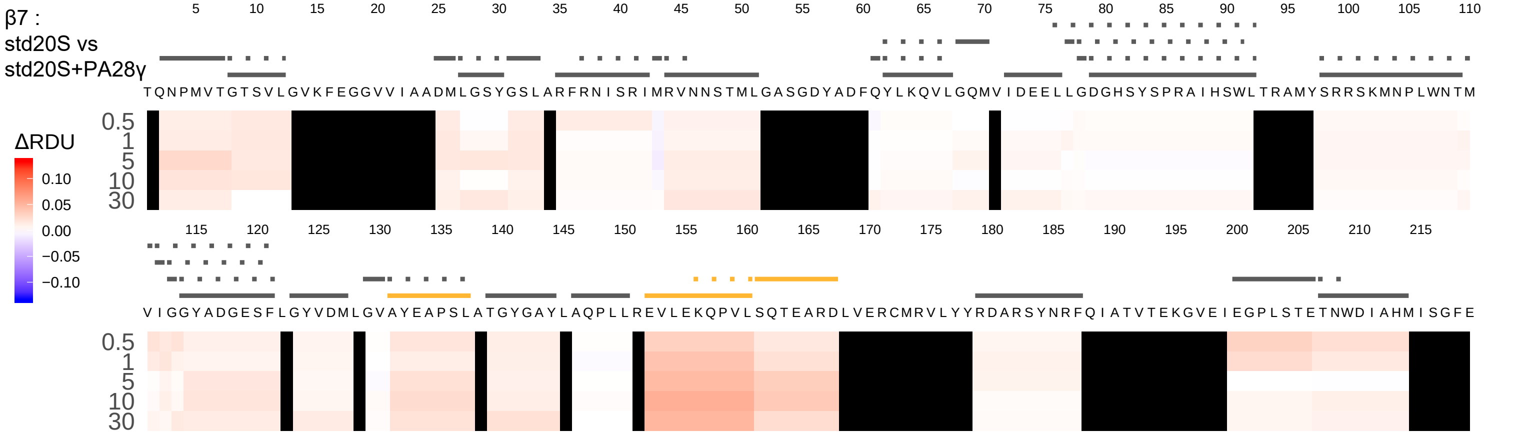
β7 :
std20S vs
std20S+PA28γ
5 10 15 20 25 30 35 40 45 50 55 60 65 70 75 80 85 90 95 100 105 110
T Q N P M V T G T S V L G V K F E G G V V I A A D M L G S Y G S L A R F R N I S R I M R V N N S T M L G A S G D Y A D F Q Y L K Q V L G Q M V I D E E L L G D G H S Y S P R A I H S W L T R A M Y S R R S K M N P L W N T M
0.5
1
5
10
30
ΔRDU
0.10
0.05
0.00
−0.05
−0.10
115 120 125 130 135 140 145 150 155 160 165 170 175 180 185 190 195 200 205 210 215
V I G G Y A D G E S F L G Y V D M L G V A Y E A P S L A T G Y G A Y L A Q P L L R E V L E K Q P V L S Q T E A R D L V E R C M R V L Y Y R D A R S Y N R F Q I A T V T E K G V E I E G P L S T E T N W D I A H M I S G F E
0.5
1
5
10
30

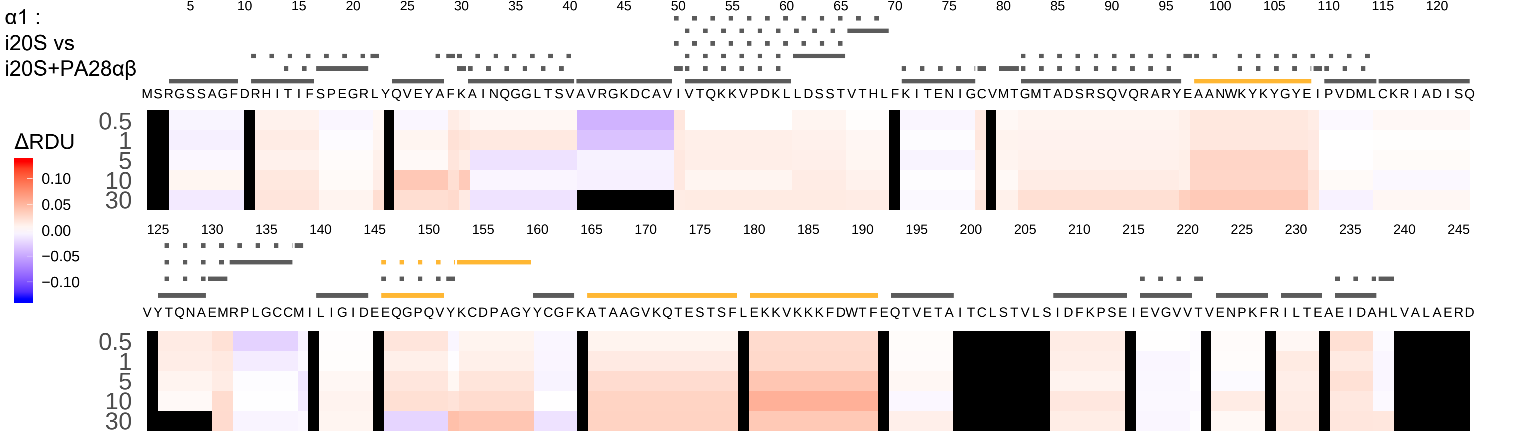
α1 :
i20S vs
i20S+PA28αβ
ΔRDU
0.10
0.05
0.00
−0.05
−0.10
0.5
1
5
10
30
MSRGSSAGFDRHITIFSPEGRLYQVEYAFKAINQGGLTSVAVRGKDCAVIVTQKKVPDKLLDSSTVTHLFKITENIGCVMTGMTADSRSQVQRARYEAANWKYKYGYEIPVDMLCKRIADISQ
VYTQNAEMRPLGCCMILIGIDEEQGPQVYKCDPAGYYCGFKATAAGVKQTESTSFLEKKVKKKFDWTFEQTVETAITCLSTVLSIDFKPSEIEVGVVTVENPKFRILTEAEIDAHLVALAERD

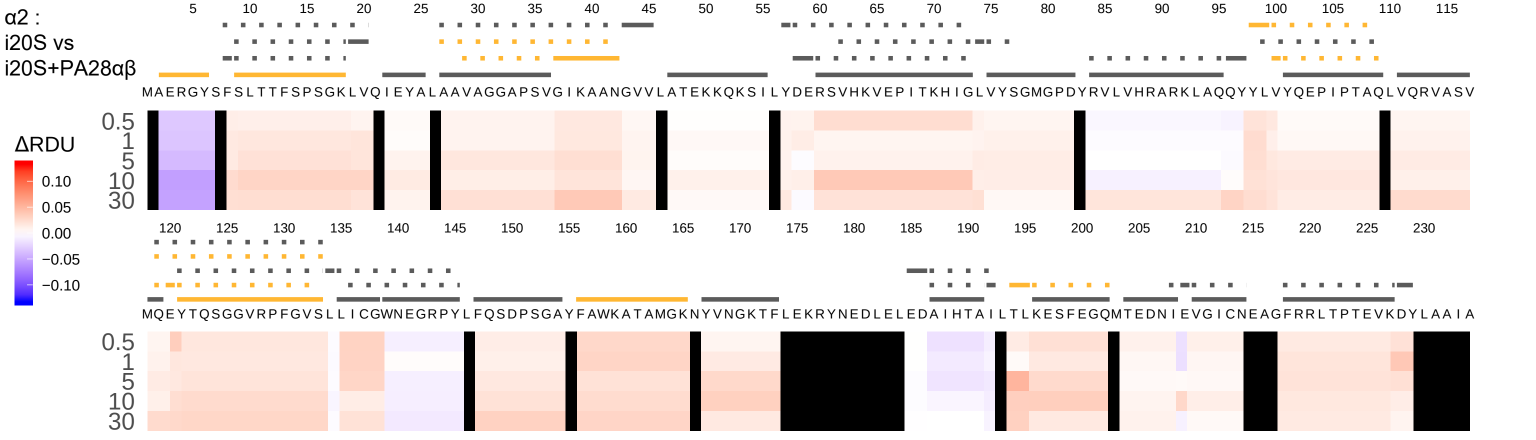

α2 :
i20S vs
i20S+PA28αβ
ΔRDU
0.10
0.05
0.00
−0.05
−0.10
0.5
1
5
10
30
5 10 15 20 25 30 35 40 45 50 55 60 65 70 75 80 85 90 95 100 105 110 115
MAERGYSFSLTTFSPSGKLVQIEYALAAVAGGAPSVGIKAANGVVLATEKKQKSILYDERSVHKVEPITKHIGLVYSGMGPDYRVLVHRARKLAQQYYLVYQEPIPTAQLVQRVASV
120 125 130 135 140 145 150 155 160 165 170 175 180 185 190 195 200 205 210 215 220 225 230
MQEYTQSGGVRPFGVSLLICGWNEGRPYLFQSDPSGAYFAWKATAMGKNYVNGKTFLEKRYNEDLELEDAIHTAILTLKESFEGQMTEDNIEVGICNEAGFRRLTPTEVKDYLAAIA

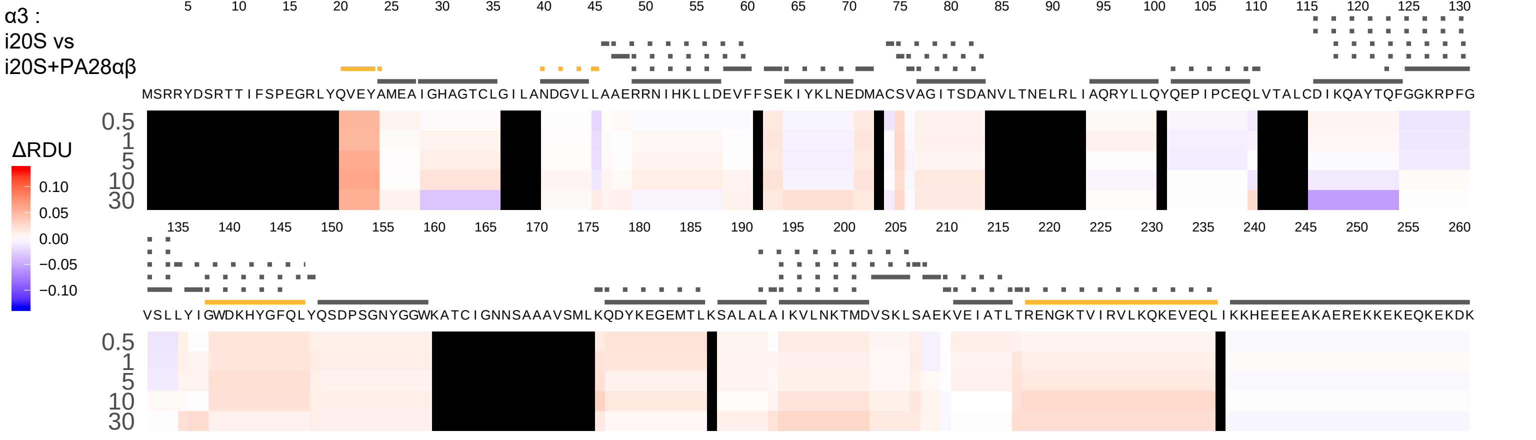

α3 :
i20S vs
i20S+PA28αβ
ΔRDU
0.10
0.05
0.00
−0.05
−0.10
0.5
1
5
10
30
5 10 15 20 25 30 35 40 45 50 55 60 65 70 75 80 85 90 95 100 105 110 115 120 125 130
MSRRYDSRTTIFSPEGRLYQVEYAMEAIGHAGTCLGILANDGVLLAAERRNIHKLLDEVFFSEKIYKLNEDMACSVAGITSDANVLTNELRLIAQRYLLQYQEPIPCEQLVTALCDIKQAYTQFGGKRPFG
135 140 145 150 155 160 165 170 175 180 185 190 195 200 205 210 215 220 225 230 235 240 245 250 255 260
VSLLYIGWDKHYGFQLYQSDPSGNYGGWKATCIGNNSAAAVSMLKQDYKEGEMTLKSALALAIKVLNKTMDVSKLSAEKVEIATLTRENGKTVIRVLKQKEVEQLIKKHEEEEAKAEREKKEKEQKEKDK

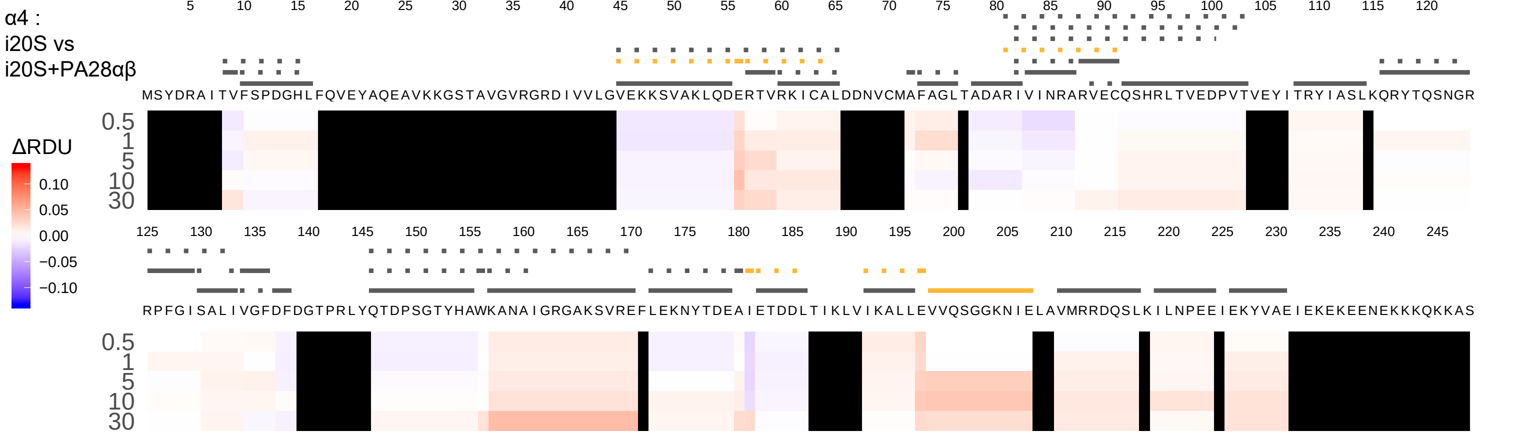

α4 :
i20S vs
i20S+PA28αβ
ΔRDU
0.10
0.05
0.00
−0.05
−0.10
0.5
1
5
10
30
MSYDRAITVFSPDGHLFQVEYAQEAVKKGSTAVGVRGRDIVVLGVEKKSVAKLQDERTVRKICALDDNVCMAFAGLTADARIVINRARVECQSHRLTVEDPVTVEYITRYIASLKQRYTQSNGR
RPFGISALIVGFDFDGTPRLYQTDPSGTYHAWKANAIGRGAKSVREFLEKNYTDEAIETDDLTIKLVIKALLEVVQSGGKNIELAVMRRDQSLKILNPEEIEKYVAEIEKEKEENEKKKQKKAS

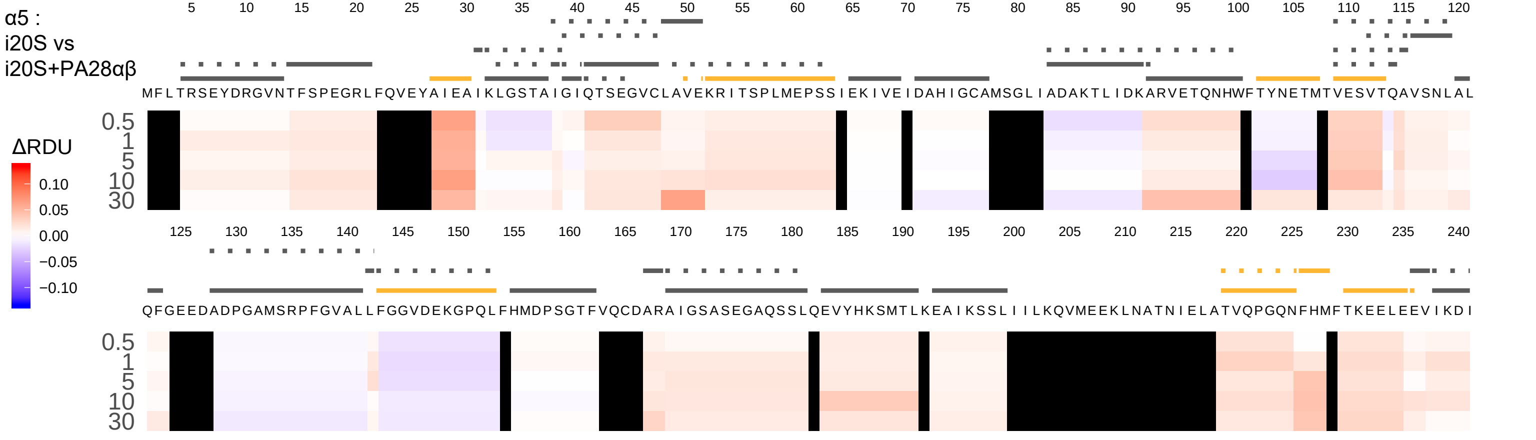

α5 :
i20S vs
i20S+PA28αβ
ΔRDU
0.10
0.05
0.00
−0.05
−0.10
0.5
1
5
10
30
MFLTRSEYDRGVNTFSPEGRLFQVEYAIEAIKLGSTAIGIQTSEGVCLAVEKRITSPLMEPSSIEKIVEIDAHIGCAMSGLIADAKTLIDKARVETQNHWFTYNETMTVESVTQAVSNLAL
QFGEEDADPGAMSRPFGVALLFGGVDEKGPQLFHMDPSGTFVQCDARAIGSASEGAQSSLQEVYHKSMTLKEAIKSSLIIILKQVMEEKLNATNIELATVQPGQNFHMFTKEELEEVIKDI

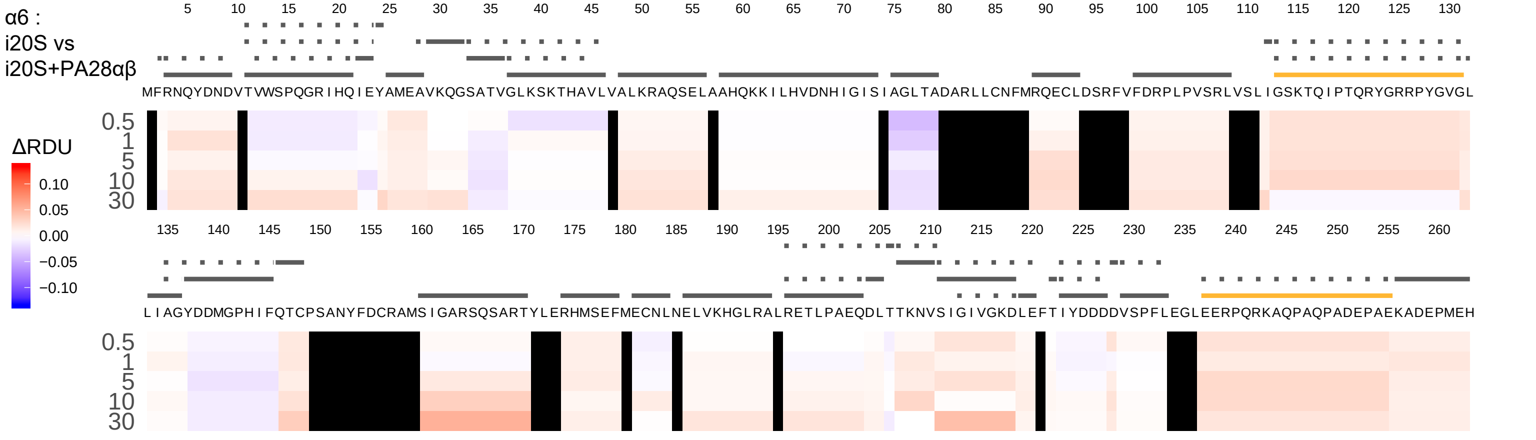

α6 :
i20S vs
i20S+PA28αβ
ΔRDU
0.10
0.05
0.00
−0.05
−0.10
5 10 15 20 25 30 35 40 45 50 55 60 65 70 75 80 85 90 95 100 105 110 115 120 125 130
MFRNQYDNDVTVWSPQGRIHQIEYAMEAVKQGSATVGLKSKTHAVLVALKRAQSELAAHQKKILHVDNHIGISIAGLTADARLLCNFMRQECLDSRFVFDRPLPVSRLVSLIGSKTQIPTQRYGRRPYGVGL
0.5
1
5
10
30
135 140 145 150 155 160 165 170 175 180 185 190 195 200 205 210 215 220 225 230 235 240 245 250 255 260
LIAGYDDMGPHIFQTCPSANYFDCRAMSIGARSQSARTYLERHMSEFMECNLNELVKHGLRALRETLPAEQDLTTKNVSIGIVGKDLEFTIYDDDDVSPFLEGLEERPQRKAQPAQPADEPAEKADEPMEH
0.5
1
5
10
30

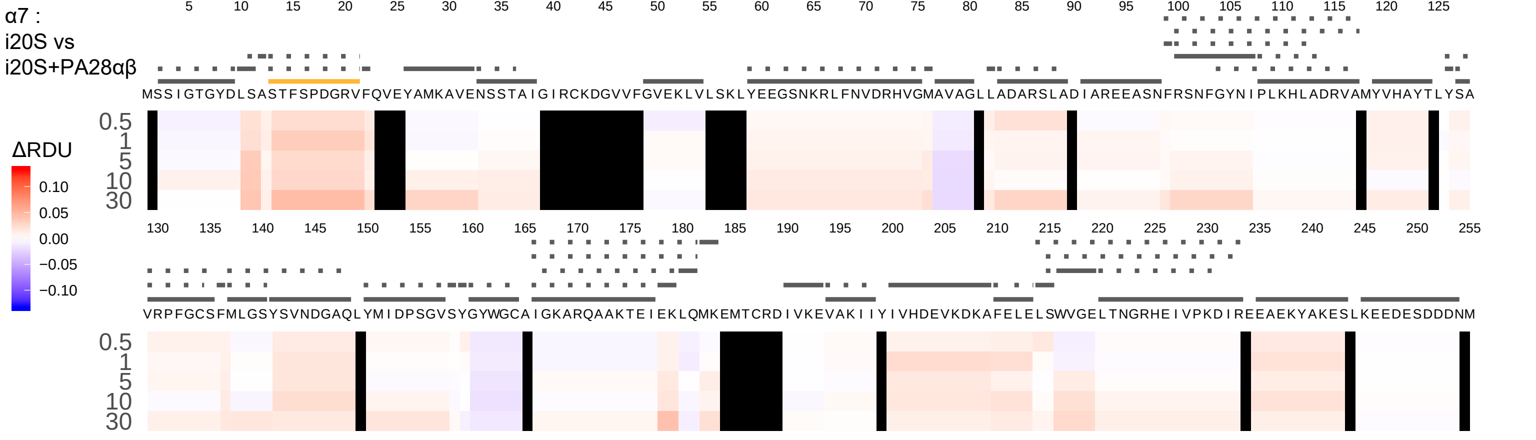

α7 :
i20S vs
i20S+PA28αβ
MSSIGTGYDLSASTFSPDGRVFQVEYAMKAVENSSTAIGIRCKDGVVFGVEKLVLSKLYEEGSNKRLFNVDRHVGMAVAGLLADARSLADIAREEASNFRSNFGYNIPLKHLADRVAMYVHAYTLYSA
VRPFGCSFMLGSYSVNDGAQLYMIDPSGVSYGYWGCAIGKARQAAKTEIEKLQMKEMTCRDIVKEVAKIIYIVHDEVKDKAFELELSWVGELTNGRHEIVPKDIREEAEKYAKESLKEEDESDDDNM
0.5
1
5
10
30
ΔRDU
0.10
0.05
0.00
−0.05
−0.10

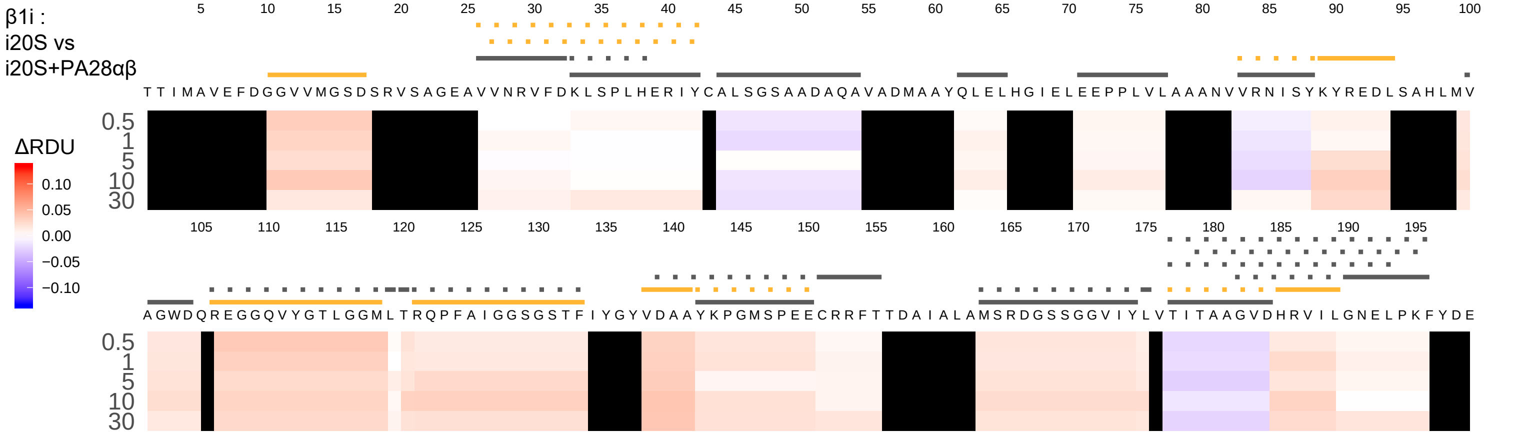
β1i :
i20S vs
i20S+PA28αβ
ΔRDU
0.10
0.05
0.00
−0.05
−0.10
0.5
1
5
10
30
T T I M A V E F D G G V V M G S D S R V S A G E A V V N R V F D K L S P L H E R I Y C A L S G S A A D A Q A V A D M A A Y Q L E L H G I E L E E P P L V L A A A N V V R N I S Y K Y R E D L S A H L M V
A G W D Q R E G G Q V Y G T L G G M L T R Q P F A I G G S G S T F I Y G Y V D A A Y K P G M S P E E C R R F T T D A I A L A M S R D G S S G G V I Y L V T I T A A G V D H R V I L G N E L P K F Y D E

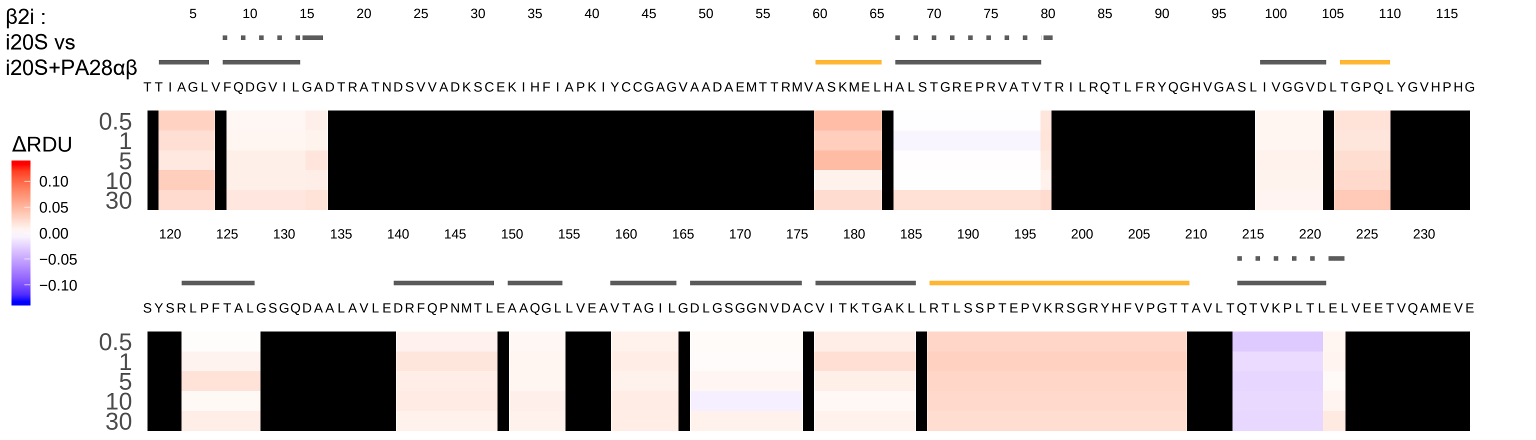
β2i :
i20S vs
i20S+PA28αβ
ΔRDU
0.10
0.05
0.00
−0.05
−0.10
0.5
1
5
10
30
TTIAGLVFQDGVILGADTRATNDSVVADKSCEKIHFIAPKIYCCGAGVAADAEMTTRMVASKMELHALSTGREPRVATVTRILRQTLFRYQGHVGASLIVGGVDLTGPQLYGVHPHG
SYSRLPFTALGSGQDAALAVLEDRFQPNMTLEAAQGLLVEAVTAGILGDLGSGGNVDACVITKTGAKLLRTLSSPTEPVKRSGRYHFVPGTTAVLTQTVKPLTLELVEETVQAMEVE

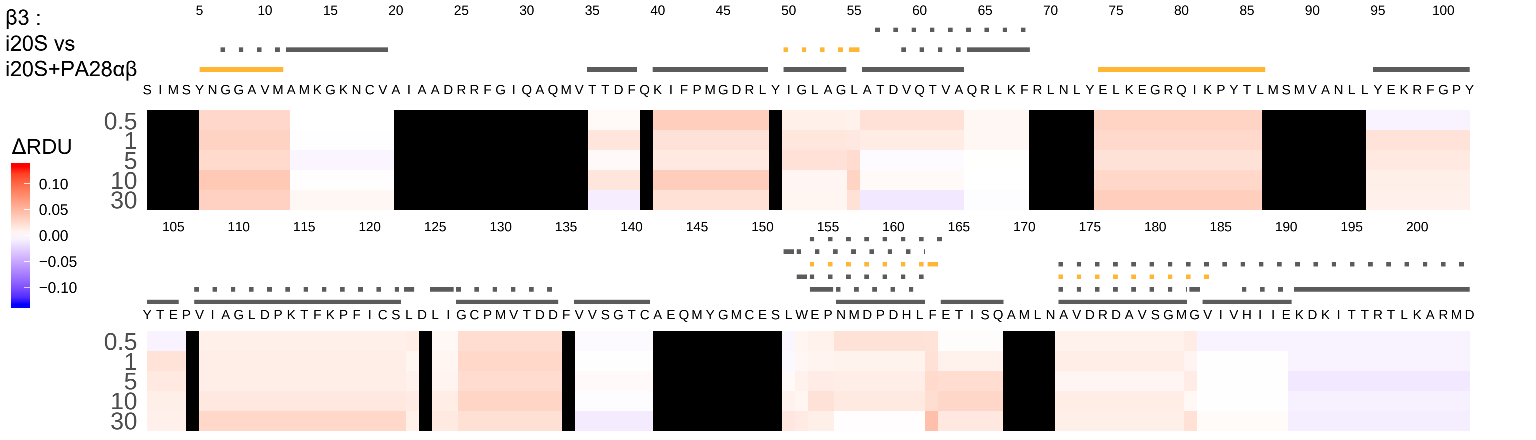
β3 :
i20S vs
i20S+PA28αβ
ΔRDU
0.10
0.05
0.00
−0.05
−0.10
0.5
1
5
10
30
S I M S Y N G G A V M A M K G K N C V A I A A D R R F G I Q A Q M V T T D F Q K I F P M G D R L Y I G L A G L A T D V Q T V A Q R L K F R L N L Y E L K E G R Q I K P Y T L M S M V A N L L Y E K R F G P Y
Y T E P V I A G L D P K T F K P F I C S L D L I G C P M V T D D F V V S G T C A E Q M Y G M C E S L W E P N M D P D H L F E T I S Q A M L N A V D R D A V S G M G V I V H I I E K D K I T T R T L K A R M D

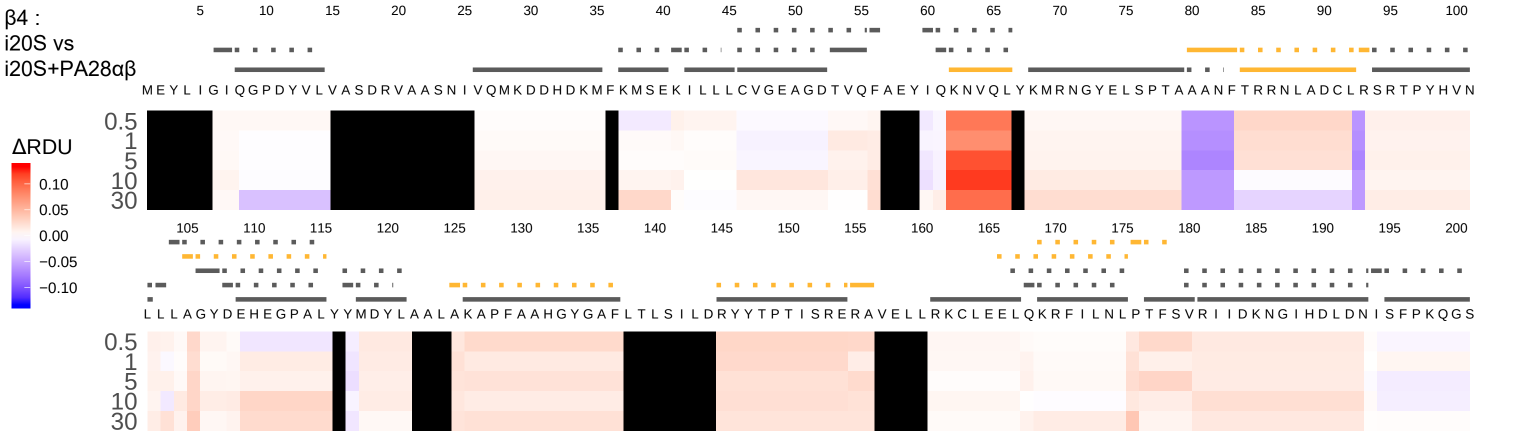
β4 :
i20S vs
i20S+PA28αβ
ΔRDU
0.10
0.05
0.00
−0.05
−0.10
0.5
1
5
10
30
MEYLIGIQGPDYVLVASDRVAASNIVQMKDDHDKMFKMSEKILLLCVGEAGDTVQFAEYIQKNVQLYKMRNGYELSPTAAANFTRRNLADCLRSRTPYHVN
LLLAGYDEHEGPALYYMDYLAALAKAPFAAHGYGAFLTLSILDRYYTPTISRERAVELLRKCLEELQKRFILNLPTFSVRIIDKNGIHDLDNISFPKQGS

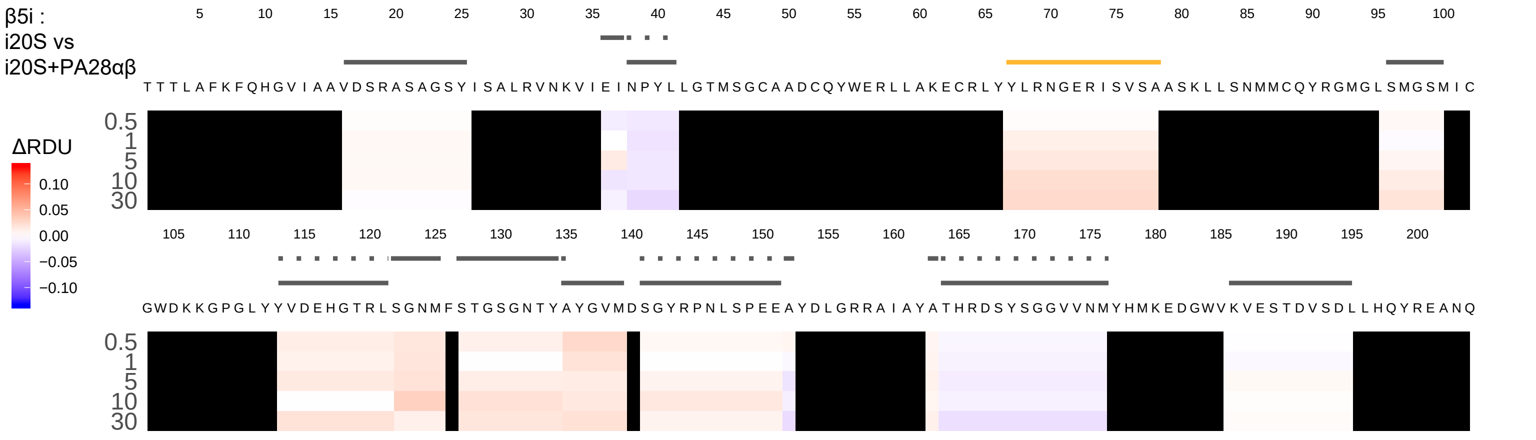

β5i :
i20S vs
i20S+PA28αβ
ΔRDU
0.10
0.05
0.00
−0.05
−0.10
0.5
1
5
10
30
TTTLAFKFQHGVIAAVDSRASAGSYISALRVNKVIEINPYLLGTMSGCAADCQYWERLLAKECRLYYLRNGERISVSAASKLLSNMMCQYRGMGLSMGSMIC
GWDKKGPGLYYVDEHGTRLSGNMFSTGSGNTYAYGVMDSGYRPNLSPEEAYDLGRRAIAYATHRDSYSGGVVNMYHMKEDGWVKVESTDVSDLLHQYREANQ

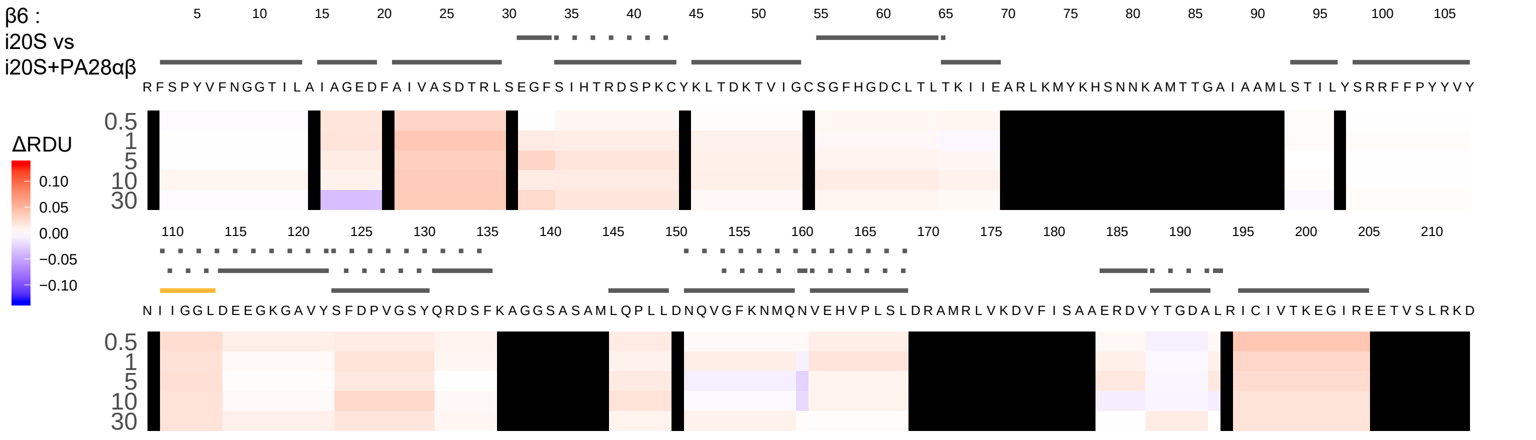
β6 :
i20S vs
i20S+PA28αβ
5 10 15 20 25 30 35 40 45 50 55 60 65 70 75 80 85 90 95 100 105
R F S P Y V F N G G T I L A I A G E D F A I V A S D T R L S E G F S I H T R D S P K C Y K L T D K T V I G C S G F H G D C L T L T K I I E A R L K M Y K H S N N K A M T T G A I A A M L S T I L Y S R R F F P Y Y V Y
0.5
1
5
10
30
ΔRDU
0.10
0.05
0.00
−0.05
−0.10
110 115 120 125 130 135 140 145 150 155 160 165 170 175 180 185 190 195 200 205 210
N I I G G L D E E G K G A V Y S F D P V G S Y Q R D S F K A G G S A S A M L Q P L L D N Q V G F K N M Q N V E H V P L S L D R A M R L V K D V F I S A A E R D V Y T G D A L R I C I V T K E G I R E E T V S L R K D
0.5
1
5
10
30

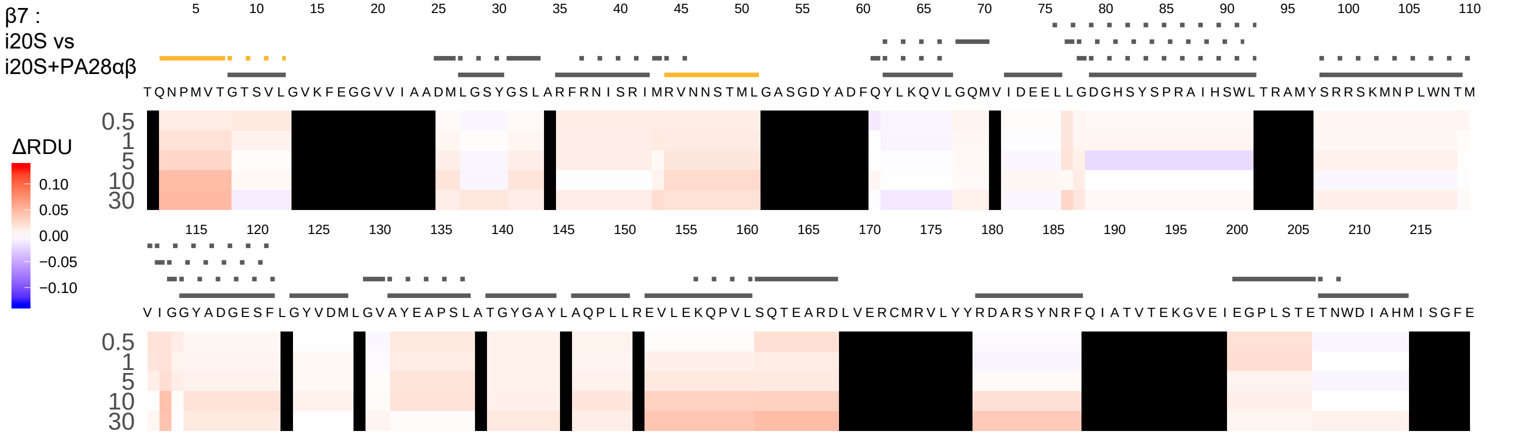
β7 :
i20S vs
i20S+PA28αβ
TQNPMVTGTSVLGVKFEGGVVIAADMLGSYGSLARFRNISRIMRVNNSTMLGASGDYADFQYLKQVLGQMVIDEELLGDGHSYSPRAIHSWLTRAMYSRRSKMNPLWNTM
VIGGYADGESFLGYVDMLGVAYEAPSLATGYGAYLAQPLLREVLEKQPVLSQTEARDLVERCMRVLYYRDARSYNRFQIATVTEKGVEIEGPLSTETNWDIAHMISGFE
ΔRDU
0.10
0.05
0.00
−0.05
−0.10
0.5
1
5
10
30

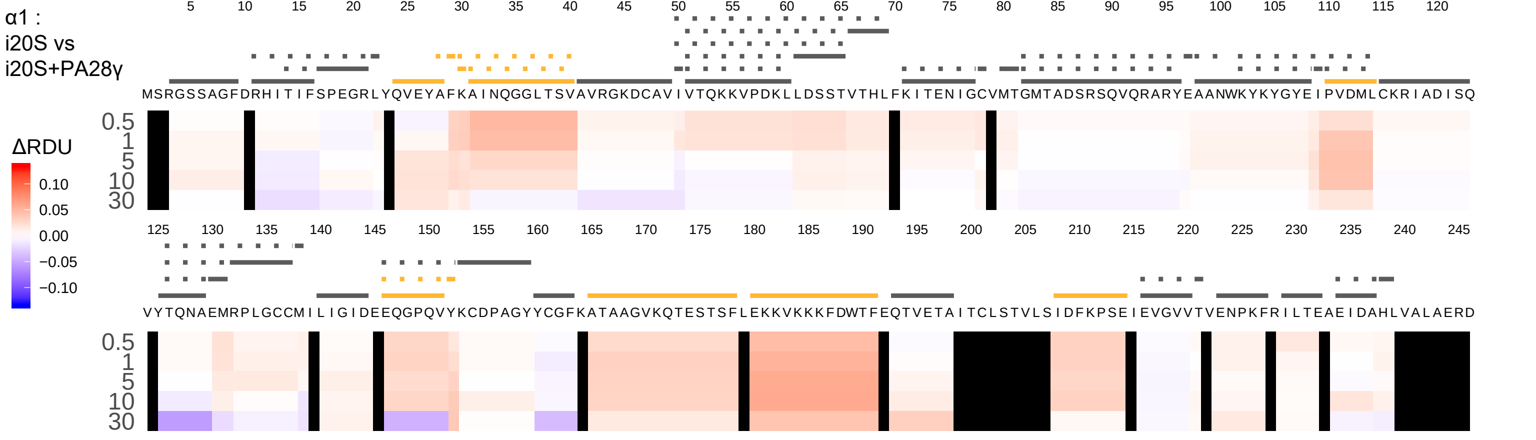

α1 :
i20S vs
i20S+PA28γ
ΔRDU
0.10
0.05
0.00
−0.05
−0.10
0.5
1
5
10
30
MSRGSSAGFDRHITIFSPEGRLYQVEYAFKAINQGGLTSVAVRGKDCAVIVTQKKVPDKLLDSSTVTHLFKITENIGCVMTGMTADSRSQVQRARYEAANWKYKYGYEIPVDMLCKRIADISQ
VYTQNAEMRPLGCCMILIGIDEEQGPQVYKCDPAGYYCGFKATAAGVKQTESTSFLEKKVKKKFDWTFEQTVETAITCLSTVLSIDFKPSEIEVGVVTVENPKFRILTEAEIDAHLVALAERD

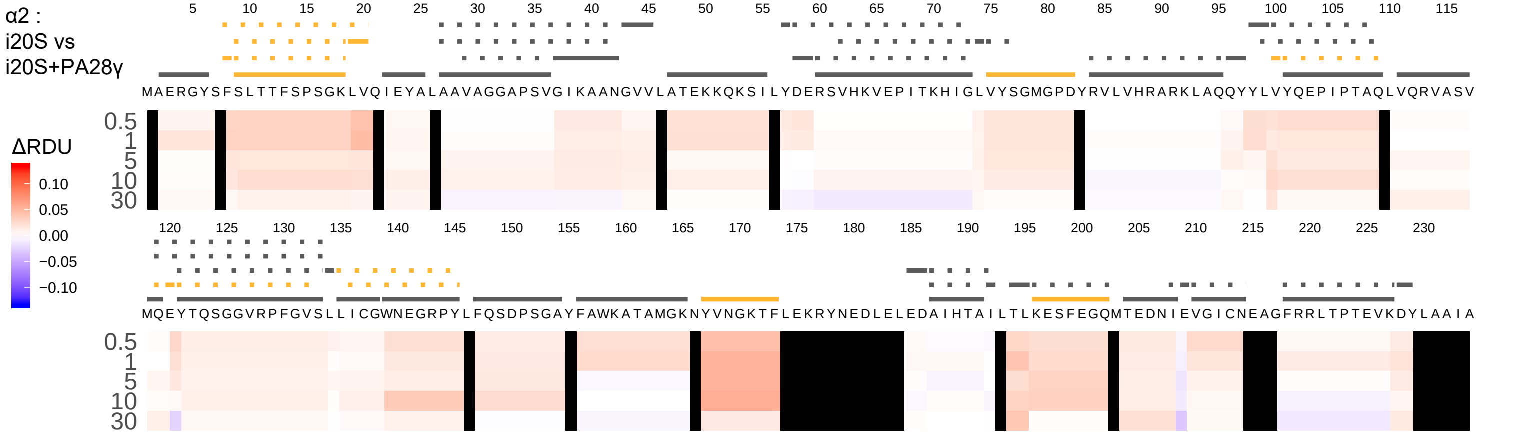
α2 :
i20S vs
i20S+PA28γ
ΔRDU
0.10
0.05
0.00
−0.05
−0.10
5 10 15 20 25 30 35 40 45 50 55 60 65 70 75 80 85 90 95 100 105 110 115
MAERGYSFSLTTFSPSGKLVQIEYALAAVAGGAPSVGIKAANGVVLATEKKQKSILYDERSVHKVEPITKHIGLVYSGMGPDYRVLVHRARKLAQQYYLVYQEPIPTAQLVQRVASV
0.5
1
5
10
30
120 125 130 135 140 145 150 155 160 165 170 175 180 185 190 195 200 205 210 215 220 225 230
MQEYTQSGGVRPFGVSLLICGWNEGRPYLFQSDPSGAYFAWKATAMGKNYVNGKTFLEKRYNEDLELEDAIHTAILTLKESFEGQMTEDNIEVGICNEAGFRRLTPTEVKDYLAAIA
0.5
1
5
10
30

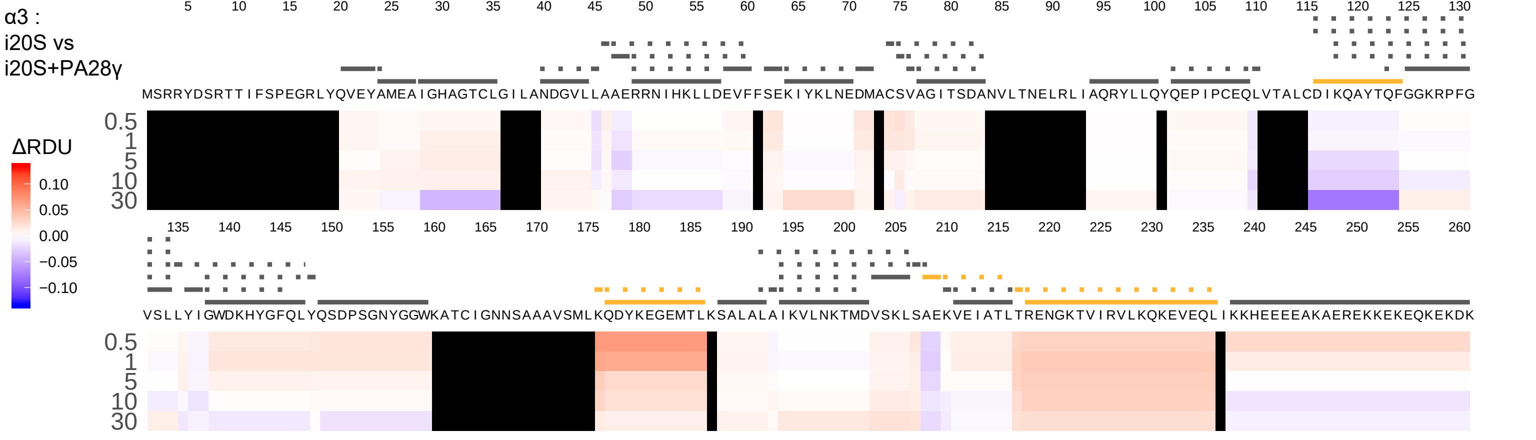
α3 :
i20S vs
i20S+PA28γ
ΔRDU
0.10
0.05
0.00
−0.05
−0.10
0.5
1
5
10
30
MSRRYDSRTTIFSPEGRLYQVEYAMEAIGHAGTCLGILANDGVLLAAERRNIHKLLDEVFFSEKIYKLNEDMACSVAGITSDANVLTNELRLIAQRYLLQYQEPIPCEQLVTALCDIKQAYTQFGGKRPFG
VSLLYIGWDKHYGFQLYQSDPSGNYGGWKATCIGNNSAAAVSMLKQDYKEGEMTLKSALALAIKVLNKTMDVSKLSAEKVEIATLTRENGKTVIRVLKQKEVEQLIKKHEEEEAKAEREKKEKEQKEKDK

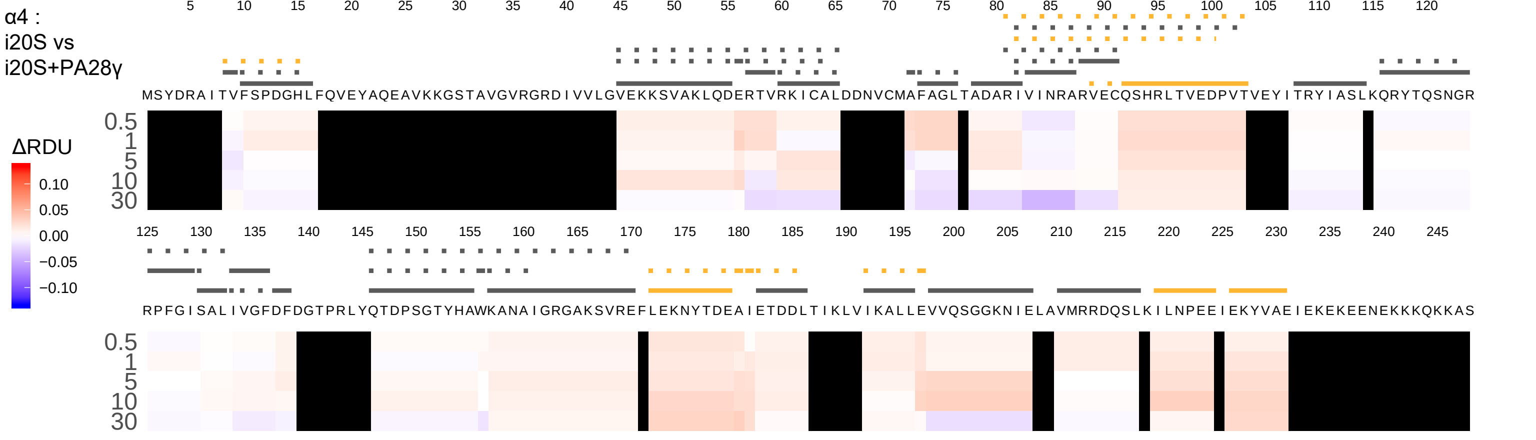
α4 :
i20S vs
i20S+PA28γ
ΔRDU
0.10
0.05
0.00
−0.05
−0.10
0.5
1
5
10
30
MSYDRAITVFSPDGHLFQVEYAQEAVKKGSTAVGVRGRDIVVLGVEKKSVAKLQDERTVRKICALDDNVCMAFAGLTADARIVINRARVECQSHRLTVEDPVTVEYITRYIASLKQRYTQSNGR
RPFGISALIVGFDFDGTPRLYQTDPSGTYHAWKANAIGRGAKSVREFLEKNYTDEAIETDDLTIKLVIKALLEVVQSGGKNIELAVMRRDQSLKILNPEEIEKYVAEIEKEKEENEKKKQKKAS

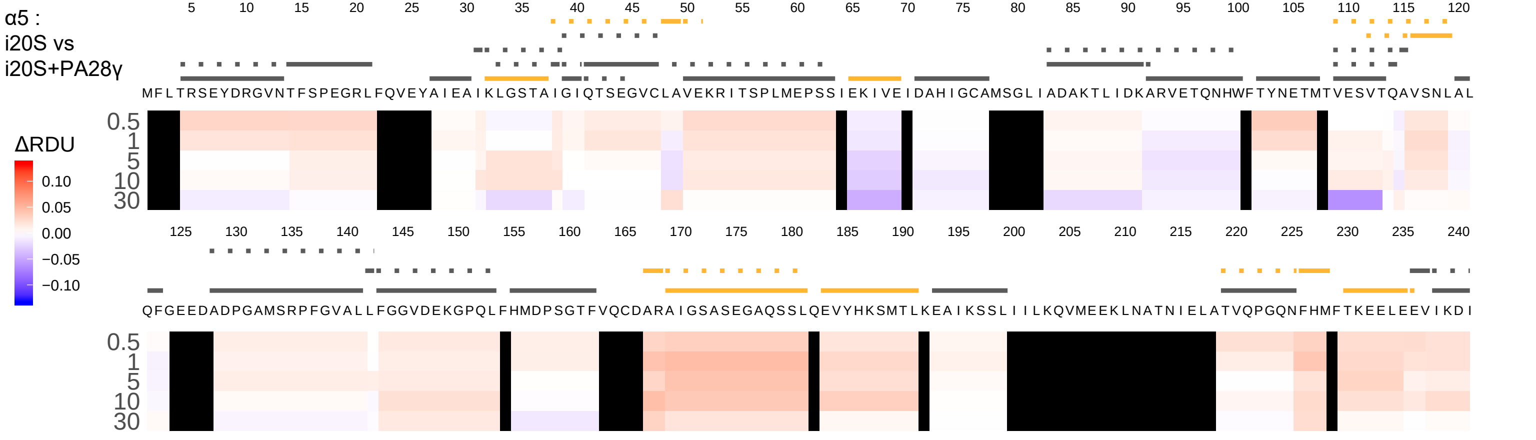

α5 :
i20S vs
i20S+PA28γ
MFLTRSEYDRGVNTFSPEGRLFQVEYAIEAIKLGSTAIGIQTSEGVCLAVEKRITSPLMEPSSIEKIVEIDAHIGCAMSGLIADAKTLIDKARVETQNHWFTYNETMTVESVTQAVSNLAL
QFGEEDADPGAMSRPFGVALLFGGVDEKGPQLFHMDPSGTFVQCDARAIGSASEGAQSSLQEVYHKSMTLKEAIKSSLIILKQVMEEKLNATNIELATVQPGQNFHMFTKEELEEVIKDI
ΔRDU
0.10
0.05
0.00
−0.05
−0.10
0.5
1
5
10
30

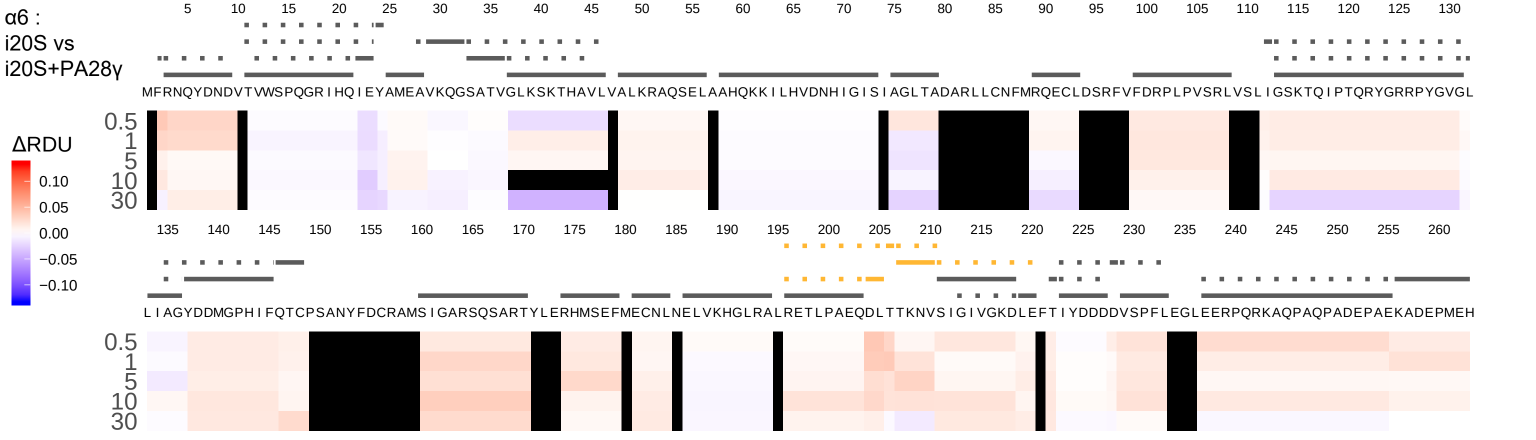
α6 :
i20S vs
i20S+PA28γ
ΔRDU
0.10
0.05
0.00
−0.05
−0.10
0.5
1
5
10
30
MFRNQYDNDVTVWSPQGRIHQIEYAMEAVKQGSATVGLKSKTHAVLVALKRAQSELAAHQKKILHVDNHIGISIAGLTADARLLCNFMRQECLDSRFVFDRPLPVSRLVSLIGSKTQIPTQRYGRRPYGVGL
LIAGYDDMGPHIFQTCPSANYFDCRAMSIGARSQSARTYLERHMSEFMECNLNELVKHGLRALRETLPAEQDLTTKNVSIGIVGKDLEFTIYDDDDVSPFLEGLEERPQRKAQPAQPADEPAEKADEPMEH

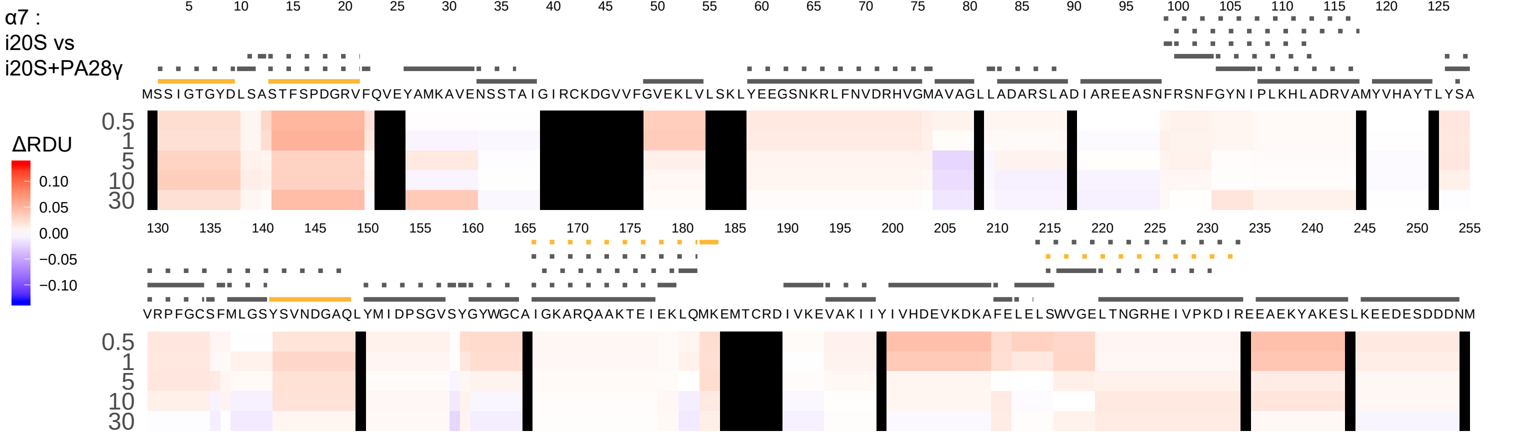

α7 :
i20S vs
i20S+PA28γ
ΔRDU
0.10
0.05
0.00
−0.05
−0.10
0.5
1
5
10
30
MSSIGTGYDLSASTFSPDGRVFQVEYAMKAVENSSTAIGIRCKDGVVFGVEKLVLSKLYEEGSNKRLFNVDRHVGMAVAGLLADARSLADIAREEASNFRSNFGYNIPLKHLADRVAMYVHAYTLYSA
VRPFGCSFMLGSYSVNDGAQLYMIDPSGVSYGYWGCAIGKARQAAKTEIEKLQMKEMTCRDIVKEVAKIIYIVHDEVKDKAFELELSWVGELTNGRHEIVPKDIREEAEKYAKESLKEEDESDDDNM

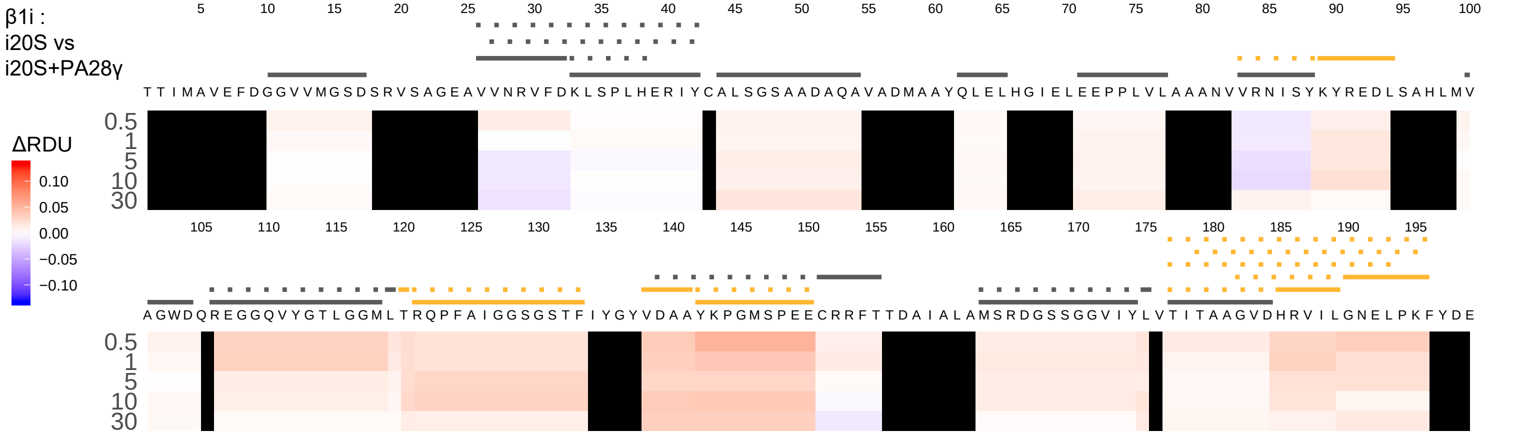

β1i :
i20S vs
i20S+PA28γ
ΔRDU
0.10
0.05
0.00
−0.05
−0.10
0.5
1
5
10
30
TTIMAVEFDGGVVMGSDSRVSAGEAVVNRVFDKLSPLHERIYCALSGSAADAQAVADMAAYQLELHGIELEEPPLVLAAANVVRNISYKYREDLSAHLMV
AGWDQREGGQVYGTLGGMLTRQPFAIGGSGSTFIYGYVDAAYKPGMSPEECRRFTTDAIALAMSRDGSSGGVIYLVTITAAGVDHRVILGNELPKFYDE

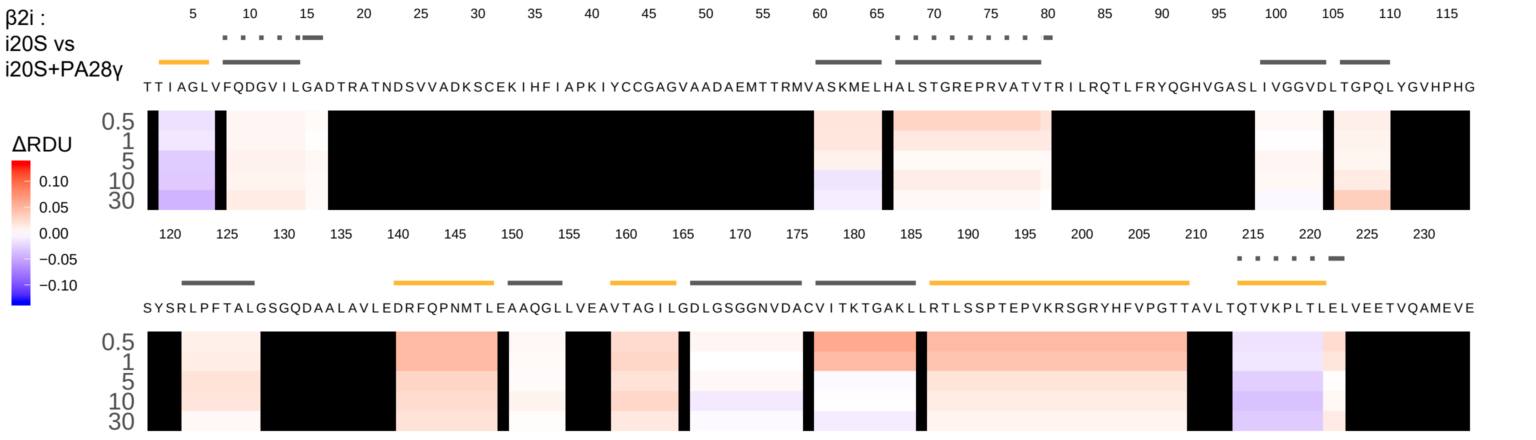

β2i :
i20S vs
i20S+PA28γ
ΔRDU
0.10
0.05
0.00
−0.05
−0.10
5 10 15 20 25 30 35 40 45 50 55 60 65 70 75 80 85 90 95 100 105 110 115
TTIAGLVFQDGVILGADTRATNDSVVADKSCEKIHFIAPKIYCCGAGVAADAEMTTRMVASKMELHALSTGREPRVATVTRILRQTLFRYQGHVGASLIVGGVDLTGPQLYGVHPHG
0.5
1
5
10
30
120 125 130 135 140 145 150 155 160 165 170 175 180 185 190 195 200 205 210 215 220 225 230
SYSRLPFTALGSGQDAALAVLEDRFQPNMTLEAAQGLLVEAVTAGILGDLGSGGNVDACVITKTGAKLLRTLSSPTEPVKRSGRYHFVPGTTAVLTQTVKPLTLELVEETVQAMEVE
0.5
1
5
10
30

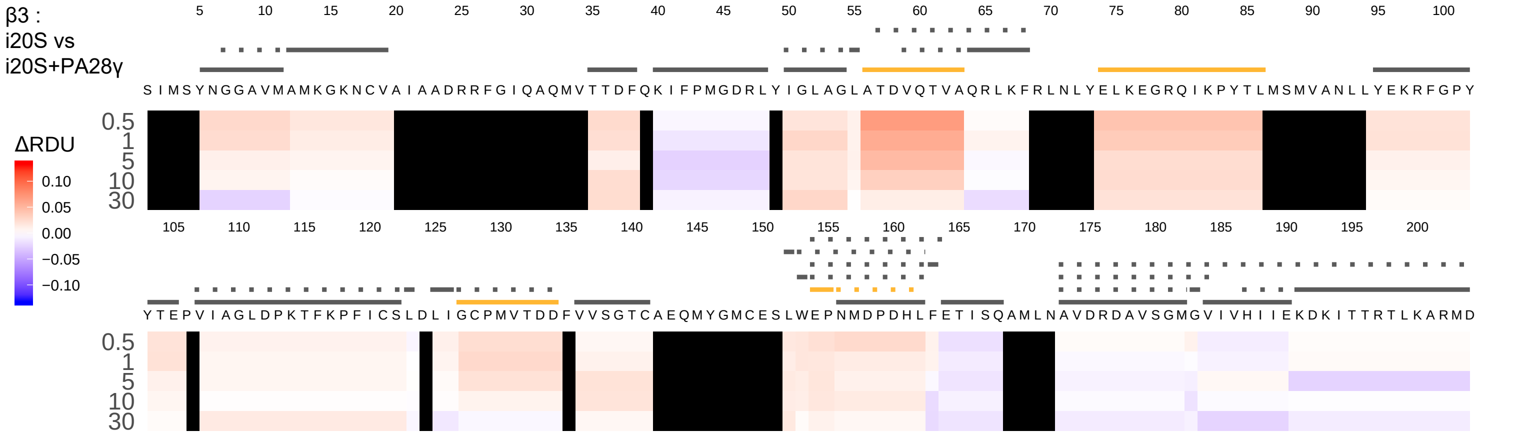
β3 :
i20S vs
i20S+PA28γ
5 10 15 20 25 30 35 40 45 50 55 60 65 70 75 80 85 90 95 100
S I M S Y N G G A V M A M K G K N C V A I A A D R R F G I Q A Q M V T T D F Q K I F P M G D R L Y I G L A G L A T D V Q T V A Q R L K F R L N L Y E L K E G R Q I K P Y T L M S M V A N L L Y E K R F G P Y
0.5
1
5
10
30
ΔRDU
0.10
0.05
0.00
−0.05
−0.10
105 110 115 120 125 130 135 140 145 150 155 160 165 170 175 180 185 190 195 200
Y T E P V I A G L D P K T F K P F I C S L D L I G C P M V T D D F V V S G T C A E Q M Y G M C E S L W E P N M D P D H L F E T I S Q A M L N A V D R D A V S G M G V I V H I I E K D K I T T R T L K A R M D

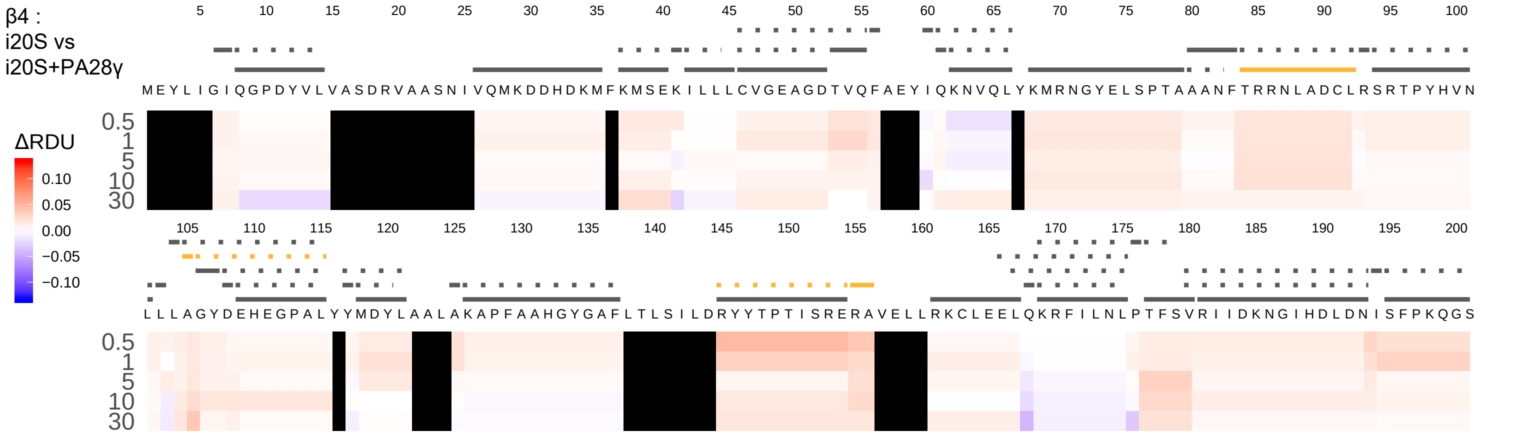
β4 :
i20S vs
i20S+PA28γ
ΔRDU
0.10
0.05
0.00
−0.05
−0.10
0.5
1
5
10
30
MEYLIGIQGPDYVLVASDRVAASNIVQMKDDHDKMFKMSEKILLLCVGEAGDTVQFAEYIQKNVQLYKMRNGYELSPTAAANFTRRNLADCLRSRTPYHVN
LLLAGYDEHEGPALYYMDYLAALAKAPFAAHGYGAFLTLSILDRYYTPTISRERAVELLRKCLEELQKRFILNLPTFSVRIIDKNGIHDLDNISFPKQGS

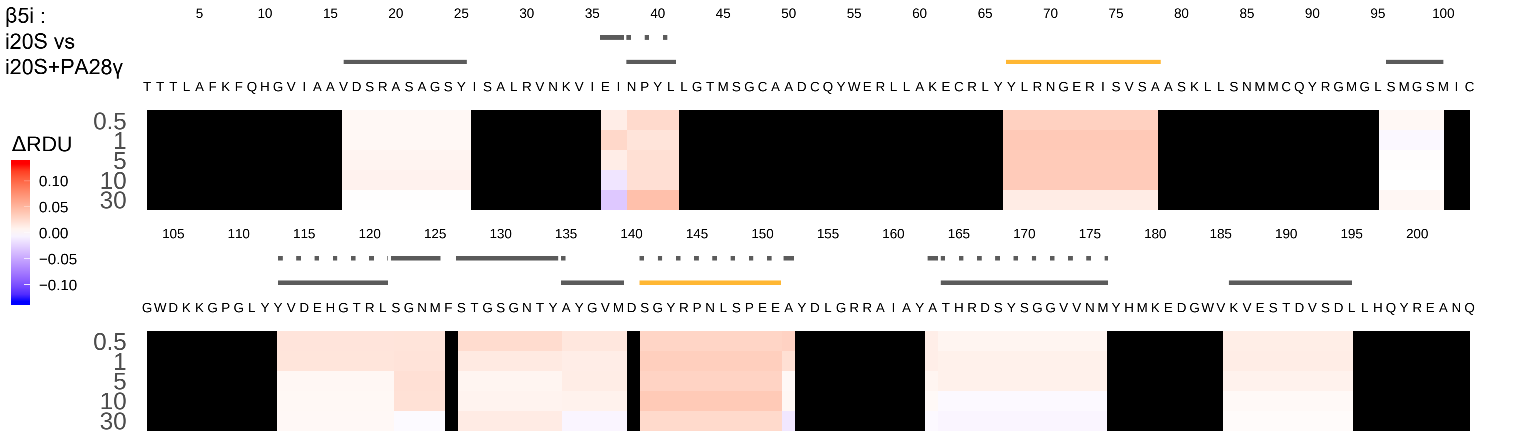
β5i :
i20S vs
i20S+PA28γ
ΔRDU
0.10
0.05
0.00
−0.05
−0.10
0.5
1
5
10
30
TTTLAFKFQHGVIAAVDSRASAGSYISALRVNKVIEINPYLLGTMSGCAADCQYWERLLAKECRLYYLRNGERISVSAASKLLSNMMCQYRGMGLSMGSMIC
GWDKKGPGLYYVDEHGTRLSGNMFSTGSGNTYAYGVMDSGYRPNLSPEEAYDLGRRAIAYATHRDSYSGGVVNMYHMKEDGWVKVESTDVSDLLHQYREANQ

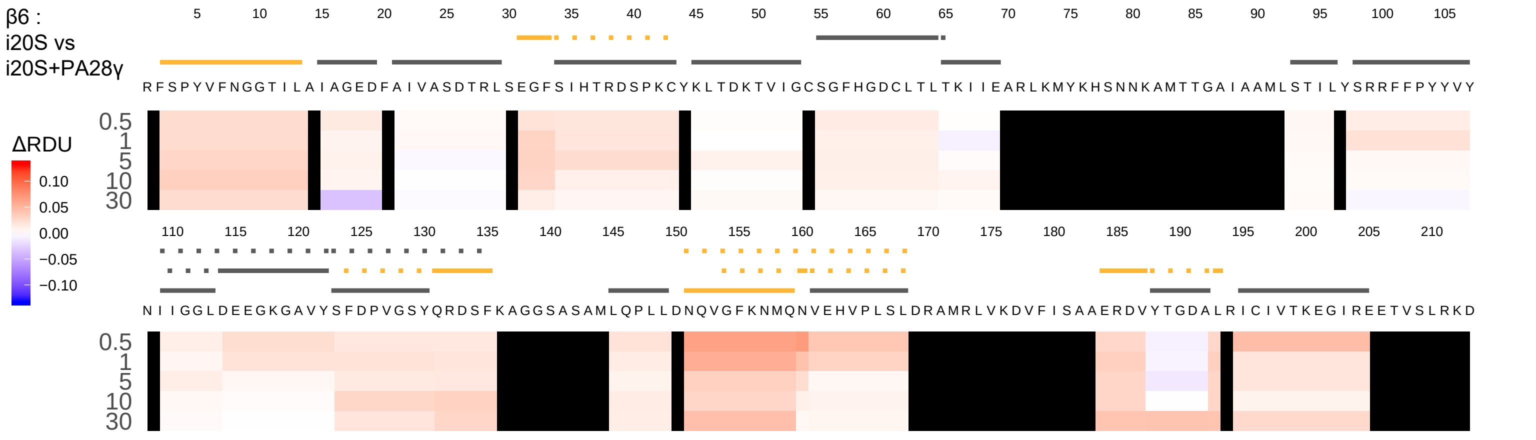

β6 :
i20S vs
i20S+PA28γ
ΔRDU
0.10
0.05
0.00
−0.05
−0.10
0.5
1
5
10
30
RFSPYVFNGGTILAIAGEDFAIVASDTRLSEGFSIHTRDSPKCYKLTDKTVIGCSGFHGDCLTLTKIIEARLKMYKHSNNKAMTTGAIAAMLSTILYSRRFFPYYVVY
NIIGGLDEEGKGAVYSFDPVGSYQRDSFKAGGSASAMLQPLLDNQVGFKNMQNVEHVPLSLDRAMRLVKDVFISAAERDVYTGDALRICIVTKEGIREETVSLRKD

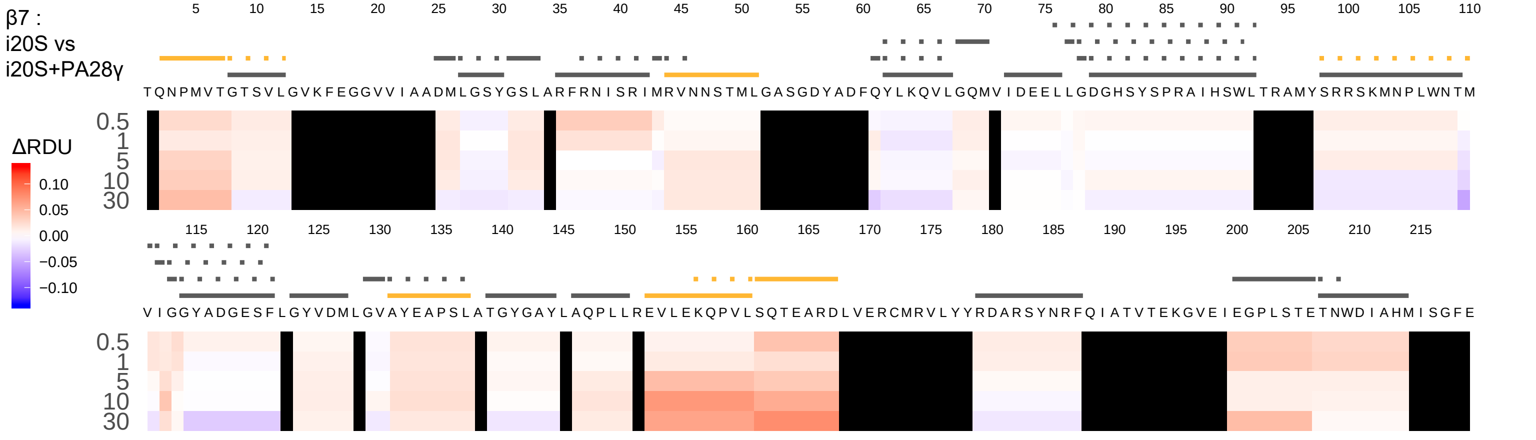

β7 :
i20S vs
i20S+PA28γ
ΔRDU
0.10
0.05
0.00
−0.05
−0.10
0.5
1
5
10
30
TQNPMVTGTSVLGVKFEGGVVIAADMLGSYGSLARFRNISRIMRVNNSTMLGASGDYADFQYLKQVLGQMVIDEELLGDGHSYSPRAIHSWLTRAMYSRRSKMNPLWNTM
VIGGYADGESFLGYVDMLGVAYEAPSLATGYGAYLAQPLLREVLEKQPVLSQTEARDLVERCMRVLYYRDARSYNRFQIATVTEKGVEIEGPLSTETNWDIAHMISGFE
