## Supplementary material for "Conformational maps of human 20S proteasomes reveal PA28- and immuno-dependent inter-ring crosstalks": Fig. S12

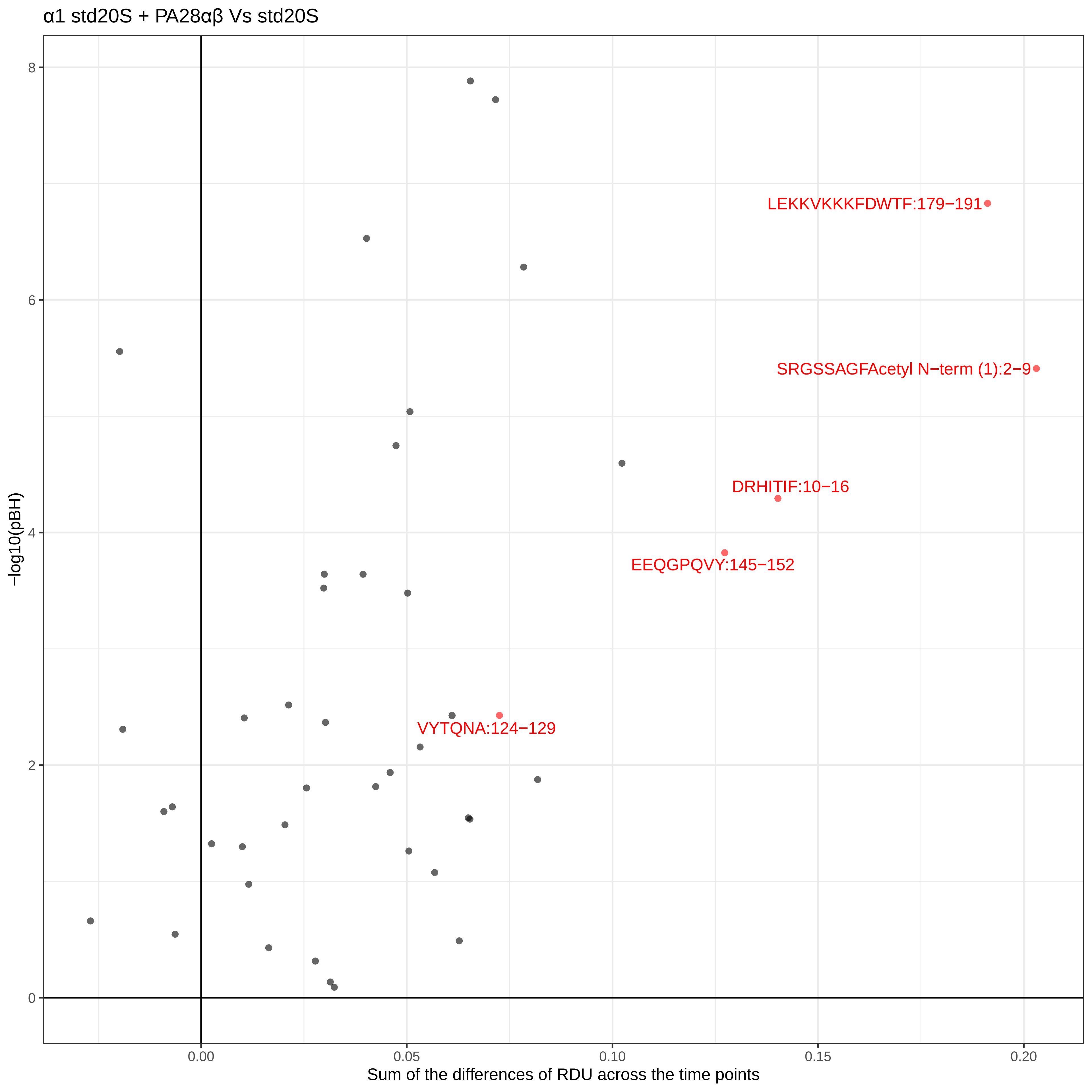
α1 std20S + PA28αβ Vs std20S
LEKKVKKKFDWTF:179−191
SRGSSAGFAcetyl N−term (1):2−9
DRHITIF:10−16
EEQGPQVY:145−152
VYTQNA:124−129
−log10(pBH)
8
6
4
2
0
0.00
0.05
0.10
0.15
0.20
Sum of the differences of RDU across the time points

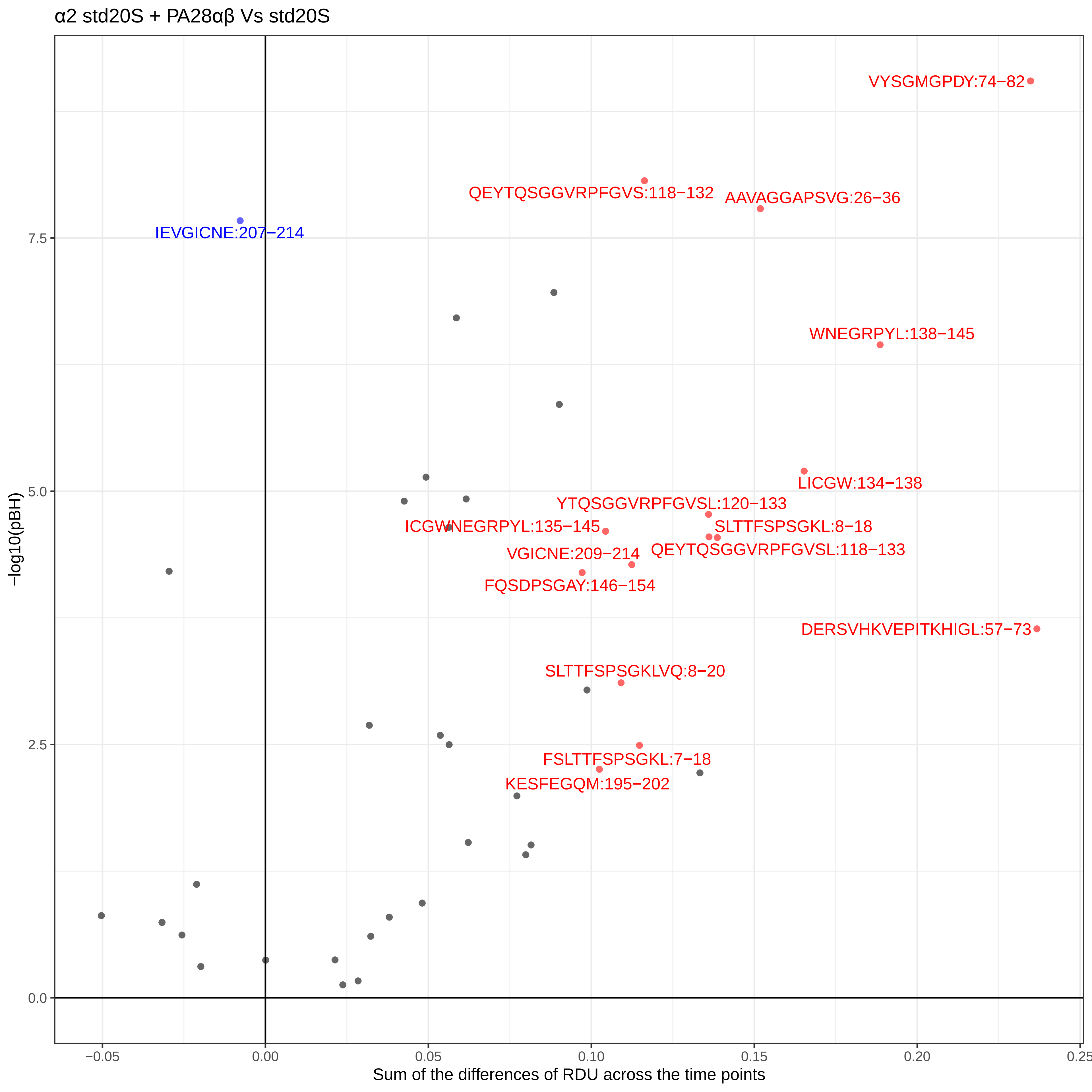
α2 std20S + PA28αβ Vs std20S
VYSGMGPDY:74−82
QEYTQSGGVRPFGVS:118−132
AAVAGGAPSVG:26−36
IEVGICNE:207−214
WNEGRPYL:138−145
LICGW:134−138
YTQSGGVRPFGVSL:120−133
ICGWNEGRPYL:135−145
SLTTFSPSGKL:8−18
VGICNE:209−214
QEYTQSGGVRPFGVSL:118−133
FQSDPSGAY:146−154
DERSVHKVEPITKHIGL:57−73
SLTTFSPSGKLVQ:8−20
FSLTTFSPSGKL:7−18
KESFEGQM:195−202
7.5
5.0
2.5
0.0
−log10(pBH)
−0.05
0.00
0.05
0.10
0.15
0.20
0.25
Sum of the differences of RDU across the time points

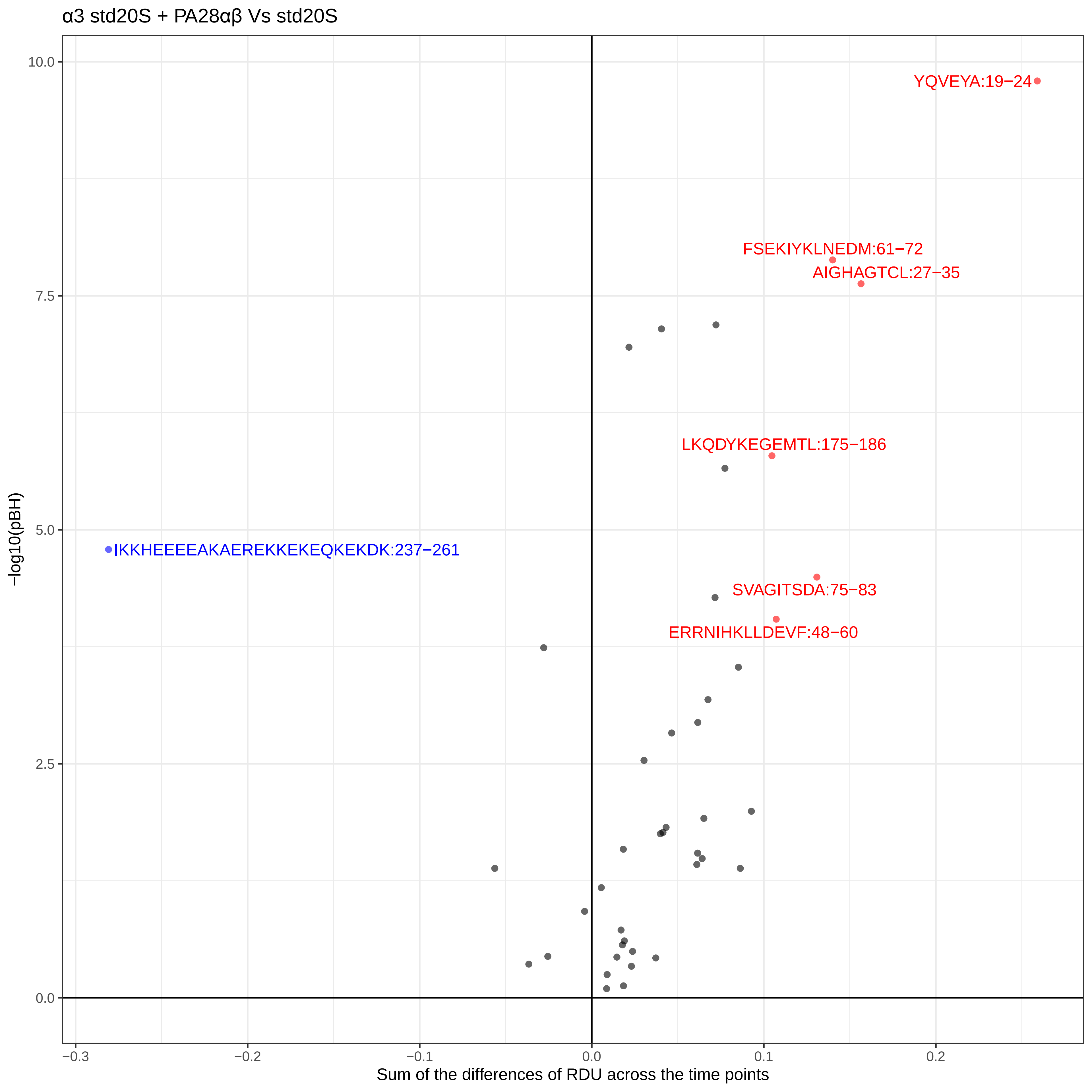
α3 std20S + PA28αβ Vs std20S
YQVEYA:19−24
FSEKIYKLNEDM:61−72
AIGHAGTCL:27−35
LKQDYKEGEMTL:175−186
IKKHEEEEAKAEREKKEKEQKEKDK:237−261
SVAGITSDA:75−83
ERRNIHKLLDEVF:48−60
−log10(pBH)
10.0
7.5
5.0
2.5
0.0
−0.3
−0.2
−0.1
0.0
0.1
0.2
Sum of the differences of RDU across the time points

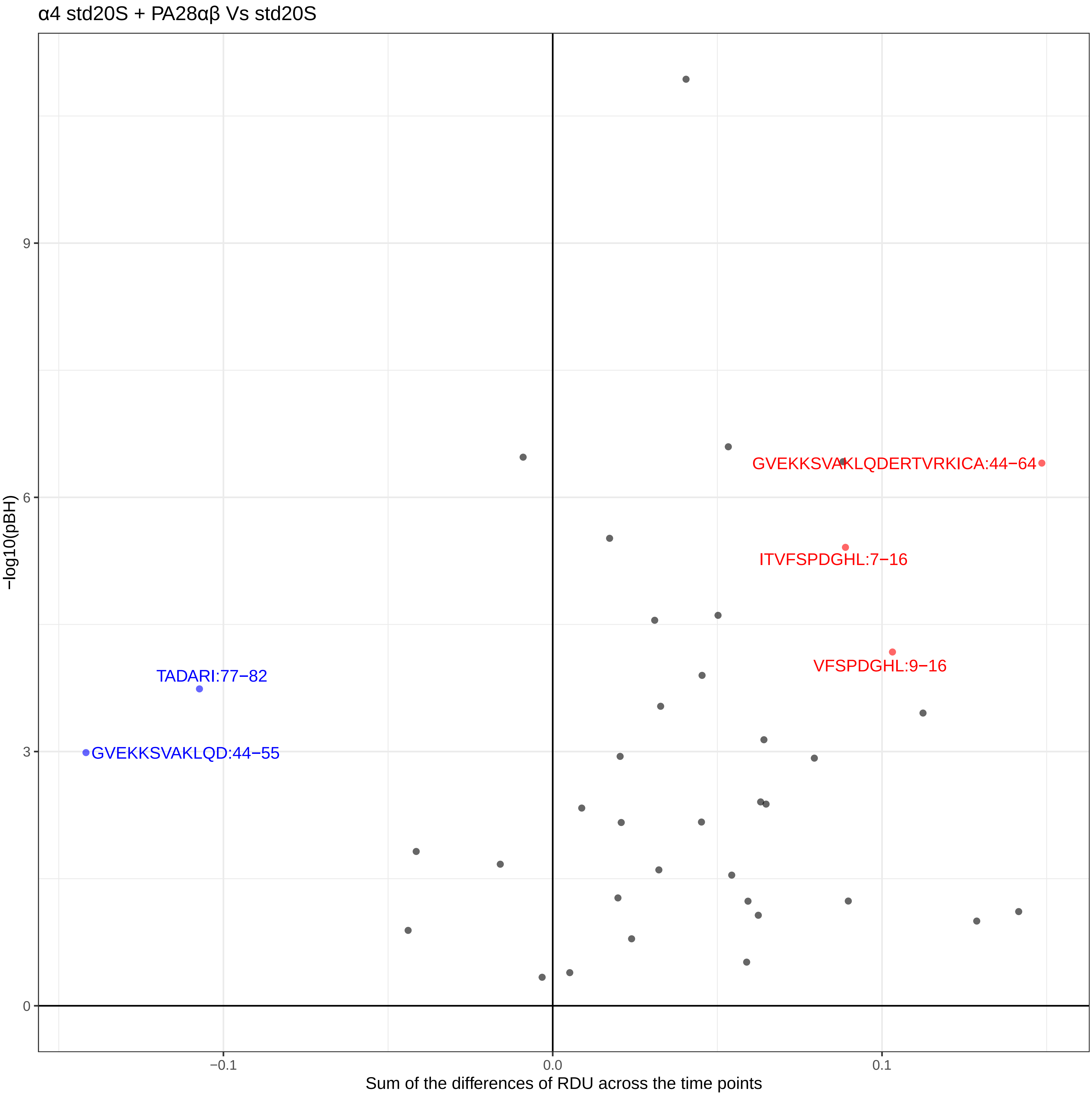

α4 std20S + PA28αβ Vs std20S
−log10(pBH)
Sum of the differences of RDU across the time points
9
6
3
0
−0.1
0.0
0.1
GVEKKSVAKLQDERTVRKICA:44−64
ITVFSPDGHL:7−16
VFSPDGHL:9−16
TADARI:77−82
GVEKKSVAKLQD:44−55

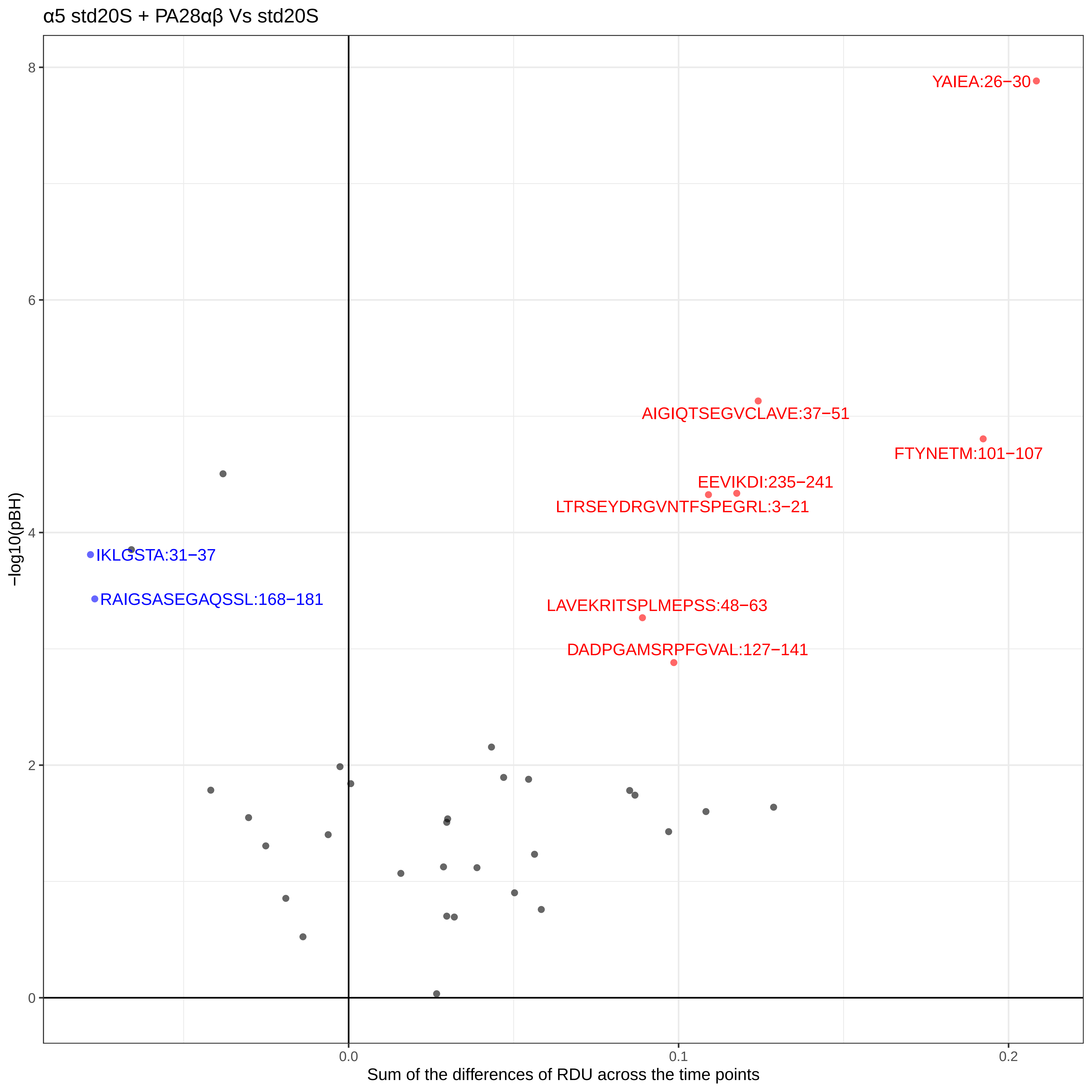
α5 std20S + PA28αβ Vs std20S
YAIEA:26−30
AIGIQTSEGVCLAVE:37−51
FTYNETM:101−107
EEVIKDI:235−241
LTRSEYDRGVNTFSPEGRL:3−21
IKLGSTA:31−37
RAIGSASEGAQSSL:168−181
LAVEKRITSPLMEPSS:48−63
DADPGAMSRPFGVAL:127−141
−log10(pBH)
Sum of the differences of RDU across the time points
0
2
4
6
8
0.0
0.1
0.2

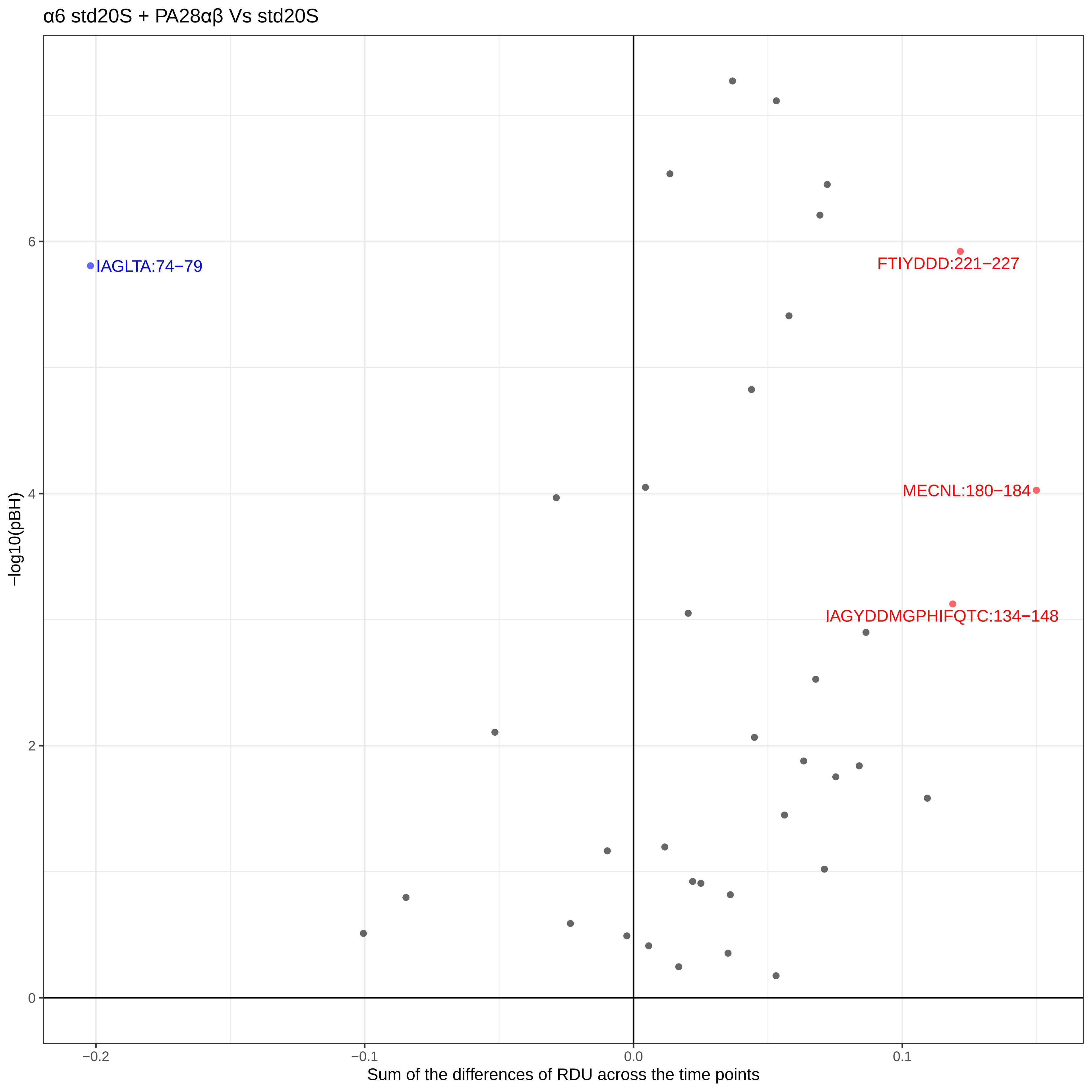
α6 std20S + PA28αβ Vs std20S
IAGLTA:74−79
FTIYDDD:221−227
MECNL:180−184
IAGYDDMGPHIFQTC:134−148
−log10(pBH)
0
2
4
6
−0.2
−0.1
0.0
0.1
Sum of the differences of RDU across the time points

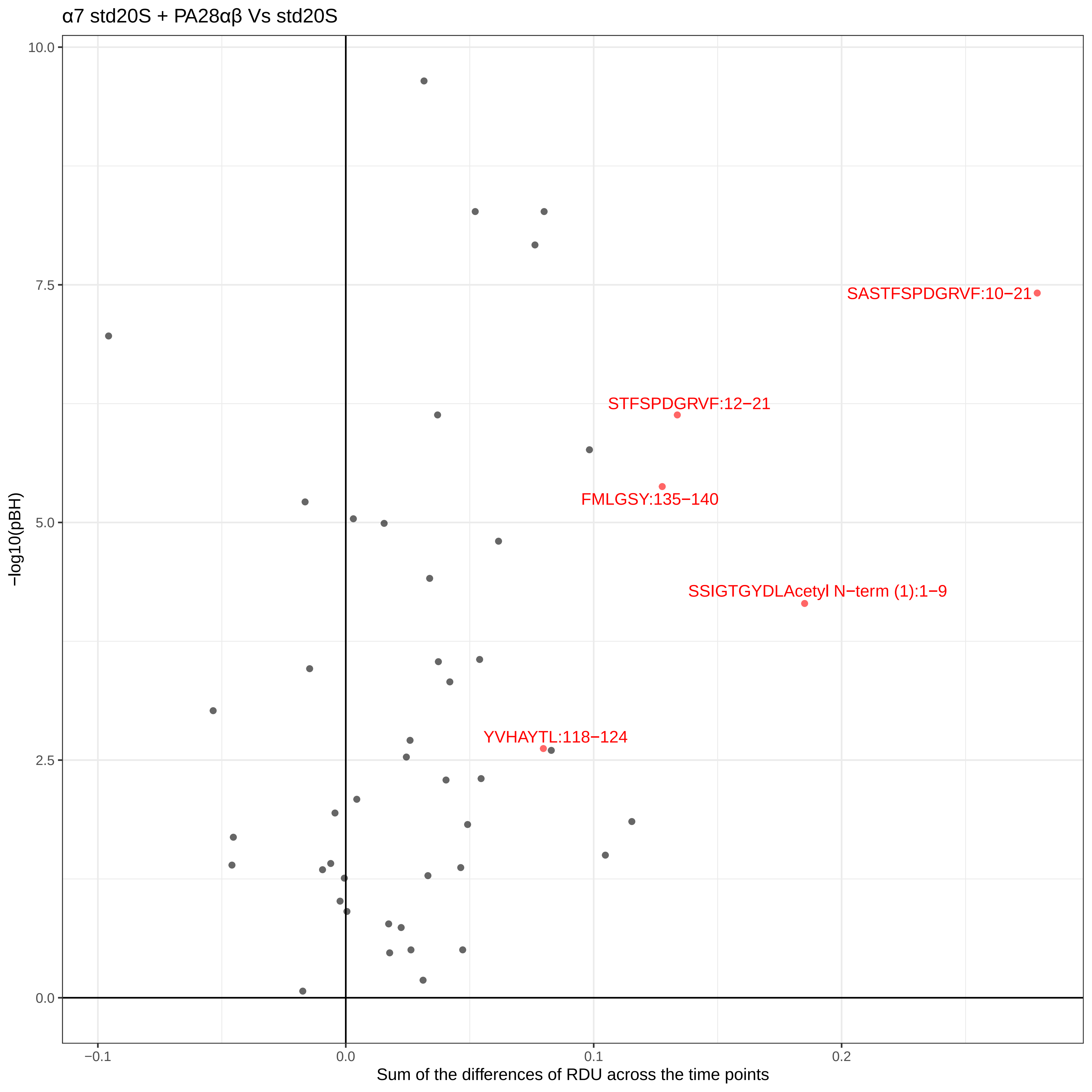

α7 std20S + PA28αβ Vs std20S
SASTFSPDGRVF:10−21
STFSPDGRVF:12−21
FMLGSY:135−140
SSIGTGYDLAcetyl N−term (1):1−9
YVHAYTL:118−124
−log10(pBH)
10.0
7.5
5.0
2.5
0.0
−0.1
0.0
0.1
0.2
Sum of the differences of RDU across the time points

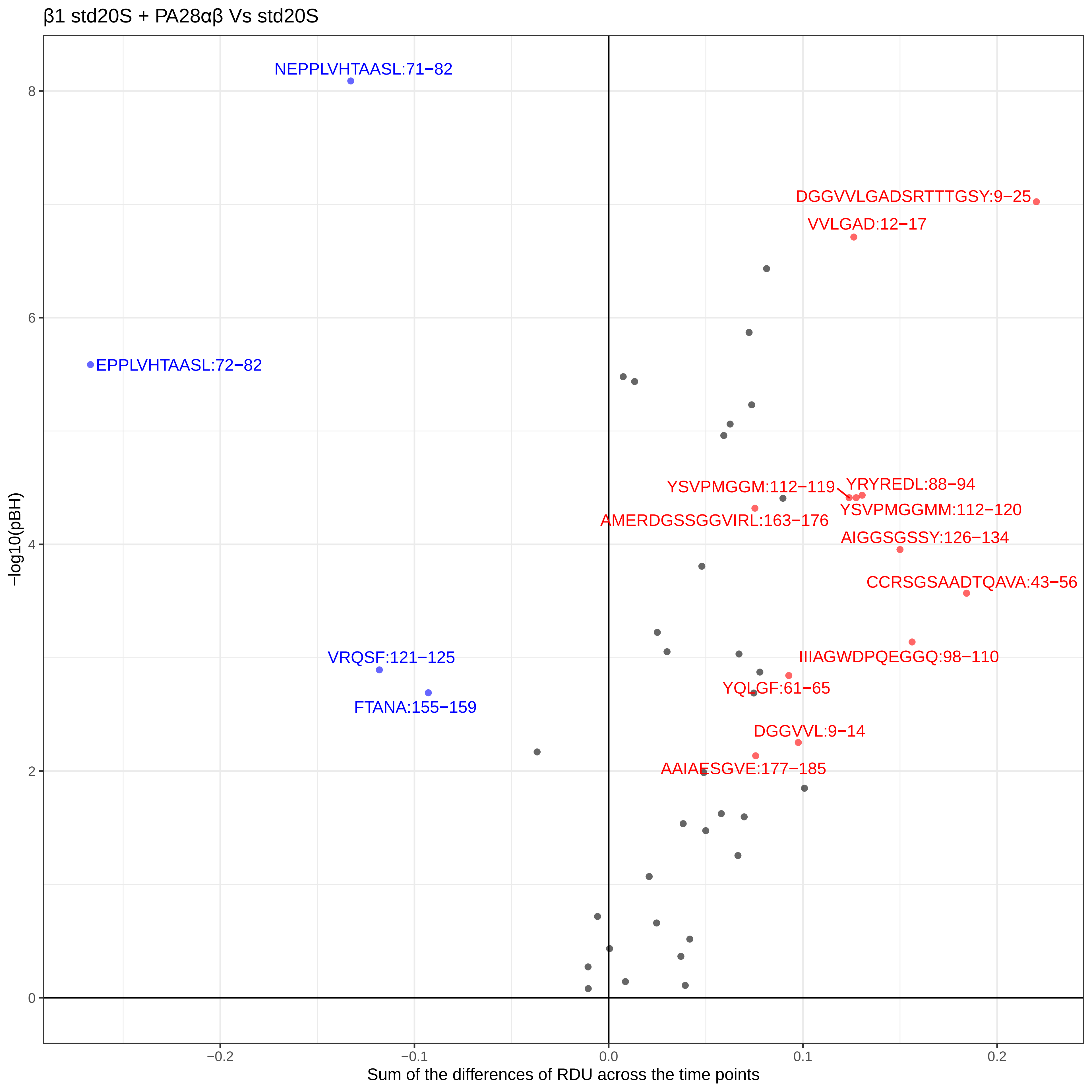

β1 std20S + PA28αβ Vs std20S
NEPPLVHTAASL:71−82
EPPLVHTAASL:72−82
VRQSF:121−125
FTANA:155−159
DGGVVLGADSRTTTGSY:9−25
VVLGAD:12−17
YSVPMGGM:112−119
YRYREDL:88−94
YSVPMGGMM:112−120
AMERDGSSGGVIRL:163−176
AIGGSGSSY:126−134
CCRSGSAADTQAVA:43−56
IIIAGWDPQEGGQ:98−110
YQLGF:61−65
DGGVVL:9−14
AAIAESGVE:177−185
−log10(pBH)
Sum of the differences of RDU across the time points
0
2
4
6
8
−0.2
−0.1
0.0
0.1
0.2

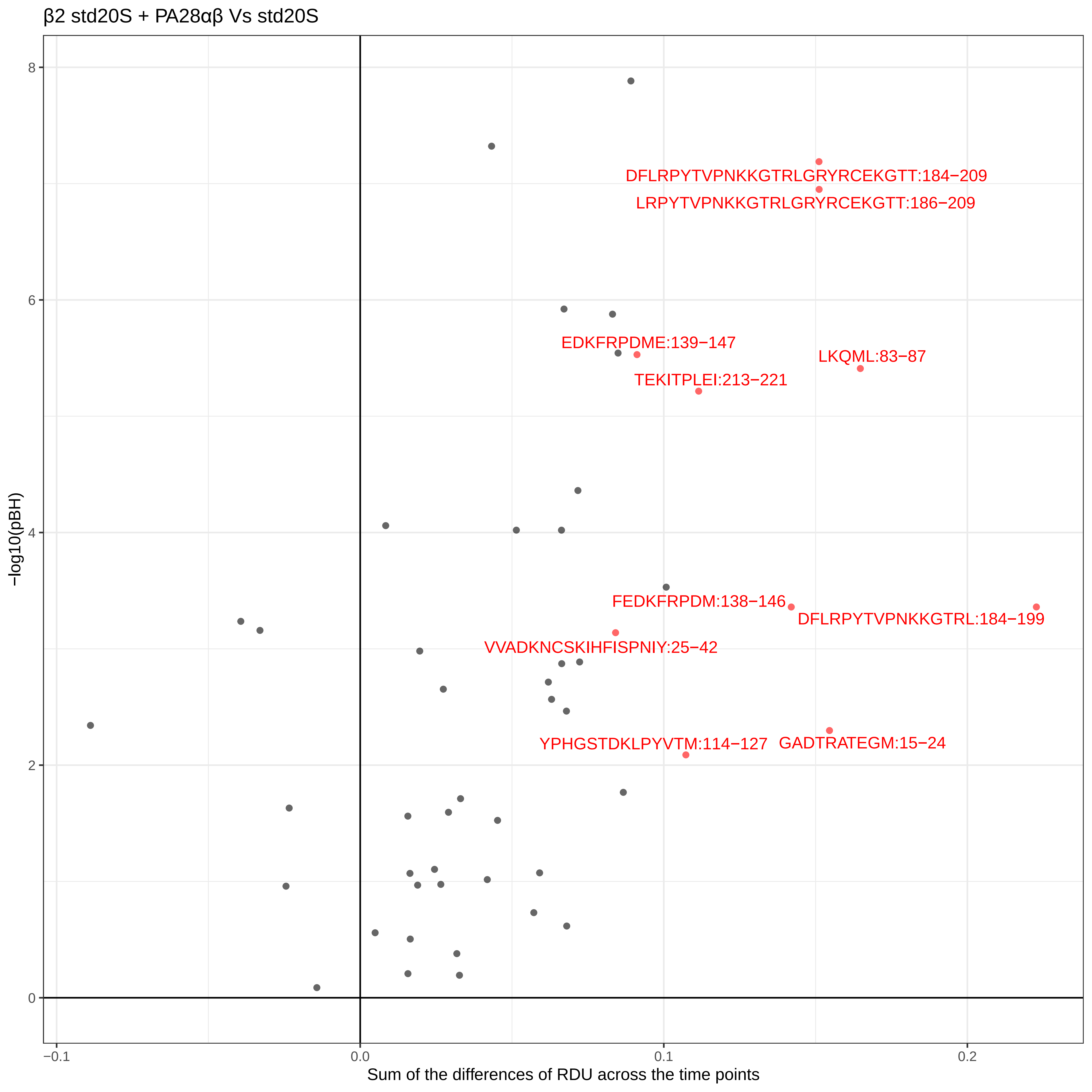

β2 std20S + PA28αβ Vs std20S
DFLRPYTVPNKKGTRLGRYRCEKGTT:184−209
LRPYTVPNKKGTRLGRYRCEKGTT:186−209
EDKFRPDME:139−147
LKQML:83−87
TEKITPLEI:213−221
FEDKFRPDM:138−146
DFLRPYTVPNKKGTRL:184−199
VVADKNCSKIHFISPNIY:25−42
YPHGSTDKLPYVTM:114−127
GADTRATEGM:15−24
−log10(pBH)
8
6
4
2
0
−0.1
0.0
0.1
0.2
Sum of the differences of RDU across the time points

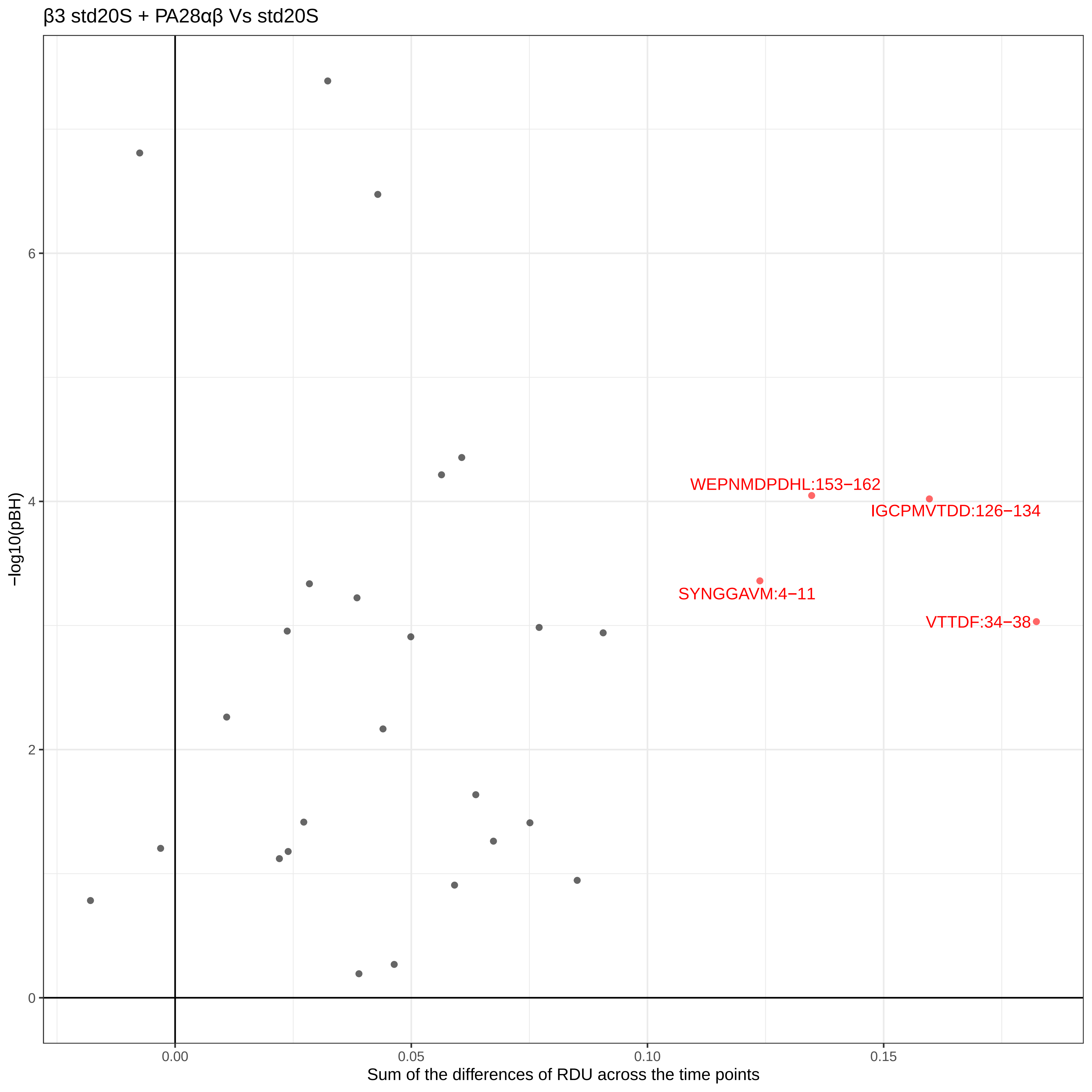

β3 std20S + PA28αβ Vs std20S
WEPNMDPDHL:153−162
IGCPMVTDD:126−134
SYNGGAVM:4−11
VTTDF:34−38
−log10(pBH)
0
2
4
6
0.00
0.05
0.10
0.15
Sum of the differences of RDU across the time points

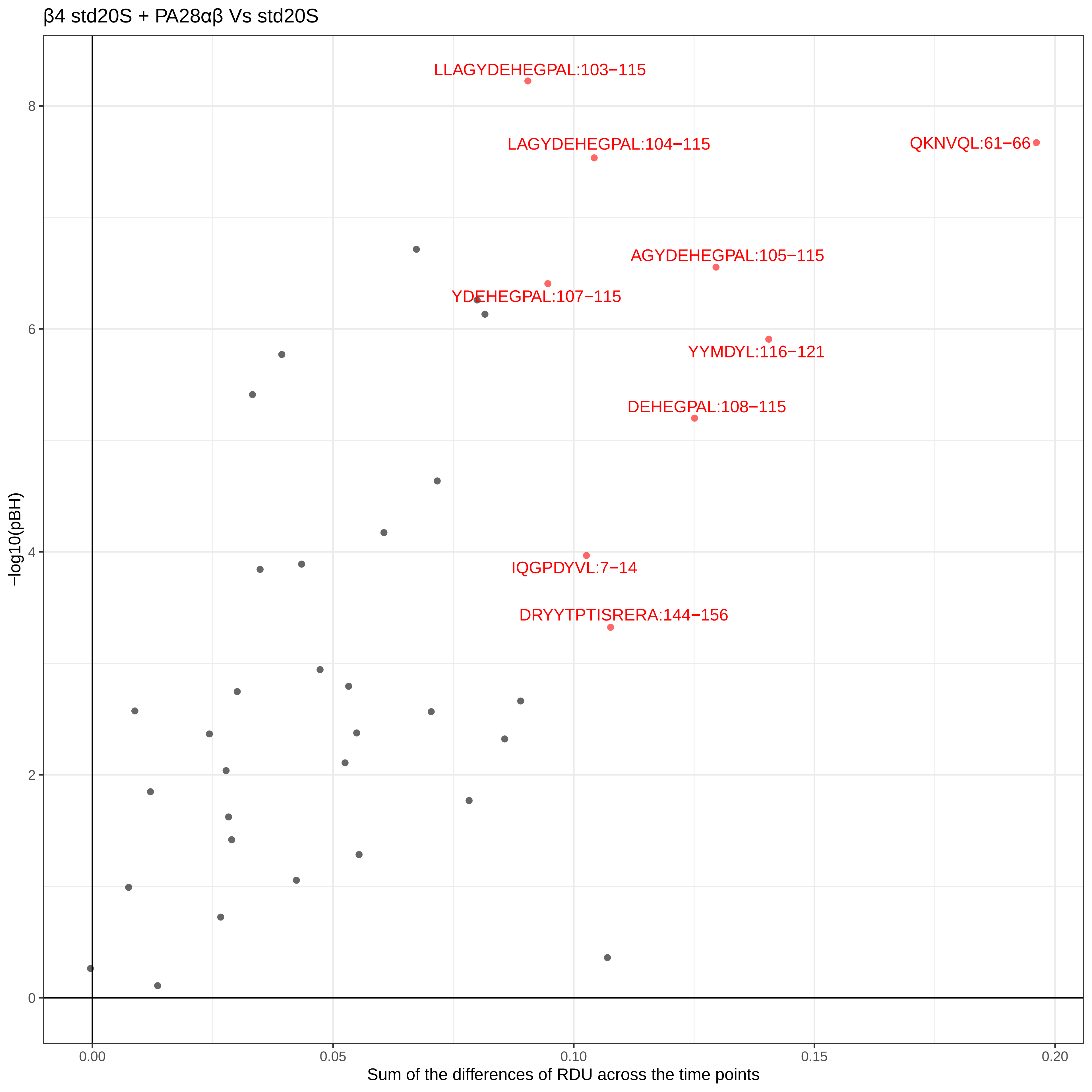

β4 std20S + PA28αβ Vs std20S
LLAGYDEHEGPAL:103−115
LAGYDEHEGPAL:104−115
QKNVQL:61−66
AGYDEHEGPAL:105−115
YDEHEGPAL:107−115
YYMDYL:116−121
DEHEGPAL:108−115
IQGPDYVL:7−14
DRYYTPTISRERA:144−156
−log10(pBH)
0
2
4
6
8
0.00
0.05
0.10
0.15
0.20
Sum of the differences of RDU across the time points

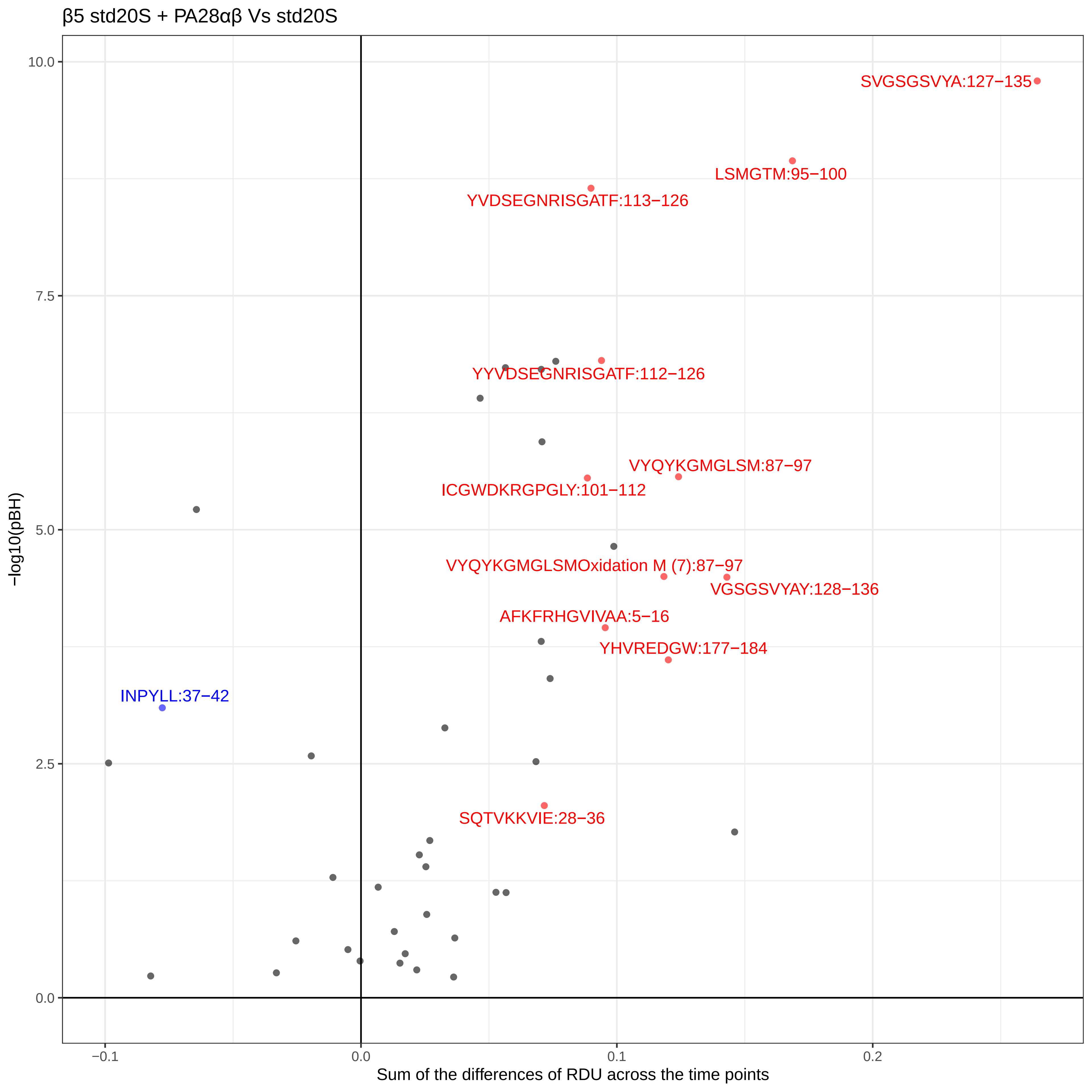
β5 std20S + PA28αβ Vs std20S
SVGSGSVYA:127−135
LSMGTM:95−100
YVDSEGNRISGATF:113−126
YYVDSEGNRISGATF:112−126
VYQYKGMGLSM:87−97
ICGWDKRGPGLY:101−112
VYQYKGMGLSMOxidation M (7):87−97
VGSGSVYAY:128−136
AFKFRHGVIVAA:5−16
YHVREDGW:177−184
INPYLL:37−42
SQTVKKVIE:28−36
−log10(pBH)
10.0
7.5
5.0
2.5
0.0
−0.1
0.0
0.1
0.2
Sum of the differences of RDU across the time points

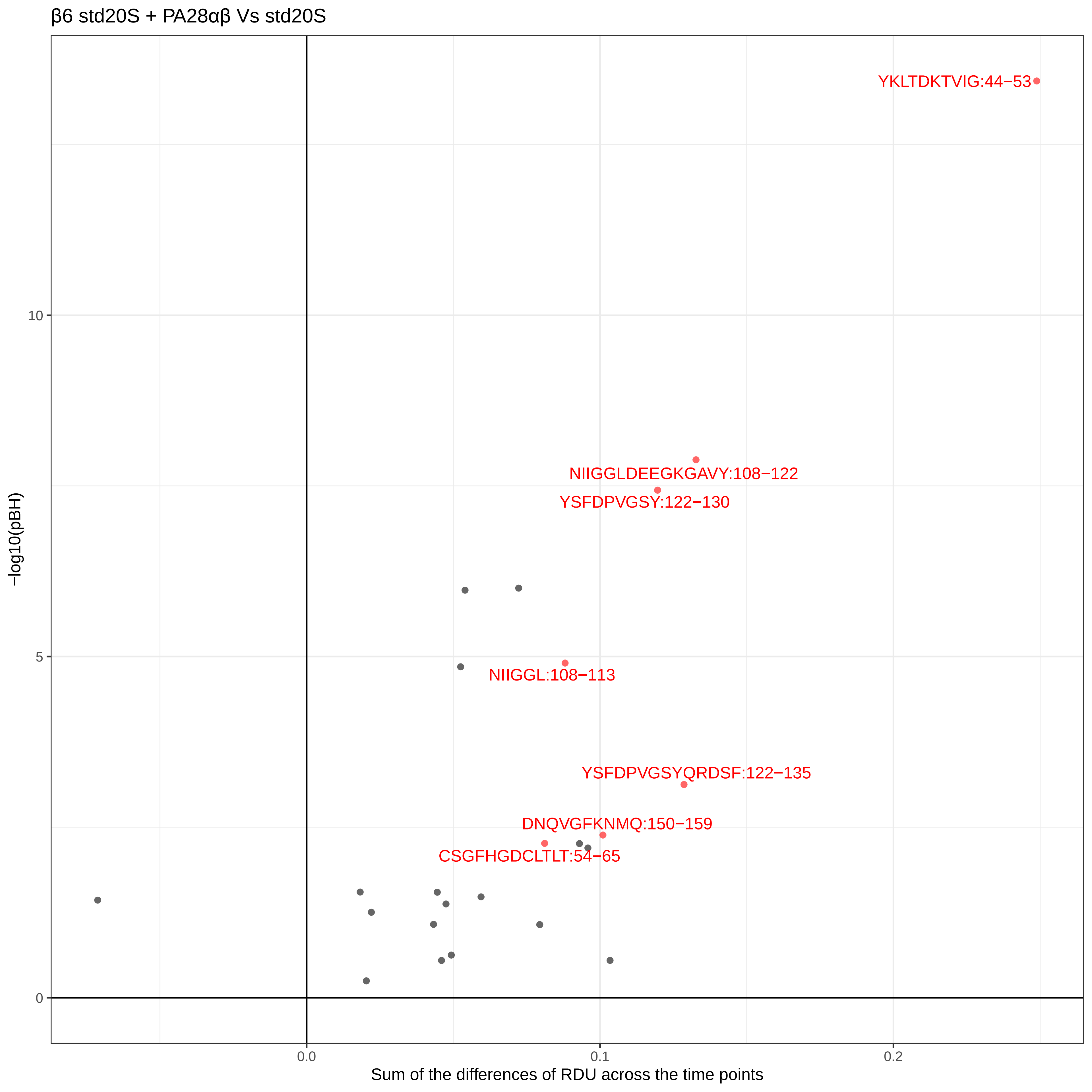

β6 std20S + PA28αβ Vs std20S
YKLTDKTVIG:44−53
NIIGGLDEEGKGAVY:108−122
YSFDPVGSY:122−130
NIIGGL:108−113
YSFDPVGSYQRDSF:122−135
DNQVGFKNMQ:150−159
CSGFHGDCLTLT:54−65
−log10(pBH)
10
5
0
0.0
0.1
0.2
Sum of the differences of RDU across the time points

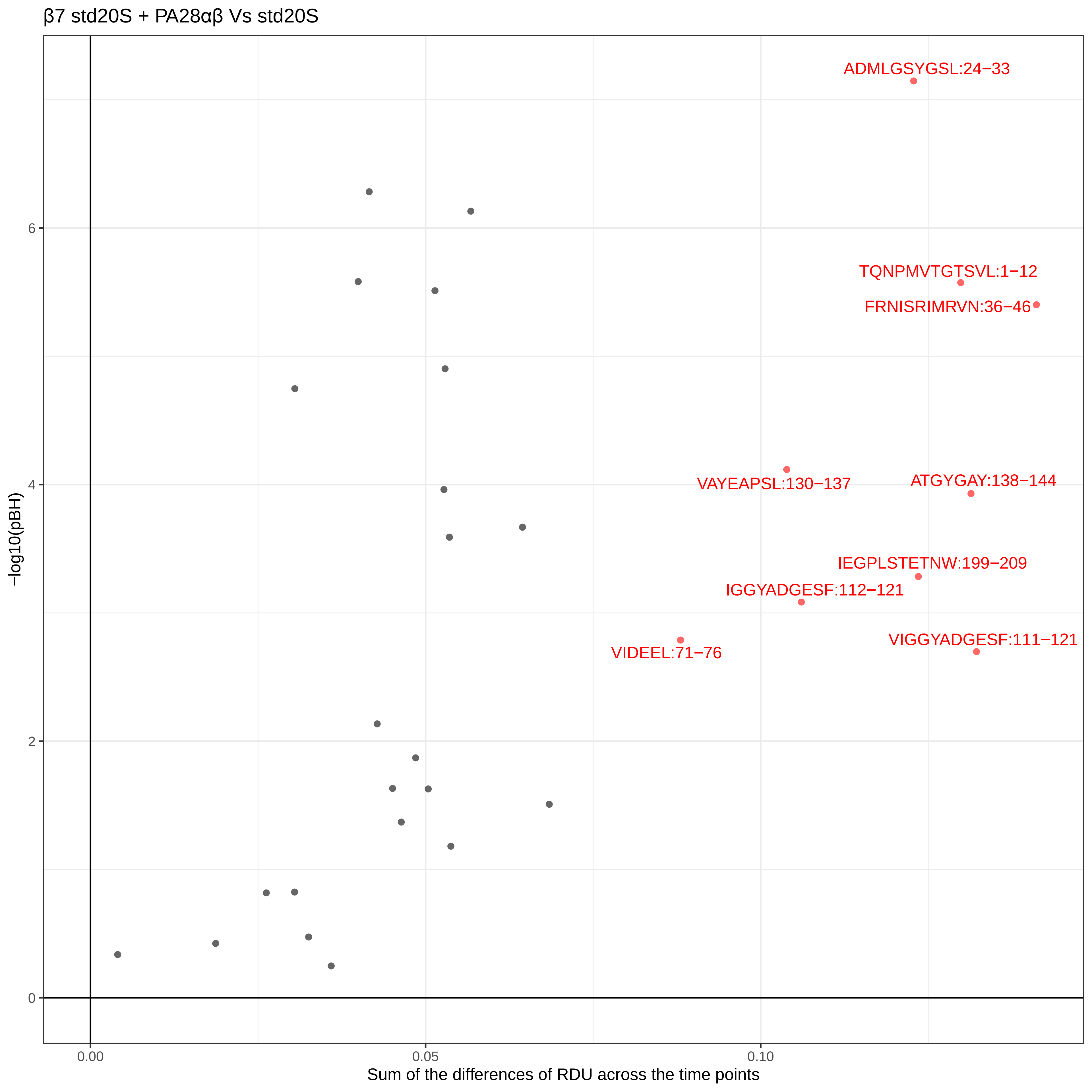
β7 std20S + PA28αβ Vs std20S
ADMLGSYGSL:24−33
TQNPMVTGTSVL:1−12
FRNISRIMRVN:36−46
VAYEAPSL:130−137
ATGYGAY:138−144
IEGPLSTETNW:199−209
IGGYADGESF:112−121
VIDEEL:71−76
VIGGYADGESF:111−121
−log10(pBH)
0
2
4
6
0.00
0.05
0.10
Sum of the differences of RDU across the time points

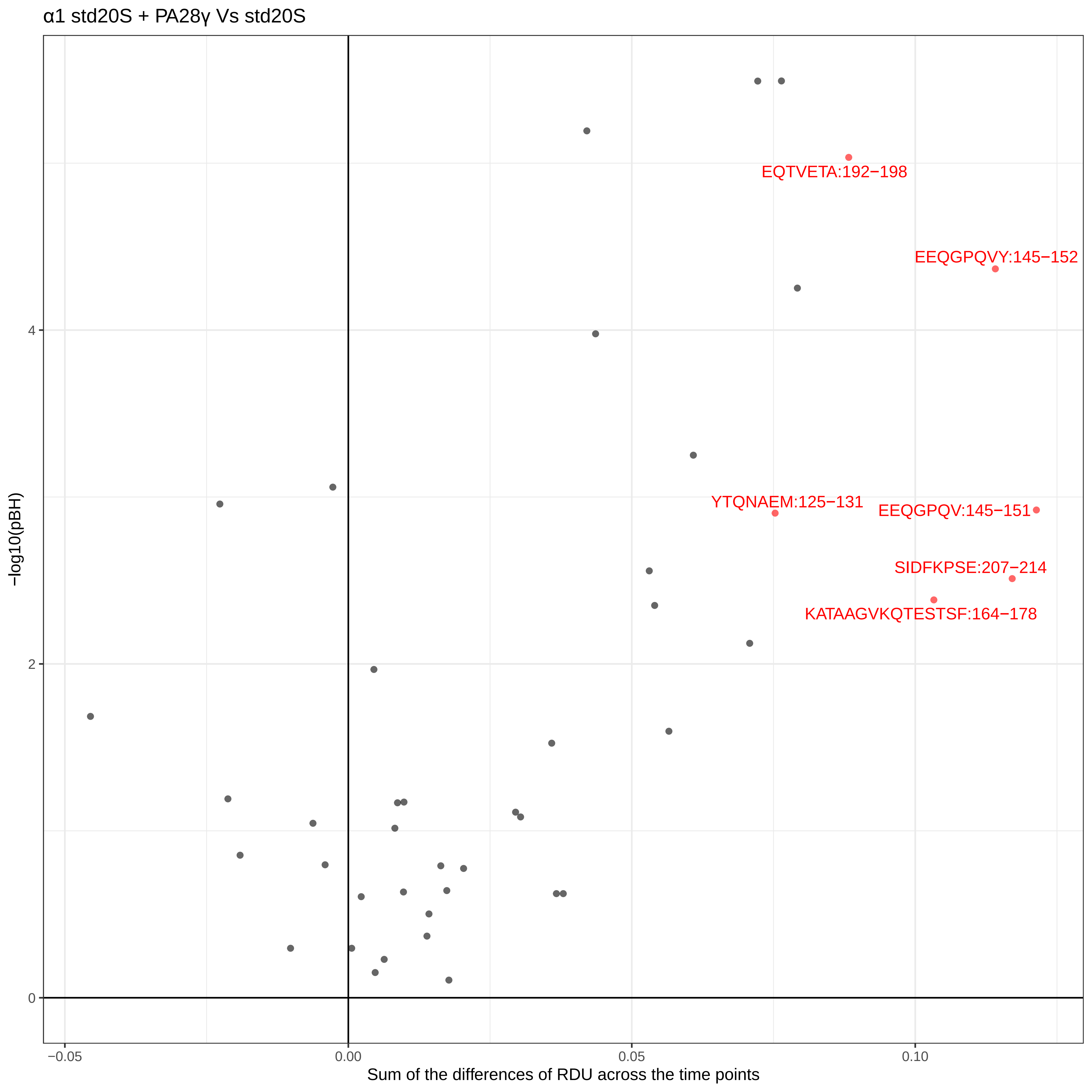

α1 std20S + PA28γ Vs std20S
EQTVETA:192−198
EEQGPQVY:145−152
YTQNAEM:125−131
EEQGPQV:145−151
SIDFKPSE:207−214
KATAAGVKQTESTSF:164−178
−log10(pBH)
4
2
0
−0.05
0.00
0.05
0.10
Sum of the differences of RDU across the time points

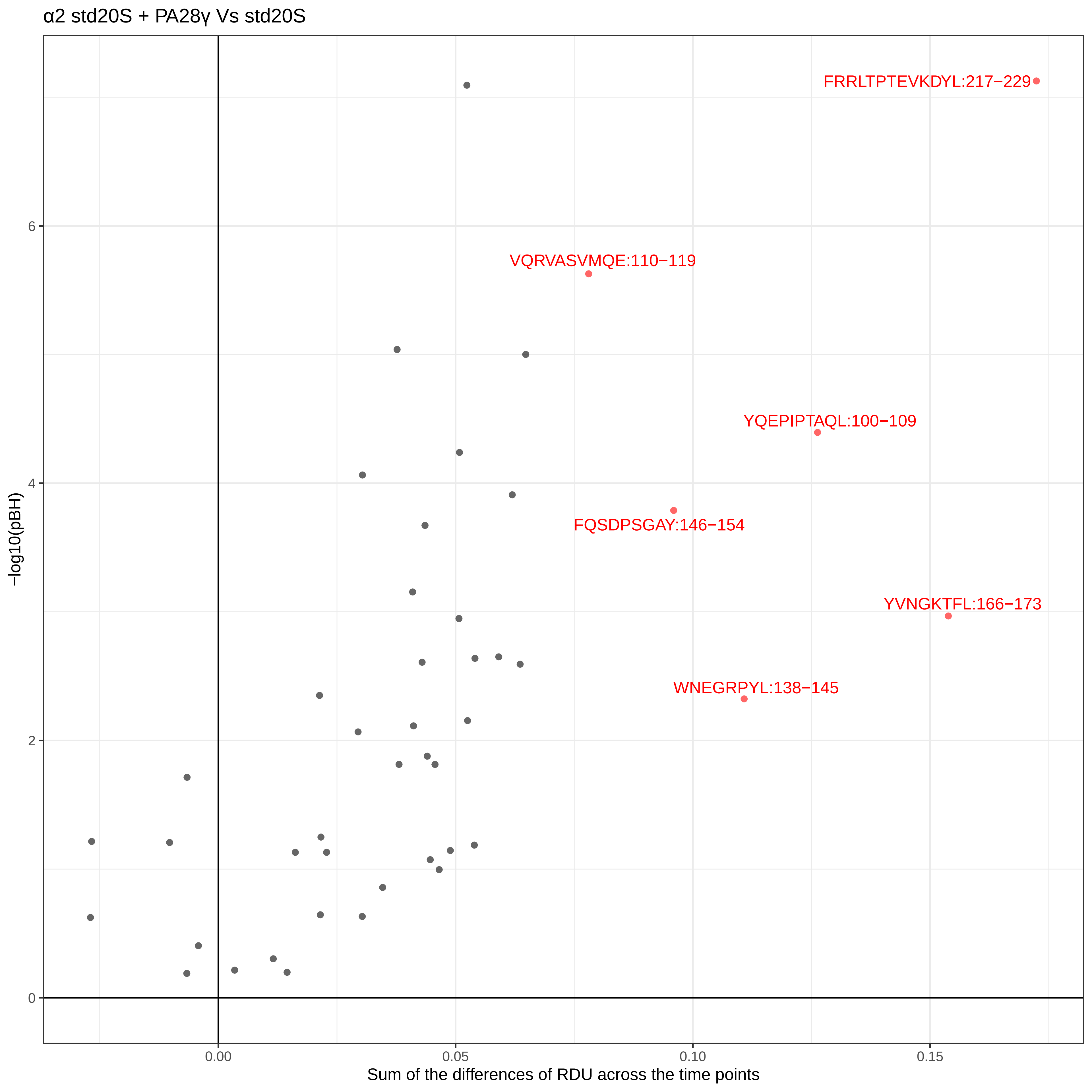
α2 std20S + PA28γ Vs std20S
FRRLTPTEVKDYL:217−229
VQRVASVMQE:110−119
YQEPIPTAQL:100−109
FQSDPSGAY:146−154
YVNGKTFL:166−173
WNEGRPYL:138−145
−log10(pBH)
6
4
2
0
0.00
0.05
0.10
0.15
Sum of the differences of RDU across the time points

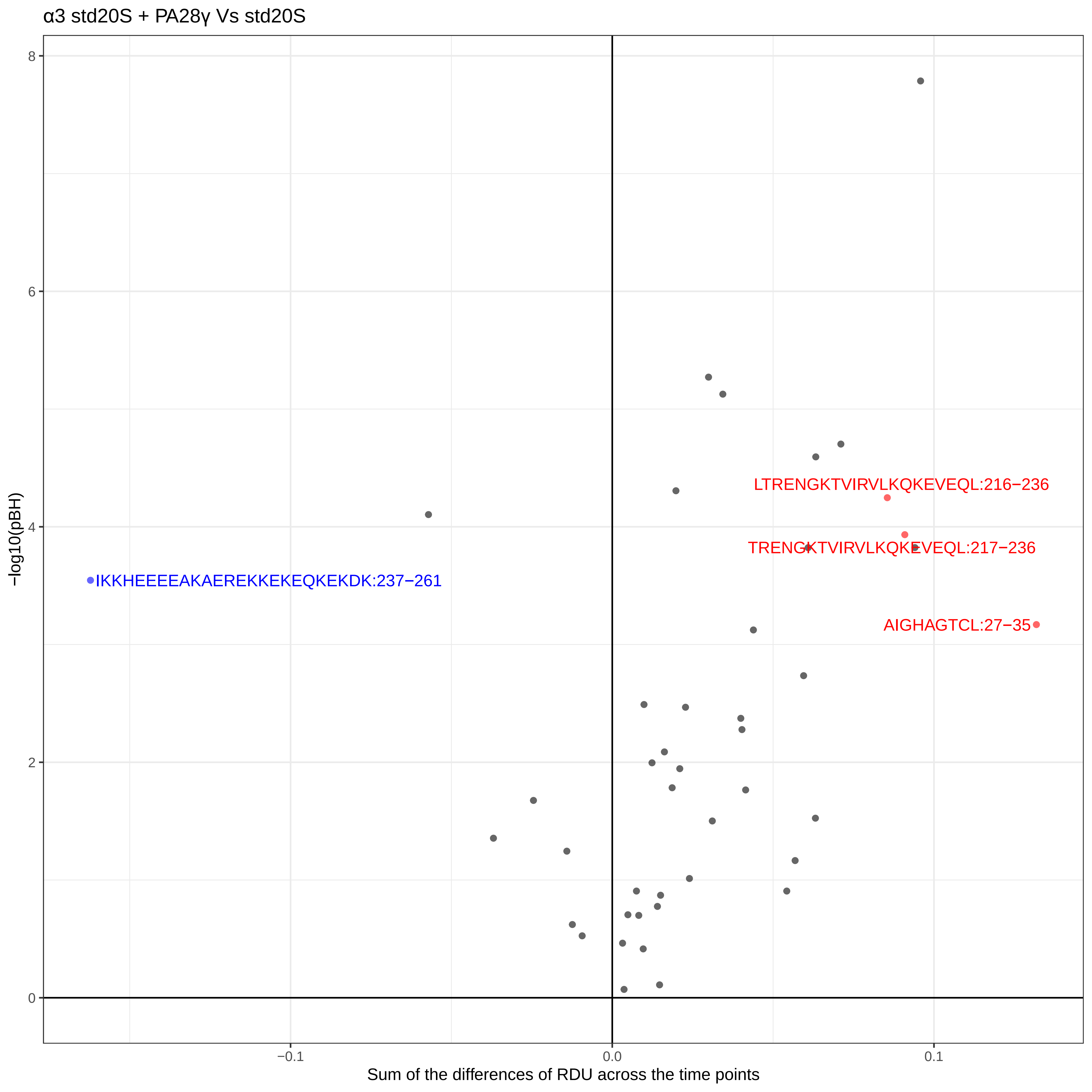
α3 std20S + PA28γ Vs std20S
−log10(pBH)
Sum of the differences of RDU across the time points
8
6
4
2
0
−0.1
0.0
0.1
LTRENGKTVIRVLKQKEVEQL:216−236
TRENGKTVIRVLKQKEVEQL:217−236
AIGHAGTCL:27−35
IKKHEEEEAKAEREKKEKEQKEKDK:237−261

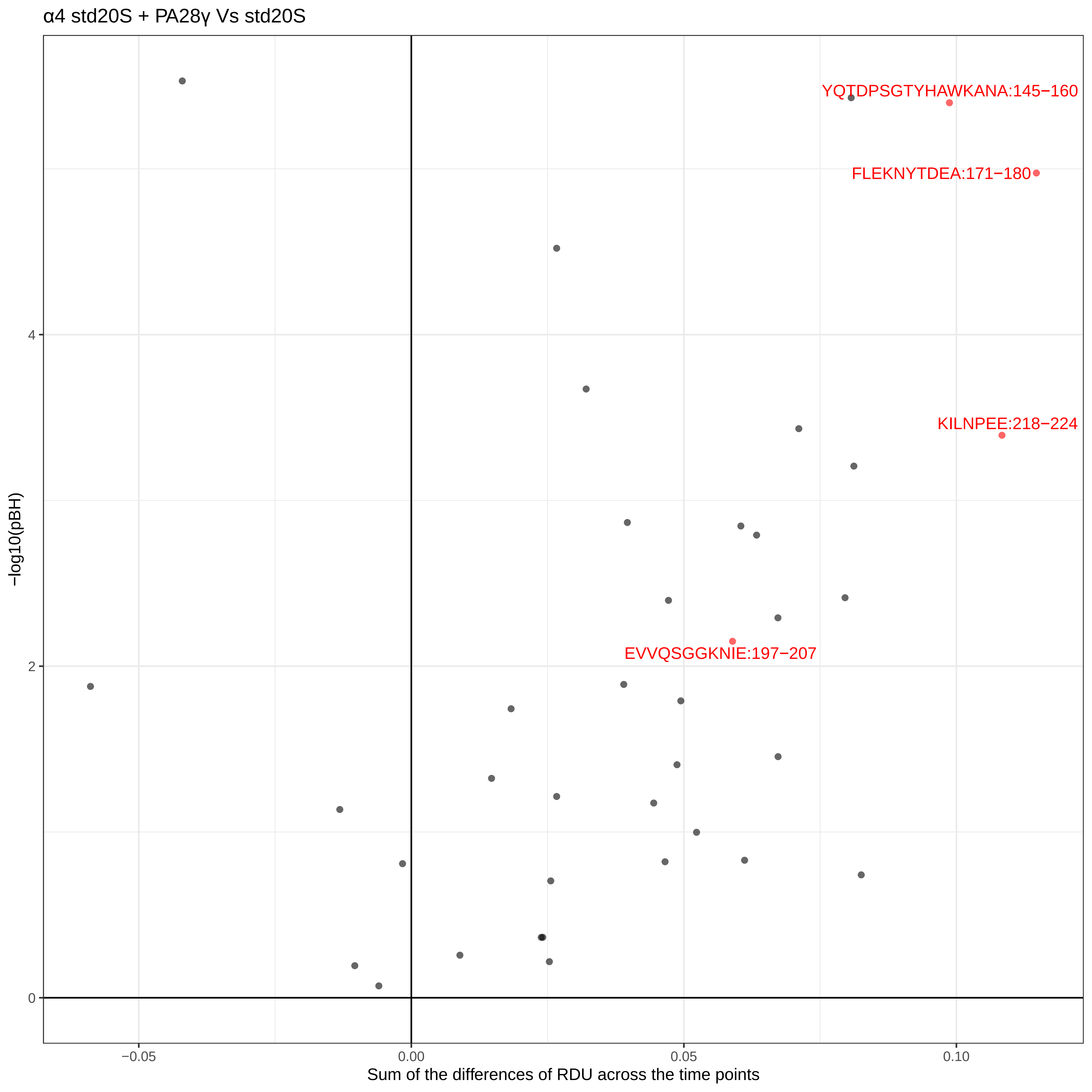

α4 std20S + PA28γ Vs std20S
YQTDPSGTYHAWKANA:145−160
FLEKNYTDEA:171−180
KILNPEE:218−224
EVVQSGGKNIE:197−207
−log10(pBH)
4
2
0
−0.05
0.00
0.05
0.10
Sum of the differences of RDU across the time points

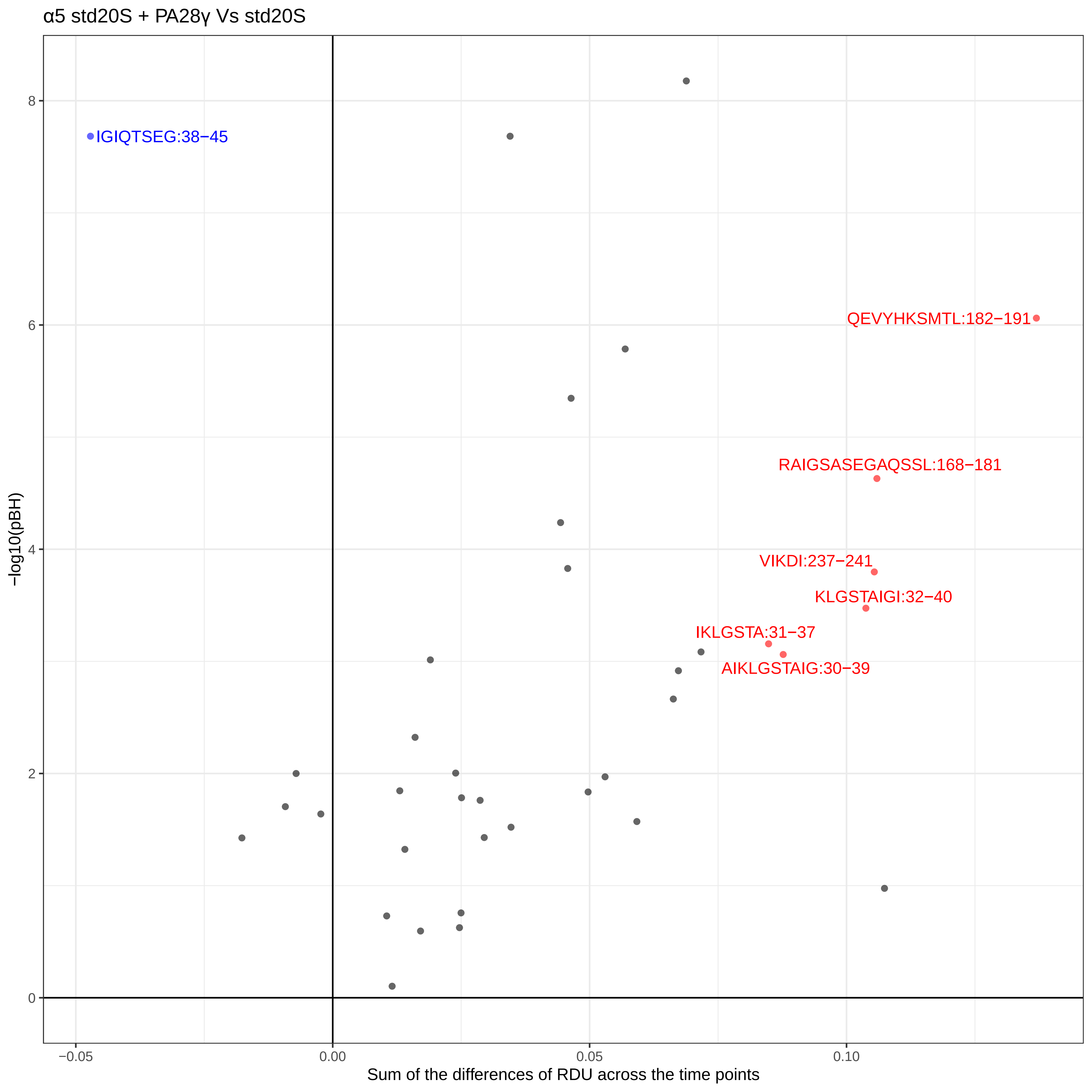

α5 std20S + PA28γ Vs std20S
IGIQTSEG:38−45
QEVYHKSMTL:182−191
RAIGSASEGAQSSL:168−181
VIKDI:237−241
KLGSTAIGI:32−40
IKLGSTA:31−37
AIKLGSTAIG:30−39
−log10(pBH)
0
2
4
6
8
−0.05
0.00
0.05
0.10
Sum of the differences of RDU across the time points

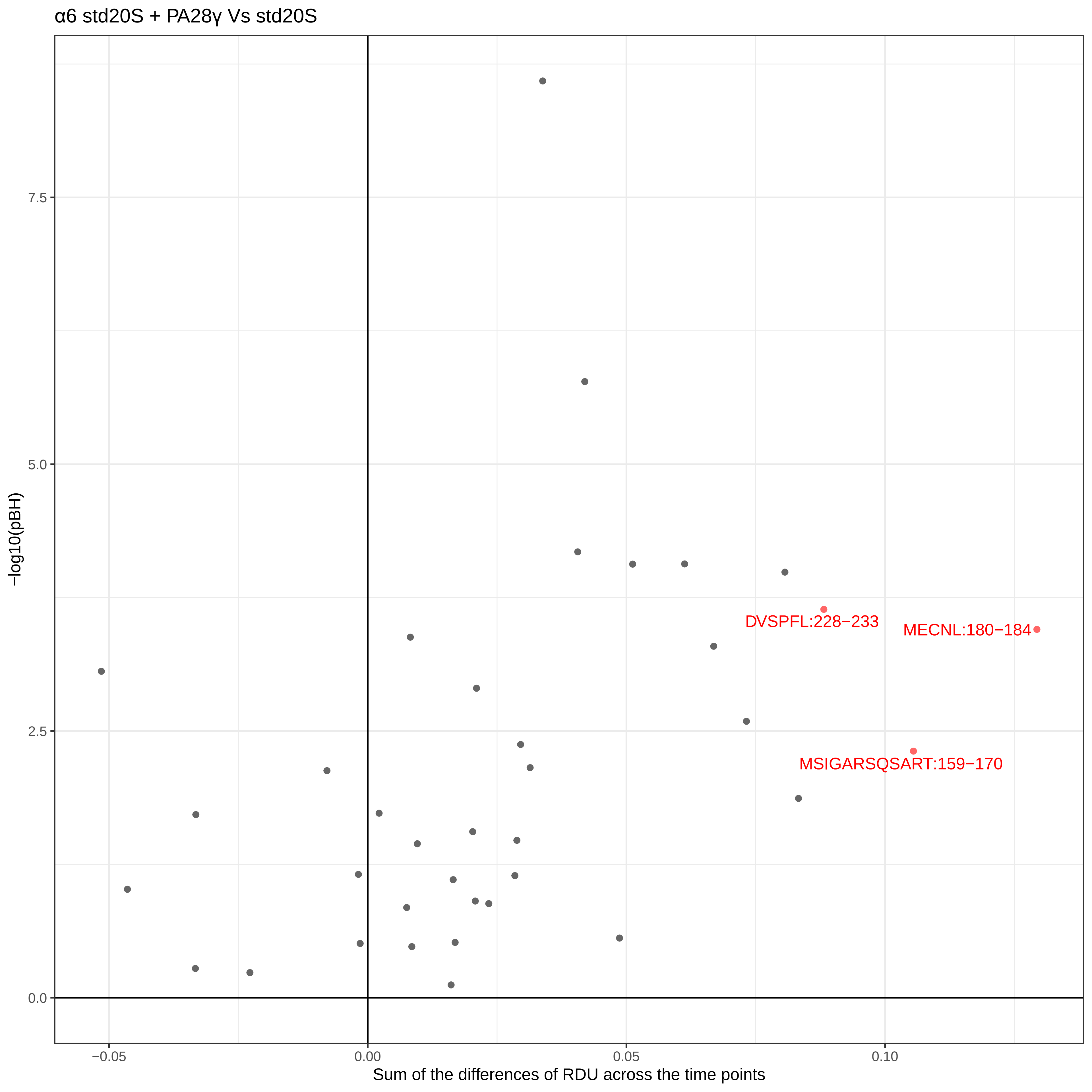
α6 std20S + PA28γ Vs std20S
−log10(pBH)
7.5
5.0
2.5
0.0
−0.05
0.00
0.05
0.10
Sum of the differences of RDU across the time points
DVSPFL:228−233
MECNL:180−184
MSIGARSQSART:159−170

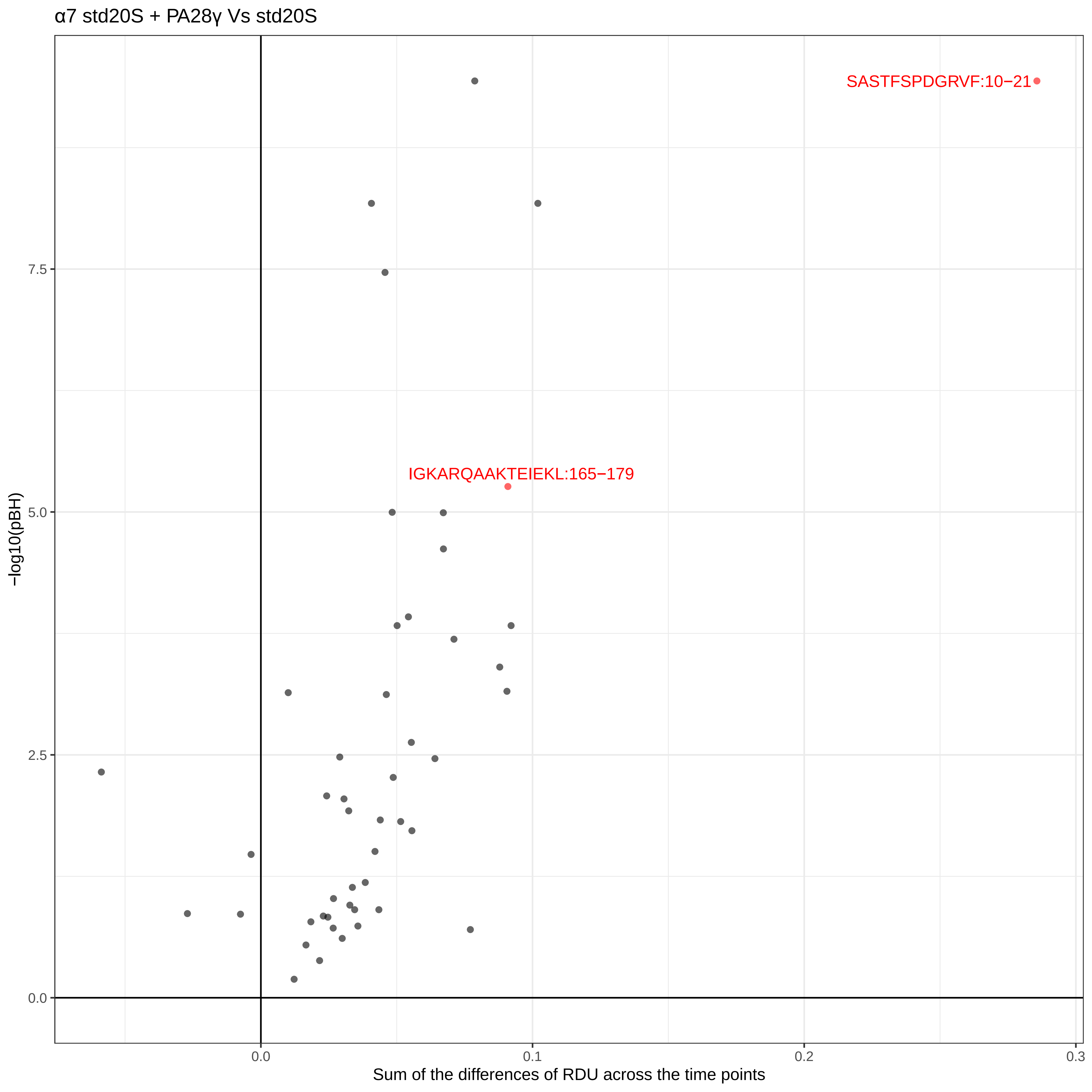
α7 std20S + PA28γ Vs std20S
SASTFSPDGRVF:10−21
IGKARQAAKTEIEKL:165−179
7.5
5.0
2.5
0.0
−log10(pBH)
0.0
0.1
0.2
0.3
Sum of the differences of RDU across the time points

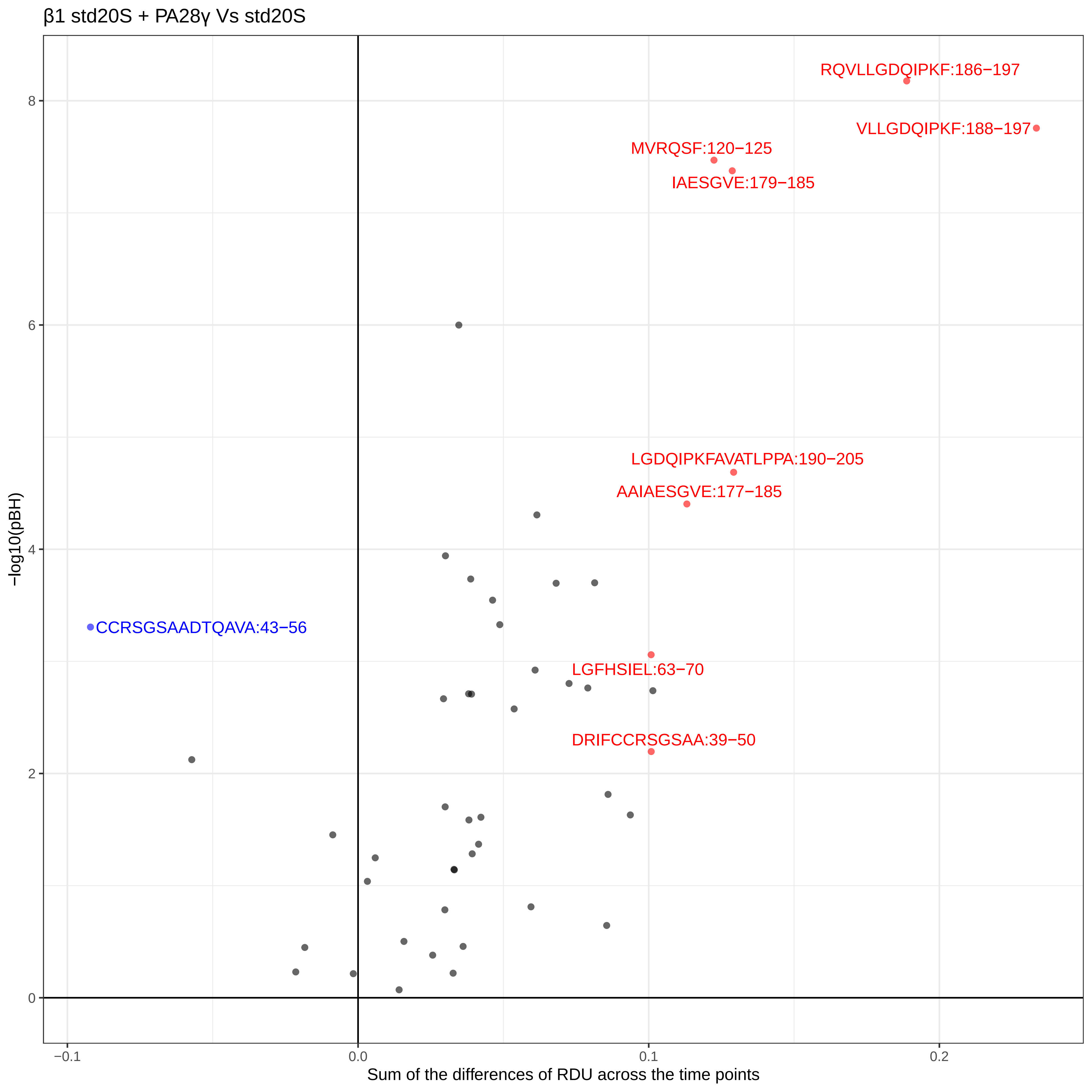

β1 std20S + PA28γ Vs std20S
RQVLLGDQIPKF:186−197
VLLGDQIPKF:188−197
MVRQSF:120−125
IAESGVE:179−185
LGDQIPKFAVATLPPA:190−205
AAIAESGVE:177−185
CCRSGSAADTQAVA:43−56
LGFHSIEL:63−70
DRIFCCRSGSAA:39−50
−log10(pBH)
8
6
4
2
0
−0.1
0.0
0.1
0.2
Sum of the differences of RDU across the time points

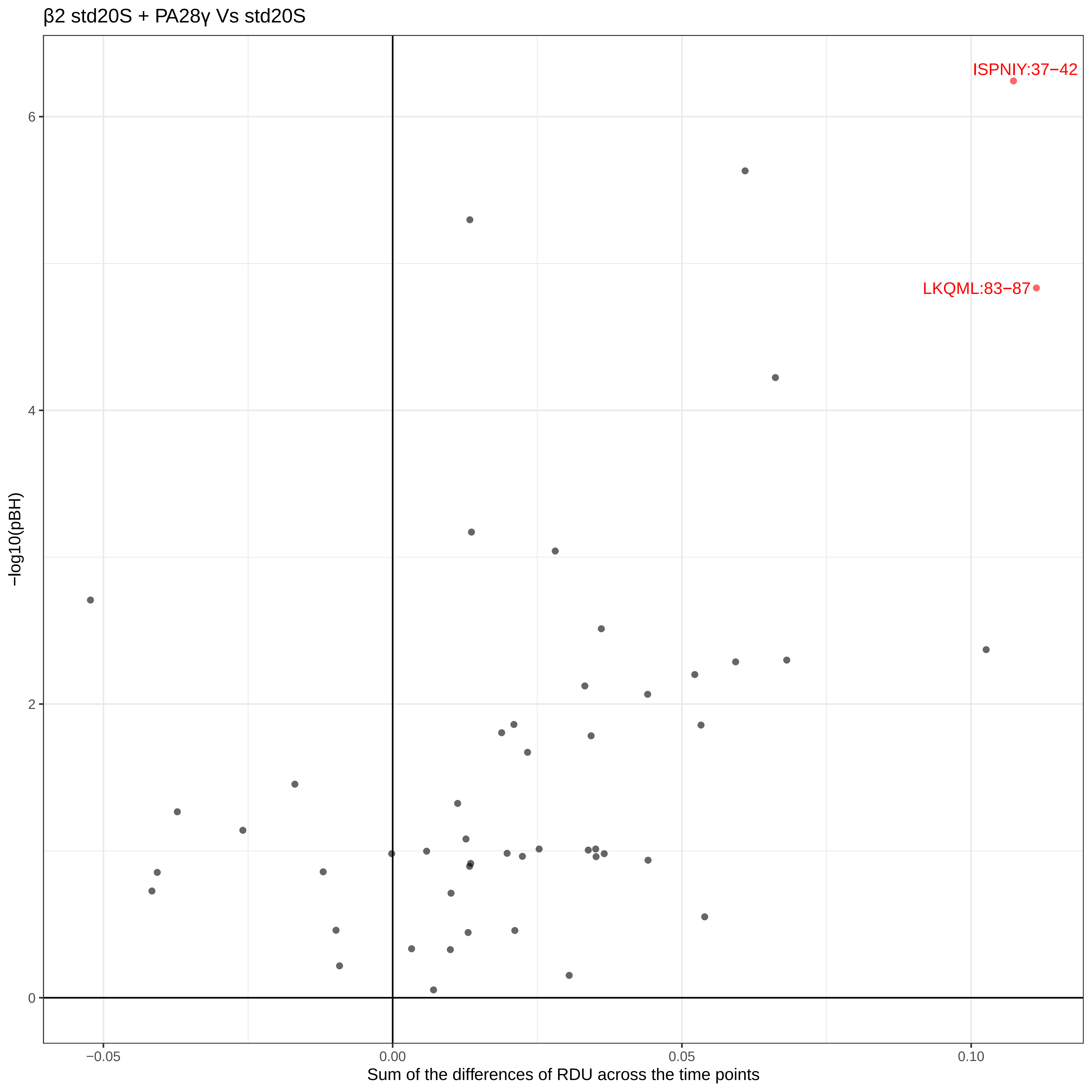
β2 std20S + PA28γ Vs std20S
ISPNIY:37−42
LKQML:83−87
−log10(pBH)
6
4
2
0
−0.05
0.00
0.05
0.10
Sum of the differences of RDU across the time points

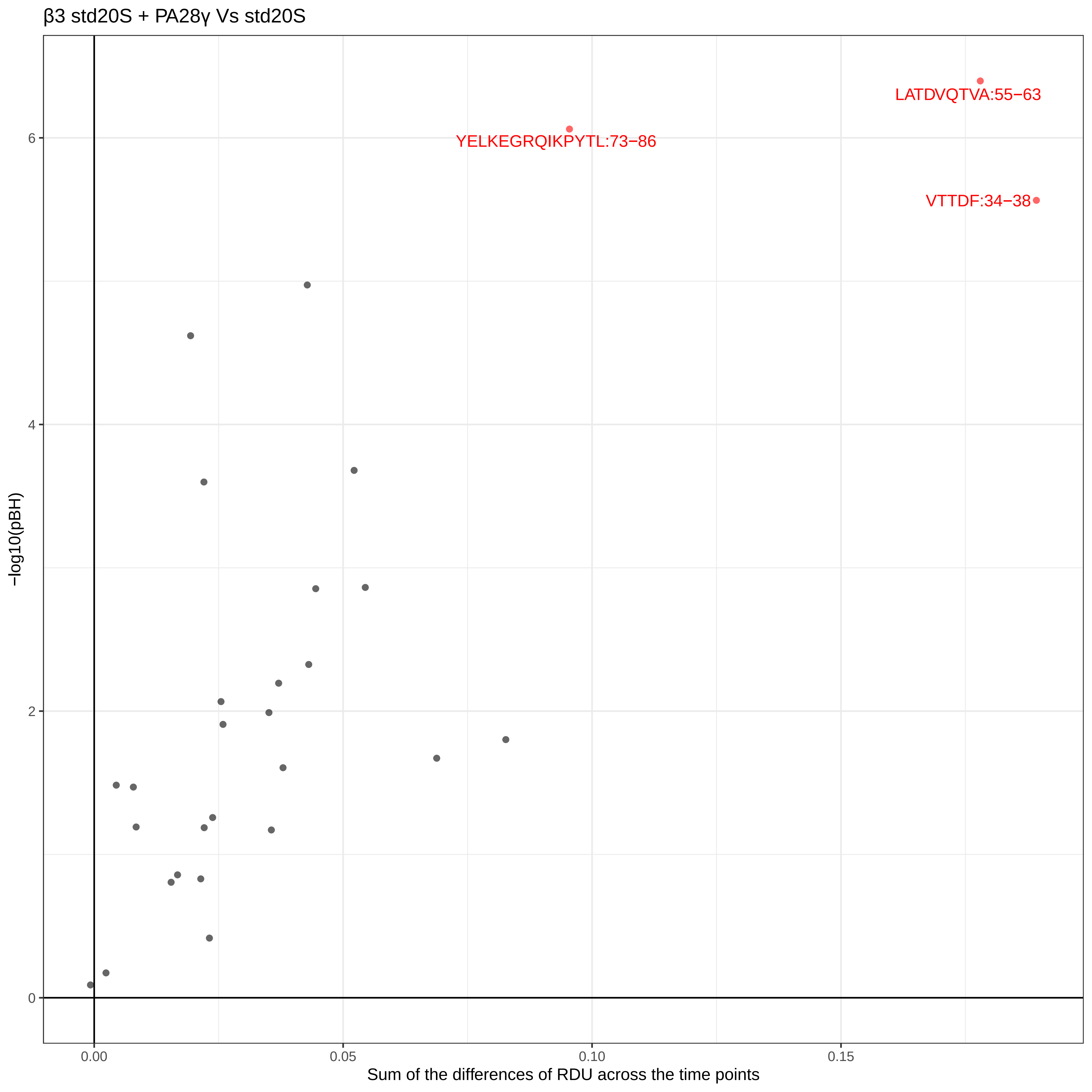

β3 std20S + PA28γ Vs std20S
LATDVQTVA:55−63
YELKEGRQIKPYTL:73−86
VTTDF:34−38
−log10(pBH)
0
2
4
6
0.00
0.05
0.10
0.15
Sum of the differences of RDU across the time points

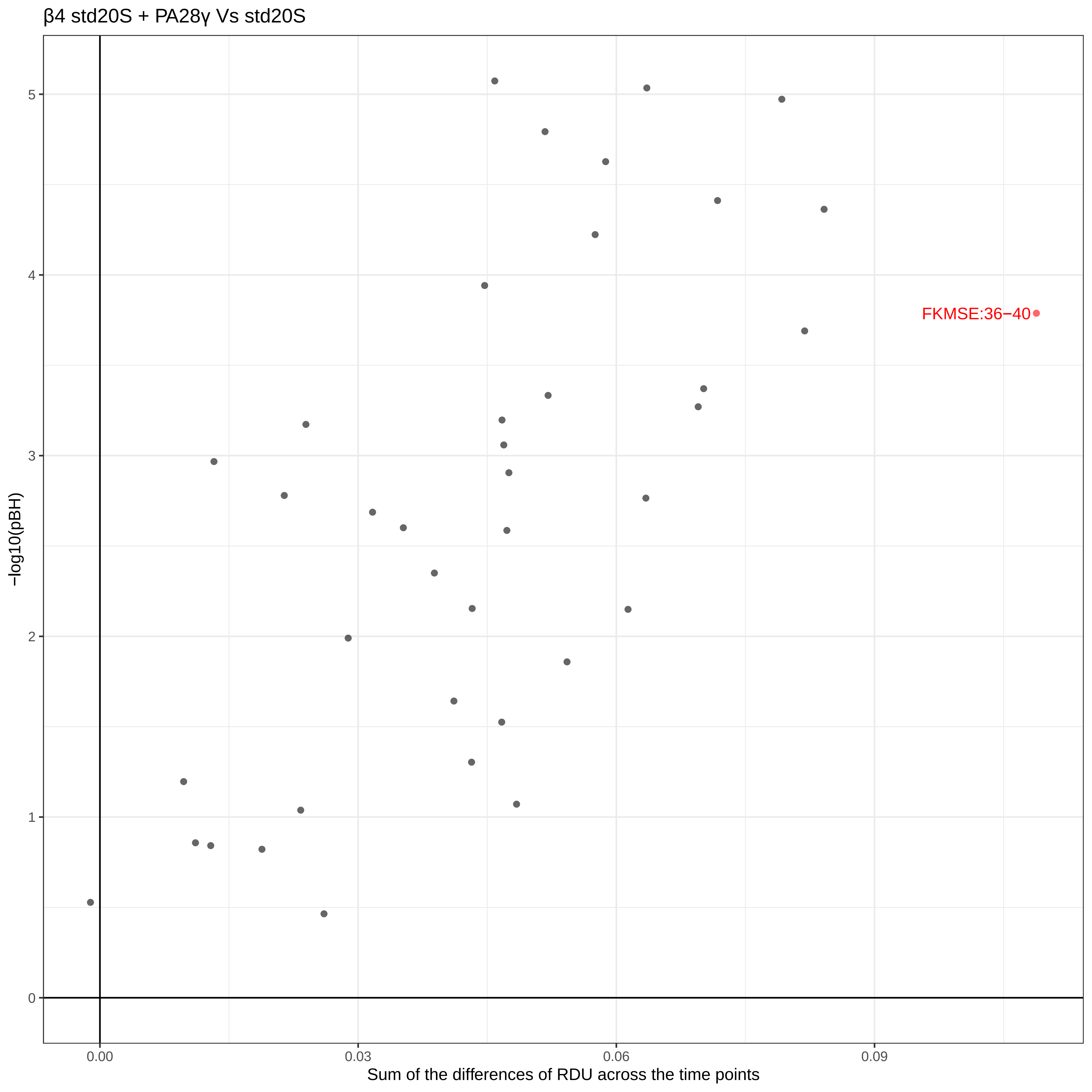

β4 std20S + PA28γ Vs std20S
−log10(pBH)
Sum of the differences of RDU across the time points
FKMSE:36−40
0
1
2
3
4
5
0.00
0.03
0.06
0.09

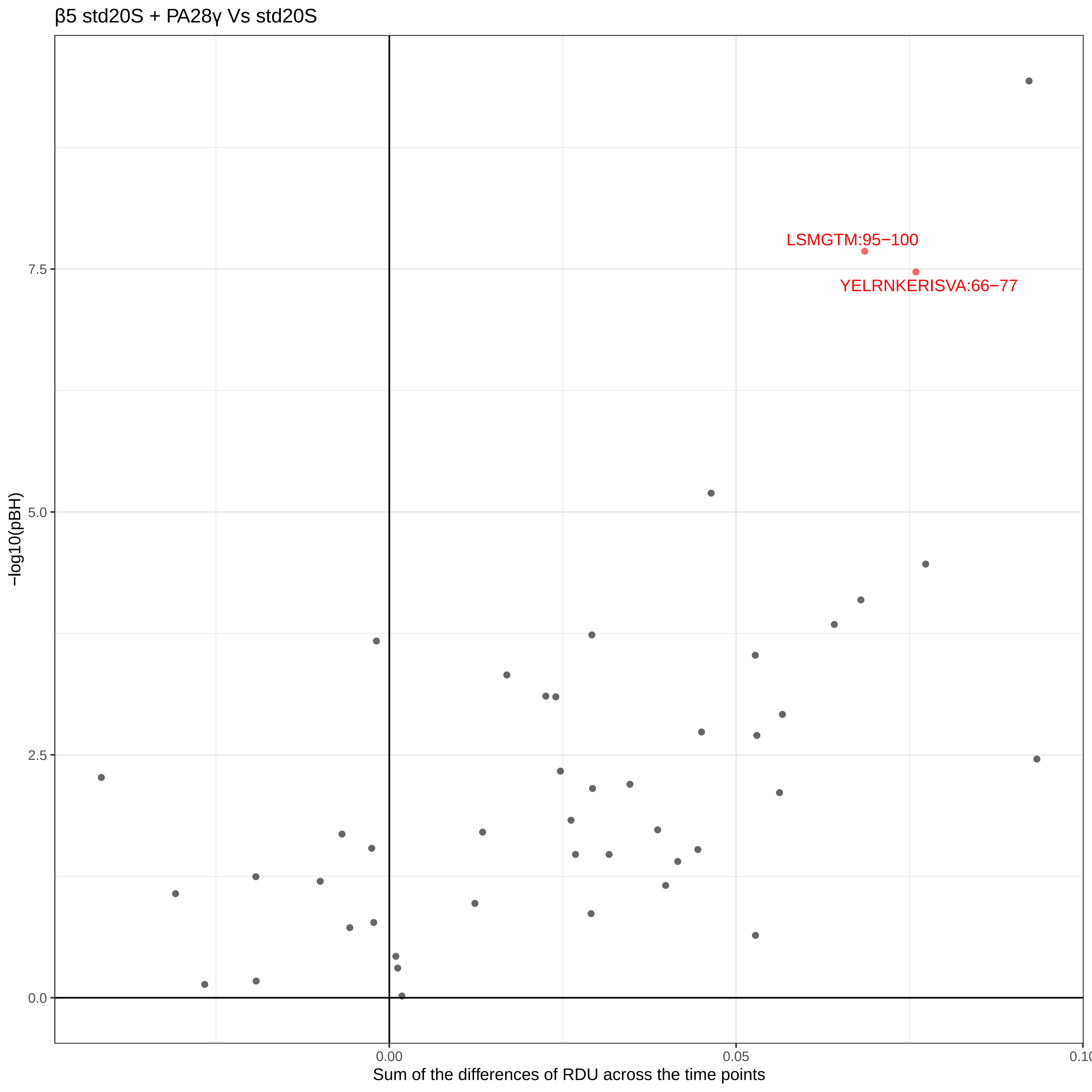

β5 std20S + PA28γ Vs std20S
LSMGTM:95−100
YELRNKERISVA:66−77
7.5
5.0
2.5
0.0
−log10(pBH)
0.00
0.05
0.10
Sum of the differences of RDU across the time points

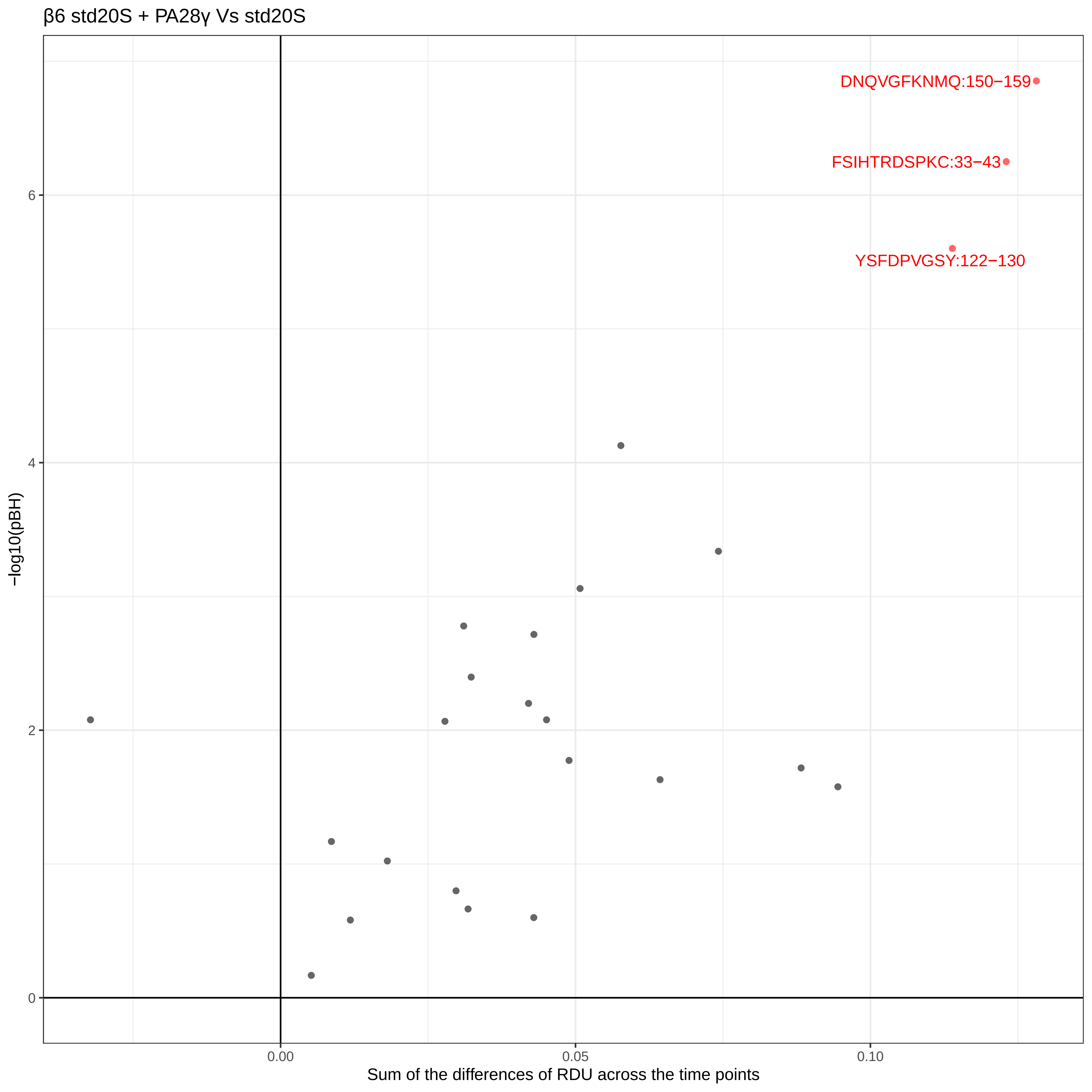

β6 std20S + PA28γ Vs std20S
DNQVGFKNMQ:150−159
FSIHTRDSPKC:33−43
YSFDPVGSY:122−130
−log10(pBH)
0
2
4
6
0.00
0.05
0.10
Sum of the differences of RDU across the time points

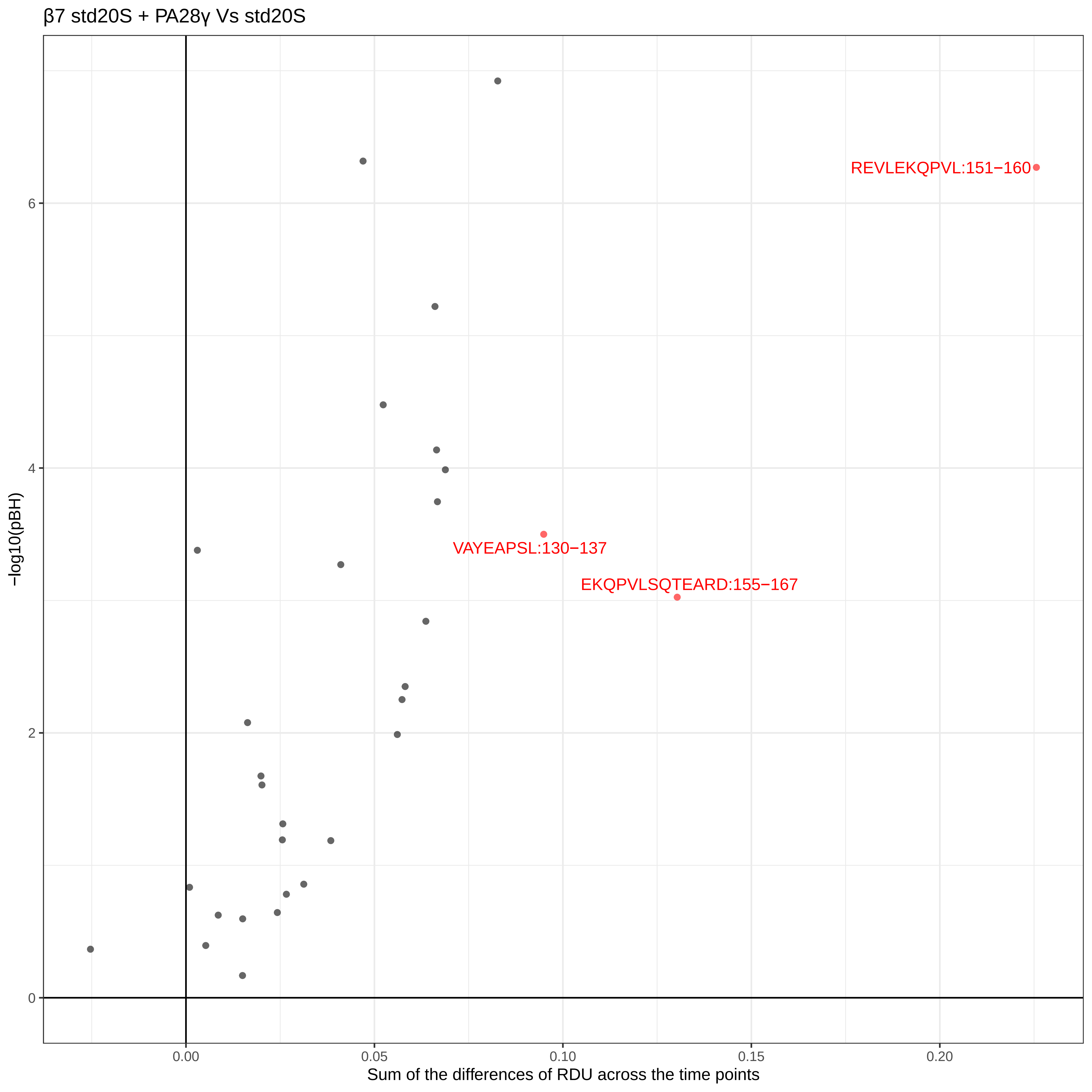

β7 std20S + PA28γ Vs std20S
REVLEKQPVL:151−160
VAYEAPSL:130−137
EKQPVLSQTEARD:155−167
−log10(pBH)
0
2
4
6
0.00
0.05
0.10
0.15
0.20
Sum of the differences of RDU across the time points

α1 i20S + PA28αβ Vs i20S

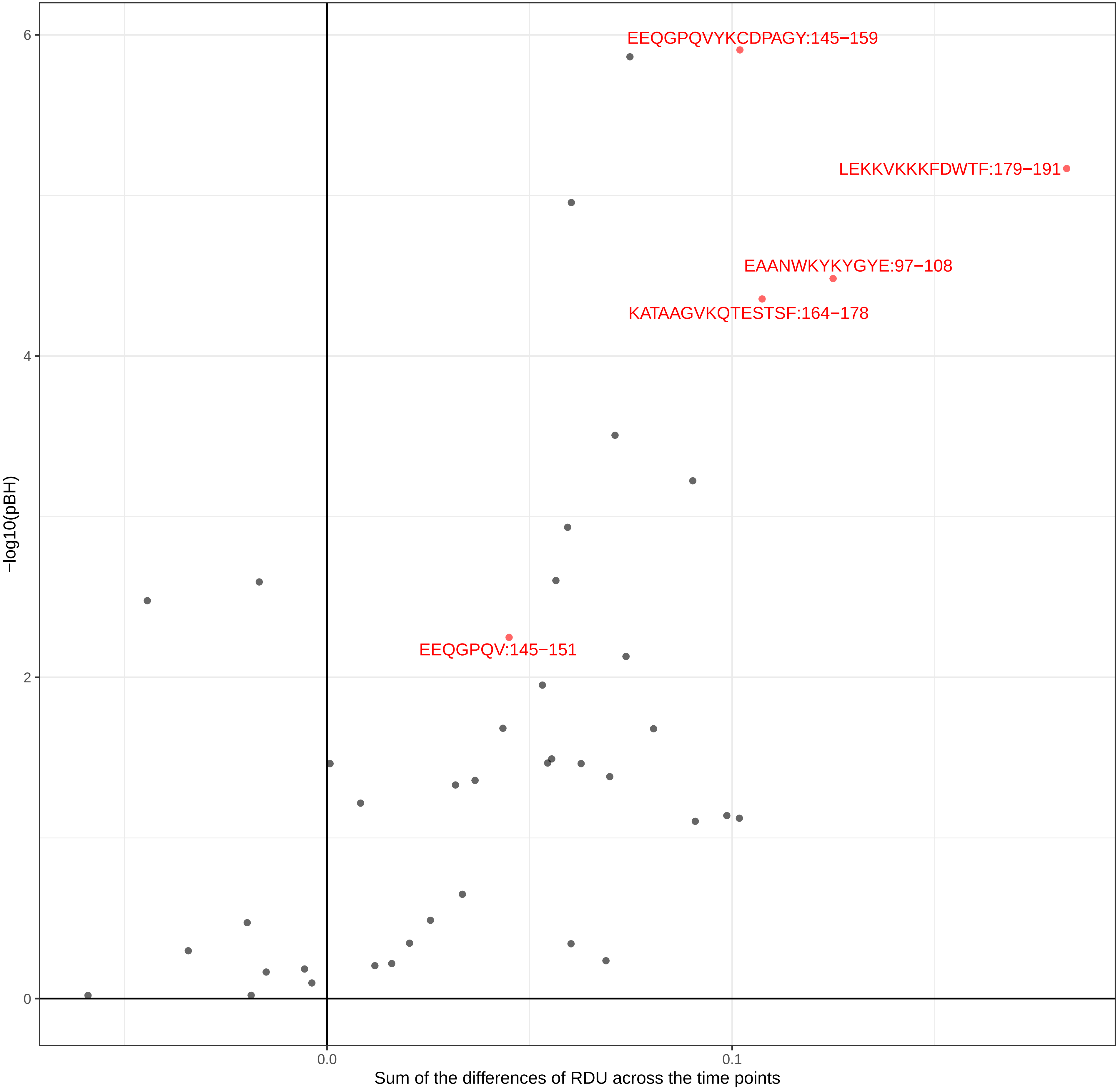

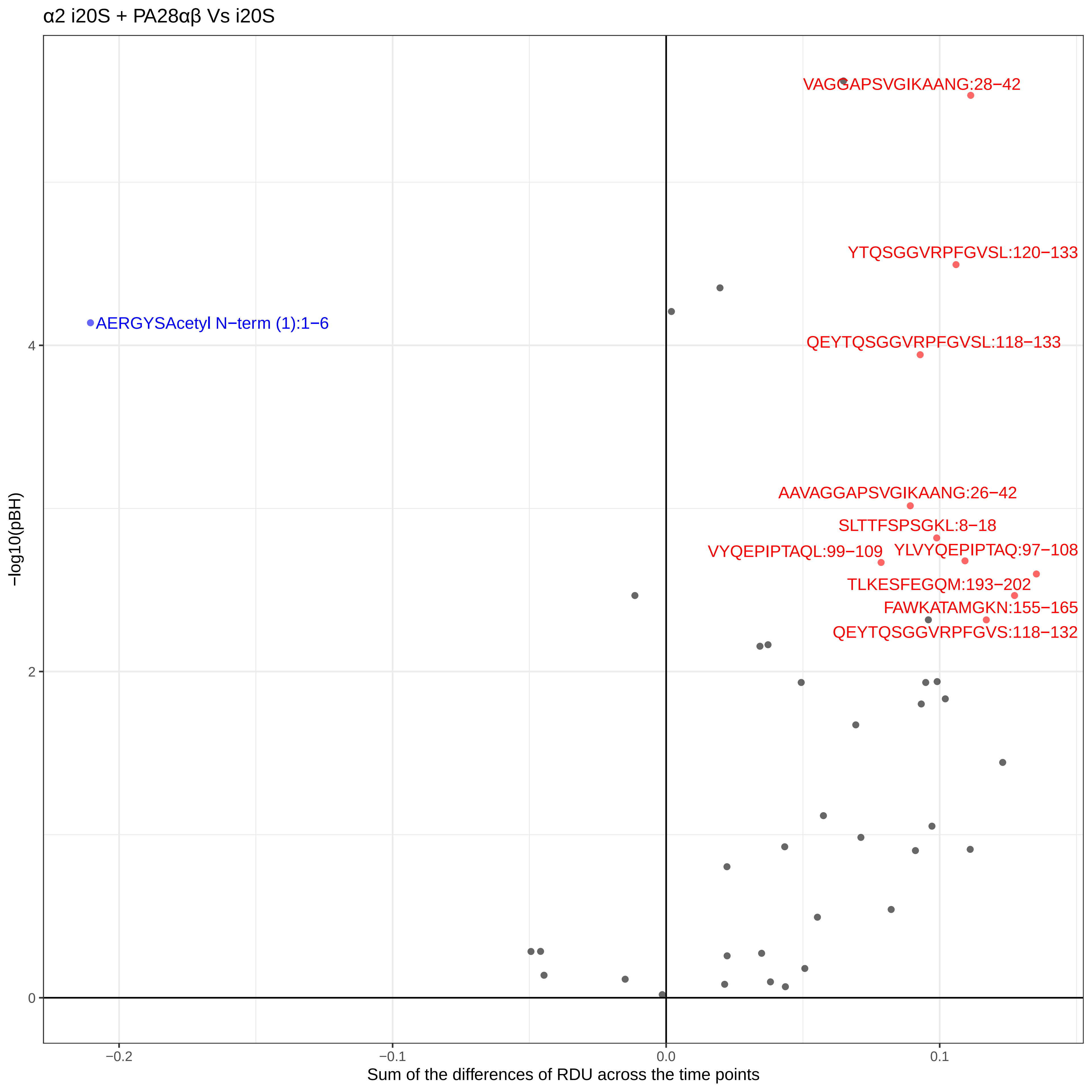
α2 i20S + PA28αβ Vs i20S
VAGGAPSVGIKAANG:28−42
YTQSGGVRPFGVSL:120−133
AERGYSAcetyl N−term (1):1−6
QEYTQSGGVRPFGVSL:118−133
AAVAGGAPSVGIKAANG:26−42
SLTTFSPSGKL:8−18
VYQEPIPTAQL:99−109
YLVYQEPIPTAQ:97−108
TLKESFEGQM:193−202
FAWKATAMGKN:155−165
QEYTQSGGVRPFGVS:118−132
−log10(pBH)
4
2
0
−0.2
−0.1
0.0
0.1
Sum of the differences of RDU across the time points

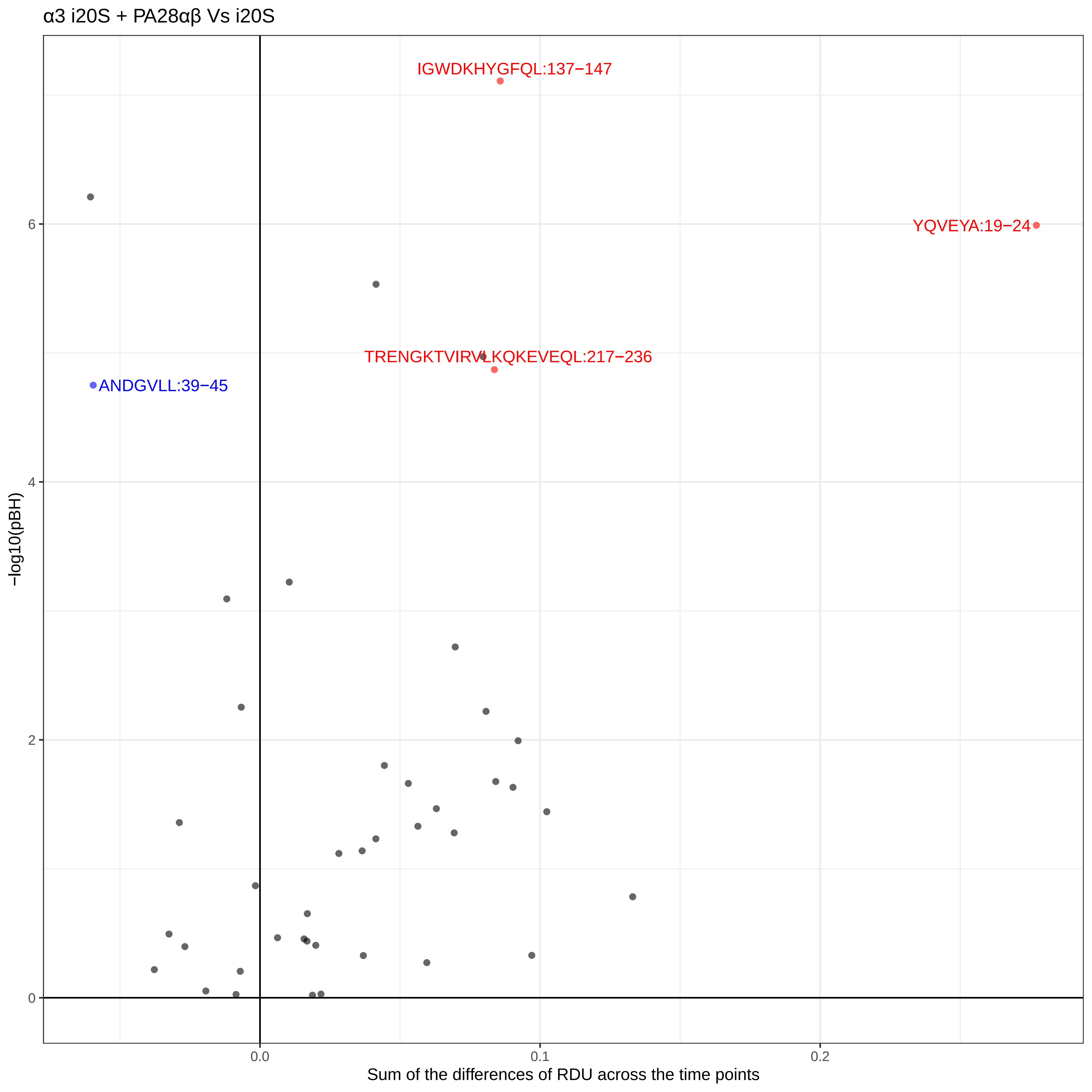

α3 i20S + PA28αβ Vs i20S
IGWDKHYGFQL:137−147
YQVEYA:19−24
TRENGKTVIRVLKQKEVEQL:217−236
ANDGVLL:39−45
−log10(pBH)
0
2
4
6
0.0
0.1
0.2
Sum of the differences of RDU across the time points

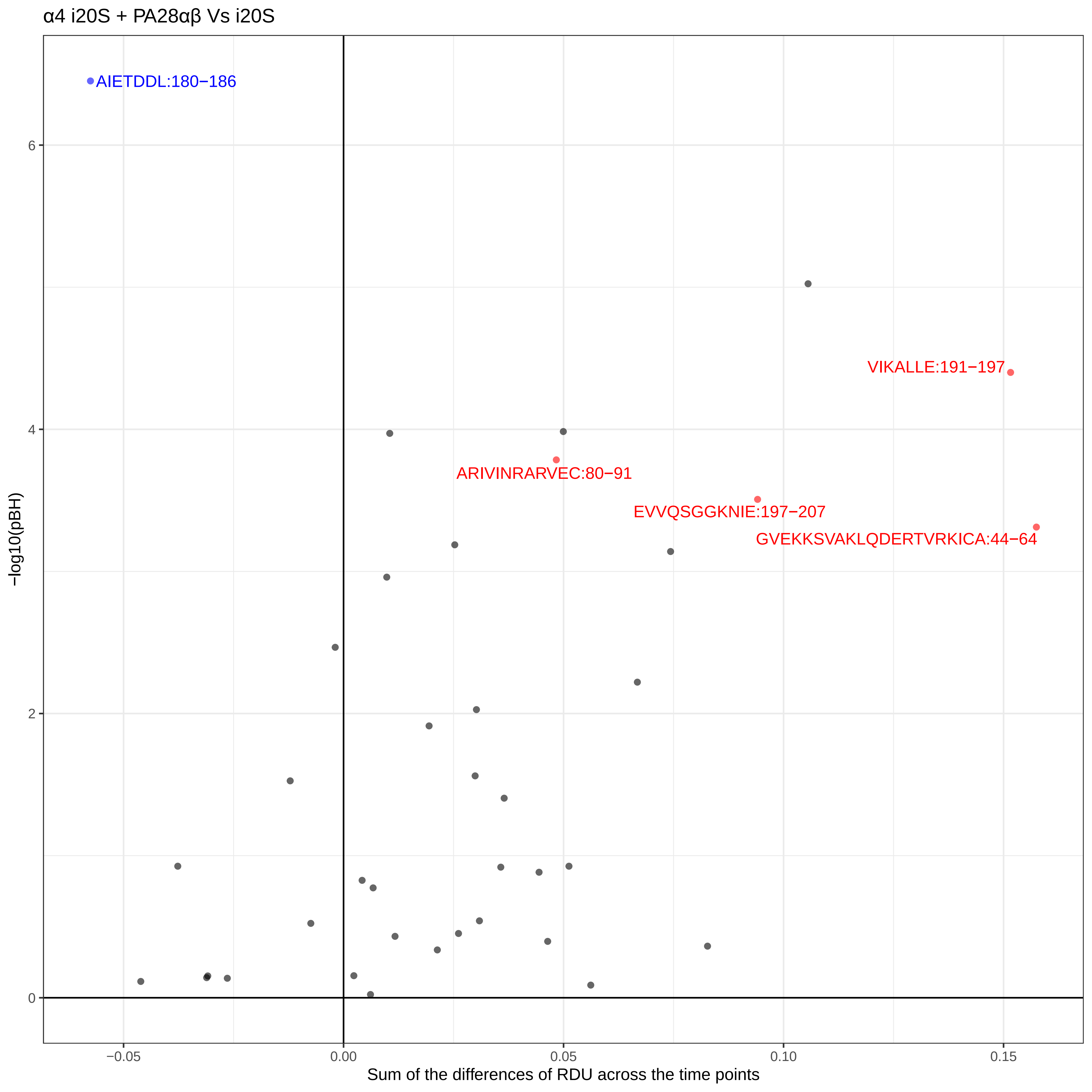

α4 i20S + PA28αβ Vs i20S
AIETDDL:180−186
VIKALLE:191−197
ARIVINRARVEC:80−91
EVVQSGGKNIE:197−207
GVEKKSVAKLQDERTVRKICA:44−64
−log10(pBH)
0
2
4
6
−0.05
0.00
0.05
0.10
0.15
Sum of the differences of RDU across the time points

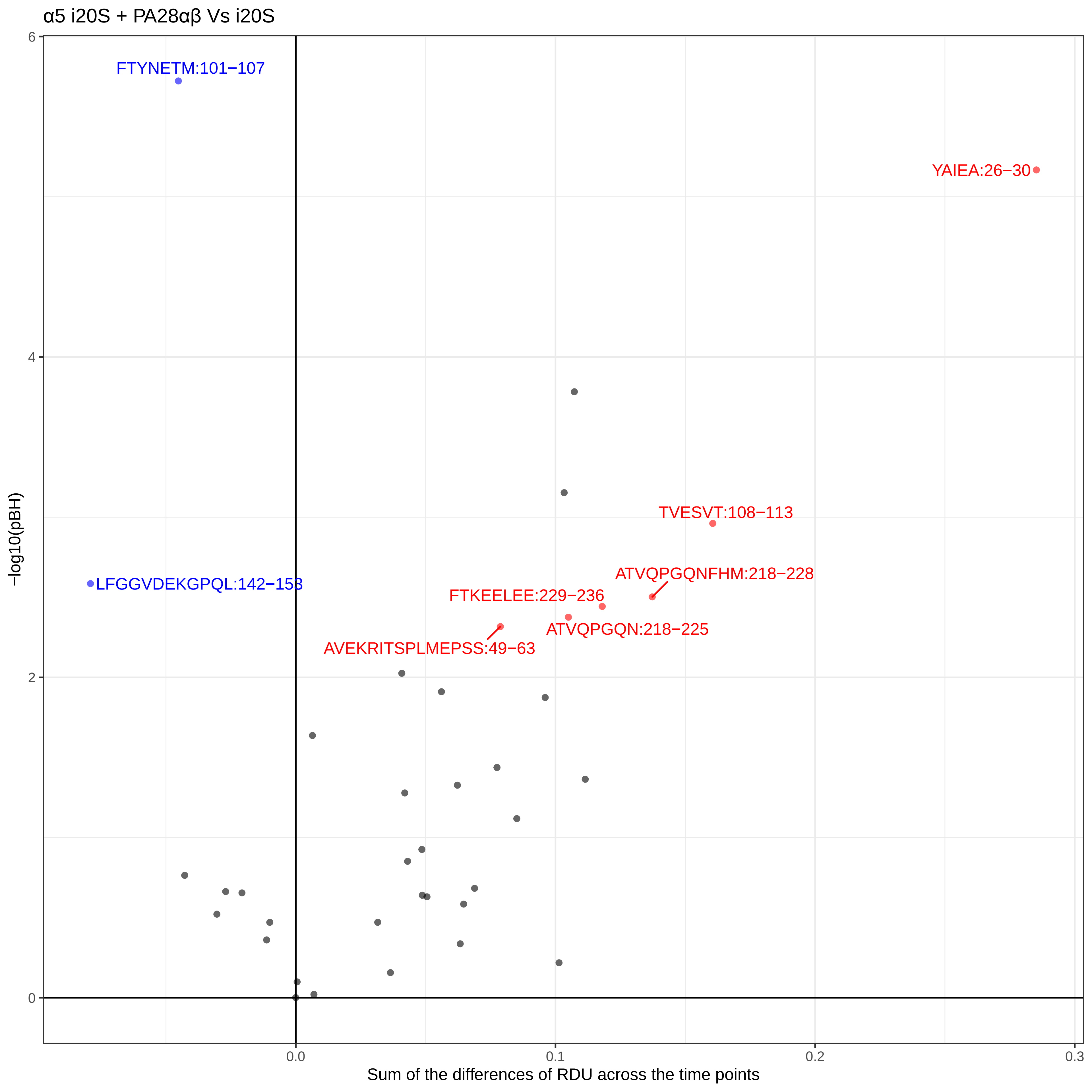
α5 i20S + PA28αβ Vs i20S
FTYNETM:101−107
YAIEA:26−30
TVESVT:108−113
ATVQPGQNFHM:218−228
LFGGVDEKGPQL:142−153
FTKEELEE:229−236
ATVQPGQN:218−225
AVEKRITSPLMEPSS:49−63
−log10(pBH)
Sum of the differences of RDU across the time points
0
2
4
6
0.0
0.1
0.2
0.3

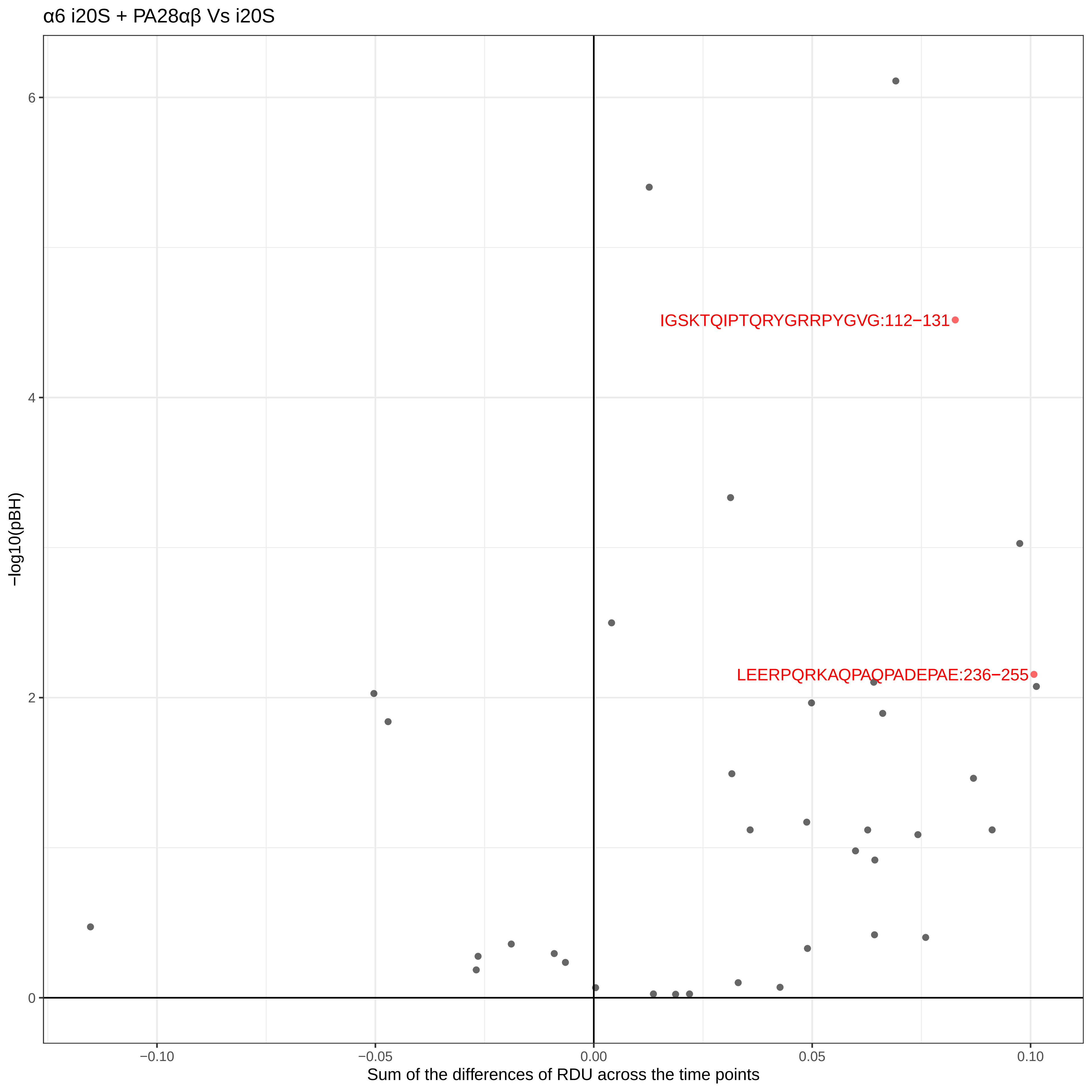
α6 i20S + PA28αβ Vs i20S
IGSKTQIPTQRYGRRPYGVG:112−131
LEERPQRKAQPAQPADEPAE:236−255
−log10(pBH)
6
4
2
0
−0.10
−0.05
0.00
0.05
0.10
Sum of the differences of RDU across the time points

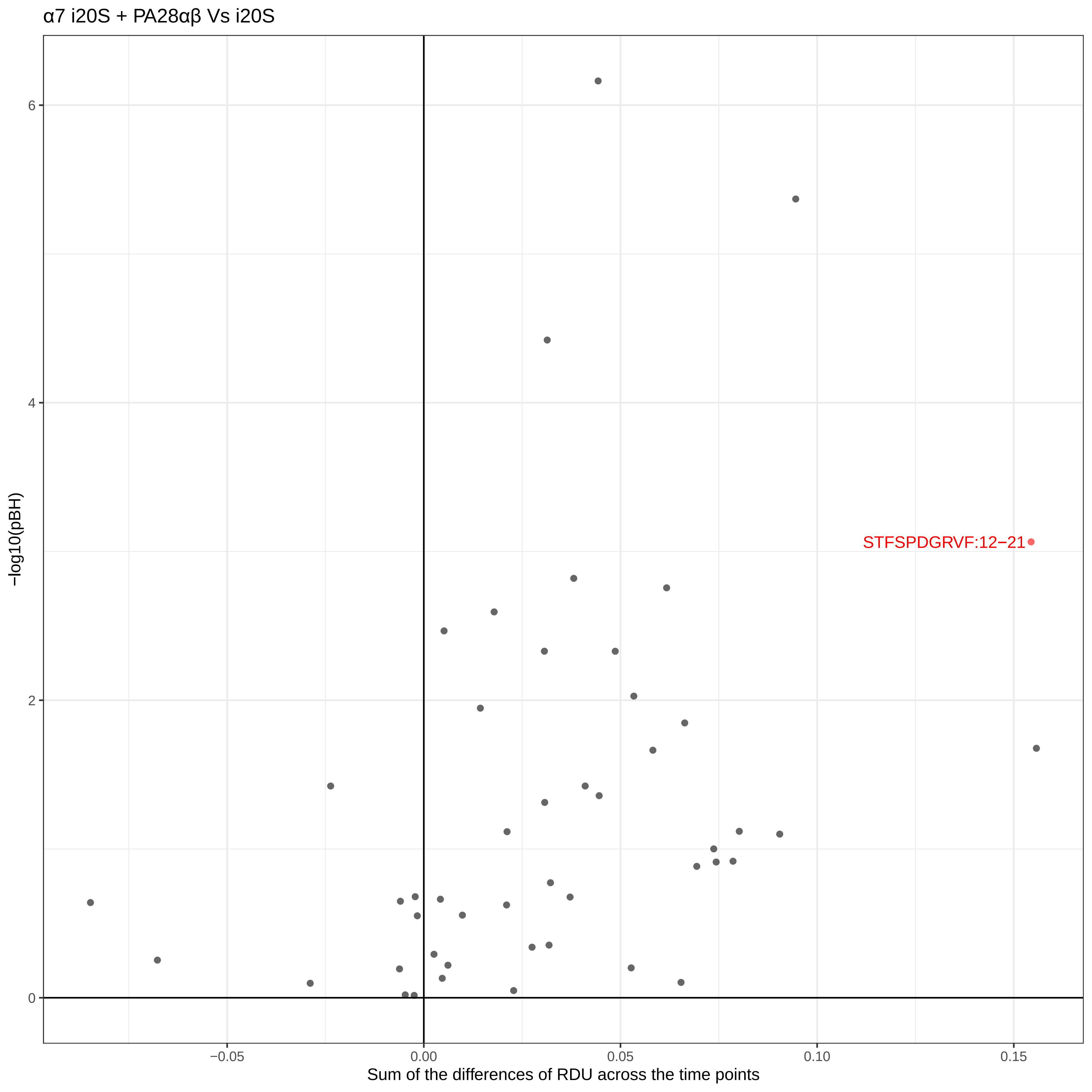

α7 i20S + PA28αβ Vs i20S
STFSPDGRVF:12−21
−log10(pBH)
6
4
2
0
−0.05
0.00
0.05
0.10
0.15
Sum of the differences of RDU across the time points

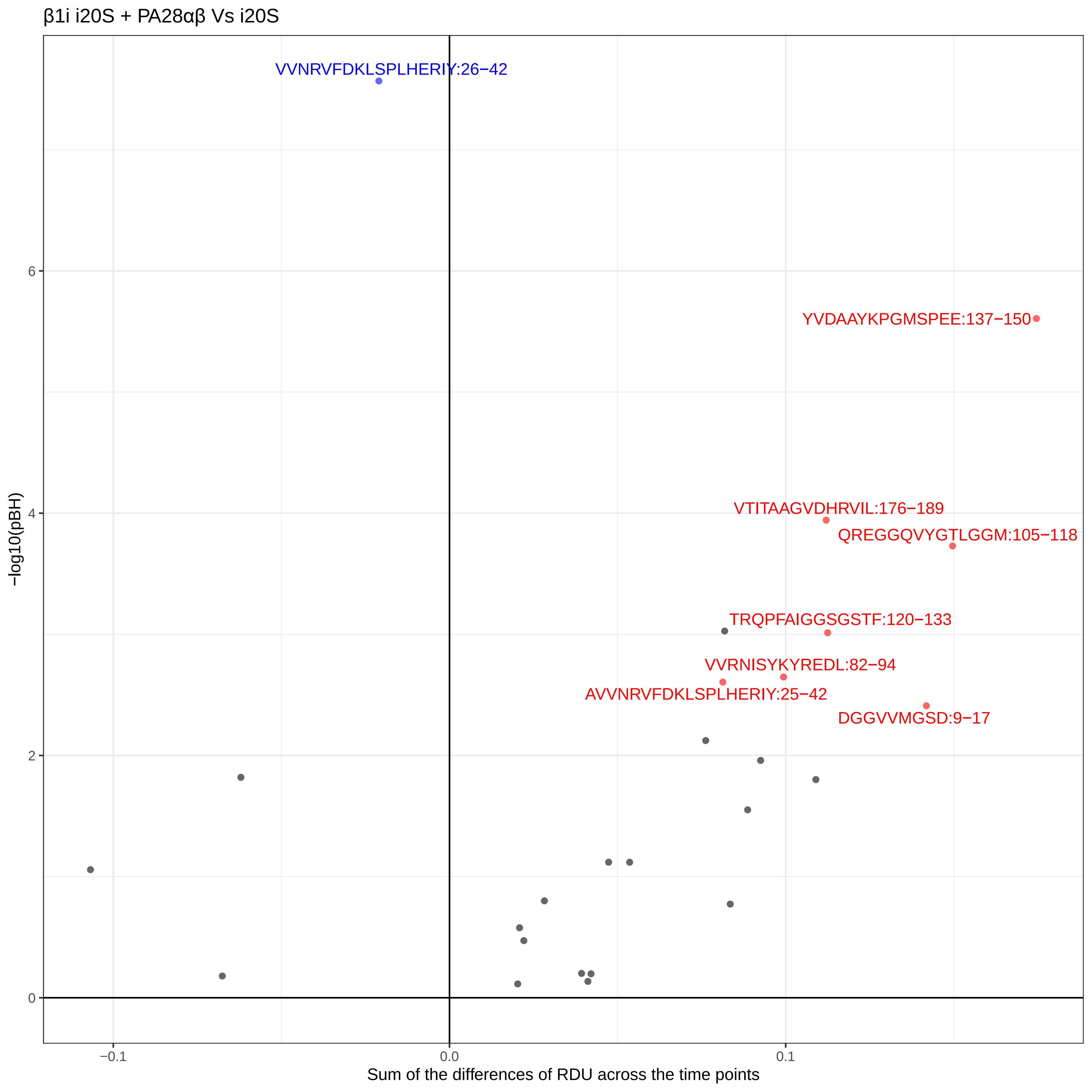
β1i i20S + PA28αβ Vs i20S
VVNRVFDKLSPLHERIY:26−42
YVDAAYKPGMSPEE:137−150
VTITAAGVDHRVIL:176−189
QREGGQVYGTLGGM:105−118
TRQPFAIGGSGSTF:120−133
VVRNISYKYREDL:82−94
AVVNRVFDKLSPLHERIY:25−42
DGGVVMGSD:9−17
−log10(pBH)
6
4
2
0
−0.1
0.0
0.1
Sum of the differences of RDU across the time points

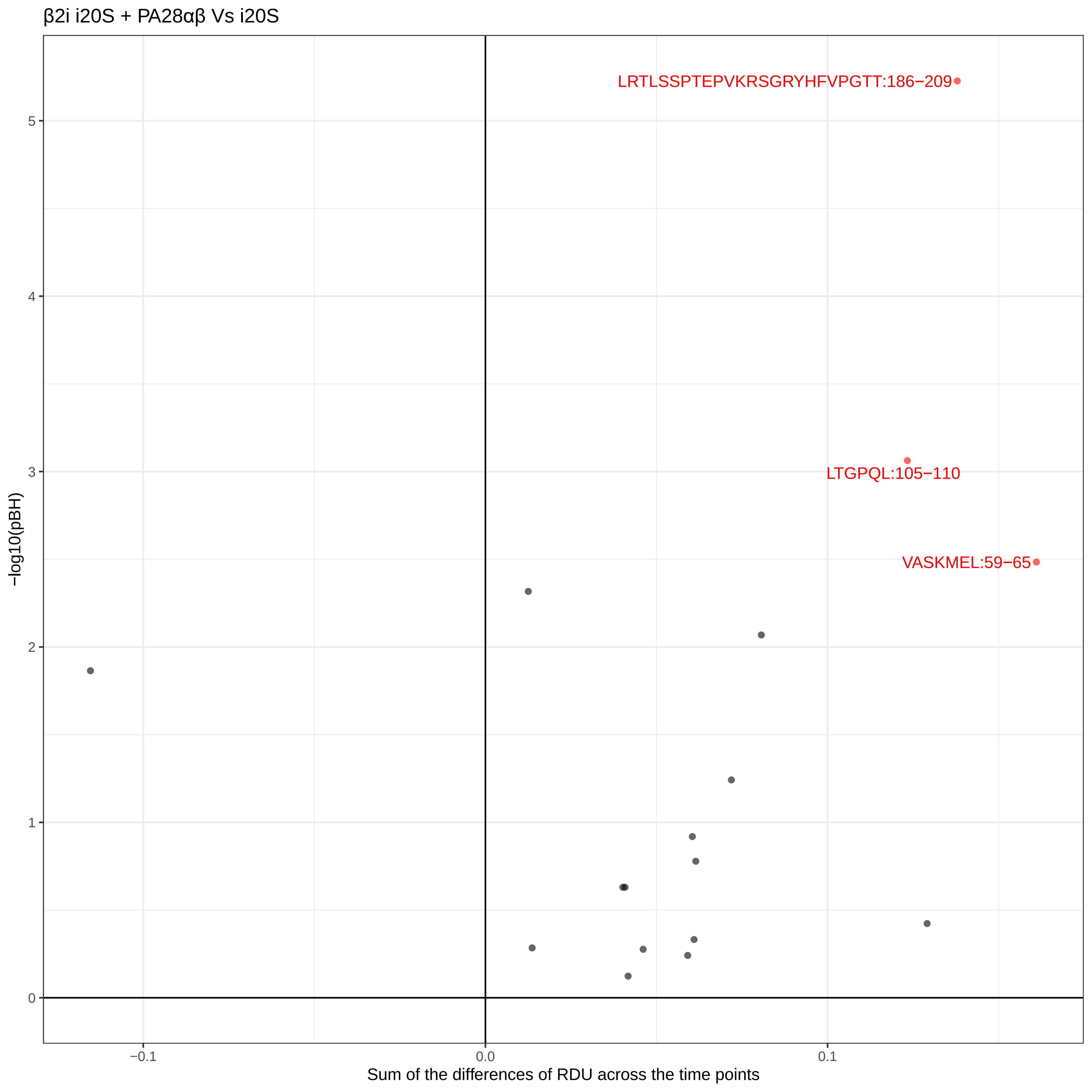
β2i i20S + PA28αβ Vs i20S
LRTLSSPTEPVKRSGRYHFVPGTT:186−209
LTGPQL:105−110
VASKMEL:59−65
−log10(pBH)
0
1
2
3
4
5
−0.1
0.0
0.1
Sum of the differences of RDU across the time points

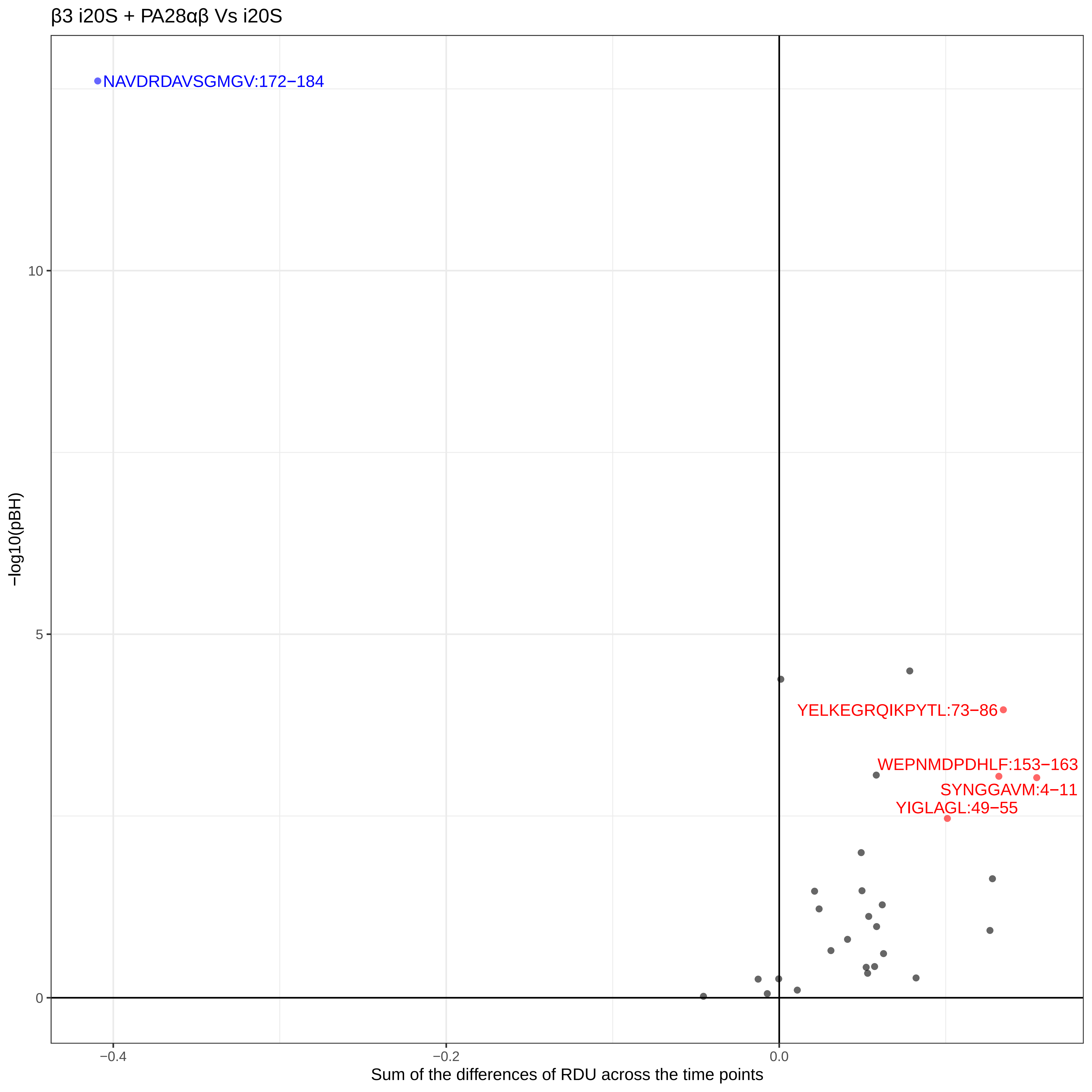
β3 i20S + PA28αβ Vs i20S
NAVDRDAVSGMGV:172−184
YELKEGRQIKPYTL:73−86
WEPNMDPDHLF:153−163
SYNGGAVM:4−11
YIGLAGL:49−55
10
5
0
−log10(pBH)
−0.4
−0.2
0.0
Sum of the differences of RDU across the time points

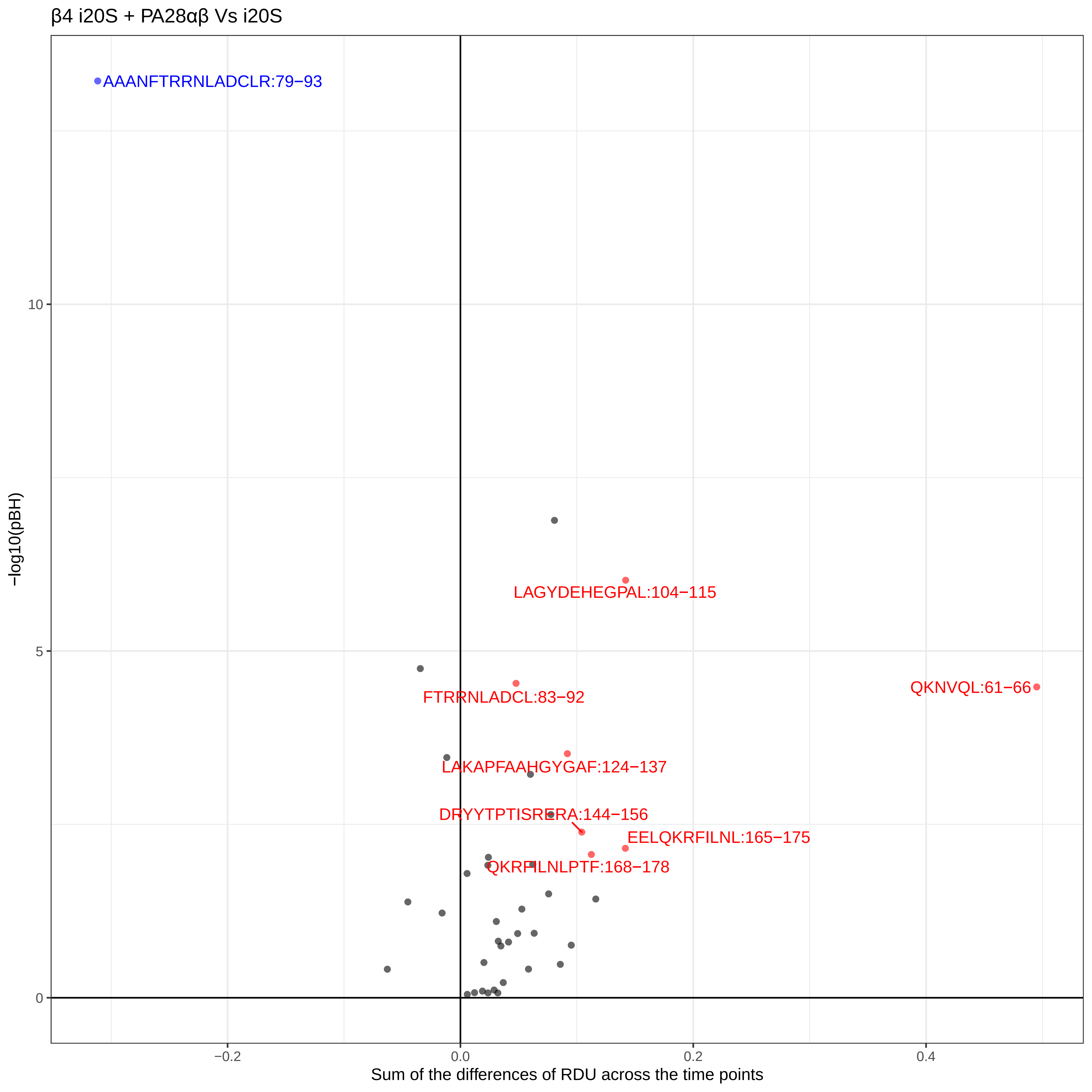
β4 i20S + PA28αβ Vs i20S
AAANFTRRNLADCLR:79−93
LAGYDEHEGPAL:104−115
FTRRNLADCL:83−92
QKNVQL:61−66
LAKAPFAAHGYGAF:124−137
DRYYTPTISRERA:144−156
EELQKRFILNL:165−175
QKRFILNLPTF:168−178
10
5
0
−0.2
0.0
0.2
0.4
−log10(pBH)
Sum of the differences of RDU across the time points

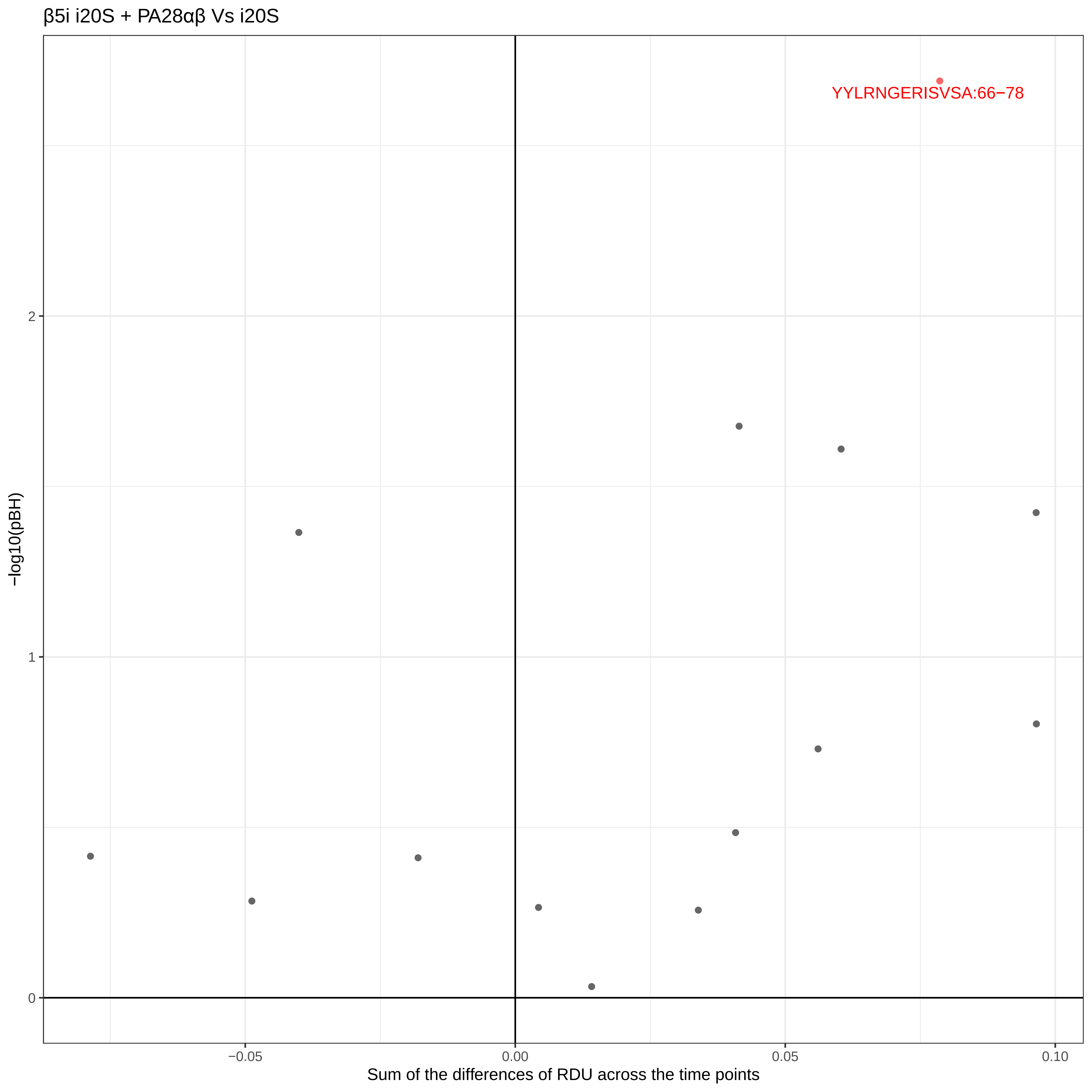
β5i i20S + PA28αβ Vs i20S
YYLRNGERISVSA:66−78
−log10(pBH)
2
1
0
−0.05
0.00
0.05
0.10
Sum of the differences of RDU across the time points

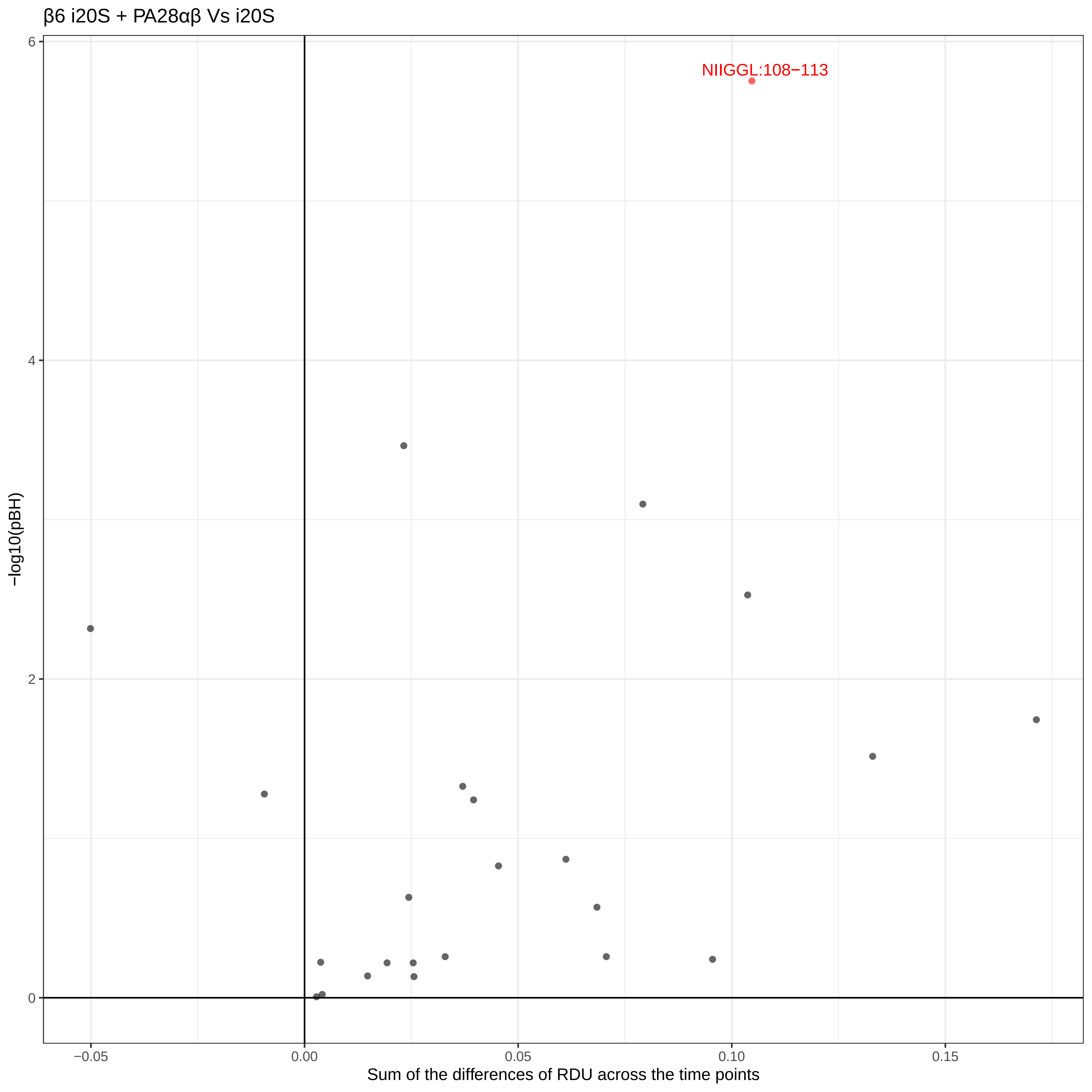
β6 i20S + PA28αβ Vs i20S
NIIGGL:108−113
−log10(pBH)
6
4
2
0
−0.05
0.00
0.05
0.10
0.15
Sum of the differences of RDU across the time points

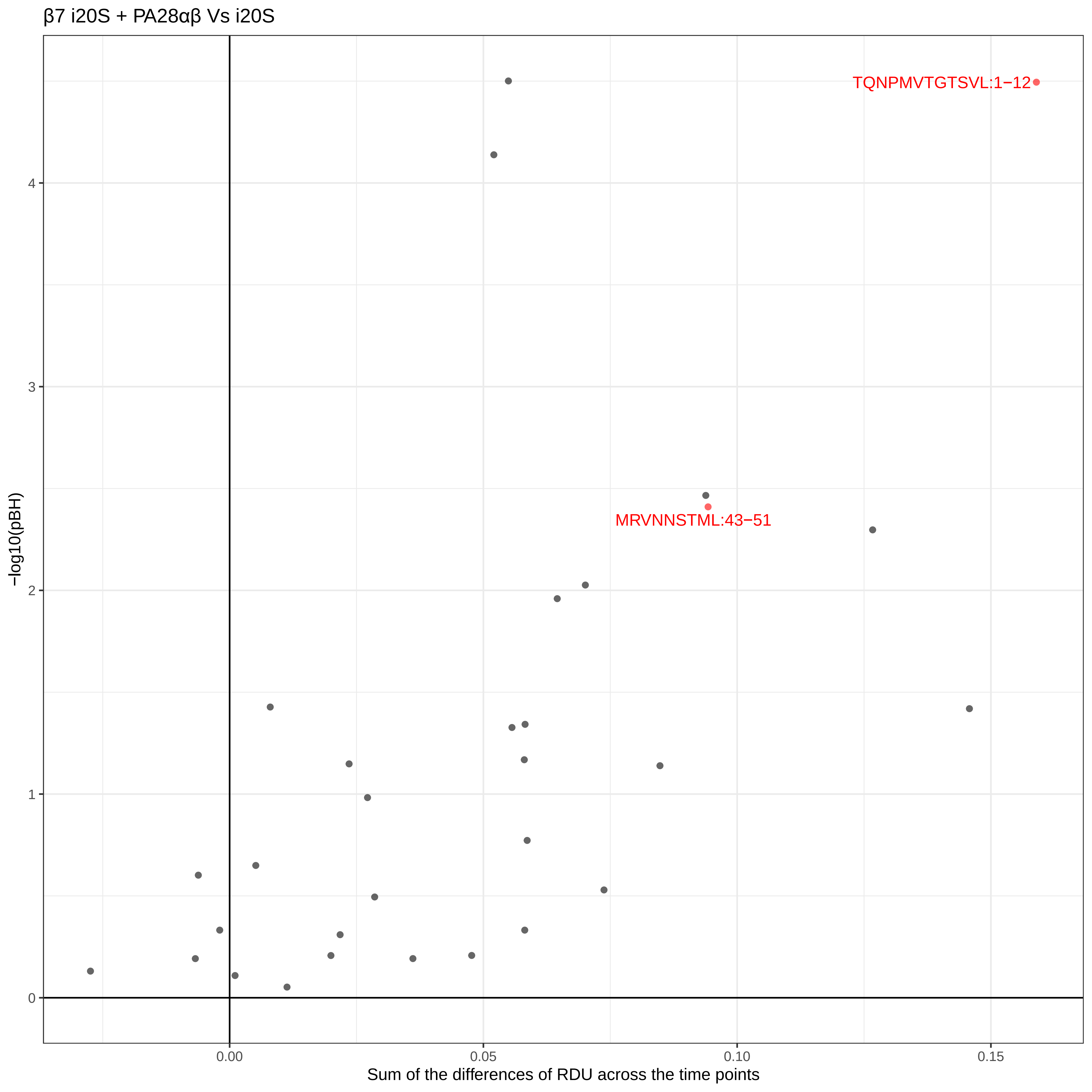
β7 i20S + PA28αβ Vs i20S
TQNPMVTGTSVL:1−12
MRVNNSTML:43−51
−log10(pBH)
0
1
2
3
4
0.00
0.05
0.10
0.15
Sum of the differences of RDU across the time points

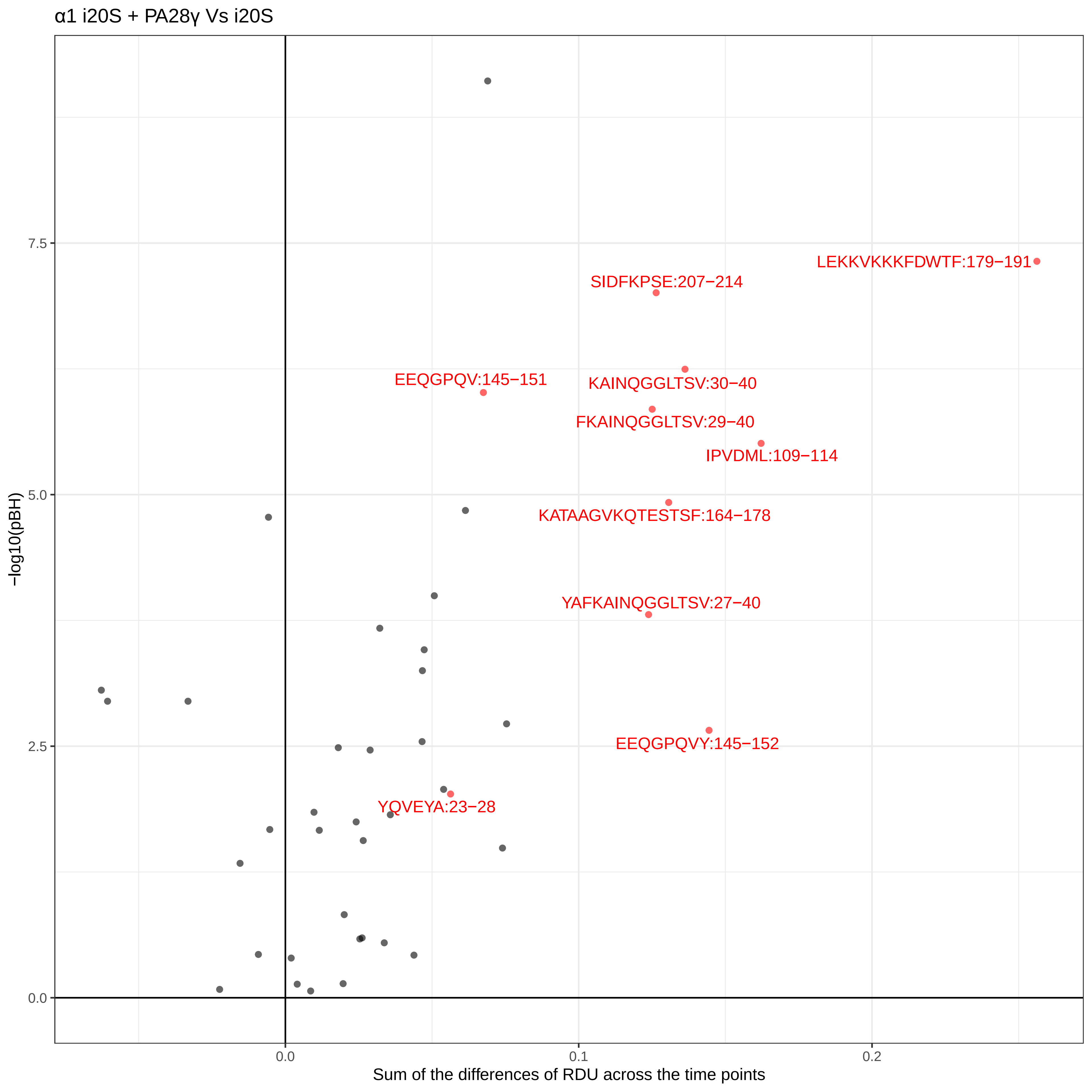
α1 i20S + PA28γ Vs i20S
−log10(pBH)
7.5
5.0
2.5
0.0
0.0
0.1
0.2
Sum of the differences of RDU across the time points
LEKKVKKKFDWTF:179−191
SIDFKPSE:207−214
EEQGPQV:145−151
KAINQGGLTSV:30−40
FKAINQGGLTSV:29−40
IPVDML:109−114
KATAAGVKQTESTSF:164−178
YAFKAINQGGLTSV:27−40
EEQGPQVY:145−152
YQVEYA:23−28

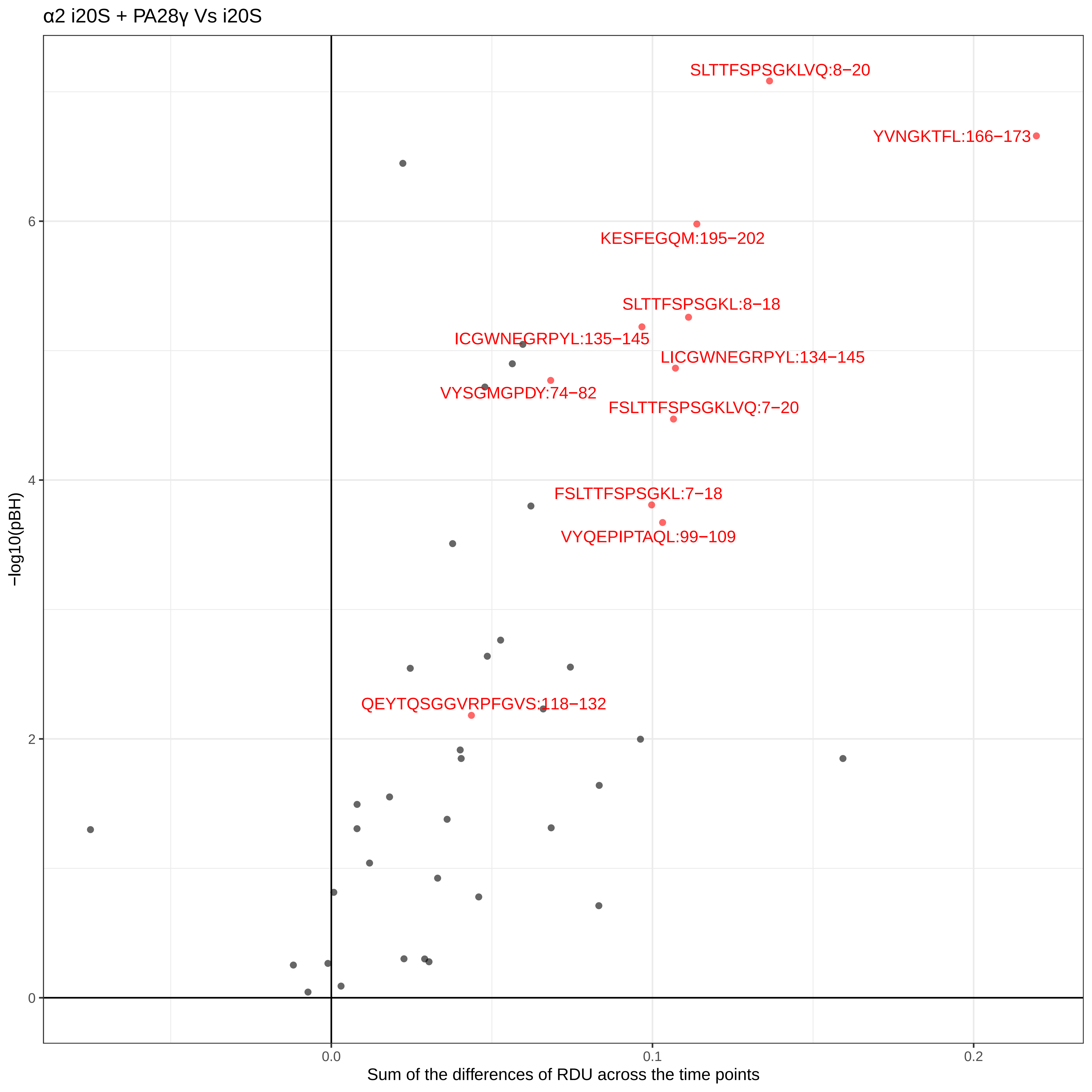

α2 i20S + PA28γ Vs i20S
SLTTFSPSGKLVQ:8−20
YVNGKTFL:166−173
KESFEGQM:195−202
SLTTFSPSGKL:8−18
ICGWNEGRPYL:135−145
LICGWNEGRPYL:134−145
VYSGMGPDY:74−82
FSLTTFSPSGKLVQ:7−20
FSLTTFSPSGKL:7−18
VYQEPIPTAQL:99−109
QEYTQSGGVRPFGVS:118−132
−log10(pBH)
0
2
4
6
0.0
0.1
0.2
Sum of the differences of RDU across the time points

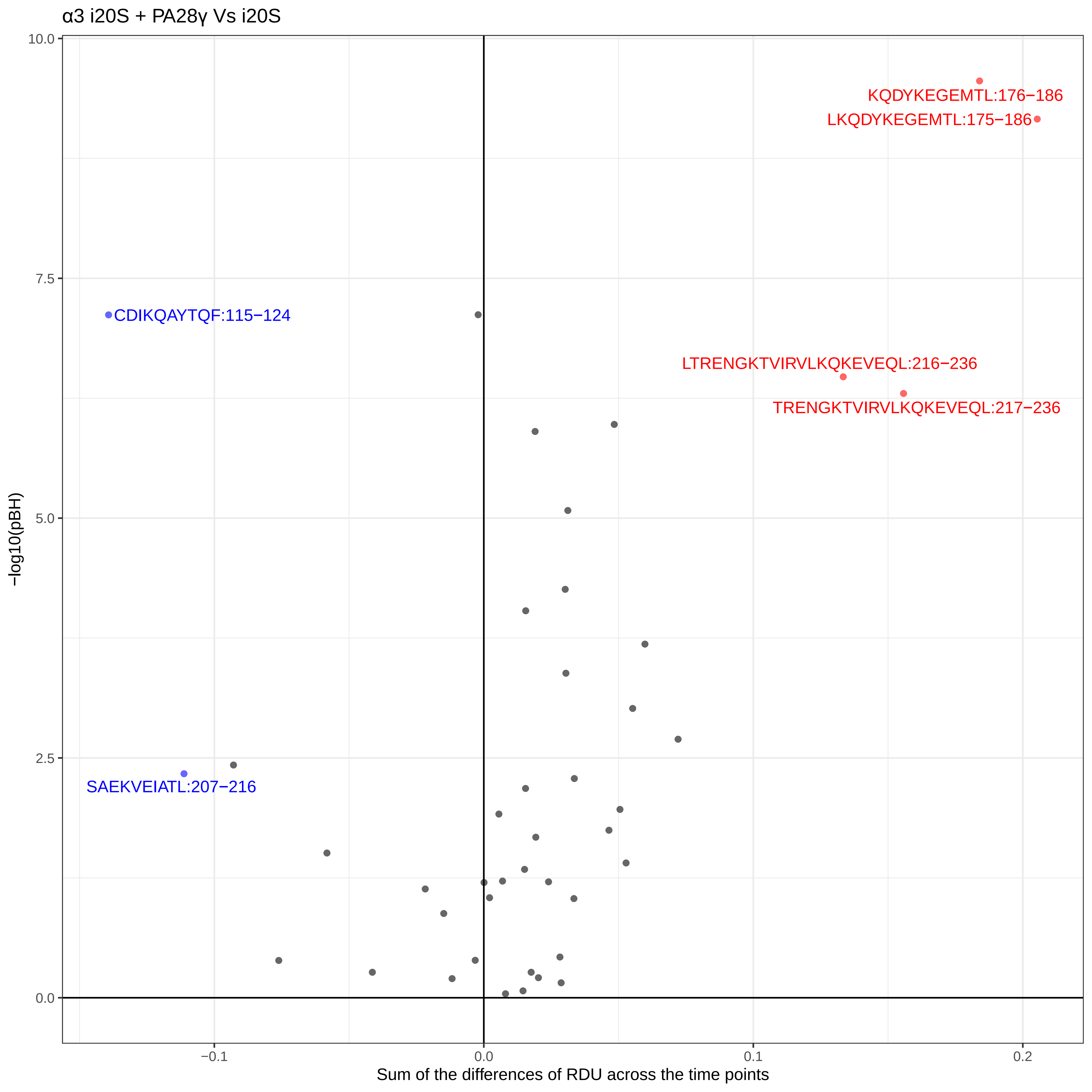
α3 i20S + PA28γ Vs i20S
KQDYKEGEMTL:176−186
LKQDYKEGEMTL:175−186
CDIKQAYTQF:115−124
LTRENGKTVIRVLKQKEVEQL:216−236
TRENGKTVIRVLKQKEVEQL:217−236
SAEKVEIATL:207−216
−log10(pBH)
10.0
7.5
5.0
2.5
0.0
−0.1
0.0
0.1
0.2
Sum of the differences of RDU across the time points

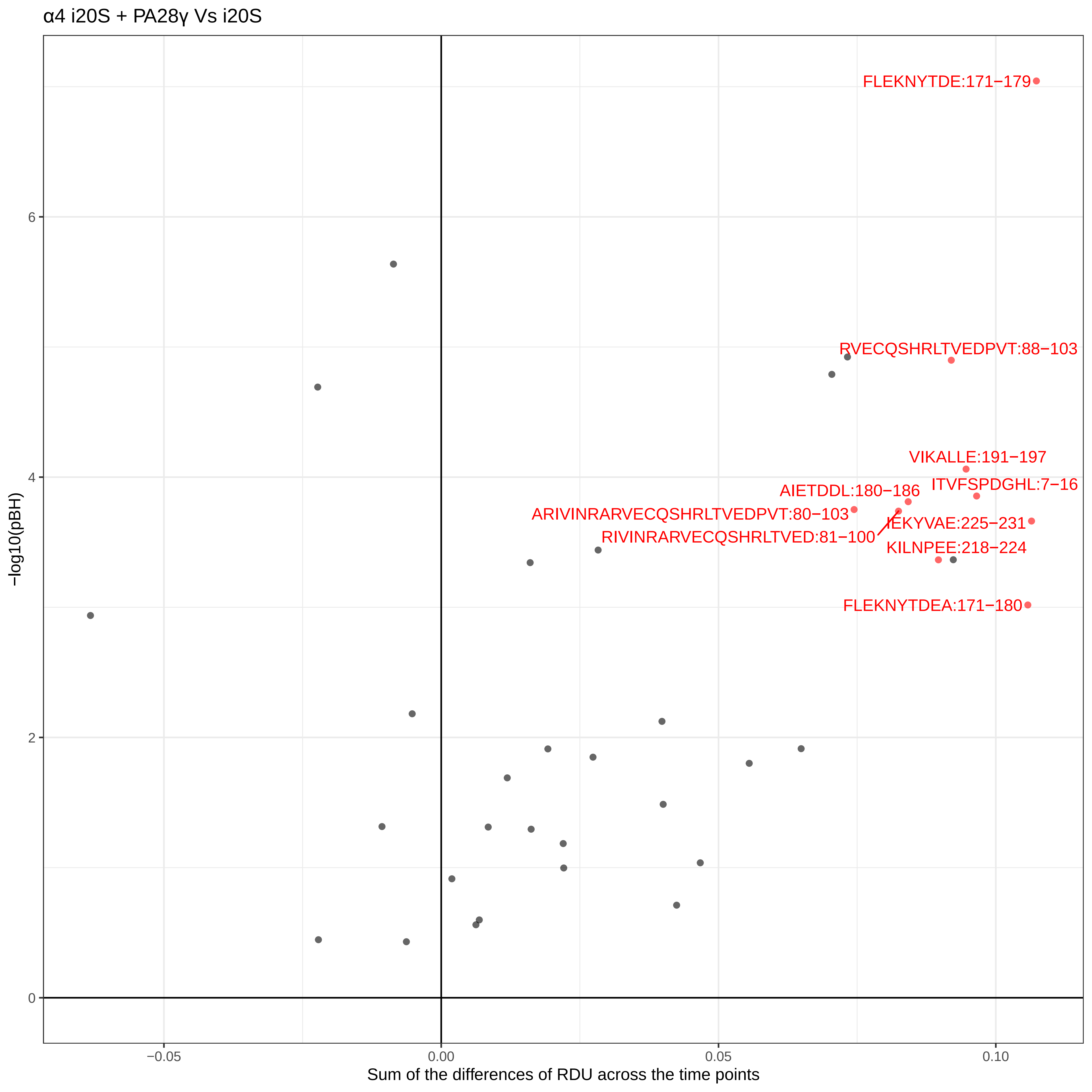

α4 i20S + PA28γ Vs i20S
FLEKNYTDE:171−179
RVECQSHRLTVEDPVT:88−103
VIKALLE:191−197
ITVFSPDGHL:7−16
AIETDDL:180−186
ARIVINRARVECQSHRLTVEDPVT:80−103
RIVINRARVECQSHRLTVED:81−100
IEKYVAE:225−231
KILNPEE:218−224
FLEKNYTDEA:171−180
6
4
2
0
−log10(pBH)
−0.05
0.00
0.05
0.10
Sum of the differences of RDU across the time points

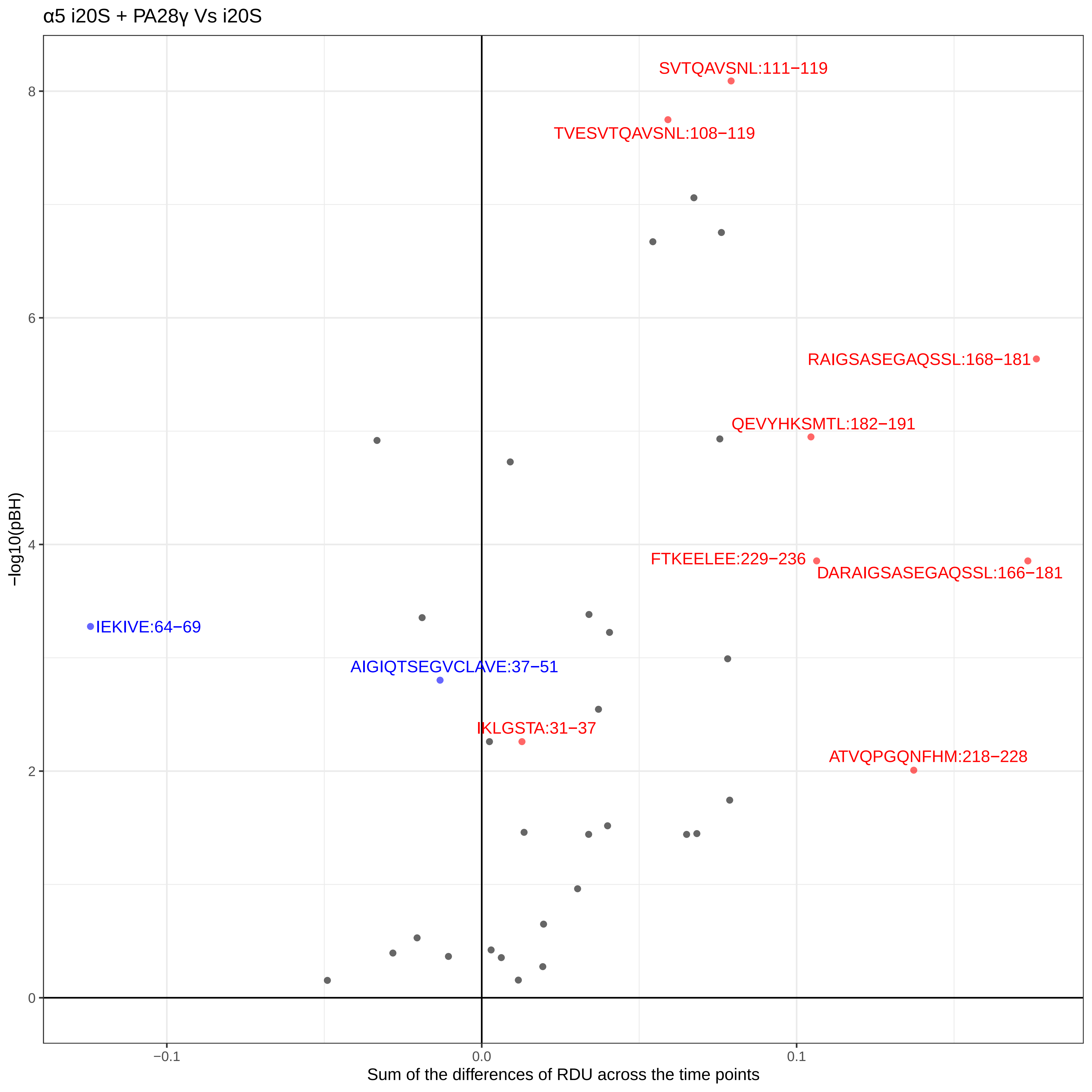
α5 i20S + PA28γ Vs i20S
SVTQAVSNL:111−119
TVESVTQAVSNL:108−119
RAIGSASEGAQSSL:168−181
QEVYHKSMTL:182−191
FTKEELEE:229−236
DARAIGSASEGAQSSL:166−181
IEKIVE:64−69
AIGIQTSEGVCLAVE:37−51
IKLGSTA:31−37
ATVQPGQNFHM:218−228
−log10(pBH)
8
6
4
2
0
−0.1
0.0
0.1
Sum of the differences of RDU across the time points

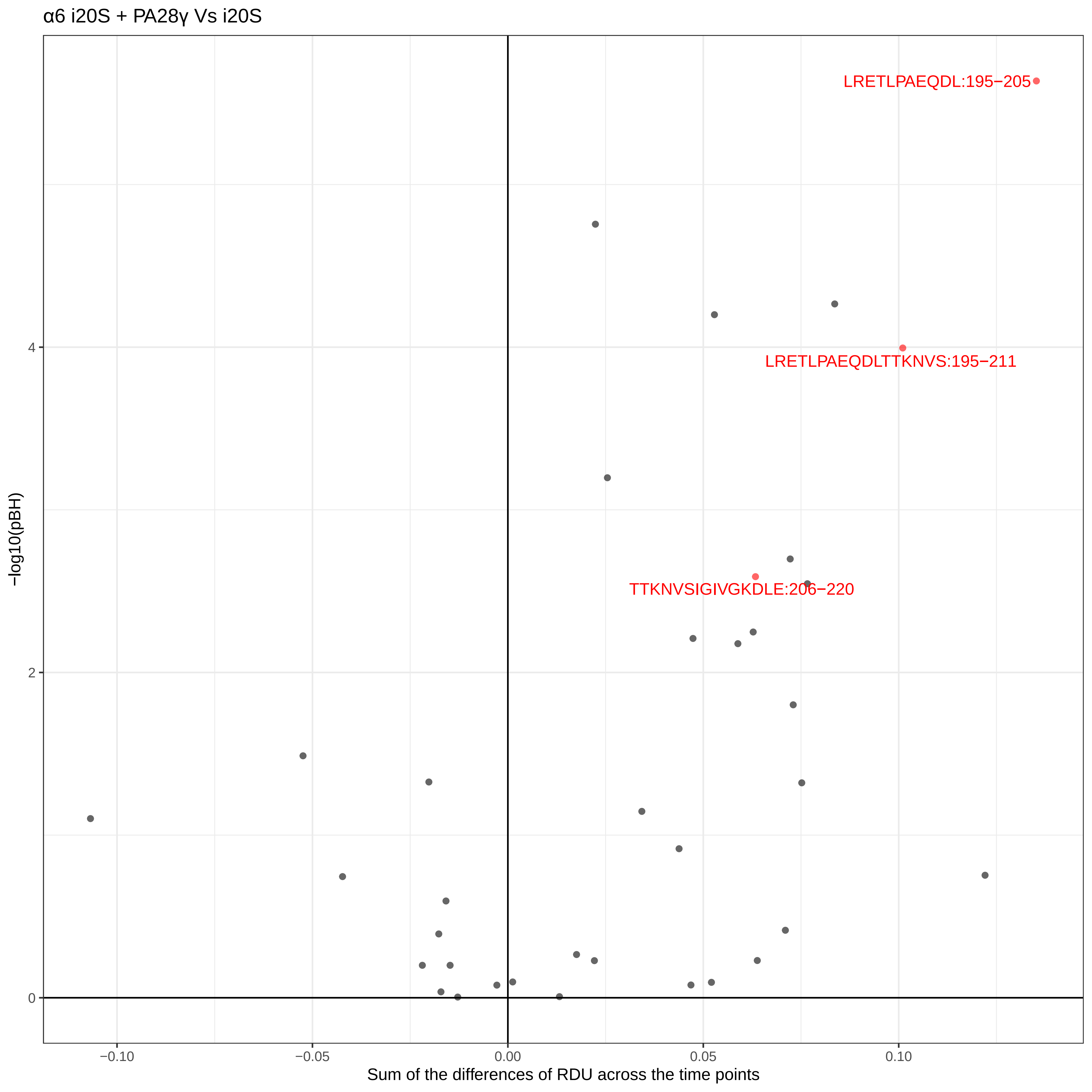
α6 i20S + PA28γ Vs i20S
LRETLPAEQDL:195−205
LRETLPAEQDLTTKNVS:195−211
TTKNVSIGIVGKDLE:206−220
−log10(pBH)
0
2
4
−0.10
−0.05
0.00
0.05
0.10
Sum of the differences of RDU across the time points

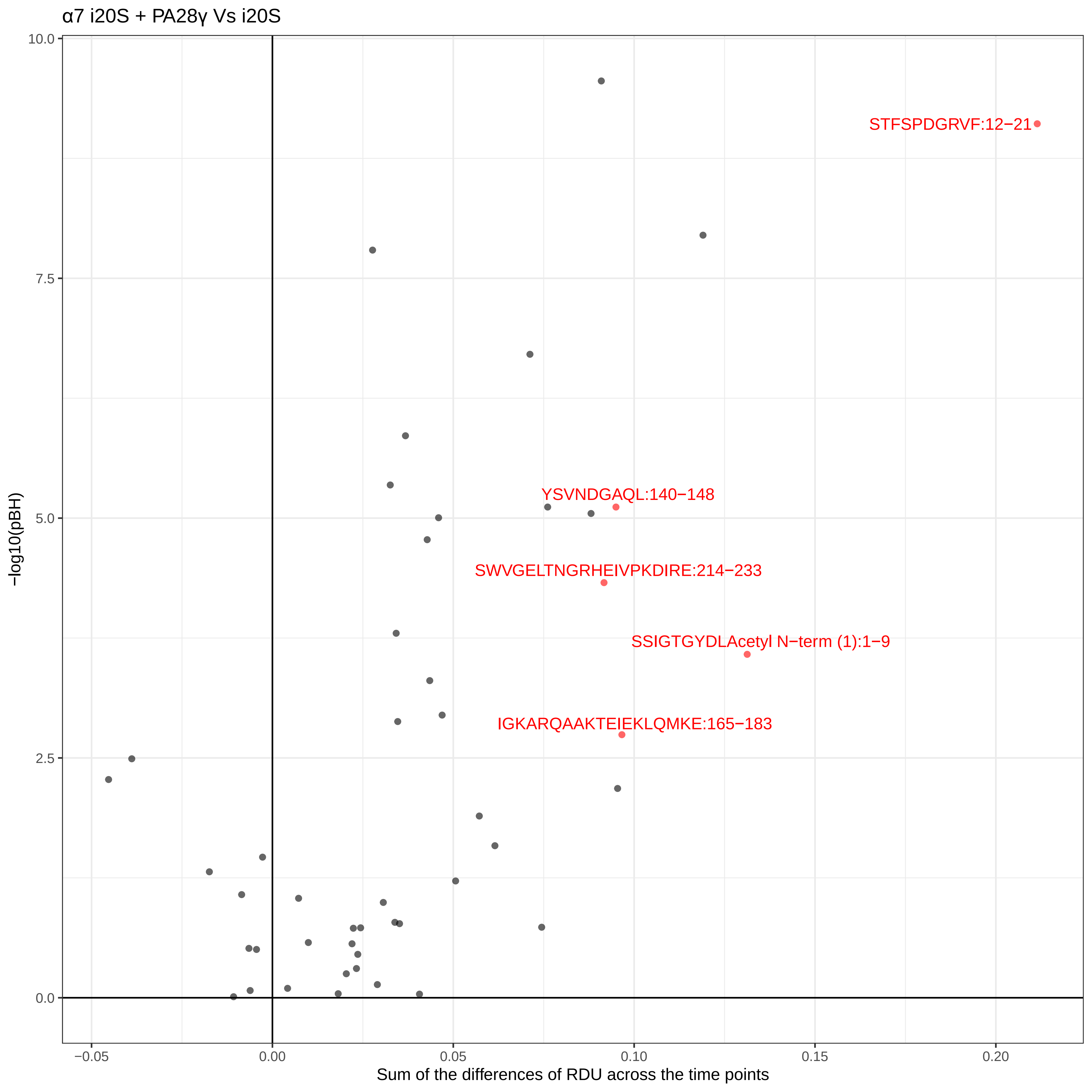

α7 i20S + PA28γ Vs i20S
−log10(pBH)
Sum of the differences of RDU across the time points
STFSPDGRVF:12−21
YSVNDGAQL:140−148
SWVGELTNGRHEIVPKDIRE:214−233
SSIGTGYDLAcetyl N−term (1):1−9
IGKARQAAKTEIEKLQMKE:165−183
10.0
7.5
5.0
2.5
0.0
−0.05
0.00
0.05
0.10
0.15
0.20

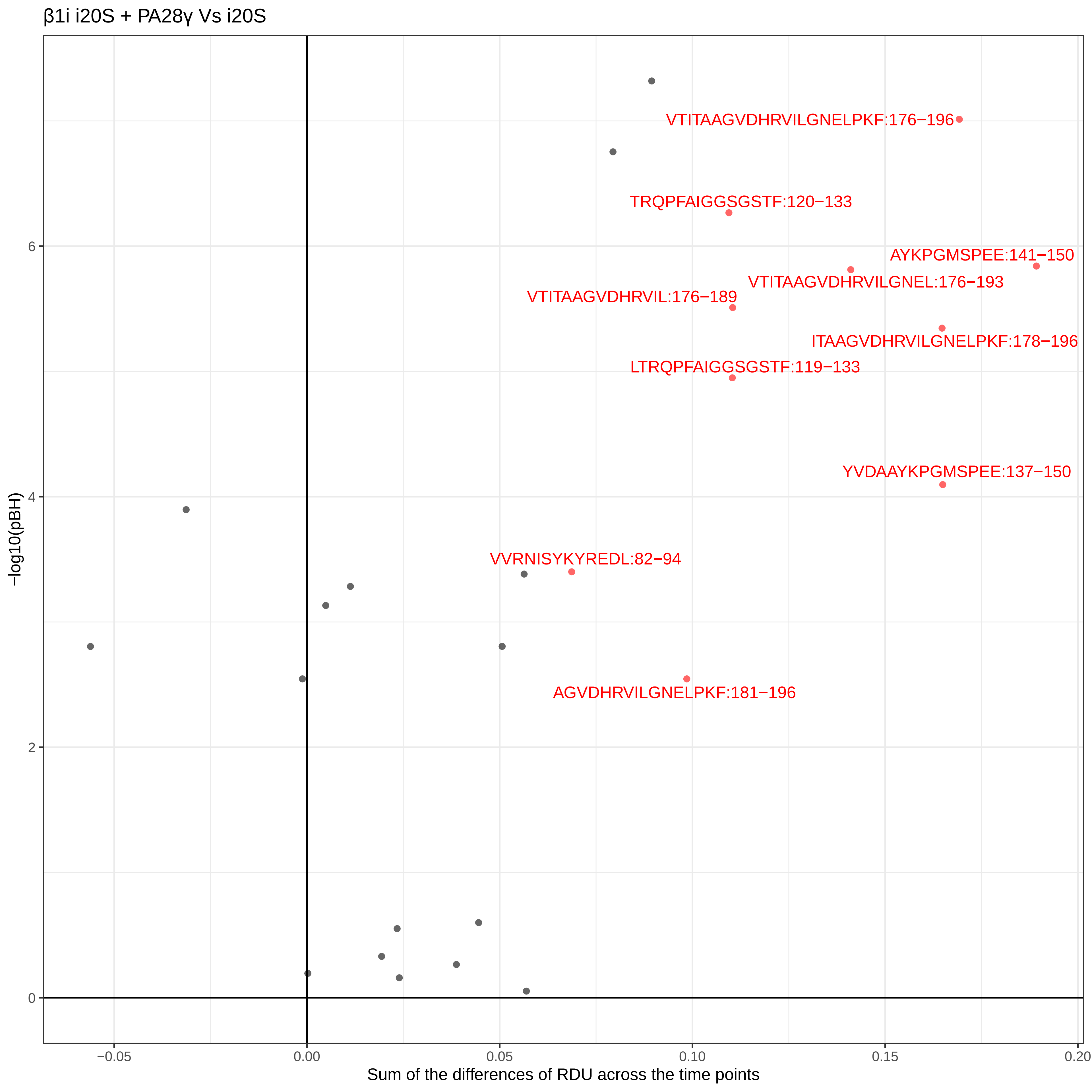

β1i i20S + PA28γ Vs i20S
VTITAAGVDHRVILGNELPKF:176−196
TRQPFAIGGSGSTF:120−133
AYKPGMSPEE:141−150
VTITAAGVDHRVILGNEL:176−193
VTITAAGVDHRVIL:176−189
ITAAGVDHRVILGNELPKF:178−196
LTRQPFAIGGSGSTF:119−133
YVDAAYKPGMSPEE:137−150
VVRNISYKYREDL:82−94
AGVDHRVILGNELPKF:181−196
−log10(pBH)
6
4
2
0
−0.05
0.00
0.05
0.10
0.15
0.20
Sum of the differences of RDU across the time points

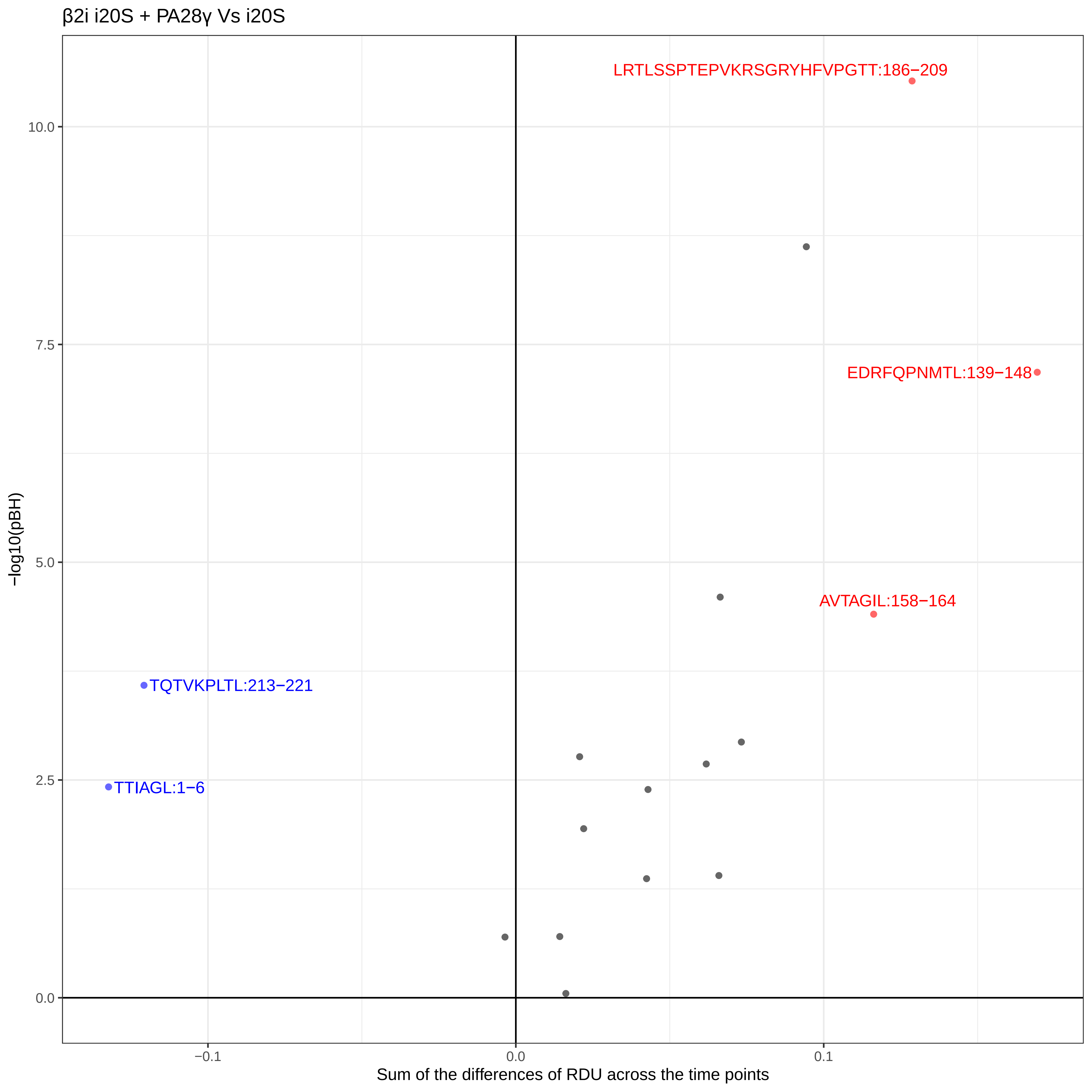
β2i i20S + PA28γ Vs i20S
LRTLSSPTEPVKRSGRYHFVPGTT:186−209
EDRFQPNMTL:139−148
AVTAGIL:158−164
TQTVKPLTL:213−221
TTIAGL:1−6
10.0
7.5
5.0
2.5
0.0
−0.1
0.0
0.1
−log10(pBH)
Sum of the differences of RDU across the time points

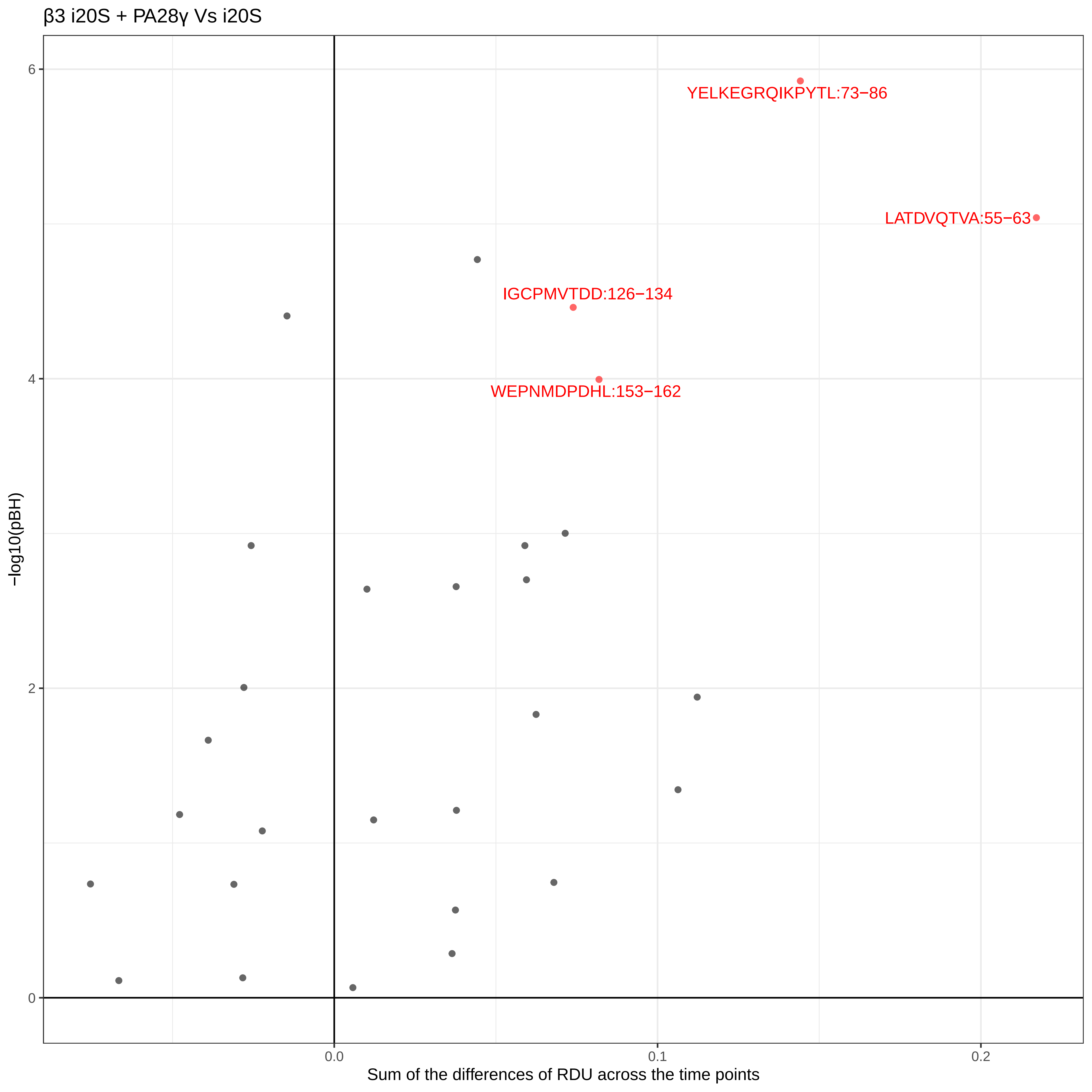

β3 i20S + PA28γ Vs i20S
YELKEGRQIKPYTL:73−86
LATDVQTVA:55−63
IGCPMVTDD:126−134
WEPNMDPDHL:153−162
6
4
2
0
−log10(pBH)
0.0
0.1
0.2
Sum of the differences of RDU across the time points

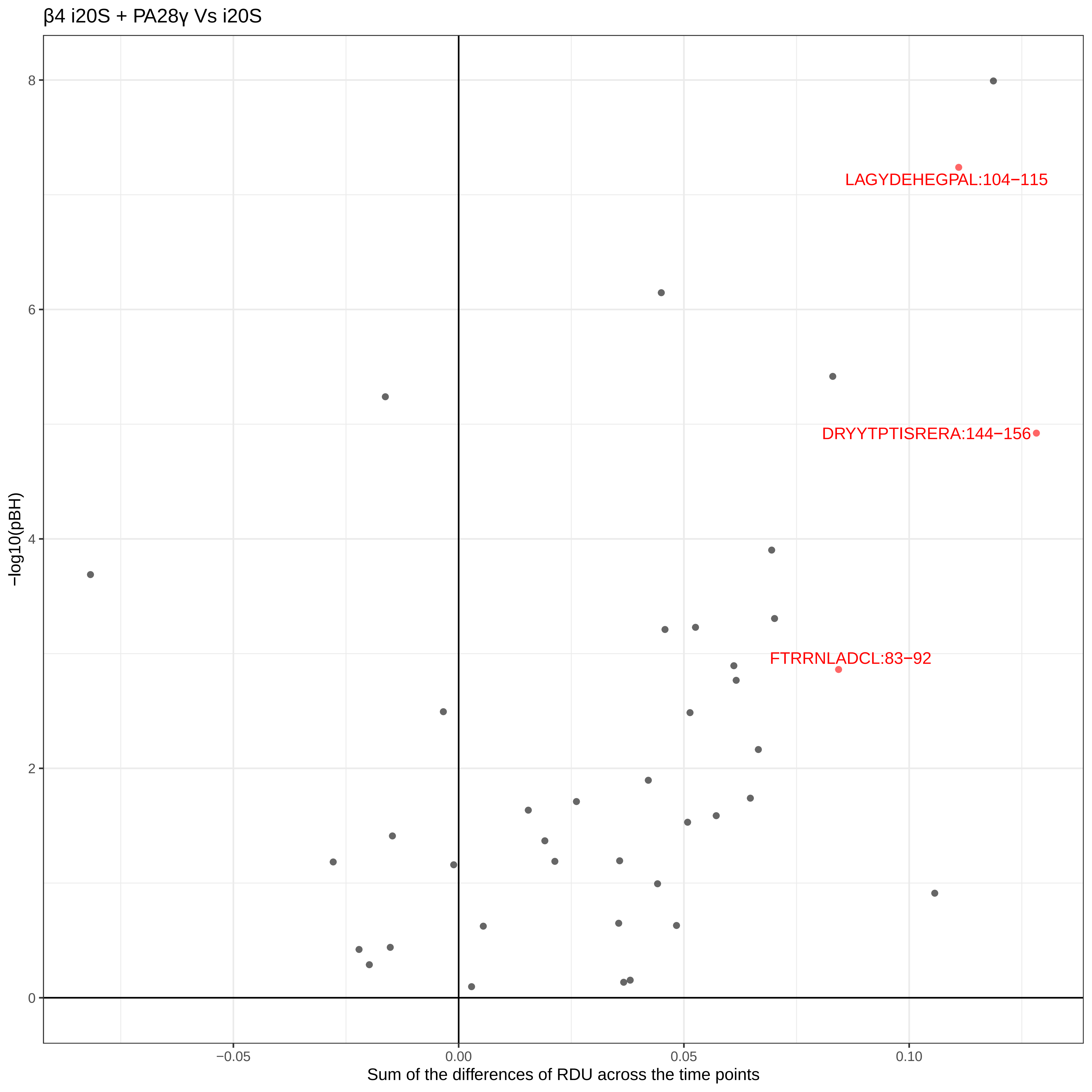
β4 i20S + PA28γ Vs i20S
LAGYDEHEGPAL:104−115
DRYYTPTISRERA:144−156
FTRRNLADCL:83−92
−log10(pBH)
8
6
4
2
0
−0.05
0.00
0.05
0.10
Sum of the differences of RDU across the time points

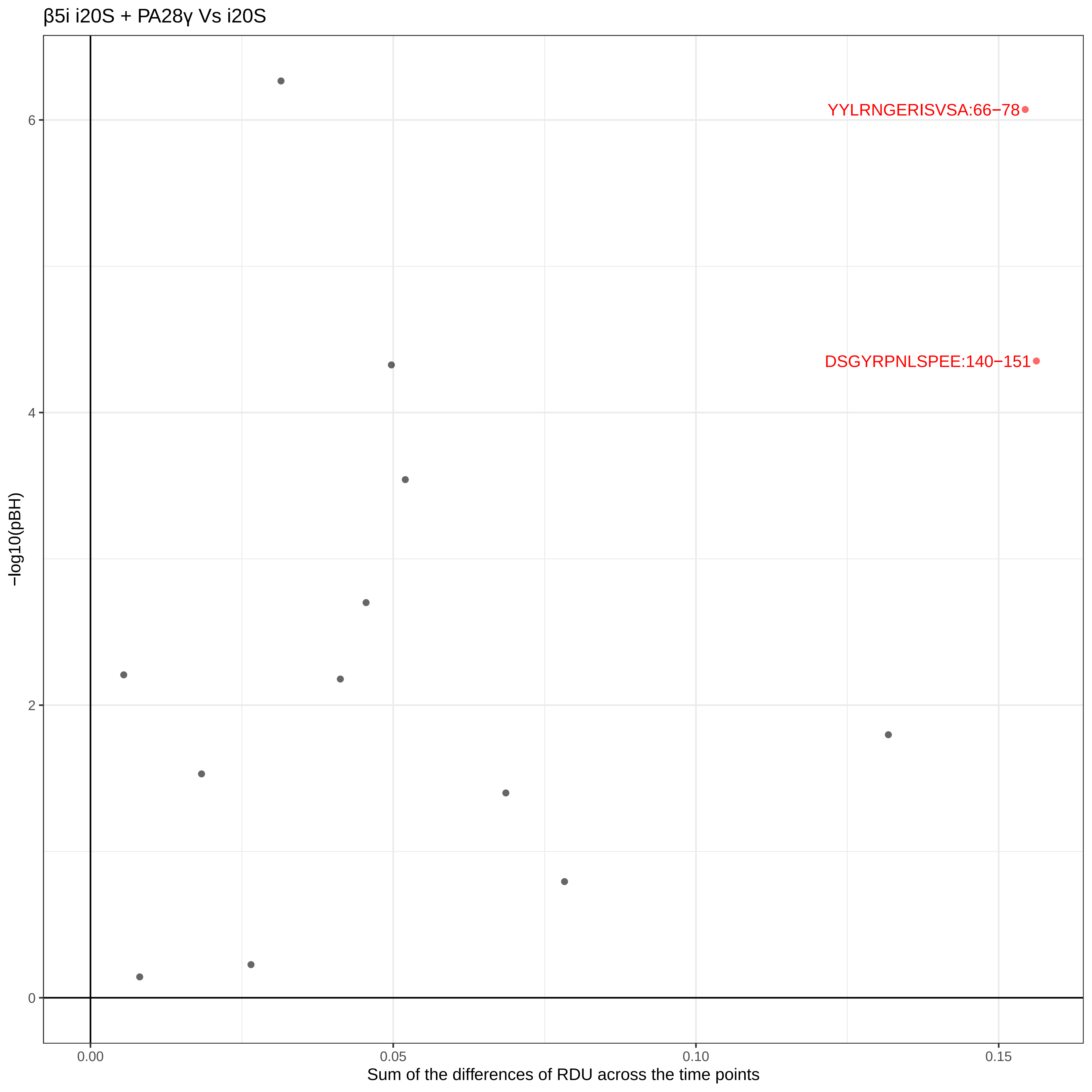

β5i i20S + PA28γ Vs i20S
YYLRNGERISVSA:66−78
DSGYRPNLSPEE:140−151
−log10(pBH)
0
2
4
6
0.00
0.05
0.10
0.15
Sum of the differences of RDU across the time points

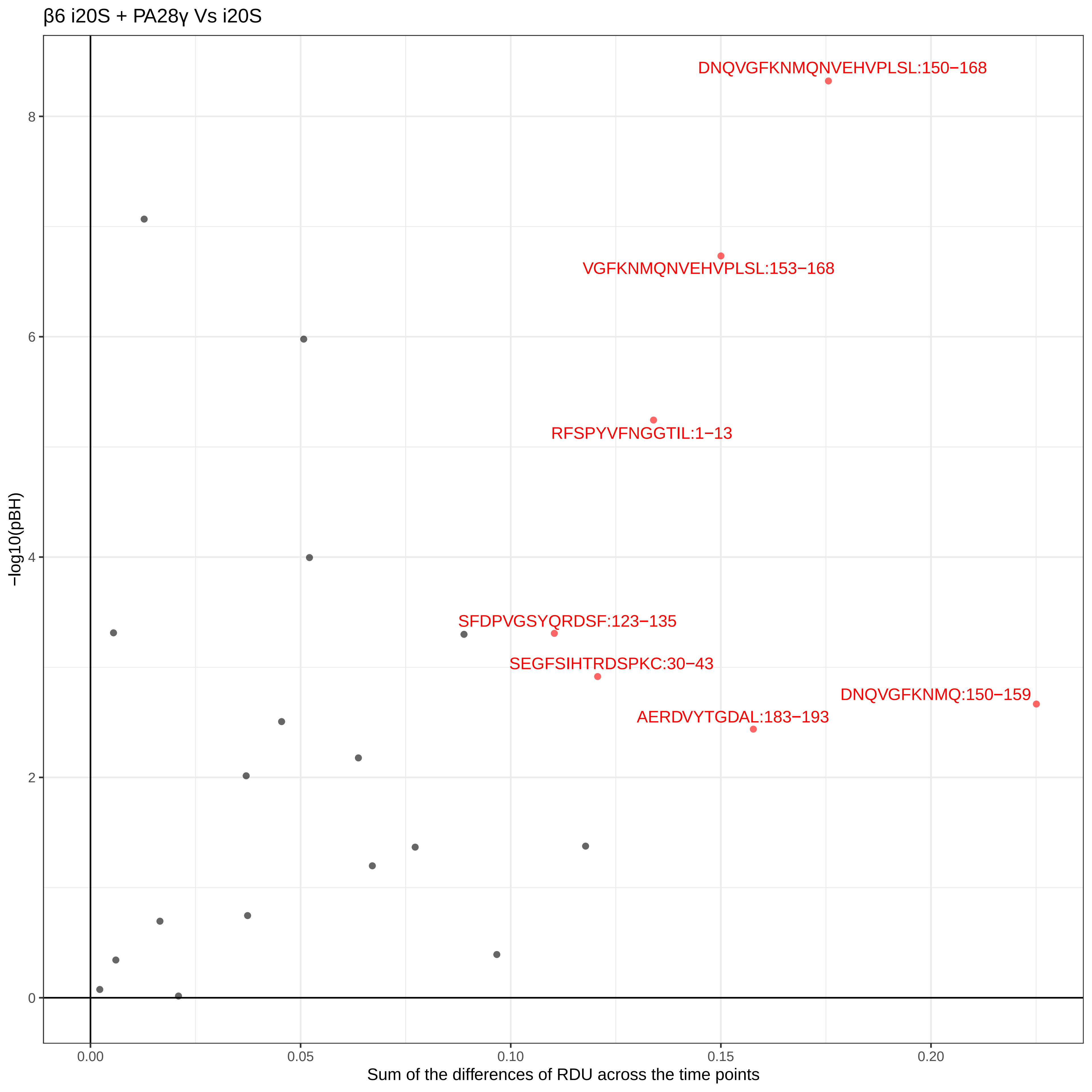
β6 i20S + PA28γ Vs i20S
DNQVGFKNMQNVEHVPLSL:150−168
VGFKNMQNVEHVPLSL:153−168
RFSPYVFNGGTIL:1−13
SFDPVGSYQRDSF:123−135
SEGFSIHTRDSPKC:30−43
DNQVGFKNMQ:150−159
AERDVYTGDAL:183−193
−log10(pBH)
0
2
4
6
8
0.00
0.05
0.10
0.15
0.20
Sum of the differences of RDU across the time points

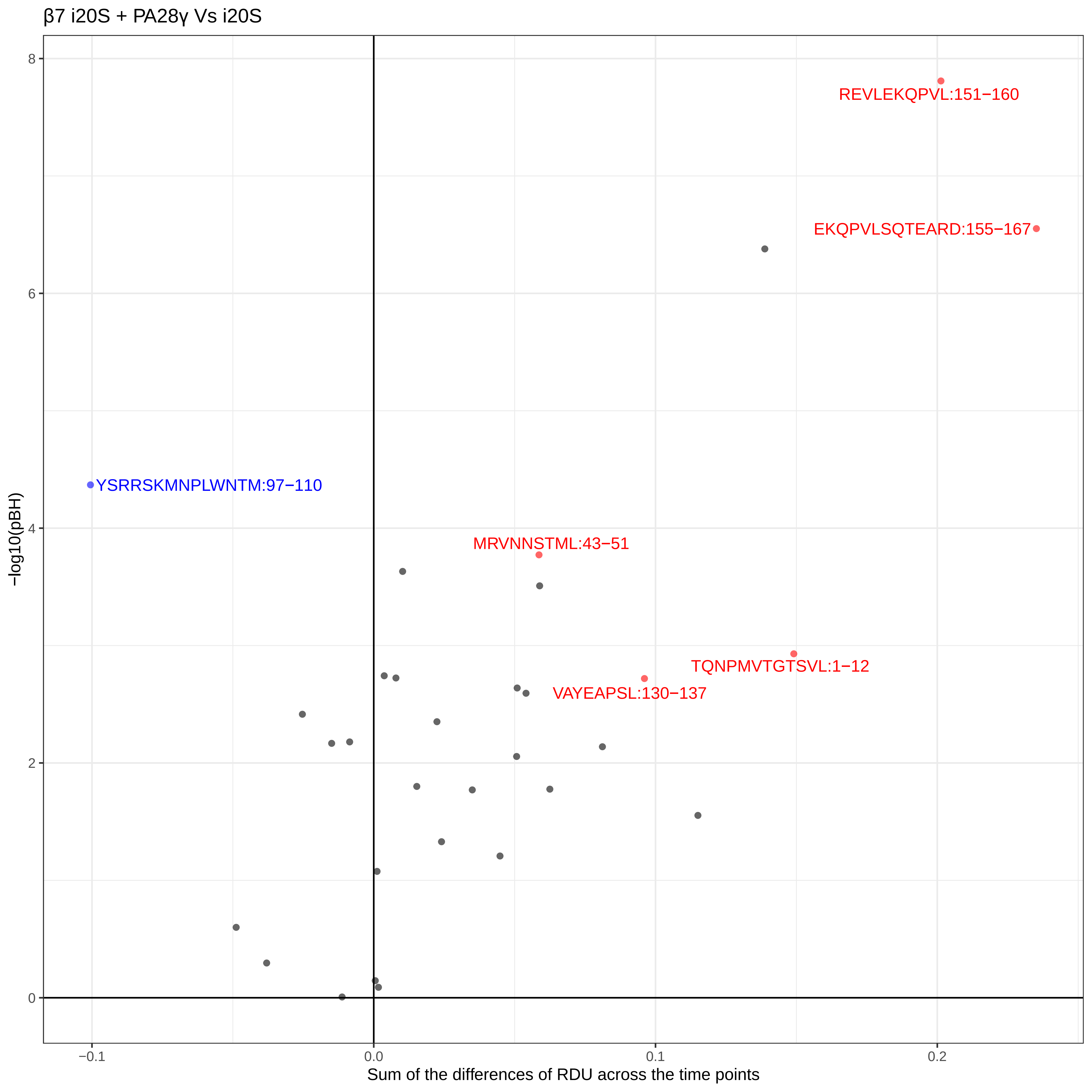
β7 i20S + PA28γ Vs i20S
REVLEKQPVL:151−160
EKQPVLSQTEARD:155−167
YSRRSKMNPLWNTM:97−110
MRVNNSTML:43−51
TQNPMVTGTSVL:1−12
VAYEAPSL:130−137
−log10(pBH)
8
6
4
2
0
−0.1
0.0
0.1
0.2
Sum of the differences of RDU across the time points
